## Supplemental File 1 for "A biophysical model of striatal microcircuits suggests delta/theta-rhythmically interleaved gamma and beta oscillations mediate periodicity in motor control": calcFRcellOut.m

```
function data = calcFRcellOut(data, varargin)
% convert struct to cell output
data = dsCalcFR(data, varargin{:});
flds = fieldnames(data);
fld = flds(~cellfun(@isempty, regexp(flds, '_FR')));
fld = fld{1}; % take first fld with '_FR' string in it
data = data.(fld);
data = {data}; % make cell output
end
```
