## Supplemental File 1 for "A biophysical model of striatal microcircuits suggests delta/theta-rhythmically interleaved gamma and beta oscillations mediate periodicity in motor control": classifyPop1.m

```
function result = classifyPop1(data,varargin)

if ischar(data) && strcmp(data, 'info')
  result = {'nan', []; % Third column for marker.
            'hfo', [];
            'gamma', [];
            'beta', [];
            'alpha', [];
            'theta', [];
            'delta', [];
            'slow', [];
            'silent', [];
            'unclassified', [];
            };
  nClasses = size(result,1);
  classColors = distinguishable_colors(nClasses);
  result(:,2) = num2cell(classColors, 2);
  return
end

if numel(data)>1
  % use dsAnalyzeStudy to recursively call classifyTCRE on each data set
  result = dsAnalyzeStudy(data,@classifyPop1, varargin{:});
  return;
end

data = dsCalcMetrics(data, varargin{:});

% NaN
if data.pop1_metrics.nanV
  result={'nan'};
  
%spiking
elseif any(data.pop1_metrics.muaFR > 0)
  if any(data.pop1_metrics.muaFR > 100)
    result = {'hfo'};
  elseif any(data.pop1_metrics.muaFR > 30)
    result = {'gamma'};
  elseif any(data.pop1_metrics.muaFR > 12)
    result = {'beta'};
  elseif any(data.pop1_metrics.muaFR > 8)
    result = {'alpha'};
  elseif any(data.pop1_metrics.muaFR > 4)
    result = {'theta'};
  elseif any(data.pop1_metrics.muaFR > 1)
    result = {'delta'};
  else
    result = {'slow'};
  end
  
% silent
elseif all(data.pop1_metrics.muaFR) == 0
  result = {'silent'};
  
% Unknown
else
  result = {'unclassified'};
end

end
```
