## Supplemental File 1 for "A biophysical model of striatal microcircuits suggests delta/theta-rhythmically interleaved gamma and beta oscillations mediate periodicity in motor control": dsDemos_classify.m

```
% %%%%%%%%%%%%%%%%%%%%%%%%%%%%%%%%%%%%%%%%%%%%%%%%%%%%%%%%%%%%%%%%%%%%%%%%
%% DynaSim Classification Demos
% %%%%%%%%%%%%%%%%%%%%%%%%%%%%%%%%%%%%%%%%%%%%%%%%%%%%%%%%%%%%%%%%%%%%%%%%
%{
Download DynaSim toolbox from https://github.com/dynasim/dynasim.
  or, download using git: git clone https://github.com/dynasim/dynasim.git
For further documentation, see tutorial.m in the demos directory
Sign up for user mailing list at: https://groups.google.com/forum/#!forum/dynasim-users.

Tip: In Matlab, you can obtain more information associated with any function "FUNCTION_NAME"
by entering "help FUNCTION_NAME" in the command window. Use the "See also" list
at the end of the help section to browse through related help documentation.
%}

% Get ready...

% Add DynaSim to path if it's not already there
if exist('setupDynaSimPath','file')
    setupDynaSimPath;
else
    error('Add the DynaSim folder to the MATLAB path - e.g. run addpath(genpath(DynaSimPath))');
end

% Set where to save outputs
output_directory = dsGetConfig('demos_path');

% move to root directory where outputs will be saved
mkdirSilent(output_directory);
cd(output_directory);

% Here we go!

% %%%%%%%%%%%%%%%%%%%%%%%%%%%%%%%%%%%%%%%%%%%%%%%%%%%%%%%%%%%%%%%%%%%%%%%%
%% Sparse Pyramidal-Interneuron-Network-Gamma (sPING)
% %%%%%%%%%%%%%%%%%%%%%%%%%%%%%%%%%%%%%%%%%%%%%%%%%%%%%%%%%%%%%%%%%%%%%%%%

% Set where to save outputs
study_dir = 'demo_sPING_classify';

% define equations of cell model (same for E and I populations)
eqns={
  'dV/dt=Iapp+@current+noise*randn(1,N_pop)';
  };
% Tip: monitor all functions of a mechanism using: monitor MECHANISM.functions

% create DynaSim specification structure
s=[];
% neural pops
s.populations(1).name='E';
s.populations(1).size=8;
s.populations(1).equations=eqns;
s.populations(1).mechanism_list={'iNa','iK'};
s.populations(1).parameters={'Iapp',0,'gNa',120,'gK',36,'noise',40};
s.populations(2).name='I';
s.populations(2).size=2;
s.populations(2).equations=eqns;
s.populations(2).mechanism_list={'iNa','iK'};
s.populations(2).parameters={'Iapp',0,'gNa',120,'gK',36,'noise',40};

% synapses
s.connections(1).direction='I->E';
s.connections(1).mechanism_list={'iGABAaScaled'};
s.connections(1).parameters={'tauD',10, 'gSYN',1, 'prob_cxn',1};
s.connections(2).direction='E->I';
s.connections(2).mechanism_list={'iAMPAScaled'};
s.connections(2).parameters={'tauD',2, 'gSYN',1, 'prob_cxn',1};

% sim parameters
time_end = 500;
dt = 0.01;

% Specify what to vary
vary={
  'E',     'Iapp',  [0:0.5:1];
  'I',     'Iapp',  [0:0.5:1];
  '(E,I)', 'noise', [0:10:30];
%   'E',     'prob_cxn', [1, 0.5, .1];
%   'I',     'prob_cxn', [1, 0.5, .1];
  'E->I',     'gSYN', [1, 0.5];
  'I->E',     'gSYN', [1, 0.5];
  }; % vary the amplitude of tonic input to E-cells

% if exist(study_dir,'dir')
%   rmdir(study_dir,'s')
% end

dsSimulate(s,'save_data_flag',1, 'save_results_flag',1, 'overwrite_flag',1, 'study_dir',study_dir, 'compile_flag',1,...
  'vary',vary, 'solver','euler', 'dt',0.01, 'verbose_flag',1, 'tspan', [0 time_end], 'downsample_factor',1/dt, 'parallel_flag',1,...
  'analysis_functions',{@classifyEI},...
  'plot_functions',{@dsPlot,@dsPlot},...
  'plot_options',{
    {'varied_filename_flag', 1, 'format', 'jpg', 'visible', 'off'},...
    {'plot_type','rastergram','varied_filename_flag', 1, 'format', 'jpg', 'visible', 'off'},...
  });

%% load and plot the saved data
% study_dir = 'demo_sPING_classify';
% if ~exist('data','var')
%   data=ImportData('demo_sPING_classify');
% end
% PlotData(data);
% PlotData(data,'plot_type','rastergram');

% xAxisVaryParamInd = 2;
% yAxisVaryParamInd = 1;
% PlotClass(study_dir, xAxisVaryParamInd, yAxisVaryParamInd);

% gvRunDS(study_dir, struct('overwrite',1))
% gvRunDS(study_dir, struct('overwrite',0))


% %%%%%%%%%%%%%%%%%%%%%%%%%%%%%%%%%%%%%%%%%%%%%%%%%%%%%%%%%%%%%%%%%%%%%%%%
%% Izhikevich neuron with noisy drive
% %%%%%%%%%%%%%%%%%%%%%%%%%%%%%%%%%%%%%%%%%%%%%%%%%%%%%%%%%%%%%%%%%%%%%%%%

% Izhikevich study of neuro-computational properties (using Syntax 1)
% based on: http://www.izhikevich.org/publications/izhikevich.m

% Set where to save outputs
study_dir = 'demo_izhikevich_classify';

% define equations of cell model
eqns={
  'a=.02; b=.2; c=-65; d=6; I=14';
  'dV/dt=.04*V^2+5*V+140-u+I; V(0)=-70';
  'du/dt=a*(b*V-u); u(0)=-20';
  'if(V>=30)(V=c;u=u+d)';
  };

% create DynaSim specification structure
s=[];
s.populations(1).name = 'pop1';
s.populations(1).size = 1;
s.populations(1).equations = eqns;

% sim parameters
time_end = 500;
dt = 0.01;

% Specify what to vary
P='pop1'; % name of population
vary={
  {P,'a',.02; P,'b',.2 ; P,'c',-50; P,'d',2;  P,'I',15} % tonic bursting
  {P,'a',.01; P,'b',.2 ; P,'c',-65; P,'d',8;  P,'I',30} % spike frequency adaptation
  {P,'a',.02; P,'b',.2 ; P,'c',-65; P,'d',6;  P,'I',7}  % spike latency
  {P,'a',.03; P,'b',.25; P,'c',-52; P,'d',0;  P,'I',0}  % rebound burst
  {P,'a',1;   P,'b',1.5; P,'c',-60; P,'d',0;  P,'I',-65}% bistability
  {P,'a',.02; P,'b',1  ; P,'c',-55; P,'d',4;  P,'I',1}  % accomodation
  {P,'a',-.02;P,'b',-1 ; P,'c',-60; P,'d',8;  P,'I',80} % inhibition-induced spiking
  {P,'a',-.026;P,'b',-1; P,'c',-45; P,'d',0;  P,'I',70} % inhibition-induced bursting
  };

dsSimulate(s,'save_data_flag',1, 'save_results_flag',1, 'overwrite_flag',1, 'study_dir',study_dir, 'compile_flag',1,...
  'vary',vary, 'solver','euler', 'dt',0.01, 'verbose_flag',1, 'tspan', [0 time_end], 'downsample_factor',1/dt, 'parallel_flag',1,...
  'analysis_functions',{@classifyPop1, @calcFRcellOut},...
  'analysis_options',{{},{'variable','*_V', 'bin_size',.99}},...
  'plot_functions',{@dsPlot},...
  'plot_options',{
    {'varied_filename_flag', 1, 'format', 'jpg', 'visible', 'off'},...
  });

%% Izhikevich neuron with noisy drive sweep
% %%%%%%%%%%%%%%%%%%%%%%%%%%%%%%%%%%%%%%%%%%%%%%%%%%%%%%%%%%%%%%%%%%%%%%%%

% Izhikevich study of neuro-computational properties (using Syntax 1)
% based on: http://www.izhikevich.org/publications/izhikevich.m

% Set where to save outputs
study_dir = 'demo_izhikevich_classify_lattice';

% define equations of cell model
eqns={
  'a=.02; b=.2; c=-65; d=6; I=14';
  'dV/dt=.04*V^2+5*V+140-u+I; V(0)=-70';
  'du/dt=a*(b*V-u); u(0)=-20';
  'if(V>=30)(V=c;u=u+d)';
  };

% create DynaSim specification structure
s=[];
s.populations(1).name = 'pop1';
s.populations(1).size = 1;
s.populations(1).equations = eqns;

% sim parameters
time_end = 1000;
dt = 0.01;

% Specify what to vary
P='pop1'; % name of population
vary={
  P,'a',linspace(-.1, 1, 5);
  P,'b',linspace(-.1, 1, 5);
  P,'c',[-45, -55, -65];
  P,'d',linspace(0, 8, 5);
  P,'I',[0, 30, 70]
  };

dsSimulate(s,'save_data_flag',1, 'save_results_flag',1, 'overwrite_flag',1, 'study_dir',study_dir, 'compile_flag',1,...
  'vary',vary, 'solver','euler', 'dt',0.01, 'verbose_flag',1, 'tspan', [0 time_end], 'downsample_factor',1/dt, 'parallel_flag',1,...
  'analysis_functions',{@classifyPop1, @calcFRcellOut},...
  'analysis_options',{{},{'variable','*_V', 'bin_size',.99}},...
  'plot_functions',{@dsPlot},...
  'plot_options',{
    {'varied_filename_flag', 1, 'format', 'jpg', 'visible', 'off'},...
  });
```
