## Supplemental File 1 for "A biophysical model of striatal microcircuits suggests delta/theta-rhythmically interleaved gamma and beta oscillations mediate periodicity in motor control": dsDemos_classify_scc.m

```
% %%%%%%%%%%%%%%%%%%%%%%%%%%%%%%%%%%%%%%%%%%%%%%%%%%%%%%%%%%%%%%%%%%%%%%%%
%% DynaSim Classification Demos
% %%%%%%%%%%%%%%%%%%%%%%%%%%%%%%%%%%%%%%%%%%%%%%%%%%%%%%%%%%%%%%%%%%%%%%%%
%{
Download DynaSim toolbox from https://github.com/dynasim/dynasim.
  or, download using git: git clone https://github.com/dynasim/dynasim.git
For further documentation, see tutorial.m in the demos directory
Sign up for user mailing list at: https://groups.google.com/forum/#!forum/dynasim-users.

Tip: In Matlab, you can obtain more information associated with any function "FUNCTION_NAME"
by entering "help FUNCTION_NAME" in the command window. Use the "See also" list
at the end of the help section to browse through related help documentation.
%}

% Get ready...

% Add DynaSim to path if it's not already there
if exist('setupDynaSimPath','file')
    setupDynaSimPath;
else
    error('Add the DynaSim folder to the MATLAB path - e.g. run addpath(genpath(DynaSimPath))');
end

% Set where to save outputs
output_directory = dsGetConfig('demos_path');

% move to root directory where outputs will be saved
mkdirSilent(output_directory);
cd(output_directory);

% Set where to save outputs
study_dir = 'demo_sPING_classify_scc';

% Here we go!

% %%%%%%%%%%%%%%%%%%%%%%%%%%%%%%%%%%%%%%%%%%%%%%%%%%%%%%%%%%%%%%%%%%%%%%%%
%% BUILDING LARGE MODELS WITH MULTIPLE POPULATIONS AND CONNECTIONS
% %%%%%%%%%%%%%%%%%%%%%%%%%%%%%%%%%%%%%%%%%%%%%%%%%%%%%%%%%%%%%%%%%%%%%%%%

%% Sparse Pyramidal-Interneuron-Network-Gamma (sPING)

% define equations of cell model (same for E and I populations)
eqns={
  'dV/dt=Iapp+@current+noise*randn(1,N_pop)';
  };
% Tip: monitor all functions of a mechanism using: monitor MECHANISM.functions

% create DynaSim specification structure
s=[];
% neural pops
s.populations(1).name='E';
s.populations(1).size=80;
s.populations(1).equations=eqns;
s.populations(1).mechanism_list={'iNa','iK'};
s.populations(1).parameters={'Iapp',0,'gNa',120,'gK',36,'noise',40};
s.populations(2).name='I';
s.populations(2).size=20;
s.populations(2).equations=eqns;
s.populations(2).mechanism_list={'iNa','iK'};

% synapses
s.connections(1).direction='I->E';
s.connections(1).mechanism_list={'iGABAaScaled'};
s.connections(1).parameters={'tauD',10, 'gSYN',1, 'prob_cxn',1};
s.connections(2).direction='E->I';
s.connections(2).mechanism_list={'iAMPAScaled'};
s.connections(2).parameters={'tauD',2, 'gSYN',1, 'prob_cxn',1};

% %%%%%%%%%%%%%%%%%%%%%%%%%%%%%%%%%%%%%%%%%%%%%%%%%%%%%%%%%%%%%%%%%%%%%%%%
%% SAVING SIMULATED DATA
% %%%%%%%%%%%%%%%%%%%%%%%%%%%%%%%%%%%%%%%%%%%%%%%%%%%%%%%%%%%%%%%%%%%%%%%%
% How to: set 'save_data_flag' to 1 and (optionally) 'study_dir' to /path/to/outputs

%% Save data from a set of simulations
time_end = 500;
dt = 0.01;

% Specify what to vary
% Tip: use 'vary' Syntax to to systematically vary a parameter
vary={
  'E',     'Iapp',  [0:.5:1];
  'I',     'Iapp',  [0:.5:1];
%   '(E,I)', 'noise', [0:25:50];
  'E->I',     'prob_cxn', [1, 0.5, .1];
  'I->E',     'prob_cxn', [1, 0.5, .1];
  'E',     'gSYN', [1, 0.5, .1];
  'I',     'gSYN', [1, 0.5, .1];
  }; % vary the amplitude of tonic input to E-cells

if exist(study_dir,'dir')
  rmdir(study_dir,'s')
end

data = dsSimulate(s,'save_data_flag',0, 'save_results_flag',1, 'overwrite_flag',1, 'study_dir',study_dir,...
  'vary',vary, 'solver','euler', 'dt',dt, 'verbose_flag',1, 'tspan', [0 time_end], 'downsample_factor',1/dt,...
  'analysis_functions',{@classifyPop1, @calcFRcellOut},...
  'analysis_options',{{},{'variable','*_V', 'bin_size',.99}},...
  'plot_functions',{@dsPlotData,@dsPlotData,@dsPlotData},...
  'plot_options',{
    {'varied_filename_flag', 0, 'format', 'jpg', 'visible', 'off'},...
    {'plot_type','rastergram','varied_filename_flag', 0, 'format', 'jpg', 'visible', 'off'},...
    {'plot_type','power', 'xlim',[0 100], 'varied_filename_flag',0, 'format','jpg', 'visible','off'},...
  },...
  'cluster_flag',1, 'qsub_mode','array', 'optimize_big_vary',1, 'sims_per_job',20);

%% load and plot the saved data

% Run this after sims finish
% gvRun(study_dir)
```
