## Supplemental File 1 for "A biophysical model of striatal microcircuits suggests delta/theta-rhythmically interleaved gamma and beta oscillations mediate periodicity in motor control": dsDemos_cluster_GUI_noVerbose.m

```
% %%%%%%%%%%%%%%%%%%%%%%%%%%%%%%%%%%%%%%%%%%%%%%%%%%%%%%%%%%%%%%%%%%%%%%%%
%% DynaSim Demos
% %%%%%%%%%%%%%%%%%%%%%%%%%%%%%%%%%%%%%%%%%%%%%%%%%%%%%%%%%%%%%%%%%%%%%%%%
%{
Download DynaSim toolbox from https://github.com/dynasim/dynasim.
  or, download using git: git clone https://github.com/dynasim/dynasim.git
For further documentation, see tutorial.m in the demos directory
Sign up for user mailing list at: https://groups.google.com/forum/#!forum/dynasim-users.

Tip: In Matlab, you can obtain more information associated with any function "FUNCTION_NAME"
by entering "help FUNCTION_NAME" in the command window. Use the "See also" list
at the end of the help section to browse through related help documentation.
%}

% Add DynaSim to path if it's not already there
if exist('setupDynaSimPath','file')
    setupDynaSimPath;
else
    error('Add the DynaSim folder to the MATLAB path - e.g.) addpath(genpath(''/path/to/dynasim''))');
end

% Set where to save outputs
output_directory = dsGetConfig('demos_path');

% move to root directory where outputs will be saved
mkdir(output_directory);
cd(output_directory);

% Here we go!

% %%%%%%%%%%%%%%%%%%%%%%%%%%%%%%%%%%%%%%%%%%%%%%%%%%%%%%%%%%%%%%%%%%%%%%%%
%% DEFINING AND SIMULATING MODELS
% %%%%%%%%%%%%%%%%%%%%%%%%%%%%%%%%%%%%%%%%%%%%%%%%%%%%%%%%%%%%%%%%%%%%%%%%

%% Lorenz equations with phase plot

% DynaSim makes it easy to simulate arbitrary systems of ordinary
% differential equations. Simply write out the system in a cell array of
% strings, separating equations into different strings or the same string
% separated by semicolons.

eqns={
  's=10; r=27; b=2.666'
  'dx/dt=s*(y-x);   x(0)=1'
  'dy/dt=r*x-y-x*z; y(0)=2'
  'dz/dt=-b*z+x*y;  z(0)=.5'
};
data=dsSimulate(eqns, 'time_limits',[0 100], 'solver','rk4');

% time_limits: time limits on integration [ms]
% solver: numerical method to use (default: rk4 = "4th-order Runge-Kutta")

% All models are numerically integrated using a DynaSim solver function
% created uniquely for a given model and stored in a directory named
% "solve". The file that solves the system (i.e,. numerically integrates
% it) is stored in data.simulator_options and can be viewed or rerun afterwards:
% edit(data.simulator_options.solve_file)

% Every component of the model is assigned to a "population", and the
% population name (default: 'pop1') is prepended to all variable and
% function names.

% Simulated data can be easily plotted using the resulting data structure:
figure; plot(data.pop1_x,data.pop1_z); % <-- Figure 1 in DynaSim paper
title('Lorenz equations'); xlabel('x'); ylabel('z')

%% Izhikevich neuron with noisy drive
% (reference: p274 of "Dynamical Systems in Neuroscience" by Izhikevich)

% The DynaSim data structure always contains the model state variables,
% time vector, and a copy of the DynaSim model structure that was
% simulated. Additionally, functions can be recorded and returned in the
% DynaSim data structure if indicated using the "monitor" keyword.
% Syntax: monitor FUNCTION

eqns={
  'C=100; vr=-60; vt=-40; k=.7; Iapp=70; ton=200; toff=800'
  'a=.03; b=-2; c=-50; d=100; vpeak=35'
  'dv/dt=(k*(v-vr)*(v-vt)-u+I(t))/C; v(0)=vr'
  'du/dt=a*(b*(v-vr)-u); u(0)=0'
  'if(v>vpeak)(v=c; u=u+d)'
  'I(t)=Iapp*(t>ton&t<toff).*(1+.5*rand(1,Npop))' % define applied input using reserved variables 't' for time and 'dt' for fixed time step of numerical integration
  'monitor I'                            % indicate to store applied input during simulation
  'monitor w(t)=u.*v'                    % defining a function of state variables to monitor (note that monitor expressions follow Matlab's syntax)
};
data=dsSimulate(eqns, 'time_limits',[0 1000]);

% plot the simulated voltage and monitored input function
figure; % <-- Figure 2 in DynaSim paper
subplot(2,1,1); plot(data.time,data.pop1_v); % plot voltage
xlabel('time (ms)'); ylabel('v'); title('Izhikevich neuron')
subplot(2,1,2); plot(data.time,data.pop1_I); % plot input function
xlabel('time (ms)'); ylabel('Iapp');

% note: "t", "dt", and "T" are special variables that can be used in model
% equations. "t" represents the current time point of the simulation.
% "dt" is the fixed time step used for numeric integration. "T" is the full
% simulated time vector defined before simulation begins.

% %%%%%%%%%%%%%%%%%%%%%%%%%%%%%%%%%%%%%%%%%%%%%%%%%%%%%%%%%%%%%%%%%%%%%%%%
%% RUNNING SETS OF SIMULATIONS
% %%%%%%%%%%%%%%%%%%%%%%%%%%%%%%%%%%%%%%%%%%%%%%%%%%%%%%%%%%%%%%%%%%%%%%%%

% 'vary' indicates the variable to vary, the values it should take, and the
% object (population or connection) whose variable should be varied.

% Syntax 1: vary={{object, variable, value1},{object, variable, value2},...}
%   - this is useful for simulating an arbitrary set of parameter values
% Syntax 2: vary={object, variable, values; ...}
%   - this is useful for varying parameters systematically (described later)

% Izhikevich study of neuro-computational properties (using Syntax 1)
% based on: http://www.izhikevich.org/publications/izhikevich.m
eqns={
  'a=.02; b=.2; c=-65; d=6; I=14'
  'dv/dt=.04*v^2+5*v+140-u+I; v(0)=-70'
  'du/dt=a*(b*v-u); u(0)=-20'
  'if(v>=30)(v=c;u=u+d)'
  };
P='pop1'; % name of population
vary={
  {P,'a',.02; P,'b',.2 ; P,'c',-50; P,'d',2;  P,'I',15} % tonic bursting
  {P,'a',.01; P,'b',.2 ; P,'c',-65; P,'d',8;  P,'I',30} % spike frequency adaptation
  {P,'a',.02; P,'b',.2 ; P,'c',-65; P,'d',6;  P,'I',7}  % spike latency
  {P,'a',.03; P,'b',.25; P,'c',-52; P,'d',0;  P,'I',0}  % rebound burst
  {P,'a',1;   P,'b',1.5; P,'c',-60; P,'d',0;  P,'I',-65}% bistability
  {P,'a',.02; P,'b',1  ; P,'c',-55; P,'d',4;  P,'I',1}  % accomodation
  {P,'a',-.02;P,'b',-1 ; P,'c',-60; P,'d',8;  P,'I',80} % inhibition-induced spiking
  {P,'a',-.026;P,'b',-1; P,'c',-45; P,'d',0;  P,'I',70} % inhibition-induced bursting
  };
data=dsSimulate(eqns, 'time_limits',[0 250], 'vary',vary);
dsPlot(data);

% Do the same simulation, but using parfor in order to speed up larger computations
data=dsSimulate(eqns, 'time_limits',[0 250], 'vary',vary, 'parfor_flag',1);

% %%%%%%%%%%%%%%%%%%%%%%%%%%%%%%%%%%%%%%%%%%%%%%%%%%%%%%%%%%%%%%%%%%%%%%%%
%% QUICKLY BUILDING LARGE MODELS FROM EXISTING "MECHANISMS"
% %%%%%%%%%%%%%%%%%%%%%%%%%%%%%%%%%%%%%%%%%%%%%%%%%%%%%%%%%%%%%%%%%%%%%%%%

% Mechanisms are predefined reusable sub-models meant to be incorporated
% in larger complete models. Examples of mechanisms in neuron models include
% ion currents, pumps, etc. Once defined, they can be easily incorporated
% into larger models by simply listing the name of the file containing their
% equations, without the need to re-write any of the mechanism equations.
% This greatly simplifies large model prototyping and re-configuration.

% Hodgkin-Huxley neuron equations (without predefined mechanisms)
eqns={
  'gNa=120; gK=36; Cm=1'
  'INa(v,m,h) = gNa.*m.^3.*h.*(v-50)'
  'IK(v,n) = gK.*n.^4.*(v+77)'
  'dv/dt = (10-INa(v,m,h)-IK(v,n))/Cm; v(0)=-65'
  'dm/dt = aM(v).*(1-m)-bM(v).*m; m(0)=.1'
  'dh/dt = aH(v).*(1-h)-bH(v).*h; h(0)=.1'
  'dn/dt = aN(v).*(1-n)-bN(v).*n; n(0)=0'
  'aM(v) = (2.5-.1*(v+65))./(exp(2.5-.1*(v+65))-1)'
  'bM(v) = 4*exp(-(v+65)/18)'
  'aH(v) = .07*exp(-(v+65)/20)'
  'bH(v) = 1./(exp(3-.1*(v+65))+1)'
  'aN(v) = (.1-.01*(v+65))./(exp(1-.1*(v+65))-1)'
  'bN(v) = .125*exp(-(v+65)/80)'
};
data=dsSimulate(eqns);

figure; plot(data.time,data.(data.labels{1}))
xlabel('time (ms)'); ylabel('membrane potential (mV)'); title('Hodgkin-Huxley neuron')

% Equivalent Hodgkin-Huxley neuron with predefined mechanisms
data=dsSimulate('dv/dt=10+@current/Cm; Cm=1; v(0)=-65; {iNa,iK}');

figure; plot(data.time,data.(data.labels{1}))
xlabel('time (ms)'); ylabel('membrane potential (mV)'); title('Hodgkin-Huxley neuron')

% View the mechanism files:
[~,eqnfile]=dsLocateModelFiles('iNa.mech'); % edit(eqnfile{1});
[~,eqnfile]=dsLocateModelFiles('iK.mech');  % edit(eqnfile{1});
% Mechanisms can be custom built; however, DynaSim does come pakaged with
% some common ones like popular ion currents (see <dynasim>/models).

% Example of a bursting neuron model using three predefined current mechanisms
eqns='dv/dt=Iapp+@current; {iNaF,iKDR,iM}; Iapp=5; gNaF=100; gKDR=5; gM=1.5; v(0)=-70';
data=dsSimulate(eqns, 'time_limits',[0 200]);

figure; plot(data.time,data.(data.labels{1})) % <-- Figure 3 in DynaSim paper
xlabel('time (ms)'); ylabel('membrane potential (mV)'); title('Intrinsically Bursting neuron using predefined mechanisms')

% Example of the same bursting neuron using a predefined population
eqns='IB'; % file name of predefined model of intrinsically bursting (IB) neuron
data=dsSimulate(eqns, 'vary',{'IB','Iapp',5}, 'time_limits',[0 200]);

figure; plot(data.time,data.(data.labels{1}))
xlabel('time (ms)'); ylabel('membrane potential (mV)'); title('Predefined Intrinsically Bursting neuron')

% Predefined populations are stored in text files (e.g., 'IB.pop') and
% simulated by passing the file name to dsSimulate (e.g., dsSimulate('IB'))
% or by equating population equations to it in the DynaSim specification
% structure (see below).

% View the predefined population file:
[~,eqnfile]=dsLocateModelFiles('IB.pop'); % edit(eqnfile{1});

% %%%%%%%%%%%%%%%%%%%%%%%%%%%%%%%%%%%%%%%%%%%%%%%%%%%%%%%%%%%%%%%%%%%%%%%%
%% BUILDING LARGE MODELS WITH MULTIPLE POPULATIONS AND CONNECTIONS
% %%%%%%%%%%%%%%%%%%%%%%%%%%%%%%%%%%%%%%%%%%%%%%%%%%%%%%%%%%%%%%%%%%%%%%%%

%% Setup Sparse Pyramidal-Interneuron-Network-Gamma (sPING)

% define equations of cell model (same for E and I populations)
eqns={
  'dv/dt=Iapp+@current+noise*randn(1,N_pop); Iapp=0; noise=0'
  'monitor iGABAa.functions, iAMPA.functions'
};
% Tip: monitor all functions of a mechanism using: monitor MECHANISM.functions

% create DynaSim specification structure
s=[];
s.populations(1).name='E';
s.populations(1).size=80;
s.populations(1).equations=eqns;
s.populations(1).mechanism_list={'iNa','iK'};
s.populations(1).parameters={'Iapp',5,'gNa',120,'gK',36,'noise',40};
s.populations(2).name='I';
s.populations(2).size=20;
s.populations(2).equations=eqns;
s.populations(2).mechanism_list={'iNa','iK'};
s.populations(2).parameters={'Iapp',0,'gNa',120,'gK',36,'noise',10};
s.connections(1).direction='I->E';
s.connections(1).mechanism_list={'iGABAa'};
s.connections(1).parameters={'tauD',10,'gSYN',.1,'netcon','ones(N_pre,N_post)'}; % connectivity matrix defined using a string that evalutes to a numeric matrix
s.connections(2).direction='E->I';
s.connections(2).mechanism_list={'iAMPA'};
s.connections(2).parameters={'tauD',2,'gSYN',.1,'netcon',ones(80,20)}; % connectivity set using a numeric matrix defined in script

%% Simulate Sparse Pyramidal-Interneuron-Network-Gamma (sPING)
data=dsSimulate(s);

dsPlot(data); % <-- Figure 4 in DynaSim paper
dsPlot(data,'variable',{'E_v','E_I_iGABAa_ISYN'});

% View the connection mechanism file:
[~,eqnfile]=dsLocateModelFiles('iAMPA.mech'); % edit(eqnfile{1});

%% Explore sPING in DynaSim GUI

dynasim(s); % Display model "s" in the DynaSim GUI

% Notes:
% - DynaSim GUI is only supported in MATLAB at this time.
% - Launching the GUI without a model (i.e., by executing "dynasim")
%   will load a default demo network.

% %%%%%%%%%%%%%%%%%%%%%%%%%%%%%%%%%%%%%%%%%%%%%%%%%%%%%%%%%%%%%%%%%%%%%%%%
%% SAVING SIMULATED DATA
% %%%%%%%%%%%%%%%%%%%%%%%%%%%%%%%%%%%%%%%%%%%%%%%%%%%%%%%%%%%%%%%%%%%%%%%%
% How to: set 'save_data_flag' to 1 and (optionally) 'study_dir' to /path/to/outputs

%% Save data from a single simulation

% Example using the previous sPING model:
data=dsSimulate(s,'save_data_flag',1,'study_dir','demo_sPING_1');

% Tip for saving large data files:
% - By default data is saved in compatible mode between Matlab and Octave ('matCompatibility_flag' set to 1, i.e., data is saved in '-v7' mat format).
%   - Unfortunately, '-v7' mat format is not able to save variables > 2GB.
%   - If compatible saving fails, data is saved in '-v7.3' format (Matlab), or in '-hdf5' format (Octave). This allows that data can be stored in all its integrity.
% - However, if any var > 2GB is anticipated, we strongly recommend to set 'matCompatibility_flag' manually to 0 to speed up data saving.
% Example using 'matCompatibility_flag' set to 0:
data=dsSimulate(s,'matCompatibility_flag',0,'save_data_flag',1,'study_dir','demo_sPING_1_varlt2GB');

%% Save data from a set of simulations

% Specify what to vary
% Tip: use 'vary' Syntax 2 to systematically vary a parameter
vary={'E','Iapp',[0 10 20]}; % vary the amplitude of tonic input to E-cells
data=dsSimulate(s, 'save_data_flag',1, 'study_dir','demo_sPING_2',...
                     'vary',vary);

% load and plot the saved data
data_from_disk = dsImport('demo_sPING_2');
dsPlot(data_from_disk);
dsPlot(data_from_disk,'variable','E_v');

% Vary a connection parameter
vary={'I->E','tauD',[5 10 15]}; % inhibition decay time constant from I to E

% Vary two parameters (run a simulation for all combinations of values)
vary={
  'E'   ,'Iapp',[0 10 20];      % amplitude of tonic input to E-cells
  'I->E','tauD',[5 10 15]       % inhibition decay time constant from I to E
  };
dsSimulate(s, 'save_data_flag', 1, 'study_dir', 'demo_sPING_3',...
                'vary', vary, 'verbose_flag', 0, 'parfor_flag',1);
data=dsImport('demo_sPING_3');
dsPlot(data);
dsPlot(data,'plot_type','rastergram'); % <-- Figure 5 in DynaSim paper
dsPlotFR(data); % examine how mean firing rate changes with Iapp and tauD

dsPlot(data,'plot_type','power');

% %%%%%%%%%%%%%%%%%%%%%%%%%%%%%%%%%%%%%%%%%%%%%%%%%%%%%%%%%%%%%%%%%%%%%%%%
%% RUNNING SIMULATIONS ON THE CLUSTER
% %%%%%%%%%%%%%%%%%%%%%%%%%%%%%%%%%%%%%%%%%%%%%%%%%%%%%%%%%%%%%%%%%%%%%%%%
% How to: set 'cluster_flag' to 1
% Requirement: you must be logged on to a cluster that recognizes 'qsub'

if 1
  % Execute the following cluster examples from a login node

  % Run three simulations in parallel jobs and save the simulated data
  eqns='dv/dt=@current+I; {iNa,iK}';
  vary={'','I',[0 10 20]};
  dsSimulate(eqns, 'save_data_flag',1, 'study_dir','demo_cluster_1',...
                     'vary',vary, 'cluster_flag',1, 'overwrite_flag',1, 'verbose_flag',0);
  % tips for checking job status:
  % !qstat -u <YOUR_USERNAME>
  % !cat ~/batchdirs/demo_cluster_1/pbsout/sim_job1.out
  % data=dsImport('demo_cluster_1');
  % dsPlot(data);

  % Repeat but also save plotted data
  eqns='dv/dt=@current+I; {iNa,iK}';
  vary={'','I',[0 10 20]};
  dsSimulate(eqns, 'save_data_flag',1, 'study_dir','demo_cluster_2',...
                     'vary',vary, 'cluster_flag',1, 'overwrite_flag',1, 'verbose_flag',0,...
                     'plot_functions',@dsPlot);
  % !cat ~/batchdirs/demo_cluster_2/pbsout/sim_job1.out

  % Save multiple plots and pass custom options to each plotting function
  eqns='dv/dt=@current+I; {iNa,iK}';
  vary={'','I',[0 10 20]};
  dsSimulate(eqns, 'save_data_flag',1, 'study_dir','demo_cluster_3',...
                     'vary',vary, 'cluster_flag',1, 'overwrite_flag',1, 'verbose_flag',0,...
                     'plot_functions',{@dsPlot,@dsPlot},...
                     'plot_options',{{},{'plot_type','power'}});
  % !cat ~/batchdirs/demo_cluster_3/pbsout/sim_job1.out

  % Post-simulation analyses can be performed similarly by passing
  % analysis function handles and options using 'analysis_functions' and
  % 'analysis_options'.

  % Note: options will be passed to plot and analysis functions in the order
  % given. You can pass handles and options for any built-in, pre-packaged,
  % or custom functions.

  % Run on cluster with compilation
  dsSimulate(eqns, 'save_data_flag',1, 'study_dir','demo_cluster_4','mex_flag',1,...
                     'vary',vary, 'cluster_flag',1, 'overwrite_flag',1, 'verbose_flag',0);
end

% %%%%%%%%%%%%%%%%%%%%%%%%%%%%%%%%%%%%%%%%%%%%%%%%%%%%%%%%%%%%%%%%%%%%%%%%
%% MORE FEATURES
% %%%%%%%%%%%%%%%%%%%%%%%%%%%%%%%%%%%%%%%%%%%%%%%%%%%%%%%%%%%%%%%%%%%%%%%%

% Simulating large models can be sped up significantly by compiling the
% simulation before running it. DynaSim makes this easy to do using the
% 'mex_flag' option in dsSimulate. Note: compiling the model can
% take several seconds to minutes; however, it only compiles the first time
% it is run and is significantly faster on subsequent runs.

data=dsSimulate(s, 'mex_flag',1, 'study_dir','demo_sPING_3_compile');
dsPlot(data);

% Now run again:
data=dsSimulate(s, 'mex_flag',1, 'study_dir','demo_sPING_3_compile');
dsPlot(data);

% Combine compilation and parallelization to maximize computational speed locally
vary={'E','Iapp',[0 10 20]};
data=dsSimulate(s, 'mex_flag',1, 'parfor_flag',1, 'vary', vary, 'study_dir','demo_sPING_3_compile_parallel');
dsPlot(data);

% %%%%%%%%%%%%%%%%%%%%%%%%%%%%%%%%%%%%%%%%%%%%%%%%%%%%%%%%%%%%%%%%%%%%%%%%
% Script-based modeling using predefined objects without writing equations.
% swapping out population models:

T=[0 200]; % ms, [beg end], simulation time limits

s=[];
s.populations.size=5;
s.populations.mechanism_list={'stim','noise'};

% Classic Hodgkin-Huxley neurons
s.populations.equations='HH';
s.populations.parameters={'stim_amp',10,'noise_amp',1e3};
dsPlot(dsSimulate(s,'time_limits',T));

% Intrinsically-Bursting neurons
s.populations.equations='IB';
s.populations.parameters={'stim_amp',5,'noise_amp',1e3};
dsPlot(dsSimulate(s,'time_limits',T));

% Morris-Lecar neurons
s.populations.equations='ML';
s.populations.parameters={'stim_amp',100,'noise_amp',1e3};
dsPlot(dsSimulate(s,'time_limits',T));

% FitzHugh-Nagumo neurons
s.populations.equations='FHN';
s.populations.parameters={'stim_amp',.5,'noise_amp',10};
dsPlot(dsSimulate(s,'time_limits',T));

% Leaky integrate-and-fire neurons
s.populations.equations='LIF';
s.populations.parameters={'stim_amp',10,'noise_amp',1e3};
dsPlot(dsSimulate(s,'time_limits',T));

% Izhikevich neurons
s.populations.equations='Izh';
s.populations.parameters={'stim_amp',200,'noise_amp',2e4};
dsPlot(dsSimulate(s,'time_limits',T));

% Wilson-Cowan oscillators
s.populations.equations='WC';
s.populations.parameters={'stim_amp',0,'noise_amp',10};
dsPlot(dsSimulate(s,'time_limits',T));

% Regular Spiking (RS) neuron from Kramer 2008:
s.populations.equations='RS';
s.populations.parameters={'stim_amp',5,'noise_amp',1e3};
dsPlot(dsSimulate(s,'time_limits',T));

% Layer 2/3 somatosensory RS/FS network (adapted from Kramer 2008):
% Note that the entire model is specified without writing any equations!
s=[];
s.populations(1).equations='RS';              % excitatory (E) population
s.populations(1).mechanism_list={'stim'};     % tonic stimulation
s.populations(1).parameters={'stim_amp',10};  % amplitude of stimulation
s.populations(2).equations='FS';              % inhibitory (I) population
s.connections(1).direction='RS->FS';          % E->I connection
s.connections(1).mechanism_list='iAMPA';      % AMPA synapse
s.connections(1).parameters={'gSYN',.5};      % synaptic weight
s.connections(2).direction='FS->RS';          % I->E connection
s.connections(2).mechanism_list='iGABAa';     % GABAa synapse
s.connections(2).parameters={'gSYN',1,'tauD',10}; % strength and time constant of inhibition
dsPlot(dsSimulate(s,'time_limits',T));

%% Multicompartment neurons

% Compartments can be specified using the "populations" field or the
% "compartments" field; the latter is an alias for the former; it was
% added for conceptual clarity in the model specification.

% Excitatory cell S driving two-comparmtent E-cell
s=[];
s.populations(1).name='S';
s.populations(1).equations='dv/dt=@current+10;{iNa,iK}';
s.compartments(1).name='Edend';
s.compartments(1).equations='dv/dt=@current;{iNa,iK}';
s.compartments(2).name='Esoma';
s.compartments(2).equations='dv/dt=@current;{iNa,iK}';
s.connections(1).direction='S->Edend';
s.connections(1).mechanism_list='iAMPA';
s.connections(2).direction='Edend->Esoma';
s.connections(2).mechanism_list='iCOM';

d=dsSimulate(s);
dsPlot(d);

% For more examples that include compartmental dimensions/geometry, see
% dynasim/demos/examples/Multicompartment_PFC_neurons

%% Networks of integrate-and-fire neurons with axonal delays and refractory period
% tau [ms]: membrane time constant (RC)
% tref [ms]: absolute refractory period
% delay [ms]: axonal delay
% tspike: reserved variable for storing past spike times when monitoring spikes

% 1 E-cell driving 1 I-cell
LIF={
    'dV/dt=(E-V+R*I+noise*randn-@isyn)/tau; V(0)=-65'
    'if(any(t<tspike+tref,1))(V=reset)'
    'tau=10; tref=10; E=-70; thresh=-55; reset=-75; R=9; I=1.55; noise=100'
    'monitor V.spikes(thresh)'
     };

iampa={
  'gSYN=.5; ESYN=0; tauD=2; tauR=0.4; delay=15'
  'f(x) = (exp(-x/tauD)-exp(-x/tauR)).*(x>0)'
  'Isyn(V) = gSYN.*sum(f(t-tspike_pre-delay)).*(V-ESYN)'
  '@isyn += Isyn(V_post)'
};

s=[];
s.populations(1).name='E';
s.populations(1).equations=LIF;
s.populations(2).name='I';
s.populations(2).equations=LIF;
s.populations(2).parameters={'I',0,'noise',0};
s.connections(1).direction='E->I';
s.connections(1).mechanism_list='iampa';
s.mechanisms(1).name='iampa';
s.mechanisms(1).equations=iampa;

data=dsSimulate(s,'time_limits',[0 200],'solver','rk1','dt',.01);
dsPlot(data);

% For more examples, including STDP, see: dynasim/demos/dsLIFnetwork.m

%% Delay differential equations (e.g., axonal delays in network of HH neurons)
% This example demonstrates (1) creating delay differential  equations with
% X_pre(t-delay), and (2) defining mechanisms (e.g., iampa) in the same
% script as the full model specification  and storing them using
% the specification.mechanisms field.

ampa_with_delay={
  'gSYN=.1; ESYN=0; tauD=2; tauR=0.2; delay=20'
  'netcon=ones(N_pre,N_post)'
  'ISYN(X,s)=gSYN.*(s*netcon).*(X-ESYN)'
  'ds/dt=-s./tauD+1/2*(1+tanh(X_pre(t-delay)/10)).*((1-s)/tauR); s(0)=.1' % 20ms delay
  '@current += -ISYN(X_post,s)'
};

s=[];
s.populations(1).name='HH';
s.populations(1).equations='dV/dt=@current+10*(t<50);{iNa,iK};V(0)=-65';
s.connections(1).direction='HH->HH';
s.connections(1).mechanism_list='iampa';
s.connections(1).parameters={'gSYN',.1};
s.mechanisms(1).name='iampa';
s.mechanisms(1).equations=ampa_with_delay;
data=dsSimulate(s,'time_limits',[0 100]);
figure; plot(data.time,data.HH_V);

%% Poisson synaptic inputs

eqns='dV/dt=@current; {iNa,iK,iPoisson}; V(0)=-65; DC=2000; gext=.01';
data=dsSimulate(eqns,'time_limits',[0 1000]);
dsPlot(data,'variable',{'V','gPoisson'});

% REFERENCE (default iPoisson parameters)
% % poisson parameters
% baseline=0;     % sp/s, baseline rate
% DC=2000;        % sp/s, steady component of the signal
% AC=0;           % sp/s, oscillatory component of the signal
% f=0;            % Hz, modulation frequency of the signal
% phi=0;          % radians, phase at which the signal begins
% onset=0;        % ms, start time of signal
% offset=inf;     % ms, stop time of signal
%
% % synaptic parameters
% gext=.01;       % max synaptic conductance
% Eext=0;         % mV, synaptic reversal potential
% tau=2;          % ms, synaptic time constant

%% Tips for efficient simulation

downsample_factor=10; % downsampling saves time by recording fewer time points
mex_flag=1;       % takes longer to compile on 1st run; faster on subsequent runs
                      % Note: compilation is most beneficial when Npop>1.
solver='euler';       % Euler integration requires fewer calculations than 4th-order Runge Kutta
dt=.01;               % increase time step as long as solution converges

eqns='dv/dt=@current+I; {iNa,iK}';

% run and compile MEX file
data=dsSimulate(eqns,'vary',{'I',10},'downsample_factor',downsample_factor,...
  'mex_flag',mex_flag,'solver',solver,'dt',dt);
dsPlot(data);

% sets of simulations:
% multinode (on a cluster): cluster_flag=1;
% multicore (local simulation): parfor_flag=1;
```
