## Supplemental File 1 for "A biophysical model of striatal microcircuits suggests delta/theta-rhythmically interleaved gamma and beta oscillations mediate periodicity in motor control": dsDemos_cluster_modes.m

```
% The tests some of the major cluster options

%% Setup

% Add DynaSim to path if it's not already there
if exist('setupDynaSimPath','file')
    setupDynaSimPath;
else
    error('Add the DynaSim folder to the MATLAB path - e.g. run addpath(genpath(DynaSimPath))');
end

% Set where to save outputs
output_directory = dsGetConfig('demos_path');

% move to root directory where outputs will be saved
mkdirSilent(output_directory);
cd(output_directory);

%% model
eqns='dv/dt = @current+I; {iNa,iK}';
vary={'','I',0:10:30};

%% options
qsub_modes = {'loop', 'array'};
% mex_flag = [0,1];
% one_solve_file_flag = [0,1];
% parfor_flag = [0,1];

N = 2;
v = (1:2^(N^2))-1;
opts = dec2bin(v)' - '0';
nOpt = size(opts,2);
for iOpt = 1:nOpt
  thisOpt = opts(:, iOpt);

  % get opt values
  qsub_mode = qsub_modes{thisOpt(1)+1};
  mex_flag = thisOpt(2);
  parfor_flag = thisOpt(3);
  one_solve_file_flag = thisOpt(4);

  % print
  fprintf('opt: %i/%i \n  qsub_mode:%s \n  mex_flag:%i \n  parfor_flag:%i \n  one_solve_file_flag:%i \n\n', iOpt, nOpt, qsub_mode, mex_flag, parfor_flag, one_solve_file_flag)

  if parfor_flag
    num_cores = 2;
  else
    num_cores = 1;
  end

  % make dir name
  study_dir = fullfile(output_directory, sprintf('study_HH_varyI_cluster_%s',qsub_mode));
  if mex_flag
    study_dir = [study_dir '_comp'];
  end
  if parfor_flag
    study_dir = [study_dir '_par'];
  end
  if one_solve_file_flag
    study_dir = [study_dir '_oneFile'];
  end

  dsSimulate(eqns,'vary',vary, 'study_dir',study_dir,'save_data_flag',1,...
  'cluster_flag',1,'verbose_flag',1,'qsub_mode',qsub_mode, 'mex_flag',mex_flag,...
  'one_solve_file_flag',one_solve_file_flag, 'parfor_flag',parfor_flag, 'num_cores',num_cores);

  % print hr
  fprintf('_____________________________________________________________________ \n')
end
```
