## Supplemental File 1 for "A biophysical model of striatal microcircuits suggests delta/theta-rhythmically interleaved gamma and beta oscillations mediate periodicity in motor control": dsDemos_covar_temp.m

```
% Get ready...

% Add DynaSim to path if it's not already there
if exist('setupDynaSimPath','file')
    setupDynaSimPath;
else
    error('Add the DynaSim folder to the MATLAB path - e.g. run addpath(genpath(DynaSimPath))');
end

% Set where to save outputs
output_directory = dsGetConfig('demos_path');

% move to root directory where outputs will be saved
mkdirSilent(output_directory);
cd(output_directory);

%% Sparse Pyramidal-Interneuron-Network-Gamma (sPING)

% define equations of cell model (same for E and I populations)
eqns={ 
  'dv/dt=Iapp+@current+noise*randn(1,N_pop)';
  'monitor iGABAa.functions, iAMPA.functions'
};
% Tip: monitor all functions of a mechanism using: monitor MECHANISM.functions

% create DynaSim specification structure
s=[];
s.populations(1).name='E';
s.populations(1).size=80;
s.populations(1).equations=eqns;
s.populations(1).mechanism_list={'iNa','iK'};
s.populations(1).parameters={'Iapp',5,'gNa',120,'gK',36,'noise',40};
s.populations(2).name='I';
s.populations(2).size=20;
s.populations(2).equations=eqns;
s.populations(2).mechanism_list={'iNa','iK'};
s.populations(2).parameters={'Iapp',0,'gNa',120,'gK',36,'noise',40};
s.connections(1).direction='I->E';
s.connections(1).mechanism_list={'iGABAa'};
s.connections(1).parameters={'tauD',10,'gSYN',.1,'netcon','ones(N_pre,N_post)'};
s.connections(2).direction='E->I';
s.connections(2).mechanism_list={'iAMPA'};
s.connections(2).parameters={'tauD',2,'gSYN',.1,'netcon',ones(80,20)};

% %%%%%%%%%%%%%%%%%%%%%%%%%%%%%%%%%%%%%%%%%%%%%%%%%%%%%%%%%%%%%%%%%%%%%%%%
% SAVING SIMULATED DATA
% %%%%%%%%%%%%%%%%%%%%%%%%%%%%%%%%%%%%%%%%%%%%%%%%%%%%%%%%%%%%%%%%%%%%%%%%
% How to: set 'save_data_flag' to 1 and (optionally) 'study_dir' to /path/to/outputs

% Save data from a single simulation
% Example using the previous sPING model:

% Save data from a set of simulations

% Specify what to vary 

% Vary a connection parameter

% Vary two parameters (run a simulation for all combinations of values)
% vary={'(E,I)','(EK1,EK2)',[-80 -60]}; %check
%       This sets modifications:
%           * E_EK1, E_EK2, I_EK1, I_EK2 = -80
%           * E_EK1, E_EK2, I_EK1, I_EK2 = -60
% vary={'(E,I)','(EK1,EK2)',[-80 -60; -85 -65]}; %check
%       This sets modifications:
%           * E_EK1, I_EK1 = -80 and E_EK2, I_EK2 = -85
%           * E_EK1, I_EK1 = -60 and E_EK2, I_EK2 = -65
% vary={'(E,I)','(EK1,EK2)',cat(3,[-80 -60], [-85 -65])}; %check
%       This sets modifications:
%           * E_EK1, E_EK2 = -80 and I_EK1, I_EK2 = -85
%           * E_EK1, E_EK2 = -60 and I_EK1, I_EK2 = -65
% vary={'(E,I)','(EK1,EK2)',cat(3, [-75 -55; -80 -60], [-85 -65; -90 -70])}; %check
%       This sets modifications:
%           * E_EK1 = -75, E_EK2 = -80, I_EK1 = -85, I_EK2 = -90.
%           * E_EK1 = -55, E_EK2 = -60, I_EK1 = -65, I_EK2 = -70.
dsSimulate(s,'save_data_flag',0,'study_dir','demo_sPING_covary',...
                'vary',vary,'verbose_flag',1);
```
