## Supplemental File 1 for "A biophysical model of striatal microcircuits suggests delta/theta-rhythmically interleaved gamma and beta oscillations mediate periodicity in motor control": dsDemos_generate_data.m

```
%% Generate data
% This is a script for generating small sets of dat for use in testing. It
% saves the data in .mat files, which seems to be a bit more efficient than
% saving as a DynaSim path


%% Set up paths 

% Add DynaSim to path if it's not already there
if exist('setupDynaSimPath','file')
    setupDynaSimPath;
else
    error('Add the DynaSim folder to the MATLAB path - e.g. run addpath(genpath(DynaSimPath))');
end

% Set where to save outputs
output_directory = dsGetConfig('demos_path');

% move to root directory where outputs will be saved
mkdirSilent(output_directory);


%% Generate a sample dataset
% Run simulation - Sparse Pyramidal-Interneuron-Network-Gamma (sPING)

% define equations of cell model (same for E and I populations)
eqns={
  'dv/dt=Iapp+@current+noise*randn(1,N_pop); Iapp=0; noise=0'
  'monitor iGABAa.functions, iAMPA.functions'
};
% Tip: monitor all functions of a mechanism using: monitor MECHANISM.functions

% create DynaSim specification structure
s=[];
s.populations(1).name='E';
s.populations(1).size=80;
s.populations(1).equations=eqns;
s.populations(1).mechanism_list={'iNa','iK'};
s.populations(1).parameters={'Iapp',5,'gNa',120,'gK',36,'noise',40};
s.populations(2).name='I';
s.populations(2).size=20;
s.populations(2).equations=eqns;
s.populations(2).mechanism_list={'iNa','iK'};
s.populations(2).parameters={'Iapp',0,'gNa',120,'gK',36,'noise',10};
s.connections(1).direction='I->E';
s.connections(1).mechanism_list={'iGABAa'};
s.connections(1).parameters={'tauD',10,'gGABAa',.1,'netcon','ones(N_pre,N_post)'}; % connectivity matrix defined using a string that evalutes to a numeric matrix
s.connections(2).direction='E->I';
s.connections(2).mechanism_list={'iAMPA'};
s.connections(2).parameters={'tauD',2,'gAMPA',.1,'netcon',ones(80,20)}; % connectivity set using a numeric matrix defined in script

% % Vary two parameters (run a simulation for all combinations of values)
% vary={
%   'E'   ,'Iapp',[0:10:10];      % amplitude of tonic input to E-cells
%   %'I'   ,'Iapp',[0 5 10];      % amplitude of tonic input to E-cells
%   'I->E','tauD',[5:5:5]       % inhibition decay time constant from I to E
%   };

% Vary two parameters (run a simulation for all combinations of values)
vary={
  'E'   ,'Iapp',[1:5];      % amplitude of tonic input to E-cells
  'I->E','tauD',[10]       % inhibition decay time constant from I to E
  };

study_dir = fullfile(output_directory,'demo_sPING_100cells_5x1');

dsSimulate(s,'save_data_flag',1,'study_dir',study_dir,...
                'vary',vary,'verbose_flag',1, 'downsample_factor', 10, ...
                'save_results_flag',1,'parfor_flag',1,'plot_functions',{@dsPlot,@dsPlot,@dsPlotFR2},'plot_options',{{'format','png','visible','off','figwidth',0.5,'figheight',0.5}, ...
                {'format','png','visible','off','plot_type','rastergram','figwidth',0.5,'figheight',0.5},...
                {'format','png','visible','off'}} );
            
zipfname = dsZipDemoData(study_dir);

%% Generate another sample dataset

% Vary two parameters (run a simulation for all combinations of values)
vary={
  'E'   ,'Iapp',[0:10:20];      % amplitude of tonic input to E-cells
  %'I'   ,'Iapp',[0 5 10];      % amplitude of tonic input to I-cells
  'I->E','tauD',[5:5:15]       % inhibition decay time constant from I to E
  };

study_dir = fullfile(output_directory,'demo_sPING_100cells_3x3');

dsSimulate(s,'save_data_flag',1,'study_dir',study_dir,...
                'vary',vary,'verbose_flag',1, 'downsample_factor', 10, ...
                'save_results_flag',1,'parfor_flag',1,'plot_functions',{@dsPlot,@dsPlot,@dsPlotFR2},'plot_options',{{'format','png','visible','off','figwidth',0.5,'figheight',0.5}, ...
                {'format','png','visible','off','plot_type','rastergram','figwidth',0.5,'figheight',0.5},...
                {'format','png','visible','off'}} );

zipfname = dsZipDemoData(study_dir);

%% ********************************** Leftover stuff not used! ********************************** 

%% Load the data

% Load data in traditional DynaSim format
data=dsImport(study_dir);

% Import plot files
data_img = dsImportPlots(study_dir);


%% Downsample number of cells if necessary
% (For saving space)

% Convert to MDD object
xp = ds2mdd(data);

% Num_cells_to_keep = 20;
downsample_factor = 2;
mydata = xp.data;

for i = 1:numel(mydata)
    if ~isempty(mydata{i})
        %mydata{i} = mydata{i}(:,1:Num_cells_to_keep);
        mydata{i} = mydata{i}(:,1:round(size(mydata{i},2)/downsample_factor));
    end
end

xp.data = mydata;

data = ds.MDD2ds(xp);

%% Package up the data

% Stores it in the DynaSim demos archive for adding to repo
zipfname = dsZipDemoData(study_dir);
% (can later recover this by running: dsUnzipDemoData(study_dir);)

%% 


% Load into DynaSim structure
[data_table,column_titles] = dsDataField2Table (data_img,'plot_files');

% Preview the contents of this table
previewTable(data_table,column_titles);

% The entries in the first column contain the paths to the figure files.
% There can be multiple figures associated with each simulation, which is
% why these are cell arrays of strings.
disp(data_table{1}{1})
disp(data_table{1}{2})

% Import the linear data into an MDD object
xp_img = MDD;
X = data_table{1}; axislabels = data_table(2:end);
xp_img = xp_img.importDataTable(X, axislabels);
xp_img = xp_img.importAxisNames(column_titles(2:end));
```
