## Supplemental File 1 for "A biophysical model of striatal microcircuits suggests delta/theta-rhythmically interleaved gamma and beta oscillations mediate periodicity in motor control": dsDemos_MDD.m

```
%% % % % % % % % % % % % % % % DEMO - Using MDD with DynaSim % % % % % %
% % % % % % % % % % % % % % % % % % % % % % % % % % % % % % % % % % % % % 

%% Set up paths 
% Get ready...

% Format
format compact
% restoredefaultpath

% Add DynaSim to path if it's not already there
if exist('setupDynaSimPath','file')
    setupDynaSimPath;
else
    error('Add the DynaSim folder to the MATLAB path - e.g. run addpath(genpath(DynaSimPath))');
end

% Set where to save outputs
output_directory = dsGetConfig('demos_path');

% Set where to save outputs
study_dir = fullfile(output_directory,'demo_sPING_100cells_3x3');
mkdirSilent(output_directory);


%% Import the data

% Make sure sample data exists; if not copy it into place
if ~exist(study_dir,'dir')
    dsUnzipDemoData(study_dir);
end

% Load data in traditional DynaSim format
data=dsImport(study_dir);


%% Convert it into an MDD object

% Extract the data in a linear table format
[data_table,column_titles,time] = dsData2Table (data);

% Preview the contents of this table
%     Note: We cannot make this one big cell array since we want to allow
%     axis labels to be either strings or numerics.
previewTable(data_table,column_titles);

% Import the linear data into an MDD object
xp = MDD;
data_column = data_table{1};                          % X holds the data that will populate the multidimensional array. Must be numeric or cell array.
axis_val_columns = data_table(2:end);                 % Each entry in X has an associated set of axis labels, which will define its location in multidimensional space. **Must be numeric or cell array of chars only**
xp = xp.importDataTable(data_column,axis_val_columns);
xp = xp.importAxisNames(column_titles(2:end));  % There should be 1 axis name for every axis, of type char.

% xp.meta stores meta data for use by the user as they see fit.
% Here we will add some custom info to xp.metadata. This can be whatever
% you want. Here, I will use this to provide information about what is
% stored in each of the matrices in the xp.data cell array. (Alternatively,
% we could also make each of these matrices an MDD object!)
meta = struct;
meta.datainfo(1:2) = MDDAxis;
meta.datainfo(1).name = 'time(ms)';
meta.datainfo(1).values = time;
meta.datainfo(2).name = 'cells';
meta.datainfo(2).values = [];
meta.dynasim.labels = data.labels;
meta.dynasim.model = data.model;
meta.dynasim.simulator_options = data.simulator_options;
meta.dynasim.time = data.time;
meta.dynasim.varied = data.varied;
xp.meta = meta;
clear meta

%% Take the full MDD tutorial
% Run the following code to be re-directed to a more indepth tutorial on
% using MDD. Otherwise, continue onward to a few plotting examples.

str = which('tutorial_MDD');
cd(fileparts(str))
edit tutorial_MDD.m

%% Plot 2D data
% Tip: don't try to understand what recursivePlot is doing - instead, try
% putting break points in the various function handles to see how this
% command works.
close all;

% Pull out a 2D subset of the data
clc
xp4 = xp(:,:,'E','v');
xp4.printAxisInfo

% Set up plotting arguments
function_handles = {@xp_subplot_grid,@xp_matrix_basicplot};   % Specifies the handles of the plotting functions
dimensions = {{'E_Iapp','I_E_tauD'},{'data'}};                % Specifies which axes of xp each function handle
                                                                % will operate on. Note that dimension 'data' refers to the 
                                                                % the contents of each element in xp.data (e.g. the matrix of
                                                                % time series data). It must come last.
function_arguments = {{},{}};	% This allows you to supply input arguments to each of the 
                                % functions in function handles. For
                                % now we'll leave this empty.
                                                                
% Run the plot. Note the "+" icons next to each plot allow zooming. 
figl; recursivePlot(xp4,function_handles,dimensions,function_arguments);


%% Load saved .png images rather than raw data
close all;

% Import plot files
data_img = dsImportPlots(study_dir);

% Load into DynaSim structure
[data_table,column_titles] = dsDataField2Table (data_img,'plot_files');

% Preview the contents of this table
previewTable(data_table,column_titles);

% The entries in the first column contain the paths to the figure files.
% There can be multiple figures associated with each simulation, which is
% why these are cell arrays of strings.
disp(data_table{1}{1})
disp(data_table{1}{2})

% Import the linear data into an MDD object
xp_img = MDD;
data_column = data_table{1}; axis_val_columns = data_table(2:end);
xp_img = xp_img.importDataTable(data_column, axis_val_columns);
xp_img = xp_img.importAxisNames(column_titles(2:end));

%% Plot the saved .png images

dimensions = {[1,2],[0]};
plotimage_options.scale = 0.5;
func_arguments = {{},{plotimage_options}};         % The 0.5 argument tells xp_plotimage to
                                                   % scale down the resolution of its
                                                   % plots by 0.5. This increases speed.

figl; recursivePlot(xp_img,{@xp_subplot_grid,@xp_plotimage},dimensions,func_arguments);


%% Use mergeDims to convert to Jason's DynaSim format
% Analogous to Reshape

% This combines the 2D "vary" sweep into a single dimension. It also
% combines all populations and variables into a single 1D list. Thus, Axis
% 1 is equivalent to Jason's structure array - data(1:9). Axis 4 is
% equivalent to the structure fields in Jason's DynaSim structure.
xp2 = xp.mergeDims([3,4]);
xp2 = xp2.mergeDims([1,2]);
xp3 = squeeze(xp2); % Squeeze out the empty dimensions.
xp3.printAxisInfo;


% Note that the variable names are not sorted the same was as in Jason's
% DynaSim structure, in that they alternate E and I cells
disp(xp3.axis(2).values);
% Can easily sort them as follows.
[~,I] = sort(xp3.axis(2).values);
xp4 = xp3(:,I);
disp(xp4.axis(2).values);
```
