## Supplemental File 1 for "A biophysical model of striatal microcircuits suggests delta/theta-rhythmically interleaved gamma and beta oscillations mediate periodicity in motor control": dsDemos_mlsolvers_temp.m

```
% %%%%%%%%%%%%%%%%%%%%%%%%%%%%%%%%%%%%%%%%%%%%%%%%%%%%%%%%%%%%%%%%%%%%%%%%
%% DynaSim Demos
% %%%%%%%%%%%%%%%%%%%%%%%%%%%%%%%%%%%%%%%%%%%%%%%%%%%%%%%%%%%%%%%%%%%%%%%%
%{
Download DynaSim toolbox from https://github.com/dynasim/dynasim.
  or, download using git: git clone https://github.com/dynasim/dynasim.git
For further documentation, see tutorial.m in the demos directory
Sign up for user mailing list at: https://groups.google.com/forum/#!forum/dynasim-users.

Tip: In Matlab, you can obtain more information associated with any function "FUNCTION_NAME"
by entering "help FUNCTION_NAME" in the command window. Use the "See also" list
at the end of the help section to browse through related help documentation.
%}

% Add DynaSim to path if it's not already there
if exist('setupDynaSimPath','file')
    setupDynaSimPath;
else
    error('Add the DynaSim folder to the MATLAB path - e.g. run addpath(genpath(DynaSimPath))');
end

% Set where to save outputs
output_directory = dsGetConfig('demos_path');

% move to root directory where outputs will be saved
mkdirSilent(output_directory)
cd(output_directory);

% Here we go!

% %%%%%%%%%%%%%%%%%%%%%%%%%%%%%%%%%%%%%%%%%%%%%%%%%%%%%%%%%%%%%%%%%%%%%%%%
%% DEFINING AND SIMULATING MODELS
% %%%%%%%%%%%%%%%%%%%%%%%%%%%%%%%%%%%%%%%%%%%%%%%%%%%%%%%%%%%%%%%%%%%%%%%%

%% Lorenz equations with phase plot

% DynaSim makes it easy to simulate arbitrary systems of ordinary
% differential equations. Simply write out the system in a cell array of
% strings, separating equations into different strings or the same string
% separated by semicolons.

eqns={
  's=10; r=27; b=2.666';
  'dx/dt=s*(y-x)';
  'dy/dt=r*x-y-x*z';
  'dz/dt=-b*z+x*y';
};
data=dsSimulate(eqns,'tspan',[0 100],'study_dir','lorenz_ml_solver', 'ic',[1 2 .5],'solver','ode45',...
  'verbose_flag',1);
dsPlot(data)

%% QUICKLY BUILDING LARGE MODELS FROM EXISTING "MECHANISMS"

% Mechanisms are predefined reusable sub-models meant to be incorporated 
% in larger complete models. Examples of mechanisms in neuron models include
% ion currents, pumps, etc. Once defined, they can be easily incorporated
% into larger models by simply listing the name of the file containing their
% equations, without the need to re-write any of the mechanism equations.
% This greatly simplifies large model prototyping and re-configuration.

% Hodgkin-Huxley neuron equations (without predefined mechanisms)
eqns={
  'gNa=120; gK=36; Cm=1';
  'INa(v,m,h) = gNa.*m.^3.*h.*(v-50)';
  'IK(v,n) = gK.*n.^4.*(v+77)';
  'dv/dt = (10-INa(v,m,h)-IK(v,n))/Cm; v(0)=-65';
  'dm/dt = aM(v).*(1-m)-bM(v).*m; m(0)=.1';
  'dh/dt = aH(v).*(1-h)-bH(v).*h; h(0)=.1';
  'dn/dt = aN(v).*(1-n)-bN(v).*n; n(0)=0';
  'aM(v) = (2.5-.1*(v+65))./(exp(2.5-.1*(v+65))-1)';
  'bM(v) = 4*exp(-(v+65)/18)';
  'aH(v) = .07*exp(-(v+65)/20)';
  'bH(v) = 1./(exp(3-.1*(v+65))+1)';
  'aN(v) = (.1-.01*(v+65))./(exp(1-.1*(v+65))-1)';
  'bN(v) = .125*exp(-(v+65)/80)';
};
data=dsSimulate(eqns,'dt',0.01, 'downsample_factor',100, 'study_dir','hh_ml_solver', 'solver','ode45',...
  'verbose_flag',1);
figure; plot(data.time,data.(data.labels{1}))
xlabel('time (ms)'); ylabel('membrane potential (mV)'); title('Hodgkin-Huxley neuron')

% % Equivalent Hodgkin-Huxley neuron with predefined mechanisms
% data=dsSimulate('dv/dt=10+@current/Cm; Cm=1; v(0)=-65; {iNa,iK}','study_dir','hh_predef_ml_solver', 'solver','ode45');
% figure; plot(data.time,data.(data.labels{1}))
% xlabel('time (ms)'); ylabel('membrane potential (mV)'); title('Hodgkin-Huxley neuron')
% 
% % Example of a bursting neuron model using three active current mechanisms:
% eqns='dv/dt=5+@current; {iNaF,iKDR,iM}; gNaF=100; gKDR=5; gM=1.5; v(0)=-70';
% data=dsSimulate(eqns,'tspan',[0 200],'study_dir','hh_active_ml_solver', 'solver','ode45');
% figure; plot(data.time,data.(data.labels{1}))
% xlabel('time (ms)'); ylabel('membrane potential (mV)'); title('Intrinsically Bursting neuron')

% %%%%%%%%%%%%%%%%%%%%%%%%%%%%%%%%%%%%%%%%%%%%%%%%%%%%%%%%%%%%%%%%%%%%%%%%
%% BUILDING LARGE MODELS WITH MULTIPLE POPULATIONS AND CONNECTIONS
% %%%%%%%%%%%%%%%%%%%%%%%%%%%%%%%%%%%%%%%%%%%%%%%%%%%%%%%%%%%%%%%%%%%%%%%%

%% Sparse Pyramidal-Interneuron-Network-Gamma (sPING)

% define equations of cell model (same for E and I populations)
eqns={ 
  'dv/dt=Iapp+@current+noise*randn(1,N_pop)';
};
% Tip: monitor all functions of a mechanism using: monitor MECHANISM.functions

% create DynaSim specification structure
s=[];
s.populations(1).name='E';
s.populations(1).size=4;
s.populations(1).equations=eqns;
s.populations(1).mechanism_list={'iNa','iK'};
s.populations(1).parameters={'Iapp',5,'gNa',120,'gK',36,'noise',40};
s.populations(2).name='I';
s.populations(2).size=2;
s.populations(2).equations=eqns;
s.populations(2).mechanism_list={'iNa','iK'};
s.populations(2).parameters={'Iapp',0,'gNa',120,'gK',36,'noise',40};
s.connections(1).direction='I->E';
s.connections(1).mechanism_list={'iGABAa'};
s.connections(1).parameters={'tauD',10,'gSYN',.1,'netcon','ones(N_pre,N_post)'};
s.connections(2).direction='E->I';
s.connections(2).mechanism_list={'iAMPA'};
s.connections(2).parameters={'tauD',2,'gSYN',.1,'netcon','ones(N_pre,N_post)'};

data=dsSimulate(s,'dt',0.01, 'downsample_factor',100, 'study_dir','sPing_ml_solver',...
  'solver','ode45', 'matlab_solver_options', {'InitialStep', 0.01}, 'mex_flag',1,...
  'verbose_flag',1);
dsPlot(data);


%% RUNNING SIMULATIONS ON THE CLUSTER
% %%%%%%%%%%%%%%%%%%%%%%%%%%%%%%%%%%%%%%%%%%%%%%%%%%%%%%%%%%%%%%%%%%%%%%%%
% How to: set 'cluster_flag' to 1
% Requirement: you must be logged on to a cluster that recognizes 'qsub'

% Run three simulations in parallel jobs and save the simulated data
eqns='dv/dt=@current+I; {iNa,iK}';
vary={'','I',[0 10 20]};

% od45
dsSimulate(eqns,'save_data_flag',1,'study_dir','demo_cluster_1_ml_solver_ode45',...
                   'vary',vary,'cluster_flag',1,'overwrite_flag',1,'verbose_flag',1,...
                   'dt',0.01, 'downsample_factor',100,'solver','ode45',...
                    'matlab_solver_options', {'InitialStep', 0.01}, 'mex_flag',0);
  
% ode45 compiled
dsSimulate(eqns,'save_data_flag',1,'study_dir','demo_cluster_1_ml_solver_ode45_compiled',...
                   'vary',vary,'cluster_flag',1,'overwrite_flag',1,'verbose_flag',1,...
                   'dt',0.01, 'downsample_factor',100,'solver','ode45',...
                    'matlab_solver_options', {'InitialStep', 0.01}, 'mex_flag',1);
               
% ode23
dsSimulate(eqns,'save_data_flag',1,'study_dir','demo_cluster_1_ml_solver_ode23s',...
                   'vary',vary,'cluster_flag',1,'overwrite_flag',1,'verbose_flag',1,...
                   'dt',0.01, 'downsample_factor',100,'solver','ode23s',...
                    'matlab_solver_options', {'InitialStep', 0.01}, 'mex_flag',0);
                  
% ode23 compiled
dsSimulate(eqns,'save_data_flag',1,'study_dir','demo_cluster_1_ml_solver_ode23s_compiled',...
                   'vary',vary,'cluster_flag',1,'overwrite_flag',1,'verbose_flag',1,...
                   'dt',0.01, 'downsample_factor',100,'solver','ode23s',...
                    'matlab_solver_options', {'InitialStep', 0.01}, 'mex_flag',1);
```
