## Supplemental File 1 for "A biophysical model of striatal microcircuits suggests delta/theta-rhythmically interleaved gamma and beta oscillations mediate periodicity in motor control": dsDemos_one_file_temp.m

```
%% Setup

% Add DynaSim to path if it's not already there
if exist('setupDynaSimPath','file')
    setupDynaSimPath;
else
    error('Add the DynaSim folder to the MATLAB path - e.g. run addpath(genpath(DynaSimPath))');
end

% Set where to save outputs
output_directory = dsGetConfig('demos_path');

% move to root directory where outputs will be saved
mkdirSilent(output_directory);
cd(output_directory);

%% ***************************************************************
%% ****************************LOCAL******************************
%% ***************************************************************

% %% Solve with one file mode
% study_dir=fullfile('.', 'study_HH_varyI_one_file');
% if exist(study_dir, 'dir')
%   rmdir(study_dir, 's')
% end
%
% eqns='dv/dt=@current+I; {iNa,iK}';
% vary={'','I',[0:10:30]};
%
% dsSimulate(eqns,'vary',vary, 'study_dir',study_dir,'save_data_flag',1,...
%   'cluster_flag',0,'verbose_flag',1,'qsub_mode','array', 'one_solve_file_flag',1,...
%   'tspan',[0 100], 'dt',0.01, 'solver','euler', 'downsample_factor', 100,...
%   'plot_functions',{@dsPlot}, 'plot_options',{{'format', 'jpg', 'visible', 'off'}});
%
% %% Solve with one file mode compile
% study_dir=fullfile('.', 'study_HH_varyI_one_file_compile');
% if exist(study_dir, 'dir')
%   rmdir(study_dir, 's')
% end
%
% eqns='dv/dt=@current+I; {iNa,iK}';
% vary={'','I',[0:10:30]};
%
% dsSimulate(eqns,'vary',vary, 'study_dir',study_dir,'save_data_flag',1,...
%   'cluster_flag',0,'verbose_flag',1,'qsub_mode','array', 'one_solve_file_flag',1,...
%   'tspan',[0 100], 'dt',0.01, 'solver','euler', 'downsample_factor', 100,...
%   'plot_functions',{@dsPlot}, 'plot_options',{{'format', 'jpg', 'visible', 'off'}},...
%   'parfor_flag',0, 'mex_flag',1);
%
%% Solve with one file mode parallel
study_dir=fullfile('.', 'study_HH_varyI_one_file_parallel');
if exist(study_dir, 'dir')
  rmdir(study_dir, 's')
end

eqns='dv/dt=@current+I; {iNa,iK}';
vary={'','I',[0:10:30]};

dsSimulate(eqns,'vary',vary, 'study_dir',study_dir,'save_data_flag',1,...
  'cluster_flag',0,'verbose_flag',1,'qsub_mode','array', 'one_solve_file_flag',1,...
  'tspan',[0 100], 'dt',0.01, 'solver','euler', 'downsample_factor', 100,...
  'plot_functions',{@dsPlot}, 'plot_options',{{'format', 'jpg', 'visible', 'off'}},...
  'parfor_flag',1, 'mex_flag',0);
%
%% Solve with one file mode parallel compile
study_dir=fullfile('.', 'study_HH_varyI_one_file_parallel_compile');
if exist(study_dir, 'dir')
  rmdir(study_dir, 's')
end

eqns='dv/dt=@current+I; {iNa,iK}';
vary={'','I',[0:10:30]};

dsSimulate(eqns,'vary',vary, 'study_dir',study_dir,'save_data_flag',1,...
  'cluster_flag',0,'verbose_flag',1,'qsub_mode','array', 'one_solve_file_flag',1,...
  'tspan',[0 100], 'dt',0.01, 'solver','euler', 'downsample_factor', 100,...
  'plot_functions',{@dsPlot}, 'plot_options',{{'format', 'jpg', 'visible', 'off'}},...
  'parfor_flag',1, 'mex_flag',1);


%% ***************************************************************
%% ****************************CLUSTER****************************
%% ***************************************************************

%% Solve with one file mode
study_dir=fullfile('.', 'study_HH_varyI_one_file_scc');
if exist(study_dir, 'dir')
  rmdir(study_dir, 's')
end

eqns='dv/dt=@current+I; {iNa,iK}';
vary={'','I',[0:10:30]};

dsSimulate(eqns,'vary',vary, 'study_dir',study_dir,'save_data_flag',1,...
  'cluster_flag',1,'verbose_flag',1,'qsub_mode','array', 'one_solve_file_flag',1,...
  'tspan',[0 100], 'dt',0.01, 'solver','euler', 'downsample_factor', 100,...
  'plot_functions',{@dsPlot}, 'plot_options',{{'format', 'jpg', 'visible', 'off'}});

%% Solve with one file mode compile
study_dir=fullfile('.', 'study_HH_varyI_one_file_compile_scc');
if exist(study_dir, 'dir')
  rmdir(study_dir, 's')
end

eqns='dv/dt=@current+I; {iNa,iK}';
vary={'','I',[0:10:30]};

dsSimulate(eqns,'vary',vary, 'study_dir',study_dir,'save_data_flag',1,...
  'cluster_flag',1,'verbose_flag',1,'qsub_mode','array', 'one_solve_file_flag',1,...
  'tspan',[0 100], 'dt',0.01, 'solver','euler', 'downsample_factor', 100,...
  'plot_functions',{@dsPlot}, 'plot_options',{{'format', 'jpg', 'visible', 'off'}},...
  'parfor_flag',0, 'mex_flag',1);

%% Solve with one file mode parallel
study_dir=fullfile('.', 'study_HH_varyI_one_file_parallel_scc');
if exist(study_dir, 'dir')
  rmdir(study_dir, 's')
end

eqns='dv/dt=@current+I; {iNa,iK}';
vary={'','I',[0:10:30]};

dsSimulate(eqns,'vary',vary, 'study_dir',study_dir,'save_data_flag',1,...
  'cluster_flag',1,'verbose_flag',1,'qsub_mode','array', 'one_solve_file_flag',1,...
  'tspan',[0 100], 'dt',0.01, 'solver','euler', 'downsample_factor', 100,...
  'plot_functions',{@dsPlot}, 'plot_options',{{'format', 'jpg', 'visible', 'off'}},...
  'parfor_flag',1, 'mex_flag',0);

%% Solve with one file mode parallel compile
study_dir=fullfile('.', 'study_HH_varyI_one_file_parallel_compile_scc');
if exist(study_dir, 'dir')
  rmdir(study_dir, 's')
end

eqns='dv/dt=@current+I; {iNa,iK}';
vary={'','I',[0:10:30]};

dsSimulate(eqns,'vary',vary, 'study_dir',study_dir,'save_data_flag',1,...
  'cluster_flag',1,'verbose_flag',1,'qsub_mode','array', 'one_solve_file_flag',1,...
  'tspan',[0 100], 'dt',0.01, 'solver','euler', 'downsample_factor', 100,...
  'plot_functions',{@dsPlot}, 'plot_options',{{'format', 'jpg', 'visible', 'off'}},...
  'parfor_flag',1, 'mex_flag',1);


% %% Sparse Pyramidal-Interneuron-Network-Gamma (sPING)
% study_dir=fullfile('.', 'study_sPING_one_file');
% if exist(study_dir, 'dir')
%   rmdir(study_dir, 's')
% end
%
% % define equations of cell model (same for E and I populations)
% eqns={
%   'dv/dt=Iapp+@current+noise*randn(1,N_pop)'
% };
% % Tip: monitor all functions of a mechanism using: monitor MECHANISM.functions
%
% % create DynaSim specification structure
% s=[];
% s.populations(1).name='E';
% s.populations(1).size=4;
% s.populations(1).equations=eqns;
% s.populations(1).mechanism_list={'iNa','iK'};
% s.populations(1).parameters={'Iapp',5,'gNa',120,'gK',36,'noise',40};
% s.populations(2).name='I';
% s.populations(2).size=2;
% s.populations(2).equations=eqns;
% s.populations(2).mechanism_list={'iNa','iK'};
% s.populations(2).parameters={'Iapp',0,'gNa',120,'gK',36,'noise',40};
% s.connections(1).direction='I->E';
% s.connections(1).mechanism_list={'iGABAa'};
% s.connections(1).parameters={'tauD',10,'gSYN',.1,'netcon','ones(N_pre,N_post)'};
% s.connections(2).direction='E->I';
% s.connections(2).mechanism_list={'iAMPA'};
% s.connections(2).parameters={'tauD',2,'gSYN',.1,'netcon','ones(N_pre,N_post)'};
%
% vary={
%   'E'   ,'(gNa,gK)',[10 20];
%   '(E,I)','Iapp',[0 10];      % amplitude of tonic input to E-cells
%   'I->E','tauD',[5 10]       % inhibition decay time constant from I to E
%   };
%
% dsSimulate(s,'vary',vary, 'study_dir',study_dir,'save_data_flag',1, 'mex_flag',1,...
%   'cluster_flag',1,'verbose_flag',1,'qsub_mode','array', 'one_solve_file_flag',1,...
%   'tspan',[0 500], 'dt',0.01, 'solver','euler', 'downsample_factor', 100,...
%   'plot_functions',{@dsPlot}, 'plot_options',{{'format', 'jpg', 'visible', 'off'}});
```
