## Supplemental File 1 for "A biophysical model of striatal microcircuits suggests delta/theta-rhythmically interleaved gamma and beta oscillations mediate periodicity in motor control": dsDemos_PlotData2.m

```
% % % % % % % % dsPlot2 tutorial % % % % % % % %
% % % % % % % % % % % % % % % % % % % % % % % % % % % % % % % % % % % % 
% Catch to prevent people from running entire script (F5)
error('Do not F5. This script is not meant to be run in its entirety.');

%% Set up paths 
% Get ready...

% Add DynaSim to path if it's not already there
if exist('setupDynaSimPath','file')
    setupDynaSimPath;
else
    error('Add the DynaSim folder to the MATLAB path - e.g. run addpath(genpath(DynaSimPath))');
end

% Set where to save outputs
output_directory = dsGetConfig('demos_path');
study_dir = fullfile(output_directory,'demo_sPING_100cells_3x3');

%% Import the data

% Make sure sample data exists; if not copy it into place
if ~exist(study_dir,'dir')
    dsUnzipDemoData(study_dir);
end

% Load data in traditional DynaSim format
data_full=dsImport(study_dir);

% % Import saved plots from huge sweep of images
% data_3D_plots = dsImportPlots(fullfile(output_directory,'demo_sPING_3b_3D'));

%% Decimate data to increase speeds

% Remove all but 5 cells at random, so as to increase speed
data = dsDecimateCells(data_full,5);

% Decimate the data by dropping every other data point
data = dsDecimateData(data,2);

%% Do some basic plots with dsPlot2

close all

% Default settings
dsPlot2(data);

% Take mean. Note that E and I cells are now collapsed into the same plot.
dsPlot2(data,'do_mean',1);

% Plot data, just E cells, with zoom turned on
dsPlot2(data,'do_zoom',1,'population','E');

% As above, but focus on only a subset of the varied data. Also, reduce the
% total number of traces displayed with the max_num_overlaid option. Note
% that varied2 is an alias for the second parameter varied, in this
% case I_E_tauD.
dsPlot2(data,'population','E','variable','v','E_Iapp',1:3,'varied2',[1,3],'do_zoom',1,'max_num_overlaid',3);


% All basic plot_types from dsPlot should work
% Options are availalbe from dsPlot:
%     {'waveform','imagesc','rastergram','raster','power'}
% And also from dsPlotFR2
%     {'heatmapFR','heatmap_sortedFR','meanFR','meanFRdens'}
dsPlot2(data_full,'population','all','plot_type','rastergram')              % Rastergram
dsPlot2(data_full,'population','I','plot_type','heatmap_sortedFR')          % Firing rate (FR) heatmap

%% Re-ordering plots

% The order by which dimensions of the data are "stacked" into dsPlot2
% figures is controlled by two options: force_last and dim_stacking

% By default, if there is only 1 cell present in each population,
% populations are overlaid in subplots
dsPlot2(data,'max_num_overlaid',1);

% However, this can be changed with the 'force_last' option. Now,
% different strenght applied currents are now overlaid within subplots, and
% each population gets its own axis.
dsPlot2(data,'max_num_overlaid',1,'force_last','E_Iapp');

% More detailed control can be obtained with the dim_stacking option, in
% which the ordering of all dimensions can be controlled. The following
% forces the varied2 parameter (synaptic decay time) to the bottom of the
% stack as the overlay, and then populations and varied1 are the subplots.
dsPlot2(data,'max_num_overlaid',1,'dim_stacking',{'variables','populations','varied1','varied2'});

% Note that, with dim_stacking, since this is essentially a permute
% operation, all dimensions present in the original data must be specified. For
% example, dsPlot2(data,'max_num_overlaid',1,'dim_stacking',{'populations','varied1','varied2'})
% will produce an error.

% Additionally, values within a dimension can be re-ordered using the 
% value_stacking option. For example, the following plots I cells first and
% E cells second.
dsPlot2(data,'max_num_overlaid',1,'force_last','E_Iapp','value_stacking',{'populations',[2,1]});
% Here, [2,1] denotes to select the second population first, and the first
% second.


%% Modifying overlaid traces

close all

% This section provides a method for stacking more data into a single
% plot.

% If there is only 1 cell from each population, dsPlot will overlay
% cells from different populations by default.
% This can be achieved by averaging across cells...
dsPlot2(data,'do_mean',1,'varied1',1:3,'varied2',2:3)

% ... or by looking at only the 1st cell in each population
dsPlot2(data,'max_num_overlaid',1,'varied1',1:3,'varied2',2:3)

close all;

% Overlays can also be doubly stacked. 
dsPlot2(data,'force_last','populations','Ndims_per_subplot',2);

% Here, Ndims_per_subplot tells dsPlot2 to assign 2
% dimensions to each subplot panel - in this case, both the populations
% dimension (E and I populations) and the cells dimension (the first 5
% cells of each population). Although it is difficult to see, both E and I
% cells are present in each individual subplot.

% This can be cleaned up by shifting groups of cells up or down with the
% do_overlay_shift option.
dsPlot2(data,'force_last','populations','do_overlay_shift',true,'Ndims_per_subplot',2);

    % Variables are stacked from top to bottom, so E cells are on top and I
    % cells underneath

% You can overlay any variable. For example, here is injected current.
% This is accomplished by setting the axis "varied1", which corresponds to
% the injected current, to be the variable that is rastered across the
% overlays.

dsPlot2(data,'do_mean',0,'varied1',1:3,'varied2',2:3,...
    'force_last','varied1','do_overlay_shift',1,'overlay_shift_val',100,'Ndims_per_subplot',2);


% Variables with very different units can be compared side-by-side by
% taking the z-score first. In this case we compare E cell membrane voltage
% to its inhibitory synaptic input.

dsPlot2(data,'population','E','variable','/v|I_iGABAa_s/','force_last','variables','do_overlay_shift',true,'overlay_shift_val',3,'do_zscore',true,'do_zoom',1,'Ndims_per_subplot',2);

% % Double stack overlays
% dsPlot2(data,'population','E','variable','/iNa_m|I_iGABAa_s/','force_last','variables','do_overlay_shift',true);
% dsPlot2(data,'population','I','variable','v');

%% Overlay different types of plots
% This code overlays a plot of E cell membrane potential with the GABA A
% state variable.

% First, plot a heatmap of GABA A state variable
h = dsPlot2(data_full,'population','E','variable','iGABAa_s','plot_type','imagesc');

% Extract the handle for the subplot_grid subplot
fig_options.suppress_newfig = true;
subplot_options.subplot_grid_handle = h.hsub{1}.hcurr;

% Normalize Vm data between 0 and 20 (values of y-axis limits)
norm_data = @(x) (x - min(x(:))) ./ (max(x(:)) - min(x(:))) * 20;
myxp = dsAll2mdd(data_full);
myxp.data = cellfun(norm_data,myxp.data,'UniformOutput',0);

% Plot this data overlaid. We include the above axis handle in the options
% structure of xp_subplot_grid.m
h = dsPlot2(myxp,'population','E','plot_type','waveform',...
    'figure_options',fig_options,...
    'subplot_options',subplot_options);


%% Import pre-saved images
close all

% Import plot files
data_img = dsImportPlots(study_dir);
dsPlot2(data_img);

% Can also reference study_dir directly
dsPlot2(study_dir)

% As above, but supersize them
dsPlot2(study_dir,'supersize_me',1)
    % See folder ./Figs for output
    % Note that the subplots will be blurry because the original images
    % were also blurry; however, this can be fixed by increasing the DPI
    % of the original images.


%% Supersize me
% load sample_data_dynasim_large.mat
% 
% dsPlot2(data);


%% Merging simulation output data

study_dir2 = fullfile(output_directory,'demo_sPING_100cells_5x1');

% Make sure sample data exists; if not copy it into place
if ~exist(study_dir2,'dir')
    dsUnzipDemoData(study_dir2);
end

% Load data in traditional DynaSim format
data_sim2=dsImport(study_dir2);

% Test plot of new data
dsPlot2(data_sim2);

% Plot the merged new data with the old data
data_merged = dsMergeData(data,data_sim2);
dsPlot2(data_merged,'do_mean',1)


%% Merge simulations' saved images

data_img_sim2=dsImportPlots(study_dir2);

% Test plot of saved images from sim2
dsPlot2(data_img_sim2);

% Merge
data_img_merged = dsMergeData(data_img,data_img_sim2);
dsPlot2(data_img_merged);


%% Recursive plots with dsPlot2

close all

% The number of subplots packed into a single figure can be substantially
% increased by recursive subplotting. This is controlled with the
% "num_embedded_subplots" flag, which can range from 1-4.

% Plot membrane voltage for E and I cells across the parameter sweep
% % (1 variable, 2 pops, varied1, varied2: Ndims = 3)
dsPlot2(data,'num_embedded_subplots',2,'do_zoom',1);  % Default
dsPlot2(data,'num_embedded_subplots',4,'do_zoom',1);  % Nested embedding

% Note that in the prev example, although we set embedded subplots to 4, only
% the first 3 are used. If excess subplots are requested and unused, they
% will be thrown out.
% 
% Setting num_embedded_subplots to provides the same data but in a different
% arrangement.
dsPlot2(data,'num_embedded_subplots',3,'do_zoom',1);

% Depending on the data being plotted, different some arrangements can be
% more useful than others. For example, this plot embeds varied as a
% sub-subplot.
% % (2 variables, 2 pops, varied2, (varied1 is fixed): Ndims = 3)
dsPlot2(data,'num_embedded_subplots',3,'variable','iNa','varied1',2)

% Lastly, 4 subplots can be nested together to view 4 dimensions
% simultaneously. This can be increased to 5 with the overlay function
% described below, and 6 if you count multiple figures. While these figures
% may get crowded, the supersize_me flag, also described below, aims to 
% address this by producing single figures occupying very large canvases,
% which can be zoomed to a high level of detail.
% % (2 vars, 2 pops, varied1, varied2: Ndims = 4)
dsPlot2(data,'num_embedded_subplots',4,'population','all','variable','/v|iNa_h/','varied1',2:3,'lock_axes',false);
```
