## Supplemental File 1 for "A biophysical model of striatal microcircuits suggests delta/theta-rhythmically interleaved gamma and beta oscillations mediate periodicity in motor control": dsDemos_PlotData2_debugging.m

```
%%
% % % % % % % % % % % % % % % % % % % % % % % % % % % % % % % % % 
% % % % % % % % Advanced testing for debugging dsPlot2 % % % % %
% % % % % % % % % % % % % % % % % % % % % % % % % % % % % % % % % 
%% Advanced testing for debugging - set up data

% Make sure data is loaded and path is ready
if ~exist('dsSimulate','file'); error('Use demos_dsPlot2.m script to set up simulation path.'); end
if ~exist('data','var'); error('Use demos_dsPlot2.m script to load data'); end

xp = ds2mdd(data);
xp_single = xp(1,1,1,'v');
xp_single_pops = xp(1,1,:,'v');
xp_single_vars = xp(1,1,1,:);
xp_row = xp(:,1,1,'v');
xp_col = xp(1,:,1,'v');
xp_col_pops = xp(2,:,:,'v');
xp_col_vars = xp(1,:,1,:);
xp_col_varspops = xp(1,:,:,:);
xp_mat_pops = xp(:,:,:,'v');
xp_mat_vars = xp(:,:,1,:);

data_single = dsMdd2ds(xp_single);
data_single_squeeze = dsMdd2ds(squeeze(xp_single));
data_single_pops = dsMdd2ds(xp_single_pops); data_single_pops=rmfield(data_single_pops,{'varied','I_E_tauD','E_Iapp'});  % This removes all the "vary" fields
data_single_vars = dsMdd2ds(xp_single_vars); data_single_vars=rmfield(data_single_vars,{'varied','I_E_tauD','E_Iapp'});  % This removes all the "vary" fields
data_row = dsMdd2ds(xp_row);
data_col = dsMdd2ds(xp_col);
data_col_pops = dsMdd2ds(xp_col_pops);
data_col_vars = dsMdd2ds(xp_col_vars);
data_col_varspops = dsMdd2ds(xp_col_varspops);
data_mat_pops = dsMdd2ds(xp_mat_pops);
data_mat_vars = dsMdd2ds(xp_mat_vars);
data_all = dsMdd2ds(xp);

vis = 'off';

%% Plot waveforms, comparing dsPlot and dsPlot2
close all; d = data_single; dsPlot2(d,'visible',vis); d = dsMdd2ds(ds2mdd(d)); dsPlot2(d,'visible',vis); %dsPlot(d,'visible',vis);
close all; d = data_single_squeeze; dsPlot2(d,'visible',vis); d = dsMdd2ds(ds2mdd(d)); dsPlot2(d,'visible',vis); %dsPlot(d,'visible',vis);
close all; d = data_single_pops; dsPlot2(d,'visible',vis); d = dsMdd2ds(ds2mdd(d)); dsPlot2(d,'visible',vis); dsPlot(d,'visible',vis);
close all; d = data_single_vars; dsPlot2(d,'visible',vis); d = dsMdd2ds(ds2mdd(d)); dsPlot2(d,'visible',vis); %dsPlot(d,'visible',vis);
close all; d = data_row; dsPlot2(d,'visible',vis); d = dsMdd2ds(ds2mdd(d)); dsPlot2(d,'visible',vis); dsPlot(d,'visible',vis);
close all; d = data_col; dsPlot2(d,'visible',vis); d = dsMdd2ds(ds2mdd(d)); dsPlot2(d,'visible',vis); dsPlot(d,'visible',vis);
close all; d = data_col_pops; dsPlot2(d,'visible',vis); d = dsMdd2ds(ds2mdd(d)); dsPlot2(d,'visible',vis); dsPlot(d,'visible',vis);
close all; d = data_col_vars; dsPlot2(d,'visible',vis); d = dsMdd2ds(ds2mdd(d)); dsPlot2(d,'visible',vis); dsPlot(d,'visible',vis);
close all; d = data_col_varspops; dsPlot2(d,'visible',vis); d = dsMdd2ds(ds2mdd(d)); dsPlot2(d,'visible',vis); dsPlot(d,'visible',vis);
close all; d = data_mat_pops; dsPlot2(d,'do_mean',1,'visible',vis); d = dsMdd2ds(ds2mdd(d)); dsPlot2(d,'do_mean',1,'visible',vis); dsPlot(d,'visible',vis);
close all; d = data_mat_vars; dsPlot2(d,'visible',vis); d = dsMdd2ds(ds2mdd(d)); dsPlot2(d,'visible',vis); dsPlot(d,'visible',vis);
close all; d = data_all; dsPlot2(d,'do_mean',1,'visible',vis); d = dsMdd2ds(ds2mdd(d)); dsPlot2(d,'do_mean',1,'visible',vis); dsPlot(d,'visible',vis);


%% Plot rastergrams, comparing dsPlot and dsPlot2
close all; d = data_single_pops; dsPlot2(d,'plot_type','rastergram','visible',vis); d = dsMdd2ds(ds2mdd(d)); dsPlot2(d,'plot_type','rastergram','visible',vis); dsPlot(d,'plot_type','rastergram','visible',vis);
close all; d = data_single_vars; dsPlot2(d,'plot_type','rastergram','visible',vis); d = dsMdd2ds(ds2mdd(d)); dsPlot2(d,'plot_type','rastergram','visible',vis); dsPlot(d,'plot_type','rastergram','visible',vis);
close all; d = data_row; dsPlot2(d,'plot_type','rastergram','visible',vis); d = dsMdd2ds(ds2mdd(d)); dsPlot2(d,'plot_type','rastergram','visible',vis); dsPlot(d,'plot_type','rastergram','visible',vis);
close all; d = data_col; dsPlot2(d,'plot_type','rastergram','visible',vis); d = dsMdd2ds(ds2mdd(d)); dsPlot2(d,'plot_type','rastergram','visible',vis); dsPlot(d,'plot_type','rastergram','visible',vis);
close all; d = data_col_pops; dsPlot2(d,'plot_type','rastergram','visible',vis); d = dsMdd2ds(ds2mdd(d)); dsPlot2(d,'plot_type','rastergram','visible',vis); dsPlot(d,'plot_type','rastergram','visible',vis);
close all; d = data_col_vars; dsPlot2(d,'plot_type','rastergram','visible',vis); d = dsMdd2ds(ds2mdd(d)); dsPlot2(d,'plot_type','rastergram','visible',vis); dsPlot(d,'plot_type','rastergram','visible',vis);
close all; d = data_col_varspops; dsPlot2(d,'plot_type','rastergram','visible',vis); d = dsMdd2ds(ds2mdd(d)); dsPlot2(d,'plot_type','rastergram','visible',vis); dsPlot(d,'plot_type','rastergram','visible',vis);
% close all; d = data_mat_pops; dsPlot2(d,'plot_type','rastergram','do_mean',1,'visible',vis); d = dsMdd2ds(ds2mdd(d)); dsPlot2(d,'plot_type','rastergram','do_mean',1,'visible',vis); dsPlot(d,'plot_type','rastergram','visible',vis);
close all; d = data_mat_vars; dsPlot2(d,'plot_type','rastergram','visible',vis); d = dsMdd2ds(ds2mdd(d)); dsPlot2(d,'plot_type','rastergram','visible',vis); dsPlot(d,'plot_type','rastergram','visible',vis);
% close all; d = data_all; dsPlot2(d,'plot_type','rastergram','do_mean',1,'visible',vis); d = dsMdd2ds(ds2mdd(d)); dsPlot2(d,'plot_type','rastergram','do_mean',1,'visible',vis); dsPlot(d,'plot_type','rastergram','visible',vis);


%% Plot FR2, comparing dsPlot and dsPlot2
close all; d = data_single_pops; dsPlot2(d,'plot_type','heatmap_sortedFR','visible',vis); d = dsMdd2ds(ds2mdd(d)); dsPlot2(d,'plot_type','heatmap_sortedFR','visible',vis); dsPlotFR2(d,'plot_type','heatmap_sorted','visible',vis);
close all; d = data_single_vars; dsPlot2(d,'plot_type','heatmap_sortedFR','visible',vis); d = dsMdd2ds(ds2mdd(d)); dsPlot2(d,'plot_type','heatmap_sortedFR','visible',vis); dsPlotFR2(d,'plot_type','heatmap_sorted','visible',vis);
close all; d = data_row; dsPlot2(d,'plot_type','heatmap_sortedFR','visible',vis); d = dsMdd2ds(ds2mdd(d)); dsPlot2(d,'plot_type','heatmap_sortedFR','visible',vis); dsPlotFR2(d,'plot_type','heatmap_sorted','visible',vis);
close all; d = data_col; dsPlot2(d,'plot_type','heatmap_sortedFR','visible',vis); d = dsMdd2ds(ds2mdd(d)); dsPlot2(d,'plot_type','heatmap_sortedFR','visible',vis); dsPlotFR2(d,'plot_type','heatmap_sorted','visible',vis);
close all; d = data_col_pops; dsPlot2(d,'plot_type','heatmap_sortedFR','visible',vis); d = dsMdd2ds(ds2mdd(d)); dsPlot2(d,'plot_type','heatmap_sortedFR','visible',vis); dsPlotFR2(d,'plot_type','heatmap_sorted','visible',vis);
close all; d = data_col_vars; dsPlot2(d,'plot_type','heatmap_sortedFR','visible',vis); d = dsMdd2ds(ds2mdd(d)); dsPlot2(d,'plot_type','heatmap_sortedFR','visible',vis); dsPlotFR2(d,'plot_type','heatmap_sorted','visible',vis);
close all; d = data_col_varspops; dsPlot2(d,'plot_type','heatmap_sortedFR','visible',vis); d = dsMdd2ds(ds2mdd(d)); dsPlot2(d,'plot_type','heatmap_sortedFR','visible',vis); dsPlotFR2(d,'plot_type','heatmap_sorted','visible',vis);
% close all; d = data_mat_pops; dsPlot2(d,'plot_type','heatmap_sortedFR','do_mean',1,'visible',vis); d = dsMdd2ds(ds2mdd(d)); dsPlot2(d,'plot_type','heatmap_sortedFR','do_mean',1,'visible',vis); dsPlotFR2(d,'plot_type','heatmap_sorted','visible',vis);
close all; d = data_mat_vars; dsPlot2(d,'plot_type','heatmap_sortedFR','visible',vis); d = dsMdd2ds(ds2mdd(d)); dsPlot2(d,'plot_type','heatmap_sortedFR','visible',vis); dsPlotFR2(d,'plot_type','heatmap_sorted','visible',vis);
% close all; d = data_all; dsPlot2(d,'plot_type','heatmap_sortedFR','do_mean',1,'visible',vis); d = dsMdd2ds(ds2mdd(d)); dsPlot2(d,'plot_type','heatmap_sortedFR','do_mean',1,'visible',vis); dsPlotFR2(d,'plot_type','heatmap_sorted','visible',vis);


keyboard

%% Advanced testing for debugging - set up data_img

xp_img = dsImg2MDD(data_img);
```
