## Supplemental File 1 for "A biophysical model of striatal microcircuits suggests delta/theta-rhythmically interleaved gamma and beta oscillations mediate periodicity in motor control": dsDemos_qsub_modes.m

```
% Below are 2 different ways to simulate on the cluster.  With larger
% simulations (>100s), array mode should be faster.

%% Setup

% Add DynaSim to path if it's not already there
if exist('setupDynaSimPath','file')
    setupDynaSimPath;
else
    error('Add the DynaSim folder to the MATLAB path - e.g. run addpath(genpath(DynaSimPath))');
end

% Set where to save outputs
output_directory = dsGetConfig('demos_path');

% move to root directory where outputs will be saved
mkdirSilent(output_directory);
cd(output_directory);

%% Solve with loop mode
study_dir = fullfile('.', 'study_HH_varyI_cluster_loop');
eqns='dv/dt=@current+I; {iNa,iK}';
vary={'','I',[0:10:50]};

if exist(study_dir, 'dir')
  rmdir(study_dir, 's')
end

dsSimulate(eqns,'vary',vary, 'study_dir',study_dir,'save_data_flag',1,...
  'cluster_flag',1,'verbose_flag',1,'qsub_mode','loop');

%% Solve with array mode
study_dir = fullfile('.', 'study_HH_varyI_cluster_array');
eqns='dv/dt=@current+I; {iNa,iK}';
vary={'','I',[0:10:50]};

if exist(study_dir, 'dir')
  rmdir(study_dir, 's')
end

dsSimulate(eqns,'vary',vary, 'study_dir',study_dir,'save_data_flag',1,...
  'cluster_flag',1,'verbose_flag',1,'qsub_mode','array');
```
