## Supplemental File 1 for "A biophysical model of striatal microcircuits suggests delta/theta-rhythmically interleaved gamma and beta oscillations mediate periodicity in motor control": dsUnitTests.m

```
% %%%%%%%%%%%%%%%%%%%%%%%%%%%%%%%%%%%%%%%%%%%%%%%%%%%%%%%%%%%%%%%%%%%%%%%%
% TO-DO: debug this script so that it runs without error. Then, use it to
% verify that all future changes to the DynaSim toolbox are in working
% order before committing them.
% %%%%%%%%%%%%%%%%%%%%%%%%%%%%%%%%%%%%%%%%%%%%%%%%%%%%%%%%%%%%%%%%%%%%%%%%

% Purpose of script: verify that future changes to the DynaSim toolbox do
% not break any of the features that preceded them. Always run this script
% before committing changes. This script should run unit tests for all
% features of the toolbox; thus, this script should be updated throughout
% development to incorporate all new features (so that they too are tested
% against future changes).

% %%%%%%%%%%%%%%%%%%%%%%%%%%%%%%%%%%%%%%%%%%%%%%%%%%%%%%%%%%%%%%%%%%%%%%%%
% path to DynaSim toolbox
dynasim_path='~/code/dynasim';
output_directory='~/code/dynasim/docs/demos/outputs';

addpath(dynasim_path);
cd(output_directory);

model=dsGenerateModel; % obtain default test model

% parameter values inserted in expressions:

% data stored in memory
!rm solve_ode.m
dsWriteDynaSimSolver(model,'disk_flag',0,'save_parameters_flag',0,'reduce_function_calls_flag',1,'solver','rk4','filename','solve_ode.m');
tic; [t,Ev,Ew,Em,Eh,En,cai,cam,Iv,sie,sei,sii,Eo,EiNa,EiNaJ,Io]=solve_ode; toc;
figure; plot(t,Ev);
!rm solve_ode.m
dsWriteDynaSimSolver(model,'disk_flag',0,'save_parameters_flag',0,'reduce_function_calls_flag',0,'solver','rk4','filename','solve_ode.m');
tic; [t,Ev,Ew,Em,Eh,En,cai,cam,Iv,sie,sei,sii,Eo,EiNa,EiNaJ,Io]=solve_ode; toc;
figure; plot(t,Ev);

% data written to disk (real-time)
!rm solve_ode.m
dsWriteDynaSimSolver(model,'disk_flag',1,'downsample_factor',10,'save_parameters_flag',0,'reduce_function_calls_flag',1,'solver','rk4','filename','solve_ode.m');
tic; solve_ode; toc
data=getfield(importdata('data.csv',','),'data');
t=data(:,1); v=data(:,2:3); figure; plot(t,v);
!rm solve_ode.m
dsWriteDynaSimSolver(model,'disk_flag',1,'downsample_factor',10,'save_parameters_flag',0,'reduce_function_calls_flag',0,'solver','rk4','filename','solve_ode.m');
tic; solve_ode; toc
data=getfield(importdata('data.csv',','),'data');
t=data(:,1); v=data(:,2:3); figure; plot(t,v);

% parameter values stored in structure on disk:

% data stored in memory
!rm solve_ode.m
dsWriteDynaSimSolver(model,'disk_flag',0,'save_parameters_flag',1,'reduce_function_calls_flag',1,'solver','rk4','filename','solve_ode.m');
tic; [t,Ev,Ew,Em,Eh,En,cai,cam,Iv,sie,sei,sii,Eo,EiNa,EiNaJ,Io]=solve_ode; toc;
figure; plot(t,Ev);
!rm solve_ode.m
dsWriteDynaSimSolver(model,'disk_flag',0,'save_parameters_flag',1,'reduce_function_calls_flag',0,'solver','rk4','filename','solve_ode.m');
tic; [t,Ev,Ew,Em,Eh,En,cai,cam,Iv,sie,sei,sii,Eo,EiNa,EiNaJ,Io]=solve_ode; toc;
figure; plot(t,Ev);

% data written to disk (real-time)
!rm solve_ode.m
dsWriteDynaSimSolver(model,'disk_flag',1,'downsample_factor',10,'save_parameters_flag',1,'reduce_function_calls_flag',1,'solver','rk4','filename','solve_ode.m');
tic; solve_ode; toc
data=getfield(importdata('data.csv',','),'data');
t=data(:,1); v=data(:,2:3); figure; plot(t,v);
!rm solve_ode.m
dsWriteDynaSimSolver(model,'disk_flag',1,'downsample_factor',10,'save_parameters_flag',1,'reduce_function_calls_flag',0,'solver','rk4','filename','solve_ode.m');
tic; solve_ode; toc
data=getfield(importdata('data.csv',','),'data');
t=data(:,1); v=data(:,2:3); figure; plot(t,v);
!rm solve_ode.m
dsWriteDynaSimSolver(model,'disk_flag',1,'downsample_factor',1,'save_parameters_flag',1,'reduce_function_calls_flag',1,'solver','rk4','filename','solve_ode.m');
tic; solve_ode; toc
data=getfield(importdata('data.csv',','),'data');
t=data(:,1); v=data(:,2:3); figure; plot(t,v);

%%
spec=[];
spec.populations(1).equations='dv/dt=current; {iNa,iK};';
spec.populations(1).size=1;
model=dsGenerateModel(spec);
data=dsSimulate(model,'mex_flag',0);
figure; plot(data.time,data.(data.labels{1}))

spec=[];
spec.populations(1).equations='dv/dt=current; {iNa,iK};';
spec.populations(1).size=2;
model=dsGenerateModel(spec);
data=dsSimulate(model,'mex_flag',0);
figure; plot(data.time,data.(data.labels{1}))

spec=[];
spec.populations(1).equations='dv/dt=current; {iNa,iK,iCa};';
spec.populations(1).size=1;
model=dsGenerateModel(spec);
data=dsSimulate(model,'mex_flag',0);
figure; plot(data.time,data.(data.labels{1}))

spec=[];
spec.populations(1).equations='dv/dt=current; {iNa,iK,iCa};';
spec.populations(1).size=2;
model=dsGenerateModel(spec);
data=dsSimulate(model,'mex_flag',0);
figure; plot(data.time,data.(data.labels{1}))

if 1
  % DEBUG (broke after adding save/dsUpdateStudy to dsSimulate)
  % problem might be same as below where the model is so equivalent that a new ODE solve file
  % is not being generate when it should be.
  % edit: probably same as reason "pause" is necessary throughout demo script

  % !rm solve_ode.m params.mat
  spec=[];
  spec.populations(1).equations='dv/dt=current; {iNa,iK,iCa,CaBuffer,iCan};';
  spec.populations(1).size=1;
  model=dsGenerateModel(spec);
  data=dsSimulate(model,'mex_flag',0);
  figure; plot(data.time,data.(data.labels{1}))

  % !rm solve_ode.m params.mat
  spec=[];
  spec.populations(1).equations='dv/dt=current; {iNa,iK,iCa,CaBuffer,iCan};';
  spec.populations(1).size=5;
  model=dsGenerateModel(spec);
  data=dsSimulate(model,'mex_flag',0);
  figure; plot(data.time,data.(data.labels{1}))

  !rm solve_ode.m params.mat
  spec=[];
  spec.populations(1).equations='dv/dt=current; {iNa,iK,iCa,CaBuffer,iCan};';
  spec.populations(1).size=5;
  spec.connections(1).mechanism_list={'GABAa'};
  model=dsGenerateModel(spec);
  data=dsSimulate(model,'mex_flag',0);
  figure; plot(data.time,data.(data.labels{1}))
end

spec=[];
spec.populations(1).equations='dv/dt=current+5; {iNa,iK,iCa,CaBuffer,iCan};';
spec.populations(1).size=5;
spec.populations(2).equations='dv/dt=current; {iNa,iK,iCa,CaBuffer,iCan};';
spec.populations(2).size=2;
spec.connections(1).source='pop1';
spec.connections(1).target='pop2';
spec.connections(1).mechanism_list={'GABAa'};
model=dsGenerateModel(spec);
data=dsSimulate(model,'mex_flag',0,'verbose_flag',1);
figure; plot(data.time,data.pop1_v,'b'); hold on; plot(data.time,data.pop2_v,'r')
data=dsSimulate(model,'mex_flag',1);
figure; plot(data.time,data.pop1_v,'b'); hold on; plot(data.time,data.pop2_v,'r')

spec=[];
spec.populations(1).equations='dv/dt=@current+10; {iNa3,iK3};';
model=dsGenerateModel(spec);
data=dsSimulate(model,'mex_flag',0);
figure; plot(data.time,data.(data.labels{1}))

spec=[];
spec.populations(1).equations='dv/dt=@M+@current+10; {iNa3@M,iK3};';
model=dsGenerateModel(spec);
data=dsSimulate(model,'mex_flag',0);
figure; plot(data.time,data.(data.labels{1}))

spec=[];
spec.populations(1).equations='dv/dt=@M+10; {iNa3@M,iK3@M};';
model=dsGenerateModel(spec);
data=dsSimulate(model,'mex_flag',0);
figure; plot(data.time,data.(data.labels{1}))

% EXAMPLE: fully modularized model specification (iNa3.mech uses 'X')
spec=[];
spec.populations(1).equations='dv/dt=@M+10; {iNa3,iK3}@M;';
model=dsGenerateModel(spec);
data=dsSimulate(model,'mex_flag',0);
figure; plot(data.time,data.(data.labels{1}))

% todo:
% add monitor func, functions, v.spikes(threshold=0) -- DONE
% add vary(...) or modify simstudy() for new model structure -- DONE

% MONITORS
spec=[];
spec.populations(1).equations='dv/dt=@M+10; monitor o=v/1000; {iNa3,iK3}@M;';
model=dsGenerateModel(spec);
% pause(1) % does this fix the bug? YES. it should if problem is solver file gets same timestamp-based name
data=dsSimulate(model,'mex_flag',0);
figure; plot(data.time,data.pop1_o)
figure; plot(data.time,data.pop1_iNa3_I) % monitor set in iNa3.mech

spec=[];
spec.populations(1).equations='dv/dt=@M+10; monitor iK3.I; {iNa3,iK3}@M;';
model=dsGenerateModel(spec);
data=dsSimulate(model,'mex_flag',0);
figure; plot(data.time,data.pop1_iK3_I)

spec=[];
spec.populations(1).equations='dv/dt=@M+10; monitor iK3_I; {iNa3,iK3}@M;';
model=dsGenerateModel(spec);
data=dsSimulate(model,'mex_flag',0);
figure; plot(data.time,data.pop1_iK3_I)

% Spike Monitor (default threshold=0)
spec=[];
spec.populations(1).equations='dv/dt=@M+10; {iNa3,iK3}@M; monitor v.spikes;';
model=dsGenerateModel(spec);
data=dsSimulate(model,'mex_flag',0);
figure; plot(data.time,data.pop1_v); hold on; plot(data.time,100*data.pop1_v_spikes,'r');

% Spike Monitor with custom threshold (=60)
spec=[];
spec.populations(1).equations=...
  'dv/dt=@M+10; {iNa3,iK3}@M; monitor v.spikes(60);';
data=dsSimulate(spec);
figure; plot(data.time,data.pop1_v); hold on; plot(data.time,100*data.pop1_v_spikes,'r');

% Spike Monitor (threshold=10) for different tonic amplitudes and max sodium conductance
amps=2:2:60; gNas=[101 120];
figure('position',[330 215 1200 680])
for j=1:length(gNas)
  spec=[];
  spec.populations.equations=...
    'dv/dt=@current+amp; amp=0; {iNa3,iK3}; monitor v.spikes(10);';
  for i=1:length(amps)
    spec.populations.parameters={'amp',amps(i),'gNa',gNas(j)};
    data=dsSimulate(spec,'tspan',[0 100],'solver','rk4','dt',.01);
    if i==1,spikes=zeros(length(data.time),length(amps));end
    spikes(:,i)=amps(i)+data.pop1_v_spikes;
  end
  subplot(1,2,j); plot(data.time,spikes,'k'); axis tight
  set(gca,'ytick',amps,'yticklabel',amps);
  xlabel('time (ms)'); ylabel('tonic amplitude [uA/cm2]');
  title(sprintf('gNa=%g',gNas(j)));
end

% test dsParseModelEquations robustness
data=dsSimulate('dv/dt=@current+10; {iNa3,iK3}');
figure; plot(data.time,data.(data.labels{1}))

data=dsSimulate('dv/dt=@M+10; {iNa3,iK3}@M');
figure; plot(data.time,data.(data.labels{1}))

% Size embedded in population equations with fully modularized mechanisms

data=dsSimulate('dx[20]/dt=@M+5*(1+randn(1,Npop)); {iNa3,iK3}@M');
figure; plot(data.time,data.(data.labels{1}))
% with different differential notation: x'=...
data=dsSimulate('x[20]''=@M+5*(1+randn(1,Npop)); {iNa3,iK3}@M');
figure; plot(data.time,data.(data.labels{1}))
% with spike monitor
data=dsSimulate(...
  'x[20]''=@M+50*(1+randn(1,Npop)); {iNa3,iK3}@M; monitor x.spikes(0)'...
  );
figure;
subplot(2,1,1); plot(data.time,data.(data.labels{1}))
subplot(2,1,2); plot(data.time,data.pop1_x_spikes+repmat(1:20,[length(data.time) 1]));


%% make back-compatible - support old DynaSim specs/models (via dsCheckModel())

cd(fullfile(dynasim_path,'models','legacy','B-LTS'));

NB=1; NLTS=1; Cm=.9; onset=500;
stimB=0;    % -16;
stimLTS=8;  % -40
noiseB=0;   % 30
noiseLTS=0; % 50
gBB=0;      % 15;
gBLTS=0;    % 8
gLTSLTS=0;  % 5
gLTSB=0;
spec=[];
spec.nodes(1).label = 'B';
spec.nodes(1).multiplicity = NB;
spec.nodes(1).dynamics = {'V''=(current)/Cm'};
spec.nodes(1).mechanisms = {'itonic','randn','iNaF','iKDR','leak'};
spec.nodes(1).parameters = {'Cm',Cm,'V_IC',-65,'IC_noise',0,'g_l',.3,'E_l',-75,...
  'stim',stimB,'onset',onset,'noise',noiseB,'gNaF',200,'gKDR',20};
spec.nodes(2).label = 'LTS';
spec.nodes(2).multiplicity = NLTS;
spec.nodes(2).dynamics = {'V''=(current)/Cm'};
spec.nodes(2).mechanisms = {'itonic','randn','iNaF','iKDR','iAR','leak'};
spec.nodes(2).parameters = {'Cm',Cm,'V_IC',-65,'IC_noise',0,'g_l',1,'E_l',-80,...
  'stim',stimLTS,'onset',onset,'noise',noiseLTS,'gNaF',200,'gKDR',10,'gAR',20};
spec.connections(1,1).label = 'B-B';
spec.connections(1,1).mechanisms = {'iSYN'};
spec.connections(1,1).parameters = {'g_SYN',gBB,'E_SYN',-75,'tauDx',5,'tauRx',.5,'IC_noise',0};
spec.connections(1,2).label = 'B-LTS';
spec.connections(1,2).mechanisms = {'iSYN'};
spec.connections(1,2).parameters = {'g_SYN',gBLTS,'E_SYN',-80,'tauDx',6,'tauRx',.5,'IC_noise',0};
spec.connections(2,2).label = 'LTS-LTS';
spec.connections(2,2).mechanisms = {'iSYN'};
spec.connections(2,2).parameters = {'g_SYN',gLTSLTS,'E_SYN',-80,'tauDx',20,'tauRx',.5,'IC_noise',0};
spec.connections(2,2).label = 'LTS-B';
spec.connections(2,2).mechanisms = {'iSYN'};
spec.connections(2,2).parameters = {'g_SYN',gLTSB,'E_SYN',-80,'tauDx',20,'tauRx',.5,'IC_noise',0};
model=dsGenerateModel(spec);
data=dsSimulate(model,'tspan',[0 1000],'mex_flag',1);
figure; plot(data.time,data.B_V,'r-',data.time,data.LTS_V,'g-')


cd(fullfile(dynasim_path,'models','legacy','sPING'));
nE=80; nI=20;
tauI=10; gI=.1; gE=.1; stim=7.5; noise=4;
spec=[];
spec.nodes(1).label = 'E';
spec.nodes(1).multiplicity = nE;
spec.nodes(1).dynamics = {'V''=(current)./Cm'};
spec.nodes(1).mechanisms = {'K','Na','leak','input','randn'};
spec.nodes(1).parameters = {'Cm',1,'V_IC',-70,'stim',stim,'noise',noise};
spec.nodes(2).label = 'I';
spec.nodes(2).multiplicity = nI;
spec.nodes(2).dynamics = {'V''=(current)./Cm'};
spec.nodes(2).mechanisms = {'K','Na','leak','input','randn'};
spec.nodes(2).parameters = {'Cm',1,'V_IC',-70,'stim',0,'noise',noise};
spec.connections(1,2).label = 'E-I';
spec.connections(1,2).mechanisms = {'AMPA2'};
spec.connections(1,2).parameters = {'tauDx',2,'g_SYN',gE*(80/nE)};
spec.connections(2,1).label = 'I-E';
spec.connections(2,1).mechanisms = {'GABAa2'};
spec.connections(2,1).parameters = {'tauDx',tauI,'g_SYN',gI*(20/nI)};
model=dsGenerateModel(spec);
data=dsSimulate(model,'tspan',[0 100],'mex_flag',0);
figure; plot(data.time,data.E_V,'b-',data.time,data.I_V,'r-')

% Sparse PING in new specification format
s=[];
s.pops(1).name='E';
s.pops(1).equations=...
  'dV[80]/dt=current; {K,Na,leak,input,randn}; stim=7.5; noise=4; V(0)=-70';
s.pops(2).name='I';
s.pops(2).equations=...
  'dV[20]/dt=current; {K,Na,leak,input,randn}; stim=0; noise=4; V(0)=-70';
s.cons(1).source='E';
s.cons(1).target='I';
s.cons(1).mechanism_list={'AMPA2'};
s.cons(1).parameters={'tauDx',2,'g_SYN',.1};
s.cons(2).source='I';
s.cons(2).target='E';
s.cons(2).mechanism_list={'GABAa2'};
s.cons(2).parameters={'tauDx',10,'g_SYN',.1};
data=dsSimulate(s,'tspan',[0 100],'solver','rk4','mex_flag',0);
figure; plot(data.time,data.E_V,'b-',data.time,data.I_V,'r-')

% Sparse PING in new specification format w/ full modularization (w/o input mechanism or pop state var IC)
s=[];
s.pops(1).name='E';
s.pops(1).equations=...
  'dy[80]/dt=@M+7.5; {K,Na,leak,randn}@M; noise=4';
s.pops(2).name='I';
s.pops(2).equations=...
  'dy[20]/dt=@M; {K,Na,leak,randn}@M; noise=4';
s.cons(1).source='E';
s.cons(1).target='I';
s.cons(1).mechanism_list={'AMPA2@M'};
s.cons(1).parameters={'tauDx',2,'g_SYN',.1};
s.cons(2).source='I';
s.cons(2).target='E';
s.cons(2).mechanism_list={'GABAa2@M'};
s.cons(2).parameters={'tauDx',10,'g_SYN',.1};
data=dsSimulate(s,'tspan',[0 100],'solver','rk4','mex_flag',0);
figure; plot(data.time,data.E_y,'b-',data.time,data.I_y,'r-')

% equivalent with explicit noise term
s=[];
s.pops(1).name='E';
s.pops(1).equations=...
  'dy[80]/dt=@M+40*randn(1,Npop)+7.5; {K,Na,leak}@M';
s.pops(2).name='I';
s.pops(2).equations=...
  'dy[20]/dt=@M+40*randn(1,Npop); {K,Na,leak}@M';
s.cons(1).source='E';
s.cons(1).target='I';
s.cons(1).mechanism_list={'AMPA2@M'};
s.cons(1).parameters={'tauDx',2,'g_SYN',.1};
s.cons(2).source='I';
s.cons(2).target='E';
s.cons(2).mechanism_list={'GABAa2@M'};
s.cons(2).parameters={'tauDx',10,'g_SYN',.1};
data=dsSimulate(s,'tspan',[0 100],'solver','rk4','mex_flag',0);
figure; plot(data.time,data.E_y,'b-',data.time,data.I_y,'r-')

% simulate with data stored on disk: method 1
m=dsGenerateModel(s);
dsWriteDynaSimSolver(m,'disk_flag',1,'filename','solve_ode.m','datafile','data.csv');
solve_ode; data=dsImport('data.csv');

% simulate with data stored on disk: method 2
data=dsSimulate(s,'tspan',[0 100],'disk_flag',1,'downsample_factor',10);
figure; plot(data.time,data.E_y,'b-',data.time,data.I_y,'r-')
%figure; plot(data.time,data.(data.labels{1}))

%% Add modifications

cd(output_directory);

data=dsSimulate('dv/dt=@current/c;c=1; {iNa3,iK3}',...
  'modifications',{'pop1','size',5});
figure; plot(data.time,data.(data.labels{1}))

% test backward-compatibility:
data=dsSimulate('dv/dt=@current/c+100; c=1; {iNa3,iK3}',...
  'modifications',{'pop1','size',5,[];'pop1','parameters','c',10});
figure; plot(data.time,data.(data.labels{1}))
s=[];
s.pops(1).equations='dv/dt=@current+100; {iNa3,iK3}';
s.pops(1).parameters={'gNa',120};
data=dsSimulate(s,'modifications',{'pop1','parameters','gNa',250});
figure; plot(data.time,data.(data.labels{1}))

if 1
  % DEBUG (broke after adding save/dsUpdateStudy to dsSimulate) - same model broken above
  % test modifying connection parameters:
  % !rm solve_ode.m params.mat
  spec=[];
  spec.pops.equations='dv/dt=current; {iNa,iK,iCa,CaBuffer,iCan};';
  spec.pops.size=5;
  spec.cons.mechanism_list={'GABAa'};
  model=dsGenerateModel(spec,'modifications',{'pop1->pop1','g_SYN',1});
  data=dsSimulate(model);
  % !rm solve_ode.m params.mat
  data=dsSimulate(spec,'modifications',{'pop1->pop1','g_SYN',1});
  figure; plot(data.time,data.(data.labels{1}))
end

% modify mechanism_list
m=dsApplyModifications('dv/dt=10+current; {iNa,iK}; v(0)=-65',...
                     {'pop1','mechanism_list','-iNa'});
%m.populations

m=dsApplyModifications('dv/dt=10+current; {iNa,iK}; v(0)=-65',...
                     {'pop1','mechanism_list','+iCa'});
%m.populations

m=dsApplyModifications('dv/dt=10+current; {iNa,iK}; v(0)=-65',...
                     {'pop1','mechanism_list','+(iCa,iCan,CaBuffer)'});
%m.populations

% modify equations
m=dsApplyModifications('dv/dt=10+current; {iNa,iK}',...
                                        {'pop1','equations','cat(dv/dt,+I)'});
%m.populations.equations
m=dsApplyModifications('dv/dt=10+current; {iNa,iK}',...
                                        {'pop1','equations','cat(ODE1,+I)'});
%m.populations.equations
m=dsApplyModifications('dv/dt=10+current; {iNa,iK}',...
                                        {'pop1','equations','cat(ODE,+I)'});
%m.populations.equations
m=dsApplyModifications('dv/dt=10+@current; du/dt=-u; {iNa,iK}',...
                     {'pop1','equations','cat(ODE2,+I)'});
m.populations.equations
m=dsApplyModifications('dv/dt=I(t)+@current; I(t)=10; {iNa,iK}',...
                     {'pop1','equations','cat(I(t),+sin(2*pi*t))'});
% m.populations.equations
m=dsApplyModifications('dv/dt=I(t)+@current; I(t)=10; {iNa,iK}',...
                     {'pop1','equations','dv/dt=10+@current'});
% m.populations.equations
% m.populations.mechanism_list
m=dsApplyModifications('dv/dt=I(t)+@current; I(t)=10; {iNa,iK}',...
                     {'pop1','equations','cat(FUNCTION,+sin(2*pi*t))'});
%m.populations.equations
m=dsApplyModifications('dv/dt=I(t)+@current; I(t)=10; J(t)=11; {iNa,iK}',...
                     {'pop1','equations','cat(FUNCTION2,+sin(2*pi*t))'});
%m.populations.equations

%% various ways of specifying (Na,K) HH-type model

eqns={
'dv/dt = 10-INa(v,m,h)-IK(v,n); v(0)=-65';
'dm/dt = aM(v).*(1-m)-bM(v).*m; m(0)=.1';
'dh/dt = aH(v).*(1-h)-bH(v).*h; h(0)=.1';
'dn/dt = aN(v).*(1-n)-bN(v).*n';
'INa(v,m,h) = 120.*m.^3.*h.*(v-50)';
'IK(v,n) = 36.*n.^4.*(v+77)';
'aM(v) = (2.5-.1*(v+65))./(exp(2.5-.1*(v+65))-1)';
'bM(v) = 4*exp(-(v+65)/18)';
'aH(v) = .07*exp(-(v+65)/20)';
'bH(v) = 1./(exp(3-.1*(v+65))+1)';
'aN(v) = (.1-.01*(v+65))./(exp(1-.1*(v+65))-1)';
'bN(v) = .125*exp(-(v+65)/80)';
};
data=dsSimulate(eqns);
figure; plot(data.time,data.(data.labels{1}))

eqns={
'dv/dt = 10-INa(v,m,h)-IK(v,n); v(0)=-65';
'dm/dt = aM(v).*(1-m)-bM(v).*m; m(0)=.1';
'dh/dt = aH(v).*(1-h)-bH(v).*h; h(0)=.1';
'dn/dt = aN(v).*(1-n)-bN(v).*n';
'INa(v,m,h) = 120.*m.^3.*h.*(v-50)';
'IK(v,n) = 36.*n.^4.*(v+77)';
'aM(v) = (2.5-.1*(v+65))./(exp(2.5-.1*(v+65))-1)';
'bM(v) = 4*exp(-(v+65)/18)';
'aH(v) = .07*exp(-(v+65)/20)';
'bH(v) = 1./(exp(3-.1*(v+65))+1)';
'aN(v) = (.1-.01*(v+65))./(exp(1-.1*(v+65))-1)';
'bN(v) = .125*exp(-(v+65)/80)';
};
data=dsSimulate(eqns);
figure; plot(data.time,data.(data.labels{1}))

eqns={};
eqns{end+1}='dv/dt = 10-INa(v,m,h)-IK(v,n); v(0)=-65';
eqns{end+1}='dm/dt = aM(v).*(1-m)-bM(v).*m; m(0)=.1';
eqns{end+1}='dh/dt = aH(v).*(1-h)-bH(v).*h; h(0)=.1';
eqns{end+1}='dn/dt = aN(v).*(1-n)-bN(v).*n';
eqns{end+1}='INa(v,m,h) = 120.*m.^3.*h.*(v-50)';
eqns{end+1}='IK(v,n) = 36.*n.^4.*(v+77)';
eqns{end+1}='aM(v) = (2.5-.1*(v+65))./(exp(2.5-.1*(v+65))-1)';
eqns{end+1}='bM(v) = 4*exp(-(v+65)/18)';
eqns{end+1}='aH(v) = .07*exp(-(v+65)/20)';
eqns{end+1}='bH(v) = 1./(exp(3-.1*(v+65))+1)';
eqns{end+1}='aN(v) = (.1-.01*(v+65))./(exp(1-.1*(v+65))-1)';
eqns{end+1}='bN(v) = .125*exp(-(v+65)/80)';
data=dsSimulate(eqns);
figure; plot(data.time,data.(data.labels{1}))

eqns=[...
'dv/dt = 10-INa(v,m,h)-IK(v,n); v(0)=-65;' ...
'dm/dt = aM(v).*(1-m)-bM(v).*m; m(0)=.1;' ...
'dh/dt = aH(v).*(1-h)-bH(v).*h; h(0)=.1;' ...
'dn/dt = aN(v).*(1-n)-bN(v).*n;' ...
'INa(v,m,h) = 120.*m.^3.*h.*(v-50);' ...
'IK(v,n) = 36.*n.^4.*(v+77);' ...
'aM(v) = (2.5-.1*(v+65))./(exp(2.5-.1*(v+65))-1);' ...
'bM(v) = 4*exp(-(v+65)/18);' ...
'aH(v) = .07*exp(-(v+65)/20);' ...
'bH(v) = 1./(exp(3-.1*(v+65))+1);' ...
'aN(v) = (.1-.01*(v+65))./(exp(1-.1*(v+65))-1);' ...
'bN(v) = .125*exp(-(v+65)/80);' ...
];
data=dsSimulate(eqns);
figure; plot(data.time,data.(data.labels{1}))

data=dsSimulate('HH.eqns');
figure; plot(data.time,data.(data.labels{1}))

data=dsSimulate('dv/dt=10+current; {iNa,iK}; v(0)=-65');
figure; plot(data.time,data.(data.labels{1}))

% with parameters and 'X':
% note: X gets replaced by the state variable of the first ODE in list
% (which is also the first state variable in data.labels)
eqns={
'gNa=120; ENa=50; gK=36; EK=-77';
'aM(X) = (2.5-.1*(X+65))./(exp(2.5-.1*(X+65))-1)';
'bM(X) = 4*exp(-(X+65)/18)';
'aH(X) = .07*exp(-(X+65)/20)';
'bH(X) = 1./(exp(3-.1*(X+65))+1)';
'aN(X) = (.1-.01*(X+65))./(exp(1-.1*(X+65))-1)';
'bN(X) = .125*exp(-(X+65)/80)';
'INa(X,m,h) = gNa.*m.^3.*h.*(X-ENa)';
'IK(X,n) = gK.*n.^4.*(X-EK)';
'dv/dt=10-INa(v,m,h)-IK(v,n); v(0)=-65';
'dm/dt = aM(X).*(1-m)-bM(X).*m';
'dh/dt = aH(X).*(1-h)-bH(X).*h';
'dn/dt = aN(X).*(1-n)-bN(X).*n';
};
model=dsGenerateModel(eqns);
data=dsSimulate(model);
figure; plot(data.time,data.(data.labels{1}))

%%

% todo v0.1:
% 1. example analysis function (@FR) -- DONE
% 2. example plotting function (@dsPlotFR) -- DONE
% 3. import host:model (infinitebrain.org) -- DONE (for mechanism models)
% other additions:
% - monitor list (e.g., "monitor iNa, iK") (edit dsParseModelEquations) -- DONE
% - monitor functions (edit dsGenerateModel) -- DONE
% - set params of lower namespace (e.g,. "Na.g=100")

% todo v0.2:
% 1. design study framework -- DONE
% ...dsSimulate() with 'vary' and BatchManager() with studyinfo -- DONE
% ...AnalyzeStudy(data,@FR) -- DONE
% ... special case in pop eqns: vary() parsed in dsSimulate -- DONE

%% plan study framework
%   ideas: have quick option recognized by dsSimulate, or include .vary
%   in model, or other; in either case, may be best to have dsSimulate
%   be the single interface to simulations (instead of creating a separate
%   function like simstudy())

clear data; figure
vary={'pop1','gNa',[50 100 200]};
mod_set=dsVary2Modifications(vary); % {{'pop1','gNa',100},{'pop1','gNa',120}}
for i=1:length(mod_set)
  data(i)=dsSimulate('dv/dt=@M+10; {iNa,iK}@M','modifications',mod_set{i});
  subplot(1,length(mod_set),i); plot(data(i).time,data(i).(data(i).labels{1}))
  title(sprintf('gNa=%g',mod_set{i}{3}));
end

% the same data set and plot can be obtained directly from dsSimulate by:
vary={'pop1','gNa',[50 100 200]};
data=dsSimulate('dv/dt=@M+10; {iNa,iK}@M','vary',vary);
% [data.pop1_gNa]

vary={'','gNa',[50 100 200]};
data=dsSimulate('dv/dt=@M+10; {iNa,iK}@M','vary',vary);

vary={[],'gNa',[50 100 200]};
data=dsSimulate('dv/dt=@M+10; {iNa,iK}@M','vary',vary);

% special case of vary statement passed in equation to dsSimulate:
data=dsSimulate('dv/dt=@M/Cm+10; {iNa,iK}@M; vary(Cm=[.2 .5 1 5])');

data=dsSimulate('dv/dt=@M+amp; {iNa,iK}@M; vary(amp=[5 10 20])');
figure
for i=1:length(data)
  subplot(1,length(data),i); plot(data(i).time,data(i).(data(i).labels{1}));
  title(sprintf('%s=%g',data(i).varied{1},data(i).(data(i).varied{1})));
  ylim([-80 40]);
end

% auto-constructed search space given special case specification of what to vary
data=dsSimulate('dv/dt=@M+10; {iNa,iK}@M; vary(gNa=[50 100 200])');
% eqivalent manual construction of search space
vary={{'pop1','gNa',50},{'pop1','gNa',100},{'pop1','gNa',200}};
data=dsSimulate('dv/dt=@M+10; {iNa,iK}@M','vary',vary);
figure
for i=1:length(data)
  subplot(1,length(data),i); plot(data(i).time,data(i).(data(i).labels{1}));
  title(sprintf('%s=%g',data(i).varied{1},data(i).(data(i).varied{1})));
  ylim([-80 40]);
end

% Spike Monitor (threshold=10) for different tonic amplitudes and max sodium conductance
vary={'pop1','amp',2:2:60;'pop1','gNa',[100 120]};
data=dsSimulate('dv/dt=@current+amp; {iNa3,iK3}; monitor v.spikes(10)','vary',vary);
figure
a=[data.pop1_amp]; amps=unique(a);
b=[data.pop1_gNa]; gNas=unique(b);
spikes=cat(2,data.pop1_v_spikes);
for i=1:length(a), spikes(:,i)=spikes(:,i)+a(i); end
subplot(1,2,1); plot(data(1).time,spikes(:,b==gNas(1)),'k'); title(sprintf('gNa=%g',gNas(1))); axis tight
subplot(1,2,2); plot(data(1).time,spikes(:,b==gNas(2)),'k'); title(sprintf('gNa=%g',gNas(2))); axis tight
xlabel('time (ms)'); ylabel('tonic amplitude [uA/cm2]');
set(gca,'ytick',amps,'yticklabel',amps);

% adaptive exponential integrate and fire neuron
C=1; gl=1; El=-60; sT=1; vT=-45; sT=1; a=.5; tau=1; vmax=0; vr=-60;
s.pops.equations='dv/dt=(-gl*(v-El)+gl*sT*exp((v-vT)/sT)-w+I)/C; dw/dt=(a*(v-El)-w)/tau; if(v>vmax)(v=vr); v(0)=-70';
s.pops.parameters={'C',C,'gl',gl,'El',El,'vT',vT,'sT',sT,'a',a,'tau',tau,'I',25,'vr',vr,'vmax',vmax};
data=dsSimulate(s);
figure; plot(data.time,data.(data.labels{1}))
% eqivalent adaptive exponential integrate and fire
if 1
  % DEBUG. problem: this model is so equivalent that a new ODE solve file
  % is not being generate when it should be.
  data=dsSimulate('dv/dt=(-gl*(v-El)+gl*sT*exp((v-vT)/sT)-w+I)/C; if(v>0)(v=-60); dw/dt=(a*(v-El)-w)/tau; v(0)=-70; gl=1; El=-60; sT=1; vT=-45; C=1; a=.5; tau=1; I=25');
  figure; plot(data.time,data.(data.labels{1}))
end
% adaptive integrate and fire
data=dsSimulate('dv/dt=-(v-El)-w+100; if(v>0)(v=-60); dw/dt=a*(v-El)-w; a=.75; El=-60;');
figure; plot(data.time,data.(data.labels{1}))


%% Add post-processing

s=[];
s.populations(1).name='E';
s.populations(1).equations='dv/dt=current+10; {iNa,iK}; v(0)=-65';
s.populations(2).name='I';
s.populations(2).equations='dv/dt=current+10; {iNa,iK}; v(0)=-65';
data=dsSimulate(s);
figure; plot(data.time,data.(data.labels{1}))
data=dsSelect(data,'timelimits',[20 80]);
data=dsCalcFR(data,'variable','*_v','bin_size',30,'bin_shift',10);
figure; plot(data.time_FR,data.E_v_FR);

vary={'pop1','gNa',[50 100 200]};
data=dsSimulate('dv/dt=@M+10; {iNa,iK}@M','vary',vary);
data=AnalyzeStudy(data,@dsCalcFR,'bin_size',30,'bin_shift',10);
figure
FR=cat(2,data.pop1_v_FR);
subplot(1,2,1); plot(data.time_FR,FR);
xlabel('time'); ylabel('FR (Hz)'); title('gNa overlaid');
subplot(1,2,2); plot([data.pop1_gNa],mean(FR,1),'o-');
xlabel('gNa (mS/cm^2)'); ylabel('FR (Hz) avg over time');

vary={'pop1','gNa',[50 100 200]};
data=dsSimulate('dv/dt=@M+10; {iNa,iK}@M','vary',vary);
data=dsCalcFR(data,'variable','*_v','bin_size',30,'bin_shift',10);

% dsPlotFR: 1 pop, 1 cell, 0 varied
data=dsSimulate('dv/dt=@M+10; {iNa,iK}@M');
dsPlotFR(data,'bin_size',50,'bin_shift',15)

% dsPlotFR: 1 pop, 5 cells, 0 varied
data=dsSimulate('dv[5]/dt=@M+10*(t>150*rand(1,Npop)); {iNa,iK}@M','tspan',[0 300]);
dsPlotFR(data,'bin_size',50,'bin_shift',15)
figure; plot(data.time,data.(data.labels{1}))

% dsPlotFR: 2 pops, 1 cell/each, 0 varied
s=[];
s.pops(1).equations='dv/dt=@M+10*(t>150*rand(1,Npop)); {iNa,iK}@M; v(0)=-65';
s.pops(2).equations='dv/dt=@M+10*(t>150*rand(1,Npop)); {iNa,iK}@M; v(0)=-65';
data=dsSimulate(s,'tspan',[0 400]);
dsPlotFR(data,'bin_size',50,'bin_shift',15,'variable','*_v')

% dsPlotFR: 2 pops, 1 cell & 5 cells, 0 varied
s=[];
s.pops(1).equations='dv/dt=@M+10*(t>150*rand(1,Npop)); {iNa,iK}@M; v(0)=-65';
s.pops(2).equations='dv[5]/dt=@M+10*(t>150*rand(1,Npop)); {iNa,iK}@M; v(0)=-65';
data=dsSimulate(s,'tspan',[0 400]);
dsPlotFR(data,'bin_size',50,'bin_shift',15,'variable','*_v')

% dsPlotFR: 2 pops, 5 cells/each, 0 varied
s=[];
s.pops(1).equations='dv[5]/dt=@M+10*(t>150*rand(1,Npop)); {iNa,iK}@M; v(0)=-65';
s.pops(2).equations='dv[5]/dt=@M+10*(t>150*rand(1,Npop)); {iNa,iK}@M; v(0)=-65';
data=dsSimulate(s,'tspan',[0 600]);
dsPlotFR(data,'bin_size',50,'bin_shift',15,'variable','*_v')

% dsPlotFR: 1 pop, 1 cell, 1 varied
vary={'pop1','gNa',[50 200 400]};
data=dsSimulate('dv/dt=@M+10*(t>150*rand(1,Npop)); {iNa,iK}@M','vary',vary);
dsPlotFR(data,'bin_size',30,'bin_shift',10);

% dsPlotFR: 1 pop, 5 cells, 1 varied
vary={'pop1','gNa',[50 200 400]};
data=dsSimulate('dv[5]/dt=@M+10*(t>150*rand(1,Npop)); {iNa,iK}@M','vary',vary);
dsPlotFR(data,'bin_size',30,'bin_shift',10);

% dsPlotFR: 2 pops, 5 cells/each, 1 varied
vary={'pop1','gNa',[50 200 400]};
s=[];
s.pops(1).equations='dv[5]/dt=@M+10*(t>150*rand(1,Npop)); {iNa,iK}@M; v(0)=-65';
s.pops(2).equations='dv[5]/dt=@M+10*(t>150*rand(1,Npop)); {iNa,iK}@M; v(0)=-65';
data=dsSimulate(s,'tspan',[0 600],'vary',vary);
dsPlotFR(data,'bin_size',50,'bin_shift',15,'variable','*_v')

% dsPlotFR: 1 pop, 1 cell, 2 varied
vary={'pop1','gNa',[50 200 400];'pop1','gK',[25 50 100]};
data=dsSimulate('dv/dt=@M+10*(t>150*rand(1,Npop)); {iNa,iK}@M','vary',vary);
dsPlotFR(data,'bin_size',30,'bin_shift',10);

% dsPlotFR: 1 pop, 5 cells, 2 varied
vary={'pop1','gNa',[50 200 400];'pop1','gK',[25 50 100]};
data=dsSimulate('dv[5]/dt=@M+10*(t>150*rand(1,Npop)); {iNa,iK}@M','vary',vary);
dsPlotFR(data,'bin_size',30,'bin_shift',10);

% dsPlotFR: 2 pops, 5 cells/each, 2 varied
vary={'pop1','gNa',[50 200 400];'pop1','gK',[25 100]};
s=[];
s.pops(1).equations='dv[5]/dt=@M+10*(t>150*rand(1,Npop)); {iNa,iK}@M; v(0)=-65';
s.pops(2).equations='dv[5]/dt=@M+10*(t>150*rand(1,Npop)); {iNa,iK}@M; v(0)=-65';
data=dsSimulate(s,'tspan',[0 600],'vary',vary);
dsPlotFR(data,'bin_size',50,'bin_shift',15,'variable','*_v')

% dsPlotFR: 1 pop, 1 cell, 3 varied
vary={'pop1','gNa',[50 400];'pop1','gK',[25 100];'pop1','amp',[10 20]};
data=dsSimulate('dv/dt=@M+amp*(t>150*rand(1,Npop)); {iNa,iK}@M','vary',vary);
dsPlotFR(data,'bin_size',30,'bin_shift',10);

% dsPlotFR: 1 pop, 5 cells, 3 varied
vary={'pop1','gNa',[50 400];'pop1','gK',[25 100];'pop1','amp',[10 20]};
data=dsSimulate('dv[5]/dt=@M+amp*(t>150*rand(1,Npop)); {iNa,iK}@M','vary',vary);
dsPlotFR(data,'bin_size',30,'bin_shift',10);

% dsPlotFR: 2 pops, 5 cells/each, 3 varied
vary={'pop1','gNa',[50 400];'pop1','gK',[25 100];'pop2','amp',[10 20]};
s=[];
s.pops(1).equations='dv[5]/dt=@M+amp*(t>150*rand(1,Npop)); {iNa,iK}@M; v(0)=-65; amp=10';
s.pops(2).equations='dv[5]/dt=@M+amp*(t>150*rand(1,Npop)); {iNa,iK}@M; v(0)=-65; amp=10';
data=dsSimulate(s,'tspan',[0 600],'vary',vary);
dsPlotFR(data,'bin_size',50,'bin_shift',15,'variable','*_v')

% ------------------------------------------------
% Super simple single-parameter rate effects
data=dsSimulate('dv/dt=@M+10; {iNa,iK}@M; vary(gNa=[50 100 200])');
dsPlotFR(data,'bin_size',30,'bin_shift',10);
% ------------------------------------------------

%% Add Experiments

if 1
  % DEBUG (broke after adding dsUpdateStudy)

  % !rm solve_ode.m params.mat
  % ------------------------------------------------
  % Super simple experiment
    data=dsSimulate('dv/dt=@M+10; {iNa,iK}@M','experiment',@dsProbeFI);
    dsPlotFR(data,'bin_size',30,'bin_shift',10);
  % ------------------------------------------------

  % !rm solve_ode.m params.mat
  s=[];
  s.pops(1).equations='dv[5]/dt=@M; {iNa,iK}@M; v(0)=-65';
  s.pops(2).equations='dv[5]/dt=@M; {iNa,iK}@M; v(0)=-65';
  data=dsSimulate(s,'experiment',@dsProbeFI,'amplitudes',-10:10:10,'tspan',[0 600]);
  dsPlotFR(data,'bin_size',50,'bin_shift',15)

end

%% infbrain:modelID
[~,host]=system('echo $HOSTNAME');
if 0 %strcmp(strtrim(host),'jason-pc') % if on developer computer
  addpath ~/code/dnsim/matlab/dependencies % (passiveFtp, @ftp)

  NaMechID=57; % Na+ mechanism in infbrain
  model=dsImportModel('infbrain:57'); % import mechanism model (Na+ channel)

  s=dsCheckSpecification('dv/dt=current; {infbrain:57,iK}');
  s.populations
  s=dsCheckSpecification('dv/dt=current; {infbrain:57}; {iK}');
  s.populations
  s=dsCheckSpecification('dv/dt=current; infbrain:{57}; {iK}');
  s.populations
  s=dsCheckSpecification('dv/dt=@M; infbrain:{57}@M; {iK}@M');
  s.populations
  s=dsCheckSpecification('dv/dt=@M; infbrain:{57}@M; {iK@M}');
  s.populations
  s=dsCheckSpecification('dv/dt=@M; infbrain:{57@M}; {iK@M}');
  s.populations
  s=dsCheckSpecification('dv/dt=@M; {infbrain:57,iK}@M');
  s.populations
  s=dsCheckSpecification('dv/dt=@M; {iK@M}; {infbrain:57@M}');
  s.populations

  data=dsSimulate('dv/dt=current; {infbrain:57,iK}');
  figure; plot(data.time,data.(data.labels{1}))

  data=dsSimulate('dv/dt=current; infbrain:{57}; {iK}');
  figure; plot(data.time,data.(data.labels{1}))

  data=dsSimulate('dv/dt=@M; {infbrain:57,iK}@M');
  figure; plot(data.time,data.(data.labels{1}))

  data=dsSimulate('dv/dt=@M; infbrain:{57}@M; {iK}@M');
  figure; plot(data.time,data.(data.labels{1}))

  data=dsSimulate('dv/dt=@M; {iK@M}; {infbrain:57@M}');
  figure; plot(data.time,data.(data.labels{1}))

  data=dsSimulate('dv/dt=@M; {iK@M}; infbrain:{57@M}');
  figure; plot(data.time,data.(data.labels{1}))

  % todo: test these (importing population model, ...)
  % syntax to support integrating mechanism models from DBs:
  % equations='...infbrain:{iNa,iK}@M'
  % equations='...{infbrain:iNa,modeldb:iK}@M'
  % equations='...{infbrain:iNa@M, modeldb:iK@N}'
  % integrating population models
  % populations.name='infbrain:RE'
  % accessing and coupling network models:
  % model1=[already created]
  % model2=dsImportModel('infbrain:PFC');
  % modifications.connections.mechanism_list='GABAa'
  % modifications.connections.source='TC'
  % modifications.connections.target='PY5'
  % model=dsCombineModels(model1,model2,'modifications',modifications);

  % todo: consider pulling experiments from remote server as well:
  % eqn='dv/dt=@M+10; ib:{iNa,iK}@M';
  % data=dsSimulate(eqn,'experiment','ib:@dsProbeFI');

  % ------------------------------------------------------------------
  % Example of concise powerful statements in DynaSim:
  % single statement to (1) specify a 4-cell noisy HH-type population model combining
  % locally and remotely-stored mechanism sub-models and (2) specify and run three
  % 200ms simulations varying a parameter gna of the remote Na+ channel mechanism (ib:57):
  data=dsSimulate('dv[4]/dt=@M+50*rand(1,Npop); {ib:57,iK}@M; vary(gna=[50 75 150])','tspan',[0 200]);
  % plot overlay of cell voltage traces for gna=75 (second value for varied parameter)
  figure; plot(data(2).time,data(2).(data(2).labels{1})); xlabel('time (ms)'); ylabel(data(2).labels{1});
  % plot Firing Rates in each of the three simulations
  for i=1:3 % loop over gna parameter values (there was one simulation per value)
    dsPlotFR(data(i),'bin_size',30,'bin_shift',10);
  end
  % plot how Firing Rate varies with gna (across elements of the data array)
  dsPlotFR(data,'bin_size',30,'bin_shift',10);
  % access simulated model components stored in higher-level specification
  data(1).model.specification.populations
  % access simulated model definition
  data(1).model
  % access simulated model parameters
  data(1).model.parameters
  % ------------------------------------------------------------------
end

%% Add dsUpdateStudy() to dsSimulate()

% test dsCheckStudyinfo:
studyinfo=dsCheckStudyinfo([])
studyinfo.simulations.sim_id=1;
studyinfo=dsCheckStudyinfo(studyinfo);
studyinfo.simulations

if 0 % no longer working ... but not necessary until dsMonitorStudy is reintroduced
  % test dsStudyinfoIO:
  %   loading: studyinfo=dsStudyinfoIO([],study_file,[id,verbose_flag])
  %   saving:  dsStudyinfoIO(studyinfo,[study_file,id,verbose_flag]);
  % verbose:
  dsStudyinfoIO(studyinfo,'',[],1); % saving studyinfo
  dsStudyinfoIO(studyinfo,'',201,1); % saving studyinfo (process ID=201)
  studyinfo=dsStudyinfoIO([],'studyinfo.mat',[],1); % loading studyinfo
  studyinfo=dsStudyinfoIO([],'studyinfo.mat',201,1); % loading studyinfo (process ID=201)
  % not verbose:
  dsStudyinfoIO(studyinfo); % saving studyinfo
  studyinfo=dsStudyinfoIO([],'studyinfo.mat'); % loading studyinfo
  studyinfo=dsStudyinfoIO([]); % loading studyinfo
  studyinfo=dsStudyinfoIO; % loading studyinfo
  study_dir=pwd;
  studyinfo=dsStudyinfoIO([],study_dir); % loading studyinfo

  studyinfo=dsCheckStudyinfo(study_dir)
  studyinfo=dsCheckStudyinfo('studyinfo.mat')
  studyinfo=dsCheckStudyinfo(study_dir,'process_id',201)
  studyinfo=dsCheckStudyinfo('studyinfo.mat','process_id',201)
end

addpath(dynasim_path)

% without using studyinfo (save_data_flag=0): #############################

% test sim: local, one sim, not compiled
data=dsSimulate('dv/dt=@M+10; {iNa,iK}@M');
data=dsSimulate('dv/dt=@M+10; {iNa,iK}@M');
dsPlotFR(data);

% test sim: local, 3 sims, not compiled
data=dsSimulate('dv/dt=@M+10; {iNa,iK}@M; vary(gNa=[50 100 200])');
data=dsSimulate('dv/dt=@M+10; {iNa,iK}@M; vary(gNa=[50 100 200])');
dsPlotFR(data);
% --------------------------------------
% ERROR --
% IMPORTANT TODO: make it so that varied params maintain consistent
% parameter name (pop1_iNa_gNa should not become pop1_gNa when gNa is varied)
% --------------------------------------

% test sim: local, one sim, compiled
data=dsSimulate('dv/dt=@M+10; {iNa,iK}@M','mex_flag',1);
data=dsSimulate('dv/dt=@M+10; {iNa,iK}@M','mex_flag',1);
dsPlotFR(data);

% test sim: local, 3 sims, compiled
data=dsSimulate('dv/dt=@M+10; {iNa,iK}@M; vary(gNa=[50 100 200])','mex_flag',1);
data=dsSimulate('dv/dt=@M+10; {iNa,iK}@M; vary(gNa=[50 100 200])','mex_flag',1);
dsPlotFR(data);

% % using studyinfo (save_data_flag=1): ###################################
%
% test sim: local, one sim, not compiled
dsSimulate('dv/dt=@M+10; {iNa,iK}@M','save_data_flag',1,'verbose_flag',1);
data=dsSimulate('dv/dt=@M+10; {iNa,iK}@M','save_data_flag',1,'verbose_flag',1);
dsPlotFR(data);

% test sim: local, one sim, not compiled -- single user-assigned study_dir
dsSimulate('dv/dt=@M+10; {iNa,iK}@M','save_data_flag',1,'study_dir','StudyA','verbose_flag',1);
% - saves data to StudyA
data=dsSimulate('dv/dt=@M+10; {iNa,iK}@M','save_data_flag',1,'study_dir','StudyA','verbose_flag',1);
% - loads existing data from StudyA
dsPlotFR(data);

% test sim: local, 3 sims, not compiled
data=dsSimulate('dv/dt=@M+10; {iNa,iK}@M; vary(gNa=[50 100 200])','save_data_flag',1,'study_dir','StudyB','verbose_flag',1);
dsPlotFR(data);
load('StudyB/studyinfo.mat','studyinfo')
studyinfo
studyinfo.simulations(1)
studyinfo.simulations(2)
studyinfo.simulations(2).machine_info

% test sim: local, one sim, compiled
dsSimulate('dv/dt=@M+10; {iNa,iK}@M','mex_flag',1,'save_data_flag',1,'study_dir','StudyC','verbose_flag',1);
data=dsSimulate('dv/dt=@M+10; {iNa,iK}@M','mex_flag',1,'save_data_flag',1,'study_dir','StudyC','verbose_flag',1);
dsPlotFR(data);

% test sim: local, 3 sims, compiled
dsSimulate('dv/dt=@M+10; {iNa,iK}@M; vary(gNa=[50 100 200])','mex_flag',1,'save_data_flag',1,'study_dir','StudyD','verbose_flag',1);
data=dsSimulate('dv/dt=@M+10; {iNa,iK}@M; vary(gNa=[50 100 200])','mex_flag',1,'save_data_flag',1,'study_dir','StudyD','verbose_flag',1);
dsPlotFR(data);

% test tspan=[0 200]
data=dsSimulate('dv/dt=@M+10; {iNa,iK}@M; vary(gNa=[50 100 200])','tspan',[0 200]);
data(1).time(end)
data=dsSimulate('dv/dt=@M+10; {iNa,iK}@M','mex_flag',1,'tspan',[0 200]);
data(1).time(end)
data=dsSimulate('dv/dt=@M+10; {iNa,iK}@M','mex_flag',1,'tspan',[0 200],'save_data_flag',1);
data(1).time(end)
data=dsSimulate('dv/dt=@M+10; {iNa,iK}@M','mex_flag',1,'tspan',[0 200],'save_data_flag',1,'study_dir','StudyE');
data(1).time(end)
dsPlotFR(data);

% ############################################################
if 1 % errors to debug and features to implement

  % IMPORTANT TODO: make it so that varied params maintain consistent
  % parameter name (pop1_iNa_gNa should not become pop1_gNa when gNa is varied)
  data=dsSimulate('dv/dt=@M+10; {iNa,iK}@M');
  data=dsSimulate('dv/dt=@M+10; {iNa,iK}@M; vary(gNa=[50 100 200])');
  data(1).model.parameters
  data(2).model.parameters
  data(3).model.parameters
  % DONE

  % NEXT: add dsCreateBatch() and test cluster sims (cluster_flag=1) -- DONE

  % todo: replace model load/save by dsImportModel() and ExportModel()
  % (in dsUpdateStudy(), dsCreateBatch(), ...)
end

if 0
  % NOT WORKING: (todo: need to redesign how model data is handled/transferred)
  model=dsImportModel('infbrain:120'); % import neuron model (Morris-Lecar)

  % todo: update browse_dnsim()
end

%% add dsCreateBatch() (cluster_flag=1) and dsMonitorStudy()

model=dsGenerateModel('dv/dt=(@M+10)/Cm; Cm=1; {iNa,iK}@M');
model.parameters

modifications_set=dsVary2Modifications({'pop1','Cm',[1 2]},model);
modifications_set{:}

% test sim: cluster, one sim, not compiled
% verbose_flag
[data,studyinfo]=dsSimulate('dv/dt=@M+10; {iNa,iK}@M',...
  'cluster_flag',1,'study_dir','ClusterTest','verbose_flag',1);
dsMonitorStudy(studyinfo.study_dir);
run(studyinfo.simulations.job_file);
dsMonitorStudy(studyinfo.study_dir);
load(fullfile(studyinfo.study_dir,'studyinfo.mat'),'studyinfo');
studyinfo.simulations
% not verbose_flag
[data,studyinfo]=dsSimulate('dv/dt=@M+10; {iNa,iK}@M',...
  'cluster_flag',1,'study_dir','ClusterTest');
run(studyinfo.simulations.job_file);
load(fullfile(studyinfo.study_dir,'studyinfo.mat'),'studyinfo');
studyinfo.simulations
data=dsImport(studyinfo.simulations.data_file);%load(studyinfo.simulations.data_file,'data')
figure; plot(data.time,data.(data.labels{1}))
dsPlotFR(data)

% test sim: cluster, 3 sims, not compiled
[data,studyinfo]=dsSimulate('dv/dt=(@M+10)/Cm; Cm=1; {iNa,iK}@M; vary(Cm=[1 2 3])',...
  'cluster_flag',1,'study_dir','ClusterTestSet','verbose_flag',1);
dsMonitorStudy(studyinfo.study_dir);
run(studyinfo.simulations(1).job_file);
dsMonitorStudy(studyinfo.study_dir);
run(studyinfo.simulations(2).job_file);
dsMonitorStudy(studyinfo.study_dir);
run(studyinfo.simulations(3).job_file);
dsMonitorStudy(studyinfo.study_dir);
studyinfo=dsMonitorStudy(studyinfo.study_dir);
[studyinfo.simulations.duration]

% -------------------
% GOOD EXAMPLE:
[data,studyinfo]=dsSimulate('dv/dt=(@M+10)/Cm; Cm=1; {iNa,iK}@M; vary(Cm=[1 2 3])',...
  'cluster_flag',1,'study_dir','ClusterTestSetb','verbose_flag',1);
if isempty(data) % will not be empty if study is ran again after completion
  run(studyinfo.simulations(1).job_file);
  run(studyinfo.simulations(2).job_file);
  run(studyinfo.simulations(3).job_file);
  data=dsImport(studyinfo);
end
dsMonitorStudy(studyinfo);
dsPlotFR(data);
% -------------------

if 1
  [data,studyinfo]=dsSimulate('dv/dt=@M+10; {iNa,iK}@M; vary(gNa=[50 100 200])',...
    'cluster_flag',1,'study_dir','ClusterTestSetc','verbose_flag',1);
  run(studyinfo.simulations(1).job_file);
end

% test sim: cluster, one sim, compiled
[data,studyinfo]=dsSimulate('dv/dt=@M+10; {iNa,iK}@M',...
  'cluster_flag',1,'mex_flag',1,'study_dir','ClusterTestMEX','verbose_flag',1);
dsMonitorStudy(studyinfo.study_dir);
run(studyinfo.simulations.job_file);
dsMonitorStudy(studyinfo.study_dir);

% test sim: cluster, 3 sims, compiled
[data,studyinfo]=dsSimulate('dv/dt=(@M+10)/Cm; Cm=1; {iNa,iK}@M; vary(Cm=[1 2 3])',...
  'cluster_flag',1,'mex_flag',1,'study_dir','TestClusterSetMEX','verbose_flag',1);
dsMonitorStudy(studyinfo.study_dir);
run(studyinfo.simulations(1).job_file);
dsMonitorStudy(studyinfo.study_dir);
run(studyinfo.simulations(2).job_file);
dsMonitorStudy(studyinfo.study_dir);
run(studyinfo.simulations(3).job_file);
dsMonitorStudy(studyinfo.study_dir);
studyinfo=dsMonitorStudy(studyinfo.study_dir);
[studyinfo.simulations.duration]

% test sim: cluster, 3 sims in 2 jobs, not compiled
% set sims_per_job=2
[data,studyinfo]=dsSimulate('dv/dt=(@M+10)/Cm; Cm=1; {iNa,iK}@M; vary(Cm=[1 2 3])',...
  'cluster_flag',1,'mex_flag',0,'sims_per_job',2,'study_dir','ClusterTestSet2','verbose_flag',1);
dsMonitorStudy(studyinfo.study_dir);
run(studyinfo.simulations(1).job_file);
studyinfo=dsMonitorStudy(studyinfo.study_dir);
exist(studyinfo.simulations(1).data_file)
exist(studyinfo.simulations(2).data_file)
exist(studyinfo.simulations(3).data_file)
run(studyinfo.simulations(2).job_file);
studyinfo=dsMonitorStudy(studyinfo.study_dir);
exist(studyinfo.simulations(1).data_file)
exist(studyinfo.simulations(2).data_file)
exist(studyinfo.simulations(3).data_file)
run(studyinfo.simulations(3).job_file);
studyinfo=dsMonitorStudy(studyinfo.study_dir);
exist(studyinfo.simulations(1).data_file)
exist(studyinfo.simulations(2).data_file)
exist(studyinfo.simulations(3).data_file)

% test sim: cluster, 3 sims in 2 jobs, compiled
% set sims_per_job=2
[data,studyinfo]=dsSimulate('dv/dt=(@M+10)/Cm; Cm=1; {iNa,iK}@M; vary(Cm=[1 2 3])',...
  'cluster_flag',1,'mex_flag',1,'sims_per_job',2,'study_dir','ClusterTestSetMEX2','verbose_flag',1);
dsMonitorStudy(studyinfo.study_dir);
run(studyinfo.simulations(1).job_file);
studyinfo=dsMonitorStudy(studyinfo.study_dir);
exist(studyinfo.simulations(1).data_file)
exist(studyinfo.simulations(2).data_file)
exist(studyinfo.simulations(3).data_file)
run(studyinfo.simulations(2).job_file);
studyinfo=dsMonitorStudy(studyinfo.study_dir);
exist(studyinfo.simulations(1).data_file)
exist(studyinfo.simulations(2).data_file)
exist(studyinfo.simulations(3).data_file)
run(studyinfo.simulations(3).job_file);
studyinfo=dsMonitorStudy(studyinfo.study_dir);
exist(studyinfo.simulations(1).data_file)
exist(studyinfo.simulations(2).data_file)
exist(studyinfo.simulations(3).data_file)

% test email notification when study simulations are complete
[data,studyinfo]=dsSimulate('dv/dt=(@M+10)/Cm; Cm=1; {iNa,iK}@M; vary(Cm=[1 2 3])',...
  'cluster_flag',1,'mex_flag',1,'sims_per_job',2,'email','','study_dir','ClusterTestSetMEX3','verbose_flag',1);
run(studyinfo.simulations(1).job_file);
run(studyinfo.simulations(3).job_file);
dsMonitorStudy(studyinfo.study_dir);

% tests with disk_flag=1:

% test sim: cluster, 3 sims, not compiled, save csv
[data,studyinfo]=dsSimulate('dv/dt=(@M+10)/Cm; Cm=1; {iNa,iK}@M; vary(Cm=[1 2 3])',...
  'cluster_flag',1,'disk_flag',1,'study_dir','TestDiskClusterSet','verbose_flag',1);
dsMonitorStudy(studyinfo.study_dir);
run(studyinfo.simulations(1).job_file);
dsMonitorStudy(studyinfo.study_dir);
run(studyinfo.simulations(2).job_file);
dsMonitorStudy(studyinfo.study_dir);
run(studyinfo.simulations(3).job_file);
dsMonitorStudy(studyinfo.study_dir);
data=dsImport('TestDiskClusterSet');
dsPlotFR(data);

% test sim: cluster, 3 sims, compiled, save csv
[data,studyinfo]=dsSimulate('dv/dt=(@M+10)/Cm; Cm=1; {iNa,iK}@M; vary(Cm=[1 2 3])',...
  'cluster_flag',1,'disk_flag',1,'mex_flag',1,'study_dir','TestDiskClusterSetMEX','verbose_flag',1);
dsMonitorStudy(studyinfo.study_dir);
run(studyinfo.simulations(1).job_file);
dsMonitorStudy(studyinfo.study_dir);
run(studyinfo.simulations(2).job_file);
dsMonitorStudy(studyinfo.study_dir);
run(studyinfo.simulations(3).job_file);
dsMonitorStudy(studyinfo.study_dir);
data=dsImport('TestDiskClusterSetMEX');
dsPlotFR(data);

% test sim: local, 3 sims, not compiled, save csv
[data,studyinfo]=dsSimulate('dv/dt=(@M+10)/Cm; Cm=1; {iNa,iK}@M; vary(Cm=[1 2 3])',...
  'cluster_flag',0,'disk_flag',1,'mex_flag',0,'study_dir','TestDiskLocalSet','verbose_flag',1);
dsPlotFR(data);
data2=dsImport('TestDiskLocalSet');
dsPlotFR(data2);

% test sim: local, 3 sims, compiled, save csv
[data,studyinfo]=dsSimulate('dv/dt=(@M+10)/Cm; Cm=1; {iNa,iK}@M; vary(Cm=[1 2 3])',...
  'cluster_flag',0,'disk_flag',1,'mex_flag',1,'study_dir','TestDiskLocalSetMEX','verbose_flag',1);
dsPlotFR(data);
data2=dsImport('TestDiskLocalSetMEX');
dsPlotFR(data2);

%% troubleshooting

% check parameter naming for user-supplied parameters
s=[];
s.pops.equations='dv/dt=(@M+10)/Cm; Cm=1; {iNa,iK}@M';
s.pops.parameters={'gNa',150,'Cm',2};
m=dsGenerateModel(s);
m.parameters
m.ODEs.pop1_v
m.functions.pop1_iNa_INa

s=[];
s.pops.equations='dv/dt=(@M+10)/Cm; Cm=1; {iNa,iK}@M';
s.pops.parameters={'gNa',150,'Cm',2};
s.cons.mechanism_list={'AMPA'};
s.cons.parameters={'tauDx',3,'g_SYN',.1};
m=dsGenerateModel(s);
m.parameters
m.ODEs.pop1_pop1_AMPA_s
m.functions.pop1_pop1_AMPA_ISYN

if 1 % todo -- DONE: FIX: data sets returned are identical and only reflect the last result
  [data,studyinfo]=dsSimulate('dv/dt=(@M+10)/Cm; Cm=1; {iNa,iK}@M; vary(gNa=[100 200])',...
  'cluster_flag',1,'mex_flag',1,'study_dir','Test4','verbose_flag',1);
  dsMonitorStudy(studyinfo.study_dir);
  run(studyinfo.simulations(1).job_file);
  dsMonitorStudy(studyinfo.study_dir);
  run(studyinfo.simulations(2).job_file);
  dsMonitorStudy(studyinfo.study_dir);
  data=dsImport('Test4');
  dsPlotFR(data);
  % should look like this:
  [data,studyinfo]=dsSimulate('dv/dt=(@M+10)/Cm; Cm=1; {iNa,iK}@M; vary(gNa=[100 200])',...
  'cluster_flag',0,'mex_flag',1,'verbose_flag',1);
  dsPlotFR(data);

  [data,studyinfo]=dsSimulate('dv/dt=(@M+10)/Cm; Cm=1; {iNa,iK}@M; vary(gNa=[100 200])',...
  'cluster_flag',1,'mex_flag',0,'study_dir','Test4b','verbose_flag',1);
  dsMonitorStudy(studyinfo.study_dir);
  run(studyinfo.simulations(1).job_file);
  dsMonitorStudy(studyinfo.study_dir);
  run(studyinfo.simulations(2).job_file);
  dsMonitorStudy(studyinfo.study_dir);
  data=dsImport('Test4b');
  dsPlotFR(data);
end

% A) * FIX: tspan not adjustable after compiling MEX -- DONE
data=dsSimulate('dv/dt=@M+10; {iNa,iK}@M; vary(gNa=[50 100 200])',...
  'tspan',[0 200],'mex_flag',1,'verbose_flag',1);
dsPlotFR(data); dsPlotFR(data(2));
data=dsSimulate('dv/dt=@M+10; {iNa,iK}@M; vary(gNa=[50 100 200])',...
  'tspan',[0 100],'mex_flag',1,'verbose_flag',1);
dsPlotFR(data); dsPlotFR(data(2));
% works

data=dsSimulate('dv/dt=@M+10; {iNa,iK}@M; vary(gNa=[50 100 200])',...
  'tspan',[0 200],'mex_flag',1,'save_data_flag',1,'verbose_flag',1);
dsPlotFR(data); dsPlotFR(data(2));
data=dsSimulate('dv/dt=@M+10; {iNa,iK}@M; vary(gNa=[50 100 200])',...
  'tspan',[0 100],'mex_flag',1,'save_data_flag',1,'verbose_flag',1);
dsPlotFR(data); dsPlotFR(data(2));
% works

data=dsSimulate('dv/dt=@M+10; {iNa,iK}@M; vary(gNa=[50 100 200])',...
  'tspan',[0 200],'mex_flag',1,'save_data_flag',1,'study_dir','StudyG','verbose_flag',1);
dsPlotFR(data); dsPlotFR(data(2));
data=dsSimulate('dv/dt=@M+10; {iNa,iK}@M; vary(gNa=[50 100 200])',...
  'tspan',[0 100],'mex_flag',1,'save_data_flag',1,'study_dir','StudyG','verbose_flag',1);
dsPlotFR(data); dsPlotFR(data(2));
% works as it should; second simulation loads the result saved from the
% first; that is why the second data does not have 100ms in it. possible
% changes if loading the existing data is not desired:  (1) add tspan to
% the data_file name, (2) compare parameters between new and old in
% addition to the data_file name before deciding whether to load existing
% data or run a new simulation.
data=dsSimulate('dv/dt=@M+10; {iNa,iK}@M; vary(gNa=[50 100 200])',...
  'tspan',[0 100],'mex_flag',1,'save_data_flag',1,'study_dir','StudyG',...
  'overwrite_flag',1,'verbose_flag',1);
dsPlotFR(data); dsPlotFR(data(2));
% works

% B) * todo: debug need for pause (see above) -- DONE
% - is an issue only 1st time a solver file is created if immediately used
% SOLVED: added milliseconds to solver file name
if 1 % original example of bug
  % adaptive exponential integrate and fire neuron
  C=1; gl=1; El=-60; sT=1; vT=-45; sT=1; a=.5; tau=1; vmax=0; vr=-60;
  s.pops.equations='dv/dt=(-gl*(v-El)+gl*sT*exp((v-vT)/sT)-w+I)/C; dw/dt=(a*(v-El)-w)/tau; if(v>vmax)(v=vr); v(0)=-70';
  s.pops.parameters={'C',C,'gl',gl,'El',El,'vT',vT,'sT',sT,'a',a,'tau',tau,'I',25,'vr',vr,'vmax',vmax};
  data=dsSimulate(s);
  figure; plot(data.time,data.(data.labels{1}))
  % eqivalent adaptive exponential integrate and fire
  data=dsSimulate('dv/dt=(-gl*(v-El)+gl*sT*exp((v-vT)/sT)-w+I)/C; if(v>0)(v=-60); dw/dt=(a*(v-El)-w)/tau; v(0)=-70; gl=1; El=-60; sT=1; vT=-45; C=1; a=.5; tau=1; I=25');
  figure; plot(data.time,data.(data.labels{1}))
end

% todo: debug experiments (might be related to pause issue)
% SOLVED

%% More tests
% 1. test ic
    % DONE
% 2. add special cases of monitor (lists, keyword "functions")
    % DONE
% 3. eliminate need to create solver m-file when solve_file is set to MEX
    % run batch with profiler (is creating solve_file limiting factor?)
    % (see below)
    % Q: why is studyinfo.simulation.duration so much longer than sim elapsed time?
    % DONE
% 4. more study/cluster stuff
    % DONE

%% test: 'ic' option in dsSimulate()
data=dsSimulate('dv/dt=-v+15*sin(2*pi*t*3)','tspan',[0 10],'ic',10);
figure; plot(data.time,data.pop1_v)

data=dsSimulate('dv[2]/dt=-v+15*sin(2*pi*t*3)','tspan',[0 10],'ic',[10 20]);
figure; plot(data.time,data.pop1_v)

data=dsSimulate('dv/dt=-v+15*sin(2*pi*t*3); dw/dt=-w+15*sin(2*pi*t*1);','tspan',[0 10],'ic',[10 20]);
figure; plot(data.time,data.pop1_v,'b'); hold on; plot(data.time,data.pop1_w,'r');

data=dsSimulate('dv[2]/dt=-v+15*sin(2*pi*t*3); dw[2]/dt=-w+15*sin(2*pi*t*1);','tspan',[0 10],'ic',[10 12 20 22]);
figure; plot(data.time,data.pop1_v,'b'); hold on; plot(data.time,data.pop1_w,'r');

%% add monitor special cases (eg., lists, keyword "functions")
% - monitor list (e.g., "monitor iNa, iK") (edit dsParseModelEquations)
% - monitor functions (edit dsGenerateModel)

model=dsGenerateModel('dv/dt=@M+10; monitor functions; {iNa3,iK3}@M;');
model.monitors
data=dsSimulate('dv/dt=@M+10; monitor functions; {iNa3,iK3}@M;');
data
figure; plot(data.time,data.pop1_iK3_I)

model=dsGenerateModel('dv/dt=@M+10; monitor iNa3.functions; {iNa3,iK3}@M;');
model.monitors
data=dsSimulate('dv/dt=@M+10; monitor iNa3.functions; {iNa3,iK3}@M;');
data
figure; plot(data.time,data.pop1_iNa3_aH)

model=dsGenerateModel('dv/dt=@M+10; monitor iNa3_functions; {iNa3,iK3}@M;');
model.monitors
data=dsSimulate('dv/dt=@M+10; monitor iNa3_functions; {iNa3,iK3}@M;');
data
figure; plot(data.time,data.pop1_iNa3_aH)

model=dsGenerateModel('dv/dt=@M+10; monitor iK3.functions; {iNa3,iK3}@M;');
model.monitors
data=dsSimulate('dv/dt=@M+10; monitor iK3.functions; {iNa3,iK3}@M;');
data
figure; plot(data.time,data.pop1_iK3_aN)

model=dsGenerateModel('dv/dt=@M+10; monitor iK3_functions; {iNa3,iK3}@M;');
model.monitors
data=dsSimulate('dv/dt=@M+10; monitor iK3_functions; {iNa3,iK3}@M;');
data
figure; plot(data.time,data.pop1_iK3_aN)

model=dsGenerateModel('dv/dt=@M+10; monitor iK3.I, iK3.aN; {iNa3,iK3}@M;');
model.monitors

model=dsGenerateModel('dv/dt=@M+10; monitor iK3.I, iK3.aN, iNa3.functions; {iNa3,iK3}@M;');
model.monitors


%% profile study batch ("profile viewer")

% test sim: cluster, 3 sims, compiled
[data,studyinfo]=dsSimulate('dv/dt=(@M+10)/Cm; Cm=1; {iNa,iK}@M; vary(Cm=[1 2 3])',...
  'cluster_flag',1,'mex_flag',1,'study_dir','TestProfileStudy','verbose_flag',1);
dsMonitorStudy(studyinfo.study_dir);
run(studyinfo.simulations(1).job_file);
dsMonitorStudy(studyinfo.study_dir);
run(studyinfo.simulations(2).job_file);
dsMonitorStudy(studyinfo.study_dir);
run(studyinfo.simulations(3).job_file);
dsMonitorStudy(studyinfo.study_dir);
studyinfo=dsMonitorStudy(studyinfo.study_dir);
[studyinfo.simulations.duration]

% * todo: eliminate need to create solver m-file when solve_file is set for a
  % MEX file (compiled_flag=1). this will probably save significant time.
% DONE

%% other study/cluster stuff
% - fix setting 'varied' for batch sims (see notes in dsImport())
  % DONE
% - troubleshoot issue at line 1103 (see above)
  % DONE
% - test cluster_flag=1 with disk_flag=1
  % DONE

%% extract population names from equations

spec=dsCheckSpecification('TC:dv/dt=0');
spec.populations
spec=dsCheckSpecification('[dv/dt=0]');
spec.populations
spec=dsCheckSpecification('[TC:dv/dt=0]');
spec.populations
spec=dsCheckSpecification('[dv/dt=@M;ib:{Na,K}@M][du/dt=@M;{Na,K,Ca}]');
spec.populations(1)
spec.populations(2)
spec=dsCheckSpecification('[TC:dv/dt=@M;ib:{Na,K}@M][RE:du/dt=@M;{Na,K,Ca}]');
spec.populations(1)
spec.populations(2)
spec=dsCheckSpecification('[dv[2]/dt=@M;ib:{Na,K}@M][RE:du[3]/dt=@M;{Na,K,Ca}]');
spec.populations(1)
spec.populations(2)
spec=dsCheckSpecification('[dv[2]/dt=@M;ib:{Na,K}@M][RE:du[3]/dt=@M;{Na,K,Ca};dw[3]/dt=10]');
spec.populations(1)
spec.populations(2)

% todo: edit dsCheckSpecification to take parameters in .equations and move
% them to .parameters...
  % DONE
spec=dsCheckSpecification('[TC:dv/dt=0; p=3]');
spec.populations

s=[];
s.pops.equations='[TC:dv/dt=p; p=3]';
s.pops.parameters={'p',4};
spec=dsCheckSpecification(s);
spec.populations
m=dsGenerateModel(spec);
m.parameters

s=[];
s.pops.equations='[TC:dv/dt=p]';
s.pops.parameters={'p',4};
spec=dsCheckSpecification(s);
spec.populations
m=dsGenerateModel(spec);
m.parameters

model=dsGenerateModel('dv/dt=@M+10; gNa=150; monitor iK3.I, iK3.aN; {iNa3,iK3}@M;');
model.parameters
data=dsSimulate(model);
data
figure;
subplot(1,2,1); plot(data.time,data.pop1_iK3_I)
model=dsGenerateModel('dv/dt=@M+10; gNa=50; monitor iK3.I, iK3.aN; {iNa3,iK3}@M;');
data=dsSimulate(model);
subplot(1,2,2); plot(data.time,data.pop1_iK3_I)

s=[];
s.pops.equations='[TC:dv/dt=@M+10;{iNa,iK}@M][RE:dv/dt=@M;{iNa,iK}@M]';
s.cons.source='TC';
s.cons.target='RE';
s.cons.mechanism_list='AMPA@M';
s.cons.parameters={'g_SYN',1};
m=dsGenerateModel(s);
m.ODEs.RE_v
d=dsSimulate(m);
figure; plot(d.time,d.TC_v,'b',d.time,d.RE_v,'r'); legend('TC','RE')

s=[];
s.pops.equations='[TC:dv/dt=@M+10;{iNa,iK}@M][RE:dv/dt=@M;{iNa,iK}@M]';
s.cons.source='TC';
s.cons.target='RE';
s.cons.mechanism_list='AMPA@M';
s.cons.parameters={'g_SYN',1};
d=dsSimulate(s);
d.model.ODEs.RE_v
figure; plot(d.time,d.TC_v,'b',d.time,d.RE_v,'r'); legend('TC','RE')


%% GO LIVE %%%%%%%%%%%%%%%%%%%%%%%%%%%%%%%%%%%%%%%%%%%%%%%%%%%%%%%%%%

% - clean functions -- DONE
% - do tests on scc2 (simply copy dynasim and run simple study from dir)
% - prepare for github and convert demo into tutorial

% github (+ installation instructions)
  % - 2nd commit: standardize function help
% readthedocs {tutorial and function reference (=help sections)}
  % - turn this demo_solvers.m into a tutorial
% mailing lists (users and developers)


%% add HDF-style loading of partial data sets
% ref: http://www.mathworks.com/help/matlab/import_export/load-parts-of-variables-from-mat-files.html
dsExportData(data(1),'filename','data.mat');
data=dsImport('data.mat','toilim',[50 100],'variables','pop1_v')

%% add parallel computing (parfor_flag=1) -- for DynaSim paper
% ref: http://www.bu.edu/tech/support/research/training-consulting/online-tutorials/matlab-pct/run-batch-on-scc
% note: SCC supports qsub submissions for up to 12 cores
% ref: http://vtchl.uiuc.edu/node/537
% ref: http://vtchl.uiuc.edu/sites/default/files/MATLAB_Report.pdf
% relevant functions: parfor (parpool, matlabpool), spmd
if 0
  % note: this works if solve_file is defined:
  % on cluster (parallel sims within a job)
  num_cores=3;
  parpool(num_cores)
  parfor i=1:6
    dsSimulate('dv/dt=@M+10; monitor iNa3_functions; {iNa3,iK3}@M;','solve_file',solve_file);
    disp(i);
  end
  delete(gcp)
end

% ######################################################
% on SCC: works
if 0
  [data,studyinfo]=dsSimulate('dv/dt=(@M+10)/Cm; Cm=1; {iNa,iK}@M; vary(Cm=[1 2 3])',...
    'cluster_flag',1,'mex_flag',1,'sims_per_job',2,'parfor_flag',1,'verbose_flag',1);
  % local test:
  [data,studyinfo]=dsSimulate('dv/dt=(@M+10)/Cm; Cm=1; {iNa,iK}@M; vary(Cm=[1 2 3])',...
    'cluster_flag',1,'mex_flag',1,'sims_per_job',2,'parfor_flag',1,'verbose_flag',1);
  dsMonitorStudy(studyinfo);
  [s,r]=system(sprintf('cat %s',studyinfo.simulations(1).job_file));
  eval(r);
  dsMonitorStudy(studyinfo);
  exist(studyinfo.simulations(1).data_file)
  exist(studyinfo.simulations(2).data_file)
  exist(studyinfo.simulations(3).data_file)
end
% ######################################################

if 0
  % on local machine (use recursive call to dsSimulate...):
  clear data
  parpool(num_cores)
  parfor i=1:6
    data(i)=dsSimulate('dv/dt=@M+10; monitor iNa3_functions; {iNa3,iK3}@M;','solve_file',solve_file);
    disp(i);
  end
  delete(gcp)
  dsPlotFR(data);
end

%poolobj=gcp('nocreate');

if 0 % todo: debug
  % ######################################################
  parpool(num_cores);
  data=dsSimulate('dv/dt=@M+i;i=0;{iNa,iK}@M;vary(i=[0 10 20])','parfor_flag',1,'verbose_flag',1);
  dsPlotFR(data);
  delete(gcp);
  % ######################################################
end

% legacy:
if 0
  matlabpool('open',num_cores);
  solve_file=fullfile(dynasim_path,'solve','solve_ode_20160118232404_080.m');
  parfor i=1:6
    dsSimulate('dv/dt=@M+10; monitor iNa3_functions; {iNa3,iK3}@M;','solve_file',solve_file);%,'verbose_flag',1);
    disp(i);
  end
  matlabpool('close');
end

% consider: compilation for standalone app without need for matlab licenses:
% see: http://www.bu.edu/tech/support/research/software-and-programming/common-languages/matlab/standalone/

%% Study post-processing

[~,studyinfo]=dsSimulate('dv/dt=@M+10; {iNa,iK}@M; vary(gNa=[50 100 200])','save_data_flag',1);
results=AnalyzeStudy(studyinfo,@dsCalcFR);

data=dsSimulate('dv/dt=@M+10; {iNa,iK}@M; vary(gNa=[50 100 200])','save_data_flag',1);
results=AnalyzeStudy(data,@dsCalcFR);

results=AnalyzeStudy('ClusterTestSet2',@dsCalcFR);

% todo: add cluster support for analyses and plotting
% results=AnalyzeStudy(data,@dsCalcFR,'cluster_flag',1);

%% add model i/o (XPP, NeuroML) -- for DynaSim paper

% XPP

% NeuroML


%% update upload/download model -- not for DynaSim paper

if 0

  addpath code/dnsim/matlab/dependencies/jsonlab

  % e.g., model=dsImportModel('infbrain:120'); % import neuron model (Morris-Lecar)

  loadjson(savejson('',model))
  model % .namespaces is altered (shouldn't affect simulation though...)

  [mech_paths,mech_files]=dsLocateModelFiles(model);

  % upload:
    % zip files (json + mechs)
    % transfer to infinitebrain.org

  % download:
    % transfer from infinitebrain.org
    % unzip files
    % ...
end

  % todo: test these (importing population model, ...)
% syntax to support integrating mechanism models from DBs:
% equations='...infbrain:{iNa,iK}@M'
% equations='...{infbrain:iNa,modeldb:iK}@M'
% equations='...{infbrain:iNa@M, modeldb:iK@N}'
% integrating population models
% populations.name='infbrain:RE'
% accessing and coupling network models:
% model1=[already created]
% model2=dsImportModel('infbrain:PFC');
% modifications.connections.mechanism_list='GABAa'
% modifications.connections.source='TC'
% modifications.connections.target='PY5'
% model=dsCombineModels(model1,model2,'modifications',modifications);

% todo: consider pulling experiments from remote server as well:
% eqn='dv/dt=@M+10; ib:{iNa,iK}@M';
% data=dsSimulate(eqn,'experiment','ib:@dsProbeFI');

% todo: update browse_dnsim() for new structures

%% add ability to submit cluster jobs from local host (NOT IMPORTANT)

% save model to .mat
% scp files (.mat + mechs) to study_dir on cluster_node (login node)
% ssh single command: matlab -r "load(modelfile); dsSimulate(model,...)"
%   - might need helper expect script
%   - get login credentials from dsSimulate options (cluster_user, cluster_host; prompt for password)

% todo: edit dsMonitorStudy() to check status of cluster jobs from local host
% (e.g., ssh: matlab -r "dsMonitorStudy(remote_studyinfo_file)"; prompt for login credentials or take as options)

%% other changes
% - fix coder.varsize() for compiling variable population sizes
% - create example optimization function
% - create visualizers (BrowseData, BrowseStudy)
% - further develop analysis stream (AnalyzeStudy, dsSelect)

% minor changes:
% - rename verbose_flag to verbose_flag_flag (in this script and all functions)


% #########################################################################

% ... (introduce ICs in equations) ...
eqns={
  's=10; r=27; b=2.666';
  'dx/dt=s*(y-x); x(0)=1';
  'dy/dt=r*x-y-x*z; y(0)=2';
  'dz/dt=-b*z+x*y; z(0)=.5';
};
data=dsSimulate(eqns,'tspan',[0 100]);
figure; plot(data.pop1_x,data.pop1_z);
xlabel('x'); ylabel('z'); title('Lorenz equations')

% Morris-Lecar equations
% ... (introduce functions) ...
eqns={
  'vk=-84;vl=-60; vca=120';
  'i=100; gk=8; gl=2; gca=4; c=20';
  'v1=-1.2; v2=18; v3=2; v4=30; phi=.04';
  'minf(v)=.5*(1+tanh((v-v1)/v2))';
  'winf(v)=.5*(1+tanh((v-v3)/v4))';
  'lamw(v)=phi*cosh((v-v3)/(2*v4))';
  'dv/dt=(i+gl*(vl-v)+gk*w*(vk-v)+gca*minf(v)*(vca-v))/c';
  'dw/dt=lamw(v)*(winf(v)-w)';
};
data=dsSimulate(eqns,'tspan',[0 1000],'ic',[-60 .015])
data.model.functions
figure; plot(data.time,data.pop1_v);
xlabel('time (ms)'); ylabel('v'); title('Morris-Lecar neuron')

% ... (introduce monitors) ...
eqns={
  'vk=-84;vl=-60; vca=120';
  'i=100; gk=8; gl=2; gca=4; c=20';
  'v1=-1.2; v2=18; v3=2; v4=30; phi=.04';
  'minf(v)=.5*(1+tanh((v-v1)/v2))';
  'winf(v)=.5*(1+tanh((v-v3)/v4))';
  'lamw(v)=phi*cosh((v-v3)/(2*v4))';
  'Ica(v)=gca*minf(v)*(v-vca)';
  'Ik(v,w)=gk*w*(v-vk)';
  'dv/dt=(i+gl*(vl-v)-Ik(v,w)-Ica(v))/c';
  'dw/dt=lamw(v)*(winf(v)-w)';
  'monitor Ica, Ik, v.spikes(0)';
};
data=dsSimulate(eqns,'tspan',[0 1000],'ic',[-60.899 .014873])
data.model.monitors
data.model.functions
data.model.functions.pop1_Ica
figure;
subplot(2,1,1); plot(data.time,data.pop1_v)
xlabel('time (ms)'); ylabel('v'); title('Morris-Lecar neuron')
  % ... add spikes to plot ...
subplot(2,1,2); plot(data.time,data.pop1_Ica)
xlabel('time (ms)'); ylabel('Ica');

% HH w/ ic simulator option
eqns={
  'gNa=120; gK=36; Cm=1';
  'INa(v,m,h) = gNa.*m.^3.*h.*(v-50)';
  'IK(v,n) = gK.*n.^4.*(v+77)';
  'aM(v) = (2.5-.1*(v+65))./(exp(2.5-.1*(v+65))-1)';
  'bM(v) = 4*exp(-(v+65)/18)';
  'aH(v) = .07*exp(-(v+65)/20)';
  'bH(v) = 1./(exp(3-.1*(v+65))+1)';
  'aN(v) = (.1-.01*(v+65))./(exp(1-.1*(v+65))-1)';
  'bN(v) = .125*exp(-(v+65)/80)';
  'dv/dt = (10-INa(v,m,h)-IK(v,n))/Cm';
  'dm/dt = aM(v).*(1-m)-bM(v).*m';
  'dh/dt = aH(v).*(1-h)-bH(v).*h';
  'dn/dt = aN(v).*(1-n)-bN(v).*n';
};
data=dsSimulate(eqns,'tspan',[0 200],'ic',[-65 .1 .1 0]);
figure; plot(data.time,data.(data.labels{1}))

% TC-RE network
% ... set 'netcon',rand(N_post,N_pre) ...

% Two-compartment PY-FS network
% ... set 'netcon',eye(N_pop) ...


% adding spike monitor to existing model structure
% method 1:
tmp=sPING_model.specification.populations(1).equations;
sPING_model.specification.populations(1).equations=[tmp 'monitor v.spikes(0)'];
m=dsGenerateModel(sPING_model.specification);
m.monitors
% method 2:
m=sPING_model;
m.monitors.E_v_spikes='0'; % []
d=dsSimulate(m)

% adding spike monitor to existing model structure
sPING_model.monitors.E_v_spikes=[]; % set to threshold (as string; eg, '0')

% #########################################################################

% Wilson-Cowan equations
% phase plane plot
% ...

% Lotka-Volterra equations
% phase plane plot
% ...

% Lorenz equations
% phase plane plot
% ...

% Izhekivich equations
% ...

% Leaky integrate-and-fire equations
% ...

% Exponential LIF equations
% ...

% Adaptive exponential LIF equations
% ...

% FitzHugh-Nagumo equations
% ...

% Morris-Lecar equations
% ...

% Morris-Lecar with mechanisms
% ... introduce namespaces and linkers ...

% Hodgkin-Huxley equations
% ...

% Hodgkin-Huxley with mechanisms
% ...

% NETWORKS

% ##############################################3
% XPP examples
% ##############################################3
% the brusselator (a nice chemical oscillator)
eqns='b=3; a=1; du/dt=a-(b+1)*u+v*u^2; dv/dt=b*u-v*u^2';
data=dsSimulate(eqns,'tspan',[0 100],'ic',[1 1.5]);
figure; plot(data.pop1_u,data.pop1_v); xlabel('u'); ylabel('v');
eqns='b=3; a=1; du/dt=a-(b+1)*u+v*u^2; dv/dt=b*u-v*u^2; u(0)=1; v(0)=1.5';
data=dsSimulate(eqns,'tspan',[0 100],'dt',.01);
figure; plot(data.pop1_u,data.pop1_v); xlabel('u'); ylabel('v');
% ##############################################3
% clock model (discontinuous ODE) - a linearly decaying spiral that is kicked out
a=.1;b=1;k=.5; % ...
% ##############################################3
% the force duffing equation (error:explodes)
eqns='a=3e-10;w=1;mu=.25;f(t)=a*cos(w*t); dx/dt=v; dv/dt=mu*v+x*(1-x^2)+f(t)';
data=dsSimulate(eqns,'tspan',[0 200],'ic',[0 0],'dt',.0001);
figure; plot(data.pop1_x,data.pop1_v); xlabel('x'); ylabel('v');
% ##############################################3
% levy flight (error:explodes)
eqns={...
'xi=.2; sig=1; mu=sig/xi; f(u)=mu+sig*(u^(-xi)-1)/xi';
'dx/dt=x+f(rand)*(-1+2*(rand>.5))';
'dy/dt=y+f(rand)*(-1+2*(rand>.5))';
};
data=dsSimulate(eqns,'tspan',[0 2000],'ic',[0 0]);
figure; plot(data.pop1_x,data.pop1_y); xlabel('x'); ylabel('y');
% ##############################################3
% Morris-Lecar model from Ermentrout chapter in Koch & Segev
% A simple membrane model.
eqns={...
'iapp=.2; phi=1';
'v1=-.01; v2=.15; v3=.1; v4=.145; gca=1.33';
'vk=-.7; vl=-.5; gk=2; gl=.5; om=1';
'minf(v)=.5*(1+tanh((v-v1)/v2))';
'ninf(v)=.5*(1+tanh((v-v3)/v4))';
'lamn(v)=phi*cosh((v-v3)/(2*v4))';
'ica(v)=gca*minf(v)*(v-1)';
'dv/dt=iapp+gl*(vl-v)+gk*w*(vk-v)-ica(v)';
'dw/dt=lamn(v)*(ninf(v)-w)';
};
data=dsSimulate(eqns,'tspan',[0 100],'ic',[0 0]);
figure; plot(data.pop1_v,data.pop1_w); xlabel('v'); ylabel('w');
% ##############################################3
```
