## Supplemental File 1 for "A biophysical model of striatal microcircuits suggests delta/theta-rhythmically interleaved gamma and beta oscillations mediate periodicity in motor control": dsTutorial.m

```
% %%%%%%%%%%%%%%%%%%%%%%%%%%%%%%%%%%%%%%%%%%%%%%%%%%%%%%%%%%%%%%%%%%%%%%%%
%% DynaSim Tutorial
% %%%%%%%%%%%%%%%%%%%%%%%%%%%%%%%%%%%%%%%%%%%%%%%%%%%%%%%%%%%%%%%%%%%%%%%%
%{
Download DynaSim toolbox from https://github.com/dynasim/dynasim.
  or, download using git: git clone https://github.com/dynasim/dynasim.git
Further documentation is available at: [readthedocs link].
Sign up for user mailing list at: https://groups.google.com/forum/#!forum/dynasim-users.

Tip: In Matlab, you can obtain more information associated with any function "FUNCTION_NAME"
by entering "help FUNCTION_NAME" in the command window. Use the "See also" list
at the end of the help section to browse through related help documentation.
%}

% Get ready...

% Add DynaSim to path if it's not already there
if exist('setupDynaSimPath','file')
    setupDynaSimPath;
else
    error('Add the DynaSim folder to the MATLAB path - e.g. run addpath(genpath(DynaSimPath))');
end

% Set where to save outputs
output_directory = dsGetConfig('demos_path');

% move to root directory where outputs will be saved
mkdir(output_directory);
cd(output_directory);

% Here we go!

% %%%%%%%%%%%%%%%%%%%%%%%%%%%%%%%%%%%%%%%%%%%%%%%%%%%%%%%%%%%%%%%%%%%%%%%%
%% DEFINING AND SIMULATING MODELS
% %%%%%%%%%%%%%%%%%%%%%%%%%%%%%%%%%%%%%%%%%%%%%%%%%%%%%%%%%%%%%%%%%%%%%%%%

%% Lorenz equations with phase plot

% DynaSim makes it easy to simulate arbitrary systems of ordinary
% differential equations. Simply write out the system in a cell array of
% strings, separating equations into different strings or the same string
% separated by semicolons.

eqns={
  's=10; r=27; b=2.666';
  'dx/dt=s*(y-x)';
  'dy/dt=r*x-y-x*z';
  'dz/dt=-b*z+x*y';
};
data=dsSimulate(eqns,'tspan',[0 100],'ic',[1 2 .5],'solver','rk4');
% tspan: time limits on integration [ms]
% ic: initial conditions
% solver: numerical method to use (default: rk4 = "4th-order Runge-Kutta")

% Simulated data are returned in a DynaSim data structure:
data
data.labels % list of state variables (and functions) that are stored in data structure
data.simulator_options % dsSimulate options used to solve the model system

% The simulated model is also stored as a DynaSim model structure:
data.model
data.model.parameters       % parameters                      (s,r,b,Npop)
data.model.state_variables  % state variables                 (x,y,z)
data.model.ODEs             % ordinary differential equations (dx/dt,dy/dt,dz/dt)
data.model.ICs              % initial conditions              (x(0),y(0),z(0))

% The model structure can be obtained without simulating it using the
% dsGenerateModel function:
model=dsGenerateModel(eqns);

% Every component of the model is assigned to a "population", and the 
% population name (default: 'pop1') is prepended to all variable and
% function names. For instance, state variable "x" becomes "pop1_x" by
% default. Reasons for this and how to adjust the population name will be 
% discussed later. The size of each population is always stored as an 
% additional parameter ("Npop").

% All models are numerically integrated using a DynaSim solver function
% created uniquely for a given model and stored in a directory named
% "solve". The file that solves the system (i.e,. numerically integrates
% it) is stored in data.simulator_options and can be viewed or rerun afterwards:
edit(data.simulator_options.solve_file)

% Simulated data can be easily plotted using the resulting data structure:
figure; plot(data.pop1_x,data.pop1_z); 
title('Lorenz equations'); xlabel('x'); ylabel('z')

% Tip: since data.labels contains a list of the recorded variables, the 
% first state variable in any system can be plotted generically using:
figure; plot(data.time,data.(data.labels{1}));
title('Lorenz equations'); xlabel('t'); ylabel('x')

%% Leaky integrate-and-fire

% DynaSim also makes it easy to incorporate conditional statements in a ODE
% system using syntax: if(condition)(action). "condition" must be a valid
% Matlab expression that evaluates to true or false. "action" can be one or
% more Matlab statements separated by semicolons.

eqns='tau=10; R=10; E=-70; dV/dt=(E-V+R*1.55)/tau; if(V>-55)(V=-75)';
data=dsSimulate(eqns,'tspan',[0 200],'ic',-75);

% view the solver file:
edit(data.simulator_options.solve_file)

% plot the simulated data:
figure; plot(data.time,data.pop1_V); 
xlabel('time (ms)'); ylabel('V'); title('Leaky integrate-and-fire (LIF) neuron')

%% Leaky integrate-and-fire with spike monitor

% The DynaSim data structure always contains the model state variables,
% time vector, and a copy of the DynaSim model structure that was
% simulated. Additionally, functions and spikes (i.e., upward threshold 
% crossings) can be recorded and returned in the DynaSim data structure
% if indicated using the "monitor" keyword.

% Syntax:
% monitor FUNCTION
% monitor VARIABLE.spikes(THRESHOLD)

% Spikes are recorded as a binary point process with 1 where spikes occur
% (i.e., where THRESHOLD is exceeded) in state variable "VARIABLE" and 0 
% everywhere else. They are stored in a field named POPULATION_VARIABLE_spikes.

% example spike monitor
eqns={
  'tau=10; R=10; E=-70; I=1.55; thresh=-55; reset=-75';
  'dV/dt=(E-V+R*I)/tau; if(V>thresh)(V=reset)';
  'monitor V.spikes(thresh)';
};
data=dsSimulate(eqns,'tspan',[0 200],'ic',-75);
% insert spike where LIF resets occur
data.pop1_V(data.pop1_V_spikes==1)=20;
% plot the LIF response with spikes
figure; plot(data.time,data.pop1_V);
xlabel('time (ms)'); ylabel('V'); title('LIF with spikes')

%% Izhikevich neuron with noisy drive 
% (reference: p274 of "Dynamical Systems in Neuroscience" by Izhikevich)

% Model functions can be incorporated in a model as easily as parameters
% and returned in the DynaSim data structure along with state variables if
% specified using the "monitor" keyword.

% The following example defines and monitors a time-varying input function 
% I(t) that turns on while t>ton and t<toff, and is scaled by a noisy 
% factor using the built-in "rand" function. The example also demonstrates 
% how initial conditions can be defined in the equations themselves, 
% instead of passing them as an option to dsSimulate.

eqns={
  'C=100; vr=-60; vt=-40; k=.7; Iapp=70; ton=200; toff=800';
  'a=.03; b=-2; c=-50; d=100; vpeak=35';
  'dv/dt=(k*(v-vr)*(v-vt)-u+I(t))/C; v(0)=vr';
  'du/dt=a*(b*(v-vr)-u); u(0)=0';
  'if(v>vpeak)(v=c; u=u+d)';
  'I(t)=Iapp*(t>ton&t<toff)*(1+.5*rand)'; % define applied input using reserved variables 't' for time and 'dt' for fixed time step of numerical integration
  'monitor I';                            % indicate to store applied input during simulation
};
data=dsSimulate(eqns,'tspan',[0 1000]);
% plot the simulated voltage and monitored input function
figure; 
subplot(2,1,1); plot(data.time,data.pop1_v); % plot voltage
xlabel('time (ms)'); ylabel('v'); title('Izhikevich neuron')
subplot(2,1,2); plot(data.time,data.pop1_I); % plot input function
xlabel('time (ms)'); ylabel('Iapp');

% note: "t", "dt", and "T" are special variables that can be used in model
% equations. "t" represents the current time point of the simulation. 
% "dt" is the fixed time step used for numeric integration. "T" is the full
% simulated time vector defined before simulation begins.

%% Hodgkin-Huxley (HH) equations

eqns={
  'gNa=120; gK=36; Cm=1';
  'INa(v,m,h) = gNa.*m.^3.*h.*(v-50)';
  'IK(v,n) = gK.*n.^4.*(v+77)';
  'dv/dt = (10-INa(v,m,h)-IK(v,n))/Cm; v(0)=-65';
  'dm/dt = aM(v).*(1-m)-bM(v).*m; m(0)=.1';
  'dh/dt = aH(v).*(1-h)-bH(v).*h; h(0)=.1';
  'dn/dt = aN(v).*(1-n)-bN(v).*n; n(0)=0';
  'aM(v) = (2.5-.1*(v+65))./(exp(2.5-.1*(v+65))-1)';
  'bM(v) = 4*exp(-(v+65)/18)';
  'aH(v) = .07*exp(-(v+65)/20)';
  'bH(v) = 1./(exp(3-.1*(v+65))+1)';
  'aN(v) = (.1-.01*(v+65))./(exp(1-.1*(v+65))-1)';
  'bN(v) = .125*exp(-(v+65)/80)';
};
data=dsSimulate(eqns);
figure; plot(data.time,data.(data.labels{1}))
xlabel('time (ms)'); ylabel('membrane potential (mV)'); title('Hodgkin-Huxley neuron')

% Equations can also be stored in a text file (e.g., 'HH.pop') and 
% simulated by passing the file name to dsSimulate:
data=dsSimulate('HH.pop');

% Model files used for simulation can be easily located:
[~,eqnfile]=dsLocateModelFiles(data); % eqnfile is a cell array of file names
  % note: dsLocateModelFiles accepts DynaSim structures (model, data, specification, or studyinfo)
  % as inputs and returns all associated model files.
[~,eqnfile]=dsLocateModelFiles('HH.pop');  
  % note: can also be used to locate .pop and .mech model files
% Open the model file:
dsEditModelFiles('HH.pop'); % compare to the above list of equations
% tip: you can use dsLocateModelFiles/dsEditModelFiles to see what model files will be used
% before simulation if you are unsure. DynaSim always searches the current
% directory first, then (sub)directories of <DynaSim>/models, then the
% complete Matlab path.

% To many modelers, the classic HH neuron may represent the point at which 
% model building becomes a tedious, error-prone process, and model reconfiguration and
% prototyping become arduous, especially when several ion currents are
% involved, or multiple compartments, cell types, and networks. DynaSim
% takes advantage of the "mechanism" concept to simplify the process of
% working with larger models like these.

%% Mechanisms

% Concept:
% Physical systems can often be decomposed intuitively into sub-systems with 
% internal states governed by dynamics; their dynamics may be attributable to
% internal mechanisms or mechanisms that connect them to other sub-systems.

% NEURON uses this concept to define separate mechanism models for ion currents,
% pumps, buffers, etc, that are stored in .modl files and "inserted" into 
% neuronal compartments (sub-systems). DynaSim uses a similar approach, 
% defining mechanisms (stored in separate model files) that are included in 
% populations (sub-systems) or used to connect two populations. In NEURON, 
% the mathematical details relating mechanisms to the full ODE system is 
% hidden and only an abstract understanding of the model composition is 
% needed to construct large models. Like NEURON, users do not need to 
% understand the details of how mechanisms work to benefit from them. 
% However, unlike NEURON, the relationship between mechanisms and the full 
% ODE system is transparent in DynaSim and easily customizable to users.

% One of the unique benefits of DynaSim is its ability to transparently
% modularize large models using mechanisms, to enable constructing large
% models from pre-existing smaller ones (i.e., mechanism or single
% population models). This also provides the unique ability to seemlessly
% transition from building small models of a few differential equations to
% large models of thousands or more equations.

% Next, we'll look at an example that uses mechanisms to simulate a more
% complicated Hodgkin-Huxley-type model. After demonstrating how mechanisms
% can simplify the modeling process, we'll walk through the process of
% decomposing a set of equations into a smaller set of mechanisms; then,
% we'll demonstrate how mechanisms can be used to simplify the process of 
% modeling large networks of biophysically-detailed neurons without needing
% to understand or create new mechanisms.

% Example of simplified specification of bursting neuron using mechanisms:
eqns='dv/dt=5+@current; {iNaF,iKDR,iM}; gNaF=100; gKDR=5; gM=1.5; v(0)=-70';
data=dsSimulate(eqns,'tspan',[0 200]);
figure; plot(data.time,data.(data.labels{1}))
xlabel('time (ms)'); ylabel('membrane potential (mV)'); title('Intrinsically Bursting neuron')
% The model uses three ion current mechanisms stored in separate files: (in <dynasim>/models)
  % iNaF.mech: fast sodium current (mechanism model)
  % iKDR.mech: delayed rectifier potassium current (mechanism model)
  % iM.mech: slow muscarinic potassium current (mechanism model)

% Other currents can be incorporated into the model simply by adding their names 
% to the list in "{}" without the need to edit/write out a lot of equations.

% ---------------------------------------------------------------------

% Decomposing a set of equations into a smaller set of mechanisms:

% Important concepts:
% 1. Defining mechanisms: group equations into sub-model that is
%       mechanistically meaningful.
% 2. Linking mechanisms: link variables and/or functions in mechanism file 
%       to equations outside the file defining them.
% 3. Namespaces: unique population and mechanism-level identifiers to prevent
%       name conflicts between parameters/variables/functions defined in different places.

% Grouping Hodgkin-Huxley equations to define mechanisms:
eqns={
  'dv/dt = (10+INa(v,m,h)+IK(v,n))/Cm; Cm=1; v(0)=-65'; % voltage
  % -- iNa.mech ----
  'INa(v,m,h) = -gNa.*m.^3.*h.*(v-50); gNa=120';  % sodium current
  'dm/dt = aM(v).*(1-m)-bM(v).*m; m(0)=.1';       % - activation
  'dh/dt = aH(v).*(1-h)-bH(v).*h; h(0)=.1';       % - inactivation
  'aM(v) = (2.5-.1*(v+65))./(exp(2.5-.1*(v+65))-1)';
  'bM(v) = 4*exp(-(v+65)/18)';
  'aH(v) = .07*exp(-(v+65)/20)';
  'bH(v) = 1./(exp(3-.1*(v+65))+1)';
  % -- iK.mech -----
  'IK(v,n) = -gK.*n.^4.*(v+77); gK=36';           % potassium current
  'dn/dt = aN(v).*(1-n)-bN(v).*n; n(0)=0';        % - activation
  'aN(v) = (.1-.01*(v+65))./(exp(1-.1*(v+65))-1)';
  'bN(v) = .125*exp(-(v+65)/80)';
};

dsEditModelFiles('iNa.mech');
dsEditModelFiles('iK.mech');
  % note: "X" is an optional reserved variable that can be used in mechanism 
  % files as an alias to the first state variable in the top-level population
  % equations (e.g., "v" in the next HH model). Alternatively, the variable
  % "v" could be used in the mechanism file; however, doing so increases
  % the chance of incompatibility (e.g., an equation defining dV/dt is 
  % incompatible with a function f(v) using a lower-case variable).
  
% Mechanisms have direct access to all state variables found in the 
% DynaSim specification for their population equations, but no access to 
% state variables in other mechanisms unless those mechanisms have provided 
% linkers for accessing their components.

% When one equation needs to access a variable or function defined
% in a different namespace (e.g., INa(v,m,h) in INa.mech needs to be used
% in dv/dt defined elsewhere), the two need to be "linked". Linking is
% achieved by indicating in mechanism files - a target IDENTIFIER into which 
% a variable or function should be inserted; and defining elsewhere - 
% equations that include the IDENTIFIER where mechanism variables and 
% functions should be inserted.
% Linking involves:
% 1. In mechanism file: specifying a target location IDENTIFIER into which a variable
% or function should be substituted by some operation (e.g., "+=", as in
% C++ compound addition). Example: "@IDENTIFIER += -INa(v,m,h)" means add the
% mechanism function INa to all locations where @IDENTIFIER appears in the same
% population.
% 2. In target equation (e.g., in another mechanism or equation passed to
% dsSimulate): include the target location IDENTIFIER in the equation
% where the variable or function should be substituted. Example:
% "dX/dt=@IDENTIFIER" means insert all functions linked to @IDENTIFIER by all
% mechanisms of the same population.

% Tip: use the '@' character to identify the target location for a variable or
% function to be linked. Then, '@' appearing in an ODE will always imply 
% that the ODE depends on variables or functions defined somewhere else
% (i.e., outside its namespace). 

% Linking Na+ and K+ current mechanisms to dv/dt in HH model:
% Mechanism linkers:
%   iNa.mech: @current += -INa(v,m,h)
%   iK.mech:  @current += -IK(v,n)
% Population equations: dv/dt=@current
% Linked equation: dv/dt=-INa(v,m,h)-IK(v,n)

% Mechanism files must include the desired equations as well as a linker
% statement indicating the target IDENTIFIER(s) for their variables and 
% functions to be incorporated in equations defined elsewhere. 

% Demonstrate mechanism-based HH model simulation:
data=dsSimulate('dv/dt=10+@current/Cm; Cm=1; v(0)=-65; {iNa,iK}');

% As models get larger and incorporate an increasing number of model files, 
% the chance of two files using the same name for a variable or function increases. 
% To avoid name conflicts in such cases, each variable and function in DynaSim is 
% prepended by a namespace that indicates the population and mechanism in 
% which it is defined.

% DynaSim parses each mechanism file in turn, adds a distinguishing prefix
% to each variable and function defined therein (designating "namespaces"),
% and links their variables and functions to corresponding targets found 
% in equations outside the mechanism file (.mech). (see dsCheckModel for more details)

% namespace = <population>_ or <population>_<mechanism>_
data.model.parameters
data.model.functions
data.model.state_variables
data.model.namespaces % columns = {old_name, new_name, namespace, type}                       % 
data

% All functions of a mechanism can be monitored using the following syntax: 
% monitor MECHANISM.functions. Example:
data=dsSimulate('dv/dt=10+@current; {iNa,iK}; monitor iNa.functions');
data.model.functions
data.model.monitors
data

%% DynaSim specification structure 
% The above approach is sufficient for building single-compartment models
% with arbitrary complexity. However, larger multicompartment and network
% models require defining multiple compartments or cell types and connecting 
% them. DynaSim simplifies building these models as well. Behind the scenes, 
% DynaSim always converts the user-supplied equations into a DynaSim "specification"
% structure that can be used to specify any model at a low or high level of
% abstraction. Learning to define specification structures can greatly
% facilitate (1) constructing large network models and (2) parameterizing
% control of model parameters and mechanism lists in Matlab scripts. 
% The latter advantage provides a higher degree of control that is recommended for conducting
% computational research without the need to frequently manipulate equation strings.

% In the above examples, DynaSim first splits the user-supplied population
% equations into a 'mechanism_list', 'parameters', and 'equations'; it then
% stores those separately in a structure (the "specification" structure)
% and assigns a 'name' and 'size' to the (implicit) population. Practically 
% speaking, this approach requires the user to store the above model 
% information in a specification structure (instead of an equation string 
% or cell array of strings). The specification structure allows multiple
% populations to be defined (e.g., different compartments or cell types)
% and connected (e.g., by synapse mechanisms from a 'source' to a 'target').

% Schema for DynaSim specification structure (select fields shown):
% .populations(i): contains info for defining independent population models (required)
%   .name           : name of population (default: 'pop1')
%   .size           : number of elements in population (i.e., # cells) (default: 1)
%   .equations      : string listing equations (required)
%   .mechanism_list : cell array listing mechanisms (default: [])
%   .parameters     : parameters to assign across all equations in the population. provide as cell array list of key/value pairs (default: [])
%                     {'param1',value1,'param2',value2,...}
% .connections(j): contains info for linking population models (default: [])
%   .source         : name of source population (required if >1 pops)
%   .target         : name of target population (required if >1 pops)
%   .mechanism_list : list of mechanisms that link two populations (required)
%   .parameters     : parameters to assign across all equations in
%                     mechanisms in this connection's mechanism_list. (default: [])

% see dsCheckSpecification for more details.

% inspect the specification structure for the Hodgkin-Huxley model:
data=dsSimulate('dv/dt=10+@current; {iNa,iK}');
data.model.specification.populations
data.model.specification.populations.equations
data.model.specification.populations.mechanism_list
data.model.specification.populations.parameters
data.model.specification.populations.size
data.model.specification.populations.name
data.model.specification.connections % no connections between populations

% define the Hodgkin-Huxley model with mechanisms in a DynaSim
% specification structure:
% method #1 -- list mechanisms in equation string
specification=[];
specification.populations.equations='dv/dt=10+@current; {iNa,iK}';
data=dsSimulate(specification);
%figure; plot(data.time,data.(data.labels{1}))

% method #2 -- list mechanisms separately in mechanism_list cell array
specification=[];
specification.populations.equations='dv/dt=10+@current';
specification.populations.mechanism_list={'iNa','iK'};
data=dsSimulate(specification);
%figure; plot(data.time,data.(data.labels{1}))
% Tip: use method #2 in script with mechanism names stored in variables
% to easily control the mechanisms that are incorporated in a model.

% Define networks using DynaSim specification:
% introduce connections --> multi-compartments and networks of populations ...

% minimal example to demonstrate connections
s=[];
s.populations(1).equations='I:dv/dt=(-70-v+15.5)/10; if(v>-55)(v=-75)'; % LIF neuron
s.populations(2).equations='E:dv/dt=-.01*v+@current';
s.connections(1).source='I';
s.connections(1).target='E';
s.connections(1).mechanism_list='iGABAa';
s.connections(1).parameters={'gGABAa',100};
data=dsSimulate(s,'tspan',[0 400]);
figure; plot(data.time,data.E_v,'b-',data.time,data.I_v,'r-'); 
title('E/I network'); xlabel('time (ms)'); ylabel('v'); legend('E (decay)','I (LIF)'); ylim([-80 -50])

%% Sparse Pyramidal-Interneuron-Network-Gamma (sPING)

% define equations of cell model (same for E and I populations)
eqns={ 
  'dv/dt=Iapp+@current/Cm+noise*randn(1,N_pop)*sqrt(dt)/dt';
  'monitor v.spikes(20), iGABAa.functions, iAMPA.functions'
};
s=[];
s.populations(1).name='E';
s.populations(1).size=80; % # of cells in population
s.populations(1).equations=eqns;
s.populations(1).mechanism_list={'iNa','iK'};
s.populations(1).parameters={'Iapp',5,'gNa',120,'gK',36,'Cm',1,'noise',4};
s.populations(2).name='I';
s.populations(2).size=20;
s.populations(2).equations=eqns;
s.populations(2).mechanism_list={'iNa','iK'};
s.populations(2).parameters={'Iapp',0,'gNa',120,'gK',36,'Cm',1,'noise',4};
s.connections(1).source='I';
s.connections(1).target='E';
s.connections(1).mechanism_list={'iGABAa'};
s.connections(1).parameters={'tauD',10,'gGABAa',.1,'netcon','ones(N_pre,N_post)'};
s.connections(2).source='E';
s.connections(2).target='I';
s.connections(2).mechanism_list={'iAMPA'};
s.connections(2).parameters={'tauD',2,'gAMPA',.1,'netcon',ones(80,20)};
data=dsSimulate(s);

% Resulting data matrices have dimensions [time x cells].
%   DynaSim data structure:
%     data.labels           : list of state variables and monitors recorded
%     data.(state_variables): state variable data matrix [time x cells]
%     data.(monitors)       : monitor data matrix [time x cells]
%     data.time             : time vector [time x 1]
%     data.simulator_options: simulator options used to generate simulated data
%     data.model            : model used to generate simulated data

figure; 
subplot(2,1,1); % voltage traces
plot(data.time,data.E_v,'b-',data.time,data.I_v,'r-')
title('Sparse Pyramidal-Interneuron-Network-Gamma (sPING)'); ylabel('membrane potential (mV)');
subplot(2,1,2); % rastergram
E_spikes=nan(size(data.E_v_spikes)); E_spikes(data.E_v_spikes==1)=1;
I_spikes=nan(size(data.I_v_spikes)); I_spikes(data.I_v_spikes==1)=1;
plot(data.time,E_spikes+repmat(1:80,[length(data.time) 1]),'bo'); hold on
plot(data.time,I_spikes+repmat(80+(1:20),[length(data.time) 1]),'ro'); axis([0 100 0 100]);
title('rastergram'); xlabel('time (ms)'); ylabel('cell index');

% store model for later use
sPING_model = data.model;

% note: N_pop is a reserved variable for the size of the population in which it appears.

% mechanisms "iAMPA" and "iGABAa" contain a fixed variable "netcon" storing
% a default connectivity matrix (all 1s for all-to-all connectivity). The
% user can specify the connectivity matrix from a source to target by
% setting "netcon" in the .parameters cell array. The dimensions of
% "netcon" should be [N_pre x N_post] where N_pre (reserved variable) is
% the size of the (presynaptic) source population and N_post (reserved
% variable) is the size of the (postsynaptic) target population. "netcon"
% can be set as a numeric matrix or a string that evaluates to a numeric
% matrix of the correct dimensions.

dsEditModelFiles('iAMPA.mech');

% equivalent compact population specification with E->I netcon=0
s=[];
s.pops(1).equations='E:dv[80]/dt=10+@current+4*randn(1,N_pop); {iNa,iK}';
s.pops(2).equations='I:dv[20]/dt= 0+@current+4*randn(1,N_pop); {iNa,iK}';
s.cons(1).source='I';
s.cons(1).target='E';
s.cons(1).mechanism_list={'iGABAa'};
s.cons(1).parameters={'tauD',10,'gGABAa',.1,'netcon','ones(N_pre,N_post)'};
s.cons(2).source='E';
s.cons(2).target='I';
s.cons(2).mechanism_list={'iAMPA'};
s.cons(2).parameters={'tauD',2,'gAMPA',.1,'netcon',0*ones(80,20)};
data=dsSimulate(s);
figure; plot(data.time,data.E_v,'b-',data.time,data.I_v,'r-')
title('sPING with E->I turned off');

% Note: the compact specification demonstrates how all components of the
% population specification (name, size, equations, mechanism_list, parameters)
% can be embedded in the equation string. DynaSim will automatically parse
% the string and split the components into their proper fields.
% Specifically:
% Specify name by starting the string with 'NAME: *' (e.g., 'E: dv/dt=-v').
% Specify size by including [SIZE] after the state variable (e.g., 'dv[5]/dt=-v').
% Specify mechanism_list by including cell array listing mechanism names
% without single quotes (e.g., 'dv/dt=@current; {iNa,iK}').

%% Simulator options:

%   solver options (provided as key/value pairs: 'option1',value1,'option2',value2,...):
%     'solver'      : solver for numerical integration (see dsGetSolveFile)
%                     {'euler','rk2','rk4'} (default: 'rk4')
%     'tspan'       : time limits of simulation [begin,end] (default: [0 100]) [ms]
%                     note: units must be consistent with dt and model equations
%     'dt'          : time step used for DynaSim solvers (default: .01) [ms]
%     'downsample_factor': downsampling applied during simulation (default: 1, no downsampling) 
%                     (only every downsample_factor-time point is stored in memory and/or written to disk)
%     'ic'          : numeric array of initial conditions, one value per state 
%                     variable (default: all zeros). overrides definition in model structure
%     'random_seed' : seed for random number generator (default: 'shuffle', set randomly) (usage: rng(options.random_seed))
%     'mex_flag': whether to compile simulation using coder instead of 
%                     interpreting Matlab {0 or 1} (default: 0)
% 
%   options for running sets of simulations:
%     'vary'        : (default: [], vary nothing): cell matrix specifying model
%                     components to vary across simulations (see NOTE 1 and dsVary2Modifications)
% 
%   options to control saved data:
%     'save_data_flag': whether to save simulated data to disk after completion {0 or 1} (default: 0)
%     'overwrite_flag': whether to overwrite existing data files {0 or 1} (default: 0)
%     'study_dir'     : relative or absolute path to output directory (default: current directory)
%     'prefix'        : string to prepend to all output file names (default: 'study')
%     'disk_flag'     : whether to write to disk during simulation instead of storing in memory {0 or 1} (default: 0)
%                 
%   options for cluster computing:
%     'cluster_flag'  : whether to run simulations on a cluster submitted 
%                     using qsub (see dsCreateBatch) {0 or 1} (default: 0)
%     'sims_per_job'  : number of simulations to run per batch job (default: 1)
%     'memory_limit'  : memory to allocate per batch job (default: '8G')
% 
%   options for parallel computing: (requires Parallel Computing Toolbox)
%     'parfor_flag' : whether to use parfor to run simulations {0 or 1} (default: 0)
%     'num_cores'     : number of cores to specify in the parallel pool
%     *note: parallel computing has been disabled for debugging...
% 
%   other options:
%     'verbose_flag'  : whether to display informative messages/logs (default: 0)
%     'modifications' : how to modify DynaSim specification structure component before simulation (see dsApplyModifications)
%     'experiment'    : function handle of experiment function (see NOTE 2)
%     'optimization'  : function handle of optimization function (see NOTE 2)

% %%%%%%%%%%%%%%%%%%%%%%%%%%%%%%%%%%%%%%%%%%%%%%%%%%%%%%%%%%%%%%%%%%%%%%%%
%% RUNNING SETS OF SIMULATIONS
% %%%%%%%%%%%%%%%%%%%%%%%%%%%%%%%%%%%%%%%%%%%%%%%%%%%%%%%%%%%%%%%%%%%%%%%%

% 'vary' indicates the variable to vary, the values it should take, and the 
% object (population or connection) whose variable should be varied. 

% Syntax 1: vary={{object, variable, value1},{object, variable, value2},...}
%   - this is useful for simulating an arbitrary set of parameter values
% Syntax 2: vary={object, variable, values; ...}
%   - this is useful for varying parameters systematically (described later)

% For instance, to vary parameter 'gNa', taking on values 100 and 120, in 
% population 'E', set vary={'E','gNa',[100 120]} (syntax 1) or 
% vary={{'E','gNa',100},{'E','gNa',120}} (syntax 2). To additionally vary 
% 'gAMPA' in the connection mechanism from 'E' to 'I', set 
% vary={'E','gNa',[100 120];'E->I','gAMPA',[0 1]}.
% Mechanism lists and equations can also be varied. (see dsVary2Modifications 
% for more details and examples).

eqns='dv/dt=@current+10; {iNa,iK}; v(0)=-60';
data=dsSimulate(eqns,'vary',{'pop1','gNa',[50 100 200]});
data=dsSimulate(eqns,'vary',{'','gNa',[50 100 200]}); % since only 1 pop
data
data(1)
[data.pop1_gNa]
[data.(data(1).varied{1})] % generic access to values used

% generic plotting of traces for each value of the varied parameter
param_name=data(1).varied{1};
param_values=[data.(param_name)];
num_values=length(param_values);
figure
for i=1:num_values
  subplot(1,num_values,i)
  plot(data(i).time,data(i).(data(i).labels{1}));
  title(sprintf('%s=%g',param_name,param_values(i)));
end
% plot how mean firing rate varies with parameter
dsPlotFR(data,'bin_size',30,'bin_shift',10); % bin_size and bin_shift in [ms]
% plot how firing rate varies over time for one data set
dsPlotFR(data(2),'bin_size',30,'bin_shift',10); % bin_size and bin_shift in [ms]
% plot waveforms
dsPlot(data,'plot_type','waveform')
% plot power spectrum
dsPlot(data,'plot_type','power')
% plot rastergram
dsPlot(data,'plot_type','rastergram');

% note: dsPlotFR accepts any DynaSim data structure or array of data
% structures and generates different plots depending on properties of the
% data set (e.g., # of populations, # of parameters varied, etc); 
% it always try to generate the most informative plots.

% generic manually calculate and plot firing rate (works with any model)
data=dsSelect(data,'time_limits',[20 80]); % extract times 20-80ms
data=dsCalcFR(data,'bin_size',30,'bin_shift',10); % calculate firing rates for each cell in each data set
FRname=data(1).results{1}; % .results contains a list of fields with results calculated in dsCalcFR
FRmean=cellfun(@mean,{data.(FRname)}); % calculate average firing rates
figure; plot(param_values,FRmean,'-o');
xlabel(param_name); ylabel('mean firing rate [Hz]');

% varying two parameters (Iapp and tauD in sPING)
% ... (introduce varying connection parameters) ...
vary={
  'E'   ,'Iapp',[0 10 20];     % amplitude of tonic input to E-cells
  'I->E','tauD',[5 10 15]   % inhibition decay time constant from I to E
  };
data=dsSimulate(sPING_model,'vary',vary);
% plot firing rates calculated from spike monitor in both populations
dsPlotFR(data,'variable','*_spikes','bin_size',30,'bin_shift',10);
% plot firing rates calculated from voltage state variables in both populations
%dsPlotFR(data,'variable','*_v','bin_size',30,'bin_shift',10);
% plot firing rates calculated from voltage state variables in E population
%dsPlotFR(data,'variable','E_v','bin_size',30,'bin_shift',10);

% Spike Monitor (threshold=10) for different tonic amplitudes and max sodium conductance
eqns='dv/dt=@current+amp; {iNa,iK}; monitor v.spikes(10)';
vary={'','amp',2:2:60;'','gNa',[100 120]};
data=dsSimulate(eqns,'vary',vary);
figure
a=[data.pop1_amp]; amps=unique(a);
b=[data.pop1_gNa]; gNas=unique(b);
spikes=cat(2,data.pop1_v_spikes);
for i=1:length(a), spikes(:,i)=spikes(:,i)+a(i); end
subplot(1,2,1); plot(data(1).time,spikes(:,b==gNas(1)),'k'); title(sprintf('gNa=%g',gNas(1))); axis tight
subplot(1,2,2); plot(data(1).time,spikes(:,b==gNas(2)),'k'); title(sprintf('gNa=%g',gNas(2))); axis tight
xlabel('time (ms)'); ylabel('tonic amplitude [uA/cm2]');
set(gca,'ytick',amps,'yticklabel',amps);

%% saving results
% A set of simulations, analyses, and plots deriving from a common base
% model are collectively called a DynaSim "study". All results for a given
% study are saved in a directory called "study_dir" which contains
% subdirectories "data" (simulated data and results derived from analyzing
% the data), "plots", and "solve" (containing m- and mex-files that were
% used for all simulations of the study).

% How to: set 'save_data_flag'=1 and optionally 'study_dir' = /path/to/outputs

study_dir='study_HH_varyI'; 
  % where results will be saved (relative or absolute path)
  % note: study_dir name cannot contain hyphens, spaces, or special characters
eqns='dv/dt=@current+I; {iNa,iK}';
vary={'','I',[0 10 20]};

[data,studyinfo]=dsSimulate(eqns,'vary',vary,'save_data_flag',1,'study_dir',study_dir,'verbose_flag',1);

%   DynaSim studyinfo structure (only showing select fields, see dsCheckStudyinfo for more details)
%     studyinfo.study_dir
%     studyinfo.base_model (=[]): original model from which a set of simulations was derived
%     studyinfo.base_simulator_options (=[])
%     studyinfo.base_solve_file (='')
%     studyinfo.simulations(k): metadata for each simulation in a set of simulations
%                           .sim_id         : unique identifier in study
%                           .modifications  : modifications made to the base model during this simulation
%                           .data_file      : full filename of eventual output file
%                           .batch_dir (=[]): directory where batch jobs were saved (if cluster_flag=1)
%                           .job_file (=[]) : m-file batch job that runs this simulation (if cluster_flag=1)
%                           .simulator_options: simulator options for this simulation
%                           .solve_file     : full filename of m- or mex-file that numerically integrated the model

studyinfo
studyinfo.study_dir
studyinfo.simulations(1)
studyinfo.simulations(1).data_file
studyinfo.simulations(1).modified_model_file
% DynaSim studyinfo structure is always saved to <study_dir>/studyinfo.mat
% <study_dir>/data: contains all output data files
% <study_dir>/models: contains a model MAT file for each simulation

% loading data saved to disk
% load one data set from data file name
data=dsImport(studyinfo.simulations(2).data_file);
dsPlotFR(data,'bin_size',30,'bin_shift',10);
% equivalent ways to load all data sets associated with studyinfo structure
data=dsImport(studyinfo);
data=dsImport(study_dir);
data=dsImport('study_HH_varyI');
dsPlotFR(data);

% re-running the simulation loads data if it already exists (see log)
[data,studyinfo]=dsSimulate(eqns,'vary',vary,'save_data_flag',1,'study_dir',study_dir,'verbose_flag',1);

%% cluster computing
% How to: set 'cluster_flag' to 1
% Requirement: you must be logged on to a cluster that recognizes 'qsub'

% DynaSim creates m-files called jobs that run dsSimulate for one or
% more simulations. Jobs are saved in ~/batchdirs/<study_dir> and are
% submitted to the cluster queue using the command 'qsub'. Standard out and
% error logs for each job are saved in ~/batchdirs/<study_dir>/pbsout.

% create 3 jobs to run 3 simulations

study_dir='study_HH_varyI_cluster'; 
eqns='dv/dt=@current+I; {iNa,iK}';
vary={'','I',[0 10 20]};

[data,studyinfo]=dsSimulate(eqns,'vary',vary,...
  'study_dir',study_dir,'save_data_flag',1,'cluster_flag',1,'verbose_flag',1);
% note: if on a cluster, jobs will be automatically submitted using "qsub"
studyinfo.simulations(1)
ls(studyinfo.simulations(1).batch_dir)
edit(studyinfo.simulations(1).job_file);
edit(studyinfo.simulations(1).solve_file);

% manually run the simulation batch jobs
if 0
  % *** CAUTION: ONLY DO THIS IF NOT ON A CLUSTER ***
  % reason: jobs created on a cluster (e.g., scc2.bu.edu) will end with 
  % the 'exit' command to end the batch job. jobs created on a local
  % machine will not end with the exit command. So, if you run this on a
  % cluster, it will close Matlab when the job finishes.
  run(studyinfo.simulations(1).job_file);
  data=dsImport(studyinfo) % 1 data set
  run(studyinfo.simulations(2).job_file);
  data=dsImport(studyinfo) % 2 data sets
  run(studyinfo.simulations(3).job_file);
  data=dsImport(studyinfo) % 3 data sets
  dsPlotFR(data);
end

% create 2 jobs to run 6 simulations
% set sims_per_job=3 (i.e., run 3 simulations per batch job; 2 jobs in parallel)

study_dir='study_HH_varyI_cluster2'; 
eqns='dv/dt=@current+I; {iNa,iK}';
vary={'','I',[0:10:50]};

[data,studyinfo]=dsSimulate(eqns,'vary',vary,...
  'study_dir',study_dir,'cluster_flag',1,'sims_per_job',3,'save_data_flag',1,'verbose_flag',1);

if 0 % manually run the simulation jobs (see caution above)
  run(studyinfo.simulations(1).job_file);
  data=dsImport(studyinfo) % 3 data sets
  run(studyinfo.simulations(4).job_file);
  data=dsImport(studyinfo) % 6 data sets
  dsPlotFR(data);
end

%% More examples

% Izhikevich study of neuro-computational properties
% based on: http://www.izhikevich.org/publications/izhikevich.m
eqns={
  'a=.02; b=.2; c=-65; d=6; I=14';
  'dv/dt=.04*v^2+5*v+140-u+I; v(0)=-70';
  'du/dt=a*(b*v-u); u(0)=-20';
  'if(v>=30)(v=c;u=u+d)';
  };
P='pop1'; % name of population
vary={
  {P,'a',.02; P,'b',.2 ; P,'c',-50; P,'d',2;  P,'I',15} % tonic bursting
  {P,'a',.01; P,'b',.2 ; P,'c',-65; P,'d',8;  P,'I',30} % spike frequency adaptation
  {P,'a',.02; P,'b',.2 ; P,'c',-65; P,'d',6;  P,'I',7}  % spike latency
  {P,'a',.03; P,'b',.25; P,'c',-52; P,'d',0;  P,'I',0}  % rebound burst
  {P,'a',1;   P,'b',1.5; P,'c',-60; P,'d',0;  P,'I',-65}% bistability
  {P,'a',.02; P,'b',1  ; P,'c',-55; P,'d',4;  P,'I',1}  % accomodation
  {P,'a',-.02;P,'b',-1 ; P,'c',-60; P,'d',8;  P,'I',80} % inhibition-induced spiking
  {P,'a',-.026;P,'b',-1; P,'c',-45; P,'d',0;  P,'I',70} % inhibition-induced bursting
  };
data=dsSimulate(eqns,'tspan',[0 250],'vary',vary);
dsPlot(data);
    
% %%%%%%%%%%%%%%%%%%%%%%%%%%%%%%%%%%%%%%%%%%%%%%%%%%%%%%%%%%%%%%%%%%%%%%%%
%% Special issues and advanced concepts
% %%%%%%%%%%%%%%%%%%%%%%%%%%%%%%%%%%%%%%%%%%%%%%%%%%%%%%%%%%%%%%%%%%%%%%%%

%% Linking variables across mechanisms

% Example: HH-type neuron with calcium-dependent potassium channels

% Sometimes mechanisms in the same population may depend on each other. For
% example, a calcium buffer depends on all calcium currents in the same
% population, and all calcium-dependent mechanisms depend on the calcium
% concentration defined in the Ca2+ buffer mechanism. In these cases, state
% variables and functions can be linked between mechanisms in the same way 
% they are linked from mechanism to population equations. 

data=dsSimulate('dv/dt=@current; {iNa,iK,iCa,iCan,CaBuffer}');
figure; plot(data.time,data.(data.labels{1}))
% linkers and linked ODEs:
% CaBuffer.mech: @cai += cai
%                dcai/dt = a*(-@ica)-(cai-c0)/tau
% iCan.mech:     @ica += I(v,m)
%                dm/dt = (minf(@cai)-m)./mtau(@cai)
% iCa.mech:      @ica += I(v)
% note: always list the link used in the population equations first then
% links used across mechanisms.
eqns={
  'dv/dt=@current; {iNa,iK,iCa,iCan,CaBuffer}';
  'monitor o=sqrt(@cai), ica=@ica, iCa.I, iCan.I';
  };
data=dsSimulate(eqns)
data.model.monitors
% plot iCa.I, iCan.I, @ica (= sum of the other two)
  
  
%% Mechanisms that call custom Matlab functions (for advanced programming)

    % trivial: model equations can call any matlab function, including custom ones
    % tips: use custom functions to obtain complicated inputs, noise
    % sources, or connectivity matrices (alternatively: define them in
    % script and pass them as parameters using the DynaSim specification).
    
%% Modularization of mechanisms (@, X, X_pre, X_post)

    % the mechanism linker target IDENTIFIER used to link mechanism variables and
    % functions to population equations can be overriden by including
    % @NEWIDENTIFIER in the equations and after the mechanism name. (e.g.,
    % 'dv/dt=@M; {iNa,iK}@M'; or .mechanism_list={'iNa@M','iK@M'}).

    eqns='dv/dt=@M+I; {iNa,iK}@M; monitor functions';
    data=dsSimulate(eqns,'vary',{'','I',[0 10 20]});
    dsPlotFR(data);

    eqns='dv/dt=@M+10; {iNa,iK}@M; monitor functions';
    data=dsSimulate(eqns,'vary',{'','gNa',[50 100 200]});
    dsPlotFR(data);

%% Special functions: Experiment [and Optimization]

    data=dsSimulate('dv/dt=@current+10; {iNa,iK}','experiment',@dsProbeFI);
    dsPlotFR(data);

%% Modifications and Vary

    % "modifications" are applied to the specification, and indicate how to
    % change the name, size, equations, mechanism_list, and/or parameters of a
    % population, or mechanism_list and/or parameters of a connection.

    % dsApplyModifications() returns the modified specification or 
    % regenerated model after modifying the specification
    
    % "vary" is a way of specifying sets of modifications
    
    % dsVary2Modifications() returns a cell array of modifications specified 
    % by the vary statement
    
    % dsSimulate() supports both "modifications" and "vary" options; if
    % the latter is provided, a set of simulations are performed and a set
    % of simulated data sets are returned and/or saved.
    
    eqns='dv/dt=@current+I; {iNa,iK}';
    modifications={'','I',10};
    data=dsSimulate(eqns,'modifications',modifications);

    eqns='dv/dt=@current+I; {iNa,iK}';
    vary={'','I',[0 10 20]};
    data=dsSimulate(eqns,'vary',vary);
    
    
    vary={'pop1','gNa',[50 100 200]};
    modifications_set=dsVary2Modifications(vary); 
    % {{'pop1','gNa',50},{'pop1','gNa',100},{'pop1','gNa',200}}
    clear data; figure
    for i=1:length(modifications_set)
      modifications=modifications_set{i};
      data(i)=dsSimulate('dv/dt=@M+10; {iNa,iK}@M','modifications',modifications);
      subplot(1,length(modifications_set),i); plot(data(i).time,data(i).(data(i).labels{1}))
      title(sprintf('gNa=%g',modifications{3}));
    end

    % auto-constructed search space given special case specification of what to vary
    data=dsSimulate('dv/dt=@M+10; {iNa,iK}@M','vary',{'','gNa',[50 100 200]});
    data=dsSimulate('dv/dt=@M+10; {iNa,iK}@M; vary(gNa=[50 100 200])');
      % note: special syntax "vary(...)" only works for varying 1 parameter in a 1-population model
    % eqivalent manual construction of search space
    vary={{'pop1','gNa',50},{'pop1','gNa',100},{'pop1','gNa',200}};
    data=dsSimulate('dv/dt=@M+10; {iNa,iK}@M','vary',vary);

    % more examples of 'vary'
    vary={'E','gNa',[100 120]};
    vary={'E','gNa',[100 120];'E->I','gAMPA',[0 1]};
    vary={'E','mechanism_list','+[iNa,iK]'};
    vary={'E','mechanism_list','-{iNa,iK}'};
    vary={'(E,I)','gNa',[100 120]};
    vary={'(E,I)','(EK1,EK2)',[-80 -60]};
    
    % more examples of single modifications:
    % modifying mechanism_list
    s=dsApplyModifications('dv/dt=10+current; {iNa,iK}; v(0)=-65',...
                         {'pop1','mechanism_list','-iNa'});
    s.populations.mechanism_list
    s=dsApplyModifications('dv/dt=10+current; {iNa,iK}; v(0)=-65',...
                         {'pop1','mechanism_list','+iCa'});
    s.populations.mechanism_list
    s=dsApplyModifications('dv/dt=10+current; {iNa,iK}; v(0)=-65',...
                         {'pop1','mechanism_list','+(iCa,iCan,CaBuffer)'});
    s.populations.mechanism_list
    
%% other

% plotting state variables returned from custom matlab functions
% 1. define custom function (saved to 'get_input.m' in Matlab path)
%   example: function input=get_input(type,N,T,f)
% 2. use function in model
%   eqns='dv/dt=-v+I(k,:); I=get_input(''rectified_sin'',Npop,T,f); f=5';
%     % note: 'k' is an internal index to the current time step during
%     % simulation; 'T' is an internal variable storing the full time array;
%     % 'Npop' is an internal variable storing the size of the population.
%   data=dsSimulate(eqns,'tspan',[0 1000]);
% 3. plot the state variable stored in the post-simulation model structure
%   figure; plot(data.time,data.model.fixed_variables.pop1_I)

  
```
