## Supplemental File 1 for "A biophysical model of striatal microcircuits suggests delta/theta-rhythmically interleaved gamma and beta oscillations mediate periodicity in motor control": dsHarmonicOscillator.m

```
% Exploring natural and resonant frequencies of harmonic oscillators.
% Samped, driven harmonic oscillators
% ref: https://en.wikipedia.org/wiki/Harmonic_oscillator

% parameters
f0 = 10/1e3;  % undamped "natural" frequency
m = 1;        % mass
l = 0;        % damping ratio
w0 = 2*pi*f0; % rad/ms, undamped angular frequency
F0 = w0^2;    % amplitude of driving force

% step input
eqns_harmonic_step={
  'du/dt=v; u(0)=.5'
  'dv/dt=-w0^2*u-2*l*w0*v+(1/m)*F0'
  };

% sinusoidal driving force
eqns_harmonic_sine={
  'du/dt=v; u(0)=.5'
  'dv/dt=-w0^2*u-2*l*w0*v+(1/m)*F0*sin(2*pi*f*t)'
  };

% step response simulations
vary={'m',m;'l',l;'w0',w0;'F0',F0};
data=dsSimulate(eqns_harmonic_step,'vary',vary,'time_limits',[0 1000]);
dsPlot(data);

% vary input strength
vary={'m',m;'l',l;'w0',w0;'F0',F0*[1 2 3]};
data=dsSimulate(eqns_harmonic_step,'vary',vary,'time_limits',[0 1000]);
dsPlot(data);

% frequency response simulations
vary={'m',m;'l',l;'w0',w0;'F0',F0;'f',[0:2:20]/1e3};
data=dsSimulate(eqns_harmonic_sine,'vary',vary,'time_limits',[0 1000]);
dsPlot(data);

% vary input strength
vary={'m',m;'l',l;'w0',w0;'F0',F0*[1 2 3];'f',[0:2:20]/1e3};
data=dsSimulate(eqns_harmonic_sine,'vary',vary,'time_limits',[0 1000]);
dsPlot(data);

% observation 1:
% - resonant frequency = undamped frequency given equal strength step input
% TRUE for all second-order linear oscillatory systems.

% observation 2:
% - resonant frequency is independent of input strength
% - response amplitude increases with input strength
```
