## Supplemental File 1 for "A biophysical model of striatal microcircuits suggests delta/theta-rhythmically interleaved gamma and beta oscillations mediate periodicity in motor control": dsLIFnetwork.m

```
% 1 E-cell driving 1 I-cell
LIF={
    'dV/dt=(E-V+R*I+noise*randn-@isyn)/tau; V(0)=-65'
    'if(any(t<tspike+tabs,1))(V=reset)'
    'tau=10; tabs=10; E=-70; thresh=-55; reset=-75; R=9; I=1.55; noise=100'
    'monitor V.spikes(thresh)'
     };

iampa={
  'gSYN=.5; ESYN=0; tauD=2; tauR=0.4; delay=15'
  'f(x) = (exp(-x/tauD)-exp(-x/tauR)).*(x>0)'
  'Isyn(V) = gSYN.*sum(f(t-tspike_pre-delay)).*(V-ESYN)'
  '@isyn += Isyn(V_post)'
};

s=[];
s.pops(1).name='E';
s.pops(1).equations=LIF;
s.pops(2).name='I';
s.pops(2).equations=LIF;
s.pops(2).parameters={'I',0,'noise',0};
s.cons(1).direction='E->I';
s.cons(1).mechanism_list='iampa';
s.mechanisms(1).name='iampa';
s.mechanisms(1).equations=iampa;

data=dsSimulate(s,'time_limits',[0 200],'solver','rk1','dt',.01);
dsPlot(data);

%% Interacting E and I populations
% tau [ms]: membrane time constant (RC)
% tabs [ms]: absolute refractory period
% delay [ms]: axonal delay
% tspike: reserved variable for storing past spike times when monitoring spikes

LIF={
    'dV/dt=(E-V+mask.*(R*I+noise*randn(1,N_pop))+@isyn)/tau; V(0)=-65'
    'if(any(t<tspike+tabs,1))(V=reset)'
    'tau=10; tabs=10; E=-70; thresh=-55; reset=-75; R=9; I=1.55; noise=100; mask=ones(1,N_pop)'
    'monitor V.spikes(thresh)'
     };
 
iampa={
  'gSYN=.5; ESYN=0; tauD=2; tauR=0.4; delay=15'
  'netcon=ones(N_pre,N_post)'
  'f(x) = 1*(exp(-x/tauD)-exp(-x/tauR)).*(x>0)'
  'Isyn(V_post) = gSYN.*(sum(f(t-tspike_pre-delay))*netcon).*(V_post-ESYN)'
  '@isyn += -Isyn(V_post)'
  'monitor Isyn'
};

% input mask
Ne=80; Ni=20;
maskE=zeros(1,Ne);
maskE(round(Ne/2))=1; % drive only a single E cell

% gaussian connectivity matrix (for spreading E activation across the layer)
sigma=.05; % 5% of the E layer
Npre=Ne; Npost=Ne; Nmax=max(Npost,Npre);
srcpos=linspace(1,Nmax,Npre)'*ones(1,Npost);
dstpos=(linspace(1,Nmax,Npost)'*ones(1,Npre))';
netconEE=exp(-(srcpos-dstpos).^2/(sigma*Npost)^2)'-eye(Npost);

% dynasim specification
s=[];
s.pops(1).name='E';
s.pops(1).size=Ne;
s.pops(1).equations=LIF;
s.pops(1).parameters={'mask',maskE};
s.pops(2).name='I';
s.pops(2).size=Ni;
s.pops(2).equations=LIF;
s.pops(2).parameters={'I',0,'noise',0};
s.cons(1).direction='E->I';
s.cons(1).mechanism_list='iampa';
s.cons(2).direction='E->E';
s.cons(2).mechanism_list='iampa';
s.cons(2).parameters={'netcon',netconEE,'delay',50,'gSYN',3};
s.mechanisms(1).name='iampa';
s.mechanisms(1).equations=iampa;

% simulation
data=dsSimulate(s,'time_limits',[0 1000],'solver','rk1','dt',.01);
figure
subplot(2,1,1); imagesc(data.time,1:Ne,data.E_V'); ylabel('E index'); title('membrane potential');
subplot(2,1,2); imagesc(data.time,1:Ni,data.I_V'); ylabel('I index'); xlabel('time (ms)');
%dsPlot(data);

%% 1 E-cell driving another with E->E spike-timing dependent plasticity (STDP)
% tau [ms]: membrane time constant (RC)
% tabs [ms]: absolute refractory period
% delay [ms]: axonal delay
% tspike: reserved variable for storing past spike times when monitoring spikes
% STDP: http://www.scholarpedia.org/article/Spike-timing_dependent_plasticity, see online implementation

LIF={
    'dV/dt=(E-V+mask.*(R*I*(t>onset)+noise*randn(1,N_pop))+@isyn)/tau; V(0)=-65'
    'if(any(t<tspike+tabs,1))(V=reset)'
    'tau=10; tabs=10; E=-70; thresh=-55; reset=-75; R=9; I=1.55; noise=0; onset=0; mask=ones(1,N_pop)'
    'monitor V.spikes(thresh)'
     };

AMPAstdp={
  % STDP
  'wmax=1; rx=1; ry=1'
  'ax(x)=1-x' % incremental increase in presynaptic trace with nearest-neighbor spike-interaction
  'ay(y)=1-y' % incremental increase in postsynaptic trace
  'Ax(w)=(wmax-w)*rx' % soft bounds
  'Ay(w)=w*ry'        % soft bounds
  'dx/dt=-x+ax(x).*sum((t-tspike_pre-delay)<dt)'
  'dy/dt=-y+ay(y).*sum((t-tspike_post-delay)<dt)'
  'dw/dt=Ax(w).*x.*sum((t-tspike_post-delay)<dt) - Ay(w).*y.*sum((t-tspike_pre-delay)<dt)'
  'x(0)=.1; y(0)=.1; w(0)=.5'
  % synaptic current
  'gSYN=1; ESYN=0; tauD=2; tauR=0.4; delay=5'
  'netcon=ones(N_pre,N_post)'
  'f(x) = 1*(exp(-x/tauD)-exp(-x/tauR)).*(x>0)'
  'Isyn(X) = w.*gSYN.*(sum(f(t-tspike_pre-delay))*netcon).*(X-ESYN)'
  '@isyn += -Isyn(X_post)'
  'monitor Isyn'
};
AMPA={
  % synaptic current
  'gSYN=1; ESYN=0; tauD=2; tauR=0.4; delay=5; w=1'
  'netcon=ones(N_pre,N_post)'
  'f(x) = 1*(exp(-x/tauD)-exp(-x/tauR)).*(x>0)'
  'Isyn(X) = w.*gSYN.*(sum(f(t-tspike_pre-delay))*netcon).*(X-ESYN)'
  '@isyn += -Isyn(X_post)'
  'monitor Isyn'
};

% dynasim specification
s=[];
s.pops(1).name='e1';
s.pops(1).size=1;
s.pops(1).equations=LIF;
s.pops(1).parameters={'I',2,'noise',0,'onset',0};
s.pops(2).name='e2';
s.pops(2).size=1;
s.pops(2).equations=LIF;
s.pops(2).parameters={'I',0,'noise',0,'onset',140};
s.cons(1).direction='e1->e2';
s.cons(1).mechanism_list='AMPAstdp';
s.cons(1).parameters={'delay',5,'gSYN',3};
s.mechanisms(1).name='AMPAstdp';
s.mechanisms(1).equations=AMPAstdp;

% simulation
tspan=[0 10000];
vary={'e1','I',[2 5 10]};
vary={'e1->e2','delay',[1 5 10]; 'e1','I',2};
vary={'e1->e2','delay',[1]; 'e2','onset',[40 50 60]; 'e2','I',[.75 1 1.25]};
vary={'e1->e2','delay',1;'(e1,e2)','noise',200*[1 2 3]; '(e1,e2)','onset',[50]; '(e1,e2)','I',.5};
data=dsSimulate(s,'time_limits',tspan,'solver','rk1','dt',.01,'verbose_flag',1,'vary',vary);
dsPlot(data)
dsPlot(data,'plot_type','rastergram');
figure; plot(data(1).time,[data.e2_e1_AMPAstdp_w])

%% Interacting E and I populations with E->E spike-timing dependent plasticity (STDP)
% tau [ms]: membrane time constant (RC)
% tabs [ms]: absolute refractory period
% delay [ms]: axonal delay
% tspike: reserved variable for storing past spike times when monitoring spikes
% STDP: http://www.scholarpedia.org/article/Spike-timing_dependent_plasticity, see online implementation

LIF={
    'dV/dt=(E-V+mask.*(R*I+noise*randn(1,N_pop))+@isyn)/tau; V(0)=-65'
    'if(any(t<tspike+tabs,1))(V=reset)'
    'tau=10; tabs=10; E=-70; thresh=-55; reset=-75; R=9; I=1.55; noise=100; mask=ones(1,N_pop)'
    'monitor V.spikes(thresh)'
     };
 
AMPAee={
  'gSYN=.5; ESYN=0; tauD=2; tauR=0.4; delay=15; tauSTDP=.2'
  'netcon=ones(N_pre,N_post)'
  'f(x) = 1*(exp(-x/tauD)-exp(-x/tauR)).*(x>0)'
  'Isyn(X) = w.*gSYN.*(sum(f(t-tspike_pre-delay))*netcon).*(X-ESYN)'
  '@isyn += -Isyn(X_post)'
  'monitor Isyn'
  % STDP
  'g(x) = 1*(exp(-x/tauSTDP)-exp(-x/tauSTDP)).*(x>0)'
  'ax(x)=x; ay(y)=y; Ax(w)=w; Ay(w)=w'
  'dx/dt=-x+ax(x).*sum(g(t-tspike_pre-delay))'
  'dy/dt=-y+ay(y).*sum(g(t-tspike_post-delay))'
  'dw/dt=Ax(w).*x.*sum(g(t-tspike_post-delay)) - Ay(w).*y.*sum(g(t-tspike_pre-delay))'
  'x(0)=.1; y(0)=.1; w(0)=1'
};
AMPAei={
  'gSYN=.5; ESYN=0; tauD=2; tauR=0.4; delay=15'
  'netcon=ones(N_pre,N_post)'
  'f(x) = 1*(exp(-x/tauD)-exp(-x/tauR)).*(x>0)'
  'Isyn(X) = gSYN.*(sum(f(t-tspike_pre-delay))*netcon).*(X-ESYN)'
  '@isyn += -Isyn(X_post)'
  'monitor Isyn'
};

% input mask
Ne=20; Ni=5;
maskE=zeros(1,Ne);
maskE(round(Ne/2))=1; % drive only a single E cell

% gaussian connectivity matrix (for spreading E activation across the layer)
sigma=.1; % 10% of the E layer
Npre=Ne; Npost=Ne; Nmax=max(Npost,Npre);
srcpos=linspace(1,Nmax,Npre)'*ones(1,Npost);
dstpos=(linspace(1,Nmax,Npost)'*ones(1,Npre))';
netconEE=exp(-(srcpos-dstpos).^2/(sigma*Npost)^2)'-eye(Npost);

% dynasim specification
s=[];
s.pops(1).name='E';
s.pops(1).size=Ne;
s.pops(1).equations=LIF;
s.pops(1).parameters={'mask',maskE};
s.pops(2).name='I';
s.pops(2).size=Ni;
s.pops(2).equations=LIF;
s.pops(2).parameters={'I',0,'noise',0};
s.cons(1).direction='E->I';
s.cons(1).mechanism_list='AMPAei';
s.cons(2).direction='E->E';
s.cons(2).mechanism_list='AMPAee';
s.cons(2).parameters={'netcon',netconEE,'delay',50,'gSYN',3};
s.mechanisms(1).name='AMPAee';
s.mechanisms(1).equations=AMPAee;
s.mechanisms(2).name='AMPAei';
s.mechanisms(2).equations=AMPAei;

% simulation
data=dsSimulate(s,'time_limits',[0 500],'solver','rk1','dt',.01);
figure; subplot(3,1,1); imagesc(data.E_V'); subplot(3,1,2); imagesc(data.I_V'); subplot(3,1,3); imagesc(data.E_E_AMPAee_w); title('w');
dsPlot(data);
```
