## Supplemental File 1 for "A biophysical model of striatal microcircuits suggests delta/theta-rhythmically interleaved gamma and beta oscillations mediate periodicity in motor control": dsReactionDiffusion2D.m

```
%% Simple example for teaching about implementing 2D pops in DynaSim
% based on https://blogs.mathworks.com/graphics/2015/03/16/how-the-tiger-got-its-stripes/
s=[];
s.pops.size=[300 300];
s.pops.equations={'dA/dt=    del2(A)-A.*B.^2+.022*(1-A); A(0)=A_IC'
                  'dB/dt=0.5*del2(B)+A.*B.^2-.073*B;     B(0)=B_IC'
                  'A_IC=ones(Npop)'
                  'B_IC=zeros(Npop); B_IC(150,125:175)=1'};
data=dsSimulate(s,'dt',.25,'tspan',[0 4000],'solver','rk1','verbose_flag',1);
dsPlot2D(data);

% Recommended DynaSim implementation
% based on https://blogs.mathworks.com/graphics/2015/03/16/how-the-tiger-got-its-stripes/
f = 0.022;
k = 0.051;

% Diffusion rates
da = 1;
db = .5;
% Size of grid
width = 300;
% 5,000 simulation seconds with 4 steps per simulated second
dt = .25;
stoptime = 2000;

%Initial conditions
A = ones(width);  % Initialize A to one
B = zeros(width); % Initialize B to zero which a clump of ones
B(width/2,width/2-25:width/2+25) = 1;

s=[];
s.pops.size=[width width];
s.pops.equations={'dA/dt=da*del2(A)-A.*B.^2+f*(1-A); A(0)=A_IC; A_IC=zeros(Npop)';
                  'dB/dt=db*del2(B)+A.*B.^2-(k+f)*B; B(0)=B_IC; B_IC=zeros(Npop)'};
s.pops.parameters={'da',da,'f',f,'db',db,'k',k,'A_IC',A,'B_IC',B};

data=dsSimulate(s,'dt',dt,'tspan',[0 stoptime],'solver','rk1','verbose_flag',1);
figure; imagesc(squeeze(data.pop1_B(end,:,:)))
dsPlot2D(data);

%% Most compact implementation possible in DynaSim
clear

eqns={'dA[300,300]/dt=    del2(A)-A.*B.^2+.022*(1-A); A(0)=ones(Npop)'
      'dB[300,300]/dt=0.5*del2(B)+A.*B.^2-.073*B;     B(0)=B_IC'
      'B_IC=zeros(Npop); B_IC(150,125:175)=1'};
data=dsSimulate(eqns,'dt',.25,'tspan',[0 4000],'solver','rk1');
dsPlot2D(data);
```
