## Supplemental File 1 for "A biophysical model of striatal microcircuits suggests delta/theta-rhythmically interleaved gamma and beta oscillations mediate periodicity in motor control": dsShuntingNetwork2D.m

```
% 2D competitive network with excitatory and inhibitory rate-based populations
% The 2D grid is vectorized into a 1D linear array; it is displayed in 2D using dsPlot2D.
% See dsReactionDiffusion2D.m for a 2D grid modeled using matrices.

aa = 40;        % Side length of square grid
N = aa*aa;      % Number of neurons
tlen = 10000;
We = (eye(N));
Wi = We;
h = 0.001;
B = 10; C = 10;
Ae = 0.6;      % Excitatory kernal max height
ke = 1;        % Excitatory kernel width
Ai = 0.02;   % Inhibitory kernal max height (Play with this parameter!)
ki = 4;        % Inhibitory kernal width
ra = 0.01;
zinh = 10;
toff = 0.4;
distan = zeros(N);
kk = 0;
pos = zeros(N,2);
for ii = 1:aa
  for jj = 1:aa
    kk = kk +1;
    pos(kk,:) = [ii jj];
  end
end
for ii = 1:N
  for jj = ii:N
    distan(ii,jj) = norm(squeeze(pos(ii,:)) - squeeze(pos(jj,:)),2);
    distan(jj,ii) = distan(ii,jj);
    Wi(ii,jj) = Ai.*exp(-(distan(ii,jj)./ki).^2);%./(ki*sqrt(2*pi));
    We(ii,jj) = Ae.*exp(-(distan(ii,jj)./ke).^2);% - Ae.*exp(-(distan(ii,jj)./(0.85*ke)).^2);%./(ki*sqrt(2*pi));
    We(jj,ii) = We(ii,jj);
    Wi(jj,ii) = Wi(ii,jj);
    Wi(ii,ii) = 0;        
  end
end

Inp = 0.1*round(max(rand(tlen,N) - 0.499,0));
Inp(0.1*tlen:tlen,:) = 0;
Inp((tlen*toff):tlen,:)=0;

eqns={'dx/dt=(B-x).*(Inp(k,:)+inh(x)*We)-(x+.2*C).*(2*(z*Wi))-x*ra; x(0)=zeros(1,Npop)';
      'dz/dt=(B-z).*inh(x)-zinh*z.*(1-sign(inh(x)));                z(0)=zeros(1,Npop)';
      'inh(x)=max(x,0); B=0; Inp=0; We=0; Wi=0; ra=0; zinh=0'};
s=[];
s.pops.size=N;
s.pops.equations=eqns;
s.pops.parameters={'B',B,'Inp',Inp,'We',We,'C',C,'Wi',Wi,'ra',ra,'zinh',zinh};
data=dsSimulate(s,'dt',h,'tspan',[0 tlen-1]*h,'solver','rk1','verbose_flag',1);
dsPlot2D(data,'input','pop1_Inp');
```
