## Supplemental File 1 for "A biophysical model of striatal microcircuits suggests delta/theta-rhythmically interleaved gamma and beta oscillations mediate periodicity in motor control": Multicompartment_PFC_neurons.m

```
% This script provides explicit DynaSim specifications for each PFC cell 
% type used in JSS PFC network models. Script created by JSS on 11-Apr-2016.

% Cell models:
% 1) [DS02PY]: two-compartment pyramidal cell (PYs,PYd) (from [DS02])
% 2) [DS02FS]: FS interneuron (from [DS02])
% 3) [DS02KS14FS]: FS interneuron (IN from [DS02] w/ FS modifications from [KS14] soma)
% 4) [DS02KS14RSNP]: RSNP interneuron (IN from [DS02] w/ RS modifications from [KS14] soma)
% 5) [DS02PYjs]: Custom two-compartment pyramidal cell (PYs,PYd) ([DS02] + [TW03] dend ih)
% 6) [DS02FSjs]: Custom FS interneuron (IN from [DS02] w/ JSS modifications based on [KS14] FS)
% 7) [DS02RSNPjs]: Custom RSNP interneuron (IN from [DS02] w/ JSS modifications based on [KS14] RS and experiments)

% References:
% [DS00] Durstewitz, D., Seamans, J. K., & Sejnowski, T. J. (2000). Dopamine-mediated stabilization of delay-period activity in a network model of prefrontal cortex. Journal of neurophysiology, 83(3), 1733-1750.
% [DS02] Durstewitz, D., & Seamans, J. K. (2002). The computational role of dopamine D1 receptors in working memory. Neural Networks, 15(4), 561-572.
% [DG07] Durstewitz, D., & Gabriel, T. (2007). Dynamical basis of irregular spiking in NMDA-driven prefrontal cortex neurons. Cerebral cortex, 17(4), 894-908.
% [KS14] Konstantoudaki, X., Papoutsi, A., Chalkiadaki, K., Poirazi, P., & Sidiropoulou, K. (2014). Modulatory effects of inhibition on persistent activity in a cortical microcircuit model. Frontiers in neural circuits, 8, 1-15.
% [TW03] Traub, R. D., Buhl, E. H., Gloveli, T., & Whittington, M. A. (2003). Fast rhythmic bursting can be induced in layer 2/3 cortical neurons by enhancing persistent Na+ conductance or by blocking BK channels. Journal of neurophysiology, 89(2), 909-921.
% Code:
% NEURON source for [DS02] PY model: ftp://ftp.cnl.salk.edu/pub/dd/pcell
% NEURON (PY,FS) and Matlab (PY only) code for [DG07] model available on ModelDB.
% NEURON (PY,FS,RSNP) code for [KS14] model available on ModelDB.

%% %%%%%%%%%%%%%%%%%%%%%%%%%%%%%%%%%%%%%%%%%%%%%%%%%%%%%%%%%%%%%%%%%%%%%%%%
% -------------------------------------------------------------------------
% 1) [DS02PY] Pyramidal cell: two compartments (PYs soma, PYd dendrite) [DS02]
% -------------------------------------------------------------------------
% Note: This implementation is an exact match to the [DS02] PY model.

% # pyramidal cells
Ne=1;
% cell morphology (cylindrical compartments)
% pyramidal soma (length and diameter chosen to match surface area and internal resistance with a sphere of diam 23 microns)
ls=28.618;  % um, length
ds=21.840;  % um, diameter
% pyramidal dendrite
ld=650;     % um, length
dd=6.5;     % um, diameter

% shared parameters
epas=-70;     % mV, passive leak reversal potential
ki=140;       % mM, intracellular potassium concentration
koinf=3.82;   % mM, steady-state extracellular potassium concentration
KAF=2e6;      % potassium accumulation factor
tauK=7;       % ms, extracellular potassium decay time constant
dshellK=70e-3;% um, depth of extracellular shell for K+ diffusion
cao=2e3;      % uM, extracellular calcium concentration
cainf=50e-3;  % uM, steady-state intracellular calcium concentration
dshellCa=2e-4;% um, depth of intracellular shell for Ca2+ diffusion
faraday=96487;% s*A/mol, faraday constant (charge per mole of ions)

% ionic mechanisms and voltage dynamics present in both compartments (see [DS00] Methods for justification)
mechanism_list={'DS00iNa','DS00iNaP','DS00iDR','DS02iKS','DS02iHVA','DS00iKCa','DS00CaDyn','DS00KDyn','pas'};
state_equations='dV/dt=(@current+Iapp)./Cm; Iapp=0; Cm=1; V(0)=-65';
spec=[];

% soma
l=ls; d=ds; A=d*l*pi; % um2, cylinder surface area without the ends
Cm=1.2e-5;            % 1.2 uF/cm2 = 1.2e-5 nF/um2
gpas=1/30e5;          % Rm=30kOhm-cm2 = 30e5 MOhm-um2, gpas=1/Rm [uS/um2]
gnaf=117e-5;          % 117 mS/cm2 = 117e-5 uS/um2
gkdr=50e-5;           % 50 mS/cm2 = 50e-5 uS/um2
gnap=1.8e-5;          % uS/um2, persistent sodium channel
gks=.08e-5;           % uS/um2, slow potassium channel
ghva=.4e-5;           % uS/um2, high-voltage-activated Ca2+ channel
gkc=2.1e-5;           % uS/um2, V- and Ca2+-dependent potassium channel
tauCa=250;            % ms, calcium decay time constant
CAF=600;              % calcium accumulation factor
VshellK=pi*dshellK*l.*(d+dshellK);    % um3, volume of extracellular shell for K+ diffusion/accumulation
VshellCa=pi*dshellCa*l.*(d-dshellCa); % um3, volume of intracellular shell for Ca2+ diffusion/accumulation
spec.populations(1).name='Es';
spec.populations(1).size=Ne;
spec.populations(1).equations=state_equations;
spec.populations(1).mechanism_list=mechanism_list;
spec.populations(1).parameters={'Iapp',0,'Cm',Cm*A,'gpas',gpas*A,'epas',epas,...
  'gnaf',gnaf*A,'gnap',gnap*A,'ghva',ghva*A,'gkdr',gkdr*A,'gks',gks*A,'gkc',gkc*A,...
  'CAF',CAF,'VshellCa',VshellCa,'cainf',cainf,'tauCa',tauCa,'cao',cao,...
  'KAF',KAF,'VshellK',VshellK,'koinf',koinf,'tauK',tauK,'ki',ki,'faraday',faraday};

% dendrite
l=ld; d=dd; A=d*l*pi; % um2, cylinder surface area without the ends
Cm=Cm*1.92;           % 2.304 uF/cm2
gpas=gpas*1.92;       % 1/(Rm/1.92)
gnaf=20e-5;           % 20 mS/cm2 = 20e-5 uS/um2
gkdr=14e-5;           % 14 mS/cm2 = 14e-5 uS/um2
gnap=.8e-5;           % uS/um2, persistent sodium channel
gks=.08e-5;           % uS/um2, slow potassium channel
ghva=.8e-5;           % uS/um2, high-voltage-activated Ca2+ channel
gkc=2.1e-5;           % uS/um2, V- and Ca2+-dependent potassium channel
tauCa=120;            % ms, calcium decay time constant
CAF=600;              % calcium accumulation factor
VshellK=pi*dshellK*l.*(d+dshellK);    % um3, volume of extracellular shell for K+ diffusion/accumulation
VshellCa=pi*dshellCa*l.*(d-dshellCa); % um3, volume of intracellular shell for Ca2+ diffusion/accumulation
spec.populations(2).name='Ed';
spec.populations(2).size=Ne;
spec.populations(2).equations=state_equations;
spec.populations(2).mechanism_list=mechanism_list;
spec.populations(2).parameters={'Iapp',0,'Cm',Cm*A,'gpas',gpas*A,'epas',epas,...
  'gnaf',gnaf*A,'gnap',gnap*A,'ghva',ghva*A,'gkdr',gkdr*A,'gks',gks*A,'gkc',gkc*A,...
  'CAF',CAF,'VshellCa',VshellCa,'cainf',cainf,'tauCa',tauCa,'cao',cao,...
  'KAF',KAF,'VshellK',VshellK,'koinf',koinf,'tauK',tauK,'ki',ki,'faraday',faraday};

% intercompartmental connections
Ri=1.5; % axial resistance [MOhm-um], Ri=150 Ohm-cm
% collect relevant info
compartments={'Es' 'Ed'};
lengths     =[ls   ld];
diameters   =[ds   dd];
connections={[1 2],[2 1]};
% add connections to specification
for c=1:length(connections)
  src=connections{c}(1);
  dst=connections{c}(2);
  spec.connections(c).direction=[compartments{src} '->' compartments{dst}];
  spec.connections(c).mechanism_list={'iCOM'};
  gCOM=1/mean(Ri*4*lengths./(pi*diameters.^2));
  spec.connections(c).parameters={'gCOM',gCOM};
end

% equivalent gCOM calculation for coupling from 2 to 1:
% g12=@(r1,L1,r2,L2)(r1*r2^2)/(Ri*L1*(L1*r2^2+L2*r1^2));
% gCOM=g12*(surface area)=g12*(2*pi*r1*L1)
% d->s: g12(ds/2,ls,dd/2,ld)*(pi*ls*ds)
% s->d: g12(dd/2,ld,ds/2,ls)*(pi*ld*dd)
% Units: [Ri]=MOhm*um, [r]=[L]=um, then [g12]=uS/um2 and [gCOM]=uS. Ra=1/gCOM
% Reference: https://en.wikipedia.org/wiki/Compartmental_modelling_of_dendrites

data=dsSimulate(spec,'tspan',[0 2000],'vary',{'Es','Iapp',.1},'solver','rk1','verbose_flag',1);
dsPlot(data,'ylim',[-100 50]);

%% %%%%%%%%%%%%%%%%%%%%%%%%%%%%%%%%%%%%%%%%%%%%%%%%%%%%%%%%%%%%%%%%%%%%%%%%
% -------------------------------------------------------------------------
% 2) [DS02FS] Fast-spiking GABAergic interneuron [DS02]
% -------------------------------------------------------------------------
% Note: This implementation is an exact match to the [DS02] FS model.

% # FS cells
Ni=1;  
% cell morphology (cylindrical-approximation to spherical somatic compartment)
% FS soma
l=42;           % um, length
d=42;           % um, diameter

epas=-70;       % mV, passive leak reversal potential
ki=140;         % mM, intracellular potassium concentration
koinf=3.82;     % mM, steady-state extracellular potassium concentration
KAF=2e6;        % potassium accumulation factor
tauK=7;         % ms, extracellular potassium decay time constant
dshellK=70e-3;  % um, depth of extracellular shell for K+ diffusion
faraday=96487;  % s*A/mol, faraday constant (charge per mole of ions)

% mechanisms and interneuron-specific modifications to their kinetics (see [DS02] Fig 2)
mechanism_list={'DS00iNa','DS00iDR','DS00KDyn','pas'};
anV0=13-10;     % 10mV more hyperpolarized than pyramidal cell
bnV0=23-10;     % 10mV more hyperpolarized than pyramidal cell
amV0=-28-10;    % 10mV more hyperpolarized than pyramidal cell
bmV0=-1-12;     % 12mV more hyperpolarized than pyramidal cell
ahV0=-43.1-10;  % 10mV more hyperpolarized than pyramidal cell
bhV0=-13.1-10;  % 10mV more hyperpolarized than pyramidal cell
hnascale=2;     % sodium inactivation sped up 2x

% soma
A=d*l*pi;       % um2, cylinder surface area without the ends
Cm=1.2e-5;      % 1.2 uF/cm2 = 1.2e-5 nF/um2
gpas=1/30e5;    % Rm=30kOhm-cm2 = 30e5 MOhm-um2, gpas=1/Rm [uS/um2]
gnaf=45e-5;     % 45 mS/cm2 = 45-5 uS/um2
gkdr=18e-5;     % 18 mS/cm2 = 18-5 uS/um2
VshellK=pi*dshellK*l.*(d+dshellK);    % um3, volume of extracellular shell for K+ diffusion

spec=[];
spec.populations(1).name='FS';
spec.populations(1).size=Ni;
spec.populations(1).equations='dV/dt=(@current+Iapp)./Cm; Iapp=0; Cm=1; V(0)=-65';
spec.populations(1).mechanism_list=mechanism_list;
spec.populations(1).parameters={'Iapp',0,'Cm',Cm*A,'gpas',gpas*A,'epas',epas,...
  'gnaf',gnaf*A,'gkdr',gkdr*A,'KAF',KAF,'VshellK',VshellK,'koinf',koinf,'tauK',tauK,'ki',ki,'faraday',faraday,...
  'anV0',anV0,'bnV0',bnV0,'hnascale',hnascale,'amV0',amV0,'bmV0',bmV0,'ahV0',ahV0,'bhV0',bhV0};

data=dsSimulate(spec,'tspan',[0 2000],'vary',{'FS','Iapp',.1},'solver','rk1','verbose_flag',1);
dsPlot(data,'ylim',[-100 50]);

%% %%%%%%%%%%%%%%%%%%%%%%%%%%%%%%%%%%%%%%%%%%%%%%%%%%%%%%%%%%%%%%%%%%%%%%%%
% -------------------------------------------------------------------------
% 3) [DS02KS14FS] Fast-spiking GABAergic interneuron: (model (2) with modifications)
%    [DS02] FS with [TW03] ih and [KS14] FS soma parameters
% -------------------------------------------------------------------------
% Differences wrt [KS14] FS: [KS14] FS also includes A-type K+ current, 
% slow K+ current, and high-threshold N-type Ca2+ current. [KS14] FS has 3
% compartments (soma, dendrite, axon); the model here has a single soma.
% However, soma geometry, RMP, ~Vthresh, and electrophysiological regime (FS) are the same.

% [KS14]
% FS:   gnaf=.135 S/cm2, gkdr=.036, gh=1e-5; soma: l=27um, d=29; target: Rin=250MOhm (PV+; chandelier and basket neurons)
% RSNP: gnaf=.075 S/cm2, gkdr=.018, gh=2e-6; soma: l=42um, d=42; target: Rin=488Mohm (CB+; double bouquet and Martinotti-type)
% Rm=10kOhm/cm2

epas=-72;       % mV, passive leak reversal potential
ki=140;         % mM, intracellular potassium concentration
koinf=3.82;     % mM, steady-state extracellular potassium concentration
KAF=2e6;        % potassium accumulation factor
tauK=7;         % ms, extracellular potassium decay time constant
dshellK=70e-3;  % um, depth of extracellular shell for K+ diffusion
faraday=96487;  % s*A/mol, faraday constant (charge per mole of ions)

% mechanisms and interneuron-specific modifications to their kinetics
mechanism_list={'DS00iNa','DS00iDR','DS00KDyn','TW03iH','pas'};
anV0=13-10;     % 10mV more hyperpolarized than pyramidal cell
bnV0=23-10;     % 10mV more hyperpolarized than pyramidal cell
amV0=-28-10;    % 10mV more hyperpolarized than pyramidal cell
bmV0=-1-12;     % 12mV more hyperpolarized than pyramidal cell
ahV0=-43.1-10;  % 10mV more hyperpolarized than pyramidal cell
bhV0=-13.1-10;  % 10mV more hyperpolarized than pyramidal cell
hnascale=2;     % Na+ inactivation sped up 2x

% soma
l=27; d=29; A=d*l*pi; % um2, cylinder surface area without the ends
Cm=1.2e-5;            % 1.2 uF/cm2 = 1.2e-5 nF/um2
gpas=1/10e5;          % Rm=30kOhm-cm2 = 30e5 MOhm-um2, gpas=1/Rm
gnaf=135e-5;          % 45 mS/cm2 = 45-5 uS/um2
gkdr=36e-5;           % 18 mS/cm2 = 18-5 uS/um2
gH=.01e-5;            % max conductance of h-channel
VshellK=pi*dshellK*l.*(d+dshellK); % um3, volume of shell for extracellular K+ diffusion

spec=[];
spec.populations(1).name='FS';
spec.populations(1).size=1;
spec.populations(1).equations='dV/dt=(@current+Iapp*(t>50&t<150))./Cm; Cm=1; V(0)=-65; Iapp=0';
spec.populations(1).mechanism_list=mechanism_list;
spec.populations(1).parameters={'Iapp',0,'Cm',Cm*A,'gpas',gpas*A,'epas',epas,'gH',gH*A,...
  'gnaf',gnaf*A,'gkdr',gkdr*A,'KAF',KAF,'VshellK',VshellK,'koinf',koinf,'tauK',tauK,'ki',ki,'faraday',faraday,...
  'anV0',anV0,'bnV0',bnV0,'hnascale',hnascale,'amV0',amV0,'bmV0',bmV0,'ahV0',ahV0,'bhV0',bhV0};

data=dsSimulate(spec,'tspan',[0 200],'vary',{'FS','Iapp',[-.1 0 .1 .2]},'solver','rk1');
dsPlot(data,'ylim',[-120 60]);

Rin_fs=(1/gpas)/A % MOhm

%% %%%%%%%%%%%%%%%%%%%%%%%%%%%%%%%%%%%%%%%%%%%%%%%%%%%%%%%%%%%%%%%%%%%%%%%%
% -------------------------------------------------------------------------
% 4) [DS02KS14RSNP] Regular Spiking Non-Pyramidal GABAergic interneuron:
%    [DS02] FS with [TW03] ih and [KS14] RSNP soma parameters
% -------------------------------------------------------------------------
% Differences wrt [KS14] RSNP: [KS14] RSNP also includes A-type K+ current 
% and low-threshold T-type Ca2+ current. [KS14] RSNP has 3 compartments 
% (soma, dendrite, axon); the model here has a single soma.
% However, soma geometry, RMP, ~Vthresh, and electrophysiological regime (RS) are the same.

% [KS14]
% FS:   gnaf=.135 S/cm2, gkdr=.036, gh=1e-5; soma: l=27um, d=29; target: Rin=250MOhm (PV+; chandelier and basket neurons)
% RSNP: gnaf=.075 S/cm2, gkdr=.018, gh=2e-6; soma: l=42um, d=42; target: Rin=488Mohm (CB+; double bouquet and Martinotti-type)
% Rm=10kOhm/cm2

epas=-63.5;     % mV, passive leak reversal potential
ki=140;         % mM, intracellular potassium concentration
koinf=3.82;     % mM, steady-state extracellular potassium concentration
KAF=2e6;        % potassium accumulation factor
tauK=7;         % ms, extracellular potassium decay time constant
dshellK=70e-3;  % um, depth of extracellular shell for K+ diffusion
faraday=96487;  % s*A/mol, faraday constant (charge per mole of ions)

% mechanisms and interneuron-specific modifications to their kinetics
mechanism_list={'DS00iNa','DS00iDR','DS00KDyn','TW03iH','pas'};
anV0=13-10;     % 10mV more hyperpolarized than pyramidal cell
bnV0=23-10;     % 10mV more hyperpolarized than pyramidal cell
amV0=-28-10;    % 10mV more hyperpolarized than pyramidal cell
bmV0=-1-12;     % 12mV more hyperpolarized than pyramidal cell
ahV0=-43.1-10;  % 10mV more hyperpolarized than pyramidal cell
bhV0=-13.1-10;  % 10mV more hyperpolarized than pyramidal cell
hnascale=2;     % Na+ inactivation sped up 2x

% soma
l=42; d=42; A=d*l*pi; % um2, cylinder surface area without the ends
Cm=1.2e-5;            % 1.2 uF/cm2 = 1.2e-5 nF/um2
gpas=1/40e5;          % Rm=30kOhm-cm2 = 30e5 MOhm-um2, gpas=1/Rm
gnaf=75e-5;           % 45 mS/cm2 = 45-5 uS/um2
gkdr=18e-5;           % 18 mS/cm2 = 18-5 uS/um2
gH=.002e-5;
VshellK=pi*dshellK*l.*(d+dshellK); % um3, volume of extracellular shell for K+ diffusion

spec=[];
spec.populations(1).name='RSNP';
spec.populations(1).size=1;
spec.populations(1).equations='dV/dt=(@current+Iapp*(t>50&t<150))./Cm; Cm=1; V(0)=-65; Iapp=0';
spec.populations(1).mechanism_list=mechanism_list;
spec.populations(1).parameters={'Iapp',0,'Cm',Cm*A,'gpas',gpas*A,'epas',epas,'gH',gH*A,...
  'gnaf',gnaf*A,'gkdr',gkdr*A,'KAF',KAF,'VshellK',VshellK,'koinf',koinf,'tauK',tauK,'ki',ki,'faraday',faraday,...
  'anV0',anV0,'bnV0',bnV0,'hnascale',hnascale,'amV0',amV0,'bmV0',bmV0,'ahV0',ahV0,'bhV0',bhV0};

data=dsSimulate(spec,'tspan',[0 200],'vary',{'RSNP','Iapp',[-.1 0 .1 .2]},'solver','rk1');
dsPlot(data,'ylim',[-120 60]);

Rin_rs=(1/gpas)/A % MOhm

%% %%%%%%%%%%%%%%%%%%%%%%%%%%%%%%%%%%%%%%%%%%%%%%%%%%%%%%%%%%%%%%%%%%%%%%%%
% -------------------------------------------------------------------------
% 5) [DS02PYjs] Custom 2-compartment Pyramidal cell (PYs,PYd) ([DS02] + [TW03] dend ih)
% -------------------------------------------------------------------------
% This is the same as model (1) and [DS02] PY with the addition of a
% dendritic h-current (ih model taken from [TW03]). The PY dendritic h-current
% may be important for explaining the reduction in network rhythm frequency
% from gamma to beta2 that occurs when h-channels are blocked (in vitro; 
% unpublished result from LeBeau lab); it may also be relevant for 
% understanding how RSNP inhibition controls PY activity.
% Experimental motivation: Day, M., Carr, D. B., Ulrich, S., Ilijic, E., Tkatch, T., & Surmeier, D. J. (2005). Dendritic excitability of mouse frontal cortex pyramidal neurons is shaped by the interaction among HCN, Kir2, and Kleak channels. The Journal of neuroscience, 25(38), 8776-8787.

% # pyramidal cells
Ne=1;
% cell morphology (cylindrical compartments)
% pyramidal soma (length and diameter chosen to match surface area and internal resistance with a sphere of diam 23 microns)
ls=28.618;  % um, length
ds=21.840;  % um, diameter
% pyramidal dendrite
ld=650;     % um, length
dd=6.5;     % um, diameter

% shared parameters
epas=-70;     % mV, passive leak reversal potential
ki=140;       % mM, intracellular potassium concentration
koinf=3.82;   % mM, steady-state extracellular potassium concentration
KAF=2e6;      % potassium accumulation factor
tauK=7;       % ms, extracellular potassium decay time constant
dshellK=70e-3;% um, depth of extracellular shell for K+ diffusion
cao=2e3;      % uM, extracellular calcium concentration
cainf=50e-3;  % uM, steady-state intracellular calcium concentration
dshellCa=2e-4;% um, depth of intracellular shell for Ca2+ diffusion
faraday=96487;% s*A/mol, faraday constant (charge per mole of ions)

% ionic mechanisms and voltage dynamics present in both compartments (see [DS00] Methods for justification)
mechanism_list={'DS00iNa','DS00iNaP','DS00iDR','DS02iKS','DS02iHVA','DS00iKCa','DS00CaDyn','DS00KDyn','TW03iH','pas'};
state_equations='dV/dt=(@current+Iapp)./Cm; Iapp=0; Cm=1; V(0)=-65';
spec=[];

% soma
l=ls; d=ds; A=d*l*pi; % um2, cylinder surface area without the ends
Cm=1.2e-5;            % 1.2 uF/cm2 = 1.2e-5 nF/um2
gpas=1/30e5;          % Rm=30kOhm-cm2 = 30e5 MOhm-um2, gpas=1/Rm [uS/um2]
gnaf=117e-5;          % 117 mS/cm2 = 117e-5 uS/um2
gkdr=50e-5;           % 50 mS/cm2 = 50e-5 uS/um2
gnap=1.8e-5;          % uS/um2, persistent sodium channel
gks=.08e-5;           % uS/um2, slow potassium channel
ghva=.4e-5;           % uS/um2, high-voltage-activated Ca2+ channel
gkc=2.1e-5;           % uS/um2, V- and Ca2+-dependent potassium channel
gH=0e-5;              % max conductance of h-channel
tauCa=250;            % ms, calcium decay time constant
CAF=600;              % calcium accumulation factor
VshellK=pi*dshellK*l.*(d+dshellK);    % um3, volume of extracellular shell for K+ diffusion/accumulation
VshellCa=pi*dshellCa*l.*(d-dshellCa); % um3, volume of intracellular shell for Ca2+ diffusion/accumulation
spec.populations(1).name='Es';
spec.populations(1).size=Ne;
spec.populations(1).equations=state_equations;
spec.populations(1).mechanism_list=mechanism_list;
spec.populations(1).parameters={'Iapp',0,'Cm',Cm*A,'gpas',gpas*A,'epas',epas,'gH',gH*A,...
  'gnaf',gnaf*A,'gnap',gnap*A,'ghva',ghva*A,'gkdr',gkdr*A,'gks',gks*A,'gkc',gkc*A,...
  'CAF',CAF,'VshellCa',VshellCa,'cainf',cainf,'tauCa',tauCa,'cao',cao,...
  'KAF',KAF,'VshellK',VshellK,'koinf',koinf,'tauK',tauK,'ki',ki,'faraday',faraday};

% dendrite
l=ld; d=dd; A=d*l*pi; % um2, cylinder surface area without the ends
Cm=Cm*1.92;           % 2.304 uF/cm2
gpas=gpas*1.92;       % 1/(Rm/1.92)
gnaf=20e-5;           % 20 mS/cm2 = 20e-5 uS/um2
gkdr=14e-5;           % 14 mS/cm2 = 14e-5 uS/um2
gnap=.8e-5;           % uS/um2, persistent sodium channel
gks=.08e-5;           % uS/um2, slow potassium channel
ghva=.8e-5;           % uS/um2, high-voltage-activated Ca2+ channel
gkc=2.1e-5;           % uS/um2, V- and Ca2+-dependent potassium channel
gH=.01e-5;            % max conductance of h-channel
tauCa=120;            % ms, calcium decay time constant
CAF=600;              % calcium accumulation factor
VshellK=pi*dshellK*l.*(d+dshellK);    % um3, volume of extracellular shell for K+ diffusion/accumulation
VshellCa=pi*dshellCa*l.*(d-dshellCa); % um3, volume of intracellular shell for Ca2+ diffusion/accumulation
spec.populations(2).name='Ed';
spec.populations(2).size=Ne;
spec.populations(2).equations=state_equations;
spec.populations(2).mechanism_list=mechanism_list;
spec.populations(2).parameters={'Iapp',0,'Cm',Cm*A,'gpas',gpas*A,'epas',epas,'gH',gH*A,...
  'gnaf',gnaf*A,'gnap',gnap*A,'ghva',ghva*A,'gkdr',gkdr*A,'gks',gks*A,'gkc',gkc*A,...
  'CAF',CAF,'VshellCa',VshellCa,'cainf',cainf,'tauCa',tauCa,'cao',cao,...
  'KAF',KAF,'VshellK',VshellK,'koinf',koinf,'tauK',tauK,'ki',ki,'faraday',faraday};

% intercompartmental connections
Ri=1.5; % axial resistance [MOhm-um], Ri=150 Ohm-cm
% collect relevant info
compartments={'Es' 'Ed'};
lengths     =[ls   ld];
diameters   =[ds   dd];
connections={[1 2],[2 1]};
% add connections to specification
for c=1:length(connections)
  src=connections{c}(1);
  dst=connections{c}(2);
  spec.connections(c).direction=[compartments{src} '->' compartments{dst}];
  spec.connections(c).mechanism_list={'iCOM'};
  gCOM=1/mean(Ri*4*lengths./(pi*diameters.^2));
  spec.connections(c).parameters={'gCOM',gCOM};
end

data=dsSimulate(spec,'tspan',[0 2000],'vary',{'Es','Iapp',.1},'solver','rk1','verbose_flag',1);
dsPlot(data,'ylim',[-100 50]);

% characterize cell intrinsic properties
if 0 
  amps=[ -.2 -.1 0 .1 .2 .5]*10; onset=50; offset=250; tspan=[0 300];
  data = ProbeCellProperties(spec,'membrane_area',A,'amplitudes',amps,'onset',onset,'offset',offset,'tspan',tspan);
  dsPlot(data,'ylim',[-120 60])
  s = CalcCellProperties(data); l=spec.populations(1).name;
  [s.(l).tau_m s.(l).RMP s.(l).V_thresh s.(l).AP_dur s.(l).Ih_abssag s.(l).AHP_time2trough]
  % PYs: [48.41   -59.96      -37.93        1.1400          0           4.1500
end

% PY taum is slower in superficial layers (L2/3 25+/-20ms vs L5/6 14+/-7ms)
% L2/3: http://www.neuroelectro.org/neuron/110/
% L5/6: http://www.neuroelectro.org/neuron/111/
% Tip: decrease Rm (or Cm) in deep layer wrt superficial layer

%% %%%%%%%%%%%%%%%%%%%%%%%%%%%%%%%%%%%%%%%%%%%%%%%%%%%%%%%%%%%%%%%%%%%%%%%%
% -------------------------------------------------------------------------
% 6) [DS02FSjs] Custom Fast-spiking GABAergic interneuron: (model (3) with modifications)
%    [DS02] FS with [TW03] ih and [KS14] FS soma parameters
% -------------------------------------------------------------------------
% Modifications wrt model (3): morphological dimensions (geometry) was
% adjusted to account for the effect of removing dendrite and axon
% compartments from [KS14] FS on the input resistance. Consequence: this model
% (6) FS cell is a better match to the experimental data than model (3).
% 
% Also, modified gkdr wrt [KS14]: increased from gkdr=36e-5 to 50e-5 to achieve
% slightly shorter spike-width and lower Vthresh. This change also produces a
% gkdr/gnaf ratio that more closely matches [DS02].

%{

[KS14]
FS:   l=27um, d=29; target: Rin=250MOhm; Vthresh=-53mV; RMP=-73mV (PV+; chandelier and basket neurons)
RSNP: l=42um, d=42; target: Rin=488Mohm; Vthresh=-51mV; RMP=-64mV (CB+; double bouquet and Martinotti-type)

Modifying FS and RSNP parameters to get more accurate input resistances 
and their relative differences:

Rin=Rm/area, and Rin is larger in a smaller cell.
Modifying area has significant effects on electrophysiological regime.
Modifying Rm changes membrane time constant (taum=Rm*Cm).

% Approach: modify Rm to achieve desired Rin and reasonable taum.

% Experimental constraints:
% Gorelova, N., Seamans, J. K., & Yang, C. R. (2002). Mechanisms of dopamine activation of fast-spiking interneurons that exert inhibition in rat prefrontal cortex. Journal of neurophysiology, 88(6), 3150-3166.
  experimental result (rat PrL): FS Rin=330MOhm, RSNP Rin=450MOhm, LTS Rin=1200MOhm
% Povysheva, Nadezhda V., et al. "Parvalbumin-positive basket interneurons in monkey and rat prefrontal cortex." Journal of neurophysiology 100.4 (2008): 2348-2360. full text: http://jn.physiology.org/content/100/4/2348.long
  experimental result (monkey DLPFC): FS taum=10ms

l=27; d=29; A=d*l*pi; % um2, cylinder surface area without the ends
gpas=1/8e5;          
Rin_fs=(1/gpas)/A % MOhm
taum_fs=(1/gpas)*Cm

l=42; d=42; A=d*l*pi; % um2, cylinder surface area without the ends
gpas=1/30e5;          
Rin_rs=(1/gpas)/A % MOhm
taum_rs=(1/gpas)*Cm

non-ideally constrained:
- CB+ cell soma (double bouquet) is not typically larger than PV+ cell soma (basket).
- taum_rs is unknown. as implemented, it is 3-4x taum_fs.

%}

epas=-72;       % mV, passive leak reversal potential
ki=140;         % mM, intracellular potassium concentration
koinf=3.82;     % mM, steady-state extracellular potassium concentration
KAF=2e6;        % potassium accumulation factor
tauK=7;         % ms, extracellular potassium decay time constant
dshellK=70e-3;  % um, depth of extracellular shell for K+ diffusion
faraday=96487;  % s*A/mol, faraday constant (charge per mole of ions)

% mechanisms and interneuron-specific modifications to their kinetics
mechanism_list={'DS00iNa','DS00iDR','DS00KDyn','TW03iH','pas'};
anV0=13-10;     % 10mV more hyperpolarized than pyramidal cell
bnV0=23-10;     % 10mV more hyperpolarized than pyramidal cell
amV0=-28-10;    % 10mV more hyperpolarized than pyramidal cell
bmV0=-1-12;     % 12mV more hyperpolarized than pyramidal cell
ahV0=-43.1-10;  % 10mV more hyperpolarized than pyramidal cell
bhV0=-13.1-10;  % 10mV more hyperpolarized than pyramidal cell
hnascale=2;     % Na+ inactivation sped up 2x

% soma
l=27; d=29; A=d*l*pi; % um2, cylinder surface area without the ends
Cm=1.2e-5;            % 1.2 uF/cm2 = 1.2e-5 nF/um2
gpas=1/8e5;          % Rm=30kOhm-cm2 = 30e5 MOhm-um2, gpas=1/Rm
gnaf=135e-5;          % 45 mS/cm2 = 45-5 uS/um2
gkdr=50e-5; % [KS14] 36e-5          % 18 mS/cm2 = 18-5 uS/um2
gH=.01e-5;           % max conductance of h-channel
VshellK=pi*dshellK*l.*(d+dshellK); % um3, volume of shell for extracellular K+ diffusion

spec=[];
spec.populations(1).name='FS';
spec.populations(1).size=1;
spec.populations(1).equations='dV/dt=(@current+Iapp*(t>50&t<150))./Cm; Cm=1; V(0)=-65; Iapp=0';
spec.populations(1).mechanism_list=mechanism_list;
spec.populations(1).parameters={'Iapp',0,'Cm',Cm*A,'gpas',gpas*A,'epas',epas,'gH',gH*A,...
  'gnaf',gnaf*A,'gkdr',gkdr*A,'KAF',KAF,'VshellK',VshellK,'koinf',koinf,'tauK',tauK,'ki',ki,'faraday',faraday,...
  'anV0',anV0,'bnV0',bnV0,'hnascale',hnascale,'amV0',amV0,'bmV0',bmV0,'ahV0',ahV0,'bhV0',bhV0};

data=dsSimulate(spec,'tspan',[0 200],'vary',{'FS','Iapp',[-.1 0 .1 .2 .5]},'solver','rk1');
dsPlot(data,'ylim',[-120 60]);

Rin_fs=(1/gpas)/A % MOhm

% characterize cell intrinsic properties
if 0 
  amps=[ -.2 -.1 0 .1 .2 .5]*10; onset=50; offset=250; tspan=[0 300];
  data = ProbeCellProperties(spec,'membrane_area',A,'amplitudes',amps,'onset',onset,'offset',offset,'tspan',tspan);
  dsPlot(data,'ylim',[-120 60])
  s = CalcCellProperties(data); l=spec.populations(1).name;
  [s.(l).tau_m s.(l).RMP s.(l).V_thresh s.(l).AP_dur s.(l).Ih_abssag s.(l).AHP_time2trough]
%gkdr=36:[8.4000   -72.8196     -48.7907    0.8300         0.9751        2.6400]
%gkdr=50:[8.1600   -73.2513     -49.3951    0.7800         0.9894        2.2700]
  % [KS14] target: Rin=250-350MOhm; Vthresh=-53mV; RMP=-73mV
end

%% %%%%%%%%%%%%%%%%%%%%%%%%%%%%%%%%%%%%%%%%%%%%%%%%%%%%%%%%%%%%%%%%%%%%%%%%
% -------------------------------------------------------------------------
% 7) [DS02RSNPjs] Custom Regular Spiking Non-Pyramidal GABAergic interneuron: (model (4) with modifications)
%    [DS02] FS with [TW03] ih and [KS14] RSNP soma parameters
% -------------------------------------------------------------------------
% Modifications wrt model (4): morphological dimensions (geometry) was
% adjusted to account for the effect of removing dendrite and axon
% compartments from [KS14] RS on the input resistance. Consequence: this model
% (6) RSNP cell is a better match to the experimental data than model (4).

epas=-63.5;     % mV, passive leak reversal potential
ki=140;         % mM, intracellular potassium concentration
koinf=3.82;     % mM, steady-state extracellular potassium concentration
KAF=2e6;        % potassium accumulation factor
tauK=7;         % ms, extracellular potassium decay time constant
dshellK=70e-3;  % um, depth of extracellular shell for K+ diffusion
faraday=96487;  % s*A/mol, faraday constant (charge per mole of ions)

% mechanisms and interneuron-specific modifications to their kinetics
mechanism_list={'DS00iNa','DS00iDR','DS00KDyn','TW03iH','pas'};
anV0=13-10;     % 10mV more hyperpolarized than pyramidal cell
bnV0=23-10;     % 10mV more hyperpolarized than pyramidal cell
amV0=-28-10;    % 10mV more hyperpolarized than pyramidal cell
bmV0=-1-12;     % 12mV more hyperpolarized than pyramidal cell
ahV0=-43.1-10;  % 10mV more hyperpolarized than pyramidal cell
bhV0=-13.1-10;  % 10mV more hyperpolarized than pyramidal cell
hnascale=2;     % Na+ inactivation sped up 2x

% soma
l=42; d=42; A=d*l*pi; % um2, cylinder surface area without the ends
Cm=1.2e-5;            % 1.2 uF/cm2 = 1.2e-5 nF/um2
gpas=1/30e5;          % Rm=30kOhm-cm2 = 30e5 MOhm-um2, gpas=1/Rm
gnaf=75e-5;           % 45 mS/cm2 = 45-5 uS/um2
gkdr=18e-5;           % 18 mS/cm2 = 18-5 uS/um2
gH=.002e-5;
VshellK=pi*dshellK*l.*(d+dshellK); % um3, volume of extracellular shell for K+ diffusion

spec=[];
spec.populations(1).name='RSNP';
spec.populations(1).size=1;
spec.populations(1).equations='dV/dt=(@current+Iapp*(t>50&t<150))./Cm; Cm=1; V(0)=-65; Iapp=0';
spec.populations(1).mechanism_list=mechanism_list;
spec.populations(1).parameters={'Iapp',0,'Cm',Cm*A,'gpas',gpas*A,'epas',epas,'gH',gH*A,...
  'gnaf',gnaf*A,'gkdr',gkdr*A,'KAF',KAF,'VshellK',VshellK,'koinf',koinf,'tauK',tauK,'ki',ki,'faraday',faraday,...
  'anV0',anV0,'bnV0',bnV0,'hnascale',hnascale,'amV0',amV0,'bmV0',bmV0,'ahV0',ahV0,'bhV0',bhV0};

data=dsSimulate(spec,'tspan',[0 200],'vary',{'RSNP','Iapp',[-.1 0 .1 .2 .5]},'solver','rk1');
dsPlot(data,'ylim',[-120 60]);

Rin_rs=(1/gpas)/A % MOhm

% characterize cell intrinsic properties
if 0 
  amps=[ -.2 -.1 0 .1 .2 .5]*10; onset=50; offset=250; tspan=[0 300];
  data = ProbeCellProperties(spec,'membrane_area',A,'amplitudes',amps,'onset',onset,'offset',offset,'tspan',tspan);
  dsPlot(data,'ylim',[-120 60])
  s = CalcCellProperties(data); l=spec.populations(1).name;
  [s.(l).tau_m s.(l).RMP s.(l).V_thresh s.(l).AP_dur s.(l).Ih_abssag s.(l).AHP_time2trough]
  % [31.4500    -65.9983  -45.8906          0.8800         0          3.7400]
end

% Additional References (experiments):
% [ZK05] Zaitsev, A. V., Gonzalez-Burgos, G., Povysheva, N. V., Kröner, S., Lewis, D. A., & Krimer, L. S. (2005). Localization of calcium-binding proteins in physiologically and morphologically characterized interneurons of monkey dorsolateral prefrontal cortex. Cerebral Cortex, 15(8), 1178-1186.
% [ZL09] (DLPFC L2/3 INs) Zaitsev, A. V., Povysheva, N. V., Gonzalez-Burgos, G., Rotaru, D., Fish, K. N., Krimer, L. S., & Lewis, D. A. (2009). Interneuron diversity in layers 2–3 of monkey prefrontal cortex. Cerebral cortex, 19(7), 1597-1615.
% [K93] (rat PrL L5 FS vs LTS) Kawaguchi, Y. A. S. U. O. (1993). Groupings of nonpyramidal and pyramidal cells with specific physiological and morphological characteristics in rat frontal cortex. Journal of neurophysiology, 69(2), 416-431.
% [GG03] Gao, W. J., Wang, Y., & Goldman-Rakic, P. S. (2003). Dopamine modulation of perisomatic and peridendritic inhibition in prefrontal cortex. The Journal of neuroscience, 23(5), 1622-1630.
% [VM09] Van Aerde, K. I., Mann, E. O., Canto, C. B., Heistek, T. S., Linkenkaer‐Hansen, K., Mulder, A. B., ... & Mansvelder, H. D. (2009). Flexible spike timing of layer 5 neurons during dynamic beta oscillation shifts in rat prefrontal cortex. The Journal of physiology, 587(21), 5177-5196. http://onlinelibrary.wiley.com/doi/10.1113/jphysiol.2009.178384/full
% Results:
  % [ZK05]: Rin [MOhm]: FS (235+/-68), RSNP (582+/-195), IS (585+/-137)
  % [K93]: LTS RMP: -64mV (same as CB RS in [KS14]) > FS RMP
  % [GG03],[VM09]: FS spike-width < nonFS spike-width
  % [ZL09]: longer AP observed in DLPFC CB+ than PV+ INs

%{
% Custom model summary (cell intrinsic properties):
[       tau_m     RMP       V_thresh      AP_dur      Ih_abssag 	AHP_time2trough]
 PYs:  [48.41   -59.96      -37.93        1.1400         0            4.1500
 PYs+d:[        -66
 FS:   [8.16    -73.25      -49.3951      0.7800         0.9894       2.2700]
 RSNP: [31.45   -66.00      -45.8906      0.8800         0            3.7400]

PY taum is slower in superficial layers (L2/3 25+/-20ms vs L5/6 14+/-7ms)
L2/3: http://www.neuroelectro.org/neuron/110/
L5/6: http://www.neuroelectro.org/neuron/111/
Tip: decrease Rm (or Cm) in deep layer wrt superficial layer
%}
```
