## Supplemental File 1 for "A biophysical model of striatal microcircuits suggests delta/theta-rhythmically interleaved gamma and beta oscillations mediate periodicity in motor control": setupDynaSimPath.m

```
% Attempts to add DynaSim to path from within DynaSim demos folder

% If DynaSim isn't already in the system path, this code attempts to guess
% the location of the main DynaSim folder. If cannot find it, it returns
% an error.

if ~exist('dsSimulate','file')                                            % If DynaSim not in path...
    % Get name of parent folder
    [parentfolder_full,~] = fileparts(pwd);
    [~,parentfolder] = fileparts(parentfolder_full);
    
    % Get name of current folder
    [currfolder_full] = pwd;
    [~,currfolder] = fileparts(currfolder_full);
    
    if ~isempty(regexpi(parentfolder,'DynaSim'))                                % Check if parent folder is DynaSim
        addpath(genpath(parentfolder_full));                                    %     If is, add to path
        rmpath(genpath(fullfile(parentfolder_full,'.git')));                    %        (Remove .git folder)
    elseif ~isempty(regexpi(currfolder,'DynaSim'))                              % Check if current folder is DynaSim
        addpath(genpath(currfolder_full));                                      %     If is, add to path
        rmpath(genpath(fullfile(currfolder_full,'.git')));                      %         (Remove .git folder)
    else
        % Otherwise, return error
        error('Failed adding the DynaSim folder to the MATLAB path automatically. Manually add the DynaSim folder to the MATLAB path - e.g. run addpath(genpath(DynaSimPath))');
    end
end
```
