## Supplemental File 1 for "A biophysical model of striatal microcircuits suggests delta/theta-rhythmically interleaved gamma and beta oscillations mediate periodicity in motor control": append_pdfs.m

```
%APPEND_PDFS Appends/concatenates multiple PDF files
%
% Example:
%   append_pdfs(output, input1, input2, ...)
%   append_pdfs(output, input_list{:})
%   append_pdfs test.pdf temp1.pdf temp2.pdf
%
% This function appends multiple PDF files to an existing PDF file, or
% concatenates them into a PDF file if the output file doesn't yet exist.
%
% This function requires that you have ghostscript installed on your
% system. Ghostscript can be downloaded from: http://www.ghostscript.com
%
% IN:
%    output - string of output file name (including the extension, .pdf).
%             If it exists it is appended to; if not, it is created.
%    input1 - string of an input file name (including the extension, .pdf).
%             All input files are appended in order.
%    input_list - cell array list of input file name strings. All input
%                 files are appended in order.

% Copyright: Oliver Woodford, 2011

% Thanks to Reinhard Knoll for pointing out that appending multiple pdfs in
% one go is much faster than appending them one at a time.

% Thanks to Michael Teo for reporting the issue of a too long command line.
% Issue resolved on 5/5/2011, by passing gs a command file.

% Thanks to Martin Wittmann for pointing out the quality issue when
% appending multiple bitmaps.
% Issue resolved (to best of my ability) 1/6/2011, using the prepress
% setting

% 26/02/15: If temp dir is not writable, use the output folder for temp
%           files when appending (Javier Paredes); sanity check of inputs

function append_pdfs(varargin)

if nargin < 2,  return;  end  % sanity check

% Are we appending or creating a new file
append = exist(varargin{1}, 'file') == 2;
output = [tempname '.pdf'];
try
    % Ensure that the temp dir is writable (Javier Paredes 26/2/15)
    fid = fopen(output,'w');
    fwrite(fid,1);
    fclose(fid);
    delete(output);
    isTempDirOk = true;
catch
    % Temp dir is not writable, so use the output folder
    [dummy,fname,fext] = fileparts(output); %#ok<ASGLU>
    fpath = fileparts(varargin{1});
    output = fullfile(fpath,[fname fext]);
    isTempDirOk = false;
end
if ~append
    output = varargin{1};
    varargin = varargin(2:end);
end
% Create the command file
if isTempDirOk
    cmdfile = [tempname '.txt'];
else
    cmdfile = fullfile(fpath,[fname '.txt']);
end
fh = fopen(cmdfile, 'w');
fprintf(fh, '-q -dNOPAUSE -dBATCH -sDEVICE=pdfwrite -dPDFSETTINGS=/prepress -sOutputFile="%s" -f', output);
fprintf(fh, ' "%s"', varargin{:});
fclose(fh);
% Call ghostscript
ghostscript(['@"' cmdfile '"']);
% Delete the command file
delete(cmdfile);
% Rename the file if needed
if append
    movefile(output, varargin{1});
end
end
```
