## Supplemental File 1 for "A biophysical model of striatal microcircuits suggests delta/theta-rhythmically interleaved gamma and beta oscillations mediate periodicity in motor control": copyfig.m

```
function fh = copyfig(fh)
%COPYFIG Create a copy of a figure, without changing the figure
%
% Examples:
%   fh_new = copyfig(fh_old)
%
% This function will create a copy of a figure, but not change the figure,
% as copyobj sometimes does, e.g. by changing legends.
%
% IN:
%    fh_old - The handle of the figure to be copied. Default: gcf.
%
% OUT:
%    fh_new - The handle of the created figure.

% Copyright (C) Oliver Woodford 2012

% 26/02/15: If temp dir is not writable, use the dest folder for temp
%           destination files (Javier Paredes)
% 15/04/15: Suppress warnings during copyobj (Dun Kirk comment on FEX page 2013-10-02)

    % Set the default
    if nargin == 0
        fh = gcf;
    end
    % Is there a legend?
    if isempty(findall(fh, 'Type', 'axes', 'Tag', 'legend'))
        % Safe to copy using copyobj
        oldWarn = warning('off'); %#ok<WNOFF>  %Suppress warnings during copyobj (Dun Kirk comment on FEX page 2013-10-02)
        fh = copyobj(fh, 0);
        warning(oldWarn);
    else
        % copyobj will change the figure, so save and then load it instead
        tmp_nam = [tempname '.fig'];
        try
            % Ensure that the temp dir is writable (Javier Paredes 26/2/15)
            fid = fopen(tmp_nam,'w');
            fwrite(fid,1);
            fclose(fid);
            delete(tmp_nam);  % cleanup
        catch
            % Temp dir is not writable, so use the current folder
            [dummy,fname,fext] = fileparts(tmp_nam); %#ok<ASGLU>
            fpath = pwd;
            tmp_nam = fullfile(fpath,[fname fext]);
        end
        hgsave(fh, tmp_nam);
        fh = hgload(tmp_nam);
        delete(tmp_nam);
    end
end
```
