## Supplemental File 1 for "A biophysical model of striatal microcircuits suggests delta/theta-rhythmically interleaved gamma and beta oscillations mediate periodicity in motor control": crop_borders.m

```
function [A, vA, vB, bb_rel] = crop_borders(A, bcol, padding, crop_amounts)
%CROP_BORDERS Crop the borders of an image or stack of images
%
%   [B, vA, vB, bb_rel] = crop_borders(A, bcol, [padding])
%
%IN:
%   A - HxWxCxN stack of images.
%   bcol - Cx1 background colour vector.
%   padding - scalar indicating how much padding to have in relation to
%             the cropped-image-size (0<=padding<=1). Default: 0
%   crop_amounts - 4-element vector of crop amounts: [top,right,bottom,left]
%             where NaN/Inf indicate auto-cropping, 0 means no cropping,
%             and any other value mean cropping in pixel amounts.
%
%OUT:
%   B - JxKxCxN cropped stack of images.
%   vA     - coordinates in A that contain the cropped image
%   vB     - coordinates in B where the cropped version of A is placed
%   bb_rel - relative bounding box (used for eps-cropping)

%{
% 06/03/15: Improved image cropping thanks to Oscar Hartogensis
% 08/06/15: Fixed issue #76: case of transparent figure bgcolor
% 21/02/16: Enabled specifying non-automated crop amounts
% 04/04/16: Fix per Luiz Carvalho for old Matlab releases
% 23/10/16: Fixed issue #175: there used to be a 1px minimal padding in case of crop, now removed
%}

    if nargin < 3
        padding = 0;
    end
    if nargin < 4
        crop_amounts = nan(1,4);  % =auto-cropping
    end
    crop_amounts(end+1:4) = NaN;  % fill missing values with NaN

    [h, w, c, n] = size(A);
    if isempty(bcol)  % case of transparent bgcolor
        bcol = A(ceil(end/2),1,:,1);
    end
    if isscalar(bcol)
        bcol = bcol(ones(c, 1));
    end

    % Crop margin from left
    if ~isfinite(crop_amounts(4))
        bail = false;
        for l = 1:w
            for a = 1:c
                if ~all(col(A(:,l,a,:)) == bcol(a))
                    bail = true;
                    break;
                end
            end
            if bail
                break;
            end
        end
    else
        l = 1 + abs(crop_amounts(4));
    end

    % Crop margin from right
    if ~isfinite(crop_amounts(2))
        bcol = A(ceil(end/2),w,:,1);
        bail = false;
        for r = w:-1:l
            for a = 1:c
                if ~all(col(A(:,r,a,:)) == bcol(a))
                    bail = true;
                    break;
                end
            end
            if bail
                break;
            end
        end
    else
        r = w - abs(crop_amounts(2));
    end

    % Crop margin from top
    if ~isfinite(crop_amounts(1))
        bcol = A(1,ceil(end/2),:,1);
        bail = false;
        for t = 1:h
            for a = 1:c
                if ~all(col(A(t,:,a,:)) == bcol(a))
                    bail = true;
                    break;
                end
            end
            if bail
                break;
            end
        end
    else
        t = 1 + abs(crop_amounts(1));
    end

    % Crop margin from bottom
    bcol = A(h,ceil(end/2),:,1);
    if ~isfinite(crop_amounts(3))
        bail = false;
        for b = h:-1:t
            for a = 1:c
                if ~all(col(A(b,:,a,:)) == bcol(a))
                    bail = true;
                    break;
                end
            end
            if bail
                break;
            end
        end
    else
        b = h - abs(crop_amounts(3));
    end

    if padding == 0  % no padding
        % Issue #175: there used to be a 1px minimal padding in case of crop, now removed
        %{
        if ~isequal([t b l r], [1 h 1 w]) % Check if we're actually croppping
            padding = 1; % Leave one boundary pixel to avoid bleeding on resize
            bcol(:) = nan;  % make the 1px padding transparent
        end
        %}
    elseif abs(padding) < 1  % pad value is a relative fraction of image size
        padding = sign(padding)*round(mean([b-t r-l])*abs(padding)); % ADJUST PADDING
    else  % pad value is in units of 1/72" points
        padding = round(padding);  % fix cases of non-integer pad value
    end

    if padding > 0  % extra padding
        % Create an empty image, containing the background color, that has the
        % cropped image size plus the padded border
        B = repmat(bcol,[(b-t)+1+padding*2,(r-l)+1+padding*2,1,n]);  % Fix per Luiz Carvalho
        % vA - coordinates in A that contain the cropped image
        vA = [t b l r];
        % vB - coordinates in B where the cropped version of A will be placed
        vB = [padding+1, (b-t)+1+padding, padding+1, (r-l)+1+padding];
        % Place the original image in the empty image
        B(vB(1):vB(2), vB(3):vB(4), :, :) = A(vA(1):vA(2), vA(3):vA(4), :, :);
        A = B;
    else  % extra cropping
        vA = [t-padding b+padding l-padding r+padding];
        A = A(vA(1):vA(2), vA(3):vA(4), :, :);
        vB = [NaN NaN NaN NaN];
    end

    % For EPS cropping, determine the relative BoundingBox - bb_rel
    bb_rel = [l-1 h-b-1 r+1 h-t+1]./[w h w h];
end

function A = col(A)
    A = A(:);
end
```
