## Supplemental File 1 for "A biophysical model of striatal microcircuits suggests delta/theta-rhythmically interleaved gamma and beta oscillations mediate periodicity in motor control": eps2pdf.m

```
function eps2pdf(source, dest, crop, append, gray, quality, gs_options)
%EPS2PDF  Convert an eps file to pdf format using ghostscript
%
% Examples:
%   eps2pdf source dest
%   eps2pdf(source, dest, crop)
%   eps2pdf(source, dest, crop, append)
%   eps2pdf(source, dest, crop, append, gray)
%   eps2pdf(source, dest, crop, append, gray, quality)
%   eps2pdf(source, dest, crop, append, gray, quality, gs_options)
%
% This function converts an eps file to pdf format. The output can be
% optionally cropped and also converted to grayscale. If the output pdf
% file already exists then the eps file can optionally be appended as a new
% page on the end of the eps file. The level of bitmap compression can also
% optionally be set.
%
% This function requires that you have ghostscript installed on your
% system. Ghostscript can be downloaded from: http://www.ghostscript.com
%
% Inputs:
%   source  - filename of the source eps file to convert. The filename is
%             assumed to already have the extension ".eps".
%   dest    - filename of the destination pdf file. The filename is assumed
%             to already have the extension ".pdf".
%   crop    - boolean indicating whether to crop the borders off the pdf.
%             Default: true.
%   append  - boolean indicating whether the eps should be appended to the
%             end of the pdf as a new page (if the pdf exists already).
%             Default: false.
%   gray    - boolean indicating whether the output pdf should be grayscale
%             or not. Default: false.
%   quality - scalar indicating the level of image bitmap quality to
%             output. A larger value gives a higher quality. quality > 100
%             gives lossless output. Default: ghostscript prepress default.
%   gs_options - optional ghostscript options (e.g.: '-dNoOutputFonts'). If
%                multiple options are needed, enclose in call array: {'-a','-b'}

% Copyright (C) Oliver Woodford 2009-2014, Yair Altman 2015-

% Suggestion of appending pdf files provided by Matt C at:
% http://www.mathworks.com/matlabcentral/fileexchange/23629

% Thank you to Fabio Viola for pointing out compression artifacts, leading
% to the quality setting.
% Thank you to Scott for pointing out the subsampling of very small images,
% which was fixed for lossless compression settings.

% 9/12/2011 Pass font path to ghostscript.
% 26/02/15: If temp dir is not writable, use the dest folder for temp
%           destination files (Javier Paredes)
% 28/02/15: Enable users to specify optional ghostscript options (issue #36)
% 01/03/15: Upon GS error, retry without the -sFONTPATH= option (this might solve
%           some /findfont errors according to James Rankin, FEX Comment 23/01/15)
% 23/06/15: Added extra debug info in case of ghostscript error; code indentation
% 04/10/15: Suggest a workaround for issue #41 (missing font path; thanks Mariia Fedotenkova)
% 22/02/16: Bug fix from latest release of this file (workaround for issue #41)

    % Intialise the options string for ghostscript
    options = ['-q -dNOPAUSE -dBATCH -sDEVICE=pdfwrite -dPDFSETTINGS=/prepress -sOutputFile="' dest '"'];
    % Set crop option
    if nargin < 3 || crop
        options = [options ' -dEPSCrop'];
    end
    % Set the font path
    fp = font_path();
    if ~isempty(fp)
        options = [options ' -sFONTPATH="' fp '"'];
    end
    % Set the grayscale option
    if nargin > 4 && gray
        options = [options ' -sColorConversionStrategy=Gray -dProcessColorModel=/DeviceGray'];
    end
    % Set the bitmap quality
    if nargin > 5 && ~isempty(quality)
        options = [options ' -dAutoFilterColorImages=false -dAutoFilterGrayImages=false'];
        if quality > 100
            options = [options ' -dColorImageFilter=/FlateEncode -dGrayImageFilter=/FlateEncode -c ".setpdfwrite << /ColorImageDownsampleThreshold 10 /GrayImageDownsampleThreshold 10 >> setdistillerparams"'];
        else
            options = [options ' -dColorImageFilter=/DCTEncode -dGrayImageFilter=/DCTEncode'];
            v = 1 + (quality < 80);
            quality = 1 - quality / 100;
            s = sprintf('<< /QFactor %.2f /Blend 1 /HSample [%d 1 1 %d] /VSample [%d 1 1 %d] >>', quality, v, v, v, v);
            options = sprintf('%s -c ".setpdfwrite << /ColorImageDict %s /GrayImageDict %s >> setdistillerparams"', options, s, s);
        end
    end
    % Enable users to specify optional ghostscript options (issue #36)
    if nargin > 6 && ~isempty(gs_options)
        if iscell(gs_options)
            gs_options = sprintf(' %s',gs_options{:});
        elseif ~ischar(gs_options)
            error('gs_options input argument must be a string or cell-array of strings');
        else
            gs_options = [' ' gs_options];
        end
        options = [options gs_options];
    end
    % Check if the output file exists
    if nargin > 3 && append && exist(dest, 'file') == 2
        % File exists - append current figure to the end
        tmp_nam = tempname;
        try
            % Ensure that the temp dir is writable (Javier Paredes 26/2/15)
            fid = fopen(tmp_nam,'w');
            fwrite(fid,1);
            fclose(fid);
            delete(tmp_nam);
        catch
            % Temp dir is not writable, so use the dest folder
            [dummy,fname,fext] = fileparts(tmp_nam); %#ok<ASGLU>
            fpath = fileparts(dest);
            tmp_nam = fullfile(fpath,[fname fext]);
        end
        % Copy the file
        copyfile(dest, tmp_nam);
        % Add the output file names
        options = [options ' -f "' tmp_nam '" "' source '"'];
        try
            % Convert to pdf using ghostscript
            [status, message] = ghostscript(options);
        catch me
            % Delete the intermediate file
            delete(tmp_nam);
            rethrow(me);
        end
        % Delete the intermediate file
        delete(tmp_nam);
    else
        % File doesn't exist or should be over-written
        % Add the output file names
        options = [options ' -f "' source '"'];
        % Convert to pdf using ghostscript
        [status, message] = ghostscript(options);
    end
    % Check for error
    if status
        % Retry without the -sFONTPATH= option (this might solve some GS
        % /findfont errors according to James Rankin, FEX Comment 23/01/15)
        orig_options = options;
        if ~isempty(fp)
            options = regexprep(options, ' -sFONTPATH=[^ ]+ ',' ');
            status = ghostscript(options);
            if ~status, return; end  % hurray! (no error)
        end
        % Report error
        if isempty(message)
            error('Unable to generate pdf. Check destination directory is writable.');
        elseif ~isempty(strfind(message,'/typecheck in /findfont'))
            % Suggest a workaround for issue #41 (missing font path)
            font_name = strtrim(regexprep(message,'.*Operand stack:\s*(.*)\s*Execution.*','$1'));
            fprintf(2, 'Ghostscript error: could not find the following font(s): %s\n', font_name);
            fpath = fileparts(mfilename('fullpath'));
            gs_fonts_file = fullfile(fpath, '.ignore', 'gs_font_path.txt');
            fprintf(2, '  try to add the font''s folder to your %s file\n\n', gs_fonts_file);
            error('export_fig error');
        else
            fprintf(2, '\nGhostscript error: perhaps %s is open by another application\n', dest);
            if ~isempty(gs_options)
                fprintf(2, '  or maybe the%s option(s) are not accepted by your GS version\n', gs_options);
            end
            fprintf(2, 'Ghostscript options: %s\n\n', orig_options);
            error(message);
        end
    end
end

% Function to return (and create, where necessary) the font path
function fp = font_path()
    fp = user_string('gs_font_path');
    if ~isempty(fp)
        return
    end
    % Create the path
    % Start with the default path
    fp = getenv('GS_FONTPATH');
    % Add on the typical directories for a given OS
    if ispc
        if ~isempty(fp)
            fp = [fp ';'];
        end
        fp = [fp getenv('WINDIR') filesep 'Fonts'];
    else
        if ~isempty(fp)
            fp = [fp ':'];
        end
        fp = [fp '/usr/share/fonts:/usr/local/share/fonts:/usr/share/fonts/X11:/usr/local/share/fonts/X11:/usr/share/fonts/truetype:/usr/local/share/fonts/truetype'];
    end
    user_string('gs_font_path', fp);
end
```
