## Supplemental File 1 for "A biophysical model of striatal microcircuits suggests delta/theta-rhythmically interleaved gamma and beta oscillations mediate periodicity in motor control": export_fig.m

```
function [imageData, alpha] = export_fig(varargin)
%EXPORT_FIG  Exports figures in a publication-quality format
%
% Examples:
%   imageData = export_fig
%   [imageData, alpha] = export_fig
%   export_fig filename
%   export_fig filename -format1 -format2
%   export_fig ... -nocrop
%   export_fig ... -c[<val>,<val>,<val>,<val>]
%   export_fig ... -transparent
%   export_fig ... -native
%   export_fig ... -m<val>
%   export_fig ... -r<val>
%   export_fig ... -a<val>
%   export_fig ... -q<val>
%   export_fig ... -p<val>
%   export_fig ... -d<gs_option>
%   export_fig ... -depsc
%   export_fig ... -<renderer>
%   export_fig ... -<colorspace>
%   export_fig ... -append
%   export_fig ... -bookmark
%   export_fig ... -clipboard
%   export_fig ... -update
%   export_fig ... -nofontswap
%   export_fig(..., handle)
%
% This function saves a figure or single axes to one or more vector and/or
% bitmap file formats, and/or outputs a rasterized version to the workspace,
% with the following properties:
%   - Figure/axes reproduced as it appears on screen
%   - Cropped borders (optional)
%   - Embedded fonts (vector formats)
%   - Improved line and grid line styles
%   - Anti-aliased graphics (bitmap formats)
%   - Render images at native resolution (optional for bitmap formats)
%   - Transparent background supported (pdf, eps, png, tif)
%   - Semi-transparent patch objects supported (png & tif only)
%   - RGB, CMYK or grayscale output (CMYK only with pdf, eps, tiff)
%   - Variable image compression, including lossless (pdf, eps, jpg)
%   - Optionally append to file (pdf, tiff)
%   - Vector formats: pdf, eps
%   - Bitmap formats: png, tiff, jpg, bmp, export to workspace
%
% This function is especially suited to exporting figures for use in
% publications and presentations, because of the high quality and
% portability of media produced.
%
% Note that the background color and figure dimensions are reproduced
% (the latter approximately, and ignoring cropping & magnification) in the
% output file. For transparent background (and semi-transparent patch
% objects), use the -transparent option or set the figure 'Color' property
% to 'none'. To make axes transparent set the axes 'Color' property to
% 'none'. PDF, EPS, TIF & PNG are the only formats that support a transparent
% background; only TIF & PNG formats support transparency of patch objects.
%
% The choice of renderer (opengl, zbuffer or painters) has a large impact
% on the quality of output. The default value (opengl for bitmaps, painters
% for vector formats) generally gives good results, but if you aren't
% satisfied then try another renderer.  Notes: 1) For vector formats (EPS,
% PDF), only painters generates vector graphics. 2) For bitmaps, only
% opengl can render transparent patch objects correctly. 3) For bitmaps,
% only painters will correctly scale line dash and dot lengths when
% magnifying or anti-aliasing. 4) Fonts may be substitued with Courier when
% using painters.
%
% When exporting to vector format (PDF & EPS) and bitmap format using the
% painters renderer, this function requires that ghostscript is installed
% on your system. You can download this from:
%   http://www.ghostscript.com
% When exporting to eps it additionally requires pdftops, from the Xpdf
% suite of functions. You can download this from:
%   http://www.foolabs.com/xpdf
%
% Inputs:
%   filename - string containing the name (optionally including full or
%              relative path) of the file the figure is to be saved as. If
%              a path is not specified, the figure is saved in the current
%              directory. If no name and no output arguments are specified,
%              the default name, 'export_fig_out', is used. If neither a
%              file extension nor a format are specified, a ".png" is added
%              and the figure saved in that format.
%   -format1, -format2, etc. - strings containing the extensions of the
%                              file formats the figure is to be saved as.
%                              Valid options are: '-pdf', '-eps', '-png',
%                              '-tif', '-jpg' and '-bmp'. All combinations
%                              of formats are valid.
%   -nocrop - option indicating that the borders of the output are not to
%             be cropped.
%   -c[<val>,<val>,<val>,<val>] - option indicating crop amounts. Must be
%             a 4-element vector of numeric values: [top,right,bottom,left]
%             where NaN/Inf indicate auto-cropping, 0 means no cropping,
%             and any other value mean cropping in pixel amounts.
%   -transparent - option indicating that the figure background is to be
%                  made transparent (png, pdf, tif and eps output only).
%   -m<val> - option where val indicates the factor to magnify the
%             on-screen figure pixel dimensions by when generating bitmap
%             outputs (does not affect vector formats). Default: '-m1'.
%   -r<val> - option val indicates the resolution (in pixels per inch) to
%             export bitmap and vector outputs at, keeping the dimensions
%             of the on-screen figure. Default: '-r864' (for vector output
%             only). Note that the -m option overides the -r option for
%             bitmap outputs only.
%   -native - option indicating that the output resolution (when outputting
%             a bitmap format) should be such that the vertical resolution
%             of the first suitable image found in the figure is at the
%             native resolution of that image. To specify a particular
%             image to use, give it the tag 'export_fig_native'. Notes:
%             This overrides any value set with the -m and -r options. It
%             also assumes that the image is displayed front-to-parallel
%             with the screen. The output resolution is approximate and
%             should not be relied upon. Anti-aliasing can have adverse
%             effects on image quality (disable with the -a1 option).
%   -a1, -a2, -a3, -a4 - option indicating the amount of anti-aliasing to
%                        use for bitmap outputs. '-a1' means no anti-
%                        aliasing; '-a4' is the maximum amount (default).
%   -<renderer> - option to force a particular renderer (painters, opengl or
%                 zbuffer). Default value: opengl for bitmap formats or
%                 figures with patches and/or transparent annotations;
%                 painters for vector formats without patches/transparencies.
%   -<colorspace> - option indicating which colorspace color figures should
%                   be saved in: RGB (default), CMYK or gray. CMYK is only
%                   supported in pdf, eps and tiff output.
%   -q<val> - option to vary bitmap image quality (in pdf, eps and jpg
%             files only).  Larger val, in the range 0-100, gives higher
%             quality/lower compression. val > 100 gives lossless
%             compression. Default: '-q95' for jpg, ghostscript prepress
%             default for pdf & eps. Note: lossless compression can
%             sometimes give a smaller file size than the default lossy
%             compression, depending on the type of images.
%   -p<val> - option to pad a border of width val to exported files, where
%             val is either a relative size with respect to cropped image
%             size (i.e. p=0.01 adds a 1% border). For EPS & PDF formats,
%             val can also be integer in units of 1/72" points (abs(val)>1).
%             val can be positive (padding) or negative (extra cropping).
%             If used, the -nocrop flag will be ignored, i.e. the image will
%             always be cropped and then padded. Default: 0 (i.e. no padding).
%   -append - option indicating that if the file (pdfs only) already
%             exists, the figure is to be appended as a new page, instead
%             of being overwritten (default).
%   -bookmark - option to indicate that a bookmark with the name of the
%               figure is to be created in the output file (pdf only).
%   -clipboard - option to save output as an image on the system clipboard.
%                Note: background transparency is not preserved in clipboard
%   -d<gs_option> - option to indicate a ghostscript setting. For example,
%                   -dMaxBitmap=0 or -dNoOutputFonts (Ghostscript 9.15+).
%   -depsc -  option to use EPS level-3 rather than the default level-2 print
%             device. This solves some bugs with Matlab's default -depsc2 device
%             such as discolored subplot lines on images (vector formats only).
%   -update - option to download and install the latest version of export_fig
%   -nofontswap - option to avoid font swapping. Font swapping is automatically
%             done in vector formats (only): 11 standard Matlab fonts are
%             replaced by the original figure fonts. This option prevents this.
%   handle -  The handle of the figure, axes or uipanels (can be an array of
%             handles, but the objects must be in the same figure) to be
%             saved. Default: gcf.
%
% Outputs:
%   imageData - MxNxC uint8 image array of the exported image.
%   alpha     - MxN single array of alphamatte values in the range [0,1],
%               for the case when the background is transparent.
%
%   Some helpful examples and tips can be found at:
%      https://github.com/altmany/export_fig
%
%   See also PRINT, SAVEAS, ScreenCapture (on the Matlab File Exchange)

%{
% Copyright (C) Oliver Woodford 2008-2014, Yair Altman 2015-

% The idea of using ghostscript is inspired by Peder Axensten's SAVEFIG
% (fex id: 10889) which is itself inspired by EPS2PDF (fex id: 5782).
% The idea for using pdftops came from the MATLAB newsgroup (id: 168171).
% The idea of editing the EPS file to change line styles comes from Jiro
% Doke's FIXPSLINESTYLE (fex id: 17928).
% The idea of changing dash length with line width came from comments on
% fex id: 5743, but the implementation is mine :)
% The idea of anti-aliasing bitmaps came from Anders Brun's MYAA (fex id:
% 20979).
% The idea of appending figures in pdfs came from Matt C in comments on the
% FEX (id: 23629)

% Thanks to Roland Martin for pointing out the colour MATLAB
% bug/feature with colorbar axes and transparent backgrounds.
% Thanks also to Andrew Matthews for describing a bug to do with the figure
% size changing in -nodisplay mode. I couldn't reproduce it, but included a
% fix anyway.
% Thanks to Tammy Threadgill for reporting a bug where an axes is not
% isolated from gui objects.
%}
%{
% 23/02/12: Ensure that axes limits don't change during printing
% 14/03/12: Fix bug in fixing the axes limits (thanks to Tobias Lamour for reporting it).
% 02/05/12: Incorporate patch of Petr Nechaev (many thanks), enabling bookmarking of figures in pdf files.
% 09/05/12: Incorporate patch of Arcelia Arrieta (many thanks), to keep tick marks fixed.
% 12/12/12: Add support for isolating uipanels. Thanks to michael for suggesting it.
% 25/09/13: Add support for changing resolution in vector formats. Thanks to Jan Jaap Meijer for suggesting it.
% 07/05/14: Add support for '~' at start of path. Thanks to Sally Warner for suggesting it.
% 24/02/15: Fix Matlab R2014b bug (issue #34): plot markers are not displayed when ZLimMode='manual'
% 25/02/15: Fix issue #4 (using HG2 on R2014a and earlier)
% 25/02/15: Fix issue #21 (bold TeX axes labels/titles in R2014b)
% 26/02/15: If temp dir is not writable, use the user-specified folder for temporary EPS/PDF files (Javier Paredes)
% 27/02/15: Modified repository URL from github.com/ojwoodford to /altmany
%           Indented main function
%           Added top-level try-catch block to display useful workarounds
% 28/02/15: Enable users to specify optional ghostscript options (issue #36)
% 06/03/15: Improved image padding & cropping thanks to Oscar Hartogensis
% 26/03/15: Fixed issue #49 (bug with transparent grayscale images); fixed out-of-memory issue
% 26/03/15: Fixed issue #42: non-normalized annotations on HG1
% 26/03/15: Fixed issue #46: Ghostscript crash if figure units <> pixels
% 27/03/15: Fixed issue #39: bad export of transparent annotations/patches
% 28/03/15: Fixed issue #50: error on some Matlab versions with the fix for issue #42
% 29/03/15: Fixed issue #33: bugs in Matlab's print() function with -cmyk
% 29/03/15: Improved processing of input args (accept space between param name & value, related to issue #51)
% 30/03/15: When exporting *.fig files, then saveas *.fig if figure is open, otherwise export the specified fig file
% 30/03/15: Fixed edge case bug introduced yesterday (commit #ae1755bd2e11dc4e99b95a7681f6e211b3fa9358)
% 09/04/15: Consolidated header comment sections; initialize output vars only if requested (nargout>0)
% 14/04/15: Workaround for issue #45: lines in image subplots are exported in invalid color
% 15/04/15: Fixed edge-case in parsing input parameters; fixed help section to show the -depsc option (issue #45)
% 21/04/15: Bug fix: Ghostscript croaks on % chars in output PDF file (reported by Sven on FEX page, 15-Jul-2014)
% 22/04/15: Bug fix: Pdftops croaks on relative paths (reported by Tintin Milou on FEX page, 19-Jan-2015)
% 04/05/15: Merged fix #63 (Kevin Mattheus Moerman): prevent tick-label changes during export
% 07/05/15: Partial fix for issue #65: PDF export used painters rather than opengl renderer (thanks Nguyenr)
% 08/05/15: Fixed issue #65: bad PDF append since commit #e9f3cdf 21/04/15 (thanks Robert Nguyen)
% 12/05/15: Fixed issue #67: exponent labels cropped in export, since fix #63 (04/05/15)
% 28/05/15: Fixed issue #69: set non-bold label font only if the string contains symbols (\beta etc.), followup to issue #21
% 29/05/15: Added informative error message in case user requested SVG output (issue #72)
% 09/06/15: Fixed issue #58: -transparent removed anti-aliasing when exporting to PNG
% 19/06/15: Added -update option to download and install the latest version of export_fig
% 07/07/15: Added -nofontswap option to avoid font-swapping in EPS/PDF
% 16/07/15: Fixed problem with anti-aliasing on old Matlab releases
% 11/09/15: Fixed issue #103: magnification must never become negative; also fixed reported error msg in parsing input params
% 26/09/15: Alert if trying to export transparent patches/areas to non-PNG outputs (issue #108)
% 04/10/15: Do not suggest workarounds for certain errors that have already been handled previously
% 01/11/15: Fixed issue #112: use same renderer in print2eps as export_fig (thanks to Jes�s Pestana Puerta)
% 10/11/15: Custom GS installation webpage for MacOS. Thanks to Andy Hueni via FEX
% 19/11/15: Fixed clipboard export in R2015b (thanks to Dan K via FEX)
% 21/02/16: Added -c option for indicating specific crop amounts (idea by Cedric Noordam on FEX)
% 08/05/16: Added message about possible error reason when groot.Units~=pixels (issue #149)
% 17/05/16: Fixed case of image YData containing more than 2 elements (issue #151)
% 08/08/16: Enabled exporting transparency to TIF, in addition to PNG/PDF (issue #168)
% 11/12/16: Added alert in case of error creating output PDF/EPS file (issue #179)
% 13/12/16: Minor fix to the commit for issue #179 from 2 days ago
%}

    if nargout
        [imageData, alpha] = deal([]);
    end
    hadError = false;
    displaySuggestedWorkarounds = true;

    % Ensure the figure is rendered correctly _now_ so that properties like axes limits are up-to-date
    drawnow;
    pause(0.05);  % this solves timing issues with Java Swing's EDT (http://undocumentedmatlab.com/blog/solving-a-matlab-hang-problem)

    % Parse the input arguments
    fig = get(0, 'CurrentFigure');
    [fig, options] = parse_args(nargout, fig, varargin{:});

    % Ensure that we have a figure handle
    if isequal(fig,-1)
        return;  % silent bail-out
    elseif isempty(fig)
        error('No figure found');
    end

    % Isolate the subplot, if it is one
    cls = all(ismember(get(fig, 'Type'), {'axes', 'uipanel'}));
    if cls
        % Given handles of one or more axes, so isolate them from the rest
        fig = isolate_axes(fig);
    else
        % Check we have a figure
        if ~isequal(get(fig, 'Type'), 'figure')
            error('Handle must be that of a figure, axes or uipanel');
        end
        % Get the old InvertHardcopy mode
        old_mode = get(fig, 'InvertHardcopy');
    end

    % Hack the font units where necessary (due to a font rendering bug in print?).
    % This may not work perfectly in all cases.
    % Also it can change the figure layout if reverted, so use a copy.
    magnify = options.magnify * options.aa_factor;
    if isbitmap(options) && magnify ~= 1
        fontu = findall(fig, 'FontUnits', 'normalized');
        if ~isempty(fontu)
            % Some normalized font units found
            if ~cls
                fig = copyfig(fig);
                set(fig, 'Visible', 'off');
                fontu = findall(fig, 'FontUnits', 'normalized');
                cls = true;
            end
            set(fontu, 'FontUnits', 'points');
        end
    end

    try
        % MATLAB "feature": axes limits and tick marks can change when printing
        Hlims = findall(fig, 'Type', 'axes');
        if ~cls
            % Record the old axes limit and tick modes
            Xlims = make_cell(get(Hlims, 'XLimMode'));
            Ylims = make_cell(get(Hlims, 'YLimMode'));
            Zlims = make_cell(get(Hlims, 'ZLimMode'));
            Xtick = make_cell(get(Hlims, 'XTickMode'));
            Ytick = make_cell(get(Hlims, 'YTickMode'));
            Ztick = make_cell(get(Hlims, 'ZTickMode'));
            Xlabel = make_cell(get(Hlims, 'XTickLabelMode')); 
            Ylabel = make_cell(get(Hlims, 'YTickLabelMode')); 
            Zlabel = make_cell(get(Hlims, 'ZTickLabelMode')); 
        end

        % Set all axes limit and tick modes to manual, so the limits and ticks can't change
        % Fix Matlab R2014b bug (issue #34): plot markers are not displayed when ZLimMode='manual'
        set(Hlims, 'XLimMode', 'manual', 'YLimMode', 'manual');
        set_tick_mode(Hlims, 'X');
        set_tick_mode(Hlims, 'Y');
        if ~using_hg2(fig)
            set(Hlims,'ZLimMode', 'manual');
            set_tick_mode(Hlims, 'Z');
        end
    catch
        % ignore - fix issue #4 (using HG2 on R2014a and earlier)
    end

    % Fix issue #21 (bold TeX axes labels/titles in R2014b when exporting to EPS/PDF)
    try
        if using_hg2(fig) && isvector(options)
            % Set the FontWeight of axes labels/titles to 'normal'
            % Fix issue #69: set non-bold font only if the string contains symbols (\beta etc.)
            texLabels = findall(fig, 'type','text', 'FontWeight','bold');
            symbolIdx = ~cellfun('isempty',strfind({texLabels.String},'\'));
            set(texLabels(symbolIdx), 'FontWeight','normal');
        end
    catch
        % ignore
    end

    % Fix issue #42: non-normalized annotations on HG1 (internal Matlab bug)
    annotationHandles = [];
    try
        if ~using_hg2(fig)
            annotationHandles = findall(fig,'Type','hggroup','-and','-property','Units','-and','-not','Units','norm');
            try  % suggested by Jes�s Pestana Puerta (jespestana) 30/9/2015
                originalUnits = get(annotationHandles,'Units');
                set(annotationHandles,'Units','norm');
            catch
            end
        end
    catch
        % should never happen, but ignore in any case - issue #50
    end

    % Fix issue #46: Ghostscript crash if figure units <> pixels
    oldFigUnits = get(fig,'Units');
    set(fig,'Units','pixels');

    % Set to print exactly what is there
    set(fig, 'InvertHardcopy', 'off');
    % Set the renderer
    switch options.renderer
        case 1
            renderer = '-opengl';
        case 2
            renderer = '-zbuffer';
        case 3
            renderer = '-painters';
        otherwise
            renderer = '-opengl'; % Default for bitmaps
    end

    % Handle transparent patches
    hasTransparency = ~isempty(findall(fig,'-property','FaceAlpha','-and','-not','FaceAlpha',1));
    hasPatches      = ~isempty(findall(fig,'type','patch'));
    if hasTransparency
        % Alert if trying to export transparent patches/areas to non-supported outputs (issue #108)
        % http://www.mathworks.com/matlabcentral/answers/265265-can-export_fig-or-else-draw-vector-graphics-with-transparent-surfaces
        % TODO - use transparency when exporting to PDF by not passing via print2eps
        msg = 'export_fig currently supports transparent patches/areas only in PNG output. ';
        if options.pdf
            warning('export_fig:transparency', '%s\nTo export transparent patches/areas to PDF, use the print command:\n print(gcf, ''-dpdf'', ''%s.pdf'');', msg, options.name);
        elseif ~options.png && ~options.tif  % issue #168
            warning('export_fig:transparency', '%s\nTo export the transparency correctly, try using the ScreenCapture utility on the Matlab File Exchange: http://bit.ly/1QFrBip', msg);
        end
    end

    try
        % Do the bitmap formats first
        if isbitmap(options)
            if abs(options.bb_padding) > 1
                displaySuggestedWorkarounds = false;
                error('For bitmap output (png,jpg,tif,bmp) the padding value (-p) must be between -1<p<1')
            end
            % Get the background colour
            if options.transparent && (options.png || options.alpha)
                % Get out an alpha channel
                % MATLAB "feature": black colorbar axes can change to white and vice versa!
                hCB = findall(fig, 'Type','axes', 'Tag','Colorbar');
                if isempty(hCB)
                    yCol = [];
                    xCol = [];
                else
                    yCol = get(hCB, 'YColor');
                    xCol = get(hCB, 'XColor');
                    if iscell(yCol)
                        yCol = cell2mat(yCol);
                        xCol = cell2mat(xCol);
                    end
                    yCol = sum(yCol, 2);
                    xCol = sum(xCol, 2);
                end
                % MATLAB "feature": apparently figure size can change when changing
                % colour in -nodisplay mode
                pos = get(fig, 'Position');
                % Set the background colour to black, and set size in case it was
                % changed internally
                tcol = get(fig, 'Color');
                set(fig, 'Color', 'k', 'Position', pos);
                % Correct the colorbar axes colours
                set(hCB(yCol==0), 'YColor', [0 0 0]);
                set(hCB(xCol==0), 'XColor', [0 0 0]);

                % The following code might cause out-of-memory errors
                try
                    % Print large version to array
                    B = print2array(fig, magnify, renderer);
                    % Downscale the image
                    B = downsize(single(B), options.aa_factor);
                catch
                    % This is more conservative in memory, but kills transparency (issue #58)
                    B = single(print2array(fig, magnify/options.aa_factor, renderer));
                end

                % Set background to white (and set size)
                set(fig, 'Color', 'w', 'Position', pos);
                % Correct the colorbar axes colours
                set(hCB(yCol==3), 'YColor', [1 1 1]);
                set(hCB(xCol==3), 'XColor', [1 1 1]);

                % The following code might cause out-of-memory errors
                try
                    % Print large version to array
                    A = print2array(fig, magnify, renderer);
                    % Downscale the image
                    A = downsize(single(A), options.aa_factor);
                catch
                    % This is more conservative in memory, but kills transparency (issue #58)
                    A = single(print2array(fig, magnify/options.aa_factor, renderer));
                end

                % Set the background colour (and size) back to normal
                set(fig, 'Color', tcol, 'Position', pos);
                % Compute the alpha map
                alpha = round(sum(B - A, 3)) / (255 * 3) + 1;
                A = alpha;
                A(A==0) = 1;
                A = B ./ A(:,:,[1 1 1]);
                clear B
                % Convert to greyscale
                if options.colourspace == 2
                    A = rgb2grey(A);
                end
                A = uint8(A);
                % Crop the background
                if options.crop
                    %[alpha, v] = crop_borders(alpha, 0, 1, options.crop_amounts);
                    %A = A(v(1):v(2),v(3):v(4),:);
                    [alpha, vA, vB] = crop_borders(alpha, 0, options.bb_padding, options.crop_amounts);
                    if ~any(isnan(vB)) % positive padding
                        B = repmat(uint8(zeros(1,1,size(A,3))),size(alpha));
                        B(vB(1):vB(2), vB(3):vB(4), :) = A(vA(1):vA(2), vA(3):vA(4), :); % ADDED BY OH
                        A = B;
                    else  % negative padding
                        A = A(vA(1):vA(2), vA(3):vA(4), :);
                    end
                end
                if options.png
                    % Compute the resolution
                    res = options.magnify * get(0, 'ScreenPixelsPerInch') / 25.4e-3;
                    % Save the png
                    imwrite(A, [options.name '.png'], 'Alpha', double(alpha), 'ResolutionUnit', 'meter', 'XResolution', res, 'YResolution', res);
                    % Clear the png bit
                    options.png = false;
                end
                % Return only one channel for greyscale
                if isbitmap(options)
                    A = check_greyscale(A);
                end
                if options.alpha
                    % Store the image
                    imageData = A;
                    % Clear the alpha bit
                    options.alpha = false;
                end
                % Get the non-alpha image
                if isbitmap(options)
                    alph = alpha(:,:,ones(1, size(A, 3)));
                    A = uint8(single(A) .* alph + 255 * (1 - alph));
                    clear alph
                end
                if options.im
                    % Store the new image
                    imageData = A;
                end
            else
                % Print large version to array
                if options.transparent
                    % MATLAB "feature": apparently figure size can change when changing
                    % colour in -nodisplay mode
                    pos = get(fig, 'Position');
                    tcol = get(fig, 'Color');
                    set(fig, 'Color', 'w', 'Position', pos);
                    A = print2array(fig, magnify, renderer);
                    set(fig, 'Color', tcol, 'Position', pos);
                    tcol = 255;
                else
                    [A, tcol] = print2array(fig, magnify, renderer);
                end
                % Crop the background
                if options.crop
                    A = crop_borders(A, tcol, options.bb_padding, options.crop_amounts);
                end
                % Downscale the image
                A = downsize(A, options.aa_factor);
                if options.colourspace == 2
                    % Convert to greyscale
                    A = rgb2grey(A);
                else
                    % Return only one channel for greyscale
                    A = check_greyscale(A);
                end
                % Outputs
                if options.im
                    imageData = A;
                end
                if options.alpha
                    imageData = A;
                    alpha = zeros(size(A, 1), size(A, 2), 'single');
                end
            end
            % Save the images
            if options.png
                res = options.magnify * get(0, 'ScreenPixelsPerInch') / 25.4e-3;
                imwrite(A, [options.name '.png'], 'ResolutionUnit', 'meter', 'XResolution', res, 'YResolution', res);
            end
            if options.bmp
                imwrite(A, [options.name '.bmp']);
            end
            % Save jpeg with given quality
            if options.jpg
                quality = options.quality;
                if isempty(quality)
                    quality = 95;
                end
                if quality > 100
                    imwrite(A, [options.name '.jpg'], 'Mode', 'lossless');
                else
                    imwrite(A, [options.name '.jpg'], 'Quality', quality);
                end
            end
            % Save tif images in cmyk if wanted (and possible)
            if options.tif
                if options.colourspace == 1 && size(A, 3) == 3
                    A = double(255 - A);
                    K = min(A, [], 3);
                    K_ = 255 ./ max(255 - K, 1);
                    C = (A(:,:,1) - K) .* K_;
                    M = (A(:,:,2) - K) .* K_;
                    Y = (A(:,:,3) - K) .* K_;
                    A = uint8(cat(3, C, M, Y, K));
                    clear C M Y K K_
                end
                append_mode = {'overwrite', 'append'};
                imwrite(A, [options.name '.tif'], 'Resolution', options.magnify*get(0, 'ScreenPixelsPerInch'), 'WriteMode', append_mode{options.append+1});
            end
        end

        % Now do the vector formats
        if isvector(options)
            % Set the default renderer to painters
            if ~options.renderer
                if hasTransparency || hasPatches
                    % This is *MUCH* slower, but more accurate for patches and transparent annotations (issue #39)
                    renderer = '-opengl';
                else
                    renderer = '-painters';
                end
            end
            options.rendererStr = renderer;  % fix for issue #112
            % Generate some filenames
            tmp_nam = [tempname '.eps'];
            try
                % Ensure that the temp dir is writable (Javier Paredes 30/1/15)
                fid = fopen(tmp_nam,'w');
                fwrite(fid,1);
                fclose(fid);
                delete(tmp_nam);
                isTempDirOk = true;
            catch
                % Temp dir is not writable, so use the user-specified folder
                [dummy,fname,fext] = fileparts(tmp_nam); %#ok<ASGLU>
                fpath = fileparts(options.name);
                tmp_nam = fullfile(fpath,[fname fext]);
                isTempDirOk = false;
            end
            if isTempDirOk
                pdf_nam_tmp = [tempname '.pdf'];
            else
                pdf_nam_tmp = fullfile(fpath,[fname '.pdf']);
            end
            if options.pdf
                pdf_nam = [options.name '.pdf'];
                try copyfile(pdf_nam, pdf_nam_tmp, 'f'); catch, end  % fix for issue #65
            else
                pdf_nam = pdf_nam_tmp;
            end
            % Generate the options for print
            p2eArgs = {renderer, sprintf('-r%d', options.resolution)};
            if options.colourspace == 1  % CMYK
                % Issue #33: due to internal bugs in Matlab's print() function, we can't use its -cmyk option
                %p2eArgs{end+1} = '-cmyk';
            end
            if ~options.crop
                % Issue #56: due to internal bugs in Matlab's print() function, we can't use its internal cropping mechanism,
                % therefore we always use '-loose' (in print2eps.m) and do our own cropping (in crop_borders)
                %p2eArgs{end+1} = '-loose';
            end
            if any(strcmpi(varargin,'-depsc'))
                % Issue #45: lines in image subplots are exported in invalid color.
                % The workaround is to use the -depsc parameter instead of the default -depsc2
                p2eArgs{end+1} = '-depsc';
            end
            try
                % Generate an eps
                print2eps(tmp_nam, fig, options, p2eArgs{:});
                % Remove the background, if desired
                if options.transparent && ~isequal(get(fig, 'Color'), 'none')
                    eps_remove_background(tmp_nam, 1 + using_hg2(fig));
                end
                % Fix colorspace to CMYK, if requested (workaround for issue #33)
                if options.colourspace == 1  % CMYK
                    % Issue #33: due to internal bugs in Matlab's print() function, we can't use its -cmyk option
                    change_rgb_to_cmyk(tmp_nam);
                end
                % Add a bookmark to the PDF if desired
                if options.bookmark
                    fig_nam = get(fig, 'Name');
                    if isempty(fig_nam)
                        warning('export_fig:EmptyBookmark', 'Bookmark requested for figure with no name. Bookmark will be empty.');
                    end
                    add_bookmark(tmp_nam, fig_nam);
                end
                % Generate a pdf
                eps2pdf(tmp_nam, pdf_nam_tmp, 1, options.append, options.colourspace==2, options.quality, options.gs_options);
                % Ghostscript croaks on % chars in the output PDF file, so use tempname and then rename the file
                try
                    % Rename the file (except if it is already the same)
                    % Abbie K's comment on the commit for issue #179 (#commitcomment-20173476)
                    if ~isequal(pdf_nam_tmp, pdf_nam)
                        movefile(pdf_nam_tmp, pdf_nam, 'f');
                    end
                catch
                    % Alert in case of error creating output PDF/EPS file (issue #179)
                    if exist(pdf_nam_tmp, 'file')
                        error(['Could not create ' pdf_nam ' - perhaps the folder does not exist, or you do not have write permissions']);
                    else
                        error('Could not generate the intermediary EPS file.');
                    end
                end
            catch ex
                % Delete the eps
                delete(tmp_nam);
                rethrow(ex);
            end
            % Delete the eps
            delete(tmp_nam);
            if options.eps
                try
                    % Generate an eps from the pdf
                    % since pdftops can't handle relative paths (e.g., '..\'), use a temp file
                    eps_nam_tmp = strrep(pdf_nam_tmp,'.pdf','.eps');
                    pdf2eps(pdf_nam, eps_nam_tmp);
                    movefile(eps_nam_tmp,  [options.name '.eps'], 'f');
                catch ex
                    if ~options.pdf
                        % Delete the pdf
                        delete(pdf_nam);
                    end
                    try delete(eps_nam_tmp); catch, end
                    rethrow(ex);
                end
                if ~options.pdf
                    % Delete the pdf
                    delete(pdf_nam);
                end
            end
        end

        % Revert the figure or close it (if requested)
        if cls || options.closeFig
            % Close the created figure
            close(fig);
        else
            % Reset the hardcopy mode
            set(fig, 'InvertHardcopy', old_mode);
            % Reset the axes limit and tick modes
            for a = 1:numel(Hlims)
                try
                    set(Hlims(a), 'XLimMode', Xlims{a}, 'YLimMode', Ylims{a}, 'ZLimMode', Zlims{a},... 
                                  'XTickMode', Xtick{a}, 'YTickMode', Ytick{a}, 'ZTickMode', Ztick{a},...
                                  'XTickLabelMode', Xlabel{a}, 'YTickLabelMode', Ylabel{a}, 'ZTickLabelMode', Zlabel{a}); 
                catch
                    % ignore - fix issue #4 (using HG2 on R2014a and earlier)
                end
            end
            % Revert the tex-labels font weights
            try set(texLabels, 'FontWeight','bold'); catch, end
            % Revert annotation units
            for handleIdx = 1 : numel(annotationHandles)
                try
                    oldUnits = originalUnits{handleIdx};
                catch
                    oldUnits = originalUnits;
                end
                try set(annotationHandles(handleIdx),'Units',oldUnits); catch, end
            end
            % Revert figure units
            set(fig,'Units',oldFigUnits);
        end

        % Output to clipboard (if requested)
        if options.clipboard
            % Delete the output file if unchanged from the default name ('export_fig_out.png')
            if strcmpi(options.name,'export_fig_out')
                try
                    fileInfo = dir('export_fig_out.png');
                    if ~isempty(fileInfo)
                        timediff = now - fileInfo.datenum;
                        ONE_SEC = 1/24/60/60;
                        if timediff < ONE_SEC
                            delete('export_fig_out.png');
                        end
                    end
                catch
                    % never mind...
                end
            end

            % Save the image in the system clipboard
            % credit: Jiro Doke's IMCLIPBOARD: http://www.mathworks.com/matlabcentral/fileexchange/28708-imclipboard
            try
                error(javachk('awt', 'export_fig -clipboard output'));
            catch
                warning('export_fig -clipboard output failed: requires Java to work');
                return;
            end
            try
                % Import necessary Java classes
                import java.awt.Toolkit
                import java.awt.image.BufferedImage
                import java.awt.datatransfer.DataFlavor

                % Get System Clipboard object (java.awt.Toolkit)
                cb = Toolkit.getDefaultToolkit.getSystemClipboard();

                % Add java class (ImageSelection) to the path
                if ~exist('ImageSelection', 'class')
                    javaaddpath(fileparts(which(mfilename)), '-end');
                end

                % Get image size
                ht = size(imageData, 1);
                wd = size(imageData, 2);

                % Convert to Blue-Green-Red format
                try
                    imageData2 = imageData(:, :, [3 2 1]);
                catch
                    % Probably gray-scaled image (2D, without the 3rd [RGB] dimension)
                    imageData2 = imageData(:, :, [1 1 1]);
                end

                % Convert to 3xWxH format
                imageData2 = permute(imageData2, [3, 2, 1]);

                % Append Alpha data (unused - transparency is not supported in clipboard copy)
                alphaData2 = uint8(permute(255*alpha,[3,2,1])); %=255*ones(1,wd,ht,'uint8')
                imageData2 = cat(1, imageData2, alphaData2);

                % Create image buffer
                imBuffer = BufferedImage(wd, ht, BufferedImage.TYPE_INT_RGB);
                imBuffer.setRGB(0, 0, wd, ht, typecast(imageData2(:), 'int32'), 0, wd);

                % Create ImageSelection object from the image buffer
                imSelection = ImageSelection(imBuffer);

                % Set clipboard content to the image
                cb.setContents(imSelection, []);
            catch
                warning('export_fig -clipboard output failed: %s', lasterr); %#ok<LERR>
            end
        end

        % Don't output the data to console unless requested
        if ~nargout
            clear imageData alpha
        end
    catch err
        % Display possible workarounds before the error message
        if displaySuggestedWorkarounds && ~strcmpi(err.message,'export_fig error')
            if ~hadError,  fprintf(2, 'export_fig error. ');  end
            fprintf(2, 'Please ensure:\n');
            fprintf(2, '  that you are using the <a href="https://github.com/altmany/export_fig/archive/master.zip">latest version</a> of export_fig\n');
            if ismac
                fprintf(2, '  and that you have <a href="http://pages.uoregon.edu/koch">Ghostscript</a> installed\n');
            else
                fprintf(2, '  and that you have <a href="http://www.ghostscript.com">Ghostscript</a> installed\n');
            end
            try
                if options.eps
                    fprintf(2, '  and that you have <a href="http://www.foolabs.com/xpdf">pdftops</a> installed\n');
                end
            catch
                % ignore - probably an error in parse_args
            end
            fprintf(2, '  and that you do not have <a href="matlab:which export_fig -all">multiple versions</a> of export_fig installed by mistake\n');
            fprintf(2, '  and that you did not made a mistake in the <a href="matlab:help export_fig">expected input arguments</a>\n');
            try
                % Alert per issue #149
                if ~strncmpi(get(0,'Units'),'pixel',5)
                    fprintf(2, '  or try to set groot''s Units property back to its default value of ''pixels'' (<a href="matlab:web(''https://github.com/altmany/export_fig/issues/149'',''-browser'');">details</a>)\n');
                end
            catch
                % ignore - maybe an old MAtlab release
            end
            fprintf(2, '\nIf the problem persists, then please <a href="https://github.com/altmany/export_fig/issues">report a new issue</a>.\n\n');
        end
        rethrow(err)
    end
end

function options = default_options()
    % Default options used by export_fig
    options = struct(...
        'name',         'export_fig_out', ...
        'crop',         true, ...
        'crop_amounts', nan(1,4), ...  % auto-crop all 4 image sides
        'transparent',  false, ...
        'renderer',     0, ...         % 0: default, 1: OpenGL, 2: ZBuffer, 3: Painters
        'pdf',          false, ...
        'eps',          false, ...
        'png',          false, ...
        'tif',          false, ...
        'jpg',          false, ...
        'bmp',          false, ...
        'clipboard',    false, ...
        'colourspace',  0, ...         % 0: RGB/gray, 1: CMYK, 2: gray
        'append',       false, ...
        'im',           false, ...
        'alpha',        false, ...
        'aa_factor',    0, ...
        'bb_padding',   0, ...
        'magnify',      [], ...
        'resolution',   [], ...
        'bookmark',     false, ...
        'closeFig',     false, ...
        'quality',      [], ...
        'update',       false, ...
        'fontswap',     true, ...
        'gs_options',   {{}});
end

function [fig, options] = parse_args(nout, fig, varargin)
    % Parse the input arguments

    % Set the defaults
    native = false; % Set resolution to native of an image
    options = default_options();
    options.im =    (nout == 1);  % user requested imageData output
    options.alpha = (nout == 2);  % user requested alpha output

    % Go through the other arguments
    skipNext = false;
    for a = 1:nargin-2
        if skipNext
            skipNext = false;
            continue;
        end
        if all(ishandle(varargin{a}))
            fig = varargin{a};
        elseif ischar(varargin{a}) && ~isempty(varargin{a})
            if varargin{a}(1) == '-'
                switch lower(varargin{a}(2:end))
                    case 'nocrop'
                        options.crop = false;
                        options.crop_amounts = [0,0,0,0];
                    case {'trans', 'transparent'}
                        options.transparent = true;
                    case 'opengl'
                        options.renderer = 1;
                    case 'zbuffer'
                        options.renderer = 2;
                    case 'painters'
                        options.renderer = 3;
                    case 'pdf'
                        options.pdf = true;
                    case 'eps'
                        options.eps = true;
                    case 'png'
                        options.png = true;
                    case {'tif', 'tiff'}
                        options.tif = true;
                    case {'jpg', 'jpeg'}
                        options.jpg = true;
                    case 'bmp'
                        options.bmp = true;
                    case 'rgb'
                        options.colourspace = 0;
                    case 'cmyk'
                        options.colourspace = 1;
                    case {'gray', 'grey'}
                        options.colourspace = 2;
                    case {'a1', 'a2', 'a3', 'a4'}
                        options.aa_factor = str2double(varargin{a}(3));
                    case 'append'
                        options.append = true;
                    case 'bookmark'
                        options.bookmark = true;
                    case 'native'
                        native = true;
                    case 'clipboard'
                        options.clipboard = true;
                        options.im = true;
                        options.alpha = true;
                    case 'svg'
                        msg = ['SVG output is not supported by export_fig. Use one of the following alternatives:\n' ...
                               '  1. saveas(gcf,''filename.svg'')\n' ...
                               '  2. plot2svg utility: http://github.com/jschwizer99/plot2svg\n' ...
                               '  3. export_fig to EPS/PDF, then convert to SVG using generic (non-Matlab) tools\n'];
                        error(sprintf(msg)); %#ok<SPERR>
                    case 'update'
                        % Download the latest version of export_fig into the export_fig folder
                        try
                            zipFileName = 'https://github.com/altmany/export_fig/archive/master.zip';
                            folderName = fileparts(which(mfilename('fullpath')));
                            targetFileName = fullfile(folderName, datestr(now,'yyyy-mm-dd.zip'));
                            urlwrite(zipFileName,targetFileName);
                        catch
                            error('Could not download %s into %s\n',zipFileName,targetFileName);
                        end

                        % Unzip the downloaded zip file in the export_fig folder
                        try
                            unzip(targetFileName,folderName);
                        catch
                            error('Could not unzip %s\n',targetFileName);
                        end
                    case 'nofontswap'
                        options.fontswap = false;
                    otherwise
                        try
                            wasError = false;
                            if strcmpi(varargin{a}(1:2),'-d')
                                varargin{a}(2) = 'd';  % ensure lowercase 'd'
                                options.gs_options{end+1} = varargin{a};
                            elseif strcmpi(varargin{a}(1:2),'-c')
                                if numel(varargin{a})==2
                                    skipNext = true;
                                    vals = str2num(varargin{a+1}); %#ok<ST2NM>
                                else
                                    vals = str2num(varargin{a}(3:end)); %#ok<ST2NM>
                                end
                                if numel(vals)~=4
                                    wasError = true;
                                    error('option -c cannot be parsed: must be a 4-element numeric vector');
                                end
                                options.crop_amounts = vals;
                                options.crop = true;
                            else  % scalar parameter value
                                val = str2double(regexp(varargin{a}, '(?<=-(m|M|r|R|q|Q|p|P))-?\d*.?\d+', 'match'));
                                if isempty(val) || isnan(val)
                                    % Issue #51: improved processing of input args (accept space between param name & value)
                                    val = str2double(varargin{a+1});
                                    if isscalar(val) && ~isnan(val)
                                        skipNext = true;
                                    end
                                end
                                if ~isscalar(val) || isnan(val)
                                    wasError = true;
                                    error('option %s is not recognised or cannot be parsed', varargin{a});
                                end
                                switch lower(varargin{a}(2))
                                    case 'm'
                                        % Magnification may never be negative
                                        if val <= 0
                                            wasError = true;
                                            error('Bad magnification value: %g (must be positive)', val);
                                        end
                                        options.magnify = val;
                                    case 'r'
                                        options.resolution = val;
                                    case 'q'
                                        options.quality = max(val, 0);
                                    case 'p'
                                        options.bb_padding = val;
                                end
                            end
                        catch err
                            % We might have reached here by raising an intentional error
                            if wasError  % intentional raise
                                rethrow(err)
                            else  % unintentional
                                error(['Unrecognized export_fig input option: ''' varargin{a} '''']);
                            end
                        end
                end
            else
                [p, options.name, ext] = fileparts(varargin{a});
                if ~isempty(p)
                    options.name = [p filesep options.name];
                end
                switch lower(ext)
                    case {'.tif', '.tiff'}
                        options.tif = true;
                    case {'.jpg', '.jpeg'}
                        options.jpg = true;
                    case '.png'
                        options.png = true;
                    case '.bmp'
                        options.bmp = true;
                    case '.eps'
                        options.eps = true;
                    case '.pdf'
                        options.pdf = true;
                    case '.fig'
                        % If no open figure, then load the specified .fig file and continue
                        if isempty(fig)
                            fig = openfig(varargin{a},'invisible');
                            varargin{a} = fig;
                            options.closeFig = true;
                        else
                            % save the current figure as the specified .fig file and exit
                            saveas(fig(1),varargin{a});
                            fig = -1;
                            return
                        end
                    case '.svg'
                        msg = ['SVG output is not supported by export_fig. Use one of the following alternatives:\n' ...
                               '  1. saveas(gcf,''filename.svg'')\n' ...
                               '  2. plot2svg utility: http://github.com/jschwizer99/plot2svg\n' ...
                               '  3. export_fig to EPS/PDF, then convert to SVG using generic (non-Matlab) tools\n'];
                        error(sprintf(msg)); %#ok<SPERR>
                    otherwise
                        options.name = varargin{a};
                end
            end
        end
    end

    % Quick bail-out if no figure found
    if isempty(fig),  return;  end

    % Do border padding with repsect to a cropped image
    if options.bb_padding
        options.crop = true;
    end

    % Set default anti-aliasing now we know the renderer
    if options.aa_factor == 0
        try isAA = strcmp(get(ancestor(fig, 'figure'), 'GraphicsSmoothing'), 'on'); catch, isAA = false; end
        options.aa_factor = 1 + 2 * (~(using_hg2(fig) && isAA) | (options.renderer == 3));
    end

    % Convert user dir '~' to full path
    if numel(options.name) > 2 && options.name(1) == '~' && (options.name(2) == '/' || options.name(2) == '\')
        options.name = fullfile(char(java.lang.System.getProperty('user.home')), options.name(2:end));
    end

    % Compute the magnification and resolution
    if isempty(options.magnify)
        if isempty(options.resolution)
            options.magnify = 1;
            options.resolution = 864;
        else
            options.magnify = options.resolution ./ get(0, 'ScreenPixelsPerInch');
        end
    elseif isempty(options.resolution)
        options.resolution = 864;
    end

    % Set the default format
    if ~isvector(options) && ~isbitmap(options)
        options.png = true;
    end

    % Check whether transparent background is wanted (old way)
    if isequal(get(ancestor(fig(1), 'figure'), 'Color'), 'none')
        options.transparent = true;
    end

    % If requested, set the resolution to the native vertical resolution of the
    % first suitable image found
    if native && isbitmap(options)
        % Find a suitable image
        list = findall(fig, 'Type','image', 'Tag','export_fig_native');
        if isempty(list)
            list = findall(fig, 'Type','image', 'Visible','on');
        end
        for hIm = list(:)'
            % Check height is >= 2
            height = size(get(hIm, 'CData'), 1);
            if height < 2
                continue
            end
            % Account for the image filling only part of the axes, or vice versa
            yl = get(hIm, 'YData');
            if isscalar(yl)
                yl = [yl(1)-0.5 yl(1)+height+0.5];
            else
                yl = [min(yl), max(yl)];  % fix issue #151 (case of yl containing more than 2 elements)
                if ~diff(yl)
                    continue
                end
                yl = yl + [-0.5 0.5] * (diff(yl) / (height - 1));
            end
            hAx = get(hIm, 'Parent');
            yl2 = get(hAx, 'YLim');
            % Find the pixel height of the axes
            oldUnits = get(hAx, 'Units');
            set(hAx, 'Units', 'pixels');
            pos = get(hAx, 'Position');
            set(hAx, 'Units', oldUnits);
            if ~pos(4)
                continue
            end
            % Found a suitable image
            % Account for stretch-to-fill being disabled
            pbar = get(hAx, 'PlotBoxAspectRatio');
            pos = min(pos(4), pbar(2)*pos(3)/pbar(1));
            % Set the magnification to give native resolution
            options.magnify = abs((height * diff(yl2)) / (pos * diff(yl)));  % magnification must never be negative: issue #103
            break
        end
    end
end

function A = downsize(A, factor)
    % Downsample an image
    if factor == 1
        % Nothing to do
        return
    end
    try
        % Faster, but requires image processing toolbox
        A = imresize(A, 1/factor, 'bilinear');
    catch
        % No image processing toolbox - resize manually
        % Lowpass filter - use Gaussian as is separable, so faster
        % Compute the 1d Gaussian filter
        filt = (-factor-1:factor+1) / (factor * 0.6);
        filt = exp(-filt .* filt);
        % Normalize the filter
        filt = single(filt / sum(filt));
        % Filter the image
        padding = floor(numel(filt) / 2);
        for a = 1:size(A, 3)
            A(:,:,a) = conv2(filt, filt', single(A([ones(1, padding) 1:end repmat(end, 1, padding)],[ones(1, padding) 1:end repmat(end, 1, padding)],a)), 'valid');
        end
        % Subsample
        A = A(1+floor(mod(end-1, factor)/2):factor:end,1+floor(mod(end-1, factor)/2):factor:end,:);
    end
end

function A = rgb2grey(A)
    A = cast(reshape(reshape(single(A), [], 3) * single([0.299; 0.587; 0.114]), size(A, 1), size(A, 2)), class(A)); % #ok<ZEROLIKE>
end

function A = check_greyscale(A)
    % Check if the image is greyscale
    if size(A, 3) == 3 && ...
            all(reshape(A(:,:,1) == A(:,:,2), [], 1)) && ...
            all(reshape(A(:,:,2) == A(:,:,3), [], 1))
        A = A(:,:,1); % Save only one channel for 8-bit output
    end
end

function eps_remove_background(fname, count)
    % Remove the background of an eps file
    % Open the file
    fh = fopen(fname, 'r+');
    if fh == -1
        error('Not able to open file %s.', fname);
    end
    % Read the file line by line
    while count
        % Get the next line
        l = fgets(fh);
        if isequal(l, -1)
            break; % Quit, no rectangle found
        end
        % Check if the line contains the background rectangle
        if isequal(regexp(l, ' *0 +0 +\d+ +\d+ +r[fe] *[\n\r]+', 'start'), 1)
            % Set the line to whitespace and quit
            l(1:regexp(l, '[\n\r]', 'start', 'once')-1) = ' ';
            fseek(fh, -numel(l), 0);
            fprintf(fh, l);
            % Reduce the count
            count = count - 1;
        end
    end
    % Close the file
    fclose(fh);
end

function b = isvector(options)
    b = options.pdf || options.eps;
end

function b = isbitmap(options)
    b = options.png || options.tif || options.jpg || options.bmp || options.im || options.alpha;
end

% Helper function
function A = make_cell(A)
    if ~iscell(A)
        A = {A};
    end
end

function add_bookmark(fname, bookmark_text)
    % Adds a bookmark to the temporary EPS file after %%EndPageSetup
    % Read in the file
    fh = fopen(fname, 'r');
    if fh == -1
        error('File %s not found.', fname);
    end
    try
        fstrm = fread(fh, '*char')';
    catch ex
        fclose(fh);
        rethrow(ex);
    end
    fclose(fh);

    % Include standard pdfmark prolog to maximize compatibility
    fstrm = strrep(fstrm, '%%BeginProlog', sprintf('%%%%BeginProlog\n/pdfmark where {pop} {userdict /pdfmark /cleartomark load put} ifelse'));
    % Add page bookmark
    fstrm = strrep(fstrm, '%%EndPageSetup', sprintf('%%%%EndPageSetup\n[ /Title (%s) /OUT pdfmark',bookmark_text));

    % Write out the updated file
    fh = fopen(fname, 'w');
    if fh == -1
        error('Unable to open %s for writing.', fname);
    end
    try
        fwrite(fh, fstrm, 'char*1');
    catch ex
        fclose(fh);
        rethrow(ex);
    end
    fclose(fh);
end

function set_tick_mode(Hlims, ax)
    % Set the tick mode of linear axes to manual
    % Leave log axes alone as these are tricky
    M = get(Hlims, [ax 'Scale']);
    if ~iscell(M)
        M = {M};
    end
    M = cellfun(@(c) strcmp(c, 'linear'), M);
    set(Hlims(M), [ax 'TickMode'], 'manual');
    %set(Hlims(M), [ax 'TickLabelMode'], 'manual');  % this hides exponent label in HG2!
end

function change_rgb_to_cmyk(fname)  % convert RGB => CMYK within an EPS file
    % Do post-processing on the eps file
    try
        % Read the EPS file into memory
        fstrm = read_write_entire_textfile(fname);

        % Replace all gray-scale colors
        fstrm = regexprep(fstrm, '\n([\d.]+) +GC\n', '\n0 0 0 ${num2str(1-str2num($1))} CC\n');
        
        % Replace all RGB colors
        fstrm = regexprep(fstrm, '\n[0.]+ +[0.]+ +[0.]+ +RC\n', '\n0 0 0 1 CC\n');  % pure black
        fstrm = regexprep(fstrm, '\n([\d.]+) +([\d.]+) +([\d.]+) +RC\n', '\n${sprintf(''%.4g '',[1-[str2num($1),str2num($2),str2num($3)]/max([str2num($1),str2num($2),str2num($3)]),1-max([str2num($1),str2num($2),str2num($3)])])} CC\n');

        % Overwrite the file with the modified contents
        read_write_entire_textfile(fname, fstrm);
    catch
        % never mind - leave as is...
    end
end
```
