## Supplemental File 1 for "A biophysical model of striatal microcircuits suggests delta/theta-rhythmically interleaved gamma and beta oscillations mediate periodicity in motor control": fix_lines.m

```
%FIX_LINES  Improves the line style of eps files generated by print
%
% Examples:
%   fix_lines fname
%   fix_lines fname fname2
%   fstrm_out = fixlines(fstrm_in)
%
% This function improves the style of lines in eps files generated by
% MATLAB's print function, making them more similar to those seen on
% screen. Grid lines are also changed from a dashed style to a dotted
% style, for greater differentiation from dashed lines.
% 
% The function also places embedded fonts after the postscript header, in
% versions of MATLAB which place the fonts first (R2006b and earlier), in
% order to allow programs such as Ghostscript to find the bounding box
% information.
%
%IN:
%   fname - Name or path of source eps file.
%   fname2 - Name or path of destination eps file. Default: same as fname.
%   fstrm_in - File contents of a MATLAB-generated eps file.
%
%OUT:
%   fstrm_out - Contents of the eps file with line styles fixed.

% Copyright: (C) Oliver Woodford, 2008-2014

% The idea of editing the EPS file to change line styles comes from Jiro
% Doke's FIXPSLINESTYLE (fex id: 17928)
% The idea of changing dash length with line width came from comments on
% fex id: 5743, but the implementation is mine :)

% Thank you to Sylvain Favrot for bringing the embedded font/bounding box
% interaction in older versions of MATLAB to my attention.
% Thank you to D Ko for bringing an error with eps files with tiff previews
% to my attention.
% Thank you to Laurence K for suggesting the check to see if the file was
% opened.

% 01/03/15: Issue #20: warn users if using this function in HG2 (R2014b+)
% 27/03/15: Fixed out of memory issue with enormous EPS files (generated by print() with OpenGL renderer), related to issue #39

function fstrm = fix_lines(fstrm, fname2)

% Issue #20: warn users if using this function in HG2 (R2014b+)
if using_hg2
    warning('export_fig:hg2','The fix_lines function should not be used in this Matlab version.');
end
    
if nargout == 0 || nargin > 1
    if nargin < 2
        % Overwrite the input file
        fname2 = fstrm;
    end
    % Read in the file
    fstrm = read_write_entire_textfile(fstrm);
end

% Move any embedded fonts after the postscript header
if strcmp(fstrm(1:15), '%!PS-AdobeFont-')
    % Find the start and end of the header
    ind = regexp(fstrm, '[\n\r]%!PS-Adobe-');
    [ind2, ind2] = regexp(fstrm, '[\n\r]%%EndComments[\n\r]+');
    % Put the header first
    if ~isempty(ind) && ~isempty(ind2) && ind(1) < ind2(1)
        fstrm = fstrm([ind(1)+1:ind2(1) 1:ind(1) ind2(1)+1:end]);
    end
end

% Make sure all line width commands come before the line style definitions,
% so that dash lengths can be based on the correct widths
% Find all line style sections
ind = [regexp(fstrm, '[\n\r]SO[\n\r]'),... % This needs to be here even though it doesn't have dots/dashes!
       regexp(fstrm, '[\n\r]DO[\n\r]'),...
       regexp(fstrm, '[\n\r]DA[\n\r]'),...
       regexp(fstrm, '[\n\r]DD[\n\r]')];
ind = sort(ind);
% Find line width commands
[ind2, ind3] = regexp(fstrm, '[\n\r]\d* w[\n\r]');
% Go through each line style section and swap with any line width commands
% near by
b = 1;
m = numel(ind);
n = numel(ind2);
for a = 1:m
    % Go forwards width commands until we pass the current line style
    while b <= n && ind2(b) < ind(a)
        b = b + 1;
    end
    if b > n
        % No more width commands
        break;
    end
    % Check we haven't gone past another line style (including SO!)
    if a < m && ind2(b) > ind(a+1)
        continue;
    end
    % Are the commands close enough to be confident we can swap them?
    if (ind2(b) - ind(a)) > 8
        continue;
    end
    % Move the line style command below the line width command
    fstrm(ind(a)+1:ind3(b)) = [fstrm(ind(a)+4:ind3(b)) fstrm(ind(a)+1:ind(a)+3)];
    b = b + 1;
end

% Find any grid line definitions and change to GR format
% Find the DO sections again as they may have moved
ind = int32(regexp(fstrm, '[\n\r]DO[\n\r]'));
if ~isempty(ind)
    % Find all occurrences of what are believed to be axes and grid lines
    ind2 = int32(regexp(fstrm, '[\n\r] *\d* *\d* *mt *\d* *\d* *L[\n\r]'));
    if ~isempty(ind2)
        % Now see which DO sections come just before axes and grid lines
        ind2 = repmat(ind2', [1 numel(ind)]) - repmat(ind, [numel(ind2) 1]);
        ind2 = any(ind2 > 0 & ind2 < 12); % 12 chars seems about right
        ind = ind(ind2);
        % Change any regions we believe to be grid lines to GR
        fstrm(ind+1) = 'G';
        fstrm(ind+2) = 'R';
    end
end

% Define the new styles, including the new GR format
% Dot and dash lengths have two parts: a constant amount plus a line width
% variable amount. The constant amount comes after dpi2point, and the
% variable amount comes after currentlinewidth. If you want to change
% dot/dash lengths for a one particular line style only, edit the numbers
% in the /DO (dotted lines), /DA (dashed lines), /DD (dot dash lines) and
% /GR (grid lines) lines for the style you want to change.
new_style = {'/dom { dpi2point 1 currentlinewidth 0.08 mul add mul mul } bdef',... % Dot length macro based on line width
             '/dam { dpi2point 2 currentlinewidth 0.04 mul add mul mul } bdef',... % Dash length macro based on line width
             '/SO { [] 0 setdash 0 setlinecap } bdef',... % Solid lines
             '/DO { [1 dom 1.2 dom] 0 setdash 0 setlinecap } bdef',... % Dotted lines
             '/DA { [4 dam 1.5 dam] 0 setdash 0 setlinecap } bdef',... % Dashed lines
             '/DD { [1 dom 1.2 dom 4 dam 1.2 dom] 0 setdash 0 setlinecap } bdef',... % Dot dash lines
             '/GR { [0 dpi2point mul 4 dpi2point mul] 0 setdash 1 setlinecap } bdef'}; % Grid lines - dot spacing remains constant

% Construct the output
% This is the original (memory-intensive) code:
%first_sec = strfind(fstrm, '% line types:'); % Isolate line style definition section
%[second_sec, remaining] = strtok(fstrm(first_sec+1:end), '/');
%[remaining, remaining] = strtok(remaining, '%');
%fstrm = [fstrm(1:first_sec) second_sec sprintf('%s\r', new_style{:}) remaining];
fstrm = regexprep(fstrm,'(% line types:.+?)/.+?%',['$1',sprintf('%s\r',new_style{:}),'%']);

% Write the output file
if nargout == 0 || nargin > 1
    read_write_entire_textfile(fname2, fstrm);
end
end
```
