## Supplemental File 1 for "A biophysical model of striatal microcircuits suggests delta/theta-rhythmically interleaved gamma and beta oscillations mediate periodicity in motor control": im2gif.m

```
%IM2GIF Convert a multiframe image to an animated GIF file
%
% Examples:
%   im2gif infile
%   im2gif infile outfile
%   im2gif(A, outfile)
%   im2gif(..., '-nocrop')
%   im2gif(..., '-nodither')
%   im2gif(..., '-ncolors', n)
%   im2gif(..., '-loops', n)
%   im2gif(..., '-delay', n) 
%   
% This function converts a multiframe image to an animated GIF.
%
% To create an animation from a series of figures, export to a multiframe
% TIFF file using export_fig, then convert to a GIF, as follows:
%
%    for a = 2 .^ (3:6)
%       peaks(a);
%       export_fig test.tif -nocrop -append
%    end
%    im2gif('test.tif', '-delay', 0.5);
%
%IN:
%   infile - string containing the name of the input image.
%   outfile - string containing the name of the output image (must have the
%             .gif extension). Default: infile, with .gif extension.
%   A - HxWxCxN array of input images, stacked along fourth dimension, to
%       be converted to gif.
%   -nocrop - option indicating that the borders of the output are not to
%             be cropped.
%   -nodither - option indicating that dithering is not to be used when
%               converting the image.
%   -ncolors - option pair, the value of which indicates the maximum number
%              of colors the GIF can have. This can also be a quantization
%              tolerance, between 0 and 1. Default/maximum: 256.
%   -loops - option pair, the value of which gives the number of times the
%            animation is to be looped. Default: 65535.
%   -delay - option pair, the value of which gives the time, in seconds,
%            between frames. Default: 1/15.

% Copyright (C) Oliver Woodford 2011

function im2gif(A, varargin)

% Parse the input arguments
[A, options] = parse_args(A, varargin{:});

if options.crop ~= 0
    % Crop
    A = crop_borders(A, A(ceil(end/2),1,:,1));
end

% Convert to indexed image
[h, w, c, n] = size(A);
A = reshape(permute(A, [1 2 4 3]), h, w*n, c);
map = unique(reshape(A, h*w*n, c), 'rows');
if size(map, 1) > 256
    dither_str = {'dither', 'nodither'};
    dither_str = dither_str{1+(options.dither==0)};
    if options.ncolors <= 1
        [B, map] = rgb2ind(A, options.ncolors, dither_str);
        if size(map, 1) > 256
            [B, map] = rgb2ind(A, 256, dither_str);
        end
    else
        [B, map] = rgb2ind(A, min(round(options.ncolors), 256), dither_str);
    end
else
    if max(map(:)) > 1
        map = double(map) / 255;
        A = double(A) / 255;
    end
    B = rgb2ind(im2double(A), map);
end
B = reshape(B, h, w, 1, n);

% Bug fix to rgb2ind
map(B(1)+1,:) = im2double(A(1,1,:));

% Save as a gif
imwrite(B, map, options.outfile, 'LoopCount', round(options.loops(1)), 'DelayTime', options.delay);
end

%% Parse the input arguments
function [A, options] = parse_args(A, varargin)
% Set the defaults
options = struct('outfile', '', ...
                 'dither', true, ...
                 'crop', true, ...
                 'ncolors', 256, ...
                 'loops', 65535, ...
                 'delay', 1/15);

% Go through the arguments
a = 0;
n = numel(varargin);
while a < n
    a = a + 1;
    if ischar(varargin{a}) && ~isempty(varargin{a})
        if varargin{a}(1) == '-'
            opt = lower(varargin{a}(2:end));
            switch opt
                case 'nocrop'
                    options.crop = false;
                case 'nodither'
                    options.dither = false;
                otherwise
                    if ~isfield(options, opt)
                        error('Option %s not recognized', varargin{a});
                    end
                    a = a + 1;
                    if ischar(varargin{a}) && ~ischar(options.(opt))
                        options.(opt) = str2double(varargin{a});
                    else
                        options.(opt) = varargin{a};
                    end
            end
        else
            options.outfile = varargin{a};
        end
    end
end

if isempty(options.outfile)
    if ~ischar(A)
        error('No output filename given.');
    end
    % Generate the output filename from the input filename
    [path, outfile] = fileparts(A);
    options.outfile = fullfile(path, [outfile '.gif']);
end

if ischar(A)
    % Read in the image
    A = imread_rgb(A);
end
end

%% Read image to uint8 rgb array
function [A, alpha] = imread_rgb(name)
% Get file info
info = imfinfo(name);
% Special case formats
switch lower(info(1).Format)
    case 'gif'
        [A, map] = imread(name, 'frames', 'all');
        if ~isempty(map)
            map = uint8(map * 256 - 0.5); % Convert to uint8 for storage
            A = reshape(map(uint32(A)+1,:), [size(A) size(map, 2)]); % Assume indexed from 0
            A = permute(A, [1 2 5 4 3]);
        end
    case {'tif', 'tiff'}
        A = cell(numel(info), 1);
        for a = 1:numel(A)
            [A{a}, map] = imread(name, 'Index', a, 'Info', info);
            if ~isempty(map)
                map = uint8(map * 256 - 0.5); % Convert to uint8 for storage
                A{a} = reshape(map(uint32(A{a})+1,:), [size(A) size(map, 2)]); % Assume indexed from 0
            end
            if size(A{a}, 3) == 4
                % TIFF in CMYK colourspace - convert to RGB
                if isfloat(A{a})
                    A{a} = A{a} * 255;
                else
                    A{a} = single(A{a});
                end
                A{a} = 255 - A{a};
                A{a}(:,:,4) = A{a}(:,:,4) / 255;
                A{a} = uint8(A(:,:,1:3) .* A{a}(:,:,[4 4 4]));
            end
        end
        A = cat(4, A{:});
    otherwise
        [A, map, alpha] = imread(name);
        A = A(:,:,:,1); % Keep only first frame of multi-frame files
        if ~isempty(map)
            map = uint8(map * 256 - 0.5); % Convert to uint8 for storage
            A = reshape(map(uint32(A)+1,:), [size(A) size(map, 2)]); % Assume indexed from 0
        elseif size(A, 3) == 4
            % Assume 4th channel is an alpha matte
            alpha = A(:,:,4);
            A = A(:,:,1:3);
        end
end
end
```
