## Supplemental File 1 for "A biophysical model of striatal microcircuits suggests delta/theta-rhythmically interleaved gamma and beta oscillations mediate periodicity in motor control": ImageSelection.java

```
/*
 * Based on code snippet from
 * http://java.sun.com/developer/technicalArticles/releases/data/
 *
 * Copyright � 2008, 2010 Oracle and/or its affiliates. All rights reserved. Use is subject to license terms.
 */

import java.awt.image.BufferedImage;
import java.awt.datatransfer.*;

public class ImageSelection implements Transferable {
    
    private static final DataFlavor flavors[] =
    {DataFlavor.imageFlavor};
    
    private BufferedImage image;
    
    public ImageSelection(BufferedImage image) {
        this.image = image;
    }
    
    // Transferable
    public Object getTransferData(DataFlavor flavor) throws UnsupportedFlavorException {
        if (flavor.equals(flavors[0]) == false) {
            throw new UnsupportedFlavorException(flavor);
        }
        return image;
    }
    
    public DataFlavor[] getTransferDataFlavors() {
        return flavors;
    }
    
    public boolean isDataFlavorSupported(DataFlavor
    flavor) {
        return flavor.equals(flavors[0]);
    }
}
```
