## Supplemental File 1 for "A biophysical model of striatal microcircuits suggests delta/theta-rhythmically interleaved gamma and beta oscillations mediate periodicity in motor control": isolate_axes.m

```
function fh = isolate_axes(ah, vis)
%ISOLATE_AXES Isolate the specified axes in a figure on their own
%
% Examples:
%   fh = isolate_axes(ah)
%   fh = isolate_axes(ah, vis)
%
% This function will create a new figure containing the axes/uipanels
% specified, and also their associated legends and colorbars. The objects
% specified must all be in the same figure, but they will generally only be
% a subset of the objects in the figure.
%
% IN:
%    ah - An array of axes and uipanel handles, which must come from the
%         same figure.
%    vis - A boolean indicating whether the new figure should be visible.
%          Default: false.
%
% OUT:
%    fh - The handle of the created figure.

% Copyright (C) Oliver Woodford 2011-2013

% Thank you to Rosella Blatt for reporting a bug to do with axes in GUIs
% 16/03/12: Moved copyfig to its own function. Thanks to Bob Fratantonio
%           for pointing out that the function is also used in export_fig.m
% 12/12/12: Add support for isolating uipanels. Thanks to michael for suggesting it
% 08/10/13: Bug fix to allchildren suggested by Will Grant (many thanks!)
% 05/12/13: Bug fix to axes having different units. Thanks to Remington Reid for reporting
% 21/04/15: Bug fix for exporting uipanels with legend/colorbar on HG1 (reported by Alvaro
%           on FEX page as a comment on 24-Apr-2014); standardized indentation & help section
% 22/04/15: Bug fix: legends and colorbars were not exported when exporting axes handle in HG2

    % Make sure we have an array of handles
    if ~all(ishandle(ah))
        error('ah must be an array of handles');
    end
    % Check that the handles are all for axes or uipanels, and are all in the same figure
    fh = ancestor(ah(1), 'figure');
    nAx = numel(ah);
    for a = 1:nAx
        if ~ismember(get(ah(a), 'Type'), {'axes', 'uipanel'})
            error('All handles must be axes or uipanel handles.');
        end
        if ~isequal(ancestor(ah(a), 'figure'), fh)
            error('Axes must all come from the same figure.');
        end
    end
    % Tag the objects so we can find them in the copy
    old_tag = get(ah, 'Tag');
    if nAx == 1
        old_tag = {old_tag};
    end
    set(ah, 'Tag', 'ObjectToCopy');
    % Create a new figure exactly the same as the old one
    fh = copyfig(fh); %copyobj(fh, 0);
    if nargin < 2 || ~vis
        set(fh, 'Visible', 'off');
    end
    % Reset the object tags
    for a = 1:nAx
        set(ah(a), 'Tag', old_tag{a});
    end
    % Find the objects to save
    ah = findall(fh, 'Tag', 'ObjectToCopy');
    if numel(ah) ~= nAx
        close(fh);
        error('Incorrect number of objects found.');
    end
    % Set the axes tags to what they should be
    for a = 1:nAx
        set(ah(a), 'Tag', old_tag{a});
    end
    % Keep any legends and colorbars which overlap the subplots
    % Note: in HG1 these are axes objects; in HG2 they are separate objects, therefore we
    %       don't test for the type, only the tag (hopefully nobody but Matlab uses them!)
    lh = findall(fh, 'Tag', 'legend', '-or', 'Tag', 'Colorbar');
    nLeg = numel(lh);
    if nLeg > 0
        set([ah(:); lh(:)], 'Units', 'normalized');
        try
            ax_pos = get(ah, 'OuterPosition'); % axes and figures have the OuterPosition property
        catch
            ax_pos = get(ah, 'Position'); % uipanels only have Position, not OuterPosition
        end
        if nAx > 1
            ax_pos = cell2mat(ax_pos(:));
        end
        ax_pos(:,3:4) = ax_pos(:,3:4) + ax_pos(:,1:2);
        try
            leg_pos = get(lh, 'OuterPosition');
        catch
            leg_pos = get(lh, 'Position');  % No OuterPosition in HG2, only in HG1
        end
        if nLeg > 1;
            leg_pos = cell2mat(leg_pos);
        end
        leg_pos(:,3:4) = leg_pos(:,3:4) + leg_pos(:,1:2);
        ax_pos = shiftdim(ax_pos, -1);
        % Overlap test
        M = bsxfun(@lt, leg_pos(:,1), ax_pos(:,:,3)) & ...
            bsxfun(@lt, leg_pos(:,2), ax_pos(:,:,4)) & ...
            bsxfun(@gt, leg_pos(:,3), ax_pos(:,:,1)) & ...
            bsxfun(@gt, leg_pos(:,4), ax_pos(:,:,2));
        ah = [ah; lh(any(M, 2))];
    end
    % Get all the objects in the figure
    axs = findall(fh);
    % Delete everything except for the input objects and associated items
    delete(axs(~ismember(axs, [ah; allchildren(ah); allancestors(ah)])));
end

function ah = allchildren(ah)
    ah = findall(ah);
    if iscell(ah)
        ah = cell2mat(ah);
    end
    ah = ah(:);
end

function ph = allancestors(ah)
    ph = [];
    for a = 1:numel(ah)
        h = get(ah(a), 'parent');
        while h ~= 0
            ph = [ph; h];
            h = get(h, 'parent');
        end
    end
end
```
