## Supplemental File 1 for "A biophysical model of striatal microcircuits suggests delta/theta-rhythmically interleaved gamma and beta oscillations mediate periodicity in motor control": pdf2eps.m

```
%PDF2EPS  Convert a pdf file to eps format using pdftops
%
% Examples:
%   pdf2eps source dest
%
% This function converts a pdf file to eps format.
%
% This function requires that you have pdftops, from the Xpdf suite of
% functions, installed on your system. This can be downloaded from:
% http://www.foolabs.com/xpdf  
%
%IN:
%   source - filename of the source pdf file to convert. The filename is
%            assumed to already have the extension ".pdf".
%   dest - filename of the destination eps file. The filename is assumed to
%          already have the extension ".eps".

% Copyright (C) Oliver Woodford 2009-2010

% Thanks to Aldebaro Klautau for reporting a bug when saving to
% non-existant directories.

function pdf2eps(source, dest)
% Construct the options string for pdftops
options = ['-q -paper match -eps -level2 "' source '" "' dest '"'];
% Convert to eps using pdftops
[status, message] = pdftops(options);
% Check for error
if status
    % Report error
    if isempty(message)
        error('Unable to generate eps. Check destination directory is writable.');
    else
        error(message);
    end
end
% Fix the DSC error created by pdftops
fid = fopen(dest, 'r+');
if fid == -1
    % Cannot open the file
    return
end
fgetl(fid); % Get the first line
str = fgetl(fid); % Get the second line
if strcmp(str(1:min(13, end)), '% Produced by')
    fseek(fid, -numel(str)-1, 'cof');
    fwrite(fid, '%'); % Turn ' ' into '%'
end
fclose(fid);
end
```
