## Supplemental File 1 for "A biophysical model of striatal microcircuits suggests delta/theta-rhythmically interleaved gamma and beta oscillations mediate periodicity in motor control": print2array.m

```
function [A, bcol] = print2array(fig, res, renderer, gs_options)
%PRINT2ARRAY  Exports a figure to an image array
%
% Examples:
%   A = print2array
%   A = print2array(figure_handle)
%   A = print2array(figure_handle, resolution)
%   A = print2array(figure_handle, resolution, renderer)
%   A = print2array(figure_handle, resolution, renderer, gs_options)
%   [A bcol] = print2array(...)
%
% This function outputs a bitmap image of the given figure, at the desired
% resolution.
%
% If renderer is '-painters' then ghostcript needs to be installed. This
% can be downloaded from: http://www.ghostscript.com
%
% IN:
%   figure_handle - The handle of the figure to be exported. Default: gcf.
%   resolution - Resolution of the output, as a factor of screen
%                resolution. Default: 1.
%   renderer - string containing the renderer paramater to be passed to
%              print. Default: '-opengl'.
%   gs_options - optional ghostscript options (e.g.: '-dNoOutputFonts'). If
%                multiple options are needed, enclose in call array: {'-a','-b'}
%
% OUT:
%   A - MxNx3 uint8 image of the figure.
%   bcol - 1x3 uint8 vector of the background color

% Copyright (C) Oliver Woodford 2008-2014, Yair Altman 2015-
%{
% 05/09/11: Set EraseModes to normal when using opengl or zbuffer
%           renderers. Thanks to Pawel Kocieniewski for reporting the issue.
% 21/09/11: Bug fix: unit8 -> uint8! Thanks to Tobias Lamour for reporting it.
% 14/11/11: Bug fix: stop using hardcopy(), as it interfered with figure size
%           and erasemode settings. Makes it a bit slower, but more reliable.
%           Thanks to Phil Trinh and Meelis Lootus for reporting the issues.
% 09/12/11: Pass font path to ghostscript.
% 27/01/12: Bug fix affecting painters rendering tall figures. Thanks to
%           Ken Campbell for reporting it.
% 03/04/12: Bug fix to median input. Thanks to Andy Matthews for reporting it.
% 26/10/12: Set PaperOrientation to portrait. Thanks to Michael Watts for
%           reporting the issue.
% 26/02/15: If temp dir is not writable, use the current folder for temp
%           EPS/TIF files (Javier Paredes)
% 27/02/15: Display suggested workarounds to internal print() error (issue #16)
% 28/02/15: Enable users to specify optional ghostscript options (issue #36)
% 10/03/15: Fixed minor warning reported by Paul Soderlind; fixed code indentation
% 28/05/15: Fixed issue #69: patches with LineWidth==0.75 appear wide (internal bug in Matlab's print() func)
% 07/07/15: Fixed issue #83: use numeric handles in HG1
% 11/12/16: Fixed cropping issue reported by Harry D.
%}

    % Generate default input arguments, if needed
    if nargin < 2
        res = 1;
        if nargin < 1
            fig = gcf;
        end
    end
    % Warn if output is large
    old_mode = get(fig, 'Units');
    set(fig, 'Units', 'pixels');
    px = get(fig, 'Position');
    set(fig, 'Units', old_mode);
    npx = prod(px(3:4)*res)/1e6;
    if npx > 30
        % 30M pixels or larger!
        warning('MATLAB:LargeImage', 'print2array generating a %.1fM pixel image. This could be slow and might also cause memory problems.', npx);
    end
    % Retrieve the background colour
    bcol = get(fig, 'Color');
    % Set the resolution parameter
    res_str = ['-r' num2str(ceil(get(0, 'ScreenPixelsPerInch')*res))];
    % Generate temporary file name
    tmp_nam = [tempname '.tif'];
    try
        % Ensure that the temp dir is writable (Javier Paredes 26/2/15)
        fid = fopen(tmp_nam,'w');
        fwrite(fid,1);
        fclose(fid);
        delete(tmp_nam);  % cleanup
        isTempDirOk = true;
    catch
        % Temp dir is not writable, so use the current folder
        [dummy,fname,fext] = fileparts(tmp_nam); %#ok<ASGLU>
        fpath = pwd;
        tmp_nam = fullfile(fpath,[fname fext]);
        isTempDirOk = false;
    end
    % Enable users to specify optional ghostscript options (issue #36)
    if nargin > 3 && ~isempty(gs_options)
        if iscell(gs_options)
            gs_options = sprintf(' %s',gs_options{:});
        elseif ~ischar(gs_options)
            error('gs_options input argument must be a string or cell-array of strings');
        else
            gs_options = [' ' gs_options];
        end
    else
        gs_options = '';
    end
    if nargin > 2 && strcmp(renderer, '-painters')
        % First try to print directly to tif file
        try
            % Print the file into a temporary TIF file and read it into array A
            [A, err, ex] = read_tif_img(fig, res_str, renderer, tmp_nam);
            if err, rethrow(ex); end
        catch  % error - try to print to EPS and then using Ghostscript to TIF
            % Print to eps file
            if isTempDirOk
                tmp_eps = [tempname '.eps'];
            else
                tmp_eps = fullfile(fpath,[fname '.eps']);
            end
            print2eps(tmp_eps, fig, 0, renderer, '-loose');
            try
                % Initialize the command to export to tiff using ghostscript
                cmd_str = ['-dEPSCrop -q -dNOPAUSE -dBATCH ' res_str ' -sDEVICE=tiff24nc'];
                % Set the font path
                fp = font_path();
                if ~isempty(fp)
                    cmd_str = [cmd_str ' -sFONTPATH="' fp '"'];
                end
                % Add the filenames
                cmd_str = [cmd_str ' -sOutputFile="' tmp_nam '" "' tmp_eps '"' gs_options];
                % Execute the ghostscript command
                ghostscript(cmd_str);
            catch me
                % Delete the intermediate file
                delete(tmp_eps);
                rethrow(me);
            end
            % Delete the intermediate file
            delete(tmp_eps);
            % Read in the generated bitmap
            A = imread(tmp_nam);
            % Delete the temporary bitmap file
            delete(tmp_nam);
        end
        % Set border pixels to the correct colour
        if isequal(bcol, 'none')
            bcol = [];
        elseif isequal(bcol, [1 1 1])
            bcol = uint8([255 255 255]);
        else
            for l = 1:size(A, 2)
                if ~all(reshape(A(:,l,:) == 255, [], 1))
                    break;
                end
            end
            for r = size(A, 2):-1:l
                if ~all(reshape(A(:,r,:) == 255, [], 1))
                    break;
                end
            end
            for t = 1:size(A, 1)
                if ~all(reshape(A(t,:,:) == 255, [], 1))
                    break;
                end
            end
            for b = size(A, 1):-1:t
                if ~all(reshape(A(b,:,:) == 255, [], 1))
                    break;
                end
            end
            bcol = uint8(median(single([reshape(A(:,[l r],:), [], size(A, 3)); reshape(A([t b],:,:), [], size(A, 3))]), 1));
            for c = 1:size(A, 3)
                A(:,[1:l-1, r+1:end],c) = bcol(c);
                A([1:t-1, b+1:end],:,c) = bcol(c);
            end
        end
    else
        if nargin < 3
            renderer = '-opengl';
        end
        % Print the file into a temporary TIF file and read it into array A
        [A, err, ex] = read_tif_img(fig, res_str, renderer, tmp_nam);
        % Throw any error that occurred
        if err
            % Display suggested workarounds to internal print() error (issue #16)
            fprintf(2, 'An error occured with Matlab''s builtin print function.\nTry setting the figure Renderer to ''painters'' or use opengl(''software'').\n\n');
            rethrow(ex);
        end
        % Set the background color
        if isequal(bcol, 'none')
            bcol = [];
        else
            bcol = bcol * 255;
            if isequal(bcol, round(bcol))
                bcol = uint8(bcol);
            else
                bcol = squeeze(A(1,1,:));
            end
        end
    end
    % Check the output size is correct
    if isequal(res, round(res))
        px = round([px([4 3])*res 3]);  % round() to avoid an indexing warning below
        if ~isequal(size(A), px)
            % Correct the output size
            A = A(1:min(end,px(1)),1:min(end,px(2)),:);
        end
    end
end

% Function to create a TIF image of the figure and read it into an array
function [A, err, ex] = read_tif_img(fig, res_str, renderer, tmp_nam)
    err = false;
    ex = [];
    % Temporarily set the paper size
    old_pos_mode    = get(fig, 'PaperPositionMode');
    old_orientation = get(fig, 'PaperOrientation');
    set(fig, 'PaperPositionMode','auto', 'PaperOrientation','portrait');
    try
        % Workaround for issue #69: patches with LineWidth==0.75 appear wide (internal bug in Matlab's print() function)
        fp = [];  % in case we get an error below
        fp = findall(fig, 'Type','patch', 'LineWidth',0.75);
        set(fp, 'LineWidth',0.5);
        % Fix issue #83: use numeric handles in HG1
        if ~using_hg2(fig),  fig = double(fig);  end
        % Print to tiff file
        print(fig, renderer, res_str, '-dtiff', tmp_nam);
        % Read in the printed file
        A = imread(tmp_nam);
        % Delete the temporary file
        delete(tmp_nam);
    catch ex
        err = true;
    end
    set(fp, 'LineWidth',0.75);  % restore original figure appearance
    % Reset the paper size
    set(fig, 'PaperPositionMode',old_pos_mode, 'PaperOrientation',old_orientation);
end

% Function to return (and create, where necessary) the font path
function fp = font_path()
    fp = user_string('gs_font_path');
    if ~isempty(fp)
        return
    end
    % Create the path
    % Start with the default path
    fp = getenv('GS_FONTPATH');
    % Add on the typical directories for a given OS
    if ispc
        if ~isempty(fp)
            fp = [fp ';'];
        end
        fp = [fp getenv('WINDIR') filesep 'Fonts'];
    else
        if ~isempty(fp)
            fp = [fp ':'];
        end
        fp = [fp '/usr/share/fonts:/usr/local/share/fonts:/usr/share/fonts/X11:/usr/local/share/fonts/X11:/usr/share/fonts/truetype:/usr/local/share/fonts/truetype'];
    end
    user_string('gs_font_path', fp);
end
```
