## Supplemental File 1 for "A biophysical model of striatal microcircuits suggests delta/theta-rhythmically interleaved gamma and beta oscillations mediate periodicity in motor control": read_write_entire_textfile.m

```
%READ_WRITE_ENTIRE_TEXTFILE Read or write a whole text file to/from memory
%
% Read or write an entire text file to/from memory, without leaving the
% file open if an error occurs.
%
% Reading:
%   fstrm = read_write_entire_textfile(fname)
% Writing:
%   read_write_entire_textfile(fname, fstrm)
%
%IN:
%   fname - Pathname of text file to be read in.
%   fstrm - String to be written to the file, including carriage returns.
%
%OUT:
%   fstrm - String read from the file. If an fstrm input is given the
%           output is the same as that input. 

function fstrm = read_write_entire_textfile(fname, fstrm)
modes = {'rt', 'wt'};
writing = nargin > 1;
fh = fopen(fname, modes{1+writing});
if fh == -1
    error('Unable to open file %s.', fname);
end
try
    if writing
        fwrite(fh, fstrm, 'char*1');
    else
        fstrm = fread(fh, '*char')';
    end
catch ex
    fclose(fh);
    rethrow(ex);
end
fclose(fh);
end
```
