## Supplemental File 1 for "A biophysical model of striatal microcircuits suggests delta/theta-rhythmically interleaved gamma and beta oscillations mediate periodicity in motor control": using_hg2.m

```
%USING_HG2 Determine if the HG2 graphics engine is used
%
%   tf = using_hg2(fig)
%
%IN:
%   fig - handle to the figure in question.
%
%OUT:
%   tf - boolean indicating whether the HG2 graphics engine is being used
%        (true) or not (false).

% 19/06/2015 - Suppress warning in R2015b; cache result for improved performance
% 06/06/2016 - Fixed issue #156 (bad return value in R2016b)

function tf = using_hg2(fig)
    persistent tf_cached
    if isempty(tf_cached)
        try
            if nargin < 1,  fig = figure('visible','off');  end
            oldWarn = warning('off','MATLAB:graphicsversion:GraphicsVersionRemoval');
            try
                % This generates a [supressed] warning in R2015b:
                tf = ~graphicsversion(fig, 'handlegraphics');
            catch
                tf = ~verLessThan('matlab','8.4');  % =R2014b
            end
            warning(oldWarn);
        catch
            tf = false;
        end
        if nargin < 1,  delete(fig);  end
        tf_cached = tf;
    else
        tf = tf_cached;
    end
end
```
