## Supplemental File 1 for "A biophysical model of striatal microcircuits suggests delta/theta-rhythmically interleaved gamma and beta oscillations mediate periodicity in motor control": arrayfunu.m

```
function varargout = arrayfunu(varargin)
%ARRAYFUNU - Wraps arrayfun, with uniform output set to zero
%
% Usage:
%   out = arrayfunu (varargin)

varargout = cell(1,nargout);
[varargout{:}] = arrayfun(varargin{:},'UniformOutput',0);

end
```
