## Supplemental File 1 for "A biophysical model of striatal microcircuits suggests delta/theta-rhythmically interleaved gamma and beta oscillations mediate periodicity in motor control": catstruct.m

```
function A = catstruct(varargin)
% CATSTRUCT   Concatenate or merge structures with different fieldnames
%   X = CATSTRUCT(S1,S2,S3,...) merges the structures S1, S2, S3 ...
%   into one new structure X. X contains all fields present in the various
%   structures. An example:
%
%     A.name = 'Me' ;
%     B.income = 99999 ;
%     X = catstruct(A,B) 
%     % -> X.name = 'Me' ;
%     %    X.income = 99999 ;
%
%   If a fieldname is not unique among structures (i.e., a fieldname is
%   present in more than one structure), only the value from the last
%   structure with this field is used. In this case, the fields are 
%   alphabetically sorted. A warning is issued as well. An axample:
%
%     S1.name = 'Me' ;
%     S2.age  = 20 ; S3.age  = 30 ; S4.age  = 40 ;
%     S5.honest = false ;
%     Y = catstruct(S1,S2,S3,S4,S5) % use value from S4
%
%   The inputs can be array of structures. All structures should have the
%   same size. An example:
%
%     C(1).bb = 1 ; C(2).bb = 2 ;
%     D(1).aa = 3 ; D(2).aa = 4 ;
%     CD = catstruct(C,D) % CD is a 1x2 structure array with fields bb and aa
%
%   The last input can be the string 'sorted'. In this case,
%   CATSTRUCT(S1,S2, ..., 'sorted') will sort the fieldnames alphabetically. 
%   To sort the fieldnames of a structure A, you could use
%   CATSTRUCT(A,'sorted') but I recommend ORDERFIELDS for doing that.
%
%   When there is nothing to concatenate, the result will be an empty
%   struct (0x0 struct array with no fields).
%
%   NOTE: To concatenate similar arrays of structs, you can use simple
%   concatenation: 
%     A = dir('*.mat') ; B = dir('*.m') ; C = [A ; B] ;

%   NOTE: This function relies on unique. Matlab changed the behavior of
%   its set functions since 2013a, so this might cause some backward
%   compatibility issues when dulpicated fieldnames are found.
%
%   See also CAT, STRUCT, FIELDNAMES, STRUCT2CELL, ORDERFIELDS

% version 4.1 (feb 2015), tested in R2014a
% (c) Jos van der Geest
%

% History
% Created in 2005
% Revisions
%   2.0 (sep 2007) removed bug when dealing with fields containing cell
%                  arrays (Thanks to Rene Willemink)
%   2.1 (sep 2008) added warning and error identifiers
%   2.2 (oct 2008) fixed error when dealing with empty structs (thanks to
%                  Lars Barring)
%   3.0 (mar 2013) fixed problem when the inputs were array of structures
%                  (thanks to Tor Inge Birkenes).
%                  Rephrased the help section as well.
%   4.0 (dec 2013) fixed problem with unique due to version differences in
%                  ML. Unique(...,'last') is no longer the deafult.
%                  (thanks to Isabel P)
%   4.1 (feb 2015) fixed warning with narginchk

narginchk(1,Inf) ;
N = nargin ;

if ~isstruct(varargin{end}),
    if isequal(varargin{end},'sorted'),
        narginchk(2,Inf) ;
        sorted = 1 ;
        N = N-1 ;
    else
        error('catstruct:InvalidArgument','Last argument should be a structure, or the string "sorted".') ;
    end
else
    sorted = 0 ;
end

sz0 = [] ; % used to check that all inputs have the same size

% used to check for a few trivial cases
NonEmptyInputs = false(N,1) ; 
NonEmptyInputsN = 0 ;

% used to collect the fieldnames and the inputs
FN = cell(N,1) ;
VAL = cell(N,1) ;

% parse the inputs
for ii=1:N,
    X = varargin{ii} ;
    if ~isstruct(X),
        error('catstruct:InvalidArgument',['Argument #' num2str(ii) ' is not a structure.']) ;
    end
    
    if ~isempty(X),
        % empty structs are ignored
        if ii > 1 && ~isempty(sz0)
            if ~isequal(size(X), sz0)
                error('catstruct:UnequalSizes','All structures should have the same size.') ;
            end
        else
            sz0 = size(X) ;
        end
        NonEmptyInputsN = NonEmptyInputsN + 1 ;
        NonEmptyInputs(ii) = true ;
        FN{ii} = fieldnames(X) ;
        VAL{ii} = struct2cell(X) ;
    end
end

if NonEmptyInputsN == 0
    % all structures were empty
    A = struct([]) ;
elseif NonEmptyInputsN == 1,
    % there was only one non-empty structure
    A = varargin{NonEmptyInputs} ;
    if sorted,
        A = orderfields(A) ;
    end
else
    % there is actually something to concatenate
    FN = cat(1,FN{:}) ;    
    VAL = cat(1,VAL{:}) ;    
    FN = squeeze(FN) ;
    VAL = squeeze(VAL) ;
    
    
    [UFN,ind] = unique(FN, 'last') ;
    % If this line errors, due to your matlab version not having UNIQUE
    % accept the 'last' input, use the following line instead
    % [UFN,ind] = unique(FN) ; % earlier ML versions, like 6.5
    
    if numel(UFN) ~= numel(FN),
        warning('catstruct:DuplicatesFound','Fieldnames are not unique between structures.') ;
        sorted = 1 ;
    end
    
    if sorted,
        VAL = VAL(ind,:) ;
        FN = FN(ind,:) ;
    end
    
    A = cell2struct(VAL, FN);
    A = reshape(A, sz0) ; % reshape into original format
end
```
