## Supplemental File 1 for "A biophysical model of striatal microcircuits suggests delta/theta-rhythmically interleaved gamma and beta oscillations mediate periodicity in motor control": cellfunu.m

```
function varargout = cellfunu (varargin)
%CELLFUNU - Wraps cellfun, with uniform output set to zero
%
% Usage:
%   out = cellfunu (varargin)

varargout = cell(1,nargout);
[varargout{1:nargout}] = cellfun(varargin{:},'UniformOutput',0);

end
```
