## Supplemental File 1 for "A biophysical model of striatal microcircuits suggests delta/theta-rhythmically interleaved gamma and beta oscillations mediate periodicity in motor control": cellstr2num.m

```
function m = cellstr2num(c)
%cellstr2num - Takes a cell matrix of strings and converts each cell into a number.

  sz = size(c);
  m = zeros(sz);
  for i=1:prod(size(c))
    m(i) = sscanf(c{i},'%f');
  end;
```
