## Supplemental File 1 for "A biophysical model of striatal microcircuits suggests delta/theta-rhythmically interleaved gamma and beta oscillations mediate periodicity in motor control": compareFigFiles2.m

```
function [diffStruct,consolidatedData1,consolidatedData2] = compareFigFiles2(name1, name2, suppressPrintBool)
% compareFigFiles2 - compare .fig files or folders containing .fig files
%
% Syntax:
%           compareFigFiles(folderName1, folderName2)
%           compareFigFiles(folderName, figFilename)
%           compareFigFiles(figFilename, folderName)
%           [diffStruct,data1,data2] = compareFigFiles(figFilename1, figFilename2)
%
% Description:
%           compareFigFiles compares *.fig files, reporting internal components
%           and properties that are different between corresponding fig files.
%
%           this second version removes problematic fields.
%
%           Inputs can be either a figure filename or folder name. When a
%           folder name is specified, then all the corresponding fig file(s)
%           in that folder will be compared to the other folder/file.
%           Note: when one of the inputs is a folder, then only files that
%           have the same name will be compared.
%
%           compareFigFiles(folderOrFilename) compares the specified input to
%           the current folder (pwd).
%
%           [diffStruct,data1,data2] = compareFigFiles(file1,file2) returns
%           a Matlab struct containing the non-matching components/properties.
%           Each of the struct fields corresponds to a specific figure handle
%           and property name, and contains a cell array of 2 values, for each
%           of the compared files. data1 and data2 contain the raw data used
%           for the comparison - a Matlab struct with fields corresponding to
%           each of the components/properties defined in the fig file.
%
%           [diffStructCells,data1,data2] = compareFigFiles(folderName1, folderName2)
%           returns a Matlab cell of structs , each containing the non-matching
%           components/properties.
%           Each of the struct fields corresponds to a specific figure handle
%           and property name, and contains a cell array of 2 values, for each
%           of the compared files. data1 and data2 contain the raw data used
%           for the comparison - a Matlab struct with fields corresponding to
%           each of the components/properties defined in the fig file.
%
% Examples:
%           compareFigFiles('C:\Yair',pwd);  % compares corresponding fig files in 2 folders
%           compareFigFiles('C:\Yair');      % (same as above)
%
%           compareFigFiles('myApp',    'hisApp');      % compare 2 FIG files
%           compareFigFiles('myApp.fig','hisApp.fig');  % (same as above)
%
%           compareFigFiles('C:\Yair\myApp');     % compare C:\Yair\myApp.fig to (pwd)\myApp.fig
%           compareFigFiles('C:\Yair\myApp',pwd); % (same as above)
%           compareFigFiles(pwd,'C:\Yair\myApp'); % (same as above)
%
% Note:
%           Doesn't work with the 'compact' mode of savefig.
%
% Technical Description:
%           See http://undocumentedmatlab.com/blog/fig-files-format/
%
% Bugs and suggestions:
%    Please send to Yair Altman (altmany at gmail dot com)
%
% Change log:
%    2013-07-02: First version posted on <a href="http://www.mathworks.com/matlabcentral/fileexchange/authors/27420">MathWorks File Exchange</a>
%    2017-04-22: added suppressPrintBool, permit diffStruct for folders, make
%     figures visisble

% License to use and modify this code is granted freely to all interested, as long as the original author is
% referenced and attributed as such. The original author maintains the right to be solely associated with this work.

% Programmed and Copyright by Yair M. Altman: altmany(at)gmail.com
% $Revision: 1.01 $  $Date: 2013/07/02 15:22:51 $
% Modified by Erik Roberts

    % Process optional input args
    if nargin < 1,   help(mfilename); return;  end
    if nargin == 1,  name2 = pwd;       end
    if nargin < 3, suppressPrintBool = false; end

    % Parse folder/file args
    if isdir(name1) && isdir(name2)
        % Treat as a folder comparison
        %clc  % this might be useful when there are numerous files being compared...
        files = dir(fullfile(name1, '*.fig'));
        for fileIdx = 1 : length(files)
            % Compare all corresponding fig files one at a time
            thisFile  = fullfile(name1, files(fileIdx).name);
            otherFile = fullfile(name2, files(fileIdx).name);
            if exist(otherFile,'file')
                [diffStruct_I{fileIdx}, consolidatedData1_I(fileIdx), consolidatedData2_I(fileIdx)] = compareSingleFigFiles(thisFile, otherFile);

                if ~isempty(diffStruct_I) && ~suppressPrintBool
                    fprintf('\n');
                end
            end
        end

    elseif isdir(name1)  % but not isdir(name2)
        % Compare name2.fig to the same file in the name1 folder
        [fpath,fname,fext] = fileparts(name2); %#ok<ASGLU>
        otherFile = fullfile(name1, [fname,fext]);
        [diffStruct_I,consolidatedData1_I,consolidatedData2_I] = compareSingleFigFiles(otherFile, name2);

    elseif isdir(name2)  % but not isdir(name1)
        % Compare name1.fig to the same file in the name2 folder
        [fpath,fname,fext] = fileparts(name1); %#ok<ASGLU>
        otherFile = fullfile(name2, [fname,fext]);
        [diffStruct_I,consolidatedData1_I,consolidatedData2_I] = compareSingleFigFiles(name1, otherFile);

    else
        % Compare single .fig files
        [diffStruct_I,consolidatedData1_I,consolidatedData2_I] = compareSingleFigFiles(name1, name2);
    end
    
    % close hidden figures
    close all;

    % Process output args
    if nargout
        diffStruct = diffStruct_I;
        consolidatedData1 = consolidatedData1_I;
        consolidatedData2 = consolidatedData2_I;
    end
    
  %% nested functions
  
  % Compare 2 FIG files, reporting their differences
  function [diffStruct,consolidatedData1,consolidatedData2] = compareSingleFigFiles(fig1Filename, fig2Filename)
    fig1Filename = normalizeFigFilename(fig1Filename);
    fig2Filename = normalizeFigFilename(fig2Filename);
    f1 = strrep(fig1Filename,'\','\\');
    f2 = strrep(fig2Filename,'\','\\');
    f1FullPath = getFullPath(fig1Filename);
    f2FullPath = getFullPath(fig2Filename);
    if ~suppressPrintBool
      fprintf(['Comparing <a href="matlab:guide(''' strrep(f1FullPath,'\','\\') ''')">' strrep(f1,'.fig','') '</a>, ' ...
        '<a href="matlab:guide(''' strrep(f2FullPath,'\','\\') ''')">' strrep(f2,'.fig','') '</a>\n']);
    end
    data1 = getFigData(fig1Filename);
    data2 = getFigData(fig2Filename);
    consolidatedData1 = processStruct([],data1,'');
    consolidatedData2 = processStruct([],data2,'');
    
    consolidatedData1 = rmIssueFlds(consolidatedData1);
    consolidatedData2 = rmIssueFlds(consolidatedData2);
    
    diffStruct = objdiff(consolidatedData1, consolidatedData2);
    %if ~nargout
    if ~suppressPrintBool
      disp(diffStruct);
    end
    %end
  end
end

%% local functions

% Read a FIG file and extract its (unconsolidated) data struct
function data = getFigData(figFilename)
    % Load the FIG file as a MAT file (see http://undocumentedmatlab.com/blog/fig-files-format/)
    data = load(figFilename,'-mat');
    
    flds = fieldnames(data);
    for fld = flds(:)'
      if ~isempty(data.(fld{1}))
        data.(fld{1}).GraphicsObjects.Format3Data.Visible = 'off';
      else
%         data = rmfield(data, fld{1});
      end
    end

    % The file contains a struct with version info (will be 080000 in HG2)
    %data = data.hgS_070000;
    fn = fieldnames(data);
    data = data.(fn{1});
end

% Remove problematic fields
function data = rmIssueFlds(data)
  flds = {'figure_axes_ApplicationData.graphics_linkaxes.LinkProp', 'figure_legend_ApplicationData'};
  for fld = flds(:)'
    eval(['data.' fld{1} '= [];']);
  end
end

% Get the full path of the specified filename
function filename = getFullPath(filename)
    f = which(filename);
    if ~isempty(f)
        filename = f;
    elseif exist(filename,'file')
        if ~any(filename==':')  % i.e., not absolute path
            f = fullfile(pwd,filename);
            if exist(filename,'file')
                filename = f;
            end
        end
    else
        %error([filename ' file was not found']);
    end
end

% Append .FIG to filenames that do not have it specified
function filename = normalizeFigFilename(filename)
    [fpath,fname,fext] = fileparts(filename);  %#ok
    if isempty(fext)
        filename = [filename '.fig'];
    end
end

% Recursively process a FIG-format struct to get consolidated data in a flat struct
function consolidatedData = processStruct(consolidatedData,data,parentTag)
    thisType  = data.type;
    dataProps = data.properties;
    try thisTag = dataProps.Tag; catch, thisTag = thisType;  end
    if isempty(thisTag),  thisTag = 'tag';  end
    thisTag = strrep(thisTag,'.','');
    thisTag = [parentTag thisTag '_'];
    if ~isempty(parentTag)
        thisTag = strrep(thisTag, 'figure1_', '');
    end

    newFieldname = [thisTag 'Type'];
    newFieldname(64:end) = '';
    consolidatedData.(newFieldname) = thisType;
    propNames = fieldnames(dataProps);
    for propIdx = 1 : length(propNames)
        propName = propNames{propIdx};
        newFieldname = [thisTag propName];
        newFieldname(64:end) = '';
        consolidatedData.(newFieldname) = dataProps.(propName);
    end

    for childIdx = 1 : length(data.children)
        consolidatedData = processStruct(consolidatedData, data.children(childIdx), thisTag);
    end
end

% The following was taken from the ObjDiff utility (FEX #14395)
% http://www.mathworks.com/matlabcentral/fileexchange/14395-objdiff-generic-object-comparator
% ===========================================================================================

%% objdiff - compares two objects & returns an object of the same type with just the different fields/values
function [objectC,IA,IB] = objdiff(objectA,objectB,varargin)
% OBJDIFF  compares two objects & returns an object of the same type with just the different fields/values.
%
%   OBJDIFF (unlike Matlab's SETDIFF or SETXOR) also compares structs, GUI
%   handles, ActiveX, Matlab & Java objects, in addition to arrays & cells.
%   OBJDIFF also allows comparison of numeric cell arrays, unlike SETDIFF/
%   SETXOR. It also accepts everything that SETDIFF/SETXOR accept.
%
%   Syntax: [objectC,IA,IB] = objdiff (objectA, objectB, options, ...)
%
%   Inputs:
%     objectA - first object to compare
%     objectB - second object to compare. Field order in opaque objects does not matter.
%               Note: If objectB is not supplied, then objectA(1) is compared to objectA(2)
%     options - optional flags as follows:
%       'rows' - see documentation for <a href="matlab:doc setxor">SETXOR</a>
%       'dontIgnoreJava' - show different instances of the same java class (default=ignore them)
%
%   Outputs:
%     objectC - object containing only the different (or new) fields, in a {old, new} pattern
%     IA,IB - index vector into objectA,objectB such that objectC = [objectA(IA),objectB(IB)] (see SETXOR)
%
%   Examples:
%     >> objectA = struct('a',3, 'b',5, 'd',9);
%     >> objectB = struct('a','ert', 'c',struct('t',pi), 'd',9);
%     >> objectC = objdiff(objectA, objectB)  % a=different, b=new in objectA, c=new in objectB, d=same
%     objectC = 
%         a: {[3]  'ert'}
%         b: {[5]  {}}
%         c: {{}  [1x1 struct]}
%
%     >> objectC = objdiff(java.awt.Color.red, java.awt.Color.blue)
%     objectC = 
%         Blue: {[0]  [255]}
%          RGB: {[-65536]  [-16776961]}
%          Red: {[255]  [0]}
%
%     >> objectC = objdiff(0,gcf)  % 0 is the root handle
%     objectC = 
%           children: {[2x1 struct]  []}
%             handle: {[0]  [1]}
%         properties: {[1x1 struct]  [1x1 struct]}
%               type: {'root'  'figure'}
%
%     >> [objectC,IA,IB] = objdiff({2,3,4,7}, {2,4,5})
%     objectC =
%          3     5     7
%     IA =
%          2     4
%     IB =
%          3
%
%   Bugs and suggestions:
%     Please send to Yair Altman (altmany at gmail dot com)
%
%   Change log:
%     2007-07-27: Fixed handling of identical objects per D. Gamble
%     2007-03-23: First version posted on <a href="http://www.mathworks.com/matlabcentral/fileexchange/loadAuthor.do?objectType=author&mfx=1&objectId=1096533#">MathWorks File Exchange</a>
%
%   See also:
%     SETDIFF, SETXOR, ISSTRUCT, ISJAVA, ISHGHANDLE, ISOBJECT, ISCELL

% Programmed by Yair M. Altman: altmany(at)gmail.com
% $Revision: 1.2 $  $Date: 2007/07/26 23:41:20 $

  % Process input args
  if (nargin<1) %|| ~isstruct(objectA) || ~isstruct(objectB)
      help objdiff
      error('YMA:OBJDIFF:NotEnoughInputs', 'Not enough input arguments');
  elseif (nargin<2) || (nargin==2 && ~strcmp(class(objectA),class(objectB)))
      if numel(objectA) < 2
          error('YMA:OBJDIFF:NotEnoughInputs', 'Not enough input arguments');
      elseif numel(objectA) > 2
          warning('YMA:OBJDIFF:TooManyInputs', 'Too many elements in objectA - only comparing first 2');
      end
      objectB = objectA(2);
      objectA = objectA(1);
      varargin = {objectB, varargin{:}};
  elseif ~strcmp(class(objectA),class(objectB))
      error('YMA:OBJDIFF:DissimilarObjects', 'Input objects must be of the same type');
  end

  % Process optional options
  ignoreJavaObjectsFlag = true;
  if ~isempty(varargin)
      ignoreJavaIdx = strmatch('dontignorejava',lower(varargin{:}));
      if ~isempty(ignoreJavaIdx)
          ignoreJavaObjectsFlag = false;
          varargin(ignoreJavaIdx) = [];
      end
  end

  % TODO: check for array of java/struct objs

  % Convert opaque objects to structs with the relevant property fields
  if ishghandle(objectA)
      objectA = handle2struct(objectA);
      objectB = handle2struct(objectB);
  else%if isjava(object(A))
      try
          % This should work for any opaque object: Java, ActiveX & Matlab
          objectA = get(objectA);
          objectB = get(objectB);
      catch
          % never mind - try to process as-is
      end
  end

  % Enable comparison of numeric cell arrays
  objectA = decell(objectA);
  objectB = decell(objectB);

  % Process based on object type
  if isstruct(objectA)
      % Structs - loop over all fields
      [objectC, IA, IB] = compareStructs(objectA, objectB, ignoreJavaObjectsFlag);
  else
      % Cells and arrays - process with the regular setdiff function
      [objectC, IA, IB] = setxor(objectA, objectB, varargin{:});
  end
end

% Compare two structs
function [objectC,IA,IB] = compareStructs(objectA,objectB,ignoreJavaObjectsFlag)
  % Ensure singleton objects are compared
  objectA = getSingleton(objectA);
  objectB = getSingleton(objectB);
  objectC = struct();

  % Get all the fieldnames
  fieldsA = fieldnames(objectA);
  fieldsB = fieldnames(objectB);
  allFields = union(fieldsA, fieldsB);

  % Loop over all fields and compare the objects
  for fieldIdx = 1 : length(allFields)
      fieldName = allFields{fieldIdx};
      if ~isfield(objectA,fieldName)
          objectC.(fieldName) = {{}, objectB.(fieldName)};
      elseif ~isfield(objectB,fieldName)
          objectC.(fieldName) = {objectA.(fieldName), {}};
      elseif isa(objectA.(fieldName),'function_handle') && isa(objectB.(fieldName),'function_handle')
          funcA = char(objectA.(fieldName));
          funcB = char(objectB.(fieldName));
          if ~strcmp(funcA,funcB)
              objectC.(fieldName) = {funcA,funcB};
          end
      elseif ~isequalwithequalnans(objectA.(fieldName), objectB.(fieldName))
          if ignoreJavaObjectsFlag && isjava(objectA.(fieldName)) && isjava(objectB.(fieldName)) && ...
                  isequalwithequalnans(objectA.(fieldName).getClass, objectB.(fieldName).getClass)
              continue;
          elseif isempty(objectA.(fieldName)) && isempty(objectB.(fieldName))  % e.g., [] vs. {}
              continue;  % treat as being the same
          end
          objectC.(fieldName) = {objectA.(fieldName), objectB.(fieldName)};
      end
  end

  % Check for empty diff struct (identical input objects)
  if isempty(fieldnames(objectC))
      objectC = struct([]);
  end

  % no use for IA,IB...
  IA = [];
  IB = [];
end

% De-cell-ize a numeric cell-array
function obj = decell(obj)
  if iscell(obj) && ~iscellstr(obj)
      obj = cell2mat(obj);
  end
end

% Ensure singleton object
function obj = getSingleton(obj)
  if numel(obj) > 1
      warning('YMA:OBJDIFF:TooManyElements', 'Too many elements in %s - only comparing the first', inputname(1));
      obj = obj(1);
  end
end
```
