## Supplemental File 1 for "A biophysical model of striatal microcircuits suggests delta/theta-rhythmically interleaved gamma and beta oscillations mediate periodicity in motor control": coverage_example.m

```
%add source code to path
addpath('src');
addpath('lib/absolutepath');

%import classses, functions
import edu.stanford.covert.test.Coverage;

%turn profiler on
profile('on', '-nohistory');

%run code
plot(rand(10, 1))
close all;

%turn profiler off
profile('off');

%generate coverage report
outFileName = 'example.coverage.xml';
report = Coverage('src', '..');
report.exportXML(outFileName);
```
