## Supplemental File 1 for "A biophysical model of striatal microcircuits suggests delta/theta-rhythmically interleaved gamma and beta oscillations mediate periodicity in motor control": absolutepath.m

```
function  abs_path = absolutepath( rel_path, act_path, throwErrorIfFileNotExist )
%ABSOLUTEPATH  returns the absolute path relative to a given startpath.
%   The startpath is optional, if omitted the current dir is used instead.
%   Both argument must be strings.
%
%   Syntax:
%      abs_path = ABSOLUTEPATH( rel_path, start_path )
%
%   Parameters:
%      rel_path           - Relative path
%      start_path         - Start for relative path  (optional, default = current dir)
%
%   Examples:
%      absolutepath( '.\data\matlab'        , 'C:\local' ) = 'c:\local\data\matlab\'
%      absolutepath( 'A:\MyProject\'        , 'C:\local' ) = 'a:\myproject\'
%
%      absolutepath( '.\data\matlab'        , cd         ) is the same as
%      absolutepath( '.\data\matlab'                     )
%
%   See also:  RELATIVEPATH PATH

%   Jochen Lenz

%   Jonathan karr 12/17/2010
%   - making compatible with linux
%   - commented out lower cases
%   - switching findstr to strfind
%   - fixing mlint warnings
%   Jonathan karr 1/11/2011
%   - Per Abel Brown's comments adding optional error checking for absolute path of directories that don't exist
%   Jonathan karr 1/12/2011
%   - fixing bugs and writing test

% 2nd parameter is optional:
if nargin < 3
    throwErrorIfFileNotExist = true;
    if  nargin < 2
        if (ispc && numel(rel_path) > 3 && rel_path(2) == ':' && (rel_path(3) == '/' || rel_path(3) == '\')) || ...
           (~ispc && numel(rel_path) > 1 && rel_path(1) == filesep)
            act_path = [];
        else
            act_path = [pwd filesep];
        end
    end
end

%build absolute path
file = java.io.File([act_path rel_path]);
abs_path = char(file.getCanonicalPath());

%check that file exists
if throwErrorIfFileNotExist && ~exist(abs_path, 'file')
    throw(MException('absolutepath:fileNotExist', 'The path %s or file %s doesn''t exist', abs_path, abs_path));
end
```
