## Supplemental File 1 for "A biophysical model of striatal microcircuits suggests delta/theta-rhythmically interleaved gamma and beta oscillations mediate periodicity in motor control": relativepath.m

```
function  rel_path = relativepath( tgt_path, act_path )
%RELATIVEPATH  returns the relative path from an actual path to the target path.
%   Both arguments must be strings with absolute paths.
%   The actual path is optional, if omitted the current dir is used instead.
%   In case the volume drive letters don't match, an absolute path will be returned.
%   If a relative path is returned, it always starts with '.\' or '..\'
%
%   Syntax:
%      rel_path = RELATIVEPATH( target_path, actual_path )
%   
%   Parameters:
%      target_path        - Path which is targetted
%      actual_path        - Start for relative path (optional, default = current dir)
%
%   Examples:
%      relativepath( 'C:\local\data\matlab' , 'C:\local' ) = '.\data\matlab\'
%      relativepath( 'A:\MyProject\'        , 'C:\local' ) = 'a:\myproject\'
%
%      relativepath( 'C:\local\data\matlab' , cd         ) is the same as
%      relativepath( 'C:\local\data\matlab'              )
%
%   See also:  ABSOLUTEPATH PATH

%   Jochen Lenz


% 2nd parameter is optional:
if  nargin < 2
   act_path = cd;
end

% Predefine return string:
rel_path = '';

% Make sure strings end by a filesep character:
if  length(act_path) == 0   |   ~isequal(act_path(end),filesep)
   act_path = [act_path filesep];
end
if  length(tgt_path) == 0   |   ~isequal(tgt_path(end),filesep)
   tgt_path = [tgt_path filesep];
end

% Convert to all lowercase:
[act_path] = fileparts( lower(act_path) );
[tgt_path] = fileparts( lower(tgt_path) );

% Create a cell-array containing the directory levels:
act_path_cell = pathparts(act_path);
tgt_path_cell = pathparts(tgt_path);

% If volumes are different, return absolute path:
if  length(act_path_cell) == 0   |   length(tgt_path_cell) == 0
   return  % rel_path = ''
else
   if  ~isequal( act_path_cell{1} , tgt_path_cell{1} )
      rel_path = tgt_path;
      return
   end
end

% Remove level by level, as long as both are equal:
while  length(act_path_cell) > 0   &   length(tgt_path_cell) > 0
   if  isequal( act_path_cell{1}, tgt_path_cell{1} )
      act_path_cell(1) = [];
      tgt_path_cell(1) = [];
   else
      break
   end
end

% As much levels down ('..\') as levels are remaining in "act_path":
for  i = 1 : length(act_path_cell)
   rel_path = ['..' filesep rel_path];
end

% Relative directory levels to target directory:
for  i = 1 : length(tgt_path_cell)
   rel_path = [rel_path tgt_path_cell{i} filesep];
end

% Start with '.' or '..' :
if  isempty(rel_path)
   rel_path = ['.' filesep];
elseif  ~isequal(rel_path(1),'.')
   rel_path = ['.' filesep rel_path];
end

return

% -------------------------------------------------

function  path_cell = pathparts(path_str)

path_str = [filesep path_str filesep];
path_cell = {};

sep_pos = findstr( path_str, filesep );
for i = 1 : length(sep_pos)-1
   path_cell{i} = path_str( sep_pos(i)+1 : sep_pos(i+1)-1 );
end

return
```
