## Supplemental File 1 for "A biophysical model of striatal microcircuits suggests delta/theta-rhythmically interleaved gamma and beta oscillations mediate periodicity in motor control": Coverage.m

```
%Coverage
%  Generates XML coverage reports of executed code in 1 format:
%  - Cobertura
%
%  Example usage:
%    report = edu.stanford.covert.test.Coverage(<source dir>, <profile data (optional)>);
%    report.exportCoberturaXML(<fileName>);
%
%  This class was inspired by the mlcovr package by David Hart at Sandia
%  National Labs
%  (http://www.mathworks.com/matlabcentral/fileexchange/24985-mlcovr-package).
%
%  Author: Jonathan Karr
%  Affiliation: Covert Lab, Bioengineering, Stanford University
%  Last Updated: 5/1/17
%   Erik Roberts: change to built in strjoin

classdef Coverage
    properties
        sources
        basePath
        timeStamp
        stats
        packages
    end
    
    methods
        %- sources is a string indicating a source file directory, or a cell
        %  array containing multiple directories.
        %- basepath is string indicaticating path relative to which fileNames
        %  should be reported
        %- Optionall profData is a struct containing profile data generated by
        %  the MATLAB profiler and constructed by calling
        %    profData = profile('info')
        %  If profData is not provided the current profile data is used.
        function this = Coverage(sources, basePath, profData)
            if ~exist('sources', 'var')
                sources = {'.' 'r'};
            end
            if ~exist('basePath', 'var')
                basePath = '.';
            end
            basePath = absolutepath(basePath);
            
            if ~iscell(sources)
                sources = {sources 'r'};
            end
            for i = 1:size(sources, 1)
                sources{i, 1} = absolutepath(sources{i,1});
                
                if ~isdir(sources{i,1})                    
                    throw(MException('Coverage:invalidSource', 'Sources must be directories'));
                end
                
                if numel(basePath) > numel(sources{i,1}) || ~strcmp(sources{i,1}(1:numel(basePath)), basePath)
                    fprintf('Base Path:%s\nSource path:%s\n', basePath, sources{i,1})
                    throw(MException('Coverage:invalidPath', 'source path must be contained within base path'));
                end
            end
            
            if ~exist('profData', 'var')
                profData = profile('info');
            end
            
            this.sources = sources;
            this.basePath = basePath;
            this.timeStamp = datestr(now, 31);
            this = this.buildReportFromDir(profData);
        end
    end
    
    methods (Access = protected)
        function this = buildReportFromDir(this, profData)
            %% get package names
            packageNames = {};
            for i = 1:size(this.sources, 1)
                packageNames = [packageNames; this.getPackageNamesFromDir(this.sources{i,1})];
            end
            packageNames = unique(packageNames);
            
            %% build report
            %packages
            packages = repmat(struct('name', [], ...
                'linesCovered', [], 'linesValid', [], 'lineRate', [], ...
                'branchesCovered', [], 'branchesValid',[], 'branchRate', [], ...
                'complexity', [], ...
                'classes', []), [0 1]); %#ok<*PROP>

            this.stats = struct('lineRate', [], 'branchRate', []);
            this.stats.linesCovered = 0;
            this.stats.linesValid = 0;
            this.stats.branchesCovered = 0;
            this.stats.branchesValid = 0;
            this.stats.complexity = 0;

            for i = 1:numel(packageNames)
                package = this.buildPackageReport(meta.package.fromName(packageNames{i}), profData);
                
                if isempty(package)
                    continue;
                end
                
                packages = [packages;
                    package];
                
                this.stats.linesCovered = this.stats.linesCovered + package.linesCovered;
                this.stats.linesValid = this.stats.linesValid + package.linesValid;
                this.stats.branchesCovered = this.stats.branchesCovered + package.branchesCovered;
                this.stats.branchesValid = this.stats.branchesValid + package.branchesValid;
                this.stats.complexity = this.stats.complexity + package.complexity;
            end
            
            %overall stats
            this.packages = packages;            
            this.stats.lineRate = this.stats.linesCovered / this.stats.linesValid;
            this.stats.branchRate = this.stats.branchesCovered / this.stats.branchesValid;
        end
        
        function package = buildPackageReport(this, metaPackage, profData)
            package = [];
            
            if isempty(metaPackage.Classes) && isempty(metaPackage.Functions)
                return;
            end
            
            package.name = metaPackage.Name;
            package.classes = [];
            package.linesCovered = 0;
            package.branchesCovered = 0;
            package.linesValid = 0;
            package.branchesValid = 0;
            package.complexity = 0;
            package.classes = [];
            
            %classes
            for i = 1:numel(metaPackage.Classes)
                class = this.buildClassReport(metaPackage.Classes{i}, profData);
                
                if isempty(class); continue; end
                
                package.classes = [
                    package.classes;
                    class];
                
                package.linesCovered = package.linesCovered + class.linesCovered;
                package.branchesCovered = package.branchesCovered + class.branchesCovered;
                package.linesValid = package.linesValid + class.linesValid;
                package.branchesValid = package.branchesValid + class.branchesValid;
                package.complexity = package.complexity + class.complexity;
            end
            
            %package functions
            for i = 1:numel(metaPackage.Functions)
                class = this.buildFunctionReport(metaPackage.Name, metaPackage.Functions{i}, profData);
                
                if isempty(class); continue; end
                
                package.classes = [
                    package.classes;
                    class];
                
                package.linesCovered = package.linesCovered + class.linesCovered;
                package.branchesCovered = package.branchesCovered + class.branchesCovered;
                package.linesValid = package.linesValid + class.linesValid;
                package.branchesValid = package.branchesValid + class.branchesValid;
                package.complexity = package.complexity + class.complexity;
            end
                
            if isempty(package.classes)
                package = [];
                return;
            end
            
            %package stats
            package.lineRate = package.linesCovered / package.linesValid;
            package.branchRate = package.branchesCovered / package.branchesValid;
        end
        
        function class = buildClassReport(this, metaClass, profData)
            class = [];
            
            %check that metadata is valid
            try
                meta.class.fromName(metaClass.Name);
            catch %#ok<CTCH>
                warning('Coverage:warning', 'Invalid meta data for class %s', metaClass.Name);
            end
            
            fileName = which(metaClass.Name);
            inSources = false;
            for i = 1:size(this.sources, 1)
                source = this.sources{i, 1};
                if ...
                        (isequal(this.sources{i, 2}, 'r') && numel(fileName) >= numel(source) && strcmp(fileName(1:numel(source)+1), [source filesep])) || ...
                        (~isequal(this.sources{i, 2}, 'r') && numel(fileName) >= numel(source) && strcmp(fileName(1:numel(source)+1), [source filesep]) && ~any(strcmp(fileName(numel(source)+2:end), filesep)))
                    inSources = true;
                    break;
                end
            end
            
            if ~inSources
                return; 
            end
            
            class.name = metaClass.Name;
            class.fileName = fileName;            
            class.linesCovered = 0;
            class.branchesCovered = 0;
            class.linesValid = 0;
            class.branchesValid = 0;
            class.complexity = 0;
            class.methods = [];
            class.lines = [];
            
            shortClassName = metaClass.Name(find(metaClass.Name == '.', 1, 'last')+1:end);
            methodsCallInfo = getcallinfo(fileName);
            runnableLineIndex = unique(callstats('file_lines', fileName));
            code = {};
            fid = fopen(fileName, 'r');
            while ~feof(fid)
                code{end+1} = fgetl(fid);
            end
            fclose(fid);
            
            %methods
            metaClass; %#ok<VUNUS> %This line needed, otherwise numel(metaClass.Methods) incorrectly returns 0
            for i = 1:numel(metaClass.Methods)
                metaMethod = metaClass.Methods{i};
                
                if ~isequal(metaMethod.DefiningClass, metaClass)
                    continue;
                end
                
%                 signature = sprintf('[%s] = %s(%s)', ...
%                     strjoin(', ', metaMethod.OutputNames{:}),...
%                     metaMethod.Name, ...
%                     strjoin(', ', metaMethod.InputNames{:}));
                  
                signature = sprintf('[%s] = %s(%s)', ...
                    strjoin(metaMethod.OutputNames, ', '),...
                    metaMethod.Name, ...
                    strjoin(metaMethod.InputNames, ', '));
                
                method = this.buildMethodReport(...
                    metaMethod.Name, signature, ...
                    shortClassName, fileName, methodsCallInfo, runnableLineIndex, code, profData);
                
                if isempty(method); continue; end;
                
                class.methods = [
                    class.methods;
                    method];
                
                class.linesCovered = class.linesCovered + sum([method.linesCovered]);
                class.branchesCovered = class.branchesCovered + sum([method.branchesCovered]);
                class.linesValid = class.linesValid + sum([method.linesValid]);
                class.branchesValid = class.branchesValid + sum([method.branchesValid]);
                class.complexity = class.complexity + sum([method.complexity]);
            end
            
            %getters
            for i = 1:numel(metaClass.Properties)
                if ~isequal(metaClass.Properties{i}.DefiningClass, metaClass)
                    continue;
                end
                
                method = this.buildMethodReport(...
                    ['get.' metaClass.Properties{i}.Name], ['value = get.' metaClass.Properties{i}.Name '(this)'], ...
                    shortClassName, fileName, methodsCallInfo, runnableLineIndex, code, profData);
                
                if isempty(method); continue; end;
                
                class.methods = [
                    class.methods;
                    method];
                
                class.linesCovered = class.linesCovered + method.linesCovered;
                class.branchesCovered = class.branchesCovered + method.branchesCovered;
                class.linesValid = class.linesValid + method.linesValid;
                class.branchesValid = class.branchesValid + method.branchesValid;
                class.complexity = class.complexity + method.complexity;
            end
            
            %setters
            for i = 1:numel(metaClass.Properties)
                if ~isequal(metaClass.Properties{i}.DefiningClass, metaClass)
                    continue;
                end
                
                method = this.buildMethodReport(...
                    ['set.' metaClass.Properties{i}.Name], ['set.' metaClass.Properties{i}.Name '(this, value)'], ...
                    shortClassName, fileName, methodsCallInfo, runnableLineIndex, code, profData);
                
                if isempty(method); continue; end;
                
                class.methods = [
                    class.methods;
                    method];
                
                class.linesCovered = class.linesCovered + method.linesCovered;
                class.branchesCovered = class.branchesCovered + method.branchesCovered;
                class.linesValid = class.linesValid + method.linesValid;
                class.branchesValid = class.branchesValid + method.branchesValid;
                class.complexity = class.complexity + method.complexity;
            end
            
            %nested functions and subfunctions
            for i = 1:numel(methodsCallInfo)
                methodCallInfo = methodsCallInfo(i);
                if ~ismember(methodCallInfo.type, {'nested-function', 'subfunction'},'legacy')
                    continue;
                end
                                
                method = this.buildMethodReport(...
                    methodCallInfo.fullname(find(methodCallInfo.fullname == '.',1,'first')+1:end), [], ...
                    shortClassName, fileName, methodsCallInfo, runnableLineIndex, code, profData);
                
                if isempty(method); continue; end;
                
                class.methods = [
                    class.methods;
                    method];
                
                class.linesCovered = class.linesCovered + method.linesCovered;
                class.branchesCovered = class.branchesCovered + method.branchesCovered;
                class.linesValid = class.linesValid + method.linesValid;
                class.branchesValid = class.branchesValid + method.branchesValid;
                class.complexity = class.complexity + method.complexity;
            end
            
            %lines            
            for i = 1:numel(class.methods)
                if ~strcmp(class.methods(i).fileName, fileName); continue; end;
                
                class.lines = [
                    class.lines;
                    class.methods(i).lines];
            end
            
            %class stats
            class.lineRate = class.linesCovered / class.linesValid;
            class.branchRate = class.branchesCovered / class.branchesValid;
        end
        
        function method = buildMethodReport(this, name, signature, shortClassName, fileName, methodsCallInfo, runnableLineIndex, code, profData)
            %initialize output
            method = [];                      
            
            methodCallInfo = [
                methodsCallInfo(strcmp({methodsCallInfo.type}, 'class method') & strcmp({methodsCallInfo.name}, name));
                methodsCallInfo(strcmp({methodsCallInfo.type}, 'nested-function') & strcmp({methodsCallInfo.fullname}, [shortClassName '>' shortClassName '.' name]));
                methodsCallInfo(strcmp({methodsCallInfo.type}, 'subfunction') & strcmp({methodsCallInfo.fullname}, [shortClassName '>' name]))];
            
            %method not defined in main class file
            if isempty(methodCallInfo)
                if ~any(fileName == '@'); return; end;
                methodFileName = [fileName(1:find(fileName == filesep, 1, 'last')) name '.m'];
                if ~exist(methodFileName, 'file'); return; end;
                
                try
                    methodCallInfo = getcallinfo(methodFileName);
                    runnableLineIndex = unique(callstats('file_lines', methodFileName));
                catch %#ok<CTCH>
                    return;
                end
                code = {};
                fid = fopen(methodFileName, 'r');
                while ~feof(fid)
                    code{end+1} = fgetl(fid);
                end
                fclose(fid);
                
                method = [];
                for i = 1:numel(methodCallInfo)
                    if i == 1
                        methodProfData = profData.FunctionTable(strcmp({profData.FunctionTable.CompleteName}, [methodFileName '>' name]));
                    else
                        methodProfData = profData.FunctionTable(strcmp({profData.FunctionTable.CompleteName}, [methodFileName '>' name '.' methodCallInfo(i).name]));
                        signature = [];
                    end
                    method = [method;
                        this.buildSubFunctionReport(methodFileName, code, runnableLineIndex, methodCallInfo(i), signature, methodProfData)];
                end
                return;
            end
            
            %method defined in main class file
            method.name = name;
            method.fileName = fileName;
            method.signature = signature;
            
            %lines
            methodProfData = profData.FunctionTable(strcmp({profData.FunctionTable.CompleteName}, [fileName '>' shortClassName '.' name]));
            [method.lines, method.linesCovered, method.linesValid, method.lineRate, ...
                method.branchesCovered, method.branchesValid, method.branchRate, method.complexity] = ...
                this.buildLinesReport(code, runnableLineIndex, methodCallInfo, methodProfData);
            
            %method stats
            if ~isempty(methodProfData)
                method.calls = methodProfData.NumCalls;
                method.time = methodProfData.TotalTime;
            else
                method.calls = 0;
                method.time = 0;
            end
        end
        
        function class = buildFunctionReport(this, packageName, metaFunction, profData)
            class = [];
            
            fileName = which([packageName '.' metaFunction.Name]);
            inSources = false;
            for i = 1:size(this.sources, 1)
                source = this.sources{i, 1};
                if ...
                        (isequal(this.sources{i, 2}, 'r') && numel(fileName) >= numel(source) && strcmp(fileName(1:numel(source)+1), [source filesep])) || ...
                        (~isequal(this.sources{i, 2}, 'r') && numel(fileName) >= numel(source) && strcmp(fileName(1:numel(source)+1), [source filesep]) && ~any(strcmp(fileName(numel(source)+2:end), filesep)))
                    inSources = true;
                    break;
                end
            end
            
            if ~inSources
                return;
            end
            
            class.name = ['_' metaFunction.Name];
            class.fileName = fileName;
            class.methods = [];
            class.lines = [];
            class.linesCovered = 0;
            class.branchesCovered = 0;
            class.linesValid = 0;
            class.branchesValid = 0;
            class.complexity = 0;
            
            methodsCallInfo = getcallinfo(fileName);
            runnableLineIndex = unique(callstats('file_lines', fileName));
            code = {};
            fid = fopen(fileName, 'r');
            while ~feof(fid)
                code{end+1} = fgetl(fid);
            end
            fclose(fid);
            
            for i = 1:numel(methodsCallInfo)
                methodCallInfo = methodsCallInfo(i);
                methodProfData = profData.FunctionTable(strcmp({profData.FunctionTable.CompleteName}, [fileName '>' methodCallInfo.name]));
                method = this.buildSubFunctionReport(fileName, code, runnableLineIndex, methodCallInfo, [], methodProfData);
                
                class.methods = [
                    class.methods;
                    method];
                
                class.linesCovered = class.linesCovered + method.linesCovered;
                class.branchesCovered = class.branchesCovered + method.branchesCovered;
                class.linesValid = class.linesValid + method.linesValid;
                class.branchesValid = class.branchesValid +method.branchesValid;
                class.complexity = class.complexity + method.complexity;
            end
            
            %lines
            class.lines = [];
            for i = 1:numel(class.methods)
                class.lines = [
                    class.lines;
                    class.methods(i).lines];
            end
            
            %stats
            class.lineRate = class.linesCovered / class.linesValid;
            class.branchRate = class.branchesCovered / class.branchesValid;
        end
        
        function method = buildSubFunctionReport(this, fileName, code, runnableLineIndex, callInfo, signature, profData)
            method.name = callInfo.name;
            method.fileName = fileName;
            method.signature = signature;
            
            %lines
            [method.lines, method.linesCovered, method.linesValid, method.lineRate, ...
                method.branchesCovered, method.branchesValid, method.branchRate, method.complexity] = ...
                this.buildLinesReport(code, runnableLineIndex, callInfo, profData);
            
            %method stats
            if ~isempty(profData)
                method.calls = profData.NumCalls;
                method.time = profData.TotalTime;
            else
                method.calls = 0;
                method.time = 0;
            end
        end
        
        function [lines, linesCovered, linesValid, lineRate, branchesCovered, branchesValid, branchRate, complexity] = ...
                buildLinesReport(~, code, runLineIndex, callInfo, profData)
            lines = repmat(struct('number', [], 'hits', [], 'branch', [], 'time', [], 'complex', []), ...
                sum(runLineIndex >= callInfo.firstline & runLineIndex <= callInfo.lastline), 1);
            
            linesCovered = 0;
            branchesCovered = 0;
            linesValid = 0;
            branchesValid = 0;
            complexity = 0;
                        
            l2 = 0;
            for l1 = callInfo.firstline:callInfo.lastline
                if ~any(runLineIndex == l1); continue; end;
                l2 = l2+1;
                
                hits = 0;
                time = 0;
                if ~isempty(profData)
                    idx = find(profData.ExecutedLines(:, 1) == l1);
                    if ~isempty(idx)
                        hits = profData.ExecutedLines(idx, 2);
                        time = profData.ExecutedLines(idx, 3);
                    end
                end
                
                branch = ~isempty(regexp(code{l1}, '\<(if|switch|try)\>', 'once', 'tokens'));
                complex = ~isempty(regexp(code{l1},'(\<otherwise\>)|(\<catch\>)|(\<elseif\>)|(\<if\>)|(\<while\>)|(\<for\>)|(\<case\>)|(\<continue\>)|(&&)|(||)','once','tokens'));
                lines(l2, 1).number = l1;
                lines(l2, 1).hits = hits;
                lines(l2, 1).branch = branch;
                lines(l2, 1).time = time;
                lines(l2, 1).complex = complex;
                
                linesValid = linesValid + 1;
                if branch; branchesValid = branchesValid + 1; end;
                if hits > 0; linesCovered = linesCovered + 1; end;
                if hits > 0 && branch; branchesCovered = branchesCovered + 1; end;
                if complex; complexity = complexity + 1; end;
            end
            
            lineRate = linesCovered / linesValid;
            branchRate = branchesCovered / branchesCovered;
        end
        
        function packageNames = getPackageNamesFromDir(this, source)
            if source(end) ~= filesep
                source = [source filesep];
            end
            packageNames = {};
            
            files = dir(source);
            for i = 1:numel(files)
                if files(i).isdir && files(i).name(1) ~= '.'
                    if files(i).name(1) == '+'
                        metaPackage = meta.package.fromName(files(i).name(2:end));
                        if isempty(metaPackage); continue; end;
                        packageNames = [packageNames;
                            this.getSubPackageNamesFromPackage(metaPackage)];
                    else
                        packageNames = [packageNames;
                            this.getPackageNamesFromDir([source files(i).name])];
                    end
                end
            end
        end
        
        function packageNames = getSubPackageNamesFromPackage(this, metaPackage)
            packageNames = {metaPackage.Name};
            
            for i = 1:numel(metaPackage.Packages)
                packageNames = [packageNames; this.getSubPackageNamesFromPackage(metaPackage.Packages{i})];
            end
        end
        
        function relativePath = relativeToBasePath(this, absolutePath)
            if numel(this.basePath) > numel(absolutePath) || ~strcmp(absolutePath(1:numel(this.basePath)), this.basePath)
                throw(MException('Coverage:invalidPath', 'path must be contained within base path'));
            end
            relativePath = absolutePath(numel(this.basePath)+1:end);
        end
    end
    
    methods
        %See also http://cobertura.sourceforge.net/xml/coverage-04.dtd
        function xml = exportXML(this, fileName)
            %initialize
            xml = sprintf('<?xml version="1.0" encoding="UTF-8"?>\n');
            xml = [xml sprintf('<!DOCTYPE coverage SYSTEM "http://cobertura.sourceforge.net/xml/coverage-04.dtd">\n')];
            
            %top-level            
            xml = [xml sprintf('<coverage\n')];
            xml = [xml sprintf('\tline-rate="%f"\n', this.stats.lineRate)];
            xml = [xml sprintf('\tbranch-rate="%f"\n', this.stats.branchRate)];
            xml = [xml sprintf('\tlines-covered="%d"\n', this.stats.linesCovered)];
            xml = [xml sprintf('\tlines-valid="%d"\n', this.stats.linesValid)];
            xml = [xml sprintf('\tbranches-covered="%d"\n', this.stats.branchesCovered)];
            xml = [xml sprintf('\tbranches-valid="%d"\n', this.stats.branchesValid)];
            xml = [xml sprintf('\tcomplexity="%f"\n', this.stats.complexity)];
            xml = [xml sprintf('\tversion="%s"\n', '0.1')];
            xml = [xml sprintf('\ttimestamp="%s">\n', this.timeStamp)];
            xml = [xml sprintf('\t<sources>\n')];
            for i = 1:size(this.sources, 1)
                xml = [xml sprintf('\t\t<source><![CDATA[%s]]></source>\n', this.sources{i, 1})];
            end
            xml = [xml sprintf('\t</sources>\n')];
            xml = [xml sprintf('\t<packages>\n')];            
            
            %packages
            for i = 1:numel(this.packages)
                package = this.packages(i);
                
                xml = [xml sprintf('\t\t<package name="%s" lines-covered="%d" lines-valid="%d" line-rate="%f" branches-covered="%d" branches-valid="%d" branch-rate="%f" complexity="%f">\n', ...
                    package.name, ...
                    package.linesCovered, package.linesValid, package.lineRate, ...
                    package.branchesCovered, package.branchesValid, package.branchRate, ...
                    package.complexity)]; %#ok<*AGROW>
                xml = [xml sprintf('\t\t\t<classes>\n')];
                %classes in package
                for j = 1:numel(package.classes)
                    class = package.classes(j);
                    
                    xml = [xml sprintf('\t\t\t\t<class name="%s" filename="%s" lines-covered="%d" lines-valid="%d" line-rate="%f" branches-covered="%d" branches-valid="%d" branch-rate="%f" complexity="%f">\n', ...
                        class.name, this.relativeToBasePath(class.fileName), ...
                        class.linesCovered, class.linesValid, class.lineRate, ...
                        class.branchesCovered, class.branchesValid, class.branchRate, ...
                        class.complexity)];
                    xml = [xml sprintf('\t\t\t\t\t<methods>\n')];
                    %methods in class
                    for k = 1:numel(class.methods)
                        method = class.methods(k);
                        
                        xml = [xml sprintf('\t\t\t\t\t\t<method name="%s" filename="%s" signature="%s" lines-covered="%d" lines-valid="%d" line-rate="%f" branches-covered="%d" branches-valid="%d" branch-rate="%f" complexity="%f" calls="%d" time="%f">\n', ...
                            method.name, this.relativeToBasePath(method.fileName), method.signature, ...
                            method.linesCovered, method.linesValid, method.lineRate, ...
                            method.branchesCovered, method.branchesValid, method.branchRate, ...
                            method.complexity, ...
                            method.calls, method.time)];
                        xml = [xml sprintf('\t\t\t\t\t\t\t<lines>\n')];
                        for l = 1:numel(method.lines)
                            line = method.lines(l);
                            xml = [xml sprintf('\t\t\t\t\t\t\t\t<line number="%d" hits="%d" branch="%d" time="%f" condition-coverage="0%% (0/0)"/>\n', ...
                                line.number, line.hits, line.branch, line.time)];
                        end
                        xml = [xml sprintf('\t\t\t\t\t\t\t</lines>\n')];
                        xml = [xml sprintf('\t\t\t\t\t\t</method>\n')];
                    end
                    xml = [xml sprintf('\t\t\t\t\t</methods>\n')];
                    
                    %lines in class
                    xml = [xml sprintf('\t\t\t\t\t<lines>\n')];
                    for k = 1:numel(class.lines)
                        line = class.lines(k);
                        xml = [xml sprintf('\t\t\t\t\t\t<line number="%d" hits="%d" branch="%d" time="%f" condition-coverage="0%% (0/0)"/>\n',...
                            line.number, line.hits, line.branch, line.time)];
                    end
                    xml = [xml sprintf('\t\t\t\t\t</lines>\n')];
                    xml = [xml sprintf('\t\t\t\t</class>\n')];
                end
                xml = [xml sprintf('\t\t\t</classes>\n')];
                xml = [xml sprintf('\t\t</package>\n')];
            end
            xml = [xml sprintf('\t</packages>\n')];
            xml = [xml sprintf('</coverage>\n')];
            
            if exist('fileName', 'var')
                fid = fopen(fileName, 'w');
                fwrite(fid, xml);
                fclose(fid);
            end
        end
    end
end
```
