## Supplemental File 1 for "A biophysical model of striatal microcircuits suggests delta/theta-rhythmically interleaved gamma and beta oscillations mediate periodicity in motor control": DataHash.m

```
function Hash = DataHash(Data, Opt)
% DATAHASH - Checksum for Matlab array of any type
% This function creates a hash value for an input of any type. The type and
% dimensions of the input are considered as default, such that UINT8([0,0]) and
% UINT16(0) have different hash values. Nested STRUCTs and CELLs are parsed
% recursively.
%
% Hash = DataHash(Data, Opt)
% INPUT:
%   Data: Array of these built-in types:
%           (U)INT8/16/32/64, SINGLE, DOUBLE, (real/complex, full/sparse)
%           CHAR, LOGICAL, CELL (nested), STRUCT (scalar or array, nested),
%           function_handle.
%   Opt:  Struct to specify the hashing algorithm and the output format.
%         Opt and all its fields are optional.
%         Opt.Method: String, known methods for Java 1.6 (Matlab 2011b):
%              'SHA-1', 'SHA-256', 'SHA-384', 'SHA-512', 'MD2', 'MD5'.
%            Call DataHash without inputs to get a list of available methods.
%            Default: 'MD5'.
%         Opt.Format: String specifying the output format:
%            'hex', 'HEX':      Lower/uppercase hexadecimal string.
%            'double', 'uint8': Numerical vector.
%            'base64':          Base64 encoded string, only printable ASCII
%                               characters, shorter than 'hex', no padding.
%            Default: 'hex'.
%         Opt.Input: Type of the input as string, not case-sensitive:
%            'array': The contents, type and size of the input [Data] are
%                     considered  for the creation of the hash. Nested CELLs
%                     and STRUCT arrays are parsed recursively. Empty arrays of
%                     different type reply different hashs.
%            'file':  [Data] is treated as file name and the hash is calculated
%                     for the files contents.
%            'bin':   [Data] is a numerical, LOGICAL or CHAR array. Only the
%                     binary contents of the array is considered, such that
%                     e.g. empty arrays of different type reply the same hash.
%            'ascii': Same as 'bin', but only the 8-bit ASCII part of the 16-bit
%                     Matlab CHARs is considered.
%            Default: 'array'.
%
% OUTPUT:
%   Hash: String, DOUBLE or UINT8 vector. The length depends on the hashing
%         method.
%
% EXAMPLES:
% % Default: MD5, hex:
%   DataHash([])                % 5b302b7b2099a97ba2a276640a192485
% % MD5, Base64:
%   Opt = struct('Format', 'base64', 'Method', 'MD5');
%   DataHash(int32(1:10), Opt)  % +tJN9yeF89h3jOFNN55XLg
% % SHA-1, Base64:
%   S.a = uint8([]);
%   S.b = {{1:10}, struct('q', uint64(415))};
%   Opt.Method = 'SHA-1';
%   Opt.Format = 'HEX';
%   DataHash(S, Opt)            % 18672BE876463B25214CA9241B3C79CC926F3093
% % SHA-1 of binary values:
%   Opt = struct('Method', 'SHA-1', 'Input', 'bin');
%   DataHash(1:8, Opt)          % 826cf9d3a5d74bbe415e97d4cecf03f445f69225
% % SHA-256, consider ASCII part only (Matlab's CHAR has 16 bits!):
%   Opt.Method = 'SHA-256';
%   Opt.Input  = 'ascii';
%   DataHash('abc', Opt)
%       % ba7816bf8f01cfea414140de5dae2223b00361a396177a9cb410ff61f20015ad
%   % Or equivalently:
%   Opt.Input = 'bin';
%   DataHash(uint8('abc'), Opt)
%
% NOTES:
%   Function handles and user-defined objects cannot be converted uniquely:
%   - The subfunction ConvertFuncHandle uses the built-in function FUNCTIONS,
%     but the replied struct can depend on the Matlab version.
%   - It is tried to convert objects to UINT8 streams in the subfunction
%     ConvertObject. A conversion by STRUCT() might be more appropriate.
%   Adjust these subfunctions on demand.
%
%   MATLAB CHARs have 16 bits! Use Opt.Input='ascii' for comparisons with e.g.
%   online hash generators.
%
%   Matt Raum suggested this for e.g. user-defined objects:
%     DataHash(getByteStreamFromArray(Data)
%   This works very well, but unfortunately getByteStreamFromArray is
%   undocumented, such that it might vanish in the future or reply different
%   output.
%
%   For arrays the calculated hash value might be changed in new versions.
%   Calling this function without inputs replies the version of the hash.
%
%   The C-Mex function GetMD5 is 2 to 100 times faster, but obtains MD5 only:
%   http://www.mathworks.com/matlabcentral/fileexchange/25921
%
% Tested: Matlab 7.7, 7.8, 7.13, 8.6, WinXP/32, Win7/64
% Author: Jan Simon, Heidelberg, (C) 2011-2016 matlab.2010(a)n(MINUS)simon.de
%
% See also: TYPECAST, CAST.
%
% Michael Kleder, "Compute Hash", no structs and cells:
%   http://www.mathworks.com/matlabcentral/fileexchange/8944
% Tim, "Serialize/Deserialize", converts structs and cells to a byte stream:
%   http://www.mathworks.com/matlabcentral/fileexchange/29457

% $JRev: R-H V:033 Sum:R+m7rAPNLvlw Date:18-Jun-2016 14:33:17 $
% $License: BSD (use/copy/change/redistribute on own risk, mention the author) $
% $File: Tools\GLFile\DataHash.m $
% History:
% 001: 01-May-2011 21:52, First version.
% 007: 10-Jun-2011 10:38, [Opt.Input], binary data, complex values considered.
% 011: 26-May-2012 15:57, Fixed: Failed for binary input and empty data.
% 014: 04-Nov-2012 11:37, Consider Mex-, MDL- and P-files also.
%      Thanks to David (author 243360), who found this bug.
%      Jan Achterhold (author 267816) suggested to consider Java objects.
% 016: 01-Feb-2015 20:53, Java heap space exhausted for large files.
%      Now files are process in chunks to save memory.
% 017: 15-Feb-2015 19:40, Collsions: Same hash for different data.
%      Examples: zeros(1,1) and zeros(1,1,0)
%                complex(0) and zeros(1,1,0,0)
%      Now the number of dimensions is included, to avoid this.
% 022: 30-Mar-2015 00:04, Bugfix: Failed for strings and [] without TYPECASTX.
%      Ross found these 2 bugs, which occur when TYPECASTX is not installed.
%      If you need the base64 format padded with '=' characters, adjust
%      fBase64_enc as you like.
% 026: 29-Jun-2015 00:13, Changed hash for STRUCTs.
%      Struct arrays are analysed field by field now, which is much faster.
% 027: 13-Sep-2015 19:03, 'ascii' input as abbrev. for Input='bin' and UINT8().
% 028: 15-Oct-2015 23:11, Example values in help section updated to v022.
% 029: 16-Oct-2015 22:32, Use default options for empty input.
% 031: 28-Feb-2016 15:10, New hash value to get same reply as GetMD5.
%      New Matlab version (at least 2015b) use a fast method for TYPECAST, such
%      that calling James Tursa's TYPECASTX is not needed anymore.
%      Matlab 6.5 not supported anymore: MException for CATCH.
% 033: 18-Jun-2016 14:28, BUGFIX: Failed on empty files.
%      Thanks to Christian (AuthorID 2918599).

% OPEN BUGS:
% Nath wrote:
% function handle refering to struct containing the function will create
% infinite loop. Is there any workaround ?
% Example:
%   d= dynamicprops();
%   addprop(d,'f');
%   d.f= @(varargin) struct2cell(d);
%   DataHash(d.f) % infinite loop
% This is caught with an error message concerning the recursion limit now.

% Main function: ===============================================================
% Default options: -------------------------------------------------------------
Method    = 'MD5';
OutFormat = 'hex';
isFile    = false;
isBin     = false;

% Check number and type of inputs: ---------------------------------------------
nArg = nargin;
if nArg == 2
   if isa(Opt, 'struct') == 0   % Bad type of 2nd input:
      Error_L('BadInput2', '2nd input [Opt] must be a struct.');
   end
   
   % Specify hash algorithm:
   if isfield(Opt, 'Method')  && ~isempty(Opt.Method)   % Short-circuiting
      Method = upper(Opt.Method);
   end
   
   % Specify output format:
   if isfield(Opt, 'Format') && ~isempty(Opt.Format)    % Short-circuiting
      OutFormat = Opt.Format;
   end
   
   % Check if the Input type is specified - default: 'array':
   if isfield(Opt, 'Input') && ~isempty(Opt.Input)      % Short-circuiting
      if strcmpi(Opt.Input, 'File')
         if ischar(Data) == 0
            Error_L('CannotOpen', '1st input FileName must be a string');
         end
         isFile = true;
         
      elseif strncmpi(Opt.Input, 'bin', 3)  % Accept 'binary' also
         if (isnumeric(Data) || ischar(Data) || islogical(Data)) == 0 || ...
               issparse(Data)
            Error_L('BadDataType', ...
               '1st input must be numeric, CHAR or LOGICAL for binary input.');
         end
         isBin = true;
         
      elseif strncmpi(Opt.Input, 'asc', 3)  % 8-bit ASCII characters
         if ~ischar(Data)
            Error_L('BadDataType', ...
               '1st input must be a CHAR for the input type ASCII.');
         end
         isBin = true;
         Data  = uint8(Data);
      end
   end
   
elseif nArg == 0  % Reply version of this function:
   R = Version_L;
   
   if nargout == 0
      disp(R);
   else
      Hash = R;
   end
   
   return;
   
elseif nArg ~= 1  % Bad number of arguments:
   Error_L('BadNInput', '1 or 2 inputs required.');
end

% Create the engine: -----------------------------------------------------------
try
   Engine = java.security.MessageDigest.getInstance(Method);
catch
   Error_L('BadInput2', 'Invalid algorithm: [%s].', Method);
end

% Create the hash value: -------------------------------------------------------
if isFile
   % Open the file:
   FID = fopen(Data, 'r');
   if FID < 0
      % Check existence of file:
      Found = FileExist_L(Data);
      if Found
         Error_L('CantOpenFile', 'Cannot open file: %s.', Data);
      else
         Error_L('FileNotFound', 'File not found: %s.', Data);
      end
   end
   
   % Read file in chunks to save memory and Java heap space:
   Chunk = 1e6;      % Fastest for 1e6 on Win7/64, HDD
   Count = Chunk;    % Dummy value to satisfy WHILE condition
   while Count == Chunk
      [Data, Count] = fread(FID, Chunk, '*uint8');
      if Count ~= 0  % Avoid error for empty file
         Engine.update(Data);
      end
   end
   fclose(FID);
   
   % Calculate the hash:
   Hash = typecast(Engine.digest, 'uint8');
   
elseif isBin             % Contents of an elementary array, type tested already:
   if isempty(Data)      % Nothing to do, Engine.update fails for empty input!
      Hash = typecast(Engine.digest, 'uint8');
   else                  % Matlab's TYPECAST is less elegant:
      if isnumeric(Data)
         if isreal(Data)
            Engine.update(typecast(Data(:), 'uint8'));
         else
            Engine.update(typecast(real(Data(:)), 'uint8'));
            Engine.update(typecast(imag(Data(:)), 'uint8'));
         end
      elseif islogical(Data)               % TYPECAST cannot handle LOGICAL
         Engine.update(typecast(uint8(Data(:)), 'uint8'));
      elseif ischar(Data)                  % TYPECAST cannot handle CHAR
         Engine.update(typecast(uint16(Data(:)), 'uint8'));
         % Bugfix: Line removed
      end
      Hash = typecast(Engine.digest, 'uint8');
   end
else                 % Array with type:
   Engine = CoreHash(Data, Engine);
   Hash   = typecast(Engine.digest, 'uint8');
end

% Convert hash specific output format: -----------------------------------------
switch OutFormat
   case 'hex'
      Hash = sprintf('%.2x', double(Hash));
   case 'HEX'
      Hash = sprintf('%.2X', double(Hash));
   case 'double'
      Hash = double(reshape(Hash, 1, []));
   case 'uint8'
      Hash = reshape(Hash, 1, []);
   case 'base64'
      Hash = fBase64_enc(double(Hash));
   otherwise
      Error_L('BadOutFormat', ...
         '[Opt.Format] must be: HEX, hex, uint8, double, base64.');
end

% return;

% ******************************************************************************
function Engine = CoreHash(Data, Engine)
% This methods uses the slower TYPECAST of Matlab

% Consider the type and dimensions of the array to distinguish arrays with the
% same data, but different shape: [0 x 0] and [0 x 1], [1,2] and [1;2],
% DOUBLE(0) and SINGLE([0,0]):
% <  v016: [class, size, data]. BUG! 0 and zeros(1,1,0) had the same hash!
% >= v016: [class, ndims, size, data]
Engine.update([uint8(class(Data)), ...
              typecast(uint64([ndims(Data), size(Data)]), 'uint8')]);
           
if issparse(Data)                    % Sparse arrays to struct:
   [S.Index1, S.Index2, S.Value] = find(Data);
   Engine                        = CoreHash(S, Engine);
elseif isstruct(Data)                % Hash for all array elements and fields:
   F = sort(fieldnames(Data));       % Ignore order of fields
   for iField = 1:length(F)          % Loop over fields
      aField = F{iField};
      Engine.update(uint8(aField));
      for iS = 1:numel(Data)         % Loop over elements of struct array
         Engine = CoreHash(Data(iS).(aField), Engine);
      end
   end
elseif iscell(Data)                  % Get hash for all cell elements:
   for iS = 1:numel(Data)
      Engine = CoreHash(Data{iS}, Engine);
   end
elseif isempty(Data)                 % Nothing to do
elseif isnumeric(Data)
   if isreal(Data)
      Engine.update(typecast(Data(:), 'uint8'));
   else
      Engine.update(typecast(real(Data(:)), 'uint8'));
      Engine.update(typecast(imag(Data(:)), 'uint8'));
   end
elseif islogical(Data)               % TYPECAST cannot handle LOGICAL
   Engine.update(typecast(uint8(Data(:)), 'uint8'));
elseif ischar(Data)                  % TYPECAST cannot handle CHAR
   Engine.update(typecast(uint16(Data(:)), 'uint8'));
elseif isa(Data, 'function_handle')
   Engine = CoreHash(ConvertFuncHandle(Data), Engine);
elseif (isobject(Data) || isjava(Data)) && ismethod(Data, 'hashCode')
   Engine = CoreHash(char(Data.hashCode), Engine);
else  % Most likely a user-defined object:
   try
      BasicData = ConvertObject(Data);
   catch ME
      error(['JSimon:', mfilename, ':BadDataType'], ...
         '%s: Cannot create elementary array for type: %s\n  %s', ...
         mfilename, class(Data), ME.message);
   end
   
   try
      Engine = CoreHash(BasicData, Engine);
   catch ME
      if strcmpi(ME.identifier, 'MATLAB:recursionLimit')
         ME = MException(['JSimon:', mfilename, ':RecursiveType'], ...
            '%s: Cannot create hash for recursive data type: %s', ...
            mfilename, class(Data));
      end
      throw(ME);
   end
end

% return;

% ******************************************************************************
function FuncKey = ConvertFuncHandle(FuncH)
%   The subfunction ConvertFuncHandle converts function_handles to a struct
%   using the Matlab function FUNCTIONS. The output of this function changes
%   with the Matlab version, such that DataHash(@sin) replies different hashes
%   under Matlab 6.5 and 2009a.
%   An alternative is using the function name and name of the file for
%   function_handles, but this is not unique for nested or anonymous functions.
%   If the MATLABROOT is removed from the file's path, at least the hash of
%   Matlab's toolbox functions is (usually!) not influenced by the version.
%   Finally I'm in doubt if there is a unique method to hash function handles.
%   Please adjust the subfunction ConvertFuncHandles to your needs.

% The Matlab version influences the conversion by FUNCTIONS:
% 1. The format of the struct replied FUNCTIONS is not fixed,
% 2. The full paths of toolbox function e.g. for @mean differ.
FuncKey = functions(FuncH);

% Include modification file time and file size. Suggested by Aslak Grinsted:
if ~isempty(FuncKey.file)
    d = dir(FuncKey.file);
    if ~isempty(d)
        FuncKey.filebytes = d.bytes;
        FuncKey.filedate  = d.datenum;
    end
end

% ALTERNATIVE: Use name and path. The <matlabroot> part of the toolbox functions
% is replaced such that the hash for @mean does not depend on the Matlab
% version.
% Drawbacks: Anonymous functions, nested functions...
% funcStruct = functions(FuncH);
% funcfile   = strrep(funcStruct.file, matlabroot, '<MATLAB>');
% FuncKey    = uint8([funcStruct.function, ' ', funcfile]);

% Finally I'm afraid there is no unique method to get a hash for a function
% handle. Please adjust this conversion to your needs.

% return;

% ******************************************************************************
function DataBin = ConvertObject(DataObj)
% Convert a user-defined object to a binary stream. There cannot be a unique
% solution, so this part is left for the user...

try    % Perhaps a direct conversion is implemented:
   DataBin = uint8(DataObj);
   
   % Matt Raum had this excellent idea - unfortunately this function is
   % undocumented and might not be supported in te future:
   % DataBin = getByteStreamFromArray(DataObj);
   
catch  % Or perhaps this is better:
   WarnS   = warning('off', 'MATLAB:structOnObject');
   DataBin = struct(DataObj);
   warning(WarnS);
end

% return;

% ******************************************************************************
function Out = fBase64_enc(In)
% Encode numeric vector of UINT8 values to base64 string.
% The intention of this is to create a shorter hash than the HEX format.
% Therefore a padding with '=' characters is omitted on purpose.

Pool = [65:90, 97:122, 48:57, 43, 47];  % [0:9, a:z, A:Z, +, /]
v8   = [128; 64; 32; 16; 8; 4; 2; 1];
v6   = [32, 16, 8, 4, 2, 1];

In  = reshape(In, 1, []);
X   = rem(floor(In(ones(8, 1), :) ./ v8(:, ones(length(In), 1))), 2);
Y   = reshape([X(:); zeros(6 - rem(numel(X), 6), 1)], 6, []);
Out = char(Pool(1 + v6 * Y));

% return;

% ******************************************************************************
function Ex = FileExist_L(FileName)
% A more reliable version of EXIST(FileName, 'file'):
dirFile = dir(FileName);
if length(dirFile) == 1
   Ex = ~(dirFile.isdir);
else
   Ex = false;
end

% return;

% ******************************************************************************
function R = Version_L()
% The output differs between versions of this function. So give the user a
% chance to recognize the version:
% 1: 01-May-2011, Initial version
% 2: 15-Feb-2015, The number of dimensions is considered in addition.
%    In version 1 these variables had the same hash:
%    zeros(1,1) and zeros(1,1,0), complex(0) and zeros(1,1,0,0)
% 3: 29-Jun-2015, Struct arrays are processed field by field and not element
%    by element, because this is much faster. In consequence the hash value
%    differs, if the input contains a struct.
% 4: 28-Feb-2016 15:20, same output as GetMD5 for MD5 sums. Therefore the
%    dimensions are casted to UINT64 at first.
R.HashVersion = 4;
R.Date        = [2016, 2, 28];

R.HashMethod  = {};
try
   Provider = java.security.Security.getProviders;
   for iProvider = 1:numel(Provider)
      S     = char(Provider(iProvider).getServices);
      Index = strfind(S, 'MessageDigest.');
      for iDigest = 1:length(Index)
         Digest       = strtok(S(Index(iDigest):end));
         Digest       = strrep(Digest, 'MessageDigest.', '');
         R.HashMethod = cat(2, R.HashMethod, {Digest});
      end
   end
catch ME
   fprintf(2, '%s\n', ME.message);
   R.HashMethod = 'error';
end

% return;

% ******************************************************************************
function Error_L(ID, varargin)

error(['JSimon:', mfilename, ':', ID], ['*** %s: ', varargin{1}], ...
   mfilename, varargin{2:nargin - 1});

% return;
```
