## Supplemental File 1 for "A biophysical model of striatal microcircuits suggests delta/theta-rhythmically interleaved gamma and beta oscillations mediate periodicity in motor control": displayError.m

```
function displayError(err)
%DISPLAYERROR - Wrapper to print MATLAB error
  fprintf('Error: %s\n',err.message);
  for i=1:length(err.stack)
    fprintf('\t in %s (line %g)\n',err.stack(i).name,err.stack(i).line);
  end
```
