## Supplemental File 1 for "A biophysical model of striatal microcircuits suggests delta/theta-rhythmically interleaved gamma and beta oscillations mediate periodicity in motor control": distinguishable_colors.m

```
function colors = distinguishable_colors(n_colors,bg,func)
% DISTINGUISHABLE_COLORS: pick colors that are maximally perceptually distinct
%
% When plotting a set of lines, you may want to distinguish them by color.
% By default, Matlab chooses a small set of colors and cycles among them,
% and so if you have more than a few lines there will be confusion about
% which line is which. To fix this problem, one would want to be able to
% pick a much larger set of distinct colors, where the number of colors
% equals or exceeds the number of lines you want to plot. Because our
% ability to distinguish among colors has limits, one should choose these
% colors to be "maximally perceptually distinguishable."
%
% This function generates a set of colors which are distinguishable
% by reference to the "Lab" color space, which more closely matches
% human color perception than RGB. Given an initial large list of possible
% colors, it iteratively chooses the entry in the list that is farthest (in
% Lab space) from all previously-chosen entries. While this "greedy"
% algorithm does not yield a global maximum, it is simple and efficient.
% Moreover, the sequence of colors is consistent no matter how many you
% request, which facilitates the users' ability to learn the color order
% and avoids major changes in the appearance of plots when adding or
% removing lines.
%
% Syntax:
%   colors = distinguishable_colors(n_colors)
% Specify the number of colors you want as a scalar, n_colors. This will
% generate an n_colors-by-3 matrix, each row representing an RGB
% color triple. If you don't precisely know how many you will need in
% advance, there is no harm (other than execution time) in specifying
% slightly more than you think you will need.
%
%   colors = distinguishable_colors(n_colors,bg)
% This syntax allows you to specify the background color, to make sure that
% your colors are also distinguishable from the background. Default value
% is white. bg may be specified as an RGB triple or as one of the standard
% "ColorSpec" strings. You can even specify multiple colors:
%     bg = {'w','k'}
% or
%     bg = [1 1 1; 0 0 0]
% will only produce colors that are distinguishable from both white and
% black.
%
%   colors = distinguishable_colors(n_colors,bg,rgb2labfunc)
% By default, distinguishable_colors uses the image processing toolbox's
% color conversion functions makecform and applycform. Alternatively, you
% can supply your own color conversion function.
%
% Example:
%   c = distinguishable_colors(25);
%   figure
%   image(reshape(c,[1 size(c)]))
%
% Example using the file exchange's 'colorspace':
%   func = @(x) colorspace('RGB->Lab',x);
%   c = distinguishable_colors(25,'w',func);

% Copyright 2010-2011 by Timothy E. Holy

  % Parse the inputs
  if (nargin < 2)
    bg = [1 1 1];  % default white background
  else
    if iscell(bg)
      % User specified a list of colors as a cell aray
      bgc = bg;
      for i = 1:length(bgc)
	bgc{i} = parsecolor(bgc{i});
      end
      bg = cat(1,bgc{:});
    else
      % User specified a numeric array of colors (n-by-3)
      bg = parsecolor(bg);
    end
  end
  
  % Generate a sizable number of RGB triples. This represents our space of
  % possible choices. By starting in RGB space, we ensure that all of the
  % colors can be generated by the monitor.
  n_grid = 30;  % number of grid divisions along each axis in RGB space
  x = linspace(0,1,n_grid);
  [R,G,B] = ndgrid(x,x,x);
  rgb = [R(:) G(:) B(:)];
  if (n_colors > size(rgb,1)/3)
    error('You can''t readily distinguish that many colors');
  end
  
  % Convert to Lab color space, which more closely represents human
  % perception
  if (nargin > 2)
    lab = func(rgb);
    bglab = func(bg);
  else
    C = makecform('srgb2lab');
    lab = applycform(rgb,C);
    bglab = applycform(bg,C);
  end

  % If the user specified multiple background colors, compute distances
  % from the candidate colors to the background colors
  mindist2 = inf(size(rgb,1),1);
  for i = 1:size(bglab,1)-1
    dX = bsxfun(@minus,lab,bglab(i,:)); % displacement all colors from bg
    dist2 = sum(dX.^2,2);  % square distance
    mindist2 = min(dist2,mindist2);  % dist2 to closest previously-chosen color
  end
  
  % Iteratively pick the color that maximizes the distance to the nearest
  % already-picked color
  colors = zeros(n_colors,3);
  lastlab = bglab(end,:);   % initialize by making the "previous" color equal to background
  for i = 1:n_colors
    dX = bsxfun(@minus,lab,lastlab); % displacement of last from all colors on list
    dist2 = sum(dX.^2,2);  % square distance
    mindist2 = min(dist2,mindist2);  % dist2 to closest previously-chosen color
    [~,index] = max(mindist2);  % find the entry farthest from all previously-chosen colors
    colors(i,:) = rgb(index,:);  % save for output
    lastlab = lab(index,:);  % prepare for next iteration
  end
end

function c = parsecolor(s)
  if ischar(s)
    c = colorstr2rgb(s);
  elseif isnumeric(s) && size(s,2) == 3
    c = s;
  else
    error('MATLAB:InvalidColorSpec','Color specification cannot be parsed.');
  end
end

function c = colorstr2rgb(c)
  % Convert a color string to an RGB value.
  % This is cribbed from Matlab's whitebg function.
  % Why don't they make this a stand-alone function?
  rgbspec = [1 0 0;0 1 0;0 0 1;1 1 1;0 1 1;1 0 1;1 1 0;0 0 0];
  cspec = 'rgbwcmyk';
  k = find(cspec==c(1));
  if isempty(k)
    error('MATLAB:InvalidColorString','Unknown color string.');
  end
  if k~=3 || length(c)==1,
    c = rgbspec(k,:);
  elseif length(c)>2,
    if strcmpi(c(1:3),'bla')
      c = [0 0 0];
    elseif strcmpi(c(1:3),'blu')
      c = [0 0 1];
    else
      error('MATLAB:UnknownColorString', 'Unknown color string.');
    end
  end
end
```
