## Supplemental File 1 for "A biophysical model of striatal microcircuits suggests delta/theta-rhythmically interleaved gamma and beta oscillations mediate periodicity in motor control": aschar.m

```
function str = aschar(obj)
%% aschar
% Author: Erik Roberts, 2018
%
% Purpose: give string representation of input suitable for using eval on
%
% TODO
%  > 2d cell
%  > 2d mat
%  > 1 el logical
%  struct array

if ischar(obj) && (sum( size(obj) ~= 1 ) <= 1)
  str = ['''' obj ''''];
elseif isstring(obj)
  str = ['"' char(obj) '"'];
elseif isnumeric(obj) && isscalar(obj)
  str = num2str(obj);
elseif isnumeric(obj) && ~isscalar(obj) && ndims(obj) <= 2
  str = num2str(obj);
  
  str(:, end+1) = ';';
  str(:, end+1) = ' ';
  
  % cat rows
  nRows = size(obj, 1);
  temp = str;
  str = '[';
  for i = 1:nRows
    str = [str temp(i, :)];
  end
  str(end-1:end) = []; % remove trailing '; '
  str = [str ']'];
  
  str = regexprep(str, '(\d)\s+', '$1, '); % replace whitespace with comma space
elseif islogical(obj) && isscalar(obj)
  if obj
    str = 'true;';
  else
    str = 'false';
  end
elseif isa(obj, 'function_handle')
  str = ['@' func2str(obj)];
elseif iscell(obj) && ndims(obj) <= 2
  nRows = size(obj, 1);
  nCols = size(obj, 2);
  
  str = '{';
  for i = 1:nRows
    for j = 1:nCols
      str = [str aschar(obj{i,j}) ', '];
    end
    str(end-1:end) = []; % remove trailing ', '
    
    str = [str '; '];
  end
  str(end-1:end) = []; % remove trailing '; '
  
  str = [str '}'];
elseif isstruct(obj) && all( size(obj) == 1 )
  str = 'struct(';
  
  flds = fieldnames(obj);
  for iFld = 1:length(flds)
    fld = flds{iFld};
    str = [str '''' fld ''',' aschar(obj.(fld)) ', '];
  end
  str(end-1:end) = []; % remove trailing ', '
  
  str = [str ')'];
else
  error('Unsupported datatype');
end

end
```
