## Supplemental File 1 for "A biophysical model of striatal microcircuits suggests delta/theta-rhythmically interleaved gamma and beta oscillations mediate periodicity in motor control": exist_mkdir.m

```
function exist_mkdir(directoryPath)
%% exist_mkdir
% Author: Erik Roberts
%
% Purpose: Makes dir if doesn't exist
%
% Usage: exist_mkdir(directoryPath)

if ~exist(directoryPath, 'dir')
  mkdir(directoryPath);
end

end
```
