## Supplemental File 1 for "A biophysical model of striatal microcircuits suggests delta/theta-rhythmically interleaved gamma and beta oscillations mediate periodicity in motor control": fileparts2.m

```
function [pathstr, name, ext] = fileparts2(file)
%% FILEPARTS2  fileparts without missing trailing slash bug
%
% Purpose: Fixes mixing trailing slash bug in fileparts. If path is a dir without
%           a trailing slash, will add the slash so fileparts gives expected pathstr.
%
% Usage: [pathstr, name, ext] = fileparts2(file)
%
% Explanation:
%   Given a directory structure: ./parent/child/
%
%   Problem:
%     fileparts(parent/child/) = parent/child % Expected
%     fileparts(parent/child) = parent % Bug
%
%   Solution:
%     fileparts2(parent/child/) = parent/child % Expected
%     fileparts2(parent/child) = parent/child % Fixed bug
%
% See also FILEPARTS
%
% Author: Erik Roberts

if isdir(file) && file(end) ~= filesep
  file(end+1) = filesep;
end

[pathstr, name, ext] = fileparts(file);

end
```
