## Supplemental File 1 for "A biophysical model of striatal microcircuits suggests delta/theta-rhythmically interleaved gamma and beta oscillations mediate periodicity in motor control": getAbsolutePath.m

```
function [pathOut, pathInAbsBool] = getAbsolutePath(pathIn)
%% getAbsolutePath
%
% Purpose: Get absolute path from given absolute or relative path
%
% Usage: pathOut = getAbsolutePath(pathIn)
%        [pathOut, pathInAbsBool] = getAbsolutePath(pathIn)
%
% Outputs:
%   pathOut: absolute path given input, pathIn
%   pathInAbsBool: logical whether pathIn is already an absolute path
%
% Author: Erik Roberts

if ((isunix || ismac) && pathIn(1) == filesep) || (ispc && pathIn(2) == ':')
  % absolute
  pathOut = pathIn;
  pathInAbsBool = true;
elseif pathIn(1) == '~'
  % relative to home
  [~,homePath] = system('echo $HOME');
  pathOut = fullfile(homePath(1:end-1), pathIn(2:end)); % need to remove return char
  pathInAbsBool = false;
else
  % relative
  pathOut = fullfile(pwd, pathIn);
  pathInAbsBool = false;
end

end
```
