## Supplemental File 1 for "A biophysical model of striatal microcircuits suggests delta/theta-rhythmically interleaved gamma and beta oscillations mediate periodicity in motor control": getCallerCode.m

```
function callerCode = getCallerCode(stackNumber)
%% getCallerCode
% Author: Erik Roberts
%
% Purpose: stores code from caller as string
%
% Usage: callerCode = getCallerCode()
%        callerCode = getCallerCode(stackNumber)
%
% Input (Optional)
%   stackNumber: integer of which level in the call stack to get code from. Top
%   level is defined as largest number. Top level called by default. Immediate
%   caller is 1.

stack = dbstack;
callerCode = fileread(stack(end).file); %top of stacks

end
```
