## Supplemental File 1 for "A biophysical model of striatal microcircuits suggests delta/theta-rhythmically interleaved gamma and beta oscillations mediate periodicity in motor control": getStackCode.m

```
function stackCode = getStackCode()
%% getStackCode
% Author: Erik Roberts
%
% Purpose: stores code from whole caller stack as cell array of strings.
% Immediate caller is defined as first cell. Higher levels are later cells.
%
% Usage: stackCode = getStackCode()

stack = dbstack;
files = {stack.file};
nFiles = length(files);
stackCode = cell(nFiles,1);
for iFile = 1:nFiles
  stackCode{iFile} = fileread(files{iFile}); %top of stacks
end

end
```
