## Supplemental File 1 for "A biophysical model of striatal microcircuits suggests delta/theta-rhythmically interleaved gamma and beta oscillations mediate periodicity in motor control": isassigned.m

```
function returnBool = isassigned(varStr)
%% isassigned
% Author: Erik Roberts
%
% Purpose: Check if variable exists and is not empty
%
% Usage: returnBool = isassigned(varStr)

  returnBool = evalin('caller', ['exist(''' varStr ''', ''var'')']) && ~isempty(evalin('caller', varStr));
  
end
```
