## Supplemental File 1 for "A biophysical model of striatal microcircuits suggests delta/theta-rhythmically interleaved gamma and beta oscillations mediate periodicity in motor control": isValidFigHandle.m

```
function validBool = isValidFigHandle(h)
%% isValidFigHandle
% Purpose: check if input exists and is a valid figure handle(s).
%
% Usage:
%   validBool = isValidFigHandle(handle)
%   validBool = isValidFigHandle(strForHandle)
%
% Input: string with handle name, or handle itself
%
% Note: will work with string containing any callable matlab indexing/object
%
% Author: Erik Roberts

if isempty(h)
  validBool = false;
  return
end

try
  if ischar(h) %string input
    validBool = evalin('caller', ['isvalid(' h ')']) || evalin('caller', ['isgraphics(' h ')']);
  elseif ismatrix(h) && numel(h) > 1
    validBool = zeros(1,length(h));
    for iH = 1:length(h)
      validBool(iH) = isValidFigHandle(h(iH));
    end
  else
    validBool = isvalid(h) || isgraphics(h);
  end
catch
  validBool = false;
end

end
```
