## Supplemental File 1 for "A biophysical model of striatal microcircuits suggests delta/theta-rhythmically interleaved gamma and beta oscillations mediate periodicity in motor control": lscell.m

```
function dirList = lscell(arg, removePathBool, relativePathBool)
%% lscell
% Author: Erik Roberts, 2018
%
% Purpose: returns cell matrix of strings with results from call to dir
%
% Usage: dirList = lscell()
%        dirList = lscell(arg)
%        dirList = lscell(arg, removePathBool)
%        dirList = lscell(arg, removePathBool, relativePathBool)
%
% Inputs (optional):
%   arg: argument to dir
%   removePathBool: logical whether to remove the path before the files/dirs (default=true)
%   relativePathBool: logical whether to convert absolute paths to relative paths (default=false)
%                     if removePathBool== true, this is ignored.
%
% Output:
%   dirList: cellstr list of arg contents from dir. Paths to folders never
%            end in trailing filesep, i.e. '/' or '\'.
%
% Tips: in order to search subdirectories, use the '**' glob character in the arg
%
% See also: DIR

% Dev Note: checked on Mac OS 10.12, Windows 10, Linux Mint with Matlab 2017b

% parse args
if ~nargin || isempty(arg)
  arg = '.';
end
if nargin < 2 || isempty(removePathBool)
  removePathBool = true; %defaults to true
end
if nargin < 3 || isempty(relativePathBool)
  relativePathBool = false; %defaults to false
end

% get dir contents
dirListS = dir(arg);

if isempty(dirListS)
  dirList = {};
  return
end

% remove first period
if strcmp(dirListS(1).name, '.')
  dirListS(1) = [];
end

% remove double period
if strcmp(dirListS(1).name, '..')
  dirListS(1) = [];
end

% convert struct to cellstr
dirList = strcat({dirListS.folder}, filesep, {dirListS.name});

% ensure column vector
dirList = dirList(:);

% remove extra cells with '..'
dirList(~cellfun(@isempty, regexp(dirList, '\.\.$'))) = [];

% remove trailing period and filesep from dirs
if isunix || ismac
  dirList = regexprep(dirList, '/\.$', '');
else
  dirList = regexprep(dirList, '\\\.$', '');
end

% remove duplicated absolute paths from glob
dirList = unique(dirList);

% dirList is cellstr with absolute paths

if relativePathBool && ~removePathBool
  regexStr = ['^' pwd filesep];
  if isunix || ismac
    dirList = regexprep(dirList, regexStr, '');
  else
    regexStr = strrep(regexStr, '\', '\\');
    dirList = regexprep(dirList, regexStr, '');
  end
end

% dirList is cellstr with absolute paths, or relative paths starting with name

if removePathBool
  dirList = cellfun(@removePath, dirList, 'Uni',0);
end
  
  % nested functions
  function thisFilename = removePath(thisPath)
    [~,name,ext] = fileparts(thisPath);
    thisFilename = [name,ext];
  end
end
```
