## Supplemental File 1 for "A biophysical model of striatal microcircuits suggests delta/theta-rhythmically interleaved gamma and beta oscillations mediate periodicity in motor control": mfileFnName.m

```
function fnName = mfileFnName
%% mfileFnName
%
% Purpose: Adds support for packages to mfilename. Will return the callable
%   function name including all namespaced prefixes.
%
% Usage: fnName = mfileFnName
%
% Author: Erik Roberts


filePath = evalin('caller', 'mfilename(''fullpath'');');

if ~isempty(strfind(filePath, '+'))
  startInd = regexp(filePath, '\+', 'start');
  packagePath = filePath( startInd(1):end);
  
  splitPath = strsplit(packagePath, filesep);
  
  reFilesep = filesep;
  if reFilesep == '\'
    reFilesep = [reFilesep reFilesep]; % escape slash for re
  end
  
  repSplitPath = regexprep(splitPath, '+(.+)', '$1\.');
  
  packageFile = strjoin(repSplitPath,'');
  
  [~,fnName] = fileparts(packageFile);
else
  [~,fnName] = fileparts(filePath);
end

end
```
