## Supplemental File 1 for "A biophysical model of striatal microcircuits suggests delta/theta-rhythmically interleaved gamma and beta oscillations mediate periodicity in motor control": mkdirSilent.m

```
function varargout = mkdirSilent(output_path,varargin)
% makes dir if doesn't exist, otherwise does nothing

suppress_output = 1;

if ~exist(output_path,'file')
  if ~suppress_output
    fprintf('Creating %s \n', output_path);
  end
  %system( ['mkdir ' output_path]);
  [varargout{1:nargout}] = mkdir(output_path,varargin{:});   % Outputs are supplied simply to suppress warning
  % message for existing folder.
else
  if ~suppress_output; fprintf('Folder %s already exists. Doing nothing.\n',output_path); end
end

end
```
