## Supplemental File 1 for "A biophysical model of striatal microcircuits suggests delta/theta-rhythmically interleaved gamma and beta oscillations mediate periodicity in motor control": rls.m

```
function fileList = rls(arg, relPathBool)
%% rls
% Author: Erik Roberts
%
% Purpose: recursive file list using unix 'find'
%
% Usage: fileList = rls(arg)
%        fileList = rls(arg, relPathBool)
%
% Inputs:
%   arg: argument string to unix 'find' program
%   relPathBool: logical whether to convert to relative paths

% default inputs
if nargin == 0
  arg = [];
end
if nargin < 2
  relPathBool = false;
end

if isunix || ismac
  [~, fileList] = system(['find ' arg]);
else
  fprintf('This function only works on linux, mac, and unix systems.\n')
  return
end

% convert from char array to cell array
fileList = strsplit(fileList);
if isempty(fileList{end})
  fileList(end) = [];
end

if relPathBool
  fileList = regexprep(fileList, pwd, '');
end

end
```
