## Supplemental File 1 for "A biophysical model of striatal microcircuits suggests delta/theta-rhythmically interleaved gamma and beta oscillations mediate periodicity in motor control": saveWithStack.m

```
function saveWithStack(varargin)
%% saveWithStack
% Author: Erik Roberts
%
% Purpose: performs save with args, and adds the code from each file in stack to
%   variable stackCode.
%
% Usage: saveWithStack(args)


%% save
args = cellfun(@(x) ['''' x ''''], varargin, 'UniformOutput', false);
args = strjoin(args,',');
evalin('caller', ['save(' args ')'])

%% add stackCode
stackCode = getStackCode;
filename = varargin{1};
fileIO = matfile(filename,'Writable', true);
fileIO.stackCode = stackCode;

end
```
