## Supplemental File 1 for "A biophysical model of striatal microcircuits suggests delta/theta-rhythmically interleaved gamma and beta oscillations mediate periodicity in motor control": thisMfileDir.m

```
function dirPath = thisMfileDir()
%% thisMfileDir
% Author: Erik Roberts
%
% Purpose: Gets current m-file dir
%
% Usage: dirPath = thisMfileDir()

stack = dbstack;
% dirPath = evalin('caller', 'fileparts(which(mfilename(''fullpath'')))');
dirPath = fileparts(which(stack(end).name));

end
```
