## Supplemental File 1 for "A biophysical model of striatal microcircuits suggests delta/theta-rhythmically interleaved gamma and beta oscillations mediate periodicity in motor control": wprintf.m

```
function wprintf(varargin)
%% wprintf
% Purpose: Warning print. Prints in red and makes beep. Separates with
%   whitespace. Formats str input with sprintf.
%
% Usage:
%   wprintf(str)
%   wprintf(str, args)
%
% Input: str - string for warning
%
% Output (to stdout): ['\nWarning: ' fortmattedStr '\n\n'].

str = sprintf(varargin{:});

fprintf(2, ['\nWarning: ' str '\n\n']);
beep

end
```
