## Supplemental File 1 for "A biophysical model of striatal microcircuits suggests delta/theta-rhythmically interleaved gamma and beta oscillations mediate periodicity in motor control": eksr.m

```
function r=eksr(x,y,h,N)
% EKSR Epanechnikov Kernel smoothing regression
%
% r=eksr(x,y) returns the Epanechnikov kernel regression in structure r such that
%   r.f(r.x) = y(x) + e
% The bandwidth and number of samples are also stored in r.h and r.n
% respectively.
%
% r=eksr(x,y,h) performs the regression using the specified bandwidth, h.
%
% r=eksr(x,y,h,n) calculates the regression in n points (default same as input).
%
% Without output, eksr(x,y) or eksr(x,y,h) will display the regression plot.
%
% Algorithm
% The kernel regression is a non-parametric approach to estimate the
% conditional expectation of a random variable:
%
% E(Y|X) = f(X)
%
% where f is a non-parametric function. Based on the kernel density
% estimation, this code implements the Nadaraya-Watson kernel regression
% using the Epanechnikov kernel as follows:
%
% f(x) = sum(kerf((x-X)/h).*Y)/sum(kerf((x-X)/h))
%
% See also ksr, gkde, ksdensity

% Example 1: smooth curve with noise
%{
x = 1:100;
y = sin(x/10)+(x/50).^2;
yn = y + 0.2*randn(1,100);
r=ksr(x,yn);
plot(x,y,'b-',x,yn,'co',r.x,r.f,'r--','linewidth',2)
legend('true','data','regression','location','northwest');
title('Gaussian kernel regression')
%}
% Example 2: with missing data
%{
x = sort(rand(1,100)*99)+1;
y = sin(x/10)+(x/50).^2;
y(round(rand(1,20)*100)) = NaN;
yn = y + 0.2*randn(1,100);
r=ksr(x,yn);
plot(x,y,'b-',x,yn,'co',r.x,r.f,'r--','linewidth',2)
legend('true','data','regression','location','northwest');
title('Gaussian kernel regression with 20% missing data')
%}

% By Yi Cao at Cranfield University on 12 March 2008.
%

% Check input and output
narginchk(2,4);
nargoutchk(0,1);
if numel(x)~=numel(y)
    error('x and y are in different sizes.');
end

x=x(:);
y=y(:);
% clean missing or invalid data points
inv=(x~=x)|(y~=y);
x(inv)=[];
y(inv)=[];

% Default parameters
if nargin<4
    % N=100;
    N = length(x);
elseif ~isscalar(N)
    error('N must be a scalar.')
end
r.n=length(x);
if nargin<3
    % optimal bandwidth suggested by Bowman and Azzalini (1997) p.31
    hx=median(abs(x-median(x)))/0.6745*(4/3/r.n)^0.2;
    hy=median(abs(y-median(y)))/0.6745*(4/3/r.n)^0.2;
    h=sqrt(hy*hx);
    if h<sqrt(eps)*N
        error('There is no enough variation in the data. Regression is meaningless.')
    end
elseif ~isscalar(h)
    error('h must be a scalar.')
end
r.h=h;

% Gaussian kernel function
% kerf = @(z)exp(-z.*z/2)/sqrt(2*pi);

% Epanechnikov kernel function
kerf = @(z)0.75*(1-z.*z);

r.x=linspace(min(x),max(x),N);
r.f=zeros(1,N);
for k=1:N
    z = kerf((r.x(k)-x)/h);
    z(z < 0) = 0;
    r.f(k) = sum(z.*y)/sum(z);
    % if k >= N/2 && k < N/2 +1
    %   figure,plot(z)
    % end
end

% Plot
if ~nargout
    figure
    plot(r.x,r.f,'r',x,y,'bo')
    ylabel('f(x)')
    xlabel('x')
    title('Epanechnikov Kernel Smoothing Regression');
end
```
