## Supplemental File 1 for "A biophysical model of striatal microcircuits suggests delta/theta-rhythmically interleaved gamma and beta oscillations mediate periodicity in motor control": fig2svg.m

```
function varargout = fig2svg(filename, id, debug, legendIcons, clippingMode, figureSize, pixelFileType, creditPrintBool)
  %  Matlab/GNU Octave FIG to SVG converter
  %
  %  Usage: fig2svg(filename, id, debug, legendIcons, clippingMode, figureSize, pixelFileType, creditPrintBool)
  %
  %         all arguments are optional
  %
  %         1. filename: name of the svg file (including extension)
  %         can be omitted (nargin < 1) or left empty ('' or []) and a system dialog will be prompted
  %
  %         2. id: graphic handle (if omitted or empty, current gcf is taken)
  %
  %         3. debug = 0 or 1 (enable to ease debugging of fig2svg functionality; 0 if omitted or empty)
  %
  %         4. legendIcons: legend pieces (necessary in new matlab versions to respect legend appearance)
  %         can be omitted (nargin < 4) or left empty ('' or [])
  %
  %         5. clippingMode = 0 (no clipping) or 1 (box on/off dependent clipping) or 2 (strict axes clipping) or 3 (axes+data dependent clipping)
  %         clippingMode is set to 1 if argument is omitted or empty
  %
  %         6. figureSize = [width,height] (actual figure size if omitted or empty)
  %
  %         7. pixelFileType = 'png' or 'jpg' (png used if omitted or empty)
  %
  %         8. creditPrintBool - logical argument for whether to display credit information (1 if omitted or empty)

  global FIG2SVG_globals
  global colorname
  release_version = '2019.01.3'; % year.month.incremental
  FIG2SVG_globals.runningIdNumber = 0;
  FIG2SVG_globals.UI = reportUI;
  FIG2SVG_globals.octave = false;
  FIG2SVG_globals.checkUserData = true;
  FIG2SVG_globals.ScreenPixelsPerInch = 96; % New default ppi
  FIG2SVG_globals.resolutionScaling = 1;
  FIG2SVG_globals.WN = 0;
  if nargin < 5 || isempty(clippingMode)
    FIG2SVG_globals.ClippingMode = 1; % nees revision -> note that it does not work fine with Inkscape pdf conversion for latex
  else
    FIG2SVG_globals.ClippingMode = clippingMode;
  end

  try
    % Rounding it to be an integer (Octave resolution is 96.024)
    FIG2SVG_globals.ScreenPixelsPerInch = round(get(0, 'ScreenPixelsPerInch'));
  catch
    % Keep the default ppi
  end
  % Octave resolution is 96.024, rounding it to 96
  FIG2SVG_globals.resolutionScaling = FIG2SVG_globals.ScreenPixelsPerInch/96;
  if nargout == 1
    varargout = {0};
  end
  if nargin < 8 || isempty(creditPrintBool) || creditPrintBool == 1
    disp(['   FIG to SVG converter version ' release_version ', maintained by Salva Ardid (https://github.com/kupiqu/fig2svg).'])
  end
  if strcmp(FIG2SVG_globals.UI,'octave')
    FIG2SVG_globals.octave = true;
  end
  if nargout > 1
    error('Function returns only one return value.')
  end

  if nargin < 2 || isempty(id) % Check if proper handle was included into function call, otherwise take current figure
    id = gcf;
  end
  if nargin < 3 || isempty(debug) % No debug mode
    FIG2SVG_globals.debugModeOn = 0;
  else
    FIG2SVG_globals.debugModeOn = debug;
  end

  f1 = id;
  objects = allchild(f1);
  if all(~ismember(get(objects,'Type'),'wordcloud'))
    % a nice way to keep the original figure safe and work on a temporary copy (does not work for wordclouds)
    copyfig = 1;
    if ~FIG2SVG_globals.octave
      warning('off', 'MATLAB:copyobj:ObjectNotCopied'); % these warnings don't seem to come from figure content
      xl = get(gca,'xlabel');
      yl = get(gca,'ylabel');
      zl = get(gca,'zlabel');
      tl = get(gca,'title');
      cmap = get(id,'Colormap');
    end
    if FIG2SVG_globals.debugModeOn % only show the copy in debug mode
      f2 = figure;
    else
      f2 = figure('visible', 'off');
    end
    try
      copyobj(get(f1,'children'),f2);
      if ~FIG2SVG_globals.octave
        paperpos = get(f1,'Position');
        set(f2,'Position',paperpos)
        if ~UIverlessthan('8.4.0')
          if ~isempty(xl.String)
           set(gca,'xlabel',xl)
          end
          if ~isempty(yl.String)
           set(gca,'ylabel',yl)
          end
          if ~isempty(zl.String)
           set(gca,'zlabel',zl)
          end
          if ~isempty(tl.String)
           set(gca,'title',tl)
          end
        else
          copyobj(xl,gca);
          copyobj(yl,gca);
          copyobj(zl,gca);
          copyobj(tl,gca);
        end
        id = f2;
        colormap(cmap);
      end
    catch
      fprintf('   Warning: Figure copy failed, fig2svg will proceed with the original figure.\n   Unnoticed rearrangements of the figure may occur. Caution must be taken.\n');
      copyfig = 0;
      close(f2);
    end
    warning('on', 'MATLAB:copyobj:ObjectNotCopied'); % restoring the warning after copying
  else
    copyfig = 0;
  end

  if nargin < 1 || isempty(filename)
    [filename, pathname] = uiputfile( {'*.svg', 'SVG File (*.svg)'},'Save Figure as SVG File');
    if ~( isequal( filename, 0) || isequal( pathname, 0))
      % yes. add backslash to path (if not already there)
      pathname = addBackSlash( pathname);
      % check, if extension is allrigth
      if ( ~strcmpi( getFileExtension( filename), '.svg'))
        filename = [ filename, '.svg'];
      end
      finalname = [pathname filename];
    else
      disp('   Cancelled as requested.')
      return
    end
  else
    finalname = filename;
  end

  if nargin < 4 % No legend icons passed
    legendIcons = [];
  end

  % needed to see annotation axes
  originalShowHiddenHandles = get(0, 'ShowHiddenHandles');
  set(0, 'ShowHiddenHandles', 'on');
  originalFigureUnits = get(id,'Units');
  set(id,'Units','pixels');   % All data in the svg-file is saved in pixels
  paperpos = get(id,'Position');
  if nargin >= 6 && ~isempty(figureSize)
    paperpos(3) = figureSize(1);
    paperpos(4) = figureSize(2);
  end
  paperpos = convertunit(paperpos, 'pixels', 'pixels');
  if (nargin < 7) || isempty(pixelFileType)
    FIG2SVG_globals.pixelFileType = 'png';
  else
    FIG2SVG_globals.pixelFileType = pixelFileType;
  end
  cmap = get(id,'Colormap');
  colorname = '';
  for i = 1:size(cmap,1)
    colorname(i,:) = sprintf('%02x%02x%02x',fix(cmap(i,1)*255),fix(cmap(i,2)*255),fix(cmap(i,3)*255));
  end

  % Open SVG-file
  [pathstr,name] = fileparts(finalname);
  %FIG2SVG_globals.basefilename = fullfile(pathstr,name);
  FIG2SVG_globals.basefilepath = pathstr;
  FIG2SVG_globals.basefilename = name;
  FIG2SVG_globals.figurenumber = 1;
  fid = fopen(finalname,'wt');   % Create a new text file
  fprintf(fid,'<?xml version = "1.0" encoding = "utf-8" standalone = "no"?><!DOCTYPE svg PUBLIC "-//W3C//DTD SVG 1.1//EN" "http://www.w3.org/Graphics/SVG/1.1/DTD/svg11.dtd">\n');    % Insert file header
  % Making transparent by default
  % fprintf(fid,'<svg preserveAspectRatio = "xMinYMin meet" width = "100%%" height = "100%%" fill = "#ffffff" viewBox = "0 0 %0.3f %0.3f" ',paperpos(3),paperpos(4));
  fprintf(fid,'<svg preserveAspectRatio = "xMinYMin meet" width = "100%%" height = "100%%" fill = "none" viewBox = "0 0 %0.3f %0.3f" ',paperpos(3),paperpos(4));
  fprintf(fid,' version = "1.1" xmlns = "http://www.w3.org/2000/svg" xmlns:xlink = "http://www.w3.org/1999/xlink"');
  %fprintf(fid,' onload = "Init(evt)"');
  fprintf(fid,'>\n');
  fprintf(fid,'  <desc>Matlab Figure Converted by FIG2SVG</desc>\n');
  %fprintf(fid,'  <script type = "text/ecmascript" xlink:href = "puzzle_script.js" />\n');
  fprintf(fid,'  <g id = "topgroup">\n');
  group = 1;
  groups = [];
  % Frame of figure
  figcolor = searchcolor(id,get(id, 'Color'));
  if ~strcmp(figcolor, 'none')
    % Draw rectangle in the background of the graphic frame to cover all
    % other graphic elements
    if strcmp(get(id,'InvertHardcopy'),'on')
      fprintf(fid,'  <rect x = "0" y = "0" width = "%0.3f" height = "%0.3f" fill = "#ffffff" stroke = "none" />\n',paperpos(3),paperpos(4));
    else
      fprintf(fid,'  <rect x = "0" y = "0" width = "%0.3f" height = "%0.3f" fill = "%s" stroke = "none" />\n',paperpos(3),paperpos(4),figcolor);
    end
  end
  % Search all axes
  ax = get(id,'Children');
  contLegend = 0;
  for j = length(ax):-1:1
    currentType = get(ax(j),'Type');
    if strcmp(currentType,'axes') || strcmp(currentType,'bar') || strcmp(currentType,'scatter')
      if FIG2SVG_globals.debugModeOn
        disp(['ax(',num2str(j),') = ', currentType]);
      end
      groups = [groups group];
      if ~FIG2SVG_globals.octave
        axYAXIS = get(ax(j),'YAxis');
        if numel(axYAXIS) == 2 % yyaxis
          yyaxis left;
          group = axes2svg(fid,id,ax(j),group,paperpos);
          yyaxis right;
          set(ax(j),'color','none'); % so it doesn't hide the left content
          group = axes2svg(fid,id,ax(j),group,paperpos);
        else
          group = axes2svg(fid,id,ax(j),group,paperpos);
        end
      else
        group = axes2svg(fid,id,ax(j),group,paperpos);
      end
    elseif strcmp(currentType,'colorbar')
      if FIG2SVG_globals.debugModeOn
        disp(['ax(',num2str(j),') = ', currentType]);
      end
      groups = [groups group];
      group = colorbar_axes2svg(fid,id,ax(j),group,paperpos);
    elseif strcmp(currentType,'legend')
      contLegend = contLegend + 1;
      legendVisible = get(ax(j),'Visible');
      if strcmp(legendVisible,'on')
        if ~UIverlessthan('9.1.0') % Not defined before Matlab 2016b
          set(ax(j),'AutoUpdate','off'); % sometimes weird things happen with legends including patches (e.g., bar graphs) if not used
        end
        legendPosition = get(ax(j),'Position');

        x = legendPosition(1)*paperpos(3);
        w = legendPosition(3)*paperpos(3);
        y = (1-(legendPosition(2)+legendPosition(4)))*paperpos(4);
        h = legendPosition(4)*paperpos(4);
        legendBoundingBox = [x y w h];

        legendLabels = get(ax(j),'String');
        legendBox = get(ax(j),'Box');
        legendColor = get(ax(j),'Color');
        legendEdgeColor = get(ax(j),'EdgeColor');
        legendFontSize = get(ax(j),'FontSize');
        legendLineWidth = get(ax(j),'LineWidth');
        legendLocation = get(ax(j),'Location');
        legendOrientation = get(ax(j),'Orientation');
        % The following is a trick to automatically get the distinct parts of a legend. However, there are some Matlab limitations, so for a proper legend, it's strongly encouraged to pass the legend icons to fig2svg (see warning and explanations below)
        if isempty(legendIcons)
          % note though that when only a subset of the graph appears in the legend, this trick selects, with the exception of text, not the proper subset of the graph, but the first items that were plotted (Matlab's fault).
          % one way to workaround the issue is to just plot things in order in the original graph, as they appear in the legend, but that's not always optimal, continue reading:
          % an additional problem comes from lines appearing under patches, which could be fixed by adding zdata (different layers of depth in the xy projection from the z-axis), but this solution is only expected to work in svg 2, once z index is introduced (limitation of current svg implementation).
          % there are two possible solutions to this:
          % 1. Encouraged 'fix': pass the legend icons to fig2svg (see below to know how)
          % 2. Discouraged 'fix': plot those lines twice, before the patch, so they appear in the legend (matlab's fault), and after the patch, so they appear on top (svg 1.1 limitation).
          warning(sprintf('\n\nFor proper results when lines and patches are present in a figure, pass the legend icons to fig2svg function as a fourth argument, e.g., fig2svg(''filename.svg'','''','''',legendIcons)\nLegend icons can be grabbed by calling legend in the following way: [lgd,legendIcons] = legend(...);\nIf more than a legend is present within a figure (e.g., in different subplots) pass the legend icons in a cell array.\n'));
          [lgd,legendIcons{contLegend}] = legend(legendLabels,'Location',legendLocation,'Orientation',legendOrientation,'FontSize',legendFontSize,'LineWidth',legendLineWidth,'Color',legendColor,'EdgeColor',legendEdgeColor,'Box',legendBox);
          ax(j) = lgd;
        elseif ~iscell(legendIcons)
          tmp_legendIcons{contLegend} = legendIcons;
          legendIcons = tmp_legendIcons;
        end
        if FIG2SVG_globals.debugModeOn
          for k = numel(legendIcons{contLegend}):-1:1
            iconType = get(legendIcons{contLegend}(k),'Type');
            disp(['legend(',num2str(k),') = ', iconType]);
          end
        end
        legendGroupax = 1;
        projection = []; % not used
        fprintf(fid,'  <g id  = "%s">\n', createId);
        legendIdString = createId;
        fprintf(fid,'  <clipPath id = "%s">\n',legendIdString);
        fprintf(fid,'    <rect x = "%0.3f" y = "%0.3f" width = "%0.3f" height = "%0.3f"/>\n',...
          legendBoundingBox(1), legendBoundingBox(2), legendBoundingBox(3), legendBoundingBox(4));
        fprintf(fid,'  </clipPath>\n');
        if strcmp(legendBox,'on')
          scolorname = searchcolor(id,legendEdgeColor);
          fcolorname = searchcolor(id,legendColor);
          fprintf(fid,'  <g id = "%s">\n',createId);
          fprintf(fid,'    <rect fill = "%s" stroke-linecap = "square" stroke-linejoin = "miter" stroke = "%s" stroke-width = "%0.3fpt" stroke-dasharray = "none" stroke-opacity = "1.000" x = "%0.3f" y = "%0.3f" width = "%0.3f" height = "%0.3f"/>\n',...
            fcolorname, scolorname, legendLineWidth, legendBoundingBox(1), legendBoundingBox(2), legendBoundingBox(3), legendBoundingBox(4));
          fprintf(fid,'  </g>\n');
        end
        axchild2svg(fid,id,legendIdString,ax(j),paperpos,legendIcons{contLegend},legendPosition,legendGroupax,projection,legendBoundingBox);
        fprintf(fid,'  </g>\n');
      end
    elseif strcmp(currentType,'uicontrol')
      if strcmp(get(ax(j),'Visible'),'on')
        control2svg(fid,id,ax(j),paperpos);
      end
    elseif strcmp(currentType, 'uicontextmenu') || ...
      strcmp(currentType, 'uimenu') || ...
      strcmp(currentType, 'hgjavacomponent') || ...
      strcmp(currentType, 'uitoolbar')
      % ignore these types
    elseif strcmp(currentType,'annotationpane')
      if FIG2SVG_globals.debugModeOn
        disp(['ax(',num2str(j),') = ', currentType]);
      end
      groups = [groups group];
      group = annotation2svg(fid,id,ax(j),group,paperpos);
    else
      disp(['   Warning: Unhandled main figure child type: ' currentType]);
    end
  end
  fprintf(fid,'  </g>\n');
  fprintf(fid,'</svg>\n');
  fclose(fid);    % close text file
  if nargout == 1
    varargout = {0};
  end
  if copyfig && ~FIG2SVG_globals.octave
    % set(id,'Units',originalFigureUnits);
    % set(0, 'ShowHiddenHandles', originalShowHiddenHandles);
    set(0,'CurrentFigure',f1)
    if ~UIverlessthan('8.4.0')
      if ~isempty(xl.String)
       set(gca,'xlabel',xl)
      end
      if ~isempty(yl.String)
       set(gca,'ylabel',yl)
      end
      if ~isempty(zl.String)
       set(gca,'zlabel',zl)
      end
      % weird but title does not come back unless I do this:
      copyobj(tl,gca);
      % if ~isempty(tl.String)
      %  set(gca,'title',tl)
      % end
    end
    if ~FIG2SVG_globals.debugModeOn % close the temporary copies
      close(f2);
    else
      set(0,'CurrentFigure',f2)
      if ~UIverlessthan('8.4.0')
        % in new matlab versions, these look like the same but don't behave internally like axes' labels, so that's why I moved the actual axes' labels from the original to the copied figure, and then back again, once the svg file was complete (here I just copy them for visual debugging)
        copyobj(xl,gca);
        copyobj(yl,gca);
        copyobj(zl,gca);
        copyobj(tl,gca);
      end
    end
  end
end

function clippingIdString = clipping2svg(fid, id, ax, paperpos, axpos, projection, clippingIdString)
  global FIG2SVG_globals
  if FIG2SVG_globals.checkUserData && isstruct(get(id,'UserData'))
      struct_data = get(id,'UserData');
      if isfield(struct_data,'svg')
          if isfield(struct_data.svg,'ClippingPath')
              clip = struct_data.svg.ClippingPath;
              if ~isempty(clip)
                  if size(clip, 2) ~= 3
                      if size(clip, 2) == 2
                          clipx = clip(:, 1);
                          clipy = clip(:, 2);
                          clipz = zeros(size(clip,1),1);
                      else
                          error('The clipping vector has to be a nx3 or nx2 matrix.');
                      end
                  else
                      clipx = clip(:, 1);
                      clipy = clip(:, 2);
                      clipz = clip(:, 3);
                  end
                  if strcmp(get(ax,'XScale'),'log')
                      clipx(clipx <= 0) = NaN;
                      clipx = log10(clipx);
                  end
                  if strcmp(get(ax,'YScale'),'log')
                      clipy(clipy <= 0) = NaN;
                      clipy = log10(clipy);
                  end
                  if strcmp(get(ax,'ZScale'),'log')
                      clipz(clipz <= 0) = NaN;
                      clipz = log10(clipz);
                  end
                  [x,y,~] = project(clipx,clipy,clipz,projection);
                  x = (x*axpos(3)+axpos(1))*paperpos(3);
                  y = (1-(y*axpos(4)+axpos(2)))*paperpos(4);
                  clippingIdString = createId;
                  fprintf(fid,'<clipPath id = "%s">\n  <polygon fill = "none" stroke = "none" points = "', clippingIdString);
                  fprintf(fid,'%0.3f,%0.3f ',[x';y']);
                  fprintf(fid,'"/>\n</clipPath>\n');
              end
          end
      end
  end
end

function [angle, align] = improvedXLabel(id, angle, align)
  global FIG2SVG_globals
  try
    angle = get(id,'XTickLabelRotation');
  catch
    if FIG2SVG_globals.checkUserData && isstruct(get(id,'UserData'))
      struct_data = get(id,'UserData');
      if isfield(struct_data,'svg')
        if isfield(struct_data.svg,'XTickLabelAngle')
          angle = struct_data.svg.XTickLabelAngle;
          % align = 'left';
        end
      end
    end
  end
end

function [angle, align] = improvedYLabel(id, angle, align)
  global FIG2SVG_globals
  try
    angle = get(id,'YTickLabelRotation');
  catch
    if FIG2SVG_globals.checkUserData && isstruct(get(id,'UserData'))
      struct_data = get(id,'UserData');
      if isfield(struct_data,'svg')
        if isfield(struct_data.svg,'YTickLabelAngle')
          angle = struct_data.svg.YTickLabelAngle;
          % align = 'left';
        end
      end
    end
  end
end

function [angle, align] = improvedZLabel(id, angle, align)
  global FIG2SVG_globals
  try
    angle = get(id,'ZTickLabelRotation');
  catch
    if FIG2SVG_globals.checkUserData && isstruct(get(id,'UserData'))
      struct_data = get(id,'UserData');
      if isfield(struct_data,'svg')
        if isfield(struct_data.svg,'ZTickLabelAngle')
          angle = struct_data.svg.ZTickLabelAngle;
          % align = 'left';
        end
      end
    end
  end
end

function animation2svg(fid, id)
  global FIG2SVG_globals
  if FIG2SVG_globals.checkUserData && isstruct(get(id,'UserData'))
      struct_data = get(id,'UserData');
      if isfield(struct_data,'svg')
          if isfield(struct_data.svg,'Animation')
              animation = struct_data.svg.Animation;
              for i = 1:length(animation)
                  if ~isfield(animation(i).SubAnimation, 'Type')
                      error('Missing field ''Type'' for animation.');
                  end
                  switch animation(i).SubAnimation.Type
                      case 'Opacity', type = 'opacity'; animationType = 0;
                      case 'Translate', type = 'translate'; animationType = 2;
                      case 'Scale', type = 'scale'; animationType = 2;
                      case 'Rotate', type = 'rotate'; animationType = 1;
                      case 'skewX', type = 'skewX'; animationType = 1;
                      case 'skewY', type = 'skewY'; animationType = 1;
                      otherwise, error(['Unknown animation type ''' animation(i).SubAnimation.Type '''.']);
                  end
                  %fprintf(fid,'  <animate attributeType = "XML" attributeName = "%s" from = "%0.3f" to = "%0.3f" dur = "%0.3fs" repeatCount = "%s" />',...
                  %   'opacity' , 0, 1, 5, 'indefinite');
                  if animationType == 0
                      fprintf(fid,'  <animate attributeType = "XML" attributeName = "%s" dur = "%0.3fs"', type, animation(i).SubAnimation.Duration);
                      fprintf(fid,' values = "');
                      fprintf(fid,'%0.3f;', animation(i).SubAnimation.Value);
                      fprintf(fid,'" keyTimes = "');
                      fprintf(fid,'%0.3f;', max(min(animation(i).SubAnimation.Key, 1), 0));
                      fprintf(fid,'" repeatCount = "%s" calcMode = "linear" />', 'indefinite');
                  elseif animationType == 1
                      fprintf(fid,'  <animateTransform attributeName = "transform" attributeType = "XML" type = "%s" dur = "%0.3fs"', type, animation(i).SubAnimation.Duration);
                      fprintf(fid,' values = "');
                      fprintf(fid,'%0.3f;', animation(i).SubAnimation.Value);
                      fprintf(fid,'" keyTimes = "');
                      fprintf(fid,'%0.3f;', max(min(animation(i).SubAnimation.Key, 1), 0));
                      fprintf(fid,'" repeatCount = "%s" calcMode = "linear" additive = "sum" />', 'indefinite');
                  elseif animationType == 2
                      fprintf(fid,'  <animateTransform attributeName = "transform" attributeType = "XML" type = "%s" dur = "%0.3fs"', type, animation(i).SubAnimation.Duration);
                      fprintf(fid,' values = "');
                      fprintf(fid,'%0.3f,%0.3f;', animation(i).SubAnimation.Value);
                      fprintf(fid,'" keyTimes = "');
                      fprintf(fid,'%0.3f;', max(min(animation(i).SubAnimation.Key, 1), 0));
                      fprintf(fid,'" repeatCount = "%s" calcMode = "linear" additive = "sum" />', 'indefinite');
                  end
              end
          end
      end
  end
end

function [filterString, boundingBox] = filter2svg(fid, id, boundingBoxAxes, boundingBoxElement)
  global FIG2SVG_globals
  filterString = '';
  boundingBox = boundingBoxAxes;
  if FIG2SVG_globals.checkUserData && isstruct(get(id,'UserData'))
      struct_data = get(id,'UserData');
      if isfield(struct_data,'svg')
          boundingBox = boundingBoxElement;
          absolute = true;
          % offset = 0;
          if isfield(struct_data.svg,'BoundingBox')
              if isfield(struct_data.svg.BoundingBox, 'Type')
                  switch struct_data.svg.BoundingBox.Type
                      case 'axes', boundingBox = boundingBoxAxes; absolute = true;
                      case 'element', boundingBox = boundingBoxElement; absolute = true;
                      case 'relative', boundingBox = boundingBoxElement; absolute = false;
                      otherwise
                          error(['Unknown bounding box type ''' struct_data.svg.BoundingBox.Type '''.']);
                  end
              end
              if isfield(struct_data.svg.BoundingBox, 'Overlap')
                  overlap = struct_data.svg.BoundingBox.Overlap;
                  if absolute
                      boundingBox(1) = boundingBox(1) - overlap;
                      boundingBox(2) = boundingBox(2) - overlap;
                      boundingBox(3) = boundingBox(3) + 2 * overlap;
                      boundingBox(4) = boundingBox(4) + 2 * overlap;
                  else
                      boundingBox(1) = boundingBox(1) - boundingBox(3) * overlap;
                      boundingBox(2) = boundingBox(2) - boundingBox(4) * overlap;
                      boundingBox(3) = boundingBox(3) + 2 * boundingBox(3) * overlap;
                      boundingBox(4) = boundingBox(4) + 2 * boundingBox(4) * overlap;
                  end
              end
              if isfield(struct_data.svg.BoundingBox, 'Visible') && strcmp(struct_data.svg.BoundingBox.Visible, 'on')
                  % This functionality is very interesting for debugging of
                  % bounding boxes of filters
                  fprintf(fid,'<rect x = "%0.3f" y = "%0.3f" width = "%0.3f" height = "%0.3f" fill = "none" stroke-linecap = "square" stroke-linejoin = "miter" stroke = "#000000" stroke-dasharray = "1,1" stroke-width = "0.2pt" />\n', boundingBox(1), boundingBox(2), boundingBox(3), boundingBox(4));
              end
          end
          if isfield(struct_data.svg,'Filter')
              % Predefined filter sources. Additional filter sources will be
              % added later.
              predefinedSources = {'SourceGraphic','SourceAlpha','BackgroundImage','BackgroundAlpha','FillPaint','StrokePaint'};
              resultStrings = predefinedSources;
              filterId = createId;
              filterString  = ['filter = "url(#' filterId ')"'];
              fprintf(fid,'<defs>\n');
              fprintf(fid,'  <filter x = "%0.3f%%" y = "%0.3f%%" width = "%0.3f%%" height = "%0.3f%%" id = "%s">\n', 0, 0, 100, 100, filterId);
              %if absolute
              %    fprintf(fid,'  <filter x = "%0.3f" y = "%0.3f" width = "%0.3f" height = "%0.3f" filterUnits = "userSpaceOnUse" id = "%s">\n', boundingBox(1), boundingBox(2), boundingBox(3), boundingBox(4), filterId);
              %else
              %    fprintf(fid,'  <filter x = "%0.3f%%" y = "%0.3f%%" width = "%0.3f%%" height = "%0.3f%%" id = "%s">\n', -(offset * 100), -(offset * 100), 100 + (offset * 200), 100 + (offset * 200), filterId);
              %    % Note: use default -10% for attribute x
              %    %       use default -10% for attribute y
              %    %       use default 120% for attribute width
              %    %       use default 120% for attribute height
              %end
              filter = struct_data.svg.Filter;
              for i = 1:length(filter)
                  if isfield(filter(i).Subfilter, 'Type')
                      fprintf(fid,'  <%s', filter(i).Subfilter.Type);
                  else
                      error('Missing field ''Type'' for filter.')
                  end
                  try
                      if isfield(filter(i).Subfilter, 'Position')
                          printAttributeArray(fid, {'x','y','width','height'}, filter(i).Subfilter, 'Position');
                      end
                      printAttributeString(fid, 'result', filter(i).Subfilter, 'Result');
                      % Add result string to the list in order to check the in
                      % strings.
                      resultStrings{length(resultStrings) + 1} = filter(i).Subfilter.Result;
                      % The strmatch below is a very inefficient search (Matlab limitation)
                      if ~isempty(strmatch(filter(i).Subfilter.Result, predefinedSources))
                          error('Usage of a predefined filter source as filter result string is not allowed.');
                      end
                      switch (filter(i).Subfilter.Type)
                          case 'feGaussianBlur'
                              printAttributeIn(fid, 'in', filter(i).Subfilter, 'Source', 'SourceGraphic', resultStrings);
                              printAttributeDouble(fid, 'stdDeviation', filter(i).Subfilter, 'Deviation');
                              fprintf(fid,' />\n');
                          case 'feImage'
                              printAttributeString(fid, 'xlink:href', filter(i).Subfilter, 'File');
                              printAttributeString(fid, 'preserveAspectRatio', filter(i).Subfilter, 'AspectRatio', 'xMidYMid meet');
                              fprintf(fid,' />\n');
                          case 'feComposite'
                              printAttributeIn(fid, 'in', filter(i).Subfilter, 'Source1', 'SourceGraphic', resultStrings);
                              printAttributeIn(fid, 'in2', filter(i).Subfilter, 'Source2', 'SourceGraphic', resultStrings);
                              printAttributeList(fid, 'operator', filter(i).Subfilter, 'Operator', {'over','in','out','atop','xor','arithmetic'}, 'over'); % 'over' | 'in' | 'out' | 'atop' | 'xor' | 'arithmetic'
                              if isfield(filter(i).Subfilter, 'Operator') && strcmp(filter(i).Subfilter.Operator, 'arithmetic')
                                  printAttributeArray(fid, {'k1','k2','k3','k4'}, filter(i).Subfilter, 'k');
                              end
                              fprintf(fid,' />\n');
                          case 'feSpecularLighting'
                              printAttributeDouble(fid, 'specularConstant', filter(i).Subfilter, 'SpecularConstant');
                              printAttributeDouble(fid, 'specularExponent', filter(i).Subfilter, 'SpecularExponent');
                              printAttributeDouble(fid, 'surfaceScale', filter(i).Subfilter, 'SurfaceScale');
                              fprintf(fid,' style = "lighting-color:white"');
                              printAttributeIn(fid, 'in', filter(i).Subfilter, 'Source', 'SourceGraphic', resultStrings);
                              fprintf(fid,' >\n');
                              if isfield(filter(i).Subfilter, 'LightType')
                                  fprintf(fid,' <%s', filter(i).Subfilter.LightType);
                                  switch filter(i).Subfilter.LightType
                                      case 'feDistantLight'
                                          printAttributeDouble(fid, 'azimuth', filter(i).Subfilter, 'Azimuth');
                                          printAttributeDouble(fid, 'elevation', filter(i).Subfilter, 'Elevation');
                                      case 'fePointLight'
                                          printAttributeArray(fid, {'x','y','z'}, filter(i).Subfilter, 'Position');
                                      case 'feSpotLight'
                                          printAttributeArray(fid, {'x','y','z'}, filter(i).Subfilter, 'Position');
                                          printAttributeArray(fid, {'pointsAtX','pointsAtY','pointsAtZ'}, filter(i).Subfilter, 'PositionsAt');
                                          printAttributeDouble(fid, 'specularExponent', filter(i).Subfilter, 'LightSpecularExponent');
                                          printAttributeDouble(fid, 'limitingConeAngle', filter(i).Subfilter, 'LimitingConeAngle');
                                      otherwise, error(['Unknown light type ''' filter(i).Subfilter.LightType ''.']);
                                  end
                                  fprintf(fid,' />\n');
                              else
                                  error('Missing field ''LightType''.');
                              end
                              fprintf(fid,'</%s>\n',filter(i).Subfilter.Type);
                          case 'feOffset'
                              printAttributeIn(fid, 'in', filter(i).Subfilter, 'Source', 'SourceGraphic', resultStrings);
                              printAttributeArray(fid, {'dx','dy'}, filter(i).Subfilter, 'Offset');
                              fprintf(fid,' />\n');
                          case 'feBlend'
                              printAttributeIn(fid, 'in', filter(i).Subfilter, 'Source1', 'SourceGraphic', resultStrings);
                              printAttributeIn(fid, 'in2', filter(i).Subfilter, 'Source2', 'SourceGraphic', resultStrings);
                              printAttributeList(fid, 'mode', filter(i).Subfilter, 'Mode', {'normal','multiply','screen','darken','lighten'}, 'normal');   % 'normal' | 'multiply' | 'screen' | 'darken' | 'lighten'
                              fprintf(fid,' />\n');
                          case 'feTurbulence'
                              printAttributeDouble(fid, 'baseFrequency', filter(i).Subfilter, 'BaseFrequency');
                              printAttributeDouble(fid, 'numOctaves', filter(i).Subfilter, 'NumOctaves');
                              printAttributeDouble(fid, 'seed', filter(i).Subfilter, 'Seed');
                              printAttributeList(fid, 'stitchTiles', filter(i).Subfilter, 'StitchTiles', {'stitch','noStitch'}, 'noStitch');   % stitch | noStitch
                              printAttributeList(fid, 'type', filter(i).Subfilter, 'TurbulenceType', {'fractalNoise','turbulence'}, 'turbulence');   % 'fractalNoise' | 'turbulence'
                              fprintf(fid,' />\n');
                          case 'feColorMatrix'
                              printAttributeIn(fid, 'in', filter(i).Subfilter, 'Source', 'SourceGraphic', resultStrings);
                              printAttributeList(fid, 'type', filter(i).Subfilter, 'ColorType', {'matrix','saturate','hueRotate','luminanceToAlpha'});   % 'matrix' | 'saturate' | 'hueRotate' | 'luminanceToAlpha'
                              if isfield(filter(i).Subfilter, 'ColorType') && strcmp(filter(i).Subfilter.ColorType, 'matrix')
                                  if isfield(filter(i).Subfilter, 'Matrix') && (lenght(filter(i).Subfilter.Matrix) == 20)
                                      fprintf(fid,' values = "');
                                      fprintf(fid,' %0.3f', filter(i).Subfilter.Matrix);
                                      fprintf(fid,'"');
                                  else
                                      error('Field ''Matrix'' is missing or not a 5x4 matrix.');
                                  end
                              end
                              if isfield(filter(i).Subfilter, 'ColorType') && (strcmp(filter(i).Subfilter.ColorType, 'saturate') || strcmp(filter(i).Subfilter.ColorType, 'hueRotate'))
                                  printAttributeDouble(fid, 'values', filter(i).Subfilter, 'Matrix');
                              end
                              fprintf(fid,' />\n');
                          case 'feFlood'
                              printAttributeColor(fid, 'flood-color', filter(i).Subfilter, 'Color');
                              printAttributeDouble(fid, 'flood-opacity', filter(i).Subfilter, 'Opacity');
                              fprintf(fid,' />\n');
                          case 'feDisplacementMap'
                              printAttributeIn(fid, 'in', filter(i).Subfilter, 'Source1', 'SourceGraphic', resultStrings);
                              printAttributeIn(fid, 'in2', filter(i).Subfilter, 'Source2', 'SourceGraphic', resultStrings);
                              printAttributeDouble(fid, 'scale', filter(i).Subfilter, 'Scale');
                              printAttributeList(fid, 'xChannelSelector', filter(i).Subfilter, 'xChannel', {'R','G','B','A'}, 'A');    % 'R' | 'G' | 'B' | 'A'
                              printAttributeList(fid, 'yChannelSelector', filter(i).Subfilter, 'yChannel', {'R','G','B','A'}, 'A');    % 'R' | 'G' | 'B' | 'A'
                              fprintf(fid,' />\n');
                          case 'feMerge'
                              printAttributeIn(fid, 'in', filter(i).Subfilter, 'Source1', 'SourceGraphic', resultStrings);
                              printAttributeIn(fid, 'in2', filter(i).Subfilter, 'Source2', 'SourceGraphic', resultStrings);
                              printAttributeList(fid, 'mode', filter(i).Subfilter, 'Mode', {'normal','multiply','screen','darken','lighten'}); % 'normal' | 'multiply' | 'screen' | 'darken' | 'lighten'
                              fprintf(fid,' />\n');
                          case 'feMorphology'
                              printAttributeIn(fid, 'in', filter(i).Subfilter, 'Source', 'SourceGraphic', resultStrings);
                              printAttributeList(fid, 'operator', filter(i).Subfilter, 'Operator', {'erode','dilate'}); % 'erode' | 'dilate'
                              printAttributeDouble(fid, 'radius', filter(i).Subfilter, 'Radius');
                              fprintf(fid,' />\n');
                          case 'feTile'
                              printAttributeIn(fid, 'in', filter(i).Subfilter, 'Source', 'SourceGraphic', resultStrings);
                              fprintf(fid,' />\n');
                          case 'feDiffuseLighting'
                              printAttributeIn(fid, 'in', filter(i).Subfilter, 'Source', 'SourceGraphic', resultStrings);
                              fprintf(fid,' style = "lighting-color:white"');
                              printAttributeDouble(fid, 'surfaceScale', filter(i).Subfilter, 'SurfaceScale');
                              printAttributeDouble(fid, 'diffuseConstant', filter(i).Subfilter, 'DiffuseConstant');
                              printAttributeDouble(fid, 'kernelUnitLength', filter(i).Subfilter, 'KernelUnitLength');
                              fprintf(fid,' />\n');

                              %                     case 'feConvolveMatrix'
                              %
                              %                         printAttributeDouble(fid, 'order', filter(i).Subfilter);
                              % kernelMatrix
                              %                         printAttributeDouble(fid, 'divisor', filter(i).Subfilter, 1.0);
                              %                         printAttributeDouble(fid, 'bias', filter(i).Subfilter, 0);
                              % targetX
                              % targetY
                              %                         printAttributeString(fid, 'edgeMode', filter(i).Subfilter, 'EdgeMode', 'duplicate');
                              %                         fprintf(fid,' kernelUnitLength = "1 1"');
                              %                         printAttributeString(fid, 'preserveAlpha', filter(i).Subfilter, 'PreserveAlpha', 'true');
                              %                         fprintf(fid,' />\n');
                          case {'feComponentTransfer','feConvolveMatrix'}
                              error('Filter not yet implemented.');
                          otherwise
                              error(['Unknown filter ''' filter(i).Subfilter.Type '''.']);
                      end
                  catch ME
                      errStr = ME.identifier;
                      if isempty(errStr)
                          errStr = ME.message;
                      end
                      error([errStr ' Error is caused by filter type ''' filter(i).Subfilter.Type '''.']);
                  end
              end
              fprintf(fid,'  </filter>\n');
              fprintf(fid,'</defs>\n');
          end
      end
  end
end

function printAttributeArray(fid, names, svgstruct, svgfield)
  if isfield(svgstruct, svgfield)
      if isnumeric(svgstruct.(svgfield))
          if length(svgstruct.(svgfield)) ~= length(names)
              error(['Length mismatch for field ''' svgfield '''.'])
          end
          for i = 1:length(names)
              fprintf(fid,' %s = "%0.3f"', names{i}, svgstruct.(svgfield)(i));
          end
      else
          error(['Field ''' svgfield ''' must be numeric.']);
      end
  else
      if nargin < 5
          error(['Missing field ''' svgfield '''.'])
      else
          for i = 1:length(names)
              fprintf(fid,' %s = "%0.3f"', names{i}, default(i));
          end
      end
  end
end

function printAttributeDouble(fid, name, svgstruct, svgfield, default)
  if isfield(svgstruct, svgfield)
      if isnumeric(svgstruct.(svgfield))
          fprintf(fid,' %s = "%0.3f"', name, svgstruct.(svgfield));
      else
          error(['Field ''' svgfield ''' must be numeric.']);
      end
  else
      if nargin < 5
          error(['Missing field ''' svgfield '''.'])
      else
          fprintf(fid,' %s = "%0.3f"', name, default);
      end
  end
end

function printAttributeIn(fid, name, svgstruct, svgfield, default, resultStrings)
  if isfield(svgstruct, svgfield)
      if ischar(svgstruct.(svgfield))
          % The strmatch below is a very inefficient search (Matlab limitation)
          if isempty(strmatch(svgstruct.(svgfield), resultStrings))
              error(['The source string ''' svgstruct.(svgfield) ''' was never a result string of a previous filter. Check for correct spelling.']);
          else
              fprintf(fid,' %s = "%s"', name, svgstruct.(svgfield));
          end
      else
          error(['Field ''' svgfield ''' must be a string.']);
      end
  else
      if nargin < 5
          error(['Missing field ''' svgfield '''.'])
      else
          fprintf(fid,' %s = "%s"', name, default);
      end
  end
end

function printAttributeString(fid, name, svgstruct, svgfield, default)
  if isfield(svgstruct, svgfield)
      if ischar(svgstruct.(svgfield))
          fprintf(fid,' %s = "%s"', name, svgstruct.(svgfield));
      else
          error(['Field ''' svgfield ''' must be a string.']);
      end
  else
      if nargin < 5
          error(['Missing field ''' svgfield '''.'])
      else
          fprintf(fid,' %s = "%s"', name, default);
      end
  end
end

function printAttributeList(fid, name, svgstruct, svgfield, list, default)
  if isfield(svgstruct, svgfield)
      if ischar(svgstruct.(svgfield))
          if isempty(strmatch(svgstruct.(svgfield), list))
              listString = strcat(list, ''' | ''');
              listString = [listString{:}];
              error(['Illegal string identifier ''' svgstruct.(svgfield) '''. Must be one out of the list: ''' listString(1:end-4) '.']);
          else
              fprintf(fid,' %s = "%s"', name, svgstruct.(svgfield));
          end
      else
          error(['Field ''' svgfield ''' must be a string.']);
      end
  else
      if nargin < 6
          error(['Missing field ''' svgfield '''.'])
      else
          fprintf(fid,' %s = "%s"', name, default);
      end
  end
end

function printAttributeColor(fid, name, svgstruct, svgfield, default)
  if isfield(svgstruct, svgfield)
      if isnumeric(svgstruct.(svgfield))
          if length(svgstruct.(svgfield)) ~= 3
              error(['Color must be a 1x3 vector for field ''' svgfield '''.'])
          else
              fprintf(fid,' %s = "%s"', name, searchcolor(gca, svgstruct.(svgfield)));
          end
      else
          error(['Field ''' svgfield ''' must be a 1x3 vector.']);
      end
  else
      if nargin < 5
          error(['Missing field ''' svgfield '''.'])
      else
          fprintf(fid,' %s = "%s"', name, default);
      end
  end
end

function frontTicks(fid, x, y, scolorname, linewidth, tick, index, edge_neighbours, c, valid_ticks, ticklength, tick_ratio, lim, drawBorder, oneSide)
  global FIG2SVG_globals
  if ~exist('oneSide','var')
    oneSide = 0;
  end
  for k = 1:length(index)
      x_tick_end1 = interp1([0 1],[x(index(k)) x(edge_neighbours(index(k),c(1)))],ticklength*tick_ratio(c(3)),'linear','extrap');
      y_tick_end1 = interp1([0 1],[y(index(k)) y(edge_neighbours(index(k),c(1)))],ticklength*tick_ratio(c(3)),'linear','extrap');
      x_tick_end2 = interp1([0 1],[x(edge_neighbours(index(k),c(2))) x(edge_neighbours(edge_neighbours(index(k),c(2)),c(1)))],ticklength*tick_ratio(c(3)),'linear','extrap');
      y_tick_end2 = interp1([0 1],[y(edge_neighbours(index(k),c(2))) y(edge_neighbours(edge_neighbours(index(k),c(2)),c(1)))],ticklength*tick_ratio(c(3)),'linear','extrap');
      xg_line_start = interp1(lim,[x(index(k)) x(edge_neighbours(index(k),c(2)))],tick);
      yg_line_start = interp1(lim,[y(index(k)) y(edge_neighbours(index(k),c(2)))],tick);
      xg_line_end = interp1(lim,[x_tick_end1 x_tick_end2],tick);
      yg_line_end = interp1(lim,[y_tick_end1 y_tick_end2],tick);
      if k == 1 || ~oneSide
        for i = valid_ticks
          line2svg(fid,[xg_line_start(i) xg_line_end(i)],[yg_line_start(i) yg_line_end(i)],scolorname,'-',linewidth)
        end
      end
      if drawBorder
          line2svg(fid,[x(index(k)) x(edge_neighbours(index(k),c(2)))],[y(index(k)) y(edge_neighbours(index(k),c(2)))],scolorname,'-',linewidth)
      end
  end
end

function gridLines(fid, x, y, scolorname, gridlinestyle, linewidth, axlim, axtick, axindex_inner, corners, c, gridAlpha)
  xg_line_start = interp1([axlim(1) axlim(2)],[x(corners(c(1))) x(corners(c(2)))], axtick);
  yg_line_start = interp1([axlim(1) axlim(2)],[y(corners(c(1))) y(corners(c(2)))], axtick);
  xg_line_end = interp1([axlim(1) axlim(2)],[x(corners(c(3))) x(corners(c(4)))], axtick);
  yg_line_end = interp1([axlim(1) axlim(2)],[y(corners(c(3))) y(corners(c(4)))], axtick);
  if nargin < 12 || isempty(gridAlpha)
      gridAlpha = 1;
  end
  for i = axindex_inner
      line2svg(fid, [xg_line_start(i) xg_line_end(i)],[yg_line_start(i) yg_line_end(i)], scolorname, gridlinestyle, linewidth, '', gridAlpha)
  end
end

function minorGridLines(fid, x, y, scolorname, minor_gridlinestyle, linewidth, axlim, minor_axtick, corners, c, gridAlpha)
  xg_line_start = interp1([axlim(1) axlim(2)],[x(corners(c(1))) x(corners(c(2)))], minor_axtick);
  yg_line_start = interp1([axlim(1) axlim(2)],[y(corners(c(1))) y(corners(c(2)))], minor_axtick);
  xg_line_end = interp1([axlim(1) axlim(2)],[x(corners(c(3))) x(corners(c(4)))], minor_axtick);
  yg_line_end = interp1([axlim(1) axlim(2)],[y(corners(c(3))) y(corners(c(4)))], minor_axtick);
  if nargin < 12 || isempty(gridAlpha)
      gridAlpha = 1;
  end
  for i = 1:length(xg_line_start)
      line2svg(fid, [xg_line_start(i) xg_line_end(i)],[yg_line_start(i) yg_line_end(i)], scolorname, minor_gridlinestyle, linewidth, '', gridAlpha)
  end
end

function group = colorbar_axes2svg(fid,id,ax,group,paperpos)
% global colorname
  global FIG2SVG_globals
  originalAxesUnits = get(ax,'Units');
  set(ax,'Units','normalized');
  axpos = get(ax,'Position');
  faces =   [1 2 4 3; 2 4 8 6; 3 4 8 7; 1 2 6 5; 1 5 7 3; 5 6 8 7];
  %           x-y    ; y-z    ; x-z    ; y-z    ; x-z    ; x-y
  corners(:,:,1) = [1 1 2 3 4; 2 1 3 2 4];
  corners(:,:,2) = [2 2 4 6 8; 3 2 6 4 8];
  corners(:,:,3) = [1 3 4 7 8; 3 3 7 4 8];
  corners(:,:,4) = [1 1 2 5 6; 3 1 5 2 6];
  corners(:,:,5) = [2 1 3 5 7; 3 1 5 3 7];
  corners(:,:,6) = [1 5 6 7 8; 2 5 7 6 8];
  edge_neighbours = [2 3 5; 1 4 6; 4 1 7; 3 2 8; 6 7 1; 5 8 2; 8 5 3; 7 6 4];
  edge_opposite = [8 7 6 5 4 3 2 1];
  nomx = [0 1 0 1 0 1 0 1];
  nomy = [0 0 1 1 0 0 1 1];
  nomz = [0 0 0 0 1 1 1 1];
  [projection,edges] = get_projection(ax,id);
  x = (edges(1,:)*axpos(3)+axpos(1))*paperpos(3);
  y = (1-(edges(2,:)*axpos(4)+axpos(2)))*paperpos(4);
  % Depth Sort of view box edges
  if size(edges,1) == 2
    edges = [edges; ones(1,size(edges,2))];
  end
  [~,edge_index] = sort(edges(3,:));
  most_back_edge_index = edge_index(1);
  % Back faces are plot box faces that are behind the plot (as seen by the
  % view point)
  back_faces = find(any(faces == most_back_edge_index,2));
  front_faces = find(all(faces ~= most_back_edge_index,2));
  groupax = group;
  axlimx = get(ax,'XLim');
  axlimy = get(ax,'YLim');
  axlimz = [0 0];
  [axinflimx, axinflimy, axinflimz] = AxesChildBounds(ax);
  axlimx(isinf(axlimx)) = axinflimx(isinf(axlimx));
  axlimy(isinf(axlimy)) = axinflimy(isinf(axlimy));
  axlimz(isinf(axlimz)) = axinflimz(isinf(axlimz));
  axlimxori = axlimx;
  axlimyori = axlimy;
  axlimzori = axlimz;
  if strcmp(get(ax,'Direction'),'reverse')
      axlimx = fliplr(axlimx);
  end
  if strcmp(get(ax,'Direction'),'reverse')
      axlimy = fliplr(axlimy);
  end
  axlimori = [axlimxori(1) axlimyori(1) axlimzori(1) axlimxori(2)-axlimxori(1) axlimyori(2)-axlimyori(1) axlimzori(2)-axlimzori(1)];
  fprintf(fid,'  <g id  = "%s">\n', createId);
  axIdString = createId;
  boundingBoxAxes = [min(x) min(y) max(x)-min(x) max(y)-min(y)];
  if strcmp(get(ax,'Visible'),'on')
      axxtick = get(ax,'XTick');
      axytick = get(ax,'YTick');
      axlabelx = get(ax,'XTickLabel');
      axlabely = get(ax,'YTickLabel');
      if ~strcmp(get(ax,'Type'),'colorbar')
        axztick = get(ax,'ZTick');
        axlabelz = get(ax,'ZTickLabel');
      end
      both_ticklength = get(ax,'TickLength');
      ticklength = both_ticklength(1);
      xy_ratio = axpos(3)*paperpos(3)/ (axpos(4)*paperpos(4));
      if xy_ratio < 1
          % disp('2D xy_ratio < 1')
          tick_ratio = [1 1/xy_ratio 1];
      else
          % disp('2D xy_ratio >= 1')
          tick_ratio = [xy_ratio 1 1];
      end
      if strcmp(get(ax,'TickDirection'),'out')
          label_distance = -60*ticklength;
      else
          label_distance = -30*ticklength;
      end
      xlabel_distance = label_distance;
      ylabel_distance = label_distance;
      % linewidth = get(ax,'LineWidth'); % linewidth = FIG2SVG_globals.resolutionScaling*get(ax,'LineWidth');
      axxindex = find((axxtick >= axlimori(1)) & (axxtick <= (axlimori(1)+axlimori(4))));
      axyindex = find((axytick >= axlimori(2)) & (axytick <= (axlimori(2)+axlimori(5))));
      % remove sticks outside of the axes (-1 of legends)
      axxtick = axxtick(axxindex);
      axytick = axytick(axyindex);
      if length(axxtick) > 1
          minor_lin_sticks = (0.2:0.2:0.8)*(axxtick(2)-axxtick(1));
          minor_axxtick = [];
          for stick = [2*axxtick(1)-axxtick(2) axxtick]
              minor_axxtick = [minor_axxtick minor_lin_sticks + stick];
          end
          minor_axxtick = minor_axxtick(minor_axxtick > min(axlimx) & minor_axxtick < max(axlimx));
      else
          minor_axxtick = [];
      end
      if length(axytick) > 1
          minor_lin_sticks = (0.2:0.2:0.8)*(axytick(2)-axytick(1));
          minor_axytick = [];
          for stick = [2*axytick(1)-axytick(2) axytick]
              minor_axytick = [minor_axytick minor_lin_sticks + stick];
          end
          minor_axytick = minor_axytick(minor_axytick > min(axlimy) & minor_axytick < max(axlimy));
      else
          minor_axytick = [];
      end
      FIG2SVG_globals.BoxOn = strcmp(get(ax,'Box'),'on');
      if strcmp(get(ax,'Box'),'on')
          axxindex_inner = find((axxtick > axlimori(1)) & (axxtick < (axlimori(1)+axlimori(4))));
          axyindex_inner = find((axytick > axlimori(2)) & (axytick < (axlimori(2)+axlimori(5))));
      else
          axxindex_inner = find((axxtick >= axlimori(1)) & (axxtick <= (axlimori(1)+axlimori(4))));
          axyindex_inner = find((axytick >= axlimori(2)) & (axytick <= (axlimori(2)+axlimori(5))));
      end
      minor_log_sticks = log10(0.2:0.1:0.9);
      if strcmp(get(ax,'TickDir'),'out')
          ticklength = -ticklength;
          valid_xsticks = 1:length(axxindex);
          valid_ysticks = 1:length(axyindex);
      else
          valid_xsticks = axxindex_inner;
          valid_ysticks = axyindex_inner;
          if ~strcmp(get(ax,'Type'),'colorbar')
            valid_zsticks = axzindex_inner;
          end
      end
      linewidth = get(ax,'LineWidth');
  end

  fprintf(fid,'    <g>\n');
  boundingBoxAxes = colorbar2svg(fid,id,axIdString,ax,paperpos,axpos,groupax,projection,boundingBoxAxes);
  fprintf(fid,'  <clipPath id = "%s">\n',axIdString);
  fprintf(fid,'    <rect x = "%0.3f" y = "%0.3f" width = "%0.3f" height = "%0.3f"/>\n',...
     boundingBoxAxes(1), boundingBoxAxes(2), boundingBoxAxes(3), boundingBoxAxes(4));
  fprintf(fid,'  </clipPath>\n');
  fprintf(fid,'    </g>\n');

  if strcmp(get(ax,'Visible'),'on')
      fprintf(fid,'    <g>\n');
      % Search axis for labeling
      if projection.xyplane
          [~, x_axis_point_index_top] = min(y);
          [~, x_axis_point_index_bottom] = max(y);
          if strcmp(get(ax,'Box'),'on')
              if strcmp(get(ax,'XAxisLocation'),'top')
                  x_axis_point_index = [x_axis_point_index_top x_axis_point_index_bottom];
              else
                  x_axis_point_index = [x_axis_point_index_bottom x_axis_point_index_top];
              end
          else
              if strcmp(get(ax,'XAxisLocation'),'top')
                  x_axis_point_index = x_axis_point_index_top;
              else
                  x_axis_point_index = x_axis_point_index_bottom;
              end
          end
          [~, y_axis_point_index_left] = min(x);
          [~, y_axis_point_index_right] = max(x);
          if strcmp(get(ax,'Box'),'on')
              if strcmp(get(ax,'YAxisLocation'),'right')
                  y_axis_point_index = [y_axis_point_index_right y_axis_point_index_left];
              else
                  y_axis_point_index = [y_axis_point_index_left y_axis_point_index_right];
              end
          else
              if strcmp(get(ax,'YAxisLocation'),'right')
                  y_axis_point_index = y_axis_point_index_right;
              else
                  y_axis_point_index = y_axis_point_index_left;
              end
          end
          [~, z_axis_point_index] = min(x);
      else
          [~, x_axis_point_index] = max(y);
          [~, y_axis_point_index] = max(y);
          [~, z_axis_point_index] = min(x);
      end
      scolorname = searchcolor(id,get(ax,'XColor'));
      % Draw 'box' of x-axis
      if projection.xyplane == false
          if strcmp(get(ax,'Box'),'on')
              edge_line_index = [edge_opposite(most_back_edge_index) edge_neighbours(edge_opposite(most_back_edge_index),1)];
              line2svg(fid,x(edge_line_index),y(edge_line_index),scolorname,'-',linewidth)
          end
      end

      % Draw x-tick marks
      if (ticklength(1) ~= 0)
          if axlimx(1) ~= axlimx(2)
              if (nomx(x_axis_point_index(1)))
                  lim = [axlimx(2) axlimx(1)];
              else
                  lim = [axlimx(1) axlimx(2)];
              end
              x_label_end1 = interp1([0 1],[x(x_axis_point_index(1)) x(edge_neighbours(x_axis_point_index(1),2))],xlabel_distance,'linear','extrap');
              y_label_end1 = interp1([0 1],[y(x_axis_point_index(1)) y(edge_neighbours(x_axis_point_index(1),2))],xlabel_distance,'linear','extrap');
              x_label_end2 = interp1([0 1],[x(edge_neighbours(x_axis_point_index(1),1)) x(edge_neighbours(edge_neighbours(x_axis_point_index(1),1),2))],xlabel_distance,'linear','extrap');
              y_label_end2 = interp1([0 1],[y(edge_neighbours(x_axis_point_index(1),1)) y(edge_neighbours(edge_neighbours(x_axis_point_index(1),1),2))],xlabel_distance,'linear','extrap');
              xg_label_end = interp1(lim,[x_label_end1 x_label_end2],axxtick);
              yg_label_end = interp1(lim,[y_label_end1 y_label_end2],axxtick);
              if axpos(3) > axpos(4)
                frontTicks(fid, x, y, scolorname, linewidth, ...
                  axxtick, x_axis_point_index, edge_neighbours, [2 1 1], ...
                  valid_xsticks,  2*ticklength, tick_ratio, lim, true, true);
              else
                frontTicks(fid, x, y, scolorname, linewidth, ...
                  axxtick, x_axis_point_index, edge_neighbours, [2 1 1], ...
                  [],  ticklength, tick_ratio, lim, true);
              end
              if ~isempty(axlabelx) && ~(iscell(axlabelx) && all(cellfun(@isempty,axlabelx)))
                  if ischar(axlabelx) && size(axlabelx, 1) == 1
                      % Special handling of 1xn char arrays -> duplicate data
                      % for all ticks. Strange behavior but follows the
                      % behavior of Matlab
                      axlabelx = repmat(axlabelx, length(axxindex), 1);
                  end
                  if (strcmp(get(ax,'XTickLabelMode'),'manual'))
                      axlabelx = axlabelx(axxindex,:);
                  end
                  if FIG2SVG_globals.octave
                    axlabelx = axlabelx(:); % SA: Octave compatibility
                  end
                  if axpos(3) > axpos(4)
                    if strcmp(get(ax,'XAxisLocation'),'top')
                      [angle, align] = improvedXLabel(ax, 0, 'center');
                      for i = 1:length(axxindex)
                        label2svg(fid,axpos,ax,xg_label_end(i),yg_label_end(i),convertString(axlabelx(i,:)),align,angle,'bottom',1,scolorname);
                      end
                    else
                      [angle, align] = improvedXLabel(ax, 0, 'center');
                      for i = 1:length(axxindex)
                        label2svg(fid,axpos,ax,xg_label_end(i),yg_label_end(i),convertString(axlabelx(i,:)),align,angle,'top',1,scolorname);
                      end
                    end
                  end
              end
          end
      end
      scolorname = searchcolor(id,get(ax,'YColor'));
      % Draw 'box' of y-axis
      if projection.xyplane == false
          if strcmp(get(ax,'Box'),'on')
              edge_line_index = [edge_opposite(most_back_edge_index) edge_neighbours(edge_opposite(most_back_edge_index),2)];
              line2svg(fid,x(edge_line_index),y(edge_line_index),scolorname,'-',linewidth)
          end
      end

      % Draw y-tick marks
      if (ticklength(1) ~= 0)
          if axlimy(1) ~= axlimy(2)
              if (nomy(y_axis_point_index(1)))
                  lim = [axlimy(2) axlimy(1)];
              else
                  lim = [axlimy(1) axlimy(2)];
              end
              x_label_end1 = interp1([0 1],[x(y_axis_point_index(1)) x(edge_neighbours(y_axis_point_index(1),1))],ylabel_distance,'linear','extrap');
              y_label_end1 = interp1([0 1],[y(y_axis_point_index(1)) y(edge_neighbours(y_axis_point_index(1),1))],ylabel_distance,'linear','extrap');
              x_label_end2 = interp1([0 1],[x(edge_neighbours(y_axis_point_index(1),2)) x(edge_neighbours(edge_neighbours(y_axis_point_index(1),2),1))],ylabel_distance,'linear','extrap');
              y_label_end2 = interp1([0 1],[y(edge_neighbours(y_axis_point_index(1),2)) y(edge_neighbours(edge_neighbours(y_axis_point_index(1),2),1))],ylabel_distance,'linear','extrap');
              xg_label_end = interp1(lim,[x_label_end1 x_label_end2],axytick);
              yg_label_end = interp1(lim,[y_label_end1 y_label_end2],axytick);
              if strcmp(get(ax,'Type'),'colorbar') && axpos(3) < axpos(4)
                  frontTicks(fid, x, y, scolorname, linewidth, ...
                  axytick, y_axis_point_index, edge_neighbours, [1 2 2], ...
                  valid_ysticks,  2*ticklength, tick_ratio, lim, true, true);
              else
                frontTicks(fid, x, y, scolorname, linewidth, ...
                  axytick, y_axis_point_index, edge_neighbours, [1 2 2], ...
                  [],  ticklength, tick_ratio, lim, true);
              end
              if ~isempty(axlabely) && ~(iscell(axlabely) && all(cellfun(@isempty,axlabely)))
                  if ischar(axlabely) && size(axlabely, 1) == 1
                      % Special handling of 1xn char arrays -> duplicate data
                      % for all ticks. Strange behavior but follows the
                      % behavior of Matlab
                      axlabely = repmat(axlabely, length(axyindex), 1);
                  end
                  if (strcmp(get(ax,'YTickLabelMode'),'manual'))
                      axlabely = axlabely(axyindex,:);
                  end
                  if FIG2SVG_globals.octave
                    axlabely = axlabely(:); % SA: Octave compatibility
                  end
                  if axpos(3) < axpos(4)
                    if strcmp(get(ax,'YAxisLocation'),'right')
                        [angle, align] = improvedYLabel(ax, 0, 'left');
                        for i = 1:length(axyindex)
                            label2svg(fid,axpos,ax,xg_label_end(i),yg_label_end(i),convertString(axlabely(i,:)),align,angle,'middle',1,scolorname);
                        end
                   else
                        [angle, align] = improvedYLabel(ax, 0, 'right');
                        for i = 1:length(axyindex)
                            label2svg(fid,axpos,ax,xg_label_end(i),yg_label_end(i),convertString(axlabely(i,:)),align,angle,'middle',1,scolorname);
                        end
                    end
                  end
              end
          end
      end
      exponent2svg(fid,axpos,paperpos,ax,axxtick,axytick,[])
      fprintf(fid,'    </g>\n');
  end
  fprintf(fid,'  </g>\n');
  set(ax,'Units',originalAxesUnits);
end

function boundingBoxAxes = colorbar2svg(fid,id,axIdString,ax,paperpos,axpos,groupax,projection,boundingBoxAxes)
  global colorname
  global FIG2SVG_globals

  cmap = get(id,'Colormap');
  if axpos(3) > axpos(4)
    pointc = 1:size(cmap,1);
  else
    pointc = (size(cmap,1):-1:1)';
  end
  filename = [FIG2SVG_globals.basefilename sprintf('%03d',FIG2SVG_globals.figurenumber) '.' FIG2SVG_globals.pixelFileType];
  imwrite(pointc,cmap,fullfile(FIG2SVG_globals.basefilepath,filename),FIG2SVG_globals.pixelFileType);
  [filterString, boundingBox] = filter2svg(fid, ax, boundingBoxAxes, boundingBoxAxes);
  lx = axpos(3)*paperpos(3);
  ly = axpos(4)*paperpos(4);
  pointsx = axpos(1)*paperpos(3);
  pointsy = (1-(axpos(4)+axpos(2)))*paperpos(4);
  [filterString, boundingBox] = filter2svg(fid, ax, boundingBoxAxes, boundingBoxAxes);
  if FIG2SVG_globals.ClippingMode
      clippingIdString = clipping2svg(fid, ax, ax, paperpos, axpos, projection, axIdString);
      fprintf(fid,'<g id = "%s" clip-path = "url(#%s)" %s>\n', createId, clippingIdString, filterString);
      if ~isempty(filterString)
          % Workaround for Inkscape filter bug
          fprintf(fid,'<rect x = "%0.3f" y = "%0.3f" width = "%0.3f" height = "%0.3f" fill = "none" stroke = "none" />\n', boundingBox(1), boundingBox(2), boundingBox(3), boundingBox(4));
      end
      fprintf(fid,'<image x = "%0.3f" y = "%0.3f" width = "%0.3f" height = "%0.3f" image-rendering = "optimizeSpeed" preserveAspectRatio = "none" xlink:href = "%s" />\n', pointsx, pointsy, lx, ly, filename); % With image-rendering = "optimizeQuality" the image appears interpolated, which might be nice, but image/imagesc don't work that way, images appear pixelated. To workaround this, use pcolor+shading, or previously interpolate with interp2
      fprintf(fid,'</g>\n');
  else
      fprintf(fid,'<g id = "%s" %s>\n', createId, filterString);
      if ~isempty(filterString)
          % Workaround for Inkscape filter bug
          fprintf(fid,'<rect x = "%0.3f" y = "%0.3f" width = "%0.3f" height = "%0.3f" fill = "none" stroke = "none" />\n', boundingBox(1), boundingBox(2), boundingBox(3), boundingBox(4));
      end
      fprintf(fid,'<image x = "%0.3f" y = "%0.3f" width = "%0.3f" height = "%0.3f" image-rendering = "optimizeSpeed" preserveAspectRatio = "none" xlink:href = "%s" />\n', pointsx, pointsy, lx, ly, filename); % With image-rendering = "optimizeQuality" the image appears interpolated, which might be nice, but image/imagesc don't work that way, images appear pixelated. To workaround this, use pcolor+shading, or previously interpolate with interp2
      fprintf(fid,'</g>\n');
  end
end

function group = annotation2svg(fid, id, ax, group, paperpos)
  global FIG2SVG_globals
  if strcmp(get(ax,'Visible'),'on')
    axIdString = createId;
    fprintf(fid,'  <g id  = "%s">\n', axIdString);
    axchild = get(ax,'Children');
    for i = numel(axchild):-1:1
      if FIG2SVG_globals.debugModeOn
        currentType = get(axchild(i),'Type');
        disp(['    axchild(',num2str(i),') = ', currentType]);
      end
      if strcmp(get(axchild(i), 'Visible'), 'off')
        % do nothing
      elseif strcmp(get(axchild(i),'Type'),'textboxshape')
        facecolor = get(axchild(i),'BackgroundColor');
        edgecolor = get(axchild(i),'EdgeColor');
        linewidth = get(axchild(i),'LineWidth'); % linewidth = FIG2SVG_globals.resolutionScaling*get(axchild(i),'LineWidth');
        linestyle = get(axchild(i),'LineStyle');
        if ischar(facecolor)
            if ~strcmp(facecolor,'none')
                error('Illegal face color for text.');
            else
                facecolorname = 'none';
            end
        else
            facecolorname = searchcolor(id, facecolor);
        end
        face_opacity = get(axchild(i),'FaceAlpha');
        if ischar(edgecolor)
            if ~strcmp(edgecolor,'none')
                error('Illegal edge color for text.');
            else
                edgecolorname = 'none';
            end
        else
            edgecolorname = searchcolor(id, edgecolor);
        end

        projection = [];

        annotationpos = get(axchild(i), 'Position');
        annotationBox = [annotationpos(1)*paperpos(3) (1 - (annotationpos(2) + annotationpos(4)) )*paperpos(4) annotationpos(3)*paperpos(3) annotationpos(4)*paperpos(4)];

        textalign = get(axchild(i),'HorizontalAlignment');
        textvalign = get(axchild(i),'VerticalAlignment');
        margin = convertunit(get(axchild(i),'Margin'),get(axchild(i),'FontUnits'),'pixels', annotationBox(4));

        switch textalign
          case 'left', x = margin + annotationpos(1)*paperpos(3);
          case 'center', x = (annotationpos(1) + 0.5*annotationpos(3))*paperpos(3);
          case 'right', x = (annotationpos(1) + annotationpos(3))*paperpos(3) - margin;
          otherwise
        end
        switch textvalign
          case 'top', y = (1- (annotationpos(2)+annotationpos(4)))*paperpos(4);
          case 'middle', y = (1 - (annotationpos(2) + 0.5*annotationpos(4)))*paperpos(4);
          case 'bottom', y = (1 - annotationpos(2)) * paperpos(4);
          otherwise
        end
        textpos = [x y annotationBox(3) annotationBox(4)];

        [filterString, boundingBoxAxes] = filter2svg(fid, axchild(i), annotationBox, annotationBox);

        box = annotationBox;

        if FIG2SVG_globals.ClippingMode ~= 0
          fprintf(fid,'<g id = "%s" clip-path = "url(#%s)" %s>\n', createId, axIdString, filterString);
        else
          fprintf(fid,'<g id = "%s" %s>\n', createId, filterString);
        end
        if ~isempty(filterString)
            % Workaround for Inkscape filter bug
            fprintf(fid,'<rect x = "%0.3f" y = "%0.3f" width = "%0.3f" height = "%0.3f" fill = "none" stroke = "none" />\n', box(1), box(2), box(3), box(4));
        end
        if ~strcmp(edgecolorname, 'none') || ~strcmp(facecolorname, 'none')
            pattern = lineStyle2svg(linestyle, linewidth);
            fprintf(fid,'<rect x = "%0.3f" y = "%0.3f" width = "%0.3f" height = "%0.3f" fill = "%s" fill-opacity = "%0.3f" stroke-linecap = "square" stroke-linejoin = "miter" stroke = "%s" stroke-width = "%0.3fpt" %s />\n', ...
                box(1), box(2), box(3), box(4), facecolorname, face_opacity, edgecolorname, linewidth, pattern);
        end
        text2svg(fid,textpos,paperpos,axchild(i),ax,projection)
        fprintf(fid,'</g>\n');
      else
        disp(['   Warning: Unhandled child type: ' get(axchild(i),'Type')]);
      end
    end
    fprintf(fid,'</g>\n');
  end
end

%%%%%%%%%%%%%%%%%%%%%%%%%%%%%%%%%%%%%%%%% SUBFUNCTIONS %%%%%
% Create axis frame and insert all children of this axis frame
function group = axes2svg(fid,id,ax,group,paperpos)
  % global colorname
  global FIG2SVG_globals
  originalAxesUnits = get(ax,'Units');
  set(ax,'Units','normalized');
  axpos = get(ax,'Position');
  faces =   [1 2 4 3; 2 4 8 6; 3 4 8 7; 1 2 6 5; 1 5 7 3; 5 6 8 7];
  %           x-y    ; y-z    ; x-z    ; y-z    ; x-z    ; x-y
  corners(:,:,1) = [1 1 2 3 4; 2 1 3 2 4];
  corners(:,:,2) = [2 2 4 6 8; 3 2 6 4 8];
  corners(:,:,3) = [1 3 4 7 8; 3 3 7 4 8];
  corners(:,:,4) = [1 1 2 5 6; 3 1 5 2 6];
  corners(:,:,5) = [2 1 3 5 7; 3 1 5 3 7];
  corners(:,:,6) = [1 5 6 7 8; 2 5 7 6 8];
  edge_neighbours = [2 3 5; 1 4 6; 4 1 7; 3 2 8; 6 7 1; 5 8 2; 8 5 3; 7 6 4];
  edge_opposite = [8 7 6 5 4 3 2 1];
  nomx = [0 1 0 1 0 1 0 1];
  nomy = [0 0 1 1 0 0 1 1];
  nomz = [0 0 0 0 1 1 1 1];
  [projection,edges] = get_projection(ax,id);
  x = (edges(1,:)*axpos(3)+axpos(1))*paperpos(3);
  y = (1-(edges(2,:)*axpos(4)+axpos(2)))*paperpos(4);
  % Depth Sort of view box edges
  if size(edges,1) == 2
    edges = [edges; ones(1,size(edges,2))];
  end
  [~,edge_index] = sort(edges(3,:));
  most_back_edge_index = edge_index(1);
  % Back faces are plot box faces that are behind the plot (as seen by the
  % view point)
  back_faces = find(any(faces == most_back_edge_index,2));
  front_faces = find(all(faces ~= most_back_edge_index,2));
  groupax = group;
  axlimx = get(ax,'XLim');
  axlimy = get(ax,'YLim');
  axlimz = get(ax,'ZLim');
  if isfield(FIG2SVG_globals,'BoxOn') % only when axis is on
    [axinflimx, axinflimy, axinflimz] = AxesChildBounds(ax);
  else
    axinflimx = axlimx;
    axinflimy = axlimy;
    axinflimz = axlimz;
  end
  axlimx(isinf(axlimx)) = axinflimx(isinf(axlimx));
  axlimy(isinf(axlimy)) = axinflimy(isinf(axlimy));
  axlimz(isinf(axlimz)) = axinflimz(isinf(axlimz));
  axlimxori = axlimx;
  axlimyori = axlimy;
  axlimzori = axlimz;
  if strcmp(get(ax,'XScale'),'log')
      axlimx = log10(axlimx);
      axlimx(isinf(axlimx)) = 0;
  end
  if strcmp(get(ax,'YScale'),'log')
      axlimy = log10(axlimy);
      axlimy(isinf(axlimy)) = 0;
  end
  if strcmp(get(ax,'ZScale'),'log')
      axlimz = log10(axlimz);
      axlimz(isinf(axlimz)) = 0;
  end
  if strcmp(get(ax,'XDir'),'reverse')
      axlimx = fliplr(axlimx);
  end
  if strcmp(get(ax,'YDir'),'reverse')
      axlimy = fliplr(axlimy);
  end
  if strcmp(get(ax,'ZDir'),'reverse')
      axlimz = fliplr(axlimz);
  end
  axlimori = [axlimxori(1) axlimyori(1) axlimzori(1) axlimxori(2)-axlimxori(1) axlimyori(2)-axlimyori(1) axlimzori(2)-axlimzori(1)];
  fprintf(fid,'  <g id  = "%s">\n', createId);
  axIdString = createId;
  boundingBoxAxes = [min(x) min(y) max(x)-min(x) max(y)-min(y)];
  if strcmp(get(ax,'Visible'),'on')
      axxtick = get(ax,'XTick');
      axytick = get(ax,'YTick');
      axztick = get(ax,'ZTick');
      axlabelx = get(ax,'XTickLabel');
      axlabely = get(ax,'YTickLabel');
      axlabelz = get(ax,'ZTickLabel');
      % SA: Octave compatibility
      if FIG2SVG_globals.octave
        for i = length(axlabelx):-1:1
          if str2num(axlabelx{i}) < axlimx(1) || str2num(axlabelx{i}) > axlimx(2)
            axlabelx(i) = [];
          end
        end
        for i = length(axlabely):-1:1
          if str2num(axlabely{i}) < axlimy(1) || str2num(axlabely{i}) > axlimy(2)
            axlabely(i) = [];
          end
        end
        for i = length(axlabelz):-1:1
          if str2num(axlabelz{i}) < axlimz(1) || str2num(axlabelz{i}) > axlimz(2)
            axlabelz(i) = [];
          end
        end
      end
      % Workaround for Octave
      if FIG2SVG_globals.octave
          if isempty(axlabelx)
              if strcmp(get(ax,'XScale'),'log')
                  axlabelx = num2str(log10(axxtick)');
              else
                  axlabelx = num2str(axxtick');
              end
          end
          if isempty(axlabely)
              if strcmp(get(ax,'YScale'),'log')
                  axlabely = num2str(log10(axytick)');
              else
                  axlabely = num2str(axytick');
              end
          end
          if isempty(axlabelz)
              if strcmp(get(ax,'ZScale'),'log')
                  axlabelz = num2str(log10(axztick)');
              else
                  axlabelz = num2str(axztick');
              end
          end
          if projection.xyplane
              axlabelz = [];
          end
      end
      gridlinestyle = get(ax,'GridLineStyle');
      minor_gridlinestyle = get(ax,'MinorGridLineStyle');
      both_ticklength = get(ax,'TickLength');
      gridBehind = true; % Default setting
      try
          if strcmp(get(ax, 'Layer'), 'top') && projection.xyplane
              gridBehind = false;
          end
      catch
          gridBehind = true;
      end
      if projection.xyplane
          ticklength = both_ticklength(1);
          xy_ratio = axpos(3)*paperpos(3)/ (axpos(4)*paperpos(4));
          if xy_ratio < 1
              % disp('2D xy_ratio < 1')
              tick_ratio = [1 1/xy_ratio 1];
          else
              % disp('2D xy_ratio >= 1')
              tick_ratio = [xy_ratio 1 1];
          end
          if strcmp(get(ax,'TickDir'),'out')
              label_distance = -(0.02 + ticklength);
          else
              label_distance = -0.02;
          end
          xlabel_distance = label_distance;
          ylabel_distance = label_distance;
          zlabel_distance = label_distance;
      else
          % disp('3D')
          ticklength = both_ticklength(2);
          tick_ratio = 1.5*[1 1 1];
          label_distance = -2*abs(ticklength);
          xlabel_distance = tick_ratio(1)*label_distance;
          ylabel_distance = tick_ratio(2)*label_distance;
          zlabel_distance = tick_ratio(3)*label_distance;
      end
      % linewidth = get(ax,'LineWidth'); % linewidth = FIG2SVG_globals.resolutionScaling*get(ax,'LineWidth');
      axxindex = find((axxtick >= axlimori(1)) & (axxtick <= (axlimori(1)+axlimori(4))));
      axyindex = find((axytick >= axlimori(2)) & (axytick <= (axlimori(2)+axlimori(5))));
      axzindex = find((axztick >= axlimori(3)) & (axztick <= (axlimori(3)+axlimori(6))));
      % remove sticks outside of the axes (-1 of legends)
      axxtick = axxtick(axxindex);
      axytick = axytick(axyindex);
      axztick = axztick(axzindex);
      if length(axxtick) > 1
          minor_lin_sticks = (0.2:0.2:0.8)*(axxtick(2)-axxtick(1));
          minor_axxtick = [];
          for stick = [2*axxtick(1)-axxtick(2) axxtick]
              minor_axxtick = [minor_axxtick minor_lin_sticks + stick];
          end
          minor_axxtick = minor_axxtick(minor_axxtick > min(axlimx) & minor_axxtick < max(axlimx));
      else
          minor_axxtick = [];
      end
      if length(axytick) > 1
          minor_lin_sticks = (0.2:0.2:0.8)*(axytick(2)-axytick(1));
          minor_axytick = [];
          for stick = [2*axytick(1)-axytick(2) axytick]
              minor_axytick = [minor_axytick minor_lin_sticks + stick];
          end
          minor_axytick = minor_axytick(minor_axytick > min(axlimy) & minor_axytick < max(axlimy));
      else
          minor_axytick = [];
      end
      if length(axztick) > 1
          minor_lin_sticks = (0.2:0.2:0.8)*(axztick(2)-axztick(1));
          minor_axztick = [];
          for stick = [2*axztick(1)-axztick(2) axztick]
              minor_axztick = [minor_axztick minor_lin_sticks + stick];
          end
          minor_axztick = minor_axztick(minor_axztick > min(axlimz) & minor_axztick < max(axlimz));
      else
          minor_axztick = [];
      end
      FIG2SVG_globals.BoxOn = strcmp(get(ax,'Box'),'on');
      if  strcmp(get(ax,'Box'),'on')
          axxindex_inner = find((axxtick > axlimori(1)) & (axxtick < (axlimori(1)+axlimori(4))));
          axyindex_inner = find((axytick > axlimori(2)) & (axytick < (axlimori(2)+axlimori(5))));
          axzindex_inner = find((axztick > axlimori(3)) & (axztick < (axlimori(3)+axlimori(6))));
      else
          axxindex_inner = find((axxtick >= axlimori(1)) & (axxtick <= (axlimori(1)+axlimori(4))));
          axyindex_inner = find((axytick >= axlimori(2)) & (axytick <= (axlimori(2)+axlimori(5))));
          axzindex_inner = find((axztick >= axlimori(3)) & (axztick <= (axlimori(3)+axlimori(6))));
      end
      minor_log_sticks = log10(0.2:0.1:0.9);
      if strcmp(get(ax,'TickDir'),'out')
          ticklength = -ticklength;
          valid_xsticks = 1:length(axxindex);
          valid_ysticks = 1:length(axyindex);
          valid_zsticks = 1:length(axzindex);
      else
          valid_xsticks = axxindex_inner;
          valid_ysticks = axyindex_inner;
          valid_zsticks = axzindex_inner;
      end
      if strcmp(get(ax,'XScale'),'log')
          axxtick = log10(get(ax,'XTick'));
          minor_axxtick = [];
          if ~isempty(axxtick)
              all_axxtick = axxtick(1):(axxtick(end) + 1);
              for stick = all_axxtick
                  minor_axxtick = [minor_axxtick minor_log_sticks + stick];
              end
          end
          minor_axxtick = minor_axxtick(minor_axxtick > min(axlimx) & minor_axxtick < max(axlimx));
      end
      if strcmp(get(ax,'YScale'),'log')
          axytick = log10(get(ax,'YTick'));
          minor_axytick = [];
          if ~isempty(axytick)
              all_axytick = axytick(1):(axytick(end) + 1);
              for stick = all_axytick
                  minor_axytick = [minor_axytick minor_log_sticks + stick];
              end
          end
          minor_axytick = minor_axytick(minor_axytick > min(axlimy) & minor_axytick < max(axlimy));
      end
      if strcmp(get(ax,'ZScale'),'log')
          axztick = log10(get(ax,'ZTick'));
          minor_axztick = [];
          if ~isempty(axztick)
              all_axztick = axztick(1):1:(axztick(end) + 1);
              for stick = all_axztick
                  minor_axztick = [minor_axztick minor_log_sticks + stick];
              end
          end
          minor_axztick = minor_axztick(minor_axztick > min(axlimz) & minor_axztick < max(axlimz));
      end
      linewidth = get(ax,'LineWidth'); % linewidth = FIG2SVG_globals.resolutionScaling*get(ax,'LineWidth');
      % Draw back faces
      if ~strcmp(get(ax,'Color'),'none')
          background_color = searchcolor(id,get(ax,'Color'));
          background_opacity = 1;
      else
          background_color = '#000000';
          background_opacity = 0;
      end
      for p = 1:size(back_faces)
          patch2svg(fid, x(faces(back_faces(p),:)), y(faces(back_faces(p),:)), background_color, '-', linewidth, 'none', background_opacity, 1.0, true)
      end
      for pindex = 1:size(back_faces)
          p = back_faces(pindex);
          for k = 1:size(corners,1)
              selectedCorners = squeeze(corners(k,:,p));
              if FIG2SVG_globals.octave || UIverlessthan('8.4.0')
                  gridAlpha = 1;
                  minorGridAlpha = 1;
              else
                  gridAlpha = get(ax, 'GridAlpha');
                  minorGridAlpha = get(ax, 'MinorGridAlpha');
              end
              switch corners(k,1,p)
                  case 1 % x
                      % Draw x-grid
                      if FIG2SVG_globals.octave || UIverlessthan('8.4.0')
                          scolorname = get(ax, 'XColor');
                      else
                          if strcmp(get(ax, 'GridColorMode'), 'auto')
                              scolorname = get(ax, 'XColor');
                          else
                              scolorname = get(ax, 'GridColor');
                          end
                      end
                      scolorname = searchcolor(id,scolorname);
                      if strcmp(get(ax,'XGrid'),'on') && gridBehind
                          if axlimx(1) ~= axlimx(2)
                              gridLines(fid, x, y, scolorname, gridlinestyle, linewidth, axlimx, axxtick, axxindex_inner, selectedCorners, [2 3 4 5], gridAlpha)
                              if strcmp(get(ax,'XTickMode'),'auto') && strcmp(get(ax,'XMinorGrid'),'on') && ~isempty(minor_axxtick)
                                  minorGridLines(fid, x, y, scolorname, minor_gridlinestyle, linewidth, axlimx, minor_axxtick, selectedCorners, [2 3 4 5], minorGridAlpha)
                              end
                          end
                      end
                      if projection.xyplane == false
                          if strcmp(get(ax,'Box'),'on')
                              line2svg(fid,[x(corners(k,2,p)) x(corners(k,3,p))],[y(corners(k,2,p)) y(corners(k,3,p))],scolorname,'-',linewidth);
                              line2svg(fid,[x(corners(k,4,p)) x(corners(k,5,p))],[y(corners(k,4,p)) y(corners(k,5,p))],scolorname,'-',linewidth);
                          else
                              if strcmp(get(ax,'XGrid'),'on')
                                  line2svg(fid,[x(corners(k,2,p)) x(corners(k,3,p))],[y(corners(k,2,p)) y(corners(k,3,p))],scolorname,gridlinestyle,linewidth);
                                  line2svg(fid,[x(corners(k,4,p)) x(corners(k,5,p))],[y(corners(k,4,p)) y(corners(k,5,p))],scolorname,gridlinestyle,linewidth);
                              end
                          end
                      end
                  case 2 % y
                      % Draw y-grid
                      if FIG2SVG_globals.octave || UIverlessthan('8.4.0')
                          scolorname = get(ax, 'YColor');
                      else
                          if strcmp(get(ax, 'GridColorMode'), 'auto')
                              scolorname = get(ax, 'YColor');
                          else
                              scolorname = get(ax, 'GridColor');
                          end
                      end
                      scolorname = searchcolor(id,scolorname);
                      if strcmp(get(ax,'YGrid'),'on') && gridBehind
                          if axlimy(1) ~= axlimy(2)
                              gridLines(fid, x, y, scolorname, gridlinestyle, linewidth, axlimy, axytick, axyindex_inner, selectedCorners, [2 3 4 5], gridAlpha)
                              if strcmp(get(ax,'YTickMode'),'auto') && strcmp(get(ax,'YMinorGrid'),'on') && ~isempty(minor_axytick)
                                  minorGridLines(fid, x, y, scolorname, minor_gridlinestyle, linewidth, axlimy, minor_axytick, selectedCorners, [2 3 4 5], minorGridAlpha)
                              end
                          end
                      end
                      if projection.xyplane == false
                          if strcmp(get(ax,'Box'),'on')
                              line2svg(fid,[x(corners(k,2,p)) x(corners(k,3,p))],[y(corners(k,2,p)) y(corners(k,3,p))],scolorname,'-',linewidth);
                              line2svg(fid,[x(corners(k,4,p)) x(corners(k,5,p))],[y(corners(k,4,p)) y(corners(k,5,p))],scolorname,'-',linewidth);
                          else
                              if strcmp(get(ax,'YGrid'),'on')
                                  line2svg(fid,[x(corners(k,2,p)) x(corners(k,3,p))],[y(corners(k,2,p)) y(corners(k,3,p))],scolorname,gridlinestyle,linewidth);
                                  line2svg(fid,[x(corners(k,4,p)) x(corners(k,5,p))],[y(corners(k,4,p)) y(corners(k,5,p))],scolorname,gridlinestyle,linewidth);
                              end
                          end
                      end
                  case 3 % z
                      % Draw z-grid
                      if FIG2SVG_globals.octave || UIverlessthan('8.4.0')
                          scolorname = get(ax, 'ZColor');
                      else
                          if strcmp(get(ax, 'GridColorMode'), 'auto')
                              scolorname = get(ax, 'ZColor');
                          else
                              scolorname = get(ax, 'GridColor');
                          end
                      end
                      scolorname = searchcolor(id,scolorname);
                      if strcmp(get(ax,'ZGrid'),'on') && gridBehind
                          if axlimz(1) ~= axlimz(2)
                              gridLines(fid, x, y, scolorname, gridlinestyle, linewidth, axlimz, axztick, axzindex_inner, selectedCorners, [2 3 4 5], gridAlpha)
                              if strcmp(get(ax,'ZTickMode'),'auto') && strcmp(get(ax,'ZMinorGrid'),'on') && ~isempty(minor_axztick)
                                  minorGridLines(fid, x, y, scolorname, minor_gridlinestyle, linewidth, axlimz, minor_axztick, selectedCorners, [2 3 4 5], minorGridAlpha)
                              end
                          end
                      end
                      if projection.xyplane == false
                          if strcmp(get(ax,'Box'),'on')
                              line2svg(fid,[x(corners(k,2,p)) x(corners(k,3,p))],[y(corners(k,2,p)) y(corners(k,3,p))],scolorname,'-',linewidth);
                              line2svg(fid,[x(corners(k,4,p)) x(corners(k,5,p))],[y(corners(k,4,p)) y(corners(k,5,p))],scolorname,'-',linewidth);
                          else
                              if strcmp(get(ax,'ZGrid'),'on')
                                  line2svg(fid,[x(corners(k,2,p)) x(corners(k,3,p))],[y(corners(k,2,p)) y(corners(k,3,p))],scolorname,gridlinestyle,linewidth);
                                  line2svg(fid,[x(corners(k,4,p)) x(corners(k,5,p))],[y(corners(k,4,p)) y(corners(k,5,p))],scolorname,gridlinestyle,linewidth);
                              end
                          end
                      end
                end
          end
      end
  end
  fprintf(fid,'    <g>\n');
  axchild = get(ax,'Children');
  if ~FIG2SVG_globals.octave && ~UIverlessthan('8.4.0')
      % Matlab h2 engine
      axchild = [axchild; ax.Title; ax.XLabel; ax.YLabel; ax.ZLabel];
  end
  boundingBoxAxes = axchild2svg(fid,id,axIdString,ax,paperpos,axchild,axpos,groupax,projection,boundingBoxAxes);
  fprintf(fid,'  <clipPath id = "%s">\n',axIdString);
  fprintf(fid,'    <rect x = "%0.3f" y = "%0.3f" width = "%0.3f" height = "%0.3f"/>\n',...
      boundingBoxAxes(1), boundingBoxAxes(2), boundingBoxAxes(3), boundingBoxAxes(4));
  fprintf(fid,'  </clipPath>\n');
  fprintf(fid,'    </g>\n');
  if strcmp(get(ax,'Visible'),'on')
      fprintf(fid,'    <g>\n');
      % Search axis for labeling
      if projection.xyplane
          [~, x_axis_point_index_top] = min(y);
          [~, x_axis_point_index_bottom] = max(y);
          if strcmp(get(ax,'Box'),'on')
              if strcmp(get(ax,'XAxisLocation'),'top')
                  x_axis_point_index = [x_axis_point_index_top x_axis_point_index_bottom];
              else
                  x_axis_point_index = [x_axis_point_index_bottom x_axis_point_index_top];
              end
          else
              if strcmp(get(ax,'XAxisLocation'),'top')
                  x_axis_point_index = x_axis_point_index_top;
              else
                  x_axis_point_index = x_axis_point_index_bottom;
              end
          end
          [~, y_axis_point_index_left] = min(x);
          [~, y_axis_point_index_right] = max(x);
          if strcmp(get(ax,'Box'),'on')
              if strcmp(get(ax,'YAxisLocation'),'right')
                  y_axis_point_index = [y_axis_point_index_right y_axis_point_index_left];
              else
                  y_axis_point_index = [y_axis_point_index_left y_axis_point_index_right];
              end
          else
              if strcmp(get(ax,'YAxisLocation'),'right')
                  y_axis_point_index = y_axis_point_index_right;
              else
                  y_axis_point_index = y_axis_point_index_left;
              end
          end
          [~, z_axis_point_index] = min(x);
      else
          v = get(ax,'view');
          if v(1) == 90 % hack: two max indexes and the relevant is second
            x_axis_point_index = 4;
            y_axis_point_index = 4;
            z_axis_point_index = 2;
          elseif v(1) == 0 % hack: two max indexes and the relevant is second
            [~, x_axis_point_index] = max(y);
            y_axis_point_index = 4;
            [~, z_axis_point_index] = min(x);
          elseif v(1) == 180 % hack: two max indexes and the relevant is second
            [~, x_axis_point_index] = max(y);
            [~, y_axis_point_index] = max(y);
            [~, z_axis_point_index] = min(x);
            z_axis_point_index = 4;
          elseif v(1) == 270 % hack: two max indexes and the relevant is the first (it takes second, most likely by resolution error)
            [~, x_axis_point_index] = max(y);
            [~, y_axis_point_index] = max(y);
            z_axis_point_index = 3;
          else
            [~, x_axis_point_index] = max(y);
            [~, y_axis_point_index] = max(y);
            [~, z_axis_point_index] = min(x);
          end
      end

      % Draw grid
      for pindex = 1:size(front_faces)
          p = front_faces(pindex);
          for k = 1:size(corners,1)
              selectedCorners = squeeze(corners(k,:,p));
              switch corners(k,1,p)
                  case 1 % x
                      % Draw x-grid
                      scolorname = searchcolor(id,get(ax,'XColor'));
                      if strcmp(get(ax,'XGrid'),'on') && gridBehind == false
                          if axlimx(1) ~= axlimx(2)
                              gridLines(fid, x, y, scolorname, gridlinestyle, linewidth, axlimx, axxtick, axxindex_inner, selectedCorners, [2 3 4 5])
                              if strcmp(get(ax,'XTickMode'),'auto') && strcmp(get(ax,'XMinorGrid'),'on') && ~isempty(minor_axxtick)
                                  minorGridLines(fid, x, y, scolorname, minor_gridlinestyle, linewidth, axlimx, minor_axxtick, selectedCorners, [2 3 4 5])
                              end
                          end
                      end
                  case 2 % y
                      % Draw y-grid
                      scolorname = searchcolor(id,get(ax,'YColor'));
                      if strcmp(get(ax,'YGrid'),'on') && gridBehind == false
                          if axlimy(1) ~= axlimy(2)
                              gridLines(fid, x, y, scolorname, gridlinestyle, linewidth, axlimy, axytick, axyindex_inner, selectedCorners, [2 3 4 5])
                              if strcmp(get(ax,'YTickMode'),'auto') && strcmp(get(ax,'YMinorGrid'),'on') && ~isempty(minor_axytick)
                                  minorGridLines(fid, x, y, scolorname, minor_gridlinestyle, linewidth, axlimy, minor_axytick, selectedCorners, [2 3 4 5])
                              end
                          end
                      end
                  case 3 % z
                      % Draw z-grid
                      scolorname = searchcolor(id,get(ax,'ZColor'));
                      if strcmp(get(ax,'ZGrid'),'on') && gridBehind == false
                          if axlimz(1) ~= axlimz(2)
                              gridLines(fid, x, y, scolorname, gridlinestyle, linewidth, axlimz, axztick, axzindex_inner, selectedCorners, [2 3 4 5])
                              if strcmp(get(ax,'ZTickMode'),'auto') && strcmp(get(ax,'ZMinorGrid'),'on') && ~isempty(minor_axztick)
                                  minorGridLines(fid, x, y, scolorname, minor_gridlinestyle, linewidth, axlimz, minor_axztick, selectedCorners, [2 3 4 5])
                              end
                          end
                      end
              end
          end
      end

      scolorname = searchcolor(id,get(ax,'XColor'));
      % Box 'on' 3-D plots don't show the frontal lines
      % % Draw 'box' of x-axis
      % if projection.xyplane == false
      %     if strcmp(get(ax,'Box'),'on')
      %         edge_line_index = [edge_opposite(most_back_edge_index) edge_neighbours(edge_opposite(most_back_edge_index),1)];
      %         line2svg(fid,x(edge_line_index),y(edge_line_index),scolorname,'-',linewidth)
      %     end
      % end
      % Draw x-tick marks
      if (ticklength(1) ~= 0)
          if axlimx(1) ~= axlimx(2)
              if (nomx(x_axis_point_index(1)))
                  lim = [axlimx(2) axlimx(1)];
              else
                  lim = [axlimx(1) axlimx(2)];
              end
              x_label_end1 = interp1([0 1],[x(x_axis_point_index(1)) x(edge_neighbours(x_axis_point_index(1),2))],xlabel_distance,'linear','extrap');
              y_label_end1 = interp1([0 1],[y(x_axis_point_index(1)) y(edge_neighbours(x_axis_point_index(1),2))],xlabel_distance,'linear','extrap');
              x_label_end2 = interp1([0 1],[x(edge_neighbours(x_axis_point_index(1),1)) x(edge_neighbours(edge_neighbours(x_axis_point_index(1),1),2))],xlabel_distance,'linear','extrap');
              y_label_end2 = interp1([0 1],[y(edge_neighbours(x_axis_point_index(1),1)) y(edge_neighbours(edge_neighbours(x_axis_point_index(1),1),2))],xlabel_distance,'linear','extrap');
              xg_label_end = interp1(lim,[x_label_end1 x_label_end2],axxtick);
              yg_label_end = interp1(lim,[y_label_end1 y_label_end2],axxtick);
              frontTicks(fid, x, y, scolorname, linewidth, ...
                  axxtick, x_axis_point_index, edge_neighbours, [2 1 1], ...
                  valid_xsticks,  ticklength, tick_ratio, lim, true);
              if strcmp(get(ax,'XTickMode'),'auto') && (strcmp(get(ax,'XMinorGrid'),'on') || strcmp(get(ax,'XScale'),'log')) && ~isempty(minor_axxtick)
                  frontTicks(fid, x, y, scolorname, linewidth, ...
                      minor_axxtick, x_axis_point_index, edge_neighbours, [2 1 1], ...
                      1:length(minor_axxtick),  0.5 * ticklength, tick_ratio, lim, false);
              end
              if ~isempty(axlabelx) && ~(iscell(axlabelx) && all(cellfun(@isempty,axlabelx)))
                  if ischar(axlabelx) && size(axlabelx, 1) == 1
                      % Special handling of 1xn char arrays -> duplicate data
                      % for all ticks. Strange behavior but follows the
                      % behavior of Matlab
                      axlabelx = repmat(axlabelx, length(axxindex), 1);
                  end
                  if (strcmp(get(ax,'XTickLabelMode'),'manual'))
                      axlabelx = axlabelx(axxindex,:);
                  end
                  if FIG2SVG_globals.octave
                    axlabelx = axlabelx(:); % SA: Octave compatibility
                  end
                  if strcmp(get(ax,'XAxisLocation'),'top') && (projection.xyplane == true)
                    [angle, align] = improvedXLabel(ax, 0, 'center');
                    for i = 1:length(axxindex)
                      label2svg(fid,axpos,ax,xg_label_end(i),yg_label_end(i),convertString(axlabelx(i,:)),align,angle,'bottom',1,scolorname);
                    end
                  elseif projection.xyplane == true
                    [angle, align] = improvedXLabel(ax, 0, 'center');
                    for i = 1:length(axxindex)
                      label2svg(fid,axpos,ax,xg_label_end(i),yg_label_end(i),convertString(axlabelx(i,:)),align,angle,'top',1,scolorname);
                    end
                  else
                    v = pi/180*get(ax,'view');
                    if sin(v(1))*cos(v(1)) > 1e-10 && ( sin(v(1)) >= 0.2695 && cos(v(1)) > -1e-10 || sin(v(1)) < -0.2695 )
                      % disp('right')
                      [angle, align] = improvedXLabel(ax, 0, 'right');
                    elseif sin(v(1))*cos(v(1)) < 1e-10 && ( sin(v(1)) <= -0.2695 || ( cos(v(1)) < 1e-10 && cos(v(1)) > -0.963 ) )
                      % disp('left')
                      [angle, align] = improvedXLabel(ax, 0, 'left');
                    else
                      % disp('center')
                      [angle, align] = improvedXLabel(ax, 0, 'center');
                    end
                    if abs(cos(v(1))) <= 0.124
                      % disp('middle')
                      for i = 1:length(axxindex)
                        label2svg(fid,axpos,ax,xg_label_end(i),yg_label_end(i),convertString(axlabelx(i,:)),align,angle,'middle',1,scolorname);
                      end
                    else
                      % disp('top')
                      for i = 1:length(axxindex)
                        label2svg(fid,axpos,ax,xg_label_end(i),yg_label_end(i),convertString(axlabelx(i,:)),align,angle,'top',1,scolorname);
                      end
                    end
                  end
              end
          end
      end

      scolorname = searchcolor(id,get(ax,'YColor'));
      % Box 'on' 3-D plots don't show the frontal lines
      % % Draw 'box' of y-axis
      % if projection.xyplane == false
      %     if strcmp(get(ax,'Box'),'on')
      %         edge_line_index = [edge_opposite(most_back_edge_index) edge_neighbours(edge_opposite(most_back_edge_index),2)];
      %         line2svg(fid,x(edge_line_index),y(edge_line_index),scolorname,'-',linewidth)
      %     end
      % end
      % Draw y-tick marks
      if (ticklength(1) ~= 0)
          if axlimy(1) ~= axlimy(2)
              if (nomy(y_axis_point_index(1)))
                  lim = [axlimy(2) axlimy(1)];
              else
                  lim = [axlimy(1) axlimy(2)];
              end
              x_label_end1 = interp1([0 1],[x(y_axis_point_index(1)) x(edge_neighbours(y_axis_point_index(1),1))],ylabel_distance,'linear','extrap');
              y_label_end1 = interp1([0 1],[y(y_axis_point_index(1)) y(edge_neighbours(y_axis_point_index(1),1))],ylabel_distance,'linear','extrap');
              x_label_end2 = interp1([0 1],[x(edge_neighbours(y_axis_point_index(1),2)) x(edge_neighbours(edge_neighbours(y_axis_point_index(1),2),1))],ylabel_distance,'linear','extrap');
              y_label_end2 = interp1([0 1],[y(edge_neighbours(y_axis_point_index(1),2)) y(edge_neighbours(edge_neighbours(y_axis_point_index(1),2),1))],ylabel_distance,'linear','extrap');
              xg_label_end = interp1(lim,[x_label_end1 x_label_end2],axytick);
              yg_label_end = interp1(lim,[y_label_end1 y_label_end2],axytick);
              frontTicks(fid, x, y, scolorname, linewidth, ...
                  axytick, y_axis_point_index, edge_neighbours, [1 2 2], ...
                  valid_ysticks,  ticklength, tick_ratio, lim, true);
              if strcmp(get(ax,'YTickMode'),'auto') && (strcmp(get(ax,'YMinorGrid'),'on') || strcmp(get(ax,'YScale'),'log')) && ~isempty(minor_axytick)
                  frontTicks(fid, x, y, scolorname, linewidth, ...
                      minor_axytick, y_axis_point_index, edge_neighbours, [1 2 2], ...
                      1:length(minor_axytick), 0.5 * ticklength, tick_ratio, lim, false);
              end
              if ~isempty(axlabely) && ~(iscell(axlabely) && all(cellfun(@isempty,axlabely)))
                  if ischar(axlabely) && size(axlabely, 1) == 1
                      % Special handling of 1xn char arrays -> duplicate data
                      % for all ticks. Strange behavior but follows the
                      % behavior of Matlab
                      axlabely = repmat(axlabely, length(axyindex), 1);
                  end
                  if (strcmp(get(ax,'YTickLabelMode'),'manual'))
                      axlabely = axlabely(axyindex,:);
                  end
                  if FIG2SVG_globals.octave
                    axlabely = axlabely(:); % SA: Octave compatibility
                  end
                  if (projection.xyplane == true)
                      if strcmp(get(ax,'YAxisLocation'),'right')
                          [angle, align] = improvedYLabel(ax, 0, 'left');
                          for i = 1:length(axyindex)
                              label2svg(fid,axpos,ax,xg_label_end(i),yg_label_end(i),convertString(axlabely(i,:)),align,angle,'middle',1,scolorname);
                          end
                     else
                          [angle, align] = improvedYLabel(ax, 0, 'right');
                          for i = 1:length(axyindex)
                              label2svg(fid,axpos,ax,xg_label_end(i),yg_label_end(i),convertString(axlabely(i,:)),align,angle,'middle',1,scolorname);
                          end
                      end
                  else
                    v = pi/180*get(ax,'view');
                    if sin(v(1))*cos(v(1)) > -1e-10 && ( cos(v(1)) <= -0.2339 && sin(v(1)) < 1e-10 || cos(v(1)) > 0.2339)
                      % disp('left')
                      [angle, align] = improvedYLabel(ax, 0, 'left');
                    elseif sin(v(1))*cos(v(1)) < -1e-10 && ( ( sin(v(1)) < 1e-10 && sin(v(1)) > -0.9723 ) || cos(v(1)) <= -0.2339 )
                      % disp('right')
                      [angle, align] = improvedYLabel(ax, 0, 'right');
                    else
                      % disp('center')
                      [angle, align] = improvedYLabel(ax, 0, 'center');
                    end
                    if abs(sin(v(1))) <= 0.124
                      % disp('middle')
                      for i = 1:length(axyindex)
                        label2svg(fid,axpos,ax,xg_label_end(i),yg_label_end(i),convertString(axlabely(i,:)),align,angle,'middle',1,scolorname);
                      end
                    else
                      % disp('top')
                      for i = 1:length(axyindex)
                        label2svg(fid,axpos,ax,xg_label_end(i),yg_label_end(i),convertString(axlabely(i,:)),align,angle,'top',1,scolorname);
                      end
                    end
                  end
              end
          end
      end

      scolorname = searchcolor(id,get(ax,'ZColor'));
      % Box 'on' 3-D plots don't show the frontal lines
      % % Draw 'box' of z-axis
      % if projection.xyplane == false
      %     if strcmp(get(ax,'Box'),'on')
      %         edge_line_index = [edge_opposite(most_back_edge_index) edge_neighbours(edge_opposite(most_back_edge_index),3)];
      %         line2svg(fid,x(edge_line_index),y(edge_line_index),scolorname,'-',linewidth)
      %     end
      % end
      % Draw z-tick marks
      if (ticklength(1) ~= 0)
          if axlimz(1) ~= axlimz(2)
              if (nomz(z_axis_point_index(1)))
                  lim = [axlimz(2) axlimz(1)];
              else
                  lim = [axlimz(1) axlimz(2)];
              end
              x_tick_end1 = interp1([0 1],[x(z_axis_point_index) x(edge_neighbours(z_axis_point_index,2))],ticklength*tick_ratio(3),'linear','extrap');
              x_tick_end2 = interp1([0 1],[x(edge_neighbours(z_axis_point_index,3)) x(edge_neighbours(edge_neighbours(z_axis_point_index,3),2))],ticklength*tick_ratio(3),'linear','extrap');
              x_label_end1 = interp1([0 1],[x(z_axis_point_index) x(edge_neighbours(z_axis_point_index,2))],zlabel_distance,'linear','extrap');
              x_label_end2 = interp1([0 1],[x(edge_neighbours(z_axis_point_index,3)) x(edge_neighbours(edge_neighbours(z_axis_point_index,3),2))],zlabel_distance,'linear','extrap');
              xg_line_start = interp1(lim,[x(z_axis_point_index) x(edge_neighbours(z_axis_point_index,3))],axztick);
              xg_line_end = interp1(lim,[x_tick_end1 x_tick_end2],axztick);
              xg_label_end = interp1(lim,[x_label_end1 x_label_end2],axztick);

              v = get(ax,'view');
              if v(1) >= 0 && v(1) < 90 || v(1) >= 180 && v(1) < 270
                tick_ratio(3) = -tick_ratio(3);
                zlabel_distance = -zlabel_distance;
              end

              y_tick_end1 = interp1([0 1],[y(z_axis_point_index) y(edge_neighbours(z_axis_point_index,2))],ticklength*tick_ratio(3),'linear','extrap');
              y_tick_end2 = interp1([0 1],[y(edge_neighbours(z_axis_point_index,3)) y(edge_neighbours(edge_neighbours(z_axis_point_index,3),2))],ticklength*tick_ratio(3),'linear','extrap');
              y_label_end1 = interp1([0 1],[y(z_axis_point_index) y(edge_neighbours(z_axis_point_index,2))],zlabel_distance,'linear','extrap');
              y_label_end2 = interp1([0 1],[y(edge_neighbours(z_axis_point_index,3)) y(edge_neighbours(edge_neighbours(z_axis_point_index,3),2))],zlabel_distance,'linear','extrap');

              yg_line_start = interp1(lim,[y(z_axis_point_index) y(edge_neighbours(z_axis_point_index,3))],axztick);
              yg_line_end = interp1(lim,[y_tick_end1 y_tick_end2],axztick);
              yg_label_end = interp1(lim,[y_label_end1 y_label_end2],axztick);

              tol = 1e-10;
              if any(abs([diff(xg_line_start) diff(xg_line_end) diff(yg_line_start) diff(yg_line_end)]) > tol) % SA: looks better without and not used anymore in Matlab
                for i = valid_zsticks
                    line2svg(fid,[xg_line_start(i) xg_line_end(i)],[yg_line_start(i) yg_line_end(i)],scolorname,'-',linewidth)
                end
              end
              line2svg(fid,[x(z_axis_point_index) x(edge_neighbours(z_axis_point_index,3))],[y(z_axis_point_index) y(edge_neighbours(z_axis_point_index,3))],scolorname,'-',linewidth)
              if ~isempty(axlabelz) && ~(iscell(axlabelz) && all(cellfun(@isempty,axlabelz)))
                  if ischar(axlabelz) && size(axlabelz, 1) == 1
                      % Special handling of 1xn char arrays -> duplicate data
                      % for all ticks. Strange behavior but follows the
                      % behavior of Matlab
                      axlabelz = repmat(axlabelz, length(axzindex), 1);
                  end
                  if (strcmp(get(ax,'ZTickLabelMode'),'manual'))
                      axlabelz = axlabelz(axzindex,:);
                  end
                  if FIG2SVG_globals.octave
                    axlabelz = axlabelz(:); % SA: Octave compatibility
                  end
                  [angle, align] = improvedZLabel(ax, 0, 'right');
                  for i = 1:length(axzindex)
                      label2svg(fid,axpos,ax,xg_label_end(i),yg_label_end(i),convertString(axlabelz(i,:)),align,angle,'middle',1,scolorname);
                  end
              end
          end
      end

      exponent2svg(fid,axpos,paperpos,ax,axxtick,axytick,axztick)
      fprintf(fid,'    </g>\n');
  end
  fprintf(fid,'  </g>\n');
  set(ax,'Units',originalAxesUnits);
end

%%%%%%%%%%%%%%%%%%%%%%%%%%%%%%%%%%%%%%%%%%%%%%%%%%%%%%%%%%%%%%%%%%%%%%%%%
% take any axis children and create objects for them
function boundingBoxAxes = axchild2svg(fid,id,axIdString,ax,paperpos,axchild,axpos,groupax,projection,boundingBoxAxes)
  global colorname
  global FIG2SVG_globals

  % trick for new bar plots
  numChildBars = sum(strcmp(get(axchild(:),'Type'),'bar'));
  currentBarSetNumber = 0;
  stackedBarYref = 0;

  for i = length(axchild):-1:1
      if FIG2SVG_globals.debugModeOn
        currentType = get(axchild(i),'Type');
        disp(['    axchild(',num2str(i),') = ', currentType]);
      end
      if strcmp(get(axchild(i), 'Visible'), 'off')
          % do nothing
      elseif strcmp(get(axchild(i),'Type'),'line') || strcmp(get(axchild(i),'Type'),'errorbar')
          scolorname = searchcolor(id,get(axchild(i),'Color'));
          linestyle = get(axchild(i),'LineStyle');
          linewidth = get(axchild(i),'LineWidth'); % linewidth = FIG2SVG_globals.resolutionScaling*get(axchild(i),'LineWidth');
          marker = get(axchild(i),'Marker');
          markeredgecolor = get(axchild(i),'MarkerEdgeColor');
          if ischar(markeredgecolor)
              switch markeredgecolor
                  case 'none',markeredgecolorname = 'none';
                  otherwise,markeredgecolorname = scolorname;  % if markeredgecolorname is 'auto' or something else set the markeredgecolorname to the line color
              end
          else
              markeredgecolorname = searchcolor(id,markeredgecolor);
          end
          markerfacecolor = get(axchild(i),'MarkerFaceColor');
          if ischar(markerfacecolor)
              switch markerfacecolor
                  case 'none',markerfacecolorname = 'none';
                  otherwise,markerfacecolorname = scolorname;  % if markerfacecolorname is 'auto' or something else set the markerfacecolorname to the line color
              end
          else
              markerfacecolorname = searchcolor(id,markerfacecolor);
          end

          try % not currently implemented, but maybe it will in the future
            markerFaceAlpha = get(axchild(i),'MarkerFaceAlpha');
            markerEdgeAlpha = get(axchild(i),'MarkerEdgeAlpha');
          catch
            markerFaceAlpha = 1;
            markerEdgeAlpha = 1;
          end

          markersize = 2/3*get(axchild(i),'MarkerSize');
          linex = get(axchild(i),'XData');
          linex = linex(:)';
          liney = get(axchild(i),'YData');
          liney = liney(:)';
          if ~strcmp(get(axchild(i),'Type'),'errorbar')
            linez = get(axchild(i),'ZData');
            linez = linez(:)';
          else
            linez = zeros(size(linex));
            xnegdelta = get(axchild(i),'XNegativeDelta');
            xposdelta = get(axchild(i),'XPositiveDelta');
            if ~isempty(xnegdelta)
              xnegdelta = linex - xnegdelta;
              xposdelta = linex + xposdelta;
            end
            ynegdelta = get(axchild(i),'YNegativeDelta');
            yposdelta = get(axchild(i),'YPositiveDelta');
            if ~isempty(ynegdelta)
              ynegdelta = liney - ynegdelta;
              yposdelta = liney + yposdelta;
            end
            capsize = get(axchild(i),'CapSize');
          end
          try
            if strcmp(get(ax,'XScale'),'log')
                linex(linex <= 0) = NaN;
                linex = log10(linex);
                if strcmp(get(axchild(i),'Type'),'errorbar')
                  xnegdelta(xnegdelta <= 0) = NaN;
                  xnegdelta = log10(xnegdelta);
                  xposdelta(xposdelta <= 0) = NaN;
                  xposdelta = log10(xposdelta);
                end
            end
            if strcmp(get(ax,'YScale'),'log')
                liney(liney <= 0) = NaN;
                liney = log10(liney);
                if strcmp(get(axchild(i),'Type'),'errorbar')
                  ynegdelta(ynegdelta <= 0) = NaN;
                  ynegdelta = log10(ynegdelta);
                  yposdelta(yposdelta <= 0) = NaN;
                  yposdelta = log10(yposdelta);
                end
            end
            if ~strcmp(get(axchild(i),'Type'),'errorbar')
              if isempty(linez)
                linez = zeros(size(linex));
              end
              if strcmp(get(ax,'ZScale'),'log')
                linez(linez <= 0) = NaN;
                linez = log10(linez);
              end
            end
            [x,y,~] = project(linex,liney,linez,projection);
            if strcmp(get(axchild(i),'Type'),'errorbar')
              if ~isempty(xnegdelta)
                xerrorbar_linex = [xnegdelta;xposdelta;nan(size(xnegdelta))];
                xerrorbar_linex = xerrorbar_linex(:)';
                xerrorbar_liney = [liney;liney;nan(size(xnegdelta))];
                xerrorbar_liney = xerrorbar_liney(:)';
                xerrorbar_linez = [linez;linez;nan(size(xnegdelta))];
                xerrorbar_linez = xerrorbar_linez(:)';
                [xerrorbar_x,xerrorbar_y,~] = project(xerrorbar_linex,xerrorbar_liney,xerrorbar_linez,projection);
              end
              if ~isempty(ynegdelta)
                yerrorbar_liney = [ynegdelta;yposdelta;nan(size(ynegdelta))];
                yerrorbar_liney = yerrorbar_liney(:)';
                yerrorbar_linex = [linex;linex;nan(size(ynegdelta))];
                yerrorbar_linex = yerrorbar_linex(:)';
                yerrorbar_linez = [linez;linez;nan(size(ynegdelta))];
                yerrorbar_linez = yerrorbar_linez(:)';
                [yerrorbar_x,yerrorbar_y,~] = project(yerrorbar_linex,yerrorbar_liney,yerrorbar_linez,projection);
              end
            end
          catch
            % children of legend objects
            x = linex;
            y = liney;
            if ~strcmp(get(axchild(i),'Type'),'errorbar')
              z = linez;
            end
          end
          x = (x*axpos(3)+axpos(1))*paperpos(3);
          y = (1-(y*axpos(4)+axpos(2)))*paperpos(4);
          if strcmp(get(axchild(i),'Type'),'errorbar')
            if ~isempty(xnegdelta)
              xerrorbar_x = (xerrorbar_x*axpos(3)+axpos(1))*paperpos(3);
              xerrorbar_y = (1-(xerrorbar_y*axpos(4)+axpos(2)))*paperpos(4);
              xcapsize_y = bsxfun(@plus,0.5*capsize*[1;-1;NaN]*ones(size(xerrorbar_y(~isnan(xerrorbar_y)))),xerrorbar_y(~isnan(xerrorbar_y)));
              xcapsize_y = xcapsize_y(:)';
              xcapsize_x = bsxfun(@times,[1;1;NaN]*ones(size(xerrorbar_x(~isnan(xerrorbar_x)))),xerrorbar_x(~isnan(xerrorbar_x)));
              xcapsize_x = xcapsize_x(:)';
            else
              xerrorbar_x = [];
              xerrorbar_y = [];
              xcapsize_y = [];
              xcapsize_x = [];
            end
            if ~isempty(ynegdelta)
              yerrorbar_x = (yerrorbar_x*axpos(3)+axpos(1))*paperpos(3);
              yerrorbar_y = (1-(yerrorbar_y*axpos(4)+axpos(2)))*paperpos(4);
              ycapsize_x = bsxfun(@plus,0.5*capsize*[1;-1;NaN]*ones(size(yerrorbar_x(~isnan(yerrorbar_x)))),yerrorbar_x(~isnan(yerrorbar_x)));
              ycapsize_x = ycapsize_x(:)';
              ycapsize_y = bsxfun(@times,[1;1;NaN]*ones(size(yerrorbar_y(~isnan(yerrorbar_y)))),yerrorbar_y(~isnan(yerrorbar_y)));
              ycapsize_y = ycapsize_y(:)';
            else
              yerrorbar_x = [];
              yerrorbar_y = [];
              ycapsize_x = [];
              ycapsize_y = [];
            end
          end

          markerOverlap = 0;
          if ~strcmp(linestyle, 'none') && ~strcmp(get(axchild(i),'Type'),'errorbar')
              markerOverlap = max(markerOverlap, convertunit(linewidth*0.5, 'points', 'pixels', axpos(4)));
          end
          if ~strcmp(marker, 'none')
              markerOverlap = max(markerOverlap, convertunit(markersize, 'points', 'pixels', axpos(4)));
          end
          % put a line into a group with its markers
          if FIG2SVG_globals.ClippingMode ~= 2
            if ~strcmp(get(axchild(i),'Type'),'errorbar')
              boundingBoxElement = [min(x)-markerOverlap min(y)-markerOverlap max(x)-min(x)+2*markerOverlap max(y)-min(y)+2*markerOverlap];
            else
              boundingBoxElement = [min([x,xerrorbar_x,xcapsize_y])-markerOverlap min([y,yerrorbar_y,ycapsize_x])-markerOverlap max([x,xerrorbar_x,xcapsize_y])-min([x,xerrorbar_x,xcapsize_y])+2*markerOverlap max([y,yerrorbar_y,ycapsize_x])-min([y,yerrorbar_y,ycapsize_x])+2*markerOverlap];
            end
          else
            boundingBoxElement = boundingBoxAxes;
          end
          [filterString, boundingBox] = filter2svg(fid, axchild(i), boundingBoxAxes, boundingBoxElement);
          if FIG2SVG_globals.ClippingMode == 1 && isfield(FIG2SVG_globals,'BoxOn') && ~FIG2SVG_globals.BoxOn
            boundingBoxAxes = [boundingBoxAxes(1) min([boundingBoxAxes(2) boundingBoxElement(2)]) max([boundingBoxAxes(3)+max([boundingBoxAxes(1)-boundingBoxElement(1),0]) boundingBoxElement(3)+max([boundingBoxElement(1)-boundingBoxAxes(1),0])]) [boundingBoxAxes(4)+max([boundingBoxAxes(2)-boundingBoxElement(2),0])]];
          elseif FIG2SVG_globals.ClippingMode == 3 || FIG2SVG_globals.ClippingMode == 1 && ~isfield(FIG2SVG_globals,'BoxOn')
            boundingBoxAxes = [min([boundingBoxAxes(1) boundingBoxElement(1)]) min([boundingBoxAxes(2) boundingBoxElement(2)]) max([boundingBoxAxes(3)+max([boundingBoxAxes(1)-boundingBoxElement(1),0]) boundingBoxElement(3)+max([boundingBoxElement(1)-boundingBoxAxes(1),0])]) max([boundingBoxAxes(4)+max([boundingBoxAxes(2)-boundingBoxElement(2),0]) boundingBoxElement(4)+max([boundingBoxElement(2)-boundingBoxAxes(2),0])])];
          end
          if strcmp(get(axchild(i),'Clipping'),'on') && FIG2SVG_globals.ClippingMode ~= 0
            clippingIdString = clipping2svg(fid, axchild(i), ax, paperpos, axpos, projection, axIdString);
            fprintf(fid,'<g id = "%s" clip-path = "url(#%s)" %s>\n', createId, clippingIdString, filterString);
          else
            fprintf(fid,'<g id = "%s" %s>\n', createId, filterString);
          end
          if ~isempty(filterString)
              % Workaround for Inkscape filter bug
              fprintf(fid,'<rect x = "%0.3f" y = "%0.3f" width = "%0.3f" height = "%0.3f" fill = "none" stroke = "none" />\n', boundingBox(1), boundingBox(2), boundingBox(3), boundingBox(4));
          end
          line2svg(fid,x,y,scolorname,linestyle,linewidth,'round')
          if strcmp(get(axchild(i),'Type'),'errorbar')
            line2svg(fid,xerrorbar_x,xerrorbar_y,scolorname,'-',linewidth,'round')
            line2svg(fid,xcapsize_x,xcapsize_y,scolorname,'-',linewidth,'round')
            line2svg(fid,yerrorbar_x,yerrorbar_y,scolorname,'-',linewidth,'round')
            line2svg(fid,ycapsize_x,ycapsize_y,scolorname,'-',linewidth,'round')
          end

          % put the markers into a subgroup of the lines
          if  ~strcmp(marker, 'none') % but only do it if we actually are drawing markers
              fprintf(fid,'<g>\n');
              switch marker
                  case 'none'
                  case '.';circle2svg(fid,x,y,markersize/3,'none',markeredgecolorname,linewidth, 1, markerEdgeAlpha);
                  case 'o',circle2svg(fid,x,y,markersize,markeredgecolorname,markerfacecolorname,linewidth, markerFaceAlpha, markerEdgeAlpha);
                  case '+',patch2svg(fid,x'*ones(1,5)+ones(length(linex),1)*[-1 1 NaN 0 0]*0.85*markersize,y'*ones(1,5)+ones(length(liney),1)*[0 0 NaN -1 1]*0.85*markersize,markeredgecolorname,'-',linewidth,markeredgecolorname, markerFaceAlpha, markerEdgeAlpha, false);
                  case '*',patch2svg(fid,x'*ones(1,11)+ones(length(linex),1)*[-1 1 NaN 0 0 NaN -0.7 0.7 NaN -0.7 0.7]*0.85*markersize,y'*ones(1,11)+ones(length(liney),1)*[0 0 NaN -1 1 NaN 0.7 -0.7 NaN -0.7 0.7]*0.85*markersize,markeredgecolorname,'-',linewidth,markeredgecolorname, markerFaceAlpha, markerEdgeAlpha, false);
                  case 'x',patch2svg(fid,x'*ones(1,5)+ones(length(linex),1)*[-0.7 0.7 NaN -0.7 0.7]*0.8*markersize,y'*ones(1,5)+ones(length(liney),1)*[0.7 -0.7 NaN -0.7 0.7]*0.8*markersize,markeredgecolorname,'-',linewidth,markeredgecolorname, markerFaceAlpha, markerEdgeAlpha, false);
                  %% Octave keeps s, d, p and h in the HandleGraphics object, for the square, diamond, pentagram, and hexagram markers, respectively -- Jakob Malm
                  case {'square', 's'},patch2svg(fid,x'*ones(1,5)+ones(length(linex),1)*[-1 -1 1 1 -1]*0.75*markersize,y'*ones(1,5)+ones(length(liney),1)*[-1 1 1 -1 -1]*0.75*markersize,markerfacecolorname,'-',linewidth,markeredgecolorname, markerFaceAlpha, markerEdgeAlpha, true);
                  case {'diamond', 'd'},patch2svg(fid,x'*ones(1,5)+ones(length(linex),1)*[-0.7071 0 0.7071 0 -0.7071]*1.35*markersize,y'*ones(1,5)+ones(length(liney),1)*[0 1 0 -1 0]*1.35*markersize,markerfacecolorname,'-',linewidth,markeredgecolorname, markerFaceAlpha, markerEdgeAlpha, true);
                  case {'pentagram', 'p'},patch2svg(fid,...
                          x'*ones(1,11)+ones(length(linex),1)*[0 0.1180 0.5 0.1910 0.3090 0 -0.3090 -0.1910 -0.5 -0.1180 0]*2*markersize,...
                          y'*ones(1,11)+ones(length(liney),1)*[-0.5257 -0.1625 -0.1625 0.0621 0.4253 0.2008 0.4253 0.0621 -0.1625 -0.1625 -0.5257]*2*markersize,markerfacecolorname,'-',linewidth,markeredgecolorname, markerFaceAlpha, markerEdgeAlpha, true);
                  case {'hexagram', 'h'},patch2svg(fid,...
                          x'*ones(1,13)+ones(length(linex),1)*[0 0.2309 0.6928 0.4619 0.6928 0.2309 0 -0.2309 -0.6928 -0.4619 -0.6928 -0.2309 0]*1.3*markersize,...
                          y'*ones(1,13)+ones(length(liney),1)*[0.8 0.4 0.4 0 -0.4 -0.4 -0.8 -0.4 -0.4 0 0.4 0.4 0.8]*1.3*markersize,markerfacecolorname,'-',linewidth,markeredgecolorname, markerFaceAlpha, markerEdgeAlpha, true);
                  case '^',patch2svg(fid,x'*ones(1,4)+ones(length(linex),1)*[-1 1 0 -1]*1.15*markersize,y'*ones(1,4)+ones(length(liney),1)*[0.577 0.577 -1.155 0.577]*1.15*markersize,markerfacecolorname,'-',linewidth,markeredgecolorname, markerFaceAlpha, markerEdgeAlpha, true);
                  case 'v',patch2svg(fid,x'*ones(1,4)+ones(length(linex),1)*[-1 1 0 -1]*1.15*markersize,y'*ones(1,4)+ones(length(liney),1)*[-0.577 -0.577 1.155 -0.577]*1.15*markersize,markerfacecolorname,'-',linewidth,markeredgecolorname, markerFaceAlpha, markerEdgeAlpha, true);
                  case '<',patch2svg(fid,x'*ones(1,4)+ones(length(linex),1)*[0.577 0.577 -1.155 0.577]*1.15*markersize,y'*ones(1,4)+ones(length(liney),1)*[-1 1 0 -1]*1.15*markersize,markerfacecolorname,'-',linewidth,markeredgecolorname, markerFaceAlpha, markerEdgeAlpha, true);
                  case '>',patch2svg(fid,x'*ones(1,4)+ones(length(linex),1)*[-0.577 -0.577 1.155 -0.577]*1.15*markersize,y'*ones(1,4)+ones(length(liney),1)*[-1 1 0 -1]*1.15*markersize,markerfacecolorname,'-',linewidth,markeredgecolorname, markerFaceAlpha, markerEdgeAlpha, true);
              end
              % close the marker group
              fprintf(fid,'</g>\n');
          end
          animation2svg(fid, axchild(i));
          % close the line group
          fprintf(fid,'</g>\n');
      elseif strcmp(get(axchild(i),'Type'),'scatter')
          linewidth = get(axchild(i),'LineWidth'); % linewidth = FIG2SVG_globals.resolutionScaling*get(axchild(i),'LineWidth');
          marker = get(axchild(i),'Marker');

          if strcmp(get(axchild(i),'MarkerEdgeColor'),'flat') || strcmp(get(axchild(i),'MarkerFaceColor'),'flat')
              cData = get(axchild(i),'CData');
              clim = get(ax,'CLim');
              cmap = get(id,'Colormap');
              pointsc = round((cData-clim(1))/(clim(2)-clim(1))*(size(cmap,1)-1)+1);
              % Limit index to smallest or biggest color index
              pointsc = max(pointsc,1);
              pointsc = min(pointsc,size(cmap,1));
          end

          if strcmp(get(axchild(i),'MarkerEdgeColor'),'flat')
              for pointc = 1:numel(pointsc)
                  c_tmp = cmap(pointsc,:);
                  markeredgecolorname{pointc} = searchcolor(id,c_tmp(pointc,:));
              end
              markeredgecolorname = markeredgecolorname';
          else
              markeredgecolor = get(axchild(i),'MarkerEdgeColor');
              markeredgecolorname = searchcolor(id,markeredgecolor);
          end

          if strcmp(get(axchild(i),'MarkerFaceColor'),'flat')
              for pointc = 1:numel(pointsc)
                  c_tmp = cmap(pointsc,:);
                  markerfacecolorname{pointc} = searchcolor(id,c_tmp(pointc,:));
              end
              markerfacecolorname = markerfacecolorname';
          else
              markerfacecolor = get(axchild(i),'MarkerFaceColor');
              markerfacecolorname = searchcolor(id,markerfacecolor);
          end

          markerFaceAlpha = get(axchild(i),'MarkerFaceAlpha');
          markerEdgeAlpha = get(axchild(i),'MarkerEdgeAlpha');

          markersize = 2/3/10*max([60, get(axchild(i),'SizeData')]);

          linex = get(axchild(i),'XData');
          linex = linex(:)';
          liney = get(axchild(i),'YData');
          liney = liney(:)';
          linez = get(axchild(i),'ZData');
          linez = linez(:)';
          try
            if strcmp(get(ax,'XScale'),'log')
                linex(linex <= 0) = NaN;
                linex = log10(linex);
            end
            if strcmp(get(ax,'YScale'),'log')
                liney(liney <= 0) = NaN;
                liney = log10(liney);
            end
            if isempty(linez)
              linez = zeros(size(linex));
            end
            if strcmp(get(ax,'ZScale'),'log')
              linez(linez <= 0) = NaN;
              linez = log10(linez);
            end
            [x,y,~] = project(linex,liney,linez,projection);
          catch
            % children of legend objects
            x = linex;
            y = liney;
            z = linez;
          end
          x = (x*axpos(3)+axpos(1))*paperpos(3);
          y = (1-(y*axpos(4)+axpos(2)))*paperpos(4);

          markerOverlap = 0;
          markerOverlap = max(markerOverlap, convertunit(linewidth*0.5, 'points', 'pixels', axpos(4)));
          markerOverlap = max(markerOverlap, convertunit(markersize, 'points', 'pixels', axpos(4)));

          % put a line into a group with its markers
          if FIG2SVG_globals.ClippingMode ~= 2
            boundingBoxElement = [min(x)-markerOverlap min(y)-markerOverlap max(x)-min(x)+2*markerOverlap max(y)-min(y)+2*markerOverlap];
          else
            boundingBoxElement = boundingBoxAxes;
          end
          [filterString, boundingBox] = filter2svg(fid, axchild(i), boundingBoxAxes, boundingBoxElement);
          if FIG2SVG_globals.ClippingMode == 1 && isfield(FIG2SVG_globals,'BoxOn') && ~FIG2SVG_globals.BoxOn
            boundingBoxAxes = [boundingBoxAxes(1) min([boundingBoxAxes(2) boundingBoxElement(2)]) max([boundingBoxAxes(3)+max([boundingBoxAxes(1)-boundingBoxElement(1),0]) boundingBoxElement(3)+max([boundingBoxElement(1)-boundingBoxAxes(1),0])]) [boundingBoxAxes(4)+max([boundingBoxAxes(2)-boundingBoxElement(2),0])]];
          elseif FIG2SVG_globals.ClippingMode == 3 || FIG2SVG_globals.ClippingMode == 1 && ~isfield(FIG2SVG_globals,'BoxOn')
            boundingBoxAxes = [min([boundingBoxAxes(1) boundingBoxElement(1)]) min([boundingBoxAxes(2) boundingBoxElement(2)]) max([boundingBoxAxes(3)+max([boundingBoxAxes(1)-boundingBoxElement(1),0]) boundingBoxElement(3)+max([boundingBoxElement(1)-boundingBoxAxes(1),0])]) max([boundingBoxAxes(4)+max([boundingBoxAxes(2)-boundingBoxElement(2),0]) boundingBoxElement(4)+max([boundingBoxElement(2)-boundingBoxAxes(2),0])])];
          end
          if strcmp(get(axchild(i),'Clipping'),'on') && FIG2SVG_globals.ClippingMode ~= 0
            clippingIdString = clipping2svg(fid, axchild(i), ax, paperpos, axpos, projection, axIdString);
            fprintf(fid,'<g id = "%s" clip-path = "url(#%s)" %s>\n', createId, clippingIdString, filterString);
          else
            fprintf(fid,'<g id = "%s" %s>\n', createId, filterString);
          end
          if ~isempty(filterString)
              % Workaround for Inkscape filter bug
              fprintf(fid,'<rect x = "%0.3f" y = "%0.3f" width = "%0.3f" height = "%0.3f" fill = "none" stroke = "none" />\n', boundingBox(1), boundingBox(2), boundingBox(3), boundingBox(4));
          end

          % put the markers into a subgroup of the lines
          if  ~strcmp(marker, 'none') % but only do it if we actually are drawing markers
              fprintf(fid,'<g>\n');
              switch marker
                  case 'none'
                  case '.';circle2svg(fid,x,y,markersize/3,'none',markeredgecolorname,linewidth, 1, markerEdgeAlpha);
                  case 'o',circle2svg(fid,x,y,markersize,markeredgecolorname,markerfacecolorname,linewidth, markerFaceAlpha, markerEdgeAlpha);
                  case '+',scatterpatch2svg(fid,x'*ones(1,5)+ones(length(linex),1)*[-1 1 NaN 0 0]*0.85*markersize,y'*ones(1,5)+ones(length(liney),1)*[0 0 NaN -1 1]*0.85*markersize,markeredgecolorname,'-',linewidth,markeredgecolorname, markerFaceAlpha, markerEdgeAlpha, false);
                  case '*',scatterpatch2svg(fid,x'*ones(1,11)+ones(length(linex),1)*[-1 1 NaN 0 0 NaN -0.7 0.7 NaN -0.7 0.7]*0.85*markersize,y'*ones(1,11)+ones(length(liney),1)*[0 0 NaN -1 1 NaN 0.7 -0.7 NaN -0.7 0.7]*0.85*markersize,markeredgecolorname,'-',linewidth,markeredgecolorname, markerFaceAlpha, markerEdgeAlpha, false);
                  case 'x',scatterpatch2svg(fid,x'*ones(1,5)+ones(length(linex),1)*[-0.7 0.7 NaN -0.7 0.7]*0.8*markersize,y'*ones(1,5)+ones(length(liney),1)*[0.7 -0.7 NaN -0.7 0.7]*0.8*markersize,markeredgecolorname,'-',linewidth,markeredgecolorname, markerFaceAlpha, markerEdgeAlpha, false);
                  %% Octave keeps s, d, p and h in the HandleGraphics object, for the square, diamond, pentagram, and hexagram markers, respectively -- Jakob Malm
                  case {'square', 's'},scatterpatch2svg(fid,x'*ones(1,5)+ones(length(linex),1)*[-1 -1 1 1 -1]*0.75*markersize,y'*ones(1,5)+ones(length(liney),1)*[-1 1 1 -1 -1]*0.75*markersize,markerfacecolorname,'-',linewidth,markeredgecolorname, markerFaceAlpha, markerEdgeAlpha, true);
                  case {'diamond', 'd'},scatterpatch2svg(fid,x'*ones(1,5)+ones(length(linex),1)*[-0.7071 0 0.7071 0 -0.7071]*1.35*markersize,y'*ones(1,5)+ones(length(liney),1)*[0 1 0 -1 0]*1.35*markersize,markerfacecolorname,'-',linewidth,markeredgecolorname, markerFaceAlpha, markerEdgeAlpha, true);
                  case {'pentagram', 'p'},scatterpatch2svg(fid,...
                          x'*ones(1,11)+ones(length(linex),1)*[0 0.1180 0.5 0.1910 0.3090 0 -0.3090 -0.1910 -0.5 -0.1180 0]*2*markersize,...
                          y'*ones(1,11)+ones(length(liney),1)*[-0.5257 -0.1625 -0.1625 0.0621 0.4253 0.2008 0.4253 0.0621 -0.1625 -0.1625 -0.5257]*2*markersize,markerfacecolorname,'-',linewidth,markeredgecolorname, markerFaceAlpha, markerEdgeAlpha, true);
                  case {'hexagram', 'h'},scatterpatch2svg(fid,...
                          x'*ones(1,13)+ones(length(linex),1)*[0 0.2309 0.6928 0.4619 0.6928 0.2309 0 -0.2309 -0.6928 -0.4619 -0.6928 -0.2309 0]*1.3*markersize,...
                          y'*ones(1,13)+ones(length(liney),1)*[0.8 0.4 0.4 0 -0.4 -0.4 -0.8 -0.4 -0.4 0 0.4 0.4 0.8]*1.3*markersize,markerfacecolorname,'-',linewidth,markeredgecolorname, markerFaceAlpha, markerEdgeAlpha, true);
                  case '^',scatterpatch2svg(fid,x'*ones(1,4)+ones(length(linex),1)*[-1 1 0 -1]*1.15*markersize,y'*ones(1,4)+ones(length(liney),1)*[0.577 0.577 -1.155 0.577]*1.15*markersize,markerfacecolorname,'-',linewidth,markeredgecolorname, markerFaceAlpha, markerEdgeAlpha, true);
                  case 'v',scatterpatch2svg(fid,x'*ones(1,4)+ones(length(linex),1)*[-1 1 0 -1]*1.15*markersize,y'*ones(1,4)+ones(length(liney),1)*[-0.577 -0.577 1.155 -0.577]*1.15*markersize,markerfacecolorname,'-',linewidth,markeredgecolorname, markerFaceAlpha, markerEdgeAlpha, true);
                  case '<',scatterpatch2svg(fid,x'*ones(1,4)+ones(length(linex),1)*[0.577 0.577 -1.155 0.577]*1.15*markersize,y'*ones(1,4)+ones(length(liney),1)*[-1 1 0 -1]*1.15*markersize,markerfacecolorname,'-',linewidth,markeredgecolorname, markerFaceAlpha, markerEdgeAlpha, true);
                  case '>',scatterpatch2svg(fid,x'*ones(1,4)+ones(length(linex),1)*[-0.577 -0.577 1.155 -0.577]*1.15*markersize,y'*ones(1,4)+ones(length(liney),1)*[-1 1 0 -1]*1.15*markersize,markerfacecolorname,'-',linewidth,markeredgecolorname, markerFaceAlpha, markerEdgeAlpha, true);
              end
              % close the marker group
              fprintf(fid,'</g>\n');
          end
          animation2svg(fid, axchild(i));
          % close the line group
          fprintf(fid,'</g>\n');
      elseif strcmp(get(axchild(i),'Type'),'contour')
          clim = get(ax,'CLim');
          cmap = get(id,'Colormap');
          c = get(axchild(i),'ContourMatrix');
          linestyle = get(axchild(i),'LineStyle');
          linewidth = get(axchild(i),'LineWidth'); % linewidth = FIG2SVG_globals.resolutionScaling*get(axchild(i),'LineWidth');
          edge_opacity = 1.0;
          face_opacity = 1.0;
          index = 1;
          while index < size(c,2)
              patchIndices = (1:c(2,index))+index;
              % Close a patch if the coordinates do not contain NaNs
              x = c(1,patchIndices);
              y = c(2,patchIndices);
              if (x(1) == x(end)) && (y(1) == y(end))
                  closed = true;
              else
                  closed = false;
              end
              [x,y,~] = project(x,y,ones(1,c(2,index))*c(1,index),projection);
              x = (x*axpos(3)+axpos(1))*paperpos(3);
              y = (1-(y*axpos(4)+axpos(2)))*paperpos(4);
              pointc = c(1,index);
              pointc = round((pointc-clim(1))/(clim(2)-clim(1))*(size(cmap,1)-1)+1);
              % Limit index to smallest or biggest color index
              pointc = max(pointc,1);
              pointc = min(pointc,size(cmap,1));
              if ischar(get(axchild(i),'LineColor'))
                  if strcmp(get(axchild(i),'LineColor'),'none')
                      edgecolorname = 'none';
                  else
                      edgecolor = c(1,index);
                      if ~isnan(edgecolor)
                          if strcmp(get(axchild(i),'LineColor'),'flat')   % Bugfix 27.01.2008
                              edgecolorname = searchcolor(id,cmap(pointc,:));
                          else
                              edgecolorname = searchcolor(id,edgecolor);
                          end
                      else
                          edgecolorname = 'none';
                      end
                  end
              else
                  edgecolorname = searchcolor(id,get(axchild(i),'EdgeColor'));
              end
              if strcmp(get(axchild(i),'Fill'),'on')
                  facecolorname = searchcolor(id,cmap(pointc,:));
              end
              patch2svg(fid, x, y, facecolorname, linestyle, linewidth, edgecolorname, face_opacity, edge_opacity, closed)
              index = index+c(2,index)+1;
          end
      elseif strcmp(get(axchild(i),'Type'),'patch')
          fromLegend = 0;
          flat_shading = 1;
          cmap = get(id,'Colormap');
          pointc = get(axchild(i),'FaceVertexCData');
          if numel(unique(pointc)) == 1
            pointc = pointc(1);
          end
          if isempty(pointc)
            % Workaround for octave
            pointc = get(axchild(i),'CData');
          end
          % Scale color if scaled color mapping is turned on
          if strcmp(get(axchild(i),'CDataMapping'),'scaled')
              try
                clim = get(ax,'CLim');
                pointc = (pointc-clim(1))/(clim(2)-clim(1))*(size(cmap,1)-1)+1;
              catch
                fromLegend = 1; % legend patch
              end
          end
          % Limit index to smallest or biggest color index
          pointc = max(pointc,1);
          pointc = min(pointc,size(cmap,1));
          if ~ischar(get(axchild(i),'FaceAlpha'))
              face_opacity = get(axchild(i),'FaceAlpha');
          else
              face_opacity = 1.0;
          end
          if ~ischar(get(axchild(i),'EdgeAlpha'))
              edge_opacity = get(axchild(i),'EdgeAlpha');
          else
              edge_opacity = 1.0;
          end
          linestyle = get(axchild(i),'LineStyle');
          linewidth = get(axchild(i),'LineWidth'); % linewidth = FIG2SVG_globals.resolutionScaling*get(axchild(i),'LineWidth');
          marker = get(axchild(i),'Marker');
          markeredgecolor = get(axchild(i),'MarkerEdgeColor');
          markersize = 2/3*get(axchild(i),'MarkerSize');

          linex = get(axchild(i),'XData');
          linex = linex(:)';
          liney = get(axchild(i),'YData');
          liney = liney(:)';
          linez = get(axchild(i),'ZData');
          linez = linez(:)';
          try
            if strcmp(get(ax,'XScale'),'log')
                linex(linex <= 0) = NaN;
                linex = log10(linex);
            end
            if strcmp(get(ax,'YScale'),'log')
                liney(liney <= 0) = NaN;
                liney = log10(liney);
            end
            if isempty(linez)
              linez = zeros(size(linex));
            end
            if strcmp(get(ax,'ZScale'),'log')
              linez(linez <= 0) = NaN;
              linez = log10(linez);
            end
            [x,y,~] = project(linex,liney,linez,projection);
          catch
            % children of legend objects
            x = linex;
            y = liney;
            z = linez;
          end

          points = get(axchild(i),'Vertices')';
          try
            if ~fromLegend && strcmp(get(ax,'XScale'),'log')
              points(1,:) = log10(points(1,:));
            end
          catch
              fromLegend = 1;
          end
          if  ~fromLegend && strcmp(get(ax,'YScale'),'log')
              points(2,:) = log10(points(2,:));
          end
          if size(points,1) == 3
              if ~fromLegend && strcmp(get(ax,'ZScale'),'log')
                  points(3,:) = log10(points(3,:));
              end
          end
          if fromLegend
              x = points(1,:);
              y = points(2,:);
          elseif size(points,1) == 3
              [x,y,z] = project(points(1,:),points(2,:),points(3,:),projection);
          else
              [x,y,~] = project(points(1,:),points(2,:),zeros(size(points(1,:))),projection);
          end
          x = (x*axpos(3)+axpos(1))*paperpos(3);
          y = (1-(y*axpos(4)+axpos(2)))*paperpos(4);
          faces = get(axchild(i),'Faces');
          face_index = 1:size(faces,1);
          if size(points,1) == 3
              if any(isnan(faces(:)))
                  MyData = zeros(size(faces,1),1);
                  for ii = 1:size(faces,1)
                      PickMe = ~isnan(faces(ii,:));
                      MyData(ii) = sum(z(faces(ii,PickMe)),2);
                  end
                  [~,face_index] = sort(MyData);
                  faces = faces(face_index,:);
              else
                  [~,face_index] = sort(sum(z(faces(:,:)),2));
                  faces = faces(face_index,:);
              end
          end
%            if size(points,1) == 3
%                [~,face_index] = sort(sum(z(faces(:,:)),2));
%                faces = faces(face_index,:);
%            end
          markerOverlap = 0;
          if ~strcmp(linestyle, 'none')
              markerOverlap = max(markerOverlap, convertunit(linewidth*0.5, 'points', 'pixels', axpos(4)));
          end
          if ~strcmp(marker, 'none')
              markerOverlap = max(markerOverlap, convertunit(markersize, 'points', 'pixels', axpos(4)));
          end
          if FIG2SVG_globals.ClippingMode ~= 2
            boundingBoxElement = [min(x)-markerOverlap min(y)-markerOverlap max(x)-min(x)+2*markerOverlap max(y)-min(y)+2*markerOverlap];
          else
            boundingBoxElement = boundingBoxAxes;
          end
          [filterString, boundingBox] = filter2svg(fid, axchild(i), boundingBoxAxes, boundingBoxElement);
          if FIG2SVG_globals.ClippingMode == 1 && isfield(FIG2SVG_globals,'BoxOn') && ~FIG2SVG_globals.BoxOn
            boundingBoxAxes = [boundingBoxAxes(1) min([boundingBoxAxes(2) boundingBoxElement(2)]) max([boundingBoxAxes(3)+max([boundingBoxAxes(1)-boundingBoxElement(1),0]) boundingBoxElement(3)+max([boundingBoxElement(1)-boundingBoxAxes(1),0])]) [boundingBoxAxes(4)+max([boundingBoxAxes(2)-boundingBoxElement(2),0])]];
          elseif FIG2SVG_globals.ClippingMode == 3 || FIG2SVG_globals.ClippingMode == 1 && ~isfield(FIG2SVG_globals,'BoxOn')
            boundingBoxAxes = [min([boundingBoxAxes(1) boundingBoxElement(1)]) min([boundingBoxAxes(2) boundingBoxElement(2)]) max([boundingBoxAxes(3)+max([boundingBoxAxes(1)-boundingBoxElement(1),0]) boundingBoxElement(3)+max([boundingBoxElement(1)-boundingBoxAxes(1),0])]) max([boundingBoxAxes(4)+max([boundingBoxAxes(2)-boundingBoxElement(2),0]) boundingBoxElement(4)+max([boundingBoxElement(2)-boundingBoxAxes(2),0])])];
          end
          if strcmp(get(axchild(i),'Clipping'),'on') && FIG2SVG_globals.ClippingMode ~= 0
            clippingIdString = clipping2svg(fid, axchild(i), ax, paperpos, axpos, projection, axIdString);
            fprintf(fid,'<g id = "%s" clip-path = "url(#%s)" %s>\n', createId, clippingIdString, filterString);
          else
            fprintf(fid,'<g id = "%s" %s>\n', createId, filterString);
          end
          if ~isempty(filterString)
              % Workaround for Inkscape filter bug
              fprintf(fid,'<rect x = "%0.3f" y = "%0.3f" width = "%0.3f" height = "%0.3f" fill = "none" stroke = "none" />\n', boundingBox(1), boundingBox(2), boundingBox(3), boundingBox(4));
          end
          for p = 1:size(faces,1)
              if ischar(get(axchild(i),'FaceColor'))
                  if strcmp(get(axchild(i),'FaceColor'),'texturemap')
                      facecolorname = 'none';   % TO DO: texture map
                  elseif strcmp(get(axchild(i),'FaceColor'),'none')
                      facecolorname = 'none';
                  else
                      if size(pointc,1) == 1
                          facecolor = pointc;
                      elseif size(pointc,1) == size(faces,1)
                          if strcmp(get(axchild(i),'FaceColor'),'flat')
                              facecolor = pointc(face_index(p),:);
                          else
                              facecolor = pointc(face_index(p),:);
                              cdata = pointc(face_index(p),:);     % TO DO: color interpolation
                              flat_shading = 0;
                          end
                      elseif size(pointc,1) == size(points,2)
                          if strcmp(get(axchild(i),'FaceColor'),'flat')
                              facecolor = pointc(faces(p,1),:);
                          else
                              facecolor = pointc(faces(p,1));
                              cdata = pointc(faces(p,:),:);
                              flat_shading = 0;
                          end
                      else
                          error('Unsupported color handling for patches.');
                      end
                      if ~isnan(facecolor)
                          if size(facecolor,2) == 1
                              facecolorname = ['#' colorname(ceil(facecolor),:)];
                          else
                              if strcmp(get(axchild(i),'FaceColor'),'flat')  % Bugfix 27.01.2008
                                  facecolorname = searchcolor(id,facecolor/64);
                              else
                                  facecolorname = searchcolor(id,facecolor);
                              end
                          end
                      else
                          facecolorname = 'none';
                      end
                  end
              else
                  facecolorname = searchcolor(id,get(axchild(i),'FaceColor'));
              end
              if ischar(get(axchild(i),'EdgeColor'))
                  if strcmp(get(axchild(i),'EdgeColor'),'none')
                      edgecolorname = 'none';
                  else
                      if size(pointc,1) == 1
                          edgecolor = pointc;
                      elseif size(pointc,1) == size(faces,1)
                          edgecolor = pointc(p,:);
                      elseif size(pointc,1) == size(points,2)
                          if strcmp(get(axchild(i),'EdgeColor'),'flat')
                              % edgecolor = pointc(faces(p,1));
                             edgecolor = pointc;
                          else
                              edgecolor = pointc(faces(p,1));     % TO DO: color interpolation
                          end
                      else
                          error('Unsupported color handling for patches.');
                      end
                      if ~isnan(edgecolor)
                          if size(edgecolor,2) == 1
                              edgecolorname = ['#' colorname(ceil(edgecolor),:)];
                          else
                              if strcmp(get(axchild(i),'EdgeColor'),'flat')   % Bugfix 27.01.2008
                                edgecolorname = {};
                                for ii = 1:size(edgecolor,1)
                                  edgecolorname{ii} = searchcolor(id,edgecolor(ii,:)/64);
                                end
                              else
                                  edgecolorname = searchcolor(id,edgecolor);
                              end
                          end
                      else
                          edgecolorname = 'none';
                      end
                  end
              else
                  edgecolorname = searchcolor(id,get(axchild(i),'EdgeColor'));
              end
              % Close a patch if the coordinates do not contain NaNs
              if isnan(x(end)) || isnan(y(end))
                closed = false;
                facecolorname = 'none';
              elseif any(isnan(x)) || any(isnan(y))
                  closed = false;
              else
                  closed = true;
              end
              CurrNodes = faces(p,:);
              CurrNodes(isnan(CurrNodes)) = [];
              if flat_shading
                  patch2svg(fid, x(CurrNodes), y(CurrNodes), facecolorname, linestyle, linewidth, edgecolorname, face_opacity, edge_opacity, closed)
              else
                  gouraud_patch2svg(fid, x(CurrNodes), y(CurrNodes), cdata, linestyle, linewidth, edgecolorname, face_opacity, edge_opacity, id)
              end
%                if flat_shading
%                    patch2svg(fid, x(faces(p,:)), y(faces(p,:)), facecolorname, linestyle, linewidth, edgecolorname, face_opacity, edge_opacity, closed)
%                else
%                    gouraud_patch2svg(fid, x(faces(p,:)), y(faces(p,:)), cdata, linestyle, linewidth, edgecolorname, face_opacity, edge_opacity, id)
%                end
              if ~strcmp(marker, 'none')
                  for q = 1:size(faces,2)
                      xmarker = x(faces(p,q));
                      ymarker = y(faces(p,q));
                      % put the markers into a subgroup of the lines
                      fprintf(fid,'<g>\n');
                      if ischar(markeredgecolor)
                          switch markeredgecolor
                              case 'none',markeredgecolorname = 'none';
                              otherwise
                                  % if markeredgecolorname is 'auto' or something
                                  % else set the markeredgecolorname to the line color
                                  markeredgecolorname = selectColor(axchild(i), id, p, q, points, pointc, ...
                                      colorname, faces, 'MarkerEdgeColor');
                          end
                      else
                          markeredgecolorname = searchcolor(id,markeredgecolor);
                      end
                      markerfacecolor = get(axchild(i),'MarkerFaceColor');
                      if ischar(markerfacecolor)
                          switch markerfacecolor
                              case 'none',markerfacecolorname = 'none';
                              otherwise
                                  markerfacecolorname = selectColor(axchild(i), id, p, q, points, pointc, ...
                                      colorname, faces,'MarkerFaceColor');
                          end
                      else
                          markerfacecolorname = searchcolor(id,markerfacecolor);
                      end

                      try % not currently implemented, but maybe it will in the future
                        markerFaceAlpha = get(axchild(i),'MarkerFaceAlpha');
                        markerEdgeAlpha = get(axchild(i),'MarkerEdgeAlpha');
                      catch
                        markerFaceAlpha = 1;
                        markerEdgeAlpha = 1;
                      end

                      switch marker
                          case 'none'
                          case '.';circle2svg(fid,xmarker,ymarker,markersize/3,'none',markeredgecolorname,linewidth, 1, markerEdgeAlpha);
                          case 'o',circle2svg(fid,xmarker,ymarker,markersize,markeredgecolorname,markerfacecolorname,linewidth, markerFaceAlpha, markerEdgeAlpha);
                          case '+',patch2svg(fid,xmarker'*ones(1,5)+ones(length(linex),1)*[-1 1 NaN 0 0]*0.85*markersize,ymarker'*ones(1,5)+ones(length(liney),1)*[0 0 NaN -1 1]*0.85*markersize,markeredgecolorname,'-',linewidth,markeredgecolorname, markerFaceAlpha, markerEdgeAlpha, false);
                          case '*',patch2svg(fid,xmarker'*ones(1,11)+ones(length(linex),1)*[-1 1 NaN 0 0 NaN -0.7 0.7 NaN -0.7 0.7]*0.85*markersize,ymarker'*ones(1,11)+ones(length(liney),1)*[0 0 NaN -1 1 NaN 0.7 -0.7 NaN -0.7 0.7]*0.85*markersize,markeredgecolorname,'-',linewidth,markeredgecolorname, markerFaceAlpha, markerEdgeAlpha, false);
                          case 'x',patch2svg(fid,xmarker'*ones(1,5)+ones(length(linex),1)*[-0.7 0.7 NaN -0.7 0.7]*0.8*markersize,ymarker'*ones(1,5)+ones(length(liney),1)*[0.7 -0.7 NaN -0.7 0.7]*0.8*markersize,markeredgecolorname,'-',linewidth,markeredgecolorname, markerFaceAlpha, markerEdgeAlpha, false);
                          %% Octave keeps s, d, p and h in the HandleGraphics object, for the square, diamond, pentagram, and hexagram markers, respectively -- Jakob Malm
                          case {'square', 's'},patch2svg(fid,xmarker'*ones(1,5)+ones(length(linex),1)*[-1 -1 1 1 -1]*0.75*markersize,ymarker'*ones(1,5)+ones(length(liney),1)*[-1 1 1 -1 -1]*0.75*markersize,markerfacecolorname,'-',linewidth,markeredgecolorname, markerFaceAlpha, markerEdgeAlpha, true);
                          case {'diamond', 'd'},patch2svg(fid,xmarker'*ones(1,5)+ones(length(linex),1)*[-0.7071 0 0.7071 0 -0.7071]*1.35*markersize,ymarker'*ones(1,5)+ones(length(liney),1)*[0 1 0 -1 0]*1.35*markersize,markerfacecolorname,'-',linewidth,markeredgecolorname, markerFaceAlpha, markerEdgeAlpha, true);
                          case {'pentagram', 'p'},patch2svg(fid,...
                                  xmarker'*ones(1,11)+ones(length(linex),1)*[0 0.1180 0.5 0.1910 0.3090 0 -0.3090 -0.1910 -0.5 -0.1180 0]*2*markersize,...
                                  ymarker'*ones(1,11)+ones(length(liney),1)*[-0.5257 -0.1625 -0.1625 0.0621 0.4253 0.2008 0.4253 0.0621 -0.1625 -0.1625 -0.5257]*2*markersize,markerfacecolorname,'-',linewidth,markeredgecolorname, markerFaceAlpha, markerEdgeAlpha, true);
                          case {'hexagram', 'h'},patch2svg(fid,...
                                  xmarker'*ones(1,13)+ones(length(linex),1)*[0 0.2309 0.6928 0.4619 0.6928 0.2309 0 -0.2309 -0.6928 -0.4619 -0.6928 -0.2309 0]*1.3*markersize,...
                                  ymarker'*ones(1,13)+ones(length(liney),1)*[0.8 0.4 0.4 0 -0.4 -0.4 -0.8 -0.4 -0.4 0 0.4 0.4 0.8]*1.3*markersize,markerfacecolorname,'-',linewidth,markeredgecolorname, markerFaceAlpha, markerEdgeAlpha, true);
                          case '^',patch2svg(fid,xmarker'*ones(1,4)+ones(length(linex),1)*[-1 1 0 -1]*1.15*markersize,ymarker'*ones(1,4)+ones(length(liney),1)*[0.577 0.577 -1.155 0.577]*1.15*markersize,markerfacecolorname,'-',linewidth,markeredgecolorname, markerFaceAlpha, markerEdgeAlpha, true);
                          case 'v',patch2svg(fid,xmarker'*ones(1,4)+ones(length(linex),1)*[-1 1 0 -1]*1.15*markersize,ymarker'*ones(1,4)+ones(length(liney),1)*[-0.577 -0.577 1.155 -0.577]*1.15*markersize,markerfacecolorname,'-',linewidth,markeredgecolorname, markerFaceAlpha, markerEdgeAlpha, true);
                          case '<',patch2svg(fid,xmarker'*ones(1,4)+ones(length(linex),1)*[0.577 0.577 -1.155 0.577]*1.15*markersize,ymarker'*ones(1,4)+ones(length(liney),1)*[-1 1 0 -1]*1.15*markersize,markerfacecolorname,'-',linewidth,markeredgecolorname, markerFaceAlpha, markerEdgeAlpha, true);
                          case '>',patch2svg(fid,xmarker'*ones(1,4)+ones(length(linex),1)*[-0.577 -0.577 1.155 -0.577]*1.15*markersize,ymarker'*ones(1,4)+ones(length(liney),1)*[-1 1 0 -1]*1.15*markersize,markerfacecolorname,'-',linewidth,markeredgecolorname, markerFaceAlpha, markerEdgeAlpha, true);
                      end
                      % close the marker group
                      fprintf(fid,'</g>\n');
                  end
              end
          end
          fprintf(fid,'</g>\n');
      elseif strcmp(get(axchild(i),'Type'),'surface')
          flat_shading = 1;
          cmap = get(id,'Colormap');
          [faces,points,pointc,alpha] = surface2patch(axchild(i));
          points = points';
          % Scale color if scaled color mapping is turned on
          if strcmp(get(axchild(i),'CDataMapping'),'scaled')
              clim = get(ax,'CLim');
              pointc = (pointc-clim(1))/(clim(2)-clim(1))*(size(cmap,1)-1)+1;
          end
          % Limit index to smallest or biggest color index
          pointc = max(pointc,1);
          if ~ischar(get(axchild(i),'FaceAlpha'))
              face_opacity = get(axchild(i),'FaceAlpha');
          elseif strcmp(get(axchild(i),'FaceAlpha'),'flat')
              face_opacity = alpha;
              switch get(axchild(i),'AlphaDataMapping')
                  case {'direct'}
                      face_opacity = 1.0; % TODO
                  case {'scaled'}
                      alim = get(ax,'ALim');
                      face_opacity = (face_opacity-alim(1))/(alim(2)-alim(1));
                  case {'none'}
                      % Clip alpha data
                      face_opacity = min(1, face_opacity);
                      face_opacity = max(0, face_opacity);
                  otherwise
                      error(['Unsupported AlphaDataMapping identifier ''' get(axchild(i),'AlphaDataMapping') '''.']);
              end
          else
              face_opacity = 1.0;
          end
          if ~ischar(get(axchild(i),'EdgeAlpha'))
              edge_opacity = get(axchild(i),'EdgeAlpha');
          else
              edge_opacity = 1.0;
          end
          pointc = min(pointc,size(cmap,1));
          linestyle = get(axchild(i),'LineStyle');
          linewidth = get(axchild(i),'LineWidth'); % linewidth = FIG2SVG_globals.resolutionScaling*get(axchild(i),'LineWidth');
          if strcmp(get(ax,'XScale'),'log')
              points(1,:) = log10(points(1,:));
          end
          if strcmp(get(ax,'YScale'),'log')
              points(2,:) = log10(points(2,:));
          end
          if size(points,1) == 3
              if strcmp(get(ax,'ZScale'),'log')
                  points(3,:) = log10(points(3,:));
              end
          end
          if size(points,1) == 3
              [x,y,z] = project(points(1,:),points(2,:),points(3,:),projection);
          else
              [x,y,~] = project(points(1,:),points(2,:),zeros(size(points(1,:))),projection);
          end
          x = (x*axpos(3)+axpos(1))*paperpos(3);
          y = (1-(y*axpos(4)+axpos(2)))*paperpos(4);
          face_index = 1:size(faces,1);
          if size(points,1) == 3
              [~,face_index] = sort(sum(z(faces(:,:)),2));
              faces = faces(face_index,:);
          end
          markerOverlap = 0;
          if ~strcmp(linestyle, 'none')
              markerOverlap = max(markerOverlap, convertunit(linewidth*0.5, 'points', 'pixels', axpos(4)));
          end
          if FIG2SVG_globals.ClippingMode ~= 2
            boundingBoxElement = [min(x)-markerOverlap min(y)-markerOverlap max(x)-min(x)+2*markerOverlap max(y)-min(y)+2*markerOverlap];
          else
            boundingBoxElement = boundingBoxAxes;
          end
          [filterString, boundingBox] = filter2svg(fid, axchild(i), boundingBoxAxes, boundingBoxElement);
          if FIG2SVG_globals.ClippingMode == 1 && isfield(FIG2SVG_globals,'BoxOn') && ~FIG2SVG_globals.BoxOn
            boundingBoxAxes = [boundingBoxAxes(1) min([boundingBoxAxes(2) boundingBoxElement(2)]) max([boundingBoxAxes(3)+max([boundingBoxAxes(1)-boundingBoxElement(1),0]) boundingBoxElement(3)+max([boundingBoxElement(1)-boundingBoxAxes(1),0])]) [boundingBoxAxes(4)+max([boundingBoxAxes(2)-boundingBoxElement(2),0])]];
          elseif FIG2SVG_globals.ClippingMode == 3 || FIG2SVG_globals.ClippingMode == 1 && ~isfield(FIG2SVG_globals,'BoxOn')
            boundingBoxAxes = [min([boundingBoxAxes(1) boundingBoxElement(1)]) min([boundingBoxAxes(2) boundingBoxElement(2)]) max([boundingBoxAxes(3)+max([boundingBoxAxes(1)-boundingBoxElement(1),0]) boundingBoxElement(3)+max([boundingBoxElement(1)-boundingBoxAxes(1),0])]) max([boundingBoxAxes(4)+max([boundingBoxAxes(2)-boundingBoxElement(2),0]) boundingBoxElement(4)+max([boundingBoxElement(2)-boundingBoxAxes(2),0])])];
          end
          if strcmp(get(axchild(i),'Clipping'),'on') && FIG2SVG_globals.ClippingMode ~= 0
            clippingIdString = clipping2svg(fid, axchild(i), ax, paperpos, axpos, projection, axIdString);
            fprintf(fid,'<g id = "%s" clip-path = "url(#%s)" %s>\n', createId, clippingIdString, filterString);
          else
            fprintf(fid,'<g id = "%s" %s>\n', createId, filterString);
          end
          if ~isempty(filterString)
              % Workaround for Inkscape filter bug
              fprintf(fid,'<rect x = "%0.3f" y = "%0.3f" width = "%0.3f" height = "%0.3f" fill = "none" stroke = "none" />\n', boundingBox(1), boundingBox(2), boundingBox(3), boundingBox(4));
          end
          for p = 1:size(faces,1)
              if ischar(get(axchild(i),'FaceColor'))
                  if strcmp(get(axchild(i),'FaceColor'),'texturemap')
                      facecolorname = 'none';   % TO DO: texture map
                  elseif strcmp(get(axchild(i),'FaceColor'),'none')
                      facecolorname = 'none';
                  else
                      if size(pointc,1) == 1
                          facecolor = pointc;
                      elseif size(pointc,1) == size(faces,1)
                          facecolor = pointc(face_index(p),:);
                      elseif size(pointc,1) == size(points,2)
                          if strcmp(get(axchild(i),'FaceColor'),'flat')
                              facecolor = pointc(faces(p,1));
                          else
                              facecolor = pointc(faces(p,1));
                              cdata = pointc(faces(p,:));
                              flat_shading = 0;
                          end
                      else
                          error('Unsupported color handling for patches.');
                      end
                      if ~isnan(facecolor)
                          if size(facecolor,2) == 1
                              facecolorname = ['#' colorname(ceil(facecolor),:)];
                          else
                              facecolorname = searchcolor(id,facecolor);
                          end
                      else
                          facecolorname = 'none';
                      end
                  end
              else
                  facecolorname = searchcolor(id,get(axchild(i),'FaceColor'));
              end
              if size(face_opacity,1) == 1
                  face_opacity_value = face_opacity;
              elseif size(face_opacity,1) == size(faces,1)
                  face_opacity_value = face_opacity(p,:);
              elseif size(face_opacity,1) == size(points,2)
                  face_opacity_value = face_opacity(faces(p,1));
              else
                  error('Unsupported face alpha value handling for patches.');
              end
              if ischar(get(axchild(i),'EdgeColor'))
                  if strcmp(get(axchild(i),'EdgeColor'),'none')
                      edgecolorname = 'none';
                  else
                      if size(pointc,1) == 1
                          edgecolor = pointc;
                      elseif size(pointc,1) == size(faces,1)
                          edgecolor = pointc(p,:);
                      elseif size(pointc,1) == size(points,2)
                          if strcmp(get(axchild(i),'EdgeColor'),'flat')
                              edgecolor = pointc(faces(p,1));
                          else
                              edgecolor = pointc(faces(p,1));     % TO DO: color interpolation
                          end
                      else
                          error('Unsupported color handling for patches.');
                      end
                      if ~isnan(edgecolor)
                          if size(edgecolor,2) == 1
                              edgecolorname = ['#' colorname(ceil(edgecolor),:)];
                          else
                              edgecolorname = searchcolor(id,edgecolor);
                          end
                      else
                          edgecolorname = 'none';
                      end
                  end
              else
                  edgecolorname = searchcolor(id,get(axchild(i),'EdgeColor'));
              end
              if flat_shading
                  patch2svg(fid, x(faces(p,:)), y(faces(p,:)), facecolorname, linestyle, linewidth, edgecolorname,  face_opacity_value, edge_opacity, false)
              else
                  gouraud_patch2svg(fid, x(faces(p,:)), y(faces(p,:)), cdata, linestyle, linewidth, edgecolorname, face_opacity_value, edge_opacity, id)
              end
          end
          fprintf(fid,'</g>\n');
      elseif strcmp(get(axchild(i),'Type'),'rectangle')
          scolorname = searchcolor(id,get(axchild(i),'EdgeColor'));
          fcolorname = searchcolor(id,get(axchild(i),'FaceColor'));
          face_opacity = get(axchild(i),'FaceAlpha');
          linewidth = get(axchild(i),'LineWidth'); % linewidth = FIG2SVG_globals.resolutionScaling*get(axchild(i),'LineWidth');
          position = get(axchild(i),'Position');
          posx = [position(1) position(1)+position(3)];
          if strcmp(get(ax,'XScale'),'log')
              posx(posx <= 0) = NaN;
              posx = log10(posx);
          end
          posy = [position(2) position(2)+position(4)];
          if strcmp(get(ax,'YScale'),'log')
              posy(posy <= 0) = NaN;
              posy = log10(posy);
          end
          posz = [0 0];
          linestyle = get(axchild(i),'LineStyle');
          if strcmp(linestyle,'none')
              scolorname = 'none';
          end
          pattern = lineStyle2svg(linestyle, linewidth);
          [x,y,~] = project(posx,posy,posz,projection);
          x = (x*axpos(3)+axpos(1))*paperpos(3);
          y = (1-(y*axpos(4)+axpos(2)))*paperpos(4);
          rect = [min(x) min(y) max(x)-min(x) max(y)-min(y)];
          curvature = get(axchild(i),'Curvature');
          curvature(1) = curvature(1)*rect(3)*0.5;
          curvature(2) = curvature(2)*rect(4)*0.5;
          markerOverlap = 0;
          if ~strcmp(linestyle, 'none')
              markerOverlap = max(markerOverlap, convertunit(linewidth*0.5, 'points', 'pixels', axpos(4)));
          end
          % put a rectangle into a group with its markers
          if FIG2SVG_globals.ClippingMode ~= 2
            boundingBoxElement = rect + [-markerOverlap -markerOverlap 2*markerOverlap 2*markerOverlap];
          else
            boundingBoxElement = boundingBoxAxes;
          end
          [filterString, boundingBox] = filter2svg(fid, axchild(i), boundingBoxAxes, boundingBoxElement);
          if isfield(FIG2SVG_globals,'BoxOn') && ~FIG2SVG_globals.BoxOn
            boundingBoxAxes = [boundingBoxAxes(1) min([boundingBoxAxes(2) boundingBoxElement(2)]) max([boundingBoxAxes(3)+max([boundingBoxAxes(1)-boundingBoxElement(1),0]) boundingBoxElement(3)+max([boundingBoxElement(1)-boundingBoxAxes(1),0])]) [boundingBoxAxes(4)+max([boundingBoxAxes(2)-boundingBoxElement(2),0])]];
          elseif FIG2SVG_globals.ClippingMode == 3 || FIG2SVG_globals.ClippingMode == 1 && ~isfield(FIG2SVG_globals,'BoxOn')
            boundingBoxAxes = [min([boundingBoxAxes(1) boundingBoxElement(1)]) min([boundingBoxAxes(2) boundingBoxElement(2)]) max([boundingBoxAxes(3)+max([boundingBoxAxes(1)-boundingBoxElement(1),0]) boundingBoxElement(3)+max([boundingBoxElement(1)-boundingBoxAxes(1),0])]) max([boundingBoxAxes(4)+max([boundingBoxAxes(2)-boundingBoxElement(2),0]) boundingBoxElement(4)+max([boundingBoxElement(2)-boundingBoxAxes(2),0])])];
          end
          if strcmp(get(axchild(i),'Clipping'),'on') && FIG2SVG_globals.ClippingMode ~= 0
            clippingIdString = clipping2svg(fid, axchild(i), ax, paperpos, axpos, projection, axIdString);
            fprintf(fid,'<g id = "%s" clip-path = "url(#%s)" %s>\n', createId, clippingIdString, filterString);
          else
            fprintf(fid,'<g id = "%s" %s>\n', createId, filterString);
          end
          if ~isempty(filterString)
              % Workaround for Inkscape filter bug
              fprintf(fid,'<rect x = "%0.3f" y = "%0.3f" width = "%0.3f" height = "%0.3f" fill = "none" stroke = "none" />\n', boundingBox(1), boundingBox(2), boundingBox(3), boundingBox(4));
          end
          fprintf(fid,'<rect x = "%0.3f" y = "%0.3f" width = "%0.3f" height = "%0.3f" rx = "%0.3f" ry = "%0.3f" fill = "%s" fill-opacity = "%0.3f" stroke-linecap = "square" stroke-linejoin = "miter" stroke = "%s" stroke-width = "%0.3fpt" %s />\n',...
              rect(1), rect(2), rect(3), rect(4), curvature(1), curvature(2), fcolorname, face_opacity, scolorname, linewidth, pattern);
          % close the rectangle group
          fprintf(fid,'</g>\n');
      elseif strcmp(get(axchild(i),'Type'),'text')
          if FIG2SVG_globals.octave
            extent = [0 0 0 0];
          else
            extent = get(axchild(i),'Extent');
          end
          margin = get(axchild(i),'Margin');
          facecolor = get(axchild(i),'BackgroundColor');
          edgecolor = get(axchild(i),'EdgeColor');
          linewidth = get(axchild(i),'LineWidth'); % linewidth = FIG2SVG_globals.resolutionScaling*get(axchild(i),'LineWidth');
          linestyle = get(axchild(i),'LineStyle');
          if ischar(facecolor)
              if ~strcmp(facecolor,'none')
                  error('Illegal face color for text.');
              else
                  facecolorname = 'none';
              end
          else
              facecolorname = searchcolor(id, facecolor);
          end
          if ischar(edgecolor)
              if ~strcmp(edgecolor,'none')
                  error('Illegal edge color for text.');
              else
                  edgecolorname = 'none';
              end
          else
              edgecolorname = searchcolor(id, edgecolor);
          end
          extentx = [extent(1) extent(1)+extent(3)];
          extenty = [extent(2) extent(2)+extent(4)];
          extentz = [0 0];
          try
            [x,y,~] = project(extentx,extenty,extentz,projection);
          catch
            x = extentx;
            y = extenty;
          end
          x = (x*axpos(3)+axpos(1))*paperpos(3);
          y = (1-(y*axpos(4)+axpos(2)))*paperpos(4);
          box = [min(x)-margin min(y)-margin max(x)-min(x)+2*margin max(y)-min(y)+2*margin];
          markerOverlap = 0;
          if ~strcmp(linestyle, 'none')
              markerOverlap = max(markerOverlap, convertunit(linewidth*0.5, 'points', 'pixels', axpos(4)));
          end
          if FIG2SVG_globals.ClippingMode ~= 2
            boundingBoxElement = [min(x)-markerOverlap min(y)-markerOverlap max(x)-min(x)+2*markerOverlap max(y)-min(y)+2*markerOverlap];
          else
            boundingBoxElement = boundingBoxAxes;
          end
          [filterString, boundingBox] = filter2svg(fid, axchild(i), boundingBoxAxes, boundingBoxElement);
          if FIG2SVG_globals.ClippingMode == 1 && isfield(FIG2SVG_globals,'BoxOn') && ~FIG2SVG_globals.BoxOn
            boundingBoxAxes = [boundingBoxAxes(1) min([boundingBoxAxes(2) boundingBoxElement(2)]) max([boundingBoxAxes(3)+max([boundingBoxAxes(1)-boundingBoxElement(1),0]) boundingBoxElement(3)+max([boundingBoxElement(1)-boundingBoxAxes(1),0])]) [boundingBoxAxes(4)+max([boundingBoxAxes(2)-boundingBoxElement(2),0])]];
          elseif FIG2SVG_globals.ClippingMode == 3 || FIG2SVG_globals.ClippingMode == 1 && ~isfield(FIG2SVG_globals,'BoxOn')
            boundingBoxAxes = [min([boundingBoxAxes(1) boundingBoxElement(1)]) min([boundingBoxAxes(2) boundingBoxElement(2)]) max([boundingBoxAxes(3)+max([boundingBoxAxes(1)-boundingBoxElement(1),0]) boundingBoxElement(3)+max([boundingBoxElement(1)-boundingBoxAxes(1),0])]) max([boundingBoxAxes(4)+max([boundingBoxAxes(2)-boundingBoxElement(2),0]) boundingBoxElement(4)+max([boundingBoxElement(2)-boundingBoxAxes(2),0])])];
          end
          if strcmp(get(axchild(i),'Clipping'),'on') && FIG2SVG_globals.ClippingMode ~= 0
            clippingIdString = clipping2svg(fid, axchild(i), ax, paperpos, axpos, projection, axIdString);
            fprintf(fid,'<g id = "%s" clip-path = "url(#%s)" %s>\n', createId, clippingIdString, filterString);
          else
            fprintf(fid,'<g id = "%s" %s>\n', createId, filterString);
          end
          if ~isempty(filterString)
              % Workaround for Inkscape filter bug
              fprintf(fid,'<rect x = "%0.3f" y = "%0.3f" width = "%0.3f" height = "%0.3f" fill = "none" stroke = "none" />\n', boundingBox(1), boundingBox(2), boundingBox(3), boundingBox(4));
          end
          if ~strcmp(edgecolorname, 'none') || ~strcmp(facecolorname, 'none')
              pattern = lineStyle2svg(linestyle, linewidth);
              fprintf(fid,'<rect x = "%0.3f" y = "%0.3f" width = "%0.3f" height = "%0.3f" fill = "%s" stroke-linecap = "square" stroke-linejoin = "miter" stroke = "%s" stroke-width = "%0.3fpt" %s />\n', ...
                  box(1), box(2), box(3), box(4), facecolorname, edgecolorname, linewidth, pattern);
          end
          text2svg(fid,axpos,paperpos,axchild(i),ax,projection)
          fprintf(fid,'</g>\n');
      elseif strcmp(get(axchild(i),'Type'),'image')
          cmap = get(id,'Colormap');
          pointx = get(axchild(i),'XData');
          pointy = get(axchild(i),'YData');
          % If the XData is a vector we only use start and stop for the image
          if (size(pointx, 1) > 2) || (size(pointx, 2) > 1)
              pointx = [pointx(1) pointx(end)];
          end
          if (size(pointy, 1) > 2) || (size(pointy, 2) > 1)
              pointy = [pointy(1) pointy(end)];
          end
          if (size(pointx, 1) > 1) && (size(pointy, 1) > 1)
              [x,y,~] = project(pointx,zeros(size(pointx)),zeros(size(pointx)),projection);
          else
              [x,~,~] = project(pointx,zeros(size(pointx)),zeros(size(pointx)),projection);
              [~,y,~] = project(zeros(size(pointy)),pointy,zeros(size(pointy)),projection);
          end
          pointc = get(axchild(i),'CData');
          %pointcclass = class(pointc);  % Bugfix proposed by Tom
          if strcmp(get(axchild(i),'CDataMapping'),'scaled')
              clim = get(ax,'CLim');
              pointc = (pointc-clim(1))/(clim(2)-clim(1))*(size(cmap,1) - 1) + 1; % Bugfix proposed by Tom
              %pointcclass = 'double'; % since range is now [0->size(cmap,1)-1]  % Bugfix proposed by Tom
          end
          data_aspect_ratio = get(ax,'DataAspectRatio');
          if length(x) == 2
              if size(pointc, 2) == 1
                  halfwidthx = abs(x(2) - x(1)) * data_aspect_ratio(1);
              else
                  halfwidthx = abs(x(2) - x(1))/(size(pointc,2) - 1);
              end
          else
              halfwidthx = data_aspect_ratio(1);
          end
          if length(y) == 2
              if size(pointc, 1) == 1
                  halfwidthy = abs(y(2) - y(1)) * data_aspect_ratio(2);
              else
                  halfwidthy = abs(y(2) - y(1))/(size(pointc,1) - 1);
              end
          else
              halfwidthy = data_aspect_ratio(2);
          end
          if length(pointx) > 1
              if xor(strcmp(get(ax,'XDir'),'reverse'), pointx(1) > pointx(2))
                  if ndims(pointc) < 3
                      pointc = fliplr(pointc);
                  elseif ndims(pointc) == 3
                      for j = 1:size(pointc,3)
                          pointc(:,:,j) = fliplr(pointc(:,:,j));
                      end
                  else
                      error('Invalid number of dimensions of data.');
                  end
              end
          end
          if length(pointy) > 1
              if xor(strcmp(get(ax,'YDir'),'reverse'), pointy(1) > pointy(2))
                  if ndims(pointc) < 3
                      pointc = flipud(pointc);
                  elseif ndims(pointc) == 3
                      for j = 1:size(pointc,3)
                          pointc(:,:,j) = flipud(pointc(:,:,j));
                      end
                  else
                      error('Invalid number of dimensions of data.');
                  end
              end
          end
          % pointc = cast(pointc,pointcclass);  % Bugfix proposed by Tom
          % Function 'cast' is not supported by old Matlab versions
          if (~isa(pointc, 'double') && ~isa(pointc, 'single'))
              if strcmp(get(axchild(i),'CDataMapping'),'scaled')
                  pointc = double(pointc);
              else
                  pointc = double(pointc) + 1;
              end
          end
          if ndims(pointc) ~= 3
              pointc = max(min(round(double(pointc)),size(cmap,1)),1);
          end
          % CameraUpVector = get(ax,'CameraUpVector');
          filename = [FIG2SVG_globals.basefilename sprintf('%03d',FIG2SVG_globals.figurenumber) '.' FIG2SVG_globals.pixelFileType];
          FIG2SVG_globals.figurenumber = FIG2SVG_globals.figurenumber + 1;
          if isempty(FIG2SVG_globals.basefilepath)
              current_path = pwd;
          else
              current_path = FIG2SVG_globals.basefilepath;
          end
          if exist(fullfile(current_path,filename),'file')
              lastwarn('');
              delete(filename);
              if strcmp(lastwarn,'File not found or permission denied.')
                  error('Cannot write image file. Make sure that no image is opened in an other program.')
              end
          end
          if ndims(pointc) < 3
              pointc = flipud(pointc);
          elseif ndims(pointc) == 3
              for j = 1:size(pointc,3)
                  pointc(:,:,j) = flipud(pointc(:,:,j));
              end
          else
              error('Invalid number of dimensions of data.');
          end
          if ndims(pointc) == 3
              % pointc is not indexed
              imwrite(pointc,fullfile(FIG2SVG_globals.basefilepath,filename),FIG2SVG_globals.pixelFileType);
          else
              % pointc is probably indexed
              if FIG2SVG_globals.octave
                pointc = max(2, pointc);
              end
              imwrite(pointc,cmap,fullfile(FIG2SVG_globals.basefilepath,filename),FIG2SVG_globals.pixelFileType);
          end
          lx = (size(pointc,2)*halfwidthx)*axpos(3)*paperpos(3);
          ly = (size(pointc,1)*halfwidthy)*axpos(4)*paperpos(4);
          if strcmp(get(ax,'DataAspectRatioMode'),'manual')
              pointsx = ((min(x) - halfwidthx/2)*axpos(3)+axpos(1))*paperpos(3);
              pointsy = (1-((max(y) + halfwidthy/2)*axpos(4)+axpos(2)))*paperpos(4);
          else
              pointsx = axpos(1)*paperpos(3);
              pointsy = (1-(axpos(4)+axpos(2)))*paperpos(4);
          end
          [filterString, boundingBox] = filter2svg(fid, axchild(i), boundingBoxAxes, boundingBoxAxes);
          if strcmp(get(axchild(i),'Clipping'),'on') && FIG2SVG_globals.ClippingMode
              clippingIdString = clipping2svg(fid, axchild(i), ax, paperpos, axpos, projection, axIdString);
              fprintf(fid,'<g id = "%s" clip-path = "url(#%s)" %s>\n', createId, clippingIdString, filterString);
              if ~isempty(filterString)
                  % Workaround for Inkscape filter bug
                  fprintf(fid,'<rect x = "%0.3f" y = "%0.3f" width = "%0.3f" height = "%0.3f" fill = "none" stroke = "none" />\n', boundingBox(1), boundingBox(2), boundingBox(3), boundingBox(4));
              end
              fprintf(fid,'<image x = "%0.3f" y = "%0.3f" width = "%0.3f" height = "%0.3f" image-rendering = "optimizeSpeed" preserveAspectRatio = "none" xlink:href = "%s" />\n', pointsx, pointsy, lx, ly, filename); % With image-rendering = "optimizeQuality" the image appears interpolated, which might be nice, but image/imagesc don't work that way, images appear pixelated. To workaround this, use pcolor+shading, or previously interpolate with interp2
              fprintf(fid,'</g>\n');
          else
              fprintf(fid,'<g id = "%s" %s>\n', createId, filterString);
              if ~isempty(filterString)
                  % Workaround for Inkscape filter bug
                  fprintf(fid,'<rect x = "%0.3f" y = "%0.3f" width = "%0.3f" height = "%0.3f" fill = "none" stroke = "none" />\n', boundingBox(1), boundingBox(2), boundingBox(3), boundingBox(4));
              end
              fprintf(fid,'<image x = "%0.3f" y = "%0.3f" width = "%0.3f" height = "%0.3f" image-rendering = "optimizeSpeed" preserveAspectRatio = "none" xlink:href = "%s" />\n', pointsx, pointsy, lx, ly, filename); % With image-rendering = "optimizeQuality" the image appears interpolated, which might be nice, but image/imagesc don't work that way, images appear pixelated. To workaround this, use pcolor+shading, or previously interpolate with interp2
              fprintf(fid,'</g>\n');
          end
      elseif strcmp(get(axchild(i),'Type'),'hggroup') || strcmp(get(axchild(i),'Type'),'bar')
          % handle group types (like error bars in matlab < 2014b)
          % FIXME: they are not yet perfectly handled, there are more options
          % that are not used
          [filterString, boundingBox] = filter2svg(fid, axchild(i), boundingBoxAxes, boundingBoxAxes);
          if strcmp(get(axchild(i),'Clipping'),'on') && FIG2SVG_globals.ClippingMode
              clippingIdString = clipping2svg(fid, axchild(i), ax, paperpos, axpos, projection, axIdString);
              fprintf(fid,'<g id = "%s" clip-path = "url(#%s)" %s>\n', createId, clippingIdString, filterString);
          else
              fprintf(fid, '<g id = "%s" %s>', createId, filterString);
          end
          if ~isempty(filterString)
              % Workaround for Inkscape filter bug
              fprintf(fid,'<rect x = "%0.3f" y = "%0.3f" width = "%0.3f" height = "%0.3f" fill = "none" stroke = "none" />\n', boundingBox(1), boundingBox(2), boundingBox(3), boundingBox(4));
          end
          if strcmp(get(axchild(i),'Type'),'hggroup')
            boundingBoxAxes = axchild2svg(fid,id,axIdString,ax,paperpos,get(axchild(i), 'Children'),axpos,groupax,projection,boundingBoxAxes);
          elseif strcmp(get(axchild(i),'Type'),'bar')
            barFaceColor = get(axchild(i),'FaceColor');
            face_opacity = get(axchild(i),'FaceAlpha');
            barEdgeColor = get(axchild(i),'EdgeColor');
            edge_opacity = get(axchild(i),'EdgeAlpha');

            if strcmp(linestyle,'none') || strcmp(barEdgeColor,'none')
              scolorname = 'none';
              edge_opacity = 1;
            end
            if ~strcmp(barEdgeColor,'flat')
              scolorname = searchcolor(id,get(axchild(i),'EdgeColor'));
            end
            if strcmp(barFaceColor,'none')
              fcolorname = 'none';
              face_opacity = 1;
            end

            linestyle = get(axchild(i),'LineStyle');
            linewidth = get(axchild(i),'LineWidth');

            barColors = get(axchild(i),'CData');
            barCenters = get(axchild(i),'XData');
            if numel(barCenters) > 1
              barSep = min(diff(barCenters));
            else
              barSep = 1;
            end
            barHeight = get(axchild(i),'YData');
            barWidth = get(axchild(i),'BarWidth');
            showBaseline = get(axchild(i),'ShowBaseline');
            barBaseline = get(axchild(i),'BaseValue');
            barStyle = get(axchild(i),'BarLayout');
            userData = get(axchild(i),'UserData');

            currentBarSetNumber = currentBarSetNumber + 1;
            if numChildBars > 1
              if strcmp(barStyle,'grouped') && ~strcmpi(userData,'ungrouped') % grouped bars unless explicitly specified by UserData property
                [barCenters,barWidth] = groupBarProperties(currentBarSetNumber,barCenters,numChildBars,barWidth,barSep);
              elseif strcmp(barStyle,'stacked')
                if barBaseline ~= 0
                  warning('Ignoring Bar Baseline Value as it cannot be use with Stacked Layout')
                end
                stackedBarYprev = stackedBarYref;
                if numel(stackedBarYprev) == 1
                  stackedBarYprev = stackedBarYprev*ones(size(barHeight));
                end
                stackedBarYref = stackedBarYref + barHeight;
              end
            end

            % let's bar with no width appear
            if barWidth == 0
              barWidth = 0.0001;
            end

            %%% bar rectangles %%%
            for iBar = 1:numel(barCenters)
              if ~strcmp(barFaceColor,'none')
                fcolorname = searchcolor(id,barColors(iBar,:));
              end
              if strcmp(barEdgeColor,'flat')
                scolorname = searchcolor(id,barColors(iBar,:));
              end

              posx = barCenters(iBar) + barSep*barWidth*([-0.5,0.5]);
              if strcmp(get(ax,'XScale'),'log')
                posx(posx <= 0) = NaN;
                posx = log10(posx);
              end
              if strcmp(barStyle,'grouped')
                posy = [barBaseline barHeight(iBar)];
              elseif strcmp(barStyle,'stacked')
                posy = [stackedBarYprev(iBar) stackedBarYref(iBar)];
              end
              if strcmp(get(ax,'YScale'),'log')
                posy(posy <= 0) = NaN;
                posy = log10(posy);
              end
              posz = [0 0];
              pattern = lineStyle2svg(linestyle, linewidth);
              [x,y,~] = project(posx,posy,posz,projection);
              x = (x*axpos(3)+axpos(1))*paperpos(3);
              y = (1-(y*axpos(4)+axpos(2)))*paperpos(4);
              rect = [min(x) min(y) max(x)-min(x) max(y)-min(y)];
              curvature = [0 0];
              fprintf(fid,'<rect x = "%0.3f" y = "%0.3f" width = "%0.3f" height = "%0.3f" rx = "%0.3f" ry = "%0.3f" fill = "%s" fill-opacity = "%0.3f" stroke-linecap = "square" stroke-linejoin = "miter" stroke = "%s" stroke-opacity = "%0.3f" stroke-width = "%0.3fpt" %s />\n', rect(1), rect(2), rect(3), rect(4), curvature(1), curvature(2), fcolorname, face_opacity, scolorname, edge_opacity, linewidth, pattern);
            end

            %%% bar baseline %%%
            if (strcmp(showBaseline,'on') || barBaseline ~= 0) && ~strcmp(barStyle,'stacked')
              posx = get(ax,'xlim');
              posy = barBaseline*([1 1]);
              posz = [0 0];
              [x,y,~] = project(posx,posy,posz,projection);
              x = (x*axpos(3)+axpos(1))*paperpos(3);
              y = (1-(y*axpos(4)+axpos(2)))*paperpos(4);
              line2svg(fid,x,y,'#000000',linestyle,linewidth);
            end
          end
          % close the group
          fprintf(fid, '</g>');
      elseif strcmp(get(axchild(i),'Type'),'hgtransform')
          if strcmpi(get(axchild(i), 'Visible'), 'on')
              [filterString, boundingBox] = filter2svg(fid, axchild(i), boundingBoxAxes, boundingBoxAxes);
              if strcmp(get(axchild(i),'Clipping'),'on') && FIG2SVG_globals.ClippingMode
                  clippingIdString = clipping2svg(fid, axchild(i), ax, paperpos, axpos, projection, axIdString);
                  fprintf(fid,'<g id = "%s" clip-path = "url(#%s)" %s>\n', createId, clippingIdString, filterString);
              else
                  fprintf(fid, '<g id = "%s" %s>', createId, filterString);
              end
              if ~isempty(filterString)
                  % Workaround for Inkscape filter bug
                  fprintf(fid,'<rect x = "%0.3f" y = "%0.3f" width = "%0.3f" height = "%0.3f" fill = "none" stroke = "none" />\n', boundingBox(1), boundingBox(2), boundingBox(3), boundingBox(4));
              end
              boundingBoxAxes = axchild2svg(fid,id,axIdString,ax,paperpos,get(axchild(i), 'Children'),axpos,groupax,projection,boundingBoxAxes);
              fprintf(fid, '</g>');
          end
      else
          disp(['   Warning: Unhandled child type: ' get(axchild(i),'Type')]);
      end
  end
end

function [barCenters,barWidth] = groupBarProperties(currentBarSetNumber,barCenters,numGroups,barWidth,barSep)
  toPower = 0.1;
  tanh_factor = 2.5;
  minGroupSep = 0.713 - 0.49*tanh(tanh_factor*(numGroups^toPower-2^toPower)/(30^toPower-2^toPower));
  groupWidth = 1-minGroupSep;
  offset = groupWidth*[-0.5:1/(numGroups-1):0.5];
  barCenters = barCenters + barSep*offset(currentBarSetNumber);
  barWidth = min(diff(offset))*barWidth;
end

function result = selectColor(axchild, id, p, q, points, pointc, colorname, faces, type)
  if size(pointc, 1) == 1
      color = pointc;
  elseif size(pointc, 1) == size(faces, 1)
      color = pointc(p,:);
  elseif size(pointc, 1) == size(points, 2)
      if strcmp(get(axchild, type), 'flat')
          color = pointc(faces(p, q));
      else
          color = pointc(faces(p, q));     % TO DO: color interpolation
      end
  else
      error('Unsupported color handling for patches.');
  end
  if ~isnan(color)
      if size(color, 2) == 1
          result = ['#' colorname(ceil(color), :)];
      else
          if strcmp(get(axchild, type), 'flat')   % Bugfix 27.01.2008
              result = searchcolor(id, color / 64);
          else
              result = searchcolor(id, color);
          end
      end
  else
      result = 'none';
  end
end

%%%%%%%%%%%%%%%%%%%%%%%%%%%%%%%%%%%%%%%%%%%%%%%%%%%%%%%%%%%%%%%%%%%%%%%%%%%%%%%%%%%%%%%%%%%%%%%%%%%%%%%%%%%%%%%%%%%%%%%%%%%%%%%%%%%%%%%%%%%%%%%%%
% create a patch (filled area)
function patch2svg(fid,xtot,ytot,scolorname,style,width, edgecolorname, face_opacity, edge_opacity, closed)
  if closed
      type = 'polygon';
  else
      type = 'polyline';
  end
  pattern = lineStyle2svg(style, width);
  if strcmp(style, 'none')
      edge_opacity = 0.0;
  end
  for i = 1:size(xtot, 1)

    x = xtot(i, :);
    y = ytot(i, :);

    if ~iscell(edgecolorname)
      if all(~isnan(x)) && all(~isnan(y))
          for j = 1:20000:length(x)
            xx = x(j:min(length(x), j + 19999));
            yy = y(j:min(length(y), j + 19999));
            if ~isempty(xx) && ~isempty(yy) && (~strcmp(edgecolorname,'none') || ~strcmp(scolorname,'none'))
              if strcmp(type,'polygon')
                if (~strcmp(edgecolorname,'none')) && ~strcmp(scolorname,'none')
                  fprintf(fid,'      <%s fill = "%s" fill-opacity = "%0.3f" stroke-linecap = "square" stroke-linejoin = "miter" stroke = "%s" stroke-width = "%0.3fpt" stroke-opacity = "%0.3f" %s points = "', type, scolorname, face_opacity, edgecolorname, width, edge_opacity, pattern);
                elseif ~strcmp(scolorname,'none')
                  fprintf(fid,'      <%s fill = "%s" fill-opacity = "%0.3f" stroke = "none" points = "', type, scolorname, face_opacity);
                else % ~strcmp(edgecolorname,'none')
                  fprintf(fid,'      <%s fill = "none" stroke-linecap = "square" stroke-linejoin = "miter" stroke = "%s" stroke-width = "%0.3fpt" stroke-opacity = "%0.3f" %s points = "', type, edgecolorname, width, edge_opacity, pattern);
                end
                fprintf(fid,'%0.3f,%0.3f ',[xx;yy]);
                fprintf(fid,'"/>\n');
              else
                if ~strcmp(edgecolorname,'none') && ~strcmp(scolorname,'none')
                  fprintf(fid,'      <%s fill = "%s" fill-opacity = "%0.3f" stroke-linecap = "square" stroke-linejoin = "round" stroke = "%s" stroke-width = "%0.3fpt" stroke-opacity = "%0.3f" %s points = "', type, scolorname, face_opacity, edgecolorname, width, edge_opacity, pattern);
                elseif ~strcmp(scolorname,'none')
                  fprintf(fid,'      <%s fill = "%s" fill-opacity = "%0.3f" stroke = "none" points = "', type, scolorname, face_opacity);
                else % ~strcmp(edgecolorname,'none')
                  fprintf(fid,'      <%s fill = "none" stroke-linecap = "square" stroke-linejoin = "round" stroke = "%s" stroke-width = "%0.3fpt" stroke-opacity = "%0.3f" %s points = "', type, edgecolorname, width, edge_opacity, pattern);
                end
                fprintf(fid,'%0.3f,%0.3f ',[xx;yy]);
                fprintf(fid,'"/>\n');
              end
          end
        end
      else
        parts = find(isnan(x) + isnan(y));
        if ~isempty(parts) && (parts(1) ~= 1)
          parts = [0 parts];
        end
        if parts(length(parts)) ~= length(x)
          parts = [parts length(x) + 1];
        end
        for j = 1:(length(parts) - 1)
          xx = x((parts(j)+1):(parts(j+1)-1));
          yy = y((parts(j)+1):(parts(j+1)-1));
          if ~isempty(xx) && ~isempty(yy) && (~strcmp(edgecolorname,'none') || ~strcmp(scolorname,'none'))
            if ~isempty(xx) && ~isempty(yy)
              if strcmp(type,'polygon')
                fprintf(fid,'      <%s fill = "%s" fill-opacity = "%0.3f" stroke-linecap = "square" stroke-linejoin = "miter" stroke = "%s" stroke-width = "%0.3fpt" stroke-opacity = "%0.3f" %s points = "',...
                  type, scolorname, face_opacity, edgecolorname, width, edge_opacity, pattern);
              else
                fprintf(fid,'      <%s fill = "%s" fill-opacity = "%0.3f" stroke-linecap = "square" stroke-linejoin = "round" stroke = "%s" stroke-width = "%0.3fpt" stroke-opacity = "%0.3f" %s points = "',...
                    type, scolorname, face_opacity, edgecolorname, width, edge_opacity, pattern);
              end
              fprintf(fid,'%0.3f,%0.3f ',[xx;yy]);
              fprintf(fid,'"/>\n');
            end
          end
        end
      end
    else
      if all(~isnan(x)) && all(~isnan(y))
        for j = 1:20000:length(x)
          xx = x(j:min(length(x), j + 19999));
          yy = y(j:min(length(y), j + 19999));
          if ~isempty(xx) && ~isempty(yy)
            if strcmp(type,'polygon')
              if ~strcmp(scolorname,'none')
                fprintf(fid,'      <%s fill = "%s" fill-opacity = "%0.3f" stroke = "none" points = "', type, scolorname, face_opacity);
                fprintf(fid,'%0.3f,%0.3f ',[xx;yy]);
                fprintf(fid,'"/>\n');
              end
              for k = 1:numel(edgecolorname)
                if k ~= numel(edgecolorname)
                  xxx = [xx(k) xx(k+1)];
                  yyy = [yy(k) yy(k+1)];
                else
                  xxx = [xx(k) xx(1)];
                  yyy = [yy(k) yy(1)];
                end
                fprintf(fid,'      <%s fill = "none" stroke-linecap = "square" stroke-linejoin = "miter" stroke = "%s" stroke-width = "%0.3fpt" stroke-opacity = "%0.3f" %s points = "', type, edgecolorname{k}, width, edge_opacity, pattern);
                fprintf(fid,'%0.3f,%0.3f ',[xxx;yyy]);
                fprintf(fid,'"/>\n');
              end
            else
              if ~strcmp(scolorname,'none')
                fprintf(fid,'      <%s fill = "%s" fill-opacity = "%0.3f" stroke = "none" points = "', type, scolorname, face_opacity);
                fprintf(fid,'%0.3f,%0.3f ',[xx;yy]);
                fprintf(fid,'"/>\n');
              end
              for k = 1:numel(edgecolorname)
                if k ~= numel(edgecolorname)
                  xxx = [xx(k) xx(k+1)];
                  yyy = [yy(k) yy(k+1)];
                else
                  xxx = [xx(k) xx(1)];
                  yyy = [yy(k) yy(1)];
                end
                fprintf(fid,'      <%s fill = "none" stroke-linecap = "square" stroke-linejoin = "round" stroke = "%s" stroke-width = "%0.3fpt" stroke-opacity = "%0.3f" %s points = "', type, edgecolorname{k}, width, edge_opacity, pattern);
                fprintf(fid,'%0.3f,%0.3f ',[xxx;yyy]);
                fprintf(fid,'"/>\n');
              end
            end
          end
        end
      else
        parts = find(isnan(x) + isnan(y));
        if ~isempty(parts) && (parts(1) ~= 1)
          parts = [0 parts];
        end
        if parts(length(parts)) ~= length(x)
          parts = [parts length(x) + 1];
        end
        for j = 1:(length(parts) - 1)
          xx = x((parts(j)+1):(parts(j+1)-1));
          yy = y((parts(j)+1):(parts(j+1)-1));
          if ~isempty(xx) && ~isempty(yy)
            fprintf(fid,'      <%s fill = "%s" fill-opacity = "%0.3f" stroke = "none" points = "', type, scolorname, face_opacity);
            if strcmp(type,'polygon')
              for k = 1:numel(edgecolorname)
                if k ~= numel(edgecolorname)
                  xxx = [xx(k) xx(k+1)];
                  yyy = [yy(k) yy(k+1)];
                else
                  xxx = [xx(k) xx(1)];
                  yyy = [yy(k) yy(1)];
                end
                fprintf(fid,'      <%s fill = "none" stroke-linecap = "square" stroke-linejoin = "miter" stroke = "%s" stroke-width = "%0.3fpt" stroke-opacity = "%0.3f" %s points = "', type, edgecolorname{k}, width, edge_opacity, pattern);
              end
              fprintf(fid,'%0.3f,%0.3f ',[xxx;yyy]);
              fprintf(fid,'"/>\n');
            else
              for k = 1:numel(edgecolorname)
                if k ~= numel(edgecolorname)
                  xxx = [xx(k) xx(k+1)];
                  yyy = [yy(k) yy(k+1)];
                else
                  xxx = [xx(k) xx(1)];
                  yyy = [yy(k) yy(1)];
                end
                fprintf(fid,'      <%s fill = "none" stroke-linecap = "square" stroke-linejoin = "round" stroke = "%s" stroke-width = "%0.3fpt" stroke-opacity = "%0.3f" %s points = "', type, edgecolorname{k}, width, edge_opacity, pattern);
                fprintf(fid,'%0.3f,%0.3f ',[xxx;yyy]);
                fprintf(fid,'"/>\n');
              end
            end
          end
        end
      end
    end
  end
end

%%%%%%%%%%%%%%%%%%%%%%%%%%%%%%%%%%%%%%%%%%%%%%%%%%%%%%%%%%%%%%%%%%%%%%%%%%%%%%%%%%%%%%%%%%%%%%%%%%%%%%%%%%%%%%%%%%%%%%%%%%%%%%%%%%%%%%%%%%%%%%%%%
% create a patch (filled area)
function gouraud_patch2svg(fid,xtot,ytot,cdata,style,width, edgecolorname, face_opacity, edge_opacity,id)
  global colorname
  pattern = lineStyle2svg(style, width);
  if strcmp(style, 'none')
      edge_opacity = 0.0;
  end
  for ii = 1:size(xtot,1)
      x = xtot(ii,:);
      y = ytot(ii,:);
      if (any(isnan(x)) || any(isnan(y)))
          % SA: commenting this out as it is overdose
          % fprintf('Warning: Found NaN in Gouraud patch.\n')
      else
          % If there are more than 2 edges always 3 edges are taken together
          % to form a triangle
          if length(x) > 2
              for j = 3:length(x)
                  coord = [x([1 j-1 j]);y([1 j-1 j])];
                  if isempty(coord)
                    continue
                  end
                  face_color = cdata(1,:);
                  face_color2 = cdata(j-1,:);
                  face_color3 = cdata(j,:);
                  delta = coord(:,3)-coord(:,2);
                  if det([delta (coord(:,1)-coord(:,2))]) ~= 0
                      if ~isnan(face_color)
                          IDstring1 = createId;
                          IDstring2 = createId;
                          if size(face_color2,2) == 1
                              face_color_name2 = ['#' colorname(ceil(face_color2),:)];
                          else
                              face_color_name2 = searchcolor(id,face_color2/64);
                          end
                          if size(face_color3,2) == 1
                              face_color_name3 = ['#' colorname(ceil(face_color3),:)];
                          else
                              face_color_name3 = searchcolor(id,face_color3/64);
                          end
                          grad_end = (delta)*(delta'*(coord(:,1)-coord(:,2)))/(delta'*delta) + coord(:,2);
                          if size(face_color,2) == 1
                              face_color_name = ['#' colorname(ceil(face_color),:)];
                          else
                              face_color_name = searchcolor(id,face_color/64);
                          end
                          fprintf(fid,'<defs>\n');
                          fprintf(fid,'<linearGradient id = "%s" gradientUnits = "userSpaceOnUse" x1 = "%0.3f" y1 = "%0.3f" x2 = "%0.3f" y2 = "%0.3f">\n',...
                              IDstring1, coord(1,2), coord(2,2), coord(1,3), coord(2,3));
                          fprintf(fid,'<stop offset = "0" stop-color = "%s" stop-opacity = "1"/>\n',face_color_name2);
                          fprintf(fid,'<stop offset = "1" stop-color = "%s" stop-opacity = "1"/>\n',face_color_name3);
                          fprintf(fid,'</linearGradient>\n');
                          fprintf(fid,'<linearGradient id = "%s" gradientUnits = "userSpaceOnUse" x1 = "%0.3f" y1 = "%0.3f" x2 = "%0.3f" y2 = "%0.3f">\n',...
                              IDstring2, coord(1,1), coord(2,1), grad_end(1), grad_end(2));
                          fprintf(fid,'<stop offset = "0" stop-color = "%s" stop-opacity = "1"/>\n',face_color_name);
                          fprintf(fid,'<stop offset = "1" stop-color = "%s" stop-opacity = "0"/>\n',face_color_name);
                          fprintf(fid,'</linearGradient>\n');
                          fprintf(fid,'</defs>\n');
                          % Open group
                          temp_string = sprintf('%0.3f,%0.3f ',coord);
                          fprintf(fid,'<g opacity = "%0.3f">\n',face_opacity);
                          fprintf(fid,'<polyline fill = "url(#%s)" stroke = "none" points = "%s"/>\n',IDstring1,temp_string);
                          fprintf(fid,'<polyline fill = "url(#%s)" stroke = "none" points = "%s"/>\n',IDstring2,temp_string);
                          % Close group
                          fprintf(fid,'</g>\n');
                      end
                  end
              end
          end
          % Last we draw the line around the patch
          if ~strcmp(edgecolorname,'none')
              fprintf(fid,'<polygon fill = "none" stroke-linecap = "square" stroke-linejoin = "miter" stroke = "%s" stroke-width = "%0.3fpt" stroke-opacity = "%0.3f" %s points = "',...
                  edgecolorname, width, edge_opacity, pattern);
              fprintf(fid,'%0.3f,%0.3f ',[x;y]);
              fprintf(fid,'"/>\n');
          end
      end
  end
end

function patternString = lineStyle2svg(lineStyle, lineWidth)
  global FIG2SVG_globals
  % Create the string for the line style
  % Note: The line style is not the same on screen as it appears in png files.
  %       Getting the screen format as reference.
  scaling = 1/FIG2SVG_globals.resolutionScaling; % max(1, lineWidth * 0.4);
  switch lineStyle
      case '--', patternString = sprintf('stroke-dasharray = "%0.3f,%0.3f"', 6*scaling, 10*scaling); % updated to be similar to Matlab
      case ':', patternString = sprintf('stroke-dasharray = "%0.3f,%0.3f"', 0.25*scaling, 6*scaling); % updated so it looks better
      case '-.', patternString = sprintf('stroke-dasharray = "%0.3f,%0.3f,%0.3f,%0.3f"', 6*scaling, 4.875*scaling, 0.25*scaling, 4.875*scaling); % updated to be similar to Matlab
      otherwise, patternString = 'stroke-dasharray = "none"';
  end
end

%%%%%%%%%%%%%%%%%%%%%%%%%%%%%%%%%%%%%%%%%%%%%%%%%%%%%%%%%%%%%%%%%%%%%%%%%%%%%%%%%%%%%%%%%%%%%%%%%%%%%%%%%%%%%%%%%%%%%%%%%%%%%%%%%%%%%%%%%%%%%%%%%
% create a line segment
% this algorthm was optimized for large segement counts
function line2svg(fid, x, y, scolorname, style, width, linejoin, strokeopacity)
  SEG_SIZE = 5000;
  if nargin < 7 || isempty(linejoin)
      linejoin = 'miter';
  end
  % tried this but does not make a difference
  % if strcmp(linejoin,'round')
  %   linecap = 'butt';
  % else
  %   linecap = 'square';
  % end
  if nargin < 8 || ~isscalar(strokeopacity)
      strokeopacity = 1;
  end
  if ~strcmp(style,'none')
      pattern = lineStyle2svg(style, width);

      skip_pts = reshape(find(isnan(x) | isnan(y)), [],1);
      start_pts = [1; skip_pts+1];
      end_pts = [skip_pts-1; numel(x)];
      k = 1;
      while k <= numel(start_pts)
          if (end_pts(k) - start_pts(k)) > SEG_SIZE
              tmp_sPts = zeros(numel(start_pts)+1,1);
              tmp_ePts = zeros(numel(start_pts)+1,1);
              tmp_sPts(1:k) = start_pts(1:k);
              tmp_ePts(1:k-1) = end_pts(1:k-1);
              tmp_ePts(k) = tmp_sPts(k)+SEG_SIZE-1;
              tmp_sPts(k+1) = tmp_sPts(k)+SEG_SIZE-1;
              tmp_ePts(k+1:end) = end_pts(k:end);
              tmp_sPts(k+2:end) = start_pts(k+1:end);
              start_pts = tmp_sPts;
              end_pts = tmp_ePts;
          end
          k = k+1;
      end
      for j = 1:numel(start_pts)
          xx = x(start_pts(j):end_pts(j));
          yy = y(start_pts(j):end_pts(j));
          if ~isempty(xx) && ~isempty(yy)
            % Scaling stroke is a better default
            % fprintf(fid,'      <polyline fill = "none" vector-effect = "non-scaling-stroke" stroke = "%s" stroke-width = "%0.3fpt" %s stroke-opacity = "%0.3f" points = "', scolorname, width, pattern, strokeopacity);
            fprintf(fid,'      <polyline fill = "none" stroke-linecap = "square" stroke-linejoin = "%s" stroke = "%s" stroke-width = "%0.3fpt" %s stroke-opacity = "%0.3f" points = "', linejoin, scolorname, width, pattern, strokeopacity);
            fprintf(fid,'%0.3f,%0.3f ',[xx;yy]);
            fprintf(fid,'"/>\n');
          end
      end
  end
end

%%%%%%%%%%%%%%%%%%%%%%%%%%%%%%%%%%%%%%%%%%%%%%%%%%%%%%%%%%%%%%%%%%%%%%%%%%%%%%%%%%%%%%%%%%%%%%%%%%%%%%%%%%%%%%%%%%%%%%%%%%%%%%%%%%%%%%%%%%%%%%%%%
% create a circle
function circle2svg(fid,x,y,radius,markeredgecolorname,markerfacecolorname,width,markerfaceopacity,markeredgeopacity)
  if numel(markerfacecolorname) == 1 && numel(markeredgecolorname) == 1 && strcmpi(markerfacecolorname, 'none') && strcmpi(markeredgecolorname, 'none')
    return
  end
  if ~isempty(x) && ~isempty(y)
    for ii = 1:length(x)
      if ~isnan(x(ii)) && ~isnan(y(ii))
        if size(markeredgecolorname,1) > 1 && size(markerfacecolorname,1) > 1
            fprintf(fid,'<circle cx = "%0.3f" cy = "%0.3f" r = "%0.3f" fill = "%s" fill-opacity = "%0.3f" stroke-linecap = "square" stroke-linejoin = "miter" stroke = "%s" stroke-opacity = "%0.3f" stroke-width = "%0.3fpt" />\n',x(ii),y(ii),radius,markerfacecolorname{ii},markerfaceopacity,markeredgecolorname{ii},markeredgeopacity,width);
        elseif size(markeredgecolorname,1) > 1
            if iscell(markerfacecolorname)
              markerfacecolorname = markerfacecolorname{1};
            end
            fprintf(fid,'<circle cx = "%0.3f" cy = "%0.3f" r = "%0.3f" fill = "%s" fill-opacity = "%0.3f" stroke-linecap = "square" stroke-linejoin = "miter" stroke = "%s" stroke-opacity = "%0.3f" stroke-width = "%0.3fpt" />\n',x(ii),y(ii),radius,markerfacecolorname,markerfaceopacity,markeredgecolorname{ii},markeredgeopacity,width);
        elseif size(markerfacecolorname,1) > 1
            if iscell(markeredgecolorname)
              markeredgecolorname = markeredgecolorname{1};
            end
            fprintf(fid,'<circle cx = "%0.3f" cy = "%0.3f" r = "%0.3f" fill = "%s" fill-opacity = "%0.3f" stroke-linecap = "square" stroke-linejoin = "miter" stroke = "%s" stroke-opacity = "%0.3f" stroke-width = "%0.3fpt" />\n',x(ii),y(ii),radius,markerfacecolorname{ii},markerfaceopacity,markeredgecolorname,markeredgeopacity,width);
        else
            if iscell(markerfacecolorname)
              markerfacecolorname = markerfacecolorname{1};
            end
            if iscell(markeredgecolorname)
              markeredgecolorname = markeredgecolorname{1};
            end
            fprintf(fid,'<circle cx = "%0.3f" cy = "%0.3f" r = "%0.3f" fill = "%s" fill-opacity = "%0.3f" stroke-linecap = "square" stroke-linejoin = "miter" stroke = "%s" stroke-opacity = "%0.3f" stroke-width = "%0.3fpt" />\n',x(ii),y(ii),radius,markerfacecolorname,markerfaceopacity,markeredgecolorname,markeredgeopacity,width);
        end
      end
    end
  end
end
%%%%%%%%%%%%%%%%%%%%%%%%%%%%%%%%%%%%%%%%%%%%%%%%%%%%%%%%%%%%%%%%%%%%%%%%%%%%%%%%%%%%%%%%%%%%%%%%%%%%%%%%%%%%%%%%%%%%%%%%%%%%%%%%%%%%%%%%%%%%%%%%%
% create a scatter patch
function scatterpatch2svg(fid, x, y, markerfacecolorname, style, width, markeredgecolorname, face_opacity, edge_opacity, closed)
  if numel(markerfacecolorname) == 1 && numel(markeredgecolorname) == 1 && strcmpi(markerfacecolorname, 'none') && strcmpi(markeredgecolorname, 'none')
    return
  end
  if ~isempty(x) && ~isempty(y)
    for ii = 1:size(x,1)
      if ~any(isnan(x(ii,:)) | isnan(y(ii,:)))
        if size(markerfacecolorname,1) > 1 && size(markeredgecolorname,1) > 1
            patch2svg(fid, x(ii,:), y(ii,:), markerfacecolorname{ii}, style, width, markeredgecolorname{ii}, face_opacity, edge_opacity, closed)
        elseif size(markeredgecolorname,1) > 1
            if iscell(markerfacecolorname)
              markerfacecolorname = markerfacecolorname{1};
            end
            patch2svg(fid, x(ii,:), y(ii,:), markerfacecolorname, style, width, markeredgecolorname{ii}, face_opacity, edge_opacity, closed)
        elseif size(markerfacecolorname,1) > 1
            if iscell(markeredgecolorname)
              markeredgecolorname = markeredgecolorname{1};
            end
            patch2svg(fid, x(ii,:), y(ii,:), markerfacecolorname{ii}, style, width, markeredgecolorname, face_opacity, edge_opacity, closed)
        else
            if iscell(markerfacecolorname)
              markerfacecolorname = markerfacecolorname{1};
            end
            if iscell(markeredgecolorname)
              markeredgecolorname = markeredgecolorname{1};
            end
            patch2svg(fid, x(ii,:), y(ii,:), markerfacecolorname, style, width, markeredgecolorname, face_opacity, edge_opacity, closed)
        end
      end
    end
  end
end

%%%%%%%%%%%%%%%%%%%%%%%%%%%%%%%%%%%%%%%%%%%%%%%%%%%%%%%%%%%%%%%%%%%%%%%%%%%%%%%%%%%%%%%%%%%%%%%%%%%%%%%%%%%%%%%%%%%%%%%%%%%%%%%%%%%%%%%%%%%%%%%%%
function control2svg(fid,id,ax,paperpos)
  global FIG2SVG_globals
  set(ax,'Units','pixels');
  if FIG2SVG_globals.octave || UIverlessthan('8.4.0')
      pos = get(ax,'Position');
  else
      pos = ax.OuterPosition;
  end
  pict = getframe(id,pos);
  if isempty(pict.colormap)
      pict.colormap = colormap;
  end
  filename = [FIG2SVG_globals.basefilename sprintf('%03d',FIG2SVG_globals.figurenumber) '.' FIG2SVG_globals.pixelFileType];
  FIG2SVG_globals.figurenumber = FIG2SVG_globals.figurenumber + 1;
  if isempty(FIG2SVG_globals.basefilepath)
      current_path = pwd;
  else
      current_path = FIG2SVG_globals.basefilepath;
  end
  if exist(fullfile(current_path,filename),'file')
      lastwarn('');
      delete(filename);
      if strcmp(lastwarn,'File not found or permission denied.')
          error('Cannot write image file. Make sure that no image is opened in an other program.')
      end
  end
  imwrite(pict.cdata,fullfile(FIG2SVG_globals.basefilepath,filename),FIG2SVG_globals.pixelFileType);
  set(ax,'Units','normalized');
  posNorm = get(ax,'Position');
  posInches(1) = posNorm(1)*paperpos(3);
  posInches(2) = posNorm(2)*paperpos(4);
  posInches(3) = posNorm(3)*paperpos(3);
  posInches(4) = posNorm(4)*paperpos(4);
  lx = posInches(3);
  ly = posInches(4);
  pointsx = posInches(1);
  pointsy = paperpos(4)-posInches(2)-posInches(4);
  fprintf(fid,'<image x = "%0.3f" y = "%0.3f" width = "%0.3f" height = "%0.3f" image-rendering = "optimizeQuality" xlink:href = "%s" />\n', pointsx, pointsy, lx, ly, filename);
end

%%%%%%%%%%%%%%%%%%%%%%%%%%%%%%%%%%%%%%%%%%%%%%%%%%%%%%%%%%%%%%%%%%%%%%%%%%%%%%%%%%%%%%%%%%%%%%%%%%%%%%%%%%%%%%%%%%%%%%%%%%%%%%%%%%%%%%%%%%%%%%%%%
% create a text in the axis frame
% the position of the text has to be adapted to the axis scaling
function text2svg(fid,axpos,paperpos,id,ax,projection)
  global FIG2SVG_globals;
  originalTextUnits = get(id,'Units');
  originalTextPosition = get(id, 'Position');

  if ~strcmp(get(ax,'Type'),'annotationpane')
    if FIG2SVG_globals.octave
      set(id,'Units','data');
    else
      set(id,'Units','Data');
    end
    textpos = get(id,'Position');
    try
      if strcmp(get(ax,'XScale'),'log')
          textpos(1) = log10(textpos(1));
      end
      if strcmp(get(ax,'YScale'),'log')
          textpos(2) = log10(textpos(2));
      end
      if strcmp(get(ax,'ZScale'),'log')
          textpos(3) = log10(textpos(3));
      end
      [x,y,~] = project(textpos(1), textpos(2), textpos(3), projection);
    catch
      x = textpos(1);
      y = textpos(2);
    end
    x = (x * axpos(3) + axpos(1))*paperpos(3);
    y = (1 - (y * axpos(4) + axpos(2)))*paperpos(4);
  else
    x = axpos(1);
    y = axpos(2);
  end

  % textfontsize = get(id,'FontSize');
  fontsize = convertunit(get(id,'FontSize'),get(id,'FontUnits'),'points', axpos(4));   % convert fontsize to inches
  % paperposOriginal = get(gcf,'Position');
  font_color = searchcolor(id,get(id,'Color'));

  textalign = get(id,'HorizontalAlignment');
  textvalign = get(id,'VerticalAlignment');

  % checking if this is a 2-D or 3-D plot: gca axes in 2-D plots can appear in z-axis at the bottom layer [default] z = -1, at the same layer [default for legends] z = 0, or at the top layer z = 1:
  if ~strcmp(get(ax,'Type'),'annotationpane') && numel(textpos) > 2 && all(abs(textpos(3)-[-1,0,1]) > 1e-10) % 3D plot
    textvalign = ['3d-',textvalign];
  end

  texttext = get(id,'String');
  if strcmp(get(ax,'Type'),'annotationpane')
    textrot = get(id,'FontAngle');
    if ischar(textrot) && strcmp(textrot,'normal')
      textrot = 0;
    else
      disp(textrot);
      error('unsupported font angle');
    end
  else
    textrot = get(id,'Rotation');
  end
  dx = sin(textrot * pi / 180) * convertunit(fontsize * 1.2, 'points', 'pixels');
  dy = cos(textrot * pi / 180) * convertunit(fontsize * 1.2, 'points', 'pixels');
  lines = max(size(get(id,'String'),1),1);
  if size(texttext,2) ~= 0
      j = 1;
      for i = 0:1:(lines - 1)
          if iscell(texttext)
              label2svg(fid, axpos, id, x + i * dx, y + i * dy, convertString(texttext{j}), textalign, textrot, textvalign, lines, font_color)
          else
              label2svg(fid, axpos, id, x + i * dx, y + i * dy, convertString(texttext(j,:)), textalign, textrot, textvalign, lines, font_color)
          end
          j = j + 1;
      end
  else
      label2svg(fid,axpos,id,x,y,'',textalign,textrot,textvalign,lines,font_color)
  end
  % SA: the following two lines don't seem to do anything and break the consistency of the figure in Matlab
  % pause
  % set(id,'Units',originalTextUnits);
  % set(id,'Position', originalTextPosition);
end

%%%%%%%%%%%%%%%%%%%%%%%%%%%%%%%%%%%%%%%%%%%%%%%%%%%%%%%%%%%%%%%%%%%%%%%%%%%%%%%%%%%%%%%%%%%%%%%%%%%%%%%%%%%%%%%%%%%%%%%%%%%%%%%%%%%%%%%%%%%%%%%%%
% adds the exponents to the axis thickmarks if needed
% MATLAB itself offers no information about this exponent scaling
% the exponent have therefore to be extracted from the thickmarks
function exponent2svg(fid,axpos,paperpos,ax,axxtick,axytick,axztick)
  global FIG2SVG_globals
  if strcmp(get(ax,'Type'),'colorbar')
      fontsize = convertunit(get(ax,'FontSize'),get(ax,'Units'),'points', axpos(4));   % convert fontsize to inches
      font_color = searchcolor(ax,get(ax,'Color'));
      axlabel = get (ax, 'TickLabels');
      if iscell(axlabel)
        % Also new Matlab versions
        % Octave stores YTickLabel in a cell array, which does not work nicely with str2num. --Jakob Malm
        numlabels = zeros(length(axlabel), 1);
        for ix = 1:length(axlabel)
          if ~isempty(axlabel{ix})
            numlabels(ix) = str2num(axlabel{ix});
          end
        end
      else
          numlabels = get(ax,'XTickLabel');
          if ~isempty(numlabels)
            numlabels_tmp = str2double(numlabels);
            if any(isnan(numlabels_tmp)) || size(numlabels,1) ~= size(numlabels_tmp,1)
              numlabels = str2num(numlabels);
            else
              numlabels = numlabels_tmp;
            end
          end
      end
      labelpos = axxtick; % get(ax,'XTick');
      numlabels = numlabels(:);
      labelpos = labelpos(:);
      indexnz = find(labelpos ~= 0);
      if (~isempty(indexnz) && ~isempty(numlabels))
          if (max(indexnz) <= length(numlabels))
              ratio = numlabels(labelpos ~= 0)./labelpos(labelpos ~= 0);
          else
              ratio = 1;
          end
          if round(log10(ratio(1))) ~= 0 && ratio(1) ~= 0
              exptext = sprintf('&#215; 10<tspan style = "font-size:65%%;baseline-shift:super">%g</tspan>', -log10(ratio(1)));
              label2svg(fid,axpos,ax,(axpos(1)+axpos(3))*paperpos(3),(1-axpos(2))*paperpos(4)+3*fontsize,exptext,'right',0,'top',1,font_color)
          end
      end
  else
    if strcmp(get(ax,'XTickLabelMode'),'auto') && strcmp(get(ax,'XScale'),'linear')
        fontsize = convertunit(get(ax,'FontSize'),get(ax,'FontUnits'),'points', axpos(4));   % convert fontsize to inches
        font_color = searchcolor(ax,get(ax,'XColor'));
        axlabelx = get (ax, 'XTickLabel');
        if iscell(axlabelx)
          % Also new Matlab versions
          % Octave stores YTickLabel in a cell array, which does not work nicely with str2num. --Jakob Malm
          numlabels = zeros(length(axlabelx), 1);
          for ix = 1:length(axlabelx)
            if ~isempty(axlabelx{ix})
              numlabels(ix) = str2num(axlabelx{ix});
            end
          end
        else
            numlabels = get(ax,'XTickLabel');
            if ~isempty(numlabels)
              numlabels_tmp = str2double(numlabels);
              if any(isnan(numlabels_tmp)) || size(numlabels,1) ~= size(numlabels_tmp,1)
                numlabels = str2num(numlabels);
              else
                numlabels = numlabels_tmp;
              end
            end
        end
        labelpos = axxtick; % get(ax,'XTick');
        numlabels = numlabels(:);
        labelpos = labelpos(:);
        indexnz = find(labelpos ~= 0);
        if (~isempty(indexnz) && ~isempty(numlabels))
            if (max(indexnz) <= length(numlabels))
                ratio = numlabels(labelpos ~= 0)./labelpos(labelpos ~= 0);
            else
                ratio = 1;
            end
            if round(log10(ratio(1))) ~= 0 && ratio(1) ~= 0
                exptext = sprintf('&#215; 10<tspan style = "font-size:65%%;baseline-shift:super">%g</tspan>', -log10(ratio(1)));
                label2svg(fid,axpos,ax,(axpos(1)+axpos(3))*paperpos(3),(1-axpos(2))*paperpos(4)+3*fontsize,exptext,'right',0,'top',1,font_color)
            end
        end
    end
    if strcmp(get(ax,'YTickLabelMode'),'auto') && strcmp(get(ax,'YScale'),'linear')
        fontsize = convertunit(get(ax,'FontSize'),get(ax,'FontUnits'),'points', axpos(4));
        font_color = searchcolor(ax,get(ax,'YColor'));
        axlabely = get(ax, 'YTickLabel');
        if iscell(axlabely)
          % Also new Matlab versions
          % Octave stores YTickLabel in a cell array, which does not work nicely with str2num. --Jakob Malm
          numlabels = zeros(length(axlabely), 1);
          for ix = 1:length(axlabely)
            if ~isempty(axlabely{ix})
              numlabels(ix) = str2num(axlabely{ix});
            end
          end
        else
            numlabels = axlabely;
            if ~isempty(numlabels)
              numlabels_tmp = str2double(numlabels);
              if any(isnan(numlabels_tmp)) || size(numlabels,1) ~= size(numlabels_tmp,1)
                numlabels = str2num(numlabels);
              else
                numlabels = numlabels_tmp;
              end
            end
        end
        labelpos = axytick; % get(ax,'YTick');
        numlabels = numlabels(:);
        labelpos = labelpos(:);
        indexnz = find(labelpos ~= 0);
        if (~isempty(indexnz) && ~isempty(numlabels))
            if (max(indexnz) <= length(numlabels))
                ratio = numlabels(labelpos ~= 0)./labelpos(labelpos ~= 0);
            else
                ratio = 1;
            end
            if round(log10(ratio(1))) ~= 0 && ratio(1) ~= 0
                exptext = sprintf('&#215; 10<tspan style = "font-size:65%%;baseline-shift:super">%g</tspan>', -log10(ratio(1)));
                label2svg(fid,axpos,ax,axpos(1)*paperpos(3),(1-(axpos(2)+axpos(4)))*paperpos(4)-0.5*fontsize,exptext,'left',0,'bottom',1,font_color)
            end
        end
    end
    if strcmp(get(ax,'ZTickLabelMode'),'auto') && strcmp(get(ax,'ZScale'),'linear')
        fontsize = convertunit(get(ax,'FontSize'),get(ax,'FontUnits'),'points', axpos(4));
        font_color = searchcolor(ax,get(ax,'ZColor'));
        axlabelz = get (ax, 'ZTickLabel');
        if iscell(axlabelz)
          % Also new Matlab versions
          % Octave stores YTickLabel in a cell array, which does not work nicely with str2num. --Jakob Malm
          numlabels = zeros(length(axlabelz), 1);
          for ix = 1:length(axlabelz)
            if ~isempty(axlabelz{ix})
              numlabels(ix) = str2num(axlabelz{ix});
            end
          end
        else
            numlabels = get(ax,'ZTickLabel');
            if ~isempty(numlabels)
              numlabels_tmp = str2double(numlabels);
              if any(isnan(numlabels_tmp)) || size(numlabels,1) ~= size(numlabels_tmp,1)
                numlabels = str2num(numlabels);
              else
                numlabels = numlabels_tmp;
              end
            end
        end
        labelpos = axztick; % get(ax,'ZTick');
        numlabels = numlabels(:);
        labelpos = labelpos(:);
        indexnz = find(labelpos ~= 0);
        if (~isempty(indexnz) && ~isempty(numlabels))
            if (max(indexnz) <= length(numlabels))
                ratio = numlabels(labelpos ~= 0)./labelpos(labelpos ~= 0);
            else
                ratio = 1;
            end
            if round(log10(ratio(1))) ~= 0 && ratio(1) ~= 0
                exptext = sprintf('&#215; 10<tspan style = "font-size:65%%;baseline-shift:super">%g</tspan>', -log10(ratio(1)));
                label2svg(fid,axpos,ax,axpos(1)*paperpos(3),(1-(axpos(2)+axpos(4)))*paperpos(4)-0.5*fontsize,exptext,'left',0,'top',1,font_color)
            end
        end
      end
  end
end

%%%%%%%%%%%%%%%%%%%%%%%%%%%%%%%%%%%%%%%%%%%%%%%%%%%%%%%%%%%%%%%%%%%%%%%%%%%%%%%%%%%%%%%%%%%%%%%%%%%%%%%%%%%%%%%%%%%%%%%%%%%%%%%%%%%%%%%%%%%%%%%%%
% create a label in the figure
% former versions of FrameMaker supported the commands FDY and FDX to shift the text
% this commands were replaced by a shift parameter that is normed by the font size
function label2svg(fid,axpos,id,x,y,tex,align,angle,valign,lines,font_color)
  global FIG2SVG_globals
  if isempty(tex)
      return;
  end
  textfontname = get(id,'FontName');
  if strcmp(textfontname, '*')
      textfontname = 'Arial';
  end
  try % ~strcmp(get(ax,'Type'),'colorbar')
    set(id,'FontUnits','points');
  catch % colorbar
    set(id,'Units','points');
  end
  textfontsize = get(id,'FontSize');
  if isfield(get(id),'Interpreter')
      if strcmp(get(id,'Interpreter'),'tex')
          latex = 1;
      elseif strcmp(get(id,'Interpreter'),'latex')
          latex = 1;
      else
          latex = 0;
      end
  else
      latex = 1;
  end
  try % ~strcmp(get(ax,'Type'),'colorbar')
    fontsize = convertunit(get(id,'FontSize'),get(id,'FontUnits'),'points', axpos(4));   % convert fontsize to inches
  catch % colorbar
    fontsize = convertunit(get(id,'FontSize'),get(id,'Units'),'points', axpos(4));   % convert fontsize to inches;
  end
  fontweight = get(id,'FontWeight');
  switch lower(fontweight)
      case 'bold', fweight = ' font-weight = "bold"';
      case 'light', fweight = ' font-weight = "lighter"';
      case 'demi', fweight = ' font-weight = "lighter"';
      otherwise, fweight = '';   % default 'normal'
  end
  fontangle = get(id,'FontAngle');
  switch lower(fontangle)
      case 'italic', fangle = ' font-style = "italic"';
      case 'oblique', fangle = ' font-style = "oblique"';
  otherwise, fangle = '';  % default 'normal'
  end
  balign = 'auto';
  y_offset = '0.3em';
  switch lower(valign)
  case 'top', y_offset = '0.9em'; % balign = 'text-before-edge';
  case '3d-top', y_offset = '0.9em'; % balign = 'middle'; % 'central';
  case '3d-bottom', y_offset = '-0.6em'; % balign = 'middle'; % 'central';
  % case 'cap', error('use top instead of cap');
  % case 'baseline', error('use bottom instead of baseline');
  case 'bottom', y_offset = '-0.2em'; % 'text-after-edge';
  % otherwise, % balign = 'middle'; % 'central';
  end
  switch lower(align)
    case 'right', anchor = 'end';
    case 'center', anchor = 'middle';
    otherwise, anchor = 'start'; % left alignment (default)
  end
  if abs(angle) > 1e-10 && ( strcmp(valign,'top') && ~strcmp(get(id,'Type'),'text') || strcmp(valign,'3d-bottom') )
    if angle > 1e-10
      anchor = 'end';
    else
      anchor = 'start';
    end
    dx = sin(angle * pi / 180) * convertunit(fontsize * (-0.5), 'points', 'pixels');
    dy = cos(angle * pi / 180) * convertunit(fontsize * 0, 'points', 'pixels');
  elseif abs(angle) > 1e-10 && ( strcmp(valign,'bottom') && ~strcmp(get(id,'Type'),'text') || strcmp(valign,'3d-top') )
    if angle > 1e-10
      anchor = 'start';
    else
      anchor = 'end';
    end
    dx = sin(angle * pi / 180) * convertunit(fontsize * 0.5, 'points', 'pixels');
    dy = cos(angle * pi / 180) * convertunit(fontsize * 0, 'points', 'pixels');
  elseif abs(angle) > 1e-10 && strcmpi(align,'left')
    dx = cos(angle * pi / 180) * convertunit(fontsize * 0, 'points', 'pixels');
    dy = sin(angle * pi / 180) * convertunit(fontsize * 0.25, 'points', 'pixels');
  elseif abs(angle) > 1e-10 && strcmpi(align,'right')
    dx = cos(angle * pi / 180) * convertunit(fontsize * (-0.25), 'points', 'pixels');
    dy = sin(angle * pi / 180) * convertunit(fontsize * 0, 'points', 'pixels');
  else
    dx = 0;
    dy = 0;
  end
  if iscellstr(tex)
      tex = strvcat(tex);
  elseif ~ ischar(tex)
      error('Invalid character type');
  end
  if latex == 1
      tex = strrep(tex,'$','');

      tex = strrep(tex,'\alpha','{&#945;}');
      tex = strrep(tex,'\beta','{&#946;}');
      tex = strrep(tex,'\gamma','{&#947;}');
      tex = strrep(tex,'\delta','{&#948;}');
      tex = strrep(tex,'\epsilon','{&#949;}');
      tex = strrep(tex,'\zeta','{&#950;}');
      tex = strrep(tex,'\eta','{&#951;}');
      tex = strrep(tex,'\theta','{&#952;}');
      tex = strrep(tex,'\vartheta','{&#977;}');
      tex = strrep(tex,'\iota','{&#953;}');
      tex = strrep(tex,'\kappa','{&#954;}');
      tex = strrep(tex,'\lambda','{&#955;}');
      tex = strrep(tex,'\mu','{&#181;}');
      tex = strrep(tex,'\nu','{&#957;}');
      tex = strrep(tex,'\xi','{&#958;}');
      tex = strrep(tex,'\pi','{&#960;}');
      tex = strrep(tex,'\rho','{&#961;}');
      tex = strrep(tex,'\sigma','{&#963;}');
      tex = strrep(tex,'\varsigma','{&#962;}');
      tex = strrep(tex,'\tau','{&#964;}');
      tex = strrep(tex,'\upsilon','{&#965;}');
      tex = strrep(tex,'\phi','{&#966;}');
      tex = strrep(tex,'\chi','{&#967;}');
      tex = strrep(tex,'\psi','{&#968;}');
      tex = strrep(tex,'\omega','{&#969;}');
      tex = strrep(tex,'\Gamma','{&#915;}');
      tex = strrep(tex,'\Delta','{&#916;}');
      tex = strrep(tex,'\Theta','{&#920;}');
      tex = strrep(tex,'\Lambda','{&#923;}');
      tex = strrep(tex,'\Xi','{&#926;}');
      tex = strrep(tex,'\Pi','{&#928;}');
      tex = strrep(tex,'\Sigma','{&#931;}');
      tex = strrep(tex,'\Tau','{&#932;}');
      tex = strrep(tex,'\Upsilon','{&#933;}');
      tex = strrep(tex,'\Phi','{&#934;}');
      tex = strrep(tex,'\Psi','{&#936;}');
      tex = strrep(tex,'\Omega','{&#937;}');
      tex = strrep(tex,'\infty','{&#8734;}');
      tex = strrep(tex,'\pm','{&#177;}');
      tex = strrep(tex,'\Im','{&#8465;}');
      tex = strrep(tex,'\Re','{&#8476;}');
      tex = strrep(tex,'\approx','{&#8773;}');
      tex = strrep(tex,'\leq','{&#8804;}');
      tex = strrep(tex,'\geq','{&#8805;}');
      tex = strrep(tex,'\times','{&#215;}');
      tex = strrep(tex,'\leftrightarrow','{&#8596;}');
      tex = strrep(tex,'\leftarrow','{&#8592;}');
      tex = strrep(tex,'\uparrow','{&#8593;}');
      tex = strrep(tex,'\rightarrow','{&#8594;}');
      tex = strrep(tex,'\downarrow','{&#8595;}');
      tex = strrep(tex,'\circ','{&#186;}');
      tex = strrep(tex,'\propto','{&#8733;}');
      tex = strrep(tex,'\partial','{&#8706;}');
      tex = strrep(tex,'\bullet','{&#8226;}');
      tex = strrep(tex,'\div','{&#247;}');

      tex = strrep(tex,'\sum','{&#8721;}');
      tex = strrep(tex,'\ast','{&#8727;}');
      tex = strrep(tex,'\sqrt','{&#8730;}');
      tex = strrep(tex,'\angle','{&#8736;}');
      tex = strrep(tex,'\wedge','{&#8743;}');
      tex = strrep(tex,'\land','{&#8743;}');
      tex = strrep(tex,'\vee','{&#8744;}');
      tex = strrep(tex,'\lor','{&#8744;}');
      tex = strrep(tex,'\cap','{&#8745;}');
      tex = strrep(tex,'\cup','{&#8746;}');
      tex = strrep(tex,'\int','{&#8747;}');
      %&there4;&#8756;
      tex = strrep(tex,'\sim','{&#8764;}');

      tex = strrep(tex,'\forall','{&#8704;}');
      tex = strrep(tex,'\partial','{&#8706;}');
      tex = strrep(tex,'\exists','{&#8707;}');
      tex = strrep(tex,'\emptyset','{&#8709;}');
      tex = strrep(tex,'\nabla','{&#8711;}');
      tex = strrep(tex,'\in','{&#8712;}');
      tex = strrep(tex,'\notin','{&#8713;}');
      tex = strrep(tex,'\ni','{&#8715;}');
      tex = strrep(tex,'\prod','{&#8719;}');

      tex = strrep(tex,'\cong','{&#8773;}');
      tex = strrep(tex,'\approx','{&#8776;}');
      tex = strrep(tex,'\neq','{&#8800;}');
      tex = strrep(tex,'\equiv','{&#8801;}');
      tex = strrep(tex,'\leq','{&#8804;}');
      tex = strrep(tex,'\geq','{&#8805;}');
      tex = strrep(tex,'\subset','{&#8834;}');
      tex = strrep(tex,'\supset','{&#8835;}');
      %&nsub;&#8836;
      tex = strrep(tex,'\subseteq','{&#8838;}');
      tex = strrep(tex,'\supseteq','{&#8839;}');
      tex = strrep(tex,'\oplus','{&#8853;}');
      tex = strrep(tex,'\otimes','{&#8855;}');
      tex = strrep(tex,'\bot','{&#8869;}');
      tex = strrep(tex,'\cdot','{&#8901;}');
      tex = strrep(tex,'\bullet','{&#8226;}');
      tex = strrep(tex,'\ldots','{&#8230;}');
      tex = strrep(tex,'\prime','{&#8242;}');
      % &#8243; double prime
      % &#8254; oline


      tex = strrep(tex,'\\','{&#92;}');
      tex = strrep(tex,'\{','{&#123;}');
      tex = strrep(tex,'\}','{&#125;}');
      tex = strrep(tex,'\_','{&#95;}');
      tex = strrep(tex,'\^','{&#94;}');

      tex = latex2svg(tex);
  else
      tex = sprintf('<tspan>%s</tspan>', tex);
  end
  if isempty(tex)
      return;
  end
  fprintf(fid,'  <g transform = "translate(%0.3f,%0.3f)">\n', x + dx, y + dy);
  fprintf(fid,'    <g transform = "rotate(%0.3f)">\n',-angle);
  fprintf(fid,'      <text x = "%0.3f" y = "%s" font-family = "%s" text-anchor = "%s" dominant-baseline = "%s" font-size = "%0.3fpt"%s%s fill = "%s">', 0, y_offset, textfontname, anchor, balign, textfontsize, fweight, fangle, font_color);
  % tex = strrep(tex, '<tspan>', '');
  % tex = strrep(tex, '</tspan>', '');
  if ~strncmp(tex,'<tspan>',7)
    fprintf(fid,'<tspan>%s</tspan>',tex);
  else
    fprintf(fid,'%s',tex);
  end
  if FIG2SVG_globals.debugModeOn
  % useful for debugging:
    fprintf('balign = "%s" y_offset = "%s" angle = "%0.3f" type = "%s" valign = "%s" tex = "%s"\n',balign, y_offset, angle, get(id,'Type'), valign, tex);
  end
  fprintf(fid,'</text>\n');
  fprintf(fid,'    </g>\n');
  fprintf(fid,'  </g>\n');
end

%%%%%%%%%%%%%%%%%%%%%%%%%%%%%%%%%%%%%%%%%%%%%%%%%%%%%%%%%%%%%%%%%%%%%%%%%%%%%%%%%%%%%%%%%%%%%%%%%%%%%%%%%%%%%%%%%%%%%%%%%%%%%%%%%%%%%%%%%%%%%%%%%
% converts LATEX strings into SVG strings
function returnvalue = latex2svg(StringText)
  returnvalue = StringText;
  try
  if ~isempty(StringText)
      bracket = find(StringText == '{' | StringText == '}');
      bracketCounter = zeros(1,length(StringText));
      bracketCounter(StringText == '{') = 1;
      bracketCounter(StringText == '}') = -1;
      bracketCounter = cumsum(bracketCounter);
      if bracketCounter(end) ~= 0
          fprintf(['Warning: Number of open and closed braces is not equal. Latex string ''' StringText ''' will not be converted.\n']);
      elseif any(bracketCounter < 0)
          fprintf(['Warning: Found a closed brace without a previously opened brace. Latex string ''' StringText ''' will not be converted.\n']);
      else
          if isempty(bracket)
              if any(StringText == '^' | StringText == '_' | StringText == '\' )
                  returnvalue = ['<tspan>' singleLatex2svg(StringText) '</tspan>'];
                  % Clean up empty tspan elements
                  % More could be done here, but with huge effort to make it
                  % match all special cases.
                  returnvalue = strrep(returnvalue, '></tspan><tspan>', '>');
              else
                  returnvalue = ['<tspan>' StringText '</tspan>'];
              end
          else
              returnvalue = '<tspan>';
              lastValidCharacter = 1;
              for i = 1:length(bracket)
                  lastValidCharacterOffset = 1;
                  if StringText(bracket(i)) == '{'
                      % Found '{'
                      removeCharacters = 1;
                      localOffset = 0;
                      if (bracket(i) > 1)
                          if StringText(bracket(i) - 1) == '_'
                              baselineShift = 'sub';
                              localFontSize = '65%%';
                              localOffset = -1;
                              removeCharacters = 2;
                          elseif StringText(bracket(i) - 1) == '^'
                              baselineShift = 'super';
                              localFontSize = '65%%';
                              localOffset = 1;
                              removeCharacters = 2;
                          end
                      end
                      returnvalue = [returnvalue singleLatex2svg(StringText(lastValidCharacter:bracket(i) - removeCharacters)) '<tspan'];
                      if localOffset ~= 0
                          returnvalue = [returnvalue ' style = "baseline-shift:' baselineShift ';font-size:' localFontSize '"'];
                      end
                      returnvalue = [returnvalue '>'];
                  else
                      % Found '}'
                      returnvalue = [returnvalue singleLatex2svg(StringText(lastValidCharacter:bracket(i) - 1)) '</tspan>'];
                  end
                  lastValidCharacter = bracket(i) + lastValidCharacterOffset;
              end
              if lastValidCharacter <= length(StringText)
                  returnvalue = [returnvalue singleLatex2svg(StringText(lastValidCharacter:end))];
              end
              returnvalue = [returnvalue '</tspan>'];
              % Clean up empty tspan elements
              % More could be done here, but with huge effort to make it
              % match all special cases.
              returnvalue = strrep(returnvalue, '></tspan><tspan>', '>');
              returnvalue = strrep(returnvalue, '>>', '>');
          end
      end
  end
  catch ME
      errStr = ME.identifier;
      if isempty(errStr)
          errStr = ME.message;
      end
      fprintf(['Warning: Error ''' errStr ''' occurred during conversion. Latex string ''' StringText ''' will not be converted.\n']);
  end
end

function StringText = singleLatex2svg(StringText)
  index = find(StringText == '_' | StringText == '^');
  if ~isempty(index)
      if index(end) == length(StringText)
          % Remove orphan '_' or '^'
          index = index(1:end-1);
      end
      for i = length(index):-1:1
          if StringText(index(i)) == '_'
              StringText = [StringText(1:index(i)-1) ...
                      '<tspan style = "baseline-shift:sub;font-size:65%%">' ...
                      StringText(index(i)+1) ...
                      '</tspan>' ...
                      StringText(index(i)+2:end)];
          else
              StringText = [StringText(1:index(i)-1) ...
                      '<tspan style = "baseline-shift:super;font-size:65%%">' ...
                      StringText(index(i)+1) ...
                      '</tspan>' ...
                      StringText(index(i)+2:end)];
          end
      end
  end
  if ~isempty(strfind(StringText, '\bf'))
      StringText = strrep(StringText, '\bf', '</tspan><tspan font-weight = "bold">');
  end
  if ~isempty(strfind(StringText, '\it'))
      StringText = strrep(StringText, '\it', '</tspan><tspan font-style = "italic">');
  end
  if ~isempty(strfind(StringText, '\sl'))
      StringText = strrep(StringText, '\sl', '</tspan><tspan font-style = "oblique">');
  end
  if ~isempty(strfind(StringText, '\rm'))
      StringText = strrep(StringText, '\rm', '</tspan><tspan font-style = "normal" font-weight = "normal">');
  end
end

%%%%%%%%%%%%%%%%%%%%%%%%%%%%%%%%%%%%%%%%%%%%%%%%%%%%%%%%%%%%%%%%%%%%%%%%%%%%%%%%%%%%%%%%%%%%%%%%%%%%%%%%%%%%%%%%%%%%%%%%%%%%%%%%%%%%%%%%%%%%%%%%%
function name = searchcolor(~,value)
  if ischar(value)
      name = value;
  else
      name = sprintf('#%02x%02x%02x',fix(value(1)*255),fix(value(2)*255),fix(value(3)*255));
  end
end

%%%%%%%%%%%%%%%%%%%%%%%%%%%%%%%%%%%%%%%%%%%%%%%%%%%%%%%%%%%%%%%%%%%%%%%%%%%%%%%%%%%%%%%%%%%%%%%%%%%%%%%%%%%%%%%%%%%%%%%%%%%%%%%%%%%%%%%%%%%%%%%%%
function rvalue = convertunit(value, from, to, parentheight)
  global FIG2SVG_globals
  % From SVG 1.1. Specification:
  % "1pt" equals "1.25px" (and therefore 1.25 user units)
  % "1pc" equals "15px" (and therefore 15 user units)
  % "1mm" would be "3.543307px" (3.543307 user units)
  % "1cm" equals "35.43307px" (and therefore 35.43307 user units)
  % "1in" equals "90px" (and therefore 90 user units)
  % Modification by Jonathon Harding:
  % MATLAB however, assumes a variable number of pixels per inch, and
  % assuming that the pixels match is dangerous.
  if nargin < 4
      parentheight = 1.25;    % Default
  end
  switch lower(from)  % convert from input unit to points
      case 'pixels', rvalue = value * 72/FIG2SVG_globals.ScreenPixelsPerInch/FIG2SVG_globals.resolutionScaling;
      case 'points', rvalue = value;
      case 'centimeters', rvalue = value / 2.54*72;
      case 'inches', rvalue = value * 72; % 72 points = 1 inch
      case 'normalized', rvalue = value * (parentheight * 0.8);
      otherwise, error(['Unknown unit ' from '.']);
  end
  switch lower(to)    % convert from points to specified unit
      case 'pixels', rvalue = rvalue * FIG2SVG_globals.ScreenPixelsPerInch/72;
      case 'points'   % do nothing
      case 'centimeters', rvalue = rvalue * 2.54 / 72;
      case 'inches', rvalue = rvalue / 72;    % 72 points = 1 inch
      case 'normalized', rvalue = value / (parentheight * 0.8);
      otherwise, error(['Unknown unit ' to '.']);
  end
end

%%%%%%%%%%%%%%%%%%%%%%%%%%%%%%%%%%%%%%%%%%%%%%%%%%%%%%%%%%%%%%%%%%%%%%%%%%%%%%%%%%%%%%%%%%%%%%%%%%%%%%%%%%%%%%%%%%%%%%%%%%%%%%%%%%%%%%%%%%%%%%%%%
function strString = addBackSlash( strSlash)
  % adds a backslash at the last position of the string (if not already there)
  if ( strSlash(end) ~= filesep)
      strString = [ strSlash filesep];
  else
      strString = strSlash;
  end
end

%%%%%%%%%%%%%%%%%%%%%%%%%%%%%%%%%%%%%%%%%%%%%%%%%%%%%%%%%%%%%%%%%%%%%%%%%%%%%%%%%%%%%%%%%%%%%%%%%%%%%%%%%%%%%%%%%%%%%%%%%%%%%%%%%%%%%%%%%%%%%%%%%
function strExt = getFileExtension( strFileName)
  % returns the file extension of a filename
  [~, ~, strExt] = fileparts( strFileName);
end

%%%%%%%%%%%%%%%%%%%%%%%%%%%%%%%%%%%%%%%%%%%%%%%%%%%%%%%%%%%%%%%%%%%%%%%%%%%%%%%%%%%%%%%%%%%%%%%%%%%%%%%%%%%%%%%%%%%%%%%%%%%%%%%%%%%%%%%%%%%%%%%%%
function StringText = convertString(StringText)
  if iscell(StringText) % Octave stores some strings in cell arrays. --Jakob Malm
      StringText = StringText{1};
  end
  if ~isempty(StringText)
      StringText = strrep(StringText,'&','&amp;');  % Do not change sequence !!
      StringText = strrep(StringText,'\\','\');
      StringText = strrep(StringText,'<','&lt;');
      StringText = strrep(StringText,'>','&gt;');
      StringText = strrep(StringText,'"','&quot;');
      % Workaround for Firefox and Inkscape
      StringText = strrep(StringText,'°','&#176;');
      %StringText = strrep(StringText,'°','&deg;');
      StringText = strrep(StringText,'±','&plusmn;');
      StringText = strrep(StringText,'µ','&#181;');
      % StringText = strrep(StringText,'µ','&micro;');
      StringText = strrep(StringText,'²','&sup2;');
      StringText = strrep(StringText,'³','&sup3;');
      StringText = strrep(StringText,'Œ','&frac14;');
      StringText = strrep(StringText,'œ','&frac12;');
      StringText = strrep(StringText,'Ÿ','&frac34;');
      StringText = strrep(StringText,'©','&copy;');
      StringText = strrep(StringText,'®','&reg;');
      if any(StringText > 190)
          StringText = strrep(StringText,'¿','&#191;');
          StringText = strrep(StringText,'À','&#192;');
          StringText = strrep(StringText,'Á','&#193;');
          StringText = strrep(StringText,'Â','&#194;');
          StringText = strrep(StringText,'Ã','&#195;');
          StringText = strrep(StringText,'Ä','&#196;');
          StringText = strrep(StringText,'Å','&#197;');
          StringText = strrep(StringText,'Æ','&#198;');
          StringText = strrep(StringText,'Ç','&#199;');
          StringText = strrep(StringText,'È','&#200;');
          StringText = strrep(StringText,'É','&#201;');
          StringText = strrep(StringText,'Ê','&#202;');
          StringText = strrep(StringText,'Ë','&#203;');
          StringText = strrep(StringText,'Ì','&#204;');
          StringText = strrep(StringText,'Í','&#205;');
          StringText = strrep(StringText,'Î','&#206;');
          StringText = strrep(StringText,'Ï','&#207;');
          StringText = strrep(StringText,'Ð','&#208;');
          StringText = strrep(StringText,'Ñ','&#209;');
          StringText = strrep(StringText,'Ò','&#210;');
          StringText = strrep(StringText,'Ó','&#211;');
          StringText = strrep(StringText,'Ô','&#212;');
          StringText = strrep(StringText,'Õ','&#213;');
          StringText = strrep(StringText,'Ö','&#214;');
          StringText = strrep(StringText,'×','&#215;');
          StringText = strrep(StringText,'Ø','&#216;');
          StringText = strrep(StringText,'Ù','&#217;');
          StringText = strrep(StringText,'Ú','&#218;');
          StringText = strrep(StringText,'Û','&#219;');
          StringText = strrep(StringText,'Ü','&#220;');
          StringText = strrep(StringText,'Ý','&#221;');
          StringText = strrep(StringText,'Þ','&#222;');
          StringText = strrep(StringText,'ß','&#223;');
          StringText = strrep(StringText,'à','&#224;');
          StringText = strrep(StringText,'á','&#225;');
          StringText = strrep(StringText,'â','&#226;');
          StringText = strrep(StringText,'ã','&#227;');
          StringText = strrep(StringText,'ä','&#228;');
          StringText = strrep(StringText,'å','&#229;');
          StringText = strrep(StringText,'æ','&#230;');
          StringText = strrep(StringText,'ç','&#231;');
          StringText = strrep(StringText,'è','&#232;');
          StringText = strrep(StringText,'é','&#233;');
          StringText = strrep(StringText,'ê','&#234;');
          StringText = strrep(StringText,'ë','&#235;');
          StringText = strrep(StringText,'ì','&#236;');
          StringText = strrep(StringText,'í','&#237;');
          StringText = strrep(StringText,'î','&#238;');
          StringText = strrep(StringText,'ï','&#239;');
          StringText = strrep(StringText,'ð','&#240;');
          StringText = strrep(StringText,'ñ','&#241;');
          StringText = strrep(StringText,'ò','&#242;');
          StringText = strrep(StringText,'ó','&#243;');
          StringText = strrep(StringText,'ô','&#244;');
          StringText = strrep(StringText,'õ','&#245;');
          StringText = strrep(StringText,'ö','&#246;');
          StringText = strrep(StringText,'÷','&#247;');
          StringText = strrep(StringText,'ø','&#248;');
          StringText = strrep(StringText,'ù','&#249;');
          StringText = strrep(StringText,'ú','&#250;');
          StringText = strrep(StringText,'û','&#251;');
          StringText = strrep(StringText,'ü','&#252;');
          StringText = strrep(StringText,'ý','&#253;');
          StringText = strrep(StringText,'þ','&#254;');
          StringText = strrep(StringText,'ÿ','&#255;');
      end
      StringText = deblank(StringText);
  end
end

%%%%%%%%%%%%%%%%%%%%%%%%%%%%%%%%%%%%%%%%%%%%%%%%%%%%%%%%%%%%%%%%%%%%%%%%%%%%%%%%%%%%%%%%%%%%%%%%%%%%%%%%%%%%%%%%%%%%%%%%%%%%%%%%%%%%%%%%%%%%%%%%%
function IdString = createId
  global FIG2SVG_globals
  IdString = ['ID' sprintf('%06d',FIG2SVG_globals.runningIdNumber)];
  FIG2SVG_globals.runningIdNumber = FIG2SVG_globals.runningIdNumber + 1;
end

%%%%%%%%%%%%%%%%%%%%%%%%%%%%%%%%%%%%%%%%%%%%%%%%%%%%%%%%%%%%%%%%%%%%%%%%%%%%%%%%%%%%%%%%%%%%%%%%%%%%%%%%%%%%%%%%%%%%%%%%%%%%%%%%%%%%%%%%%%%%%%%%%
function [projection, edges] = get_projection(ax,id)
  global FIG2SVG_globals
  if ~strcmp(get(ax,'Type'),'colorbar')
    xc = get(ax,'CameraTarget');
    phi = get(ax,'CameraViewAngle');
    vi = get(ax,'View');
    xi = get(ax,'XLim');
    yi = get(ax,'YLim');
    zi = get(ax,'ZLim');
    projection.aspect_scaling = get(ax,'DataAspectRatio');
    if isfield(FIG2SVG_globals,'BoxOn') % only when axis is on
      [xinfi, yinfi, zinfi] = AxesChildBounds(ax);
    else
      xinfi = xi;
      yinfi = yi;
      zinfi = zi;
    end
    xi(isinf(xi)) = xinfi(isinf(xi));
    yi(isinf(yi)) = yinfi(isinf(yi));
    zi(isinf(zi)) = zinfi(isinf(zi));
    if strcmp(get(ax,'XScale'),'log')
        if strcmp(get(ax,'XLimMode'),'manual') && any(get(ax,'XLim') == 0)
            % Fix illegal scalings set by the user
            % -> replace all 0 with automatic calculated values (child limits)
            xlimM = get(ax,'XLim');
            set(ax,'XLimMode','auto');
            xlimA = get(ax,'XLim');
            xlimM(xlimM == 0) = xlimA(xlimM == 0);
            set(ax,'XLimMode','manual');
            set(ax,'XLim', xlimM);
        end
        xi = log10(get(ax,'XLim'));
    end
    if strcmp(get(ax,'YScale'),'log')
        if strcmp(get(ax,'YLimMode'),'manual') && any(get(ax,'YLim') == 0)
            % Fix illegal scalings set by the user
            % -> replace all 0 with automatic calculated values (child limits)
            ylimM = get(ax,'YLim');
            set(ax,'YLimMode','auto');
            ylimA = get(ax,'YLim');
            ylimM(ylimM == 0) = ylimA(ylimM == 0);
            set(ax,'YLimMode','manual');
            set(ax,'YLim', ylimM);
        end
        yi = log10(get(ax,'YLim'));
    end
    if strcmp(get(ax,'ZScale'),'log')
        if strcmp(get(ax,'ZLimMode'),'manual') && any(get(ax,'ZLim') == 0)
            % Fix illegal scalings set by the user
            % -> replace all 0 with automatic calculated values (child limits)
            zlimM = get(ax,'ZLim');
            set(ax,'ZLimMode','auto');
            zlimA = get(ax,'ZLim');
            zlimM(zlimM == 0) = zlimA(zlimM == 0);
            set(ax,'ZLimMode','manual');
            set(ax,'ZLim', zlimM);
        end
        zi = log10(get(ax,'ZLim'));
    end
    projection.xi = xi;
    projection.yi = yi;
    projection.zi = zi;
    xc(1) = (xc(1) - xi(1))/(xi(2)-xi(1));
    xc(2) = (xc(2) - yi(1))/(yi(2)-yi(1));
    xc(3) = (xc(3) - zi(1))/(zi(2)-zi(1));
    if strcmp(get(ax,'XScale'),'log')
        x = [xi(1) xi(2) xi(1) xi(2) xi(1) xi(2) xi(1) xi(2)] - log10(projection.aspect_scaling(1));
    else
        x = [xi(1) xi(2) xi(1) xi(2) xi(1) xi(2) xi(1) xi(2)]/projection.aspect_scaling(1);
    end
    if strcmp(get(ax,'YScale'),'log')
        y = [yi(1) yi(1) yi(2) yi(2) yi(1) yi(1) yi(2) yi(2)] - log10(projection.aspect_scaling(2));
    else
        y = [yi(1) yi(1) yi(2) yi(2) yi(1) yi(1) yi(2) yi(2)]/projection.aspect_scaling(2);
    end
    if strcmp(get(ax,'ZScale'),'log')
        z = [zi(1) zi(1) zi(1) zi(1) zi(2) zi(2) zi(2) zi(2)] - log10(projection.aspect_scaling(3));
    else
        z = [zi(1) zi(1) zi(1) zi(1) zi(2) zi(2) zi(2) zi(2)]/projection.aspect_scaling(3);
    end
    if FIG2SVG_globals.octave
      if strcmp(get(ax,'Projection'),'orthographic')
        projection.A = [cos(vi(1)*pi/180) sin(vi(1)*pi/180) 0 0; -sin(vi(1)*pi/180)*sin(vi(2)*pi/180) cos(vi(1)*pi/180)*sin(vi(2)*pi/180) cos(vi(2)*pi/180) 0; sin(vi(1)*pi/180)*cos(vi(2)*pi/180) -cos(vi(1)*pi/180)*cos(vi(2)*pi/180) sin(vi(2)*pi/180) 0; 0 0 0 1];
      else
        error('Unknown projection: %s', get(ax,'Projection'))
      end
    else
        if strcmp(get(ax,'Projection'),'orthographic')
            projection.A = viewmtx(vi(1),vi(2));
        else
            projection.A = viewmtx(vi(1),vi(2),phi,xc);
        end
    end
    if (vi(1) == 0) && (mod(vi(2),90) == 0)
        projection.xyplane = true;
    else
        projection.xyplane = false;
    end
    axpos = get(ax,'Position');
    figpos = get(id,'Position');
    [m,n] = size(x);
    x4d = [x(:),y(:),z(:),ones(m*n,1)]';
    x2d = projection.A*x4d;
    x2 = zeros(m,n);
    y2 = zeros(m,n);
    z2 = zeros(m,n);
    x2(:) = x2d(1,:)./x2d(4,:);
    y2(:) = x2d(2,:)./x2d(4,:);
    projection.ax = ax;
    projection.xrange = max(x2) - min(x2);
    projection.yrange = max(y2) - min(y2);
    projection.xoffset = (max(x2) + min(x2))/2;
    projection.yoffset = (max(y2) + min(y2))/2;
    if (strcmp(get(ax,'PlotBoxAspectRatioMode'),'manual') || strcmp(get(ax,'DataAspectRatioMode'),'manual'))
          if (projection.xrange*axpos(4)*figpos(4) < projection.yrange*axpos(3)*figpos(3))
              projection.xrange = projection.yrange*axpos(3)*figpos(3)/axpos(4)/figpos(4);
          else
              projection.yrange = projection.xrange*axpos(4)*figpos(4)/axpos(3)/figpos(3);
          end
    end
    x2(:) = (x2d(1,:)./x2d(4,:) - projection.xoffset)/projection.xrange + 0.5;
    y2(:) = (x2d(2,:)./x2d(4,:) - projection.yoffset)/projection.yrange + 0.5;
    z2(:) = x2d(3,:);
    edges = [x2; y2; z2];
  else % colorbar
    xi = get(ax,'XLim');
    yi = get(ax,'YLim');
    vi = [0 90];
    [xinfi, yinfi, ~] = AxesChildBounds(ax);
    xi(isinf(xi)) = xinfi(isinf(xi));
    yi(isinf(yi)) = yinfi(isinf(yi));
    projection.xi = xi;
    projection.yi = yi;
    x = [xi(1) xi(2) xi(1) xi(2) xi(1) xi(2) xi(1) xi(2)];
    y = [yi(1) yi(1) yi(2) yi(2) yi(1) yi(1) yi(2) yi(2)];
    projection.A = eye(3);
    projection.xyplane = true;
    axpos = get(ax,'Position');
    figpos = get(id,'Position');
    [m,n] = size(x);
    x3d = [x(:),y(:),ones(m*n,1)]';
    x2d = projection.A*x3d;
    x2 = zeros(m,n);
    y2 = zeros(m,n);
    x2(:) = x2d(1,:)./x2d(3,:);
    y2(:) = x2d(2,:)./x2d(3,:);
    projection.ax = ax;
    projection.xrange = max(x2) - min(x2);
    projection.yrange = max(y2) - min(y2);
    projection.xoffset = (max(x2) + min(x2))/2;
    projection.yoffset = (max(y2) + min(y2))/2;
    x2(:) = (x2d(1,:)./x2d(3,:) - projection.xoffset)/projection.xrange + 0.5;
    y2(:) = (x2d(2,:)./x2d(3,:) - projection.yoffset)/projection.yrange + 0.5;
    edges = [x2; y2];
  end
end

%%%%%%%%%%%%%%%%%%%%%%%%%%%%%%%%%%%%%%%%%%%%%%%%%%%%%%%%%%%%%%%%%%%%%%%%%%%%%%%%%%%%%%%%%%%%%%%%%%%%%%%%%%%%%%%%%%%%%%%%%%%%%%%%%%%%%%%%%%%%%%%%%
function [x2,y2,z2] = project(x,y,z,projection)
  [m,n] = size(x);
  if strcmp(get(projection.ax,'XDir'),'reverse')
      xi = projection.xi;
      x = (1 - (x - xi(1)) / (xi(2) - xi(1))) * (xi(2) - xi(1)) + xi(1);
  end
  if strcmp(get(projection.ax,'YDir'),'reverse')
      yi = projection.yi;
      y = (1 - (y - yi(1)) / (yi(2) - yi(1))) * (yi(2) - yi(1)) + yi(1);
  end
  if strcmp(get(projection.ax,'XScale'),'log')
      x = x - log10(projection.aspect_scaling(1));
  else
      x = x/projection.aspect_scaling(1);
  end
  if strcmp(get(projection.ax,'YScale'),'log')
      y = y - log10(projection.aspect_scaling(2));
  else
      y = y/projection.aspect_scaling(2);
  end
  if strcmp(get(projection.ax,'ZScale'),'log')
      z = z - log10(projection.aspect_scaling(3));
  else
      z = z/projection.aspect_scaling(3);
  end
  x4d = [x(:), y(:), z(:), ones(m*n,1)]';
  x2d = projection.A*x4d;
  x2 = zeros(m,n); y2 = zeros(m,n); z2 = zeros(m,n);
  x2(:) = (x2d(1,:)./x2d(4,:) - projection.xoffset)/projection.xrange + 0.5;
  y2(:) = (x2d(2,:)./x2d(4,:) - projection.yoffset)/projection.yrange + 0.5;
  z2(:) = x2d(3,:);
  %x = [0 1 0 1 0 1 0 1];
  %y = [0 0 1 1 0 0 1 1];
  %z = [0 0 0 0 1 1 1 1];
end

function [f, v, fvc, fva] = surface2patch(s)
  x = get(s, 'xdata');
  y = get(s, 'ydata');
  z = get(s, 'zdata');
  c = get(s, 'cdata');
  a = get(s, 'AlphaData');
  if ~isempty(x) && ~isequal(size(x),size(z))
      x = repmat(x(:)',size(z,1),1);
  end
  if ~isempty(y) && ~isequal(size(y),size(z))
    y = repmat(y(:),1,size(z,2));
  end
  [m,n] = size(z);
  if isempty(x)
      [x,y] = meshgrid(1:n, 1:m);
  end
  [cm,cn,cp] = size(c);
  [am,an,ap] = size(a);
  %if cm == (m-1) & cn == (n-1)
  %    cmode = 'f';
  %elseif cm == m & cn == n
  %    cmode = 'v';
  %else
  %    cmode = '';
  %end
  v = [x(:) y(:) z(:)];
  q = (1:m*n-m-1)';
  q(m:m:end) = [];
  fvc = reshape(c, [cm*cn cp]);
  fva = reshape(a, [am*an ap]);
  f = [q q+m q+m+1 q+1];
end

%%%%%%%%%%%%%%%%%%%%%%%%%%%%%%%%%%%%%%%%%%%%%%%%%%%%%%%%%%%%%%%%%%%%%%%%%%%%
% Code by Jonathon Harding to detect axes child limits
function [xlims, ylims, zlims] = AxesChildBounds(ax)
  % Get all the direct children of the axes that are not also axes (i.e.
  % old style legends)
  children = findobj(ax, '-depth', 1, '-not', 'Type', 'axes');

  % Now get all children of those objects that have data we can analyze
  dataObjs = findobj(children, 'Type', 'line', ...
      '-or', 'Type', 'patch', '-or', 'Type', 'Rectangle', '-or', 'Type', 'Surface', '-or', 'Type', 'image','-or', 'Type', 'bar','-or', 'Type', 'errorbar');

  % SA: remove data that is outside of the axis limits (see below)
  for j = 1:numel(dataObjs)
    RemoveDataOffLimits(dataObjs(j),ax);
  end

  % Generate default limits if no objects are found
  xlims = [0 1];
  ylims = [0 1];
  zlims = [0 1];
  if numel(dataObjs) == 0
      return;
  end

  % Iterate through each axis one at a time
  if any(strcmpi(get(dataObjs,'Type'),'image')) || any(strcmpi(get(dataObjs,'Type'),'bar')) || any(strcmpi(get(dataObjs,'Type'),'errorbar'))
    axisData = {'XData', 'YData'};
  else
    axisData = {'XData', 'YData', 'ZData'};
  end
  for i = 1:numel(axisData)
      % Set extreme bounds that will always be overridden
      lims = [inf -inf];
      for j = 1:numel(dataObjs)
          % For each object, get the data for the appropriate axis
          data = reshape(get(dataObjs(j), axisData{i}), [], 1);
          % Remove data that is not displayed
          data(isinf(data) | isnan(data)) = [];
          % If any data remains, update the limits
          if ~isempty(data)
              lims(1) = min(lims(1), min(data));
              lims(2) = max(lims(2), max(data));
          end
      end
      % If the limits are not infinite (i.e. no data found), then update
      % the apropriate axis limits
      if ~any(isinf(lims))
          switch axisData{i}
              case 'XData'
                  xlims = lims;
              case 'YData'
                  ylims = lims;
              case 'ZData'
                  zlims = lims;
          end
      end
  end
end

%%%%%%%%%%%%%%%%%%%%%%%%%%%%%%%%%%%%%%%%%%%%%%%%%%%%%%%%%%%%%%%%%%%%%%%%%%%%
% SA: remove data that is outside axis limits (generates lighter figures and
% it is fundamental for apps such as scribus)
function RemoveDataOffLimits(dataObj,ax)
  axlim(1,:) = get(ax,'XLim');
  axlim(2,:) = get(ax,'YLim');
  axlim(3,:) = get(ax,'ZLim');
  typesData = {'XData', 'YData', 'ZData', 'CData'};
  if strcmpi(get(dataObj,'Type'),'surface')
    % Get the data
    for i = 1:numel(typesData)
      data{i} = get(dataObj, typesData{i});
      sizeData{i} = size(data{i});
    end
    % Trunk the data off the limits
    for i = 1:length(data)
      if ~strcmp(typesData{i},'CData')
        % Remove data that is not displayed
        data_i = data{i};
        [a,b] = find(data_i<axlim(i,1) | data_i>axlim(i,2));
        % a trick to discard points that we don't want to show
        if any(a>1) && any(b>1)
          data{3}(data_i<axlim(i,1) | data_i>axlim(i,2)) = nan;
          data{4}(data_i<axlim(i,1) | data_i>axlim(i,2)) = nan;
        elseif any(a>1)
          data{i}(a) = nan;
        elseif any(b>1)
          data{i}(b) = nan;
        end
      end
    end
    % Set the new data
    set(dataObj,'XData',data{1},'YData',data{2},'ZData',data{3},'CData',data{4});
  elseif strcmpi(get(dataObj,'Type'),'patch')
    % Get the data
    setDataStr = '';
    cont = 1;
    for i = 1:numel(typesData)
      try % 3D patch
        data(cont,:,:) = get(dataObj, typesData{i});
        datatype{cont} = typesData{i};
        cont = cont + 1;
      catch
        try % 2D patch
          data(cont,:) = get(dataObj, typesData{i})';
          datatype{cont} = typesData{i};
          cont = cont + 1;
        catch % different dimension for either ZData or CData
        end
      end
    end
    sizeData = size(data);
    % Trunk the data off the limits
    for i = 1:size(data,1)
      if ~strcmp(datatype{i},'CData')
        % Remove data that is not displayed
        if length(sizeData) == 2 % 2D patch
          % display('2D patch')
          if length(unique(data(end,:))) > 1 % hggroup (e.g., scatter plot)
            data(i,data(i,:)<axlim(i,1)) = nan;
            data(i,data(i,:)>axlim(i,2)) = nan;
          else % actual 2D patch
            data(i,data(i,:)<axlim(i,1)) = axlim(i,1);
            data(i,data(i,:)>axlim(i,2)) = axlim(i,2);
          end
          setDataStr = [setDataStr,'datatype{',num2str(i),'},data(',num2str(i),',:)'','];
        else % 3D patch
          % display('3D patch')
          data_i = squeeze(data(i,:,:));
          data(:,data_i<axlim(i,1) | data_i>axlim(i,2)) = nan;
          setDataStr = [setDataStr,'datatype{',num2str(i),'},squeeze(data(',num2str(i),',:,:)),'];
        end
      end
    end
    % Set the new data
    if length(sizeData) == 3 && sizeData(1) == 4
     setDataStr = [setDataStr,'datatype{4},squeeze(data(4,:,:))'];
    else
     setDataStr = setDataStr(1:end-1);
    end
    eval(['set(dataObj,',setDataStr,');']);
  elseif strcmpi(get(dataObj,'Type'),'image')
    % Get the data
    setDataStr = '';
    cont = 1;
    for i = 1:numel(typesData)
      try
        data(cont,:,:) = get(dataObj, typesData{i});
        datatype{cont} = typesData{i};
        cont = cont + 1;
      catch % different dimension for CData
        if strcmp(typesData{i},'CData')
          cdata = get(dataObj, typesData{i});
          datatype{cont} = typesData{i};
        end
      end
    end
    if any(size(cdata)) == 1 % colorbar
      cdim = find(size(cdata) == 1);
      if ~isempty(cdim)
        sqdataprev = squeeze(data);
        % Trunk the data off the limits
        for i = 1:size(data,1)
          % a trick to discard points that we don't want to show
          data(i,1,data(i,1,:)<axlim(i,1)) = axlim(i,1);
          data(i,1,data(i,1,:)>axlim(i,2)) = axlim(i,2);
          setDataStr = [setDataStr,'datatype{',num2str(i),'},squeeze(data(',num2str(i),',1,:)),'];
        end
        sqdata = squeeze(data);
        if sqdata(cdim,1) ~= sqdataprev(cdim,1) || sqdata(cdim,end) ~= sqdataprev(cdim,end)
          b1 = round(0.5+(length(cdata)-1)*(sqdata(cdim,1)-sqdataprev(cdim,1))/(sqdataprev(cdim,end)-sqdataprev(cdim,1)));
          b2 = round(0.5+(length(cdata)-1)*(sqdata(cdim,end)-sqdataprev(cdim,1))/(sqdataprev(cdim,end)-sqdataprev(cdim,1)));
          cdata = cdata(b1:b2);
        end
      end
    else % 2-D image (limited to integer values, actual axes' bounds set to integer values + [-0.5,0.5])
      % Trunk the data off the limits
      for i = 1:size(data,1)
        % a trick to discard points that we don't want to show
        data(i,1,data(i,1,:)<axlim(i,1)) = ceil(axlim(i,1))
        data(i,1,data(i,1,:)>axlim(i,2)) = floor(axlim(i,2))
        setDataStr = [setDataStr,'datatype{',num2str(i),'},squeeze(data(',num2str(i),',1,:)),'];
      end
      sqdata = squeeze(data);
      if size(cdata,1) ~= 1+ sqdata(2,end)-sqdata(2,1)
       cdata = cdata(sqdata(2,1):sqdata(2,end),:);
      end
      if size(cdata,2) ~= 1+ sqdata(1,end)-sqdata(1,1)
       cdata = cdata(:,sqdata(1,1):sqdata(1,end));
      end
    end
    % setDataStr = setDataStr(1:end-1);
    setDataStr = [setDataStr,'datatype{end},cdata'];
    eval(['set(dataObj,',setDataStr,');']);
  else % 2D regular plot
    % Get the data
    setDataStr = '';
    cont = 1;
    for i = 1:numel(typesData)
      try
        data(cont,:,:) = get(dataObj, typesData{i});
        datatype{cont} = typesData{i};
        cont = cont + 1;
      catch % different dimension for either ZData or CData
      end
    end
    % Trunk the data off the limits
    for i = 1:size(data,1)
      if ~strcmp(datatype{i},'CData')
        % display('2D regular plot')
        % a trick to discard points that we don't want to show
        if length(data(i,1,:)) == 2 % workaround for reference lines that consist of only two points. otherwise when plots are zoomed in, these lines disappear, but by doing what follows, the two points are moved to the extremes of the zoom region
          data(i,1,data(i,1,:)<axlim(i,1)) = axlim(i,1);
          data(i,1,data(i,1,:)>axlim(i,2)) = axlim(i,2);
        else
          data(:,1,data(i,1,:)<axlim(i,1) | data(i,1,:)>axlim(i,2)) = nan;
        end
        setDataStr = [setDataStr,'datatype{',num2str(i),'},squeeze(data(',num2str(i),',1,:)),'];
      end
    end
    setDataStr = setDataStr(1:end-1);
    eval(['set(dataObj,',setDataStr,');']);
  end
end
```
