## Supplemental File 1 for "A biophysical model of striatal microcircuits suggests delta/theta-rhythmically interleaved gamma and beta oscillations mediate periodicity in motor control": getLinearIndependent.m

```
function [Abasis, Abasisi, Asub]= getLinearIndependent(A,ignore_constant_shift)
% [Abasis, Abasisi, Asub]= getLinearIndependent(A,ignore_constant_shift)
%
% 
% Purpose: Takes in a matrix of column vectors and identifies
% the subset of columns that forms a linear independent basis (Abasis). 
% 
% 
% It also clusters columns in the original matrix into groups of those that
% share linear dependence (Asub). (e.g. If col2 = 2*col1, then col1 and
% col2 would be grouped together).
% 
% 
% Usage:
%   [Abasis, Abasisi, Asub]= getLinearIndependent(A)
%   [Abasis, Abasisi, Asub]= getLinearIndependent(A,ignore_constant_shift)
%
% Inputs:
%   A: Input matrix of numerics
%
% Inputs (Optional):
%   ignore_constant_shift: Flag (true/false [default]) for ignoring constant term
%   in determining independence (e.g. if col2 = 10-col1, col1 and col2 will
%   be grouped together if true; otherwise separately if false).
%
% Outputs:
%   Abasis: The subset of linearly independent vectors in A that form a
%   basis of A.
% 
%   Abasisi: Index locations of original basis vectors in A, such
%   that Abasis = A(:,Abasisi).
% 
%   Asub: Cell array with one element for each basis vector in A. Each cell
%   in Asub identifies clusters of columns in the original matrix A that
%   share linear dependence.
% 
% Examples:
% 
%     % -----------------------
%     % % % % Example 1: % % % 
%     % -----------------------
% 
%     A = [1, 9, 2, 3, 2; 2, 8, 2, 3, 4; 3, 7, 3, 4 6; 4, 6, 4, 5, 8 ; 5, 5, 5, 6, 10];
% 
%     % A =
%     %      1     9     2     3     2
%     %      2     8     2     3     4
%     %      3     7     3     4     6
%     %      4     6     4     5     8
%     %      5     5     5     6    10
%     %
%     % Note that: col2 = 10 - col1
%     %            col4 = col3 + 1
%     %            col5 = col1*2
% 
%     [Abasis, Abasisi, Asub]= getLinearIndependent(A, true)
% 
%     % -----------------------
%     % % % % Result 1: % % % 
%     % -----------------------
%     % Abasis =                           % Subset of basis vectors
%     %      1     2
%     %      2     2
%     %      3     3
%     %      4     4
%     %      5     5
%     % Abasisi =                         % Indices of basis vectors
%     %      1     3
%     % Asub =
%     %   1x2 cell array
%     %     [1x3 double]    [1x2 double]
%     % Asub{1} : [1, 2, 5]               % Subset of columns described by 1st basis vector
%     % Asub{2} : [3, 4]                  % Subset of columns described by 2nd basis vector
%     %
%     % -----------------------
%     % % % % Example 2: % % % 
%     % -----------------------
% 
%     A2 = [ [2,2,2,2,2]', A]
% 
%     % A2 =
%     %  2     1     9     2     3     2
%     %  2     2     8     2     3     4
%     %  2     3     7     3     4     6
%     %  2     4     6     4     5     8
%     %  2     5     5     5     6    10
%     % 
% 
%     [Abasis, Abasisi, Asub]= getLinearIndependent(A2, false)
% 
%     % -----------------------
%     % % % % Result 2: % % % 
%     % -----------------------
%     % Abasis =
%     %      1     2     2
%     %      2     2     2
%     %      3     3     2
%     %      4     4     2
%     %      5     5     2
%     % Abasisi =
%     %  1     2     4
%     % Asub =
%     %   1x3 cell array
%     %     [1x3 double]    [1x2 double]    [4]
%     % Asub{1} : [1, 3, 5]
%     % Asub{2} : [2, 6]
%     % Asub{3} : [4]
% 
%
% Author: David Stanley, Boston University, 2017
%
% See also: getLinearIndependentCell, rref

if nargin < 2
    ignore_constant_shift = false;
end

sz = size(A);

% Add a column of 1's to allow columns to be linear functions of
% eachother, offset by a constant
if ignore_constant_shift
    A = [ones(sz(1),1), A];
end

% Put A in reduced row-echelon form.
[R,jb] = rref(A);

% Remove the 1st column if we inserted a constant column
Nnz = length(jb);           % Number of non-zero rows in R matrix
if ignore_constant_shift
    R = R(:,2:end);     % Drop first column, corresponding to "constant" column
    R(1:Nnz,:) = R(circshift(1:Nnz,-1),:);        % Move row corresponding to "constant" column pivot point down to end; shift everything else up by one.
    jb = jb(2:end);     % Ignore the 1st column pivot point
    jb = jb - 1;        % Subtract 1 since we are dropping the 1st column
    A = A(:,2:end);     % Remove the constant column which we added artificially above.
end

% Start identifying clusters
Asub = cell(1,Nnz);
available_ind = true(1,sz(2));
for i = 1:Nnz
    curr_cluster = R(i,:);
    Asub{i} = find(abs(curr_cluster) > 0 & available_ind);
    available_ind(Asub{i}) = false;                         % Remove from set of availble columns
end

% Since we artificially added in a column of constants, if there were
% no other columns linearly dependent on a constant value, then
% Asub{end} will be empty and we can drop it.
if ignore_constant_shift
    if isempty(Asub{Nnz})
        Asub = Asub(1:end-1);
    end
end


Abasisi = zeros(1,length(Asub));
for i = 1:length(Asub)
    Abasisi(i) = Asub{i}(1);
end
Abasis = A(:,Abasisi);

end
```
