## Supplemental File 1 for "A biophysical model of striatal microcircuits suggests delta/theta-rhythmically interleaved gamma and beta oscillations mediate periodicity in motor control": getLinearIndependentCell.m

```
function [Abasis, Abasisi, Asubs] = getLinearIndependentCell(A,ignore_constant_shift)
% [Abasis, Abasisi, Asubs] = getLinearIndependentCell(A,ignore_constant_shift)
%
% 
% Purpose: Takes in a matrix or cell array and identifies a
% subset of independent columns. It also clusters subsets of columns
% that are dependent upon each other. 
% 
% 
% Details: If input is a matrix, behavior is exacly the same as
% getLinearIndependent. That is, for numeric inputs, "dependence" is
% determined based on linear dependence (e.g. If col2 = 2*col1, then col1 and
% col2 would be considered dependent. For non-numeric inputs, columns are
% automatically considered independent unless they are equivalent (see
% examples).
% 
% Usage:
%   [Abasis, Abasisi, Asubs] = getLinearIndependentCell(A)
%   [Abasis, Abasisi, Asubs] = getLinearIndependentCell(A,ignore_constant_shift)
%
% Inputs:
%   A: Input matrix or cell array. If a matrix is supplied, behavor is the
%   same as getLinearIndependent.
%
% Inputs (Optional):
%   ignore_constant_shift: Flag (true / false [default]) for ignoring a constant term
%   in determining independence (e.g. if col2 = 10-col1, col1 and col2 will
%   be grouped together if true; otherwise separately if false).
%
% Outputs:
%   Abasis: The set of linearly independent basis vectors in A
% 
%   Abasisi: Index locations of original basis vectors in A, such
%   that Abasis = A(:,Abasisi).
% 
%   Asub: Cell array with one element for each basis vector in A. Each cell
%   in Asub identifies clusters of columns in the original matrix A that
%   share linear dependence.
% 
%     % Example:
% 
%     A = {'a','b','c';'a','b','c';'d','e','f';1,2,3;1,2,5;5,3,@plot;5,3,@plot;2,4,6}';
% 
%     % A = 
%     %     'a'    'a'    'd'    [1]    [1]    [  5]    [  5]    [2]
%     %     'b'    'b'    'e'    [2]    [2]    [  3]    [  3]    [4]
%     %     'c'    'c'    'f'    [3]    [5]    @plot    @plot    [6]
% 
%     [Abasis, Abasisi, Asubs] = getLinearIndependentCell(A)
% 
%     % Results:
%     % 
%     % % % % % % % % % Abasis - the subset of basis columns % % % % % % % % %
%     % 
%     % horzcat(Abasis{:})
%     %   3x5 cell array
%     %     [1]    [1]    'a'    'd'    [  5]
%     %     [2]    [2]    'b'    'e'    [  3]
%     %     [3]    [5]    'c'    'f'    @plot
%     % 
%     % % % % Abasisi - the indices of these columns in the original matrix % % % 
%     % 
%     % Abasisi =
%     %   1x5 cell array
%     %     [4]    [5]    [1]    [3]    [6]
%     % 
%     % % % % % % % % % Asubs - subsets ofl linearly dependent columns % % % % % % % % %
%     % 
%     % Asubs =
%     %   1x5 cell array
%     %     [1x2 double]    [5]    [1x2 double]    [3]    [1x2 double]
%     % Asubs{1} : [4, 8]
%     % Asubs{2} : [5]
%     % Asubs{3} : [1, 2]
%     % Asubs{4} : [3]
%     % Asubs{5} : [6, 7]
% 
% 
% Submodules: getLinearIndependent, uniqueCellGeneralized, iscellnum
%
% Author: David Stanley, Boston University, 2017
%
% See also: getLinearIndependent, rref, unique

%%Code%%

if nargin < 2
    ignore_constant_shift = false;
end

% Make sure working with a cell array initially
if isnumeric(A); A = num2cell(A); end

% Identify all columns that are numeric, cell str, or others
sz = size(A);
numeric_cols = false(1,sz(2));
cellstr_cols = false(1,sz(2));
unknown_cols = false(1,sz(2));
for j = 1:sz(2)
    if iscellnum(A(:,j)); numeric_cols(j) = true;
    elseif iscellstr(A(:,j)); cellstr_cols(j) = true;
    else; unknown_cols(j) = true;
    end
end

% ID clustering within numeric columns. Check for any form of linear
% dependence
if any(numeric_cols)
    Anum = cell2mat(A(:,numeric_cols));
    [~,~,covaried1] = getLinearIndependent(Anum,ignore_constant_shift);
    for i = 1:length(covaried1); covaried1{i} = convert_submat_indices_to_fullmat_indices(covaried1{i},find(numeric_cols)); end
else
    covaried1 = {};
end

% ID clustering within non-numeric columns
ind = cellstr_cols | unknown_cols;              % The getLICellOnly function can handle any type of content (not just strings); it relies on the isequal() function to determine equality
if any(ind)
    Aother = A(:,ind);
    covaried2 = getLICellOnly(Aother);
    for i = 1:length(covaried2); covaried2{i} = convert_submat_indices_to_fullmat_indices(covaried2{i},find(ind)); end
else
    covaried2 = {};
end

Asubs=vertcat(covaried1(:),covaried2(:))';

Abasisi = {};
Abasis = {};
for i = 1:length(Asubs)
    Abasisi{i} = Asubs{i}(1);
    Abasis{i} = A(:,Abasisi{i});
end

    
end

% function covaried = idCovariedNumerics_Princomp(vary_numerics)
%     % This method needs some modification to get it working. I believe
%     % that using reduced row-echelon form will be simpler.
%     [coef,score,latent] = princomp(zscore(vary_numerics));
%     
%     % Identify independent principle components in vary_params
%     thresh = 0.01;
%     coef = abs(coef) > thresh;          % Threshold columns of vary_params for being governed by each principle component. Each column in coef denotes a principle component, and denotes which columns in vary_params are associated with that pc (1 for yes, 0 for no)
%     latent = abs(latent) > thresh;
%     
%     pc = find(latent);
%     
%     covaried = cell(1,length(pc));
%     for i = 1:length(pc)
%         covaried{i} = find(coef(:,i));
%     end
%     
% end

function inds_out = convert_submat_indices_to_fullmat_indices(inds_in,mapping)
    inds_out = mapping(inds_in);

end

function covaried = getLICellOnly(vary_cell)
    
    A = nestCellColumns(vary_cell);
    [C,ia,ic] = uniqueCellGeneralized(A);
    
    covaried = cell(1,length(C));
    for i = 1:length(C)
        covaried{i} = find(ic == i);
    end
end


function out = nestCellColumns(in)
    if ~ismatrix(in); error('input must be of size NxM'); end
    if ~iscell(in); error('input must be a cell'); end
    
    sz = size(in);
    for j = 1:sz(2)
        out{j} = in(:,j)';
    end
end
```
