## Supplemental File 1 for "A biophysical model of striatal microcircuits suggests delta/theta-rhythmically interleaved gamma and beta oscillations mediate periodicity in motor control": githubSync.m

```
function githubSync(dependencies, varargin)
% This function takes a cell array of url's to hithub repositories, loop through
% them and ensure they exist on the path, or clone them to your local machine.
%
% Example input:
%
% dependencies={...
% 	'https://github.com/drbenvincent/mcmc-utils-matlab',...
% 	'https://github.com/altmany/export_fig'};
%
% Written by Ben Vincent
% https://github.com/drbenvincent/github-sync-matlab

%% Process inputs
p = inputParser;
p.FunctionName = mfilename;
p.addRequired('dependencies',@iscellstr);
p.addParameter('exclude',[], @isvector);
p.addParameter('selfUpdate',[], @islogical);
p.parse(dependencies, varargin{:});

% deal with row or column inputs
if iscolumn(p.Results.dependencies)
	dependencies = p.Results.dependencies'; 
end

% Return to original path on cleanup
originalPath = cd;
returnToOrginalDir = onCleanup(@() cd(originalPath));

% Optionally exclude dependencies
if ~isempty(p.Results.exclude)
	assert(numel(dependencies)==numel(p.Results.exclude),...
		'exclude and dependencies must be same length')
	dependencies = dependencies(p.Results.exclude ~= true);
	if isempty(dependencies)
		return
	end
end

% Shall we update the updater? Recursive black-magic
if p.Results.selfUpdate
	githubSync({'https://github.com/drbenvincent/github-sync-matlab'})
end

%% Main algorithm
for url=dependencies
	cloneOrUpdateDependency(url{:});
end

end


function cloneOrUpdateDependency(url)
displayDependencyToCommandWindow(url);
repoName = getRepoNameFromUrl(url);
if ~isRepoFolderOnPath(repoName)
	addpath(fullfile(defineInstallPath(),repoName)); % TODO: check this... on first install, the folder won't exist
	cloneGitHubRepo(url, defineInstallPath());
else
	updateGitHubRepo(findPathContainingRepo(repoName),repoName);
end
end

function displayDependencyToCommandWindow(url)
disp( makeHyperlink(url, makeWeblinkCode(url)) )
end

function repoName = getRepoNameFromUrl(url)
[~,repoName] = fileparts(url);
end

function installPath = defineInstallPath()
% installPath will be the Matlab userpath (eg /Users/Username/Documents/MATLAB)
if isempty(userpath)
	userpath('reset')
end
installPath = userpath;
% Fix the trailing ":" which only sometimes appears
installPath = removeTrailingColon(installPath);
end

function str = removeTrailingColon(str)
if str(end)==':'
	str(end)='';
end
end

function onPath = isRepoFolderOnPath(repoName)
	onPath = exist(repoName,'dir')==7;
end

function cloneGitHubRepo(repoAddress, installPath)
	try
		cd(installPath)
		command = sprintf('git clone %s.git', repoAddress);
		system(command);
	catch
		error('git clone failed')
	end
end

function updateGitHubRepo(installPath,repoName)
try
	% TODO: confirm this works when the repo is private.
	cd(fullfile(installPath,repoName))
	system('git pull');
catch
	warning('Unable to update GitHub repository')
end
end

function repoPath = findPathContainingRepo(repoName)
% requested repoName was found as a folder on the path. This function
% returns the path of the folder containing that repo.
allPaths = strsplit(path, systemDelimiter());
isMatch = cell2mat(cellfun(@isTargetPath, allPaths, 'UniformOutput', false));
if sum(isMatch)==1
		repoPath = fileparts(allPaths{ find(isMatch,1) });
		return
elseif sum(isMatch)==0
	error('previous found repo on path, but failed to locate path')
elseif sum(isMatch)>1
	disp(allPaths{isMatch==1})
	error('found multiple possible folders that could correspond to target repo')
end

	function isTarget = isTargetPath(p)
		isTarget = strcmp(repoName, getFolderFromPath(p));
	end
	function foldername = getFolderFromPath(p)
		[~, foldername] = fileparts(p);
	end
end

function delimiter = systemDelimiter()
if ismac
		delimiter = ':';
	elseif ispc
		delimiter = ';';
end
end
```
