## Supplemental File 1 for "A biophysical model of striatal microcircuits suggests delta/theta-rhythmically interleaved gamma and beta oscillations mediate periodicity in motor control": README.md

# GitHub repository synchronization for Matlab
If your Matlab project uses other GitHub repositories, then you can use this function to ensure those those dependencies are installed (cloned) and updated.

## Installation

Clone this repository to somewhere relevant.

```matlab
cd('~/Documents/Matlab')
system('git clone https://github.com/drbenvincent/github-sync-matlab.git')
```

## Example use

Make sure the folder is on the matlab path
```matlab
addpath('~/Documents/Matlab/github-sync-matlab')
```

Use as follows. *Warning this will download or update the repositories to your machine.*
```matlab
dependencies={
    'https://github.com/drbenvincent/mcmc-utils-matlab',
    'https://github.com/altmany/export_fig'};
githubSync(dependencies)
```
## Optional input arguments
### Selective updating
Optionally provide a vector (same length as `dependencies`) indicating which dependencies to exclude from updates. The vector can be logical (`true` or `false`) or binary (`0` or `1`).
```matlab
githubSync(dependencies, 'exclude', [false true])
```

### Update the updater
Get any updates to this updating code ;) Uses recursive black magic.
```matlab
githubSync(dependencies, 'selfUpdate', true)
```

## Using this code in a project you deploy to others
If you want to just use this code to keep your repo's up to date for your own work then it's fairly straightforward. However, you may also want to deploy some code, to be used by other people. While this repo will help keep dependencies up to data, there is a bit of a chicken and egg problem. How does an end user first install this code?

If you just include this code (along with the licence please) then that should be fine. But if you wanted it to be in it's own repo then you can include the code below in setup code in your deployed package. It will attempt to update the repo, but it will fail the first time (due to it not being installed) and then clone it. Every subsequent time, it will just update.

```matlab
startDir = cd;
repoURL = 'https://github.com/drbenvincent/github-sync-matlab';
repoName = 'github-sync-matlab';
try
	% Attempt to pull latest verion
	cd(fullfile(defineInstallPath(),repoName))
	addpath(cd)
	system('git pull');
catch
	% If this fails, we assume repo is not present, therefore clone
	cd(defineInstallPath())
	system( sprintf('git clone %s.git', repoURL) )
end
cd(startDir)
```
