## Supplemental File 1 for "A biophysical model of striatal microcircuits suggests delta/theta-rhythmically interleaved gamma and beta oscillations mediate periodicity in motor control": isfunction.m

```
function [TF, ID] = isfunction(FUN)
% ISFUNCTION - true for valid matlab functions
%
%   TF = ISFUNCTION(FUN) returns 1 if FUN is a valid matlab function, and 0
%   otherwise. Matlab functions can be strings or function handles.
%
%   [TF, ID] = ISFUNCTION(FUN) also returns an identier ID. ID can take the
%   following values:
%      1  : FUN is a function string
%      2  : FUN is a function handle
%      0  : FUN is not a function, but no further specification
%     -1  : FUN is script
%     -2  : FUN is not a valid function m-file (e.g., a matfile)
%     -3  : FUN does not exist (as a function)
%     -4  : FUN is not a function but something else (a variable)
%
%   FUN can also be a cell array, TF and ID will then be arrays.
%
%   Examples:
%     tf = isfunction('lookfor') 
%        % tf = 1
%     [tf, id] = isfunction({@isfunction, 'sin','qrqtwrxxy',1:4, @clown.jpg})
%        % -> tf = [ 1  1  0  0  0 ]
%        %    id = [ 2  1 -2 -4 -3 ]
%
%   See also FUNCTION, SCRIPT, EXIST, 
%            ISA, WHICH, NARGIN, FUNCTION_HANDLE

% version 3.1 (feb 2014)
% (c) Jos van der Geest
%
%
% History:
% 1.0 (dec 2011) created for strings only
% 2.0 (apr 2013) accepts cell arrays
% 3.0 (feb 2014) implemented identier based on catched error
% 3.1 (feb 2014) added lots of help and documentation, inspired to post on
%                FEX by a recent Question/Answer thread

if ~iscell(FUN)
    % we use cellfun, so convert to cells
    FUN = {FUN} ;
end
ID = cellfun(@local_isfunction,FUN) ; % get the identifier for each "function"
TF = ID > 0 ; % valid matlab functions have a positive identifier

% = = = = = = = = = = = = = = = = = = = = =

function ID = local_isfunction(FUNNAME)
try    
    nargin(FUNNAME) ; % nargin errors when FUNNAME is not a function
    ID = 1  + isa(FUNNAME, 'function_handle') ; % 1 for m-file, 2 for handle
catch ME
    % catch the error of nargin
    switch (ME.identifier)        
        case 'MATLAB:nargin:isScript'
            ID = -1 ; % script
        case 'MATLAB:narginout:notValidMfile'
            ID = -2 ; % probably another type of file, or it does not exist
        case 'MATLAB:narginout:functionDoesnotExist'
            ID = -3 ; % porbably a handle, but not to a function
        case 'MATLAB:narginout:BadInput'
            ID = -4 ; % probably a variable or an array
        otherwise
            ID = 0 ; % unknown cause for error
    end
end
```
