## Supplemental File 1 for "A biophysical model of striatal microcircuits suggests delta/theta-rhythmically interleaved gamma and beta oscillations mediate periodicity in motor control": lsmooth.m

```
function ys=lsmooth(y,span)
% function that performs local smoothing (moving window average)

  if mod (span,2) == 0
    error ('smooth: span must be odd.')
  end

  ys = y;

  if span == 1
    return;
  end

  L = length(y);
  l = (span-1)/2;

  for i = 1:l
    ys(i)=mean(y(1:i+l));
  end

  % so sloooow
  % for i = l+1:L-l
  %   ys(i)=mean(y(i-l:i+l));
  % end

  % much faster way to do it for i in l+1:L-l
  if span < inf % cumsum method is always faster than conv and filt methods in Octave (it might be slower than conv and filt in Matlab for low span; see https://stackoverflow.com/questions/26981478/how-can-i-efficiently-compute-a-moving-average-of-a-vector)
    cs = cumsum(y)/span;
    ys(l+1) = cs(span);
    ys(l+2:L-l) = cs(span+1:end) - cs(1:end-span);
  elseif span < 0
    ys(l+1:L-l) = conv(y,ones(1,span),'valid')/span;
  else
    tmp = filter(ones(1,span), 1, y)/span;
    ys(l+1:L-l) = tmp(span:end);
  end

  for i = L-l+1:L
    ys(i)=mean(y(i-l:L));
  end
```
