## Supplemental File 1 for "A biophysical model of striatal microcircuits suggests delta/theta-rhythmically interleaved gamma and beta oscillations mediate periodicity in motor control": char.m

```
function s = char(tpl)
%TEMPLATE Convert a template object in a one line description string
%  S = CHAR(TPL) is a class convertor from Template to a string, used
%  in online display.
%  
%  See also DISPLAY

%  Copyright (C) 2003 Guillaume Flandin <>
%  $Revision: 1.0 $Date: 2003/05/05 22:19:51 $

s = ['Template Object: root ''',...
		tpl.root,''', ',...
		num2str(length(tpl.file)), ' files, ',...
		num2str(length(tpl.varkeys)), ' keys, ',...
		tpl.unknowns, ' unknowns.'];
```
