## Supplemental File 1 for "A biophysical model of striatal microcircuits suggests delta/theta-rhythmically interleaved gamma and beta oscillations mediate periodicity in motor control": display.m

```
function display(tpl)
%TEMPLATE Display a template object in Matlab window
%  DISPLAY(TPL) displays informations about the content of template 
%  object TPL:
%     Template Object: root '.', 2 files, 9 keys, comment unknowns.
%  root element of template files, number of template files, number of 
%  keywords defined and the way of handling unknowns tags.

%  Copyright (C) 2003 Guillaume Flandin <>
%  $Revision: 1.0 $Date: 2003/05/05 22:19:51 $

disp(' ');
disp([inputname(1),' = ']);
disp(' ');
for i=1:prod(size(tpl))
	disp([blanks(length(inputname(1))+3) char(tpl(i))]);
end
disp(' ');
```
