## Supplemental File 1 for "A biophysical model of striatal microcircuits suggests delta/theta-rhythmically interleaved gamma and beta oscillations mediate periodicity in motor control": get.m

```
function varargout = get(tpl,action,varargin)
%TEMPLATE/GET Access data stored in a Template object
%  TPL = GET(TPL,ACTION,VARARGIN)
%     ACTION 'var'
%     ACTION 'undefined'

%  Copyright (C) 2003 Guillaume Flandin <>
%  $Revision: 1.0 $Date: 2003/05/05 22:19:51 $

narginchk(2,3);

switch lower(action)
	case 'var'
		narginchk(2,3);
		if nargin == 2
			varargout{1} = tpl.varvals;
		elseif iscellstr(varargin{1})
			varargout{1} = {};
			for i=1:length(varargin{1})
				key = find(ismember(tpl.varkeys,varargin{1}{i}));
				if isempty(key)
					%error('[Template] No such variable name.');
					varargout{1}{end+1} = '';
				else
					varargout{1}{end+1} = tpl.varvals{key};
				end
			end
		elseif ischar(varargin{1})
			varargout{1} = char(get(tpl,'var',cellstr(varargin{1})));
		else
			varargout{1} = '';
		end
	case 'undefined'
		narginchk(3,3);
		tpl = loadtpl(tpl,varargin{1});
		str = get(tpl,'var',varargin{1});
		varargout{1} = {};
		
		%%%%%%%%%%%%%%%%%%%%%%%% WIH REGEXP ONLY %%%%%%%%%%%%%%%%%%%%
		% [b, e] = regexp(str,'{[^ \t\r\n}]+}');
		% for i=1:length(b)
		% 	if ~any(ismember(tpl.varkeys,str(b(i)+1:e(i)-1)))
		% 		varargout{1}{end+1} = str(b(i)+1:e(i)-1);
		%	end
		% end
		%%%%%%%%%%%%%%%%%%%%%%%%%%%%%%%%%%%%%%%%%%%%%%%%%%%%%%%%%%%%%

	otherwise
		varargout{1} = finish(get(tpl,'var',action),tpl.unknowns);
end
```
