## Supplemental File 1 for "A biophysical model of striatal microcircuits suggests delta/theta-rhythmically interleaved gamma and beta oscillations mediate periodicity in motor control": parse.m

```
function [tpl, str] = parse(tpl,target,handle,append)
%TEMPLATE/PARSE Fill in replacement fields with the class properties
%  [TPL, STR] = PARSE(TPL,TARGET,HANDLE) fills in the replacement field
%  HANDLE using previously defined variables of template TPL and store
%  it in field TARGET. HANDLE can also be a cell array of field names.
%  Output is also provided in output STR (content of TARGET).
%  [TPL, STR] = PARSE(TPL,TARGET,HANDLE,APPEND) allows to specify if
%  TARGET field is reseted before being filled or if new content is
%  appended to the previous one.

%  Copyright (C) 2003 Guillaume Flandin <>
%  $Revision: 1.0 $Date: 2003/05/05 22:19:51 $

narginchk(3,4);
if nargin == 3
	append = 0;
end

if iscellstr(handle)
	for i=1:length(handle)
		[tpl, str] = subst(tpl,handle{i});
		tpl = set(tpl,'var',target,str);
	end
elseif ischar(handle)
	[tpl, str] = subst(tpl,handle);
	if append
		tpl = set(tpl,'var',target,[get(tpl,'var',target) str]);
	else
		tpl = set(tpl,'var',target,str);
	end
else
	error('[Template] Badly formed handle.');
end
```
