## Supplemental File 1 for "A biophysical model of striatal microcircuits suggests delta/theta-rhythmically interleaved gamma and beta oscillations mediate periodicity in motor control": finish.m

```
function str = finish(str,unknowns)
%TEMPLATE/FINISH Apply given strategy to unknown fields in a string
%  STR = FINISH(STR,UNKNOWNS) applies on string STR the strategy defined
%  in UNKNOWNS to unknowns fields '{UNKNOWNS_FIELDS}'.
%  UNKNOWNS may be:
%    * 'keep' to do nothing
%    * 'remove' to remove all undefined fields
%    * 'comment' to replace undefined fields by a warning HTML comment.
%  This function uses Matlab REGEXPREP function coming with R13. If you
%  hold an older version, please comment lines 38 and 42: then you can 
%  only apply the 'keep' strategy.

%  Copyright (C) 2003 Guillaume Flandin <>
%  $Revision: 1.0 $Date: 2003/05/05 22:19:51 $

narginchk(2,2);

switch lower(unknowns)
	case 'keep'
		%- do nothing
	case 'remove'
		%%%%%%%%%%%%%%%%%%%%%%%% WIH REGEXP ONLY %%%%%%%%%%%%%%%%%%%%
		% str = regexprep(str,'{[^ \t\r\n}]+}','');
		%%%%%%%%%%%%%%%%%%%%%%%%%%%%%%%%%%%%%%%%%%%%%%%%%%%%%%%%%%%%%
	case 'comment'
		%%%%%%%%%%%%%%%%%%%%%%%% WIH REGEXP ONLY %%%%%%%%%%%%%%%%%%%%
		% str = regexprep(str,'{[^ \t\r\n}]+}','<!-- Template variable undefined -->');
		%%%%%%%%%%%%%%%%%%%%%%%%%%%%%%%%%%%%%%%%%%%%%%%%%%%%%%%%%%%%%
	otherwise
		error('[Template] Unknown action.');
end
```
