## Supplemental File 1 for "A biophysical model of striatal microcircuits suggests delta/theta-rhythmically interleaved gamma and beta oscillations mediate periodicity in motor control": loadtpl.m

```
function tpl = loadtpl(tpl,handle)
%TEMPLATE/LOADTPL Read a template from file
%  TPL = LOADTPL(TPL,HANDLE) read the template file associated with the
%  handle HANDLE in the template TPL and store it in the variable HANDLE.

%  Copyright (C) 2003 Guillaume Flandin <>
%  $Revision: 1.0 $Date: 2003/05/05 22:19:51 $

if ~isempty(get(tpl,'var',handle))
	return;
else
	ind = find(ismember(tpl.handles,handle));
	if isempty(ind)
		error('[Template] No such template handle.');
	else
		filename = tpl.file{ind};
		[fid, errmsg] = fopen(filename,'rt');
		if ~isempty(errmsg)
			error(sprintf('Cannot open template file %s.',filename));
		end
		tpl = set(tpl,'var',handle,fscanf(fid,'%c'));
		fclose(fid);
	end
end
```
