## Supplemental File 1 for "A biophysical model of striatal microcircuits suggests delta/theta-rhythmically interleaved gamma and beta oscillations mediate periodicity in motor control": subst.m

```
function [tpl, str] = subst(tpl,handle)
%TEMPLATE/SUBST Substitute a replacement field by its value
%  STR = SUBST(TPL,HANDLE) substitute all the known fields of variable HANDLE
%  in the template TPL.

%  Copyright (C) 2003 Guillaume Flandin <>
%  $Revision: 1.0 $Date: 2003/05/05 22:19:51 $

tpl = loadtpl(tpl,handle);

str = get(tpl,'var',handle);
for i=1:length(tpl.varkeys)
	str = strrep(str, strcat('{',tpl.varkeys{i},'}'), tpl.varvals{i});
end
```
