## Supplemental File 1 for "A biophysical model of striatal microcircuits suggests delta/theta-rhythmically interleaved gamma and beta oscillations mediate periodicity in motor control": set.m

```
function tpl = set(tpl,action,varargin)
%TEMPLATE/SET Edit data stored in a Template object
%  TPL = SET(TPL,ACTION,VARARGIN)
%     ACTION 'root'
%     ACTION 'unknowns'
%     ACTION 'file'
%     ACTION 'block'
%     ACTION 'var'

%  Copyright (C) 2003 Guillaume Flandin <>
%  $Revision: 1.0 $Date: 2003/05/05 22:19:51 $

narginchk(3,5);

switch lower(action)
	case 'root'
		narginchk(3,3);
		if exist(varargin{1},'dir')
			tpl.root = varargin{1};
		else
			error('[Template] No such directory.');
		end
	case 'unknowns'
		narginchk(3,3);
		if ismember(varargin{1},{'remove' 'comment' 'keep'})
			tpl.unknowns = varargin{1};
		else
			error('[Template] Unknowns: ''remove'', ''comment'' or ''keep''.');
		end
	case 'file'
		narginchk(4,4);
		if iscellstr(varargin{1})
			for i=1:length(varargin{1})
				ind = find(ismember(tpl.handles,varargin{1}{i}));
				if isempty(ind)
					tpl.handles{end+1} = varargin{1}{i};
					if strcmp(varargin{2}{i}(1),filesep) %- absolute path (Unix)
						tpl.file{end+1} = varargin{2}{i};
					else %- relative path
						tpl.file{end+1} = fullfile(tpl.root,varargin{2}{i});
					end
				else
					if strcmp(varargin{2}{i}(1),filesep) %- absolute path (Unix)
						tpl.file{ind} = varargin{2}{i};
					else %- relative path
						tpl.file{ind} = fullfile(tpl.root,varargin{2}{i});
					end
				end
			end
		elseif ischar(varargin{1})
			tpl = set(tpl,'file',cellstr(varargin{1}),cellstr(varargin{2}));
		else
			error('[Template] Badly formed handles.');
		end
	case 'block'
		narginchk(4,5);
		tpl = loadtpl(tpl,varargin{1});
		if nargin == 4
			name = varargin{2};
		else
			name = varargin{3};
		end
		str = get(tpl,'var',varargin{1});
		blk = '';
		%- look like this (keep the same number (1) of spaces between characters!)
		%  <!-- BEGIN ??? -->
		%  <!-- END ??? -->
		
		%%%%%%%%%%%%%%%%%%%%%%%%% WIH REGEXP %%%%%%%%%%%%%%%%%%%%%%%%
		% reg = ['<!--\s+BEGIN ' varargin{2} '\s+-->(.*)\n\s*<!--\s+END ' varargin{2} '\s+-->'];
		% [b, e] = regexp(str,reg,'once');
		% if ~isempty(b), blk = str(b:e); end %- should also remove BEGIN and END comments
		% str = regexprep(str,reg,['{' name '}']);
		%%%%%%%%%%%%%%%%%%%%%%%%%%%%%%%%%%%%%%%%%%%%%%%%%%%%%%%%%%%%%
		
		%%%%%%%%%%%%%%%%%%%%%% WIHTOUT REGEXP %%%%%%%%%%%%%%%%%%%%%%%
		indbegin = findstr(str,['<!-- BEGIN ' varargin{2} ' -->']);
		indend   = findstr(str,['<!-- END ' varargin{2} ' -->']);
		if ~isempty(indbegin) & ~isempty(indend)
		   blk = str(indbegin+length(['<!-- BEGIN ' varargin{2} ' -->'])+1:indend-1);
		   str = [str(1:indbegin-1) '{' name '}' str(indend+length(['<!-- END ' varargin{2} ' -->'])+1:end)];
		end
		%%%%%%%%%%%%%%%%%%%%%%%%%%%%%%%%%%%%%%%%%%%%%%%%%%%%%%%%%%%%%
		
		tpl = set(tpl,'var',varargin{2},blk);
		tpl = set(tpl,'var',varargin{1},str);
	case 'var'
		narginchk(3,4);
		if iscellstr(varargin{1})
			for i=1:length(varargin{1})
				ind = find(ismember(tpl.varkeys,varargin{1}{i}));
				if isempty(ind)
					tpl.varkeys{end+1} = varargin{1}{i};
					if nargin == 4
						tpl.varvals{end+1} = varargin{2}{i};
					else
						tpl.varvals{end+1} = '';
					end
				else
					tpl.varvals{ind} = varargin{2}{i};
				end
			end
		elseif ischar(varargin{1})
			tpl = set(tpl,'var',cellstr(varargin{1}),cellstr(varargin{2}));
		else
			error('[Template] Badly formed variable names.');
		end
	otherwise
		error('[Template] Unknown action to perform.');
end
```
