## Supplemental File 1 for "A biophysical model of striatal microcircuits suggests delta/theta-rhythmically interleaved gamma and beta oscillations mediate periodicity in motor control": template.m

```
function tpl = template(root,unknowns)
%TEMPLATE HTML Template Toolbox Constructor
%  TPL = TEMPLATE returns a template object using default values for the
%  root path of the template files ('.') and for the way of handling unknown
%  replacement fields (default is 'remove').
%  TPL = TEMPLATE(ROOT) allows to specify the root path of the template files
%  that will then be provided relative to this path.
%  TPL = TEMPLATE(ROOT,UNKNOWNS) also allows to specify the strategy to apply
%  to unkown fields. UNKNOWNS may be:
%    * 'keep' to do nothing
%    * 'remove' to remove all undefined fields
%    * 'comment' to replace undefined fields by a warning HTML comment.
%
%  The template class allows you to keep your HTML code in some external 
%  files which are completely free of Matlab code, but contain replacement 
%  fields. The class provides you with functions which can fill in the 
%  replacement fields with arbitrary strings. These strings can become very 
%  large, e.g. entire tables.
%  See the PHPLib: <http://www.sanisoft.com/phplib/manual/template.php>
%  See also GET, SET, PARSE

%  Copyright (C) 2003 Guillaume Flandin <>
%  $Revision: 1.0 $Date: 2003/05/05 22:19:51 $

narginchk(0,2);

switch nargin
	case 0
		tpl = struct('root','.',...
					 'file',{{}},...
					 'handles',{{}},...
					 'varkeys',{{}},...
					 'varvals',{{}},...
					 'unknowns','remove');
		tpl = class(tpl,'template');
	case 1
		if isa(root,'template')
			tpl = root;
		else
			tpl = template;
			tpl = set(tpl,'root',root);
		end
	case 2
		tpl = template;
		tpl = set(tpl,'root',root);
		tpl = set(tpl,'unknowns',unknowns);
end
```
