## Supplemental File 1 for "A biophysical model of striatal microcircuits suggests delta/theta-rhythmically interleaved gamma and beta oscillations mediate periodicity in motor control": Changelog

M2HTML Release 1.5 (2005/05/01):
================================

New features:
  - added 'helptocxml' fields in options to create a 'helptoc.xml' file used
    by Matlab documentation system 
  - some directories can be ignored when searching M-files (created by versioning systems for example)

Bug fixes:
  - updated list of mexfile extensions (MacIntosh and 64bits processors)
  - dealing with space characters in front of 'function xxx'

M2HTML Release 1.4 (2004/05/05):
================================

Changes:
  - a warning is printed if the HTML index file is going to overwrite an HTML M-file
  - 'load' parameter can be a MAT-file (m2html.mat) or the HTML output directory
  - added 'rootdir' and 'language' fields in options, for further use 
    (=> previously saved mat-files may appear incomplete)

New features:
  - PHP search engine available (but slow search index generation) in beta version.
    <private/doxyread.m> works as a Matlab offline search engine
  - full dependency graph output when 'graph' and 'global' options are 'on'
  - Graphical User Interface in beta version <mwizard.m>

Bug fixes:
  - corrected the checking of template files (<exist>...) (thanks Peter)
  - added a call to <entity> when writing "purpose" to escape HTML entities
  - detected a bug with <copyfile> on Matlab R13: display of a warning
  - added quotes around 'dot_exec' to handle white spaces (thanks Mattias)
  - a default 'mex' icon is displayed even if no source code is found
  - replaced strtok delimiter [9:13 32 '('] by [9:13 32 40]
  
M2HTML Release 1.3 (2003/10/26):
================================

Changes:
  - default input M-files ('mFiles') are now the M-files found in the direct
    subdirectories of the current directory (pwd).
  - default output directory ('htmlDir') is now 'doc'
  - added link to online tutorial in m2html help
  - modified <master.tpl> for an optional search engine
  - added a javascript to redirect to frame index in frame's <mfile.tpl>

New features:
  - added an optional download link for each M-file
  - added <private/doxyread.m> for a future search engine
  - added <private/doxywrite.m> (idem)
  - added <private/doxysearch.m> (idem)
  - added <search.tpl> and <doxysearch.php> in templates (idem)
  - added <pcode.png> in templates for Pre-Parsed Pseudo-code files (P-files)
  - added <mwizard.m> and <private/m2htmltoolbarimages.mat> for a future GUI front-end

Bug fixes:
  - added 't' permission when reading/writing files
  - rewrote subfunction <getmfiles> to handle <exist> behaviour
  - return a specific warning if an argument of 'mFiles' is a full path
  - handle white spaces in directories name
  
M2HTML Release 1.2 (2003/08/31):
================================

Available for download on Matlab Central (2003-10-05):
<http://www.mathworks.com/matlabcentral/fileexchange/loadFile.do?objectId=4039>
