## Supplemental File 1 for "A biophysical model of striatal microcircuits suggests delta/theta-rhythmically interleaved gamma and beta oscillations mediate periodicity in motor control": Contents.m

```
% M2HTML Toolbox - A Documentation Generator for Matlab in HTML
% Version 1.5 01-May-2005
% 
% M2HTML main functions.
%   m2html   - Documentation System for Matlab M-files in HTML.
%   mdot     - Wrapper to GraphViz's <mdot> for dependency graphs.
%   mwizard  - Graphical user interface for m2html.
%   private  - Internal functions.
%
% Template toolbox.
%   @template  - HTML template class
%
% Templates files.
%   templates/blue   - Default HTML template.
%   templates/frame  - Identical to <blue> but using frames.
%   templates/brain  - Another frames-enabled template
%
% Others.
%   Changelog, GPL, INSTALL, LICENSE, README, TODO.

%   Copyright (C) 2003-2005 Guillaume Flandin <>
%   $Revision: 1.5 $Date: 2005/05/01 16:15:30 $
```
