## Supplemental File 1 for "A biophysical model of striatal microcircuits suggests delta/theta-rhythmically interleaved gamma and beta oscillations mediate periodicity in motor control": INSTALL

 M2HTML - Documentation System for Matlab .m files in HTML
 =========================================================

 0/ Requirements:
      - Matlab 5.3 or above
	  - Operating system: any.

 1/ Download the latest m2html.zip on this website:
         <http://www.artefact.tk/software/matlab/m2html/>
 
 2/ Extract files in your Matlab Repository /home/foo/matlab/ :
       unzip m2html.zip

 3/ Add the m2html directory in your Matlab path:
       addpath /home/foo/matlab/m2html/
   
 5/ Ready to use !
       help m2html
   
 Please read the README file.
 A tutorial and a FAQ (Frequently Asked Questions) are available online at:
         <http://www.artefact.tk/software/matlab/m2html/tutorial.php>
		 <http://www.artefact.tk/software/matlab/m2html/faq.php>
   
 If you want to generate dependency graphs, you need to install GraphViz (a
 graph visualization software), free and cross-platform, from:
         <http://www.graphviz.org/>
   
 -------------------------------------------------------------------------------
 Matlab is a Registered Trademark of The Mathworks, Inc.

 Copyright (C) 2003-2005 Guillaume Flandin <>
