## Supplemental File 1 for "A biophysical model of striatal microcircuits suggests delta/theta-rhythmically interleaved gamma and beta oscillations mediate periodicity in motor control": LICENSE

 M2HTML - Documentation System for Matlab .m files in HTML
 Copyright (C) 2003-2005 Guillaume Flandin <>

 This program is free software; you can redistribute it and/or
 modify it under the terms of the GNU General Public License
 as published by the Free Software Foundation; either version 2
 of the License, or any later version.

 This program is distributed in the hope that it will be useful,
 but WITHOUT ANY WARRANTY; without even the implied warranty of
 MERCHANTABILITY or FITNESS FOR A PARTICULAR PURPOSE.  See the
 GNU General Public License for more details.

 You should have received a copy of the GNU General Public License
 along with this program; if not, write to the Free Software
 Foundation Inc, 59 Temple Pl. - Suite 330, Boston, MA 02111-1307, USA.
