## Supplemental File 1 for "A biophysical model of striatal microcircuits suggests delta/theta-rhythmically interleaved gamma and beta oscillations mediate periodicity in motor control": m2html.m

```
function m2html(varargin)
%M2HTML - Documentation Generator for Matlab M-files and Toolboxes in HTML
%  M2HTML by itself generates an HTML documentation of the Matlab M-files found
%  in the direct subdirectories of the current directory. HTML files are 
%  written in a 'doc' directory (created if necessary). All the others options
%  are set to default (in brackets in the following).
%  M2HTML('PropertyName1',PropertyValue1,'PropertyName2',PropertyValue2,...)
%  sets multiple option values. The list of option names and default values is:
%    o mFiles - Cell array of strings or character array containing the
%       list of M-files and/or directories of M-files for which an HTML
%       documentation will be built (use relative paths without backtracking).
%       Launch M2HTML one directory above the directory your wanting to
%       generate documentation for  [ <all direct subdirectories> ]
%    o htmlDir - Top level directory for generated HTML files [ 'doc' ]
%    o recursive - Process subdirectories recursively [ on | {off} ]
%    o source - Include Matlab source code in the generated documentation
%                               [ {on} | off ]
%    o download - Add a link to download each M-file separately [ on | {off} ]
%    o syntaxHighlighting - Source Code Syntax Highlighting [ {on} | off ]
%    o tabs - Replace '\t' (horizontal tab) in source code by n white space
%        characters [ 0 ... {4} ... n ]
%    o globalHypertextLinks - Hypertext links among separate Matlab 
%        directories [ on | {off} ]
%    o todo - Create a TODO list in each directory summarizing all the
%        '% TODO %' lines found in Matlab code [ on | {off}]
%    o graph - Compute a dependency graph using GraphViz [ on | {off}]
%        'dot' required, see <http://www.graphviz.org/>
%    o indexFile - Basename of the HTML index file [ 'index' ]
%    o extension - Extension of generated HTML files [ '.html' ]
%    o template - HTML template name to use [ {'blue'} | 'frame' | ... ]
%    o search - Add a PHP search engine [ on | {off}] - beta version!
%    o ignoredDir - List of directories to be ignored [ {'.svn' 'cvs'} ]
%    o save - Save current state after M-files parsing in 'm2html.mat' 
%        in directory htmlDir [ on | {off}]
%    o load - Load a previously saved '.mat' M2HTML state to generate HTML 
%        files once again with possibly other options [ <none> ]
%    o verbose - Verbose mode [ {on} | off ]
%
%  For more information, please read the M2HTML tutorial and FAQ at:
%    <http://www.artefact.tk/software/matlab/m2html/>
%
%  Examples:
%    >> m2html('mfiles','matlab', 'htmldir','doc');
%    >> m2html('mfiles',{'matlab/signal' 'matlab/image'}, 'htmldir','doc');
%    >> m2html('mfiles','matlab', 'htmldir','doc', 'recursive','on');
%    >> m2html('mfiles','mytoolbox', 'htmldir','doc', 'source','off');
%    >> m2html('mfiles','matlab', 'htmldir','doc', 'global','on');
%    >> m2html( ... , 'template','frame', 'index','menu');
%
%  See also MWIZARD, MDOT, TEMPLATE.

%  Copyright (C) 2005 Guillaume Flandin <>
%  $Revision: 1.5 $Date: 2005/05/01 16:15:30 $

%  This program is free software; you can redistribute it and/or
%  modify it under the terms of the GNU General Public License
%  as published by the Free Software Foundation; either version 2
%  of the License, or any later version.
% 
%  This program is distributed in the hope that it will be useful,
%  but WITHOUT ANY WARRANTY; without even the implied warranty of
%  MERCHANTABILITY or FITNESS FOR A PARTICULAR PURPOSE.  See the
%  GNU General Public License for more details.
% 
%  You should have received a copy of the GNU General Public License
%  along with this program; if not, write to the Free Software
%  Foundation Inc, 59 Temple Pl. - Suite 330, Boston, MA 02111-1307, USA.

%  Suggestions for improvement and fixes are always welcome, although no
%  guarantee is made whether and when they will be implemented.
%  Send requests to <>

%  For tips on how to write Matlab code, see:
%     * MATLAB Programming Style Guidelines, by R. Johnson:
%       <http://www.datatool.com/prod02.htm>
%     * For tips on creating help for your m-files 'type help.m'.
%     * Matlab documentation on M-file Programming:
%  <http://www.mathworks.com/access/helpdesk/help/techdoc/matlab_prog/ch_funh8.html>

%  This function uses the Template class so that you can fully customize 
%  the output. You can modify .tpl files in templates/blue/ or create new
%  templates in a new directory.
%  See the template class documentation for more details.
%  <http://www.artefact.tk/software/matlab/template/>

%  Latest information on M2HTML is available on the web through:
%  <http://www.artefact.tk/software/matlab/m2html/>

%  Other Matlab to HTML converters available on the web:
%  1/ mat2html.pl, J.C. Kantor, in Perl, 1995: 
%     <http://fresh.t-systems-sfr.com/unix/src/www/mat2html>
%  2/ htmltools, B. Alsberg, in Matlab, 1997:
%     <http://www.mathworks.com/matlabcentral/fileexchange/loadFile.do?objectId=175>
%  3/ mtree2html2001, H. Pohlheim, in Perl, 1996, 2001:
%     <http://www.pohlheim.com/perl_main.html#matlabdocu>
%  4/ MatlabCodeColorizer, S. Faridani, in C#, 2005:
%     <http://www.pitchup.com/matlabcolorizer/>
%  5/ Highlight, G. Flandin, in Matlab, 2003:
%     <http://www.artefact.tk/software/matlab/highlight/>
%  6/ mdoc, P. Brinkmann, in Matlab, 2003:
%     <http://www.math.uiuc.edu/~brinkman/software/mdoc/>
%  7/ Ocamaweb, Miriad Technologies, in Ocaml, 2002:
%     <http://ocamaweb.sourceforge.net/>
%  8/ Matdoc, M. Kaminsky, in Perl, 2003:
%     <http://www.mathworks.com/matlabcentral/fileexchange/loadFile.do?objectId=3498>
%  9/ Matlab itself, The Mathworks Inc, with HELPWIN, DOC and PUBLISH (R14)

%-------------------------------------------------------------------------------
%- Set up options and default parameters
%-------------------------------------------------------------------------------
t0 = clock; % for statistics
msgInvalidPair = 'Bad value for argument: ''%s''';

options = struct('verbose', 1,...
				 'mFiles', {{'.'}},...
				 'htmlDir', 'doc',...
				 'recursive', 0,...
				 'source', 1,...
				 'download',0,...
				 'syntaxHighlighting', 1,...
				 'tabs', 4,...
				 'globalHypertextLinks', 0,...
				 'graph', 0,...
				 'todo', 0,...
				 'load', 0,...
				 'save', 0,...
				 'search', 0,...
				 'helptocxml', 0,...
				 'indexFile', 'index',...
				 'extension', '.html',...
				 'template', 'blue',...
                 'rootdir', pwd,...
				 'ignoredDir', {{'.svn' 'cvs'}}, ...
                 'language', 'english');

if nargin == 1 & isstruct(varargin{1})
	paramlist = [ fieldnames(varargin{1}) ...
				  struct2cell(varargin{1}) ]';
	paramlist = { paramlist{:} };
else
	if mod(nargin,2)
		error('Invalid parameter/value pair arguments.');
	end
	paramlist = varargin;
end

optionsnames = lower(fieldnames(options));
for i=1:2:length(paramlist)
	pname = paramlist{i};
	pvalue = paramlist{i+1};
	ind = strmatch(lower(pname),optionsnames);
	if isempty(ind)
		error(['Invalid parameter: ''' pname '''.']);
	elseif length(ind) > 1
		error(['Ambiguous parameter: ''' pname '''.']);
	end
	switch(optionsnames{ind})
		case 'verbose'
			if strcmpi(pvalue,'on')
				options.verbose = 1;
			elseif strcmpi(pvalue,'off')
				options.verbose = 0;
			else
				error(sprintf(msgInvalidPair,pname));
			end
		case 'mfiles'
			if iscellstr(pvalue)
				options.mFiles = pvalue;
			elseif ischar(pvalue)
				options.mFiles = cellstr(pvalue);
			else
				error(sprintf(msgInvalidPair,pname));
			end
			options.load = 0;
		case 'htmldir'
			if ischar(pvalue)
				if isempty(pvalue),
					options.htmlDir = '.';
				else
					options.htmlDir = pvalue;
				end
			else
				error(sprintf(msgInvalidPair,pname));
			end
		case 'recursive'
			if strcmpi(pvalue,'on')
				options.recursive = 1;
			elseif strcmpi(pvalue,'off')
				options.recursive = 0;
			else
				error(sprintf(msgInvalidPair,pname));
			end
			options.load = 0;
		case 'source'
			if strcmpi(pvalue,'on')
				options.source = 1;
			elseif strcmpi(pvalue,'off')
				options.source = 0;
			else
				error(sprintf(msgInvalidPair,pname));
			end
		case 'download'
			if strcmpi(pvalue,'on')
				options.download = 1;
			elseif strcmpi(pvalue,'off')
				options.download = 0;
			else
				error(sprintf(msgInvalidPair,pname));
			end
		case 'syntaxhighlighting'
			if strcmpi(pvalue,'on')
				options.syntaxHighlighting = 1;
			elseif strcmpi(pvalue,'off')
				options.syntaxHighlighting = 0;
			else
				error(sprintf(msgInvalidPair,pname));
			end
		case 'tabs'
			if pvalue >= 0
				options.tabs = pvalue;
			else
				error(sprintf(msgInvalidPair,pname));
			end
		case 'globalhypertextlinks'
			if strcmpi(pvalue,'on')
				options.globalHypertextLinks = 1;
			elseif strcmpi(pvalue,'off')
				options.globalHypertextLinks = 0;
			else
				error(sprintf(msgInvalidPair,pname));
			end
			options.load = 0;
		case 'graph'
			if strcmpi(pvalue,'on')
				options.graph = 1;
			elseif strcmpi(pvalue,'off')
				options.graph = 0;
			else
				error(sprintf(msgInvalidPair,pname));
			end
		case 'todo'
			if strcmpi(pvalue,'on')
				options.todo = 1;
			elseif strcmpi(pvalue,'off')
				options.todo = 0;
			else
				error(sprintf(msgInvalidPair,pname));
			end
		case 'load'
			if ischar(pvalue)
                if exist(pvalue) == 7 % directory provided
                    pvalue = fullfile(pvalue,'m2html.mat');
                end         
				try
					load(pvalue);
				catch
					error(sprintf('Unable to load %s.', pvalue));
				end
				options.load = 1;
				[dummy options.template] = fileparts(options.template);
			else
				error(sprintf(msgInvalidPair,pname));
			end
		case 'save'
			if strcmpi(pvalue,'on')
				options.save = 1;
			elseif strcmpi(pvalue,'off')
				options.save = 0;
			else
				error(sprintf(msgInvalidPair,pname));
			end
		case 'search'
			if strcmpi(pvalue,'on')
				options.search = 1;
			elseif strcmpi(pvalue,'off')
				options.search = 0;
			else
				error(sprintf(msgInvalidPair,pname));
			end
		case 'helptocxml'
			if strcmpi(pvalue,'on')
				options.helptocxml = 1;
			elseif strcmpi(pvalue,'off')
				options.helptocxml = 0;
			else
				error(sprintf(msgInvalidPair,pname));
			end
		case 'indexfile'
			if ischar(pvalue)
				options.indexFile = pvalue;
			else
				error(sprintf(msgInvalidPair,pname));
			end
		case 'extension'
			if ischar(pvalue) & pvalue(1) == '.'
				options.extension = pvalue;
			else
				error(sprintf(msgInvalidPair,pname));
			end
		case 'template'
			if ischar(pvalue)
				options.template = pvalue;
			else
				error(sprintf(msgInvalidPair,pname));
			end
		case 'ignoreddir'
            if iscellstr(pvalue)
				options.ignoredDir = pvalue;
			elseif ischar(pvalue)
				options.ignoredDir = cellstr(pvalue);
			else
				error(sprintf(msgInvalidPair,pname));
			end
        case 'language'
            if ischar(pvalue)
				options.language = pvalue;
			else
				error(sprintf(msgInvalidPair,pname));
			end
		otherwise
			error(['Invalid parameter: ''' pname '''.']);
	end
end

%-------------------------------------------------------------------------------
%- Get template files location
%-------------------------------------------------------------------------------
s = fileparts(which(mfilename));
options.template = fullfile(s,'templates',options.template);
if exist(options.template) ~= 7
	error('[Template] Unknown template.');
end

%-------------------------------------------------------------------------------
%- Get list of M-files
%-------------------------------------------------------------------------------
if ~options.load
	if strcmp(options.mFiles,'.')
		d = dir(pwd); d = {d([d.isdir]).name};
		options.mFiles = {d{~ismember(d,{'.' '..' options.ignoredDir{:}})}};
	end
	mfiles = getmfiles(options.mFiles,{},options.recursive,options.ignoredDir);
	if ~length(mfiles), fprintf('Nothing to be done.\n'); return; end
	if options.verbose,
		fprintf('Found %d M-files.\n',length(mfiles));
	end
	mfiles = sort(mfiles); % sort list of M-files in dictionary order
end

%-------------------------------------------------------------------------------
%- Get list of (unique) directories and (unique) names
%-------------------------------------------------------------------------------
if ~options.load
	mdirs = {};
	names = {};
	for i=1:length(mfiles)
		[mdirs{i}, names{i}] = fileparts(mfiles{i});
		if isempty(mdirs{i}), mdirs{i} = '.'; end
	end

	mdir = unique(mdirs);
	if options.verbose,
		fprintf('Found %d unique Matlab directories.\n',length(mdir));
	end

	name = names;
	%name = unique(names); % output is sorted
	%if options.verbose,
	%	fprintf('Found %d unique Matlab files.\n',length(name));
	%end
end

%-------------------------------------------------------------------------------
%- Create output directory, if necessary
%-------------------------------------------------------------------------------
if isempty(dir(options.htmlDir))										       
	%- Create the top level output directory							       
	if options.verbose  												       
		fprintf('Creating directory %s...\n',options.htmlDir);  		       
	end 																       
	if options.htmlDir(end) == filesep, 								       
		options.htmlDir(end) = [];  									       
	end 																       
	[pathdir, namedir] = fileparts(options.htmlDir);					       
	if isempty(pathdir) 												       
		[status, msg] = mkdir(escapeblank(namedir)); 								       
	else																       
		[status, msg] = mkdir(escapeblank(pathdir), escapeblank(namedir));						       
	end 																       
	if ~status, error(msg); end 														       
end 																	       

%-------------------------------------------------------------------------------
%- Get synopsis, H1 line, script/function, subroutines, cross-references, todo
%-------------------------------------------------------------------------------
if ~options.load
	synopsis   = cell(size(mfiles));
	h1line     = cell(size(mfiles));
	subroutine = cell(size(mfiles));
	hrefs      = sparse(length(mfiles), length(mfiles));
	todo       = struct('mfile',[], 'line',[], 'comment',{{}});
	ismex      = zeros(length(mfiles), length(mexexts));
	statlist   = {};
	statinfo   = sparse(1,length(mfiles));
	kw         = cell(size(mfiles));
	freq       = cell(size(mfiles));

	for i=1:length(mfiles)
		if options.verbose
			fprintf('Processing file %s...',mfiles{i});
		end
		s = mfileparse(mfiles{i}, mdirs, names, options);
		synopsis{i}   = s.synopsis;
		h1line{i}     = s.h1line;
		subroutine{i} = s.subroutine;
		hrefs(i,:)    = s.hrefs;
		todo.mfile    = [todo.mfile repmat(i,1,length(s.todo.line))];
		todo.line     = [todo.line s.todo.line];
		todo.comment  = {todo.comment{:} s.todo.comment{:}};
		ismex(i,:)    = s.ismex;
		if options.search
			if options.verbose, fprintf('search...'); end
			[kw{i}, freq{i}] = searchindex(mfiles{i});
			statlist = union(statlist, kw{i});
		end
		if options.verbose, fprintf('\n'); end
	end
	hrefs = hrefs > 0;
	if options.search
		if options.verbose
			fprintf('Creating the search index...');
		end
		statinfo = sparse(length(statlist),length(mfiles));
		for i=1:length(mfiles)
			i1 = find(ismember(statlist, kw{i}));
			i2 = repmat(i,1,length(i1));
			if ~isempty(i1)
				statinfo(sub2ind(size(statinfo),i1,i2)) = freq{i};
			end
			if options.verbose, fprintf('.'); end
		end
		clear kw freq;
		if options.verbose, fprintf('\n'); end
	end
end

%-------------------------------------------------------------------------------
%- Save M-filenames and cross-references for further analysis
%-------------------------------------------------------------------------------
matfilesave = 'm2html.mat';

if options.save
	if options.verbose
		fprintf('Saving MAT file %s...\n',matfilesave);
	end
	save(fullfile(options.htmlDir,matfilesave), ...
		'mfiles', 'names', 'mdirs', 'name', 'mdir', 'options', ...
		'hrefs', 'synopsis', 'h1line', 'subroutine', 'todo', 'ismex', ...
		'statlist', 'statinfo');
end

%-------------------------------------------------------------------------------
%- Setup the output directories
%-------------------------------------------------------------------------------
for i=1:length(mdir)
	if exist(fullfile(options.htmlDir,mdir{i})) ~= 7
		ldir = splitpath(mdir{i});
		for j=1:length(ldir)
			if exist(fullfile(options.htmlDir,ldir{1:j})) ~= 7
				%- Create the output directory
				if options.verbose
					fprintf('Creating directory %s...\n',...
							fullfile(options.htmlDir,ldir{1:j}));
				end
				if j == 1
					[status, msg] = mkdir(escapeblank(options.htmlDir), ...
						escapeblank(ldir{1}));
				else
					[status, msg] = mkdir(escapeblank(options.htmlDir), ...
						escapeblank(fullfile(ldir{1:j})));
				end
				error(msg);
			end
		end
	end
end

%-------------------------------------------------------------------------------
%- Write the master index file
%-------------------------------------------------------------------------------
tpl_master = 'master.tpl';
tpl_master_identifier_nbyline = 4;
php_search = 'search.php';
dotbase = 'graph';

%- Create the HTML template
tpl = template(options.template,'remove');
tpl = set(tpl,'file','TPL_MASTER',tpl_master);
tpl = set(tpl,'block','TPL_MASTER','rowdir','rowdirs');
tpl = set(tpl,'block','TPL_MASTER','idrow','idrows');
tpl = set(tpl,'block','idrow','idcolumn','idcolumns');
tpl = set(tpl,'block','TPL_MASTER','search','searchs');
tpl = set(tpl,'block','TPL_MASTER','graph','graphs');

%- Open for writing the HTML master index file
curfile = fullfile(options.htmlDir,[options.indexFile options.extension]);
if options.verbose
	fprintf('Creating HTML file %s...\n',curfile);
end
fid = openfile(curfile,'w');

%- Set some template variables
tpl = set(tpl,'var','DATE',[datestr(now,8) ' ' datestr(now,1) ' ' ...
							datestr(now,13)]);
tpl = set(tpl,'var','MASTERPATH', './');
tpl = set(tpl,'var','DIRS', sprintf('%s ',mdir{:}));

%- Print list of unique directories
for i=1:length(mdir)
	tpl = set(tpl,'var','L_DIR',...
			  fullurl(mdir{i},[options.indexFile options.extension]));
	tpl = set(tpl,'var','DIR',mdir{i});
	tpl = parse(tpl,'rowdirs','rowdir',1);
end

%- Print full list of M-files (sorted by column)
[sortnames, ind] = sort(names);
m_mod = mod(length(sortnames), tpl_master_identifier_nbyline);
ind = [ind zeros(1,tpl_master_identifier_nbyline-m_mod)];
m_floor = floor(length(ind) / tpl_master_identifier_nbyline);
ind = reshape(ind,m_floor,tpl_master_identifier_nbyline)';

for i=1:prod(size(ind))
	if ind(i)
		tpl = set(tpl,'var','L_IDNAME',...
			fullurl(mdirs{ind(i)},[names{ind(i)} options.extension]));
		tpl = set(tpl,'var','T_IDNAME',mdirs{ind(i)});
		tpl = set(tpl,'var','IDNAME',names{ind(i)});
		tpl = parse(tpl,'idcolumns','idcolumn',1);
	else
		tpl = set(tpl,'var','L_IDNAME','');
		tpl = set(tpl,'var','T_IDNAME','');
		tpl = set(tpl,'var','IDNAME','');
		tpl = parse(tpl,'idcolumns','idcolumn',1);
	end
	if mod(i,tpl_master_identifier_nbyline) == 0
		tpl = parse(tpl,'idrows','idrow',1);
		tpl = set(tpl,'var','idcolumns','');
	end
end

%- Add a search form if necessary
tpl = set(tpl,'var','searchs','');
if options.search
	tpl = set(tpl,'var','PHPFILE',php_search);
	tpl = parse(tpl,'searchs','search',1);
end

%- Link to a full dependency graph, if necessary
tpl = set(tpl,'var','graphs','');
if options.graph & options.globalHypertextLinks & length(mdir) > 1
    tpl = set(tpl,'var','LGRAPH',[dotbase options.extension]);
    tpl = parse(tpl,'graphs','graph',1);
end

%- Print the template in the HTML file
tpl = parse(tpl,'OUT','TPL_MASTER');
fprintf(fid,'%s',get(tpl,'OUT'));
fclose(fid);

%-------------------------------------------------------------------------------
%- Copy template files (CSS, images, ...)
%-------------------------------------------------------------------------------
% Get list of files
d = dir(options.template);
d = {d(~[d.isdir]).name};
% Copy files
for i=1:length(d)
	[p, n, ext] = fileparts(d{i});
	if ~strcmp(ext,'.tpl') ... % do not copy .tpl files
       & ~strcmp([n ext],'Thumbs.db') % do not copy this Windows generated file
		if isempty(dir(fullfile(options.htmlDir,d{i})))
			if options.verbose
				fprintf('Copying template file %s...\n',d{i});
			end
			%- there is a bug with <copyfile> in Matlab 6.5 :
			%   http://www.mathworks.com/support/solutions/data/1-1B5JY.html
			%- and <copyfile> does not overwrite files even if newer...
			[status, errmsg] = copyfile(fullfile(options.template,d{i}),...
										options.htmlDir);
			%- If you encounter this bug, please uncomment one of the following lines
			% eval(['!cp -rf ' fullfile(options.template,d{i}) ' ' options.htmlDir]);
			% eval(['!copy ' fullfile(options.template,d{i}) ' ' options.htmlDir]);
			% status = 1;
			if ~status
				if ~isempty(errmsg)
					error(errmsg)
				else
					warning(sprintf(['<copyfile> failed to do its job...\n' ...
				'This is a known bug in Matlab 6.5 (R13).\n' ...
				'See http://www.mathworks.com/support/solutions/data/1-1B5JY.html']));
				end
			end
		end
	end
end

%-------------------------------------------------------------------------------
%- Search engine (index file and PHP script)
%-------------------------------------------------------------------------------
tpl_search = 'search.tpl';
idx_search = 'search.idx';

% TODO % improving the fill in of 'statlist' and 'statinfo'
% TODO % improving the search template file and update the CSS file

if options.search
	%- Write the search index file in output directory
	if options.verbose
		fprintf('Creating Search Index file %s...\n', idx_search);
	end
	docinfo = cell(length(mfiles),2);
	for i=1:length(mfiles)
		docinfo{i,1} = h1line{i};
		docinfo{i,2} = fullurl(mdirs{i}, [names{i} options.extension]);
	end
	doxywrite(fullfile(options.htmlDir,idx_search),statlist,statinfo,docinfo);
	
	%- Create the PHP template
	tpl = template(options.template,'remove');
	tpl = set(tpl,'file','TPL_SEARCH',tpl_search);
	
	%- Open for writing the PHP search script
	curfile = fullfile(options.htmlDir, php_search); 
	if options.verbose
		fprintf('Creating PHP script %s...\n',curfile);
	end
	fid = openfile(curfile,'w');
	
	%- Set template fields
	tpl = set(tpl,'var','INDEX',[options.indexFile options.extension]);
	tpl = set(tpl,'var','MASTERPATH','./');
	tpl = set(tpl,'var','DATE',[datestr(now,8) ' ' datestr(now,1) ' ' ...
								datestr(now,13)]);
	tpl = set(tpl,'var','IDXFILE',idx_search);
	tpl = set(tpl,'var','PHPFILE',php_search);
	
	%- Print the template in the HTML file
	tpl = parse(tpl,'OUT','TPL_SEARCH');
	fprintf(fid,'%s',get(tpl,'OUT'));
	fclose(fid);
end

%-------------------------------------------------------------------------------
%- Create <helptoc.xml> needed to display hierarchical entries in Contents panel
%-------------------------------------------------------------------------------
% See http://www.mathworks.com/access/helpdesk/help/techdoc/matlab_env/guiref16.html
% and http://www.mathworks.com/support/solutions/data/1-18U6Q.html?solution=1-18U6Q

% TODO % display directories in TOC hierarchically instead of linearly
if options.helptocxml
	curfile = fullfile(options.htmlDir, 'helptoc.xml');
	if options.verbose
		fprintf('Creating XML Table-Of-Content %s...\n',curfile);
	end
	fid = openfile(curfile,'w');
	fprintf(fid,'<?xml version=''1.0'' encoding=''ISO-8859-1'' ?>\n');
	fprintf(fid,'<!-- $Date: %s $ -->\n\n', datestr(now,31));
	fprintf(fid,'<toc version="1.0">\n\n');
	fprintf(fid,['<tocitem target="%s" ',...
		'image="$toolbox/matlab/icons/book_mat.gif">%s\n'], ...
		[options.indexFile options.extension],'Toolbox');
	for i=1:length(mdir)
		fprintf(fid,['<tocitem target="%s" ',...
			'image="$toolbox/matlab/icons/reficon.gif">%s\n'], ...
			fullfile(mdir{i}, ...
				[options.indexFile options.extension]),mdir{i});
		if options.graph
			fprintf(fid,['\t<tocitem target="%s" ',...
			'image="$toolbox/matlab/icons/simulinkicon.gif">%s</tocitem>\n'], ...
				fullfile(mdir{i},...
				[dotbase options.extension]),'Dependency Graph');
		end
		if options.todo
			if ~isempty(intersect(find(strcmp(mdir{i},mdirs)),todo.mfile))
				fprintf(fid,['\t<tocitem target="%s" ',...
				'image="$toolbox/matlab/icons/demoicon.gif">%s</tocitem>\n'], ...
					fullfile(mdir{i},...
					['todo' options.extension]),'Todo list');
			end
		end
		for j=1:length(mdirs)
			if strcmp(mdirs{j},mdir{i})
				curfile = fullfile(mdir{i},...
					[names{j} options.extension]);
				fprintf(fid,'\t<tocitem target="%s">%s</tocitem>\n', ...
					curfile,names{j});
			end
		end
		fprintf(fid,'</tocitem>\n');
	end
	fprintf(fid,'</tocitem>\n');
	fprintf(fid,'\n</toc>\n');
	fclose(fid);
end

%-------------------------------------------------------------------------------
%- Write an index for each output directory
%-------------------------------------------------------------------------------
tpl_mdir = 'mdir.tpl';
tpl_mdir_link = '<a href="%s">%s</a>';
%dotbase defined earlier

%- Create the HTML template
tpl = template(options.template,'remove');
tpl = set(tpl,'file','TPL_MDIR',tpl_mdir);
tpl = set(tpl,'block','TPL_MDIR','row-m','rows-m');
tpl = set(tpl,'block','row-m','mexfile','mex');
tpl = set(tpl,'block','TPL_MDIR','othermatlab','other');
tpl = set(tpl,'block','othermatlab','row-other','rows-other');
tpl = set(tpl,'block','TPL_MDIR','subfolder','subfold');
tpl = set(tpl,'block','subfolder','subdir','subdirs');
tpl = set(tpl,'block','TPL_MDIR','todolist','todolists');
tpl = set(tpl,'block','TPL_MDIR','graph','graphs');
tpl = set(tpl,'var','DATE',[datestr(now,8) ' ' datestr(now,1) ' ' ...
							datestr(now,13)]);

for i=1:length(mdir)
	%- Open for writing each output directory index file
	curfile = fullfile(options.htmlDir,mdir{i},...
					   [options.indexFile options.extension]);
	if options.verbose
		fprintf('Creating HTML file %s...\n',curfile);
	end
	fid = openfile(curfile,'w');

	%- Set template fields
	tpl = set(tpl,'var','INDEX',     [options.indexFile options.extension]);
	tpl = set(tpl,'var','MASTERPATH',backtomaster(mdir{i}));
	tpl = set(tpl,'var','MDIR',      mdir{i});
	
	%- Display Matlab m-files, their H1 line and their Mex status
	tpl = set(tpl,'var','rows-m','');
	for j=1:length(mdirs)
		if strcmp(mdirs{j},mdir{i})
			tpl = set(tpl,'var','L_NAME', [names{j} options.extension]);
			tpl = set(tpl,'var','NAME',   names{j});
			tpl = set(tpl,'var','H1LINE', h1line{j});
			if any(ismex(j,:))
				tpl = parse(tpl,'mex','mexfile');
			else
				tpl = set(tpl,'var','mex','');
			end
			tpl = parse(tpl,'rows-m','row-m',1);
		end
	end
	
	%- Display other Matlab-specific files (.mat,.mdl,.p)
	tpl = set(tpl,'var','other','');
	tpl = set(tpl,'var','rows-other','');
	w = what(mdir{i}); w = w(1);
	w = {w.mat{:} w.mdl{:} w.p{:}};
	for j=1:length(w)
		tpl = set(tpl,'var','OTHERFILE',w{j});
		tpl = parse(tpl,'rows-other','row-other',1);
	end
	if ~isempty(w)
		tpl = parse(tpl,'other','othermatlab');
	end
	
	%- Display subsequent directories and classes
	tpl = set(tpl,'var','subdirs','');
	tpl = set(tpl,'var','subfold','');
	d = dir(mdir{i});
	d = {d([d.isdir]).name};
	d = {d{~ismember(d,{'.' '..' options.ignoredDir{:}})}};
	for j=1:length(d)
		if ismember(fullfile(mdir{i},d{j}),mdir)
			tpl = set(tpl,'var','SUBDIRECTORY',...
				sprintf(tpl_mdir_link,...
				fullurl(d{j},[options.indexFile options.extension]),d{j}));
		else
			tpl = set(tpl,'var','SUBDIRECTORY',d{j});
		end
		tpl = parse(tpl,'subdirs','subdir',1);
	end
	if ~isempty(d)
		tpl = parse(tpl,'subfold','subfolder');
	end
	
	%- Link to the TODO list if necessary
	tpl = set(tpl,'var','todolists','');
	if options.todo
		if ~isempty(intersect(find(strcmp(mdir{i},mdirs)),todo.mfile))
			tpl = set(tpl,'var','LTODOLIST',['todo' options.extension]);
			tpl = parse(tpl,'todolists','todolist',1);
		end
	end
	
	%- Link to the dependency graph if necessary
	tpl = set(tpl,'var','graphs','');
	if options.graph
		tpl = set(tpl,'var','LGRAPH',[dotbase options.extension]);
		tpl = parse(tpl,'graphs','graph',1);
	end
	
	%- Print the template in the HTML file
	tpl = parse(tpl,'OUT','TPL_MDIR');
	fprintf(fid,'%s',get(tpl,'OUT'));
	fclose(fid);
end

%-------------------------------------------------------------------------------
%- Write a TODO list file for each output directory, if necessary
%-------------------------------------------------------------------------------
tpl_todo = 'todo.tpl';

if options.todo
	%- Create the HTML template
	tpl = template(options.template,'remove');
	tpl = set(tpl,'file','TPL_TODO',tpl_todo);
	tpl = set(tpl,'block','TPL_TODO','filelist','filelists');
	tpl = set(tpl,'block','filelist','row','rows');
	tpl = set(tpl,'var','DATE',[datestr(now,8) ' ' datestr(now,1) ' ' ...
								datestr(now,13)]);

	for i=1:length(mdir)
		mfilestodo = intersect(find(strcmp(mdir{i},mdirs)),todo.mfile);
		if ~isempty(mfilestodo)
			%- Open for writing each TODO list file
			curfile = fullfile(options.htmlDir,mdir{i},...
							   ['todo' options.extension]);
			if options.verbose
				fprintf('Creating HTML file %s...\n',curfile);
			end
			fid = openfile(curfile,'w');
			
			%- Set template fields
			tpl = set(tpl,'var','INDEX',[options.indexFile options.extension]);
			tpl = set(tpl,'var','MASTERPATH', backtomaster(mdir{i}));
			tpl = set(tpl,'var','MDIR',       mdir{i});
			tpl = set(tpl,'var','filelists',  '');
	
			for k=1:length(mfilestodo)
				tpl = set(tpl,'var','MFILE',names{mfilestodo(k)});
				tpl = set(tpl,'var','rows','');
				nbtodo = find(todo.mfile == mfilestodo(k));
				for l=1:length(nbtodo)
					tpl = set(tpl,'var','L_NBLINE',...
						[names{mfilestodo(k)} ...
							options.extension ...
							'#l' num2str(todo.line(nbtodo(l)))]);
					tpl = set(tpl,'var','NBLINE',num2str(todo.line(nbtodo(l))));
					tpl = set(tpl,'var','COMMENT',todo.comment{nbtodo(l)});
					tpl = parse(tpl,'rows','row',1);
				end
				tpl = parse(tpl,'filelists','filelist',1);
			end
	
			%- Print the template in the HTML file
			tpl = parse(tpl,'OUT','TPL_TODO');
			fprintf(fid,'%s',get(tpl,'OUT'));
			fclose(fid);
		end
	end
end

%-------------------------------------------------------------------------------
%- Create dependency graphs using GraphViz, if requested
%-------------------------------------------------------------------------------
tpl_graph = 'graph.tpl';
% You may have to modify the following line with Matlab7 (R14) to specify
% the full path to where GraphViz is installed
dot_exec  = 'dot';
%dotbase defined earlier

if options.graph
	%- Create the HTML template
	tpl = template(options.template,'remove');
	tpl = set(tpl,'file','TPL_GRAPH',tpl_graph);
	tpl = set(tpl,'var','DATE',[datestr(now,8) ' ' datestr(now,1) ' ' ...
								datestr(now,13)]);
	
    %- Create a full dependency graph for all directories if possible
    if options.globalHypertextLinks & length(mdir) > 1
        mdotfile = fullfile(options.htmlDir,[dotbase '.dot']);
        if options.verbose
			fprintf('Creating full dependency graph %s...',mdotfile);
		end
        mdot({hrefs, names, options, mdirs}, mdotfile); %mfiles
        calldot(dot_exec, mdotfile, ...
                fullfile(options.htmlDir,[dotbase '.map']), ...
                fullfile(options.htmlDir,[dotbase '.png']));
        if options.verbose, fprintf('\n'); end
        fid = openfile(fullfile(options.htmlDir, [dotbase options.extension]),'w');
        tpl = set(tpl,'var','INDEX',[options.indexFile options.extension]);
	    tpl = set(tpl,'var','MASTERPATH', './');
	    tpl = set(tpl,'var','MDIR',       'the whole toolbox');
        tpl = set(tpl,'var','GRAPH_IMG',  [dotbase '.png']);
        try % if <dot> failed...
	    	fmap = openfile(fullfile(options.htmlDir,[dotbase '.map']),'r');
	    	tpl = set(tpl,'var','GRAPH_MAP',  fscanf(fmap,'%c'));
		    fclose(fmap);
        end
		tpl = parse(tpl,'OUT','TPL_GRAPH');
		fprintf(fid,'%s', get(tpl,'OUT'));
        fclose(fid);
    end
    
    %- Create a dependency graph for each output directory
	for i=1:length(mdir)
		mdotfile = fullfile(options.htmlDir,mdir{i},[dotbase '.dot']);
		if options.verbose
			fprintf('Creating dependency graph %s...',mdotfile);
		end
		ind = find(strcmp(mdirs,mdir{i}));
		href1 = zeros(length(ind),length(hrefs));
		for j=1:length(hrefs), href1(:,j) = hrefs(ind,j); end
		href2 = zeros(length(ind));
		for j=1:length(ind), href2(j,:) = href1(j,ind); end
		mdot({href2, {names{ind}}, options}, mdotfile); %{mfiles{ind}}
        calldot(dot_exec, mdotfile, ...
                fullfile(options.htmlDir,mdir{i},[dotbase '.map']), ...
                fullfile(options.htmlDir,mdir{i},[dotbase '.png']));
		if options.verbose, fprintf('\n'); end
		fid = openfile(fullfile(options.htmlDir,mdir{i},...
			[dotbase options.extension]),'w');
		tpl = set(tpl,'var','INDEX',[options.indexFile options.extension]);
		tpl = set(tpl,'var','MASTERPATH', backtomaster(mdir{i}));
		tpl = set(tpl,'var','MDIR',       mdir{i});
		tpl = set(tpl,'var','GRAPH_IMG',  [dotbase '.png']);
        try % if <dot> failed, no '.map' file has been created
	    	fmap = openfile(fullfile(options.htmlDir,mdir{i},[dotbase '.map']),'r');
	    	tpl = set(tpl,'var','GRAPH_MAP',  fscanf(fmap,'%c'));
		    fclose(fmap);
        end
		tpl = parse(tpl,'OUT','TPL_GRAPH');
		fprintf(fid,'%s', get(tpl,'OUT'));
		fclose(fid);
	end
end

%-------------------------------------------------------------------------------
%- Write an HTML file for each M-file
%-------------------------------------------------------------------------------
%- List of Matlab keywords (output from iskeyword)
matlabKeywords = {'break', 'case', 'catch', 'continue', 'elseif', 'else', ...
				  'end', 'for', 'function', 'global', 'if', 'otherwise', ...
				  'persistent', 'return', 'switch', 'try', 'while'};
                  %'keyboard', 'pause', 'eps', 'NaN', 'Inf'

tpl_mfile = 'mfile.tpl';

tpl_mfile_code     = '<a href="%s" class="code" title="%s">%s</a>';
tpl_mfile_keyword  = '<span class="keyword">%s</span>';
tpl_mfile_comment  = '<span class="comment">%s</span>';
tpl_mfile_string   = '<span class="string">%s</span>';
tpl_mfile_aname    = '<a name="%s" href="#_subfunctions" class="code">%s</a>';
tpl_mfile_line     = '%04d %s\n';

%- Delimiters used in strtok: some of them may be useless (% " .), removed '.'
strtok_delim = sprintf(' \t\n\r(){}[]<>+-*~!|\\@&/,:;="''%%');

%- Create the HTML template
tpl = template(options.template,'remove');
tpl = set(tpl,'file','TPL_MFILE',tpl_mfile);
tpl = set(tpl,'block','TPL_MFILE','pathline','pl');
tpl = set(tpl,'block','TPL_MFILE','mexfile','mex');
tpl = set(tpl,'block','TPL_MFILE','script','scriptfile');
tpl = set(tpl,'block','TPL_MFILE','crossrefcall','crossrefcalls');
tpl = set(tpl,'block','TPL_MFILE','crossrefcalled','crossrefcalleds');
tpl = set(tpl,'block','TPL_MFILE','subfunction','subf');
tpl = set(tpl,'block','subfunction','onesubfunction','onesubf');
tpl = set(tpl,'block','TPL_MFILE','source','thesource');
tpl = set(tpl,'block','TPL_MFILE','download','downloads');
tpl = set(tpl,'var','DATE',[datestr(now,8) ' ' datestr(now,1) ' ' ...
							datestr(now,13)]);

nblinetot = 0;
for i=1:length(mdir)
	for j=1:length(mdirs)
		if strcmp(mdirs{j},mdir{i})
		
			curfile = fullfile(options.htmlDir,mdir{i},...
							   [names{j} options.extension]);
							   
			%- Copy M-file for download, if necessary
			if options.download
				if options.verbose
					fprintf('Copying M-file %s.m to %s...\n',names{j},...
						fullfile(options.htmlDir,mdir{i}));
				end
				[status, errmsg] = copyfile(mfiles{j},...
					fullfile(options.htmlDir,mdir{i}));
				error(errmsg);
			end
			
			%- Open for writing the HTML file
			if options.verbose
				fprintf('Creating HTML file %s...\n',curfile);
			end
			fid = openfile(curfile,'w');
			if strcmp(names{j},options.indexFile)
                fprintf(['Warning: HTML index file %s will be ' ...
                        'overwritten by Matlab function %s.\n'], ...
                        [options.indexFile options.extension], mfiles{j});
            end
            
			%- Open for reading the M-file
			fid2 = openfile(mfiles{j},'r');
			
			%- Set some template fields
			tpl = set(tpl,'var','INDEX', [options.indexFile options.extension]);
			tpl = set(tpl,'var','MASTERPATH',       backtomaster(mdir{i}));
			tpl = set(tpl,'var','MDIR',             mdirs{j});
			tpl = set(tpl,'var','NAME',             names{j});
			tpl = set(tpl,'var','H1LINE',           entity(h1line{j}));
			tpl = set(tpl,'var','scriptfile',       '');
			if isempty(synopsis{j})
				tpl = set(tpl,'var','SYNOPSIS',get(tpl,'var','script'));
			else
				tpl = set(tpl,'var','SYNOPSIS', synopsis{j});
			end
			s = splitpath(mdir{i});
			tpl = set(tpl,'var','pl','');
			for k=1:length(s)
				c = cell(1,k); for l=1:k, c{l} = filesep; end
				cpath = {s{1:k};c{:}}; cpath = [cpath{:}];
				if ~isempty(cpath), cpath = cpath(1:end-1); end
				if ismember(cpath,mdir)
					tpl = set(tpl,'var','LPATHDIR',[repmat('../',...
						1,length(s)-k) options.indexFile options.extension]);
				else
					tpl = set(tpl,'var','LPATHDIR','#');
				end
				tpl = set(tpl,'var','PATHDIR',s{k});
				tpl = parse(tpl,'pl','pathline',1);
			end
			
			%- Handle mex files
			tpl = set(tpl,'var','mex', '');
			samename = dir(fullfile(mdir{i},[names{j}	'.*']));
			samename = {samename.name};
			tpl = set(tpl,'var','MEXTYPE', 'mex');
			for k=1:length(samename)
				[dummy, dummy, ext] = fileparts(samename{k});
				switch ext
					case '.c'
						tpl = set(tpl,'var','MEXTYPE', 'c');
					case {'.cpp' '.c++' '.cxx' '.C'}
						tpl = set(tpl,'var','MEXTYPE', 'c++');
					case {'.for' '.f' '.FOR' '.F'}
						tpl = set(tpl,'var','MEXTYPE', 'fortran');
					otherwise
						%- Unknown mex file source
				end
			end
			[exts, platform] = mexexts;
			mexplatforms = sprintf('%s, ',platform{find(ismex(j,:))});
			if ~isempty(mexplatforms)
				tpl = set(tpl,'var','PLATFORMS', mexplatforms(1:end-2));
				tpl = parse(tpl,'mex','mexfile');
			end
			
			%- Set description template field
			descr = '';
			flagsynopcont = 0;
			flag_seealso  = 0;
			while 1
				tline = fgets(fid2);
				if ~ischar(tline), break, end
				tline = entity(fliplr(deblank(fliplr(tline))));
				%- Synopsis line
				if ~isempty(strmatch('function',tline))
					if ~isempty(strmatch('...',fliplr(deblank(tline))))
						flagsynopcont = 1;
					end
				%- H1 line and description
				elseif ~isempty(strmatch('%',tline))
					%- Hypertext links on the "See also" line
					ind = findstr(lower(tline),'see also');
					if ~isempty(ind) | flag_seealso
						%- "See also" only in files in the same directory
						indsamedir = find(strcmp(mdirs{j},mdirs));
						hrefnames = {names{indsamedir}};
						r = deblank(tline);
						flag_seealso = 1; %(r(end) == ',');
						tline = '';
						while 1
							[t,r,q] = strtok(r,sprintf(' \t\n\r.,;%%'));
							tline = [tline q];
							if isempty(t), break, end;
							ii = strcmpi(hrefnames,t);
							if any(ii)
								jj = find(ii);
								tline = [tline sprintf(tpl_mfile_code,...
									[hrefnames{jj(1)} options.extension],...
									synopsis{indsamedir(jj(1))},t)];
							else
								tline = [tline t];
							end
						end
						tline = sprintf('%s\n',tline);
					end
					descr = [descr tline(2:end)];
				elseif isempty(tline)
					if ~isempty(descr), break, end;
				else
					if flagsynopcont
						if isempty(strmatch('...',fliplr(deblank(tline))))
							flagsynopcont = 0;
						end
					else
						break;
					end
				end
			end
			tpl = set(tpl,'var','DESCRIPTION',...
				horztab(descr,options.tabs));
			
			%- Set cross-references template fields:
			%  Function called
			ind = find(hrefs(j,:) == 1);
			tpl = set(tpl,'var','crossrefcalls','');
			for k=1:length(ind)
				if strcmp(mdirs{j},mdirs{ind(k)})
					tpl = set(tpl,'var','L_NAME_CALL', ...
						[names{ind(k)} options.extension]);
				else
					tpl = set(tpl,'var','L_NAME_CALL', ...
							  fullurl(backtomaster(mdirs{j}), ...
							  		   mdirs{ind(k)}, ...
									   [names{ind(k)} options.extension]));
				end
				tpl = set(tpl,'var','SYNOP_CALL',   synopsis{ind(k)});
				tpl = set(tpl,'var','NAME_CALL',   names{ind(k)});
				tpl = set(tpl,'var','H1LINE_CALL', h1line{ind(k)});
				tpl = parse(tpl,'crossrefcalls','crossrefcall',1);
			end
			%  Callers
			ind = find(hrefs(:,j) == 1);
			tpl = set(tpl,'var','crossrefcalleds','');
			for k=1:length(ind)
				if strcmp(mdirs{j},mdirs{ind(k)})
					tpl = set(tpl,'var','L_NAME_CALLED', ...
						[names{ind(k)} options.extension]);
				else
					tpl = set(tpl,'var','L_NAME_CALLED', ...
						fullurl(backtomaster(mdirs{j}),...
							mdirs{ind(k)}, ...
							[names{ind(k)} options.extension]));
				end
				tpl = set(tpl,'var','SYNOP_CALLED',   synopsis{ind(k)});
				tpl = set(tpl,'var','NAME_CALLED',   names{ind(k)});
				tpl = set(tpl,'var','H1LINE_CALLED', h1line{ind(k)});
				tpl = parse(tpl,'crossrefcalleds','crossrefcalled',1);
			end
			
			%- Set subfunction template field
			tpl = set(tpl,'var',{'subf' 'onesubf'},{'' ''});
			if ~isempty(subroutine{j}) & options.source
				for k=1:length(subroutine{j})
					tpl = set(tpl, 'var', 'L_SUB', ['#_sub' num2str(k)]);
					tpl = set(tpl, 'var', 'SUB',   subroutine{j}{k});
					tpl = parse(tpl, 'onesubf', 'onesubfunction',1);
				end
				tpl = parse(tpl,'subf','subfunction');
			end
			subname = extractname(subroutine{j});
			
			%- Link to M-file (for download)
			tpl = set(tpl,'var','downloads','');
			if options.download
				tpl = parse(tpl,'downloads','download',1);
			end
			
			%- Display source code with cross-references
			if options.source & ~strcmpi(names{j},'contents')
				fseek(fid2,0,-1);
				it = 1;
				matlabsource = '';
				nbsubroutine = 1;
				%- Get href function names of this file
				indhrefnames = find(hrefs(j,:) == 1);
				hrefnames = {names{indhrefnames}};
				%- Loop over lines
				while 1
					tline = fgetl(fid2);
					if ~ischar(tline), break, end
					myline = '';
					splitc = splitcode(entity(tline));
					for k=1:length(splitc)
						if isempty(splitc{k})
						elseif ~isempty(strmatch('function', ...
										fliplr(deblank(fliplr(splitc{k})))))
							if isspace(splitc{k}(1))
								nbs = find(diff(isspace(splitc{k}))==-1);
								myline = [myline splitc{k}(1:nbs(1))];
								splitc{k} = splitc{k}(nbs(1)+1:end);
							end
							%- Subfunctions definition
							myline = [myline ...
								sprintf(tpl_mfile_aname,...
									['_sub' num2str(nbsubroutine-1)],splitc{k})];
							nbsubroutine = nbsubroutine + 1;
						elseif splitc{k}(1) == ''''
							myline = [myline ...
								sprintf(tpl_mfile_string,splitc{k})];
						elseif splitc{k}(1) == '%'
							myline = [myline ...
								sprintf(tpl_mfile_comment,deblank(splitc{k}))];
						elseif ~isempty(strmatch('...',splitc{k}))
							myline = [myline sprintf(tpl_mfile_keyword,'...')];
							if ~isempty(splitc{k}(4:end))
								myline = [myline ...
									sprintf(tpl_mfile_comment,splitc{k}(4:end))];
							end
						else
							%- Look for keywords
							r = splitc{k};
							while 1
								[t,r,q] = strtok(r,strtok_delim);
								myline = [myline q];
								if isempty(t), break, end;
								%- Highlight Matlab keywords &
								%  cross-references on known functions
								if options.syntaxHighlighting & ...
										any(strcmp(matlabKeywords,t))
									if strcmp('end',t)
										rr = fliplr(deblank(fliplr(r)));
										icomma = strmatch(',',rr);
										isemicolon = strmatch(';',rr);
										if ~(isempty(rr) | ~isempty([icomma isemicolon]))
											myline = [myline t];
										else
											myline = [myline sprintf(tpl_mfile_keyword,t)];
										end
									else
										myline = [myline sprintf(tpl_mfile_keyword,t)];
									end
								elseif any(strcmp(hrefnames,t))
									indt = indhrefnames(logical(strcmp(hrefnames,t)));
									flink = [t options.extension];
									ii = ismember({mdirs{indt}},mdirs{j});
									if ~any(ii)
										% take the first one...
										flink = fullurl(backtomaster(mdirs{j}),...
													 	 mdirs{indt(1)}, flink);
									else
										indt = indt(logical(ii));
									end
									myline = [myline sprintf(tpl_mfile_code,...
											  flink, synopsis{indt(1)}, t)];
								elseif any(strcmp(subname,t))
									ii = find(strcmp(subname,t));
									myline = [myline sprintf(tpl_mfile_code,...
										['#_sub' num2str(ii)],...
										['sub' subroutine{j}{ii}],t)];
								else
									myline = [myline t];
								end
							end
						end
					end
					matlabsource = [matlabsource sprintf(tpl_mfile_line,it,myline)];
					it = it + 1;
				end
                nblinetot = nblinetot + it - 1;
				tpl = set(tpl,'var','SOURCECODE',...
						  horztab(matlabsource,options.tabs));
				tpl = parse(tpl,'thesource','source');
			else
				tpl = set(tpl,'var','thesource','');
			end
			tpl = parse(tpl,'OUT','TPL_MFILE');
			fprintf(fid,'%s',get(tpl,'OUT'));
			fclose(fid2);
			fclose(fid);
		end
	end
end

%-------------------------------------------------------------------------------
%- Display Statistics
%-------------------------------------------------------------------------------
if options.verbose
    prnbline = '';
    if options.source
        prnbline = sprintf('(%d lines) ', nblinetot);
    end
    fprintf('Stats: %d M-files %sin %d directories documented in %d s.\n', ...
            length(mfiles), prnbline, length(mdir), round(etime(clock,t0)));
end

%===============================================================================
function mfiles = getmfiles(mdirs, mfiles, recursive, ignoredDir)
	%- Extract M-files from a list of directories and/or M-files

	if nargin < 4, ignoredDir = {}; end
	for i=1:length(mdirs)
		currentdir = fullfile(pwd, mdirs{i});
		if exist(currentdir) == 2 % M-file
			mfiles{end+1} = mdirs{i};
		elseif exist(currentdir) == 7 % Directory
			d = dir(fullfile(currentdir, '*.m'));
			d = {d(~[d.isdir]).name};
			for j=1:length(d)
				%- don't take care of files containing ','
				%  probably a sccs file...
				if isempty(findstr(',',d{j}))
					mfiles{end+1} = fullfile(mdirs{i}, d{j});
				end
			end
			if recursive
				d = dir(currentdir);
				d = {d([d.isdir]).name};
				d = {d{~ismember(d,{'.' '..' ignoredDir{:}})}};
				for j=1:length(d)
					mfiles = getmfiles(cellstr(fullfile(mdirs{i},d{j})), ...
									   mfiles, recursive, ignoredDir);
				end
			end
		else
			fprintf('Warning: Unprocessed file %s.\n',mdirs{i});
			if ~isempty(strmatch('/',mdirs{i})) | findstr(':',mdirs{i})
				fprintf('         Use relative paths in ''mfiles'' option\n');
			end 
		end
	end

%===============================================================================
function calldot(dotexec, mdotfile, mapfile, pngfile, opt)
	%- Draw a dependency graph in a PNG image using <dot> from GraphViz

    if nargin == 4, opt = ''; end
    try
        %- See <http://www.graphviz.org/>
        %  <dot> must be in your system path, see M2HTML FAQ:
        %  <http://www.artefact.tk/software/matlab/m2html/faq.php>

%         eval(['!"' dotexec '" ' opt ' -Tcmap -Tpng "' mdotfile ...
%               '" -o "' mapfile ... 
%               '" -o "' pngfile '"']);
        system([dotexec opt ' -Tcmap -Tpng "' mdotfile ...
              '" -o "' mapfile ... 
              '" -o "' pngfile '"']);
        % use '!' rather than 'system' for backward compability with Matlab 5.3
    catch % use of '!' prevents errors to be catched...
        fprintf('<dot> failed.');
    end
    
%===============================================================================
function s = backtomaster(mdir)
	%- Provide filesystem path to go back to the root folder

	ldir = splitpath(mdir);
	s = repmat('../',1,length(ldir));
	
%===============================================================================
function ldir = splitpath(p)
	%- Split a filesystem path into parts using filesep as separator

	ldir = {};
	p = deblank(p);
	while 1
		[t,p] = strtok(p,filesep);
		if isempty(t), break; end
		if ~strcmp(t,'.')
			ldir{end+1} = t;
		end
	end
	if isempty(ldir)
		ldir{1} = '.'; % should be removed
	end

%===============================================================================
function name = extractname(synopsis)
	%- Extract function name in a synopsis

	if ischar(synopsis), synopsis = {synopsis}; end
	name = cell(size(synopsis));
	for i=1:length(synopsis)
		ind = findstr(synopsis{i},'=');
		if isempty(ind)
			ind = findstr(synopsis{i},'function');
			s = synopsis{i}(ind(1)+8:end);
		else
			s = synopsis{i}(ind(1)+1:end);
		end
		name{i} = strtok(s,[9:13 32 40]); % white space characters and '('
	end
	if length(name) == 1, name = name{1}; end

%===============================================================================
function f = fullurl(varargin)
	%- Build full url from parts (using '/' and not filesep)
	
	f = strrep(fullfile(varargin{:}),'\','/');

%===============================================================================
function str = escapeblank(str)
	%- Escape white spaces using '\'
	
	str = deblank(fliplr(deblank(fliplr(str))));
	str = strrep(str,' ','\ ');

%===============================================================================
function str = entity(str)
	%- Escape HTML special characters
	%- See http://www.w3.org/TR/html4/charset.html#h-5.3.2
	
	str = strrep(str,'&','&amp;');
	str = strrep(str,'<','&lt;');
	str = strrep(str,'>','&gt;');
	str = strrep(str,'"','&quot;');
	
%===============================================================================
function str = horztab(str,n)
	%- For browsers, the horizontal tab character is the smallest non-zero 
	%- number of spaces necessary to line characters up along tab stops that are
	%- every 8 characters: behaviour obtained when n = 0.
	
	if n > 0
		str = strrep(str,sprintf('\t'),blanks(n));
	end
```
