## Supplemental File 1 for "A biophysical model of striatal microcircuits suggests delta/theta-rhythmically interleaved gamma and beta oscillations mediate periodicity in motor control": mdot.m

```
function mdot(mmat, dotfile,f)
%MDOT - Export a dependency graph into DOT language
%  MDOT(MMAT, DOTFILE) loads a .mat file generated by M2HTML using option
%  ('save','on') and writes an ascii file using the DOT language that can
%  be drawn using <dot> or <neato> .
%  MDOT(MMAT, DOTFILE,F) builds the graph containing M-file F and its
%  neighbors only.
%  See the following page for more details:
%  <http://www.graphviz.org/>
%
%  Example:
%    mdot('m2html.mat','m2html.dot');
%    !dot -Tps m2html.dot -o m2html.ps
%    !neato -Tps m2html.dot -o m2html.ps
%
%  See also M2HTML

%  Copyright (C) 2004 Guillaume Flandin <>
%  $Revision: 1.1 $Date: 2004/05/05 17:14:09 $

narginchk(2,3);

if ischar(mmat)
	load(mmat);
elseif iscell(mmat)
	hrefs  = mmat{1};
	names  = mmat{2};
	options = mmat{3};
	if nargin == 3, mfiles = mmat{4}; end
    mdirs = cell(size(names));
    [mdirs{:}] = deal('');
    if nargin == 2 & length(mmat) > 3, 
        mdirs = mmat{4};
    end;
else
	error('[mdot] Invalid argument: mmat.');
end

fid = fopen(dotfile,'wt');
if fid == -1, error(sprintf('[mdot] Cannot open %s.',dotfile)); end

fprintf(fid,'/* Created by mdot for Matlab */\n');
fprintf(fid,'digraph m2html {\n');

% if 'names' contains '.' then they should be surrounded by '"'

if nargin == 2
	for i=1:size(hrefs,1)
		n = find(hrefs(i,:) == 1);
		m{i} = n;
		for j=1:length(n)
			fprintf(fid,['  ' names{i} ' -> ' names{n(j)} ';\n']);
		end
	end
	%m = unique([m{:}]);
	fprintf(fid,'\n');
	for i=1:size(hrefs,1)
		fprintf(fid,['  ' names{i} ' [URL="' ...
            fullurl(mdirs{i},[names{i} options.extension]) '"];\n']);
	end
else
	i = find(strcmp(f,mfiles));
	if length(i) ~= 1
		error(sprintf('[mdot] Cannot find %s.',f));
	end
	n = find(hrefs(i,:) == 1);
	for j=1:length(n)
		fprintf(fid,['  ' names{i} ' -> ' names{n(j)} ';\n']);
	end
	m = find(hrefs(:,i) == 1);
	for j=1:length(m)
		if n(j) ~= i
			fprintf(fid,['  ' names{m(j)} ' -> ' names{i} ';\n']);
		end
	end
	n = unique([n(:)' m(:)']);
	fprintf(fid,'\n');
	for i=1:length(n)
		fprintf(fid,['  ' names{n(i)} ' [URL="' fullurl(mdirs{i}, ...
            [names{n(i)} options.extension]) '"];\n']);
	end
end

fprintf(fid,'}');

fid = fclose(fid);
if fid == -1, error(sprintf('[mdot] Cannot close %s.',dotfile)); end

%===========================================================================
function f = fullurl(varargin)
	%- Build full url from parts (using '/' and not filesep)
	
	f = strrep(fullfile(varargin{:}),'\','/');
```
