## Supplemental File 1 for "A biophysical model of striatal microcircuits suggests delta/theta-rhythmically interleaved gamma and beta oscillations mediate periodicity in motor control": mwizard2.m

```
function mwizard(file)
%MWIZARD - M2HTML Graphical User Interface
%  MWIZARD launches a Matlab GUI front-end to edit parameters
%  that are then used by M2HTML to generate HTML documentation.
%  MWIZARD(FILE) allows to specify a mat-file FILE from which
%  default parameters are extracted and can be updated.  
%
%  For more information, please read the M2HTML tutorial and FAQ at:
%    <http://www.artefact.tk/software/matlab/m2html/>
%
%  See also M2HTML

%  Copyright (C) 2004 Guillaume Flandin <>
%  $Revision: 0.5 $Date: 2004/05/24 20:12:17 $

%  This program is free software; you can redistribute it and/or
%  modify it under the terms of the GNU General Public License
%  as published by the Free Software Foundation; either version 2
%  of the License, or any later version.
% 
%  This program is distributed in the hope that it will be useful,
%  but WITHOUT ANY WARRANTY; without even the implied warranty of
%  MERCHANTABILITY or FITNESS FOR A PARTICULAR PURPOSE.  See the
%  GNU General Public License for more details.
% 
%  You should have received a copy of the GNU General Public License
%  along with this program; if not, write to the Free Software
%  Foundation Inc, 59 Temple Pl. - Suite 330, Boston, MA 02111-1307, USA.

%  Suggestions for improvement and fixes are always welcome, although no
%  guarantee is made whether and when they will be implemented.
%  Send requests to

narginchk(0,1);

disp('This is a beta version of mwizard.');
disp('Please use the online version m2html instead.');

h = initWindow;

initOptions(h);

buildWindow(h);

setappdata(h, 'handles', guihandles(h));
% M. Krauski
% setappdata(h, 'pwd',     pwd);

if nargin == 0
	setappdata(h, 'file', '');
	setappdata(h, 'needsave', 1);
else
	setappdata(h, 'file', file);
	setappdata(h, 'needsave', 0);
	opt = load(file, 'options');
	setappdata(h, 'options', opt.options);
    refreshOptions(h);
end

set(h, 'HandleVisibility', 'callback');

%===============================================================================

function h = initWindow

h = figure('Resize',      'on',...
		   'MenuBar',     'none',...
		   'NumberTitle', 'off',...
		   'Name',        ':: M2HTML Wizard ::',...
		   'Position',    [200 200 500 650],...
		   'Tag',         mfilename);
		   
set(h, 'CloseRequestFcn', {@doClose,h});

%===============================================================================

function buildWindow(h)

wincolor = struct('bg',    [0.9 0.9 0.9], ...
                  'fg',    [0.8 0.8 0.8], ...
                  'title', [0.8 0.8 0.9]);

set(h, 'Color', wincolor.bg);
              
%-------------------------------------------------------------------------------
%- Menu
%-------------------------------------------------------------------------------

icons = load(fullfile(fileparts(which(mfilename)),'private', ...
            'm2htmltoolbarimages.mat'));

uipushtool('CData',icons.newIcon,...
	'enable','on',...
	'Separator','off',...
	'ToolTipString','New File',...
	'ClickedCallback',{@doNewFile,h},...
	'Tag','NewTool');

uipushtool('CData',icons.openIcon,...
	'enable','on',...
	'Separator','off',...
	'ToolTipString','Open File',...
	'ClickedCallback',{@doOpenFile,h},...
	'Tag','OpenTool');

uipushtool('CData',icons.saveIcon,...
	'enable','on',...
	'Separator','off',...
	'ToolTipString','Save File',...
	'ClickedCallback',{@doSaveFile,h},...
	'Tag','SaveTool');

uipushtool('CData',icons.saveAsIcon,...
	'enable','on',...
	'Separator','off',...
	'ToolTipString','Save File As',...
	'ClickedCallback',{@doSaveAsFile,h},...
	'Tag','SaveAsTool');
	
uipushtool('CData',icons.wheelIcon,...
	'enable','on',...
	'Separator','on',...
	'ToolTipString','Save and Run M2HTML',...
	'ClickedCallback',{@doRunFile,h},...
	'Tag','RunTool');

uipushtool('CData',icons.webIcon,...
	'enable','on',...
	'Separator','on',...
	'ToolTipString','Online Tutorial',...
	'ClickedCallback',...
		'web(''http://www.artefact.tk/software/matlab/m2html/'')',...
	'Tag','WebTool');

uipushtool('CData',icons.helpIcon,...
	'enable','on',...
	'Separator','off',...
	'ToolTipString','Help',...
	'ClickedCallback',{@doHelp,h},...
	'Tag','HelpTool');

%-------------------------------------------------------------------------------
%- Title
%-------------------------------------------------------------------------------

uicontrol('Style','Frame',...
	'Units','Normalized',...
	'Position',[0.02,0.92,0.96,0.06],...
	'BackgroundColor',wincolor.title);

uicontrol('Style','Text',...
	'Units','Normalized',...
	'String','M2HTML Wizard',...
	'FontSize',18,...
	'HorizontalAlignment','center',...
	'Position',[0.03,0.93,0.94,0.038],...
	'BackgroundColor',wincolor.title);

%-------------------------------------------------------------------------------
%- Input
%-------------------------------------------------------------------------------

uicontrol('Style','Frame',...
	'Units','Normalized',...
	'Position',[0.02,0.74,0.96,0.16],...
	'BackgroundColor',wincolor.fg);
	
uicontrol('Style','Frame',...
	'Units','Normalized',...
	'HorizontalAlignment','center',...
	'Position',[0.02,0.87,0.96,0.03],...
	'BackgroundColor',wincolor.title);
	
uicontrol('Style','Text',...
	'Units','Normalized',...
	'String','M-Files Input',...
	'HorizontalAlignment','left',...
	'Position',[0.03,0.875,0.94,0.02],...
	'BackgroundColor',wincolor.title);

uicontrol('Style','Text',...
	'Units','Normalized',...
	'String','Root directory:',...
    'FontAngle','oblique',...
	'HorizontalAlignment','left',...
	'Position',[0.04,0.825,0.6,0.03],...
	'BackgroundColor',wincolor.fg);

% M. Krauski 7/20/04
% changed and resized control to work with browse push button

% uicontrol('Style','edit',...
% 	'Units','Normalized',...
% 	'Position',[0.21,0.83,0.74,0.03],...
% 	'String',pwd,...
% 	'Enable','inactive',...
% 	'HorizontalAlignment','left',...
% 	'Callback','uigetfile;',...%uigetdir
% 	'BackgroundColor',wincolor.bg,...
% 	'Tag','rootdir');

uicontrol('Style','edit',...
	'Units','Normalized',...
	'Position',[0.21,0.83,0.59,0.03],...
	'String',pwd,... %this will be over written in refreshOptions
	'Enable','inactive',...
	'HorizontalAlignment','left',...
	'BackgroundColor',wincolor.bg,...
	'Tag','rootdir');

% M. Krauski 7/20/04
% added control
uicontrol('Style','pushbutton',...
	'Units','Normalized',...
	'Position',[0.81,0.83,0.14,0.03],...
	'String','Browse...',...
	'HorizontalAlignment','left',...
    'Callback',{@doBrowseButton,h},...
	'BackgroundColor',wincolor.bg,...
	'Tag','browsebutton');

uicontrol('Style','Text',...
	'Units','Normalized',...
	'String','Relative pathes:',...
	'HorizontalAlignment','left',...
	'Position',[0.04,0.785,0.6,0.03],...
	'BackgroundColor',wincolor.fg);

% M. Krauski 7/20/04
% changed and resized control to work with select push button

% uicontrol('Style','edit',...
% 	'Units','Normalized',...
% 	'Position',[0.21,0.79,0.74,0.03],...
% 	'String','',...
% 	'HorizontalAlignment','left',...
% 	'Callback',{@doSetMfiles,h},...
%     'CreateFcn',{@doInitMfiles,h},...
% 	'BackgroundColor',wincolor.bg,...
% 	'Tag','mfiles');

uicontrol('Style','edit',...
	'Units','Normalized',...
    'Position',[0.21,0.79,0.59,0.03],...
	'String','',...
    'Enable','inactive',...
	'HorizontalAlignment','left',...
    'CreateFcn',{@doInitMfiles,h},...
	'BackgroundColor',wincolor.bg,...
	'Tag','mfiles');

uicontrol('Style','pushbutton',...
	'Units','Normalized',...
	'Position',[0.81,0.79,0.14,0.03],...
	'String','Select...',...
	'HorizontalAlignment','left',...
    'Callback',{@doSelectMfiles,h},...
	'BackgroundColor',wincolor.bg,...
	'Tag','selectbutton');

uicontrol('Style','CheckBox',...
	'Units','Normalized',...
	'Position',[0.04,0.749,0.42,0.032],...
	'String',' Recursive',...
	'HorizontalAlignment','left',...
	'Callback',{@doSetRecursive,h},...
	'Value',0,...
	'BackgroundColor',wincolor.bg,...
	'Tag','recursive');

%-------------------------------------------------------------------------------
%- Output
%-------------------------------------------------------------------------------

uicontrol('Style','Frame',...
	'Units','Normalized',...
	'Position',[0.02, 0.56,0.96,0.16],...
	'BackgroundColor',wincolor.fg);

uicontrol('Style','Frame',...
	'Units','Normalized',...
	'HorizontalAlignment','center',...
	'Position',[0.02,0.69,0.96,0.03],...
	'BackgroundColor',wincolor.title);
	
uicontrol('Style','Text',...
	'Units','Normalized',...
	'String','HTML Output',...
	'HorizontalAlignment','left',...
	'Position',[0.03,0.695,0.94,0.02],...
	'BackgroundColor',wincolor.title);

uicontrol('Style','Text',...
	'Units','Normalized',...
	'String','Output Directory:',...
	'HorizontalAlignment','left',...
	'Position',[0.04,0.645,0.6,0.03],...
	'BackgroundColor',wincolor.fg);

uicontrol('Style','edit',...
	'Units','Normalized',...
	'Position',[0.21,0.65,0.74,0.03],...
	'String','',...
	'HorizontalAlignment','left',...
	'Callback',{@doSetOutputDir,h},...
    'CreateFcn',{@doInitHTMLDir,h},...
	'BackgroundColor',wincolor.bg,...
	'Tag','htmldir');

uicontrol('Style','Text',...
	'Units','Normalized',...
	'String','HTML Index:',...
	'HorizontalAlignment','left',...
	'Position',[0.04,0.605,0.6,0.03],...
	'BackgroundColor',wincolor.fg);

uicontrol('Style','edit',...
	'Units','Normalized',...
	'Position',[0.21,0.61,0.25,0.03],...
	'String','index',...
	'HorizontalAlignment','left',...
	'Callback',{@doSetIndex,h},...
	'BackgroundColor',wincolor.bg,...
	'Tag','index');

uicontrol('Style','Text',...
	'Units','Normalized',...
	'String','Extension:',...
	'HorizontalAlignment','left',...
	'Position',[0.53,0.605,0.3,0.03],...
	'BackgroundColor',wincolor.fg);

uicontrol('Style','edit',...
	'Units','Normalized',...
	'Position',[0.70,0.61,0.25,0.03],...
	'String','html',...
	'HorizontalAlignment','left',...
	'Callback',{@doSetExtension,h},...
	'BackgroundColor',wincolor.bg,...
	'Tag','extension');

uicontrol('Style','Text',...
	'Units','Normalized',...
	'String','Template:',...
	'HorizontalAlignment','left',...
	'Position',[0.04,0.565,0.3,0.03],...
	'BackgroundColor',wincolor.fg);

uicontrol('Style','popupmenu',...
	'Units','Normalized',...
	'Position',[0.21,0.57,0.25,0.03],...
	'String','',...
	'HorizontalAlignment','center',...
	'Callback',{@doSetTemplate,h},...
	'CreateFcn',{@doInitTpl,h},...
	'BackgroundColor',wincolor.bg,...
	'Tag','template');

%-------------------------------------------------------------------------------
%- Other options
%-------------------------------------------------------------------------------

uicontrol('Style','Frame',...
	'Units','Normalized',...
	'Position',[0.02,0.24,0.96,0.30],...
	'BackgroundColor',wincolor.fg);

uicontrol('Style','Frame',...
	'Units','Normalized',...
	'HorizontalAlignment','center',...
	'Position',[0.02,0.51,0.96,0.03],...
	'BackgroundColor',wincolor.title);
	
uicontrol('Style','Text',...
	'Units','Normalized',...
	'String','Other Options',...
	'HorizontalAlignment','left',...
	'Position',[0.03,0.515,0.94,0.02],...
	'BackgroundColor',wincolor.title);

uicontrol('Style','checkbox',...
	'Units','Normalized',...
	'Position',[0.04,0.464,0.42,0.032],...
	'String',' Include Source Code',...
	'HorizontalAlignment','left',...
	'Callback',{@doSetSource,h},...
	'Value',1,...
	'TooltipString','Include Source Code of each M-file',...
	'BackgroundColor',wincolor.bg,...
	'Tag','source');

uicontrol('Style','checkbox',...
	'Units','Normalized',...
	'Position',[0.53,0.464,0.42,0.032],...
	'String',' Syntax Highlighting',...
	'HorizontalAlignment','left',...
	'Callback',{@doSetHighlight,h},...
	'Value',1,...
	'TooltipString','Source Code Syntax Highlighting',...
	'BackgroundColor',wincolor.bg,...
	'Tag','highlight');

uicontrol('Style','checkbox',...
	'Units','Normalized',...
	'Position',[0.04,0.42,0.42,0.032],...
	'String',' Create Dependency Graphs',...
	'HorizontalAlignment','left',...
	'Callback',{@doSetGraph,h},...
    'CreateFcn',{@doInitGraphs,h},...
	'Value',0,...
	'TooltipString','Compute a Dependency Graph using GraphViz',...
	'BackgroundColor',wincolor.bg,...
	'Tag','graph');

uicontrol('Style','checkbox',...
	'Units','Normalized',...
	'Position',[0.53,0.42,0.42,0.032],...
	'String',' PHP Search Engine',...
	'HorizontalAlignment','left',...
	'Callback',{@doSetSearch,h},...
	'Value',0,...
	'TooltipString','Create an Index for a PHP Search Engine',...
	'BackgroundColor',wincolor.bg,...
	'Tag','search');

uicontrol('Style','checkbox',...
	'Units','Normalized',...
	'Position',[0.04,0.378,0.42,0.032],...
	'String',' Global Hyperlinks',...
	'HorizontalAlignment','left',...
	'Callback',{@doSetGlobal,1},...
	'Value',0,...
	'TooltipString','Hypertext links among separate Matlab Directories',...
	'BackgroundColor',wincolor.bg,...
	'Tag','globalhypertext');

uicontrol('Style','checkbox',...
	'Units','Normalized',...
	'Position',[0.53,0.378,0.42,0.032],...
	'String',' Downloadable M-files',...
	'HorizontalAlignment','left',...
	'Callback',{@doSetDownload,h},...
	'TooltipString','Add a link to download each M-file separately',...
	'Value',0,...
	'BackgroundColor',wincolor.bg,...
	'Tag','download');

uicontrol('Style','checkbox',...
	'Units','Normalized',...
	'Position',[0.04,0.336,0.42,0.032],...
	'String',' To Do List',...
	'HorizontalAlignment','left',...
	'Callback',{@doSetTodo,h},...
	'TooltipString',['Create a TODO list in each directory summarizing'...
	' all the ''% TODO %'' lines found in Matlab code'],...
	'Value',0,...
	'BackgroundColor',wincolor.bg,...
	'Tag','todo');

uicontrol('Style','checkbox',...
	'Units','Normalized',...
	'Position',[0.53,0.336,0.42,0.032],...
	'String',' Verbose Mode',...
	'HorizontalAlignment','left',...
	'Callback',{@doSetVerbose,h},...
	'TooltipString','Verbose mode',...
	'Value',1,...
	'BackgroundColor',wincolor.bg,...
	'Tag','verbose');

uicontrol('Style','checkbox',...
	'Units','Normalized',...
	'Position',[0.04,0.294,0.42,0.032],...
	'String',' Save M-files Parsing',...
	'HorizontalAlignment','left',...
	'Callback',{@doSetSaveAsMat,h},...
	'TooltipString',['Save current state after M-files parsing in '...
	'''m2html.mat'' in the Output directory'],...
	'Value',0,...
	'BackgroundColor',wincolor.bg,...
	'Tag','save');

uicontrol('Style','Text',...
	'Units','Normalized',...
	'String','Load File:',...
	'HorizontalAlignment','left',...
	'Position',[0.53,0.289,0.3,0.03],...
	'BackgroundColor',wincolor.fg);

uicontrol('Style','edit',...
	'Units','Normalized',...
	'Position',[0.70,0.294,0.25,0.03],...
	'String','',...
	'HorizontalAlignment','left',...
	'Callback',{@doSetLoadMat,h},...
	'TooltipString',['Load a previously saved MAT file '...
	'to generate HTML files once again'],...
	'BackgroundColor',wincolor.bg,...
	'Tag','load');

uicontrol('Style','Text',...
	'Units','Normalized',...
	'String','Tabs Length:',...
	'HorizontalAlignment','left',...
	'Position',[0.04,0.247,0.3,0.03],...
	'BackgroundColor',wincolor.fg);

uicontrol('Style','edit',...
	'Units','Normalized',...
	'Position',[0.21,0.252,0.25,0.03],...
	'String','4',...
	'HorizontalAlignment','right',...
	'Callback',{@doSetTabs,h},...
	'TooltipString',['Replace horizontal tabs in source code '...
	'by N white space characters'],...
	'BackgroundColor',wincolor.bg,...
	'Tag','tabs');

uicontrol('Style','Text',...
	'Units','Normalized',...
	'String','Nb Columns:',...
    'FontAngle','oblique',...
	'HorizontalAlignment','left',...
	'Position',[0.53,0.247,0.3,0.03],...
	'BackgroundColor',wincolor.fg);

uicontrol('Style','edit',...
	'Units','Normalized',...
	'Position',[0.70,0.252,0.25,0.03],...
	'String','4',...
	'HorizontalAlignment','right',...
	'Callback',{@doSetNbColumns,h},...
	'TooltipString','Number of columns for M-files output - not available',...
    'Enable','inactive',...
	'BackgroundColor',wincolor.bg,...
	'Tag','column');


%-------------------------------------------------------------------------------
%- Space available
%-------------------------------------------------------------------------------

% uicontrol('Style','Frame',...
% 	'Units','Normalized',...
% 	'Position',[0.02,0.07,0.96,0.14],...
% 	'BackgroundColor',wincolor.fg);

% simulate a frame using an axes
% http://www.mathworks.com/support/solutions/data/1-15P9E.html
axes('Color',wincolor.fg,...
    'XTick',[],'YTick',[],...
    'Units','Normalized',...
    'Box','on',...
    'Position',[0.02,0.07,0.9585,0.14]);

uicontrol('Style','Frame',...
	'Units','Normalized',...
	'HorizontalAlignment','center',...
	'Position',[0.02,0.19,0.96,0.03],...
	'BackgroundColor',wincolor.title);

uicontrol('Style','Text',...
	'Units','Normalized',...
	'String','M2HTML status',...
	'HorizontalAlignment','left',...
	'Position',[0.03,0.195,0.94,0.02],...
	'BackgroundColor',wincolor.title);

uicontrol('Style','Text',...
	'Units','Normalized',...
	'String','Click on the wheel in the toolbar to launch M2HTML...',...
	'HorizontalAlignment','left',... % center
	'Position',[0.12,0.135,0.76,0.02],...
    'Visible','on',...
	'BackgroundColor',wincolor.fg,...
	'Tag','textmisc');

axes('XLim',[0 100],...
    'YLim',[0 1],...
    'Box','on', ...
    'Units','Normalized',...
    'Position',[0.07,0.09,0.86,0.03],...
    'XTickMode','manual',...
    'YTickMode','manual',...
    'layer','top',...
    'XTick',[],...
    'YTick',[],...
    'XTickLabelMode','manual',...
    'XTickLabel',[],...
    'YTickLabelMode','manual',...
    'Visible','on',...
    'YTickLabel',[],...
    'Color',wincolor.bg);

x = 0; % between 0 and 100
xpatch = [0 x x 0];
ypatch = [0 0 1 1];
  
p = patch(xpatch,ypatch,'r',...
    'EdgeColor','r',...
    'Visible','on',...
    'EraseMode','none',...
	'Tag','waitbarmisc');
  
l = line([100 0 0 100 100], [0 0 1 1 0], ...
    'EraseMode','none', ...
    'Visible','on',...
    'Color',get(gca,'XColor'));
  
% for i=10:5:100
%     set(p,'Xdata',[0 i i 0]); pause(0.02);
% end
% set(p,'EraseMode','normal');
% set(p,'Xdata',[0 0 0 0]);
% set(p,'EraseMode','none');

%-------------------------------------------------------------------------------
%- Footnote
%-------------------------------------------------------------------------------

uicontrol('Style','Frame',...
	'Units','Normalized',...
	'Position',[0.02,0.02,0.96,0.03],...
	'BackgroundColor',[0.8 0.8 0.9]);

uicontrol('Style','Text',...
	'Units','Normalized',...
	'String',['M2HTML � 2004 Guillaume Flandin <>'],...
	'HorizontalAlignment','right',...
	'Position',[0.03,0.025,0.94,0.02],...
	'BackgroundColor',[0.8 0.8 0.9]);

%===============================================================================

function doClose(fig,evd,h)
	status = doCheckSave(h);
	if status
		delete(h);
	end
	
function doNewFile(fig,evd,h)
	status = doCheckSave(h);
	if status
		initOptions(h);
		setappdata(h, 'needsave', 1);
		% refresh options in GUI...
        refreshOptions(h);
	end

function doOpenFile(fig,evd,h)
	status = doCheckSave(h);
	if status
		[filename, pathname] = uigetfile('*.mat','Open File');
		if ~(isequal(filename,0)|isequal(pathname,0))
			opt = load(fullfile(pathname,filename),'options');
			setappdata(h,'options',opt.options);
			setappdata(h,'file',fullfile(pathname,filename));
		end
	end
    % refresh options in GUI...
    refreshOptions(h);

function status = doSaveFile(fig,evd,h)
	file = getappdata(h,'file');
	status = 1;
	if isempty(file)
		status = doSaveAsFile(fig,evd,h);
	else
		options = getappdata(h,'options');
		save(file, 'options');
    end
	setappdata(h,'needsave',0);

function status = doSaveAsFile(fig,evd,h)
    opt = getappdata(h,'options');
	[filename, pathname] = uiputfile(strcat(opt.rootdir,filesep,'matlab.mat'),...
        'Save File as');
	if ~(isequal(filename,0)|isequal(pathname,0))
		setappdata(h,'file',fullfile(pathname,filename));
		status = doSaveFile(fig,evd,h);
	else
		status = 0;
	end

function doRunFile(fig,evd,h)
	status = doSaveFile(fig,evd,h);
	if status
		opt = getappdata(h,'options');
        file = getappdata(h,'file');
        r = {'off' 'on'};
        % opt could be directly given to m2html (no need for file saving)
        % just need to convert on/off using opt.param = r{opt.param+1}
		m2html('load',file,'recursive',r{opt.recursive+1});
        % 'recursive' is specified to force m2html to parse M-files
	end
	
function status = doCheckSave(h)
	file = getappdata(h,'file');
	if isempty(file), file = 'Untitled'; end
	needsave = getappdata(h,'needsave');
	status = 1;
	if needsave
		button = questdlg(sprintf('Save changes to %s?',file),...
			'Mwizard','Yes','No','Cancel','Yes');
		if strcmp(button,'Yes')
			status = doSaveFile([],[],h);
		elseif strcmp(button,'Cancel')
			status = 0;
		end
	end

function doHelp(fig,evd,h)
	helpdlg(sprintf(['M2HTML by Guillaume Flandin\n'...
		'Copyright � 2003-2004\\n'...
		'<http://www.artefact.tk/>']),'M2HTML Wizard');

%===============================================================================

%-------------------------------------------------------------------------------
%- Default parameters
%-------------------------------------------------------------------------------

function varargout = initOptions(h)
	options = struct('verbose', 1,...
		'mFiles', {{''}},...
		'htmlDir', 'doc',...
		'recursive', 0,...
		'source', 1,...
		'download',0,...
		'syntaxHighlighting', 1,...
		'tabs', 4,...
		'globalHypertextLinks', 0,...
		'graph', 0,...
		'todo', 0,...
		'load', 0,...
		'save', 0,...
		'search', 0,...
		'helptocxml', 0,...
		'indexFile', 'index',...
		'extension', '.html',...
		'template', 'blue',...
        'rootdir', pwd,...
		'ignoredDir', {{'.svn' 'cvs'}}, ...
        'language','english');
	
    if nargin == 1,
	    setappdata(h,'options',options);
    else
        varargout{1} = options;    
    end

function refreshOptions(h)
    opt = getappdata(h,'options');
    handles = getappdata(h,'handles');
    
    doInitTpl(handles.template,    0, h);
    doInitMfiles(handles.mfiles,   0, h);
    doInitHTMLDir(handles.htmldir, 0, h)
    
    set(handles.recursive,       'Value',  opt.recursive);
    set(handles.graph,           'Value',  opt.graph); %doInitGraphs(handles.graph,0,h);
    set(handles.save,            'Value',  opt.save);
    set(handles.verbose,         'Value',  opt.verbose);
    set(handles.todo,            'Value',  opt.todo);
    set(handles.download,        'Value',  opt.download);
    set(handles.search,          'Value',  opt.search);
    set(handles.highlight,       'Value',  opt.syntaxHighlighting);
    set(handles.source,          'Value',  opt.source);
    set(handles.globalhypertext, 'Value',  opt.globalHypertextLinks);
    
    set(handles.index,           'String', opt.indexFile);
    set(handles.extension,       'String', opt.extension(2:end)); %remove the '.'
    set(handles.tabs,            'String', num2str(opt.tabs));
%     if ~strcmp(opt.rootdir, pwd)
%         warning('[M2HTML] You should ''cd %s'' before...',opt.rootdir);    
%     end
    set(handles.rootdir,         'String', opt.rootdir); % need to 'cd' if different...
    set(handles.column,          'String', num2str(4)); %- not saved... default here
    if ischar(opt.load)
        set(handles.load,        'String', opt.load);
    else
        set(handles.load,        'String', '');
    end
    
    set(handles.textmisc,        'String', ...
        'Click on the wheel in the toolbar to launch M2HTML...'); %- not saved... default here
    set(handles.waitbarmisc,     'EraseMode','normal');
    set(handles.waitbarmisc,     'Xdata',[0 0 0 0]);
    set(handles.waitbarmisc,     'EraseMode','none');


%-------------------------------------------------------------------------------
%- CreateFcn Callbacks
%-------------------------------------------------------------------------------

function doInitHTMLDir(fig,evd,h)
    opt = getappdata(h,'options');
    [path,name] = fileparts(opt.htmlDir);
    if isempty(path)
        opt.htmlDir = fullfile(opt.rootdir, opt.htmlDir);
    end
    set(fig,'String',opt.htmlDir);
    setappdata(h,'options',opt);

function doInitTpl(fig,evd,h)
    %- problems when templates are still in full format
    opt = getappdata(h,'options');
	d = dir(fullfile(fileparts(which(mfilename)),'templates'));
	d = {d([d.isdir]).name};
	d = {d{~ismember(d,{'.' '..'})}};
	if ~isempty(d)
		tpl = sprintf('%s|',d{:});
		set(fig,'String',tpl(1:end-1));
		i = strmatch(opt.template,d,'exact');
		if ~isempty(i)
			set(fig,'Value',i(1));
        else
            %- where is the default template ?
            warning('[M2HTML] Default template ''%s'' not found.',opt.template);
            set(fig,'Value',1);
            opt.template = d{1};
            setappdata(h,'options',opt);
            warning('[M2HTML] Using template ''%s'' instead.',opt.template);
        end
	else
		error('[M2HTML] No template found.');
	end

 function doInitMfiles(fig,evd,h)
    opt = getappdata(h,'options');
    if ~isempty(opt.mFiles{1})
        s = sprintf('''%s'', ',opt.mFiles{:}); s = s(1:end-2);
        set(fig,'String',['{' s '}']);
        return;
    end
    d = dir(opt.rootdir); d = {d([d.isdir]).name};
    d = {d{~ismember(d,{'.' '..'})}};
    if length(d) == 0
        warning('[M2HTML] No subsequent directory found. Check your cwd.');
        set(fig,'String',''); %- maybe open a uigetdir ?
        opt.mFiles = {''};
    elseif length(d) == 1
        set(fig,'String',d{1});
        opt.mFiles = d;
    else
        s = sprintf('''%s'', ',d{:}); s = s(1:end-2);
        set(fig,'String',['{' s '}']);
        opt.mFiles = d;
    end
    setappdata(h,'options',opt);
    
function doInitGraphs(fig,evd,h)
    opt = getappdata(h,'options');
    [s, w] = system('dot -V');
    if s
        disp('GraphViz not installed: Generation of dependency graphs desactivated.');
        disp('See http://www.graphviz.org/ to get ''dot'' tool.');
        set(fig,'FontAngle','Oblique','Enable','inactive');
        set(fig,'Value',0);
        opt.graph = 0;
        setappdata(h,'options',opt);
    else
        set(fig,'Value',opt.graph);
    end


%===============================================================================

%-------------------------------------------------------------------------------
%- M-Files Input Callbacks
%-------------------------------------------------------------------------------

function doBrowseButton(fig,evd,h)
    opt = getappdata(h,'options');
    handles = getappdata(h,'handles');
    d = uigetdir(pwd,'Select the Root directory');
    if ~d, return; end;
    opt.rootdir = d;
    set(handles.rootdir,'String',d);
    
    % set mfiles
    d = dir(opt.rootdir); d = {d([d.isdir]).name};
    d = {d{~ismember(d,{'.' '..'})}};
    if length(d) == 0
        warning('[M2HTML] No subsequent directory found. Check your cwd.');
        set(handles.mfiles,'String','');
        opt.mFiles = {''};
    elseif length(d) == 1
        set(handles.mfiles,'String',d{1});
        opt.mFiles = d;
    else
        s = sprintf('''%s'', ',d{:}); s = s(1:end-2);
        set(handles.mfiles,'String',['{' s '}']);
        opt.mFiles = d;
    end
    
    % set htmldir
    [path,name] = fileparts(opt.htmlDir);
    opt.htmlDir = fullfile(opt.rootdir, name);
    set(handles.htmldir,'String',opt.htmlDir);
    
    setappdata(h,'options',opt);
        
function doSelectMfiles(fig,evd,h)
    opt = getappdata(h,'options');
    handles = getappdata(h,'handles');
    
    d = dir(opt.rootdir); d = {d([d.isdir]).name};
    d = {d{~ismember(d,{'.' '..'})}};
    
    [i,v] = listdlg('ListString',d,...
        'PromptString','Select folder(s):',...
        'Name',':: M2HTML :: M-files',...
        'SelectionMode','multiple');
    if v == 1
        d = {d{i}};
        s = sprintf('''%s'', ',d{:}); s = s(1:end-2);
        set(handles.mfiles,'String',['{' s '}']);
    end
    opt.mFiles = d;

    setappdata(h,'options',opt);

function doSetRecursive(fig,evd,h)
	opt = getappdata(h,'options');
	opt.recursive = get(fig,'Value');
	setappdata(h,'options',opt);

%-------------------------------------------------------------------------------
%- HTML Output Callbacks
%-------------------------------------------------------------------------------

function doSetOutputDir(fig,evd,h)
	opt = getappdata(h,'options');
	opt.htmlDir = get(fig,'String');
	setappdata(h,'options',opt);

function doSetIndex(fig,evd,h)
	opt = getappdata(h,'options');
	opt.indexFile = get(fig,'String');
	setappdata(h,'options',opt);
	
function doSetExtension(fig,evd,h)
	opt = getappdata(h,'options');
	e = get(fig,'String');
	if ~isempty(e) & e(1) ~= '.'
		e = ['.' e];
	end
	opt.extension = e;
	setappdata(h,'options',opt);
	
function doSetTemplate(fig,evd,h)
    opt = getappdata(h,'options');
    s = get(fig,'String');
    v = get(fig,'Value');
    opt.template = deblank(s(v,:));
    setappdata(h,'options',opt);

%-------------------------------------------------------------------------------
%- Options Callbacks
%-------------------------------------------------------------------------------

function doSetSource(fig,evd,h)
	opt = getappdata(h,'options');
	opt.source = get(fig,'Value');
	setappdata(h,'options',opt);

function doSetHighlight(fig,evd,h)
	opt = getappdata(h,'options');
	opt.syntaxHighlighting = get(fig,'Value');
	setappdata(h,'options',opt);

function doSetGraph(fig,evd,h)
	opt = getappdata(h,'options');
	opt.graph = get(fig,'Value');
	setappdata(h,'options',opt);

function doSetSearch(fig,evd,h)
	opt = getappdata(h,'options');
	opt.search = get(fig,'Value');
	setappdata(h,'options',opt);

function doSetGlobal(fig,evd,h)
	opt = getappdata(h,'options');
	opt.globalHypertextLinks = get(fig,'Value');
	setappdata(h,'options',opt);

function doSetDownload(fig,evd,h)
	opt = getappdata(h,'options');
	opt.download = get(fig,'Value');
	setappdata(h,'options',opt);

function doSetTodo(fig,evd,h)
	opt = getappdata(h,'options');
	opt.todo = get(fig,'Value');
	setappdata(h,'options',opt);

function doSetVerbose(fig,evd,h)
	opt = getappdata(h,'options');
	opt.verbose = get(fig,'Value');
	setappdata(h,'options',opt);

function doSetSaveAsMat(fig,evd,h)
	opt = getappdata(h,'options');
	opt.save = get(fig,'Value');
	setappdata(h,'options',opt);

function doSetLoadMat(fig,evd,h)
	opt = getappdata(h,'options');
	[fname, pname, findex] = uigetfile('m2html.mat',...
        'Load a m2html MAT-file');
    if findex
        opt.load = fullfile(pname,fname);
        set(fig,'String',fullfile(pname,fname));
    end
	setappdata(h,'options',opt);

function doSetTabs(fig,evd,h)
	opt = getappdata(h,'options');
	t = str2num(get(fig,'String'));
	if t >= 0 & length(t) == 1 
		opt.tabs = t; 
	else
		set(fig,'String',num2str(opt.tabs));
	end
	setappdata(h,'options',opt);

function doSetNbColumns(fig,evd,h)
	opt = getappdata(h,'options');
	disp 'Not available';
	setappdata(h,'options',opt);
    
%===============================================================================

function text2 = shortenText(text, l)

    if nargin == 1, l = 64; end
    m = length(text);
    text2 = text;
    if m > l
        s = floor((l - 3) / 2);
        text2 = [text(1:s) '...' text(end-(l-s-3)+1:end)];
    end
```
