## Supplemental File 1 for "A biophysical model of striatal microcircuits suggests delta/theta-rhythmically interleaved gamma and beta oscillations mediate periodicity in motor control": doxyread.m

```
function [statlist, docinfo] = doxyread(filename)
%DOXYREAD Read a 'search.idx' file generated by DOXYGEN
%  STATLIST = DOXYREAD(FILENAME) reads FILENAME (Doxygen search.idx
%  format) and returns the list of keywords STATLIST as a cell array.
%  [STATLIST, DOCINFO] = DOXYREAD(FILENAME) also returns a cell array
%  containing details for each keyword (frequency in each file where it
%  appears and the URL).
%
%  See also DOXYSEARCH, DOXYWRITE

%  Copyright (C) 2003 Guillaume Flandin <>
%  $Revision: 1.0 $Date: 2003/05/10 17:41:21 $

%  This program is free software; you can redistribute it and/or
%  modify it under the terms of the GNU General Public License
%  as published by the Free Software Foundation; either version 2
%  of the License, or any later version.
% 
%  This program is distributed in the hope that it will be useful,
%  but WITHOUT ANY WARRANTY; without even the implied warranty of
%  MERCHANTABILITY or FITNESS FOR A PARTICULAR PURPOSE.  See the
%  GNU General Public License for more details.
% 
%  You should have received a copy of the GNU General Public License
%  along with this program; if not, write to the Free Software
%  Foundation Inc, 59 Temple Pl. - Suite 330, Boston, MA 02111-1307, USA.

%  Suggestions for improvement and fixes are always welcome, although no
%  guarantee is made whether and when they will be implemented.
%  Send requests to <>

%  See <http://www.doxygen.org/> for more details.

narginchk(0,1);
if nargin == 0,
	filename = 'search.idx';
end

%- Open the search index file
[fid, errmsg] = fopen(filename,'r','ieee-be');
if fid == -1, error(errmsg); end

%- 4 byte header (DOXS)
header = char(fread(fid,4,'uchar'))';

%- 256*256*4 byte index
idx = fread(fid,256*256,'uint32');
idx = reshape(idx,256,256);

%- Extract list of words
i = find(idx);
statlist = cell(0,2);	
for j=1:length(i) 
	fseek(fid, idx(i(j)), 'bof');	
	statw    = readString(fid);
	while ~isempty(statw)
		statidx  = readInt(fid);
		statlist{end+1,1} = statw; % word
		statlist{end,2}   = statidx; % index
		statw   = readString(fid);
	end
end
	
%- Extract occurence frequency of each word and docs info (name and url)
docinfo = cell(size(statlist,1),1);
for k=1:size(statlist,1)
	fseek(fid, statlist{k,2}, 'bof');
	numdoc = readInt(fid);
	docinfo{k} = cell(numdoc,4);
	for m=1:numdoc
		docinfo{k}{m,1} = readInt(fid); % idx
		docinfo{k}{m,2} = readInt(fid); % freq
	end
	for m=1:numdoc
		fseek(fid, docinfo{k}{m,1}, 'bof');
		docinfo{k}{m,3} = readString(fid); % name
		docinfo{k}{m,4} = readString(fid); % url
	end
	docinfo{k} = reshape({docinfo{k}{:,2:4}},numdoc,[]);
end

%- Close the search index file
fclose(fid);

%- Remove indexes
statlist = {statlist{:,1}}';

%===========================================================================
function s = readString(fid)

	s = '';
	while 1
		w = fread(fid,1,'uchar');
		if w == 0, break; end
		s(end+1) = char(w);
	end

%===========================================================================
function i = readInt(fid)

	i = fread(fid,1,'uint32');
```
