## Supplemental File 1 for "A biophysical model of striatal microcircuits suggests delta/theta-rhythmically interleaved gamma and beta oscillations mediate periodicity in motor control": doxysearch.m

```
function result = doxysearch(query,filename)
%DOXYSEARCH Search a query in a 'search.idx' file
%  RESULT = DOXYSEARCH(QUERY,FILENAME) looks for request QUERY
%  in FILENAME (Doxygen search.idx format) and returns a list of
%  files responding to the request in RESULT.
%
%  See also DOXYREAD, DOXYWRITE

%  Copyright (C) 2004 Guillaume Flandin <>
%  $Revision: 1.1 $Date: 2004/05/05 14:33:55 $

%  This program is free software; you can redistribute it and/or
%  modify it under the terms of the GNU General Public License
%  as published by the Free Software Foundation; either version 2
%  of the License, or any later version.
% 
%  This program is distributed in the hope that it will be useful,
%  but WITHOUT ANY WARRANTY; without even the implied warranty of
%  MERCHANTABILITY or FITNESS FOR A PARTICULAR PURPOSE.  See the
%  GNU General Public License for more details.
% 
%  You should have received a copy of the GNU General Public License
%  along with this program; if not, write to the Free Software
%  Foundation Inc, 59 Temple Pl. - Suite 330, Boston, MA 02111-1307, USA.

%  Suggestions for improvement and fixes are always welcome, although no
%  guarantee is made whether and when they will be implemented.
%  Send requests to <>

%  See <http://www.doxygen.org/> for more details.

narginchk(1,2);
if nargin == 1,
	filename = 'search.idx';
end

%- Open the search index file
[fid, errmsg] = fopen(filename,'r','ieee-be');
if fid == -1, error(errmsg); end

%- 4 byte header (DOXS)
header = char(fread(fid,4,'uchar'))';
if ~all(header == 'DOXS')
    error('[doxysearch] Header of index file is invalid!');
end

%- many thanks to <doxyread.m> and <doxysearch.php>
r = query;
requiredWords  = {};
forbiddenWords = {};
foundWords     = {};
res            = {};
while 1
    % extract each word of the query
    [t,r] = strtok(r);
    if isempty(t), break, end;
    if t(1) == '+'
        t = t(2:end); requiredWords{end+1} = t;
    elseif t(1) == '-'
        t = t(2:end); forbiddenWords{end+1} = t;
    end
    if ~ismember(t,foundWords)
        foundWords{end+1} = t;
        res = searchAgain(fid,t,res);
    end
end

%- Filter and sort results
docs = combineResults(res);
filtdocs = filterResults(docs,requiredWords,forbiddenWords);
filtdocs = normalizeResults(filtdocs);
res = sortResults(filtdocs);

%- 
if nargout
    result = res;
else
    for i=1:size(res,1)
        fprintf('   %d. %s - %s\n      ',i,res{i,1},res{i,2});
        for j=1:size(res{i,4},1)
            fprintf('%s ',res{i,4}{j,1});
        end
        fprintf('\n');
    end
end

%- Close the search index file
fclose(fid);

%===========================================================================
function res = searchAgain(fid, word,res)

	i = computeIndex(word);
	if i > 0
        
        fseek(fid,i*4+4,'bof'); % 4 bytes per entry, skip header
        start = size(res,1);
        idx = readInt(fid);
        
        if idx > 0
            
            fseek(fid,idx,'bof');
            statw = readString(fid);
		    while ~isempty(statw)
			    statidx  = readInt(fid);
                if length(statw) >= length(word) & ...
                    strcmp(statw(1:length(word)),word)
			        res{end+1,1} = statw;   % word
                    res{end,2}   = word;    % match
			        res{end,3}   = statidx; % index
                    res{end,4}   = (length(statw) == length(word)); % full
                    res{end,5}   = {};      % doc
                end
			    statw = readString(fid);
        	end
        
            totalfreq = 0;
            for j=start+1:size(res,1)
                fseek(fid,res{j,3},'bof');
                numdoc = readInt(fid);
                docinfo = {};
                for m=1:numdoc
			        docinfo{m,1} = readInt(fid); % idx
			        docinfo{m,2} = readInt(fid); % freq
                    docinfo{m,3} = 0;            % rank
                    totalfreq = totalfreq + docinfo{m,2};
                    if res{j,2}, 
                        totalfreq = totalfreq + docinfo{m,2};
                    end;
		        end
		        for m=1:numdoc
			        fseek(fid, docinfo{m,1}, 'bof');
			        docinfo{m,4} = readString(fid); % name
			        docinfo{m,5} = readString(fid); % url
		        end
                res{j,5} = docinfo;
            end
        
            for j=start+1:size(res,1)
                for m=1:size(res{j,5},1)
                    res{j,5}{m,3} = res{j,5}{m,2} / totalfreq;
                end
            end
            
        end % if idx > 0
        
	end % if i > 0

%===========================================================================
function docs = combineResults(result)

	docs = {};
	for i=1:size(result,1)
        for j=1:size(result{i,5},1)
            key = result{i,5}{j,5};
            rank = result{i,5}{j,3};
            if ~isempty(docs) & ismember(key,{docs{:,1}})
                l = find(ismember({docs{:,1}},key));
                docs{l,3} = docs{l,3} + rank;
                docs{l,3} = 2 * docs{l,3};
            else
                l = size(docs,1)+1;
                docs{l,1} = key; % key
                docs{l,2} = result{i,5}{j,4}; % name
                docs{l,3} = rank; % rank
                docs{l,4} = {}; %words
            end
            n = size(docs{l,4},1);
            docs{l,4}{n+1,1} = result{i,1}; % word
            docs{l,4}{n+1,2} = result{i,2}; % match
            docs{l,4}{n+1,3} = result{i,5}{j,2}; % freq
        end
	end

%===========================================================================
function filtdocs = filterResults(docs,requiredWords,forbiddenWords)

	filtdocs = {};
	for i=1:size(docs,1)
        words = docs{i,4};
        c = 1;
        j = size(words,1);
        % check required
        if ~isempty(requiredWords)
            found = 0;
            for k=1:j
                if ismember(words{k,1},requiredWords)
                    found = 1; 
                    break;  
                end
            end
            if ~found, c = 0; end
        end
        % check forbidden
        if ~isempty(forbiddenWords)
            for k=1:j
                if ismember(words{k,1},forbiddenWords)
                    c = 0;
                    break;
                end
            end
        end
        % keep it or not
        if c, 
            l = size(filtdocs,1)+1;
            filtdocs{l,1} = docs{i,1};
            filtdocs{l,2} = docs{i,2};
            filtdocs{l,3} = docs{i,3};
            filtdocs{l,4} = docs{i,4};
        end;
	end

%===========================================================================
function docs = normalizeResults(docs);

    m = max([docs{:,3}]);
    for i=1:size(docs,1)
        docs{i,3} = 100 * docs{i,3} / m;
    end

%===========================================================================
function result = sortResults(docs);

    [y, ind] = sort([docs{:,3}]);
    result = {};
    ind = fliplr(ind);
    for i=1:size(docs,1)
        result{i,1} = docs{ind(i),1};
        result{i,2} = docs{ind(i),2};
        result{i,3} = docs{ind(i),3};
        result{i,4} = docs{ind(i),4};
    end

%===========================================================================
function i = computeIndex(word)

    if length(word) < 2,
       i = -1;
    else
        i = double(word(1)) * 256 + double(word(2));
    end
    
%===========================================================================
function s = readString(fid)

	s = '';
	while 1
		w = fread(fid,1,'uchar');
		if w == 0, break; end
		s(end+1) = char(w);
	end

%===========================================================================
function i = readInt(fid)

	i = fread(fid,1,'uint32');
```
