## Supplemental File 1 for "A biophysical model of striatal microcircuits suggests delta/theta-rhythmically interleaved gamma and beta oscillations mediate periodicity in motor control": doxywrite.m

```
function doxywrite(filename, kw, statinfo, docinfo)
%DOXYWRITE Write a 'search.idx' file compatible with DOXYGEN
%  DOXYWRITE(FILENAME, KW, STATINFO, DOCINFO) writes file FILENAME
%  (Doxygen search.idx. format) using the cell array KW containing the
%  word list, the sparse matrix (nbword x nbfile) with non-null values
%  in (i,j) indicating the frequency of occurence of word i in file j
%  and the cell array (nbfile x 2) containing the list of urls and names
%  of each file.
%
%  See also DOXYREAD

%  Copyright (C) 2003 Guillaume Flandin <>
%  $Revision: 1.0 $Date: 2003/23/10 15:52:56 $

%  This program is free software; you can redistribute it and/or
%  modify it under the terms of the GNU General Public License
%  as published by the Free Software Foundation; either version 2
%  of the License, or any later version.
% 
%  This program is distributed in the hope that it will be useful,
%  but WITHOUT ANY WARRANTY; without even the implied warranty of
%  MERCHANTABILITY or FITNESS FOR A PARTICULAR PURPOSE.  See the
%  GNU General Public License for more details.
% 
%  You should have received a copy of the GNU General Public License
%  along with this program; if not, write to the Free Software
%  Foundation Inc, 59 Temple Pl. - Suite 330, Boston, MA 02111-1307, USA.

%  Suggestions for improvement and fixes are always welcome, although no
%  guarantee is made whether and when they will be implemented.
%  Send requests to <>

%  See <http://www.doxygen.org/> for more details.

narginchk(4,4);

%- Open the search index file
[fid, errmsg] = fopen(filename,'w','ieee-be');
if fid == -1, error(errmsg); end

%- Write 4 byte header (DOXS)
fwrite(fid,'DOXS','uchar');
pos = ftell(fid);

%- Write 256 * 256 header
idx = zeros(256);
writeInt(fid, idx);

%- Write word lists
i = 1;
idx2 = zeros(1,length(kw));
while 1
	s = kw{i}(1:2);
	idx(double(s(2)+1), double(s(1)+1)) = ftell(fid);
	while i <= length(kw) & strmatch(s, kw{i})
		writeString(fid,kw{i});
		idx2(i) = ftell(fid);
		writeInt(fid,0);
		i = i + 1;
	end
	fwrite(fid, 0, 'int8');
	if i > length(kw), break; end
end

%- Write extra padding bytes
pad = mod(4 - mod(ftell(fid),4), 4);
for i=1:pad, fwrite(fid,0,'int8'); end
pos2 = ftell(fid);

%- Write 256*256 header again
  fseek(fid, pos, 'bof');
  writeInt(fid, idx);

% Write word statistics
fseek(fid,pos2,'bof');
idx3 = zeros(1,length(kw));
for i=1:length(kw)
	idx3(i) = ftell(fid);
	[ia, ib, v] = find(statinfo(i,:));
	counter = length(ia); % counter
	writeInt(fid,counter);
	for j=1:counter
		writeInt(fid,ib(j)); % index
		writeInt(fid,v(j));  % freq
	end
end
pos3 = ftell(fid);

%- Set correct handles to keywords
  for i=1:length(kw)
  	fseek(fid,idx2(i),'bof');
	writeInt(fid,idx3(i));
  end

% Write urls
fseek(fid,pos3,'bof');
idx4 = zeros(1,length(docinfo));
for i=1:length(docinfo)
	idx4(i) = ftell(fid);
	writeString(fid, docinfo{i,1}); % name
	writeString(fid, docinfo{i,2}); % url
end

%- Set corrext handles to word statistics
fseek(fid,pos2,'bof');
for i=1:length(kw)
	[ia, ib, v] = find(statinfo(i,:));
	counter = length(ia);
	fseek(fid,4,'cof'); % counter
	for m=1:counter
		writeInt(fid,idx4(ib(m)));% index
		fseek(fid,4,'cof'); % freq
	end
end

%- Close the search index file
fclose(fid);

%===========================================================================
function writeString(fid, s)

	fwrite(fid,s,'uchar');
	fwrite(fid,0,'int8');

%===========================================================================
function writeInt(fid, i)
	
	fwrite(fid,i,'uint32');
```
