## Supplemental File 1 for "A biophysical model of striatal microcircuits suggests delta/theta-rhythmically interleaved gamma and beta oscillations mediate periodicity in motor control": mexexts.m

```
function [ext, platform] = mexexts
%MEXEXTS List of Mex files extensions
%  MEXEXTS returns a cell array containing the Mex files platform
%  dependent extensions and another cell array containing the full names
%  of the corresponding platforms.
%
%  See also MEX, MEXEXT

%  Copyright (C) 2003 Guillaume Flandin <>
%  $Revision: 1.0 $Date: 2003/29/04 17:33:43 $

ext = {'.mexsol' '.mexhpux' '.mexhp7' '.mexglx' '.mexa64' '.mexi64' '.mexmac' '.dll' '.mexw32' '.mexw64'};

platform = {'Sun Solaris' 'HP-UX' 'HP-UX' 'Linux PC' 'Linux AMD Opteron' 'Linux Intel Itanium2' 'MacIntosh' 'Windows' 'Windows 32' 'Windows 64'};
```
