## Supplemental File 1 for "A biophysical model of striatal microcircuits suggests delta/theta-rhythmically interleaved gamma and beta oscillations mediate periodicity in motor control": mfileparse.m

```
function s = mfileparse(mfile, mdirs, names, options)
%MFILEPARSE Parsing of an M-file to obtain synopsis, help and references
%  S = MFILEPARSE(MFILE, MDIRS, NAMES, OPTIONS) parses the M-file MFILE looking
%  for synopsis (function), H1 line, subroutines and todo tags (if any).
%  It also fills in a boolean array indicating whether MFILE calls M-files 
%  defined by MDIRS (M-files directories) AND NAMES (M-file names).
%  The input OPTIONS comes from M2HTML: fields used are 'verbose', 'global'
%  and 'todo'.
%  Output S is a structure whose fields are:
%     o synopsis: char array (empty if MFILE is a script)
%     o h1line: short one-line description into the first help line
%     o subroutine: cell array of char containing subroutines synopsis
%     o hrefs: boolean array with hrefs(i) = 1 if MFILE calls mdirs{i}/names{i}
%     o todo: structure containing information about potential todo tags
%
%  See also M2HTML

%  Copyright (C) 2003 Guillaume Flandin <>
%  $Revision: 1.0 $Date: 2003/29/04 17:33:43 $

narginchk(3,4);
if nargin == 3,
	options = struct('verbose',1, 'globalHypertextLinks',0, 'todo',0);
end

%- Delimiters used in strtok: some of them may be useless (% " .), removed '.'
strtok_delim = sprintf(' \t\n\r(){}[]<>+-*~!|\\@&/,:;="''%%');

%- Open for reading the M-file
fid = openfile(mfile,'r');
it = 0; % line number

%- Initialize Output
s = struct('synopsis',   '', ...
		   'h1line',     '', ...
		   'subroutine', {{}}, ...
		   'hrefs',      sparse(1,length(names)), ...
		   'todo',       struct('line',[],'comment',{{}}), ...
		   'ismex',      zeros(size(mexexts)));

%- Initialize flag for synopsis cont ('...')
flagsynopcont = 0;
%- Look for synopsis and H1 line
%  Help is the first set of contiguous comment lines in an m-file
%  The H1 line is a short one-line description into the first help line
while 1
	tline = fgetl(fid);
	if ~ischar(tline), break, end
	it = it + 1;
	tline = deblank(fliplr(deblank(fliplr(tline))));
	%- Synopsis line
	if ~isempty(strmatch('function',tline))
		s.synopsis = tline;
		if ~isempty(strmatch('...',fliplr(tline)))
			flagsynopcont = 1;
			s.synopsis = deblank(s.synopsis(1:end-3));
		end
	%- H1 Line
	elseif ~isempty(strmatch('%',tline))
		% allow for the help lines to be before the synopsis
		if isempty(s.h1line)
			s.h1line = fliplr(deblank(tline(end:-1:2)));
		end
		if ~isempty(s.synopsis), break, end
	%- Go through empty lines
	elseif isempty(tline)
		
	%- Code found. Stop.
	else
		if flagsynopcont
			if isempty(strmatch('...',fliplr(tline)))
				s.synopsis = [s.synopsis tline];
				flagsynopcont = 0;
			else
				s.synopsis = [s.synopsis deblank(tline(1:end-3))];
			end
		else
			break;
		end
	end
end

%- Global Hypertext Links option
%  If false, hypertext links are done only among functions in the same
%  directory.
if options.globalHypertextLinks
	hrefnames = names;
else
	indhref = find(strcmp(fileparts(mfile),mdirs));
	hrefnames = {names{indhref}};
end

%- Compute cross-references and extract subroutines
%  hrefs(i) is 1 if mfile calls mfiles{i} and 0 otherwise
while ischar(tline)
	% Remove blanks at both ends
	tline = deblank(fliplr(deblank(fliplr(tline))));
	
	% Split code into meaningful chunks
	splitc = splitcode(tline);
	for j=1:length(splitc)
		if isempty(splitc{j}) | ...
			splitc{j}(1) == '''' | ...
			~isempty(strmatch('...',splitc{j}))
			% Forget about empty lines, char strings or conts
		elseif splitc{j}(1) == '%'
			% Cross-references are not taken into account in comments
			% Just look for potential TODO or FIXME line
			if options.todo
				if ~isempty(strmatch('% TODO',splitc{j})) | ...
				   ~isempty(strmatch('% FIXME',splitc{j}))
					s.todo.line   = [s.todo.line it];
					s.todo.comment{end+1} = splitc{j}(9:end);
				end
			end
		else
			% detect if this line is a declaration of a subroutine
			if ~isempty(strmatch('function',splitc{j}))
				s.subroutine{end+1} = splitc{j};
			else
				% get list of variables and functions
				symbol = {};
				while 1
					[t,splitc{j}] = strtok(splitc{j},strtok_delim);
					if isempty(t), break, end;
					symbol{end+1} = t;
				end
				if options.globalHypertextLinks
					s.hrefs = s.hrefs + ismember(hrefnames,symbol);
				else
					if ~isempty(indhref)
						s.hrefs(indhref) = s.hrefs(1,indhref) + ...
										   ismember(hrefnames,symbol);
					end
				end
			end
		end
	end
	tline = fgetl(fid);
	it = it + 1;
end	

fclose(fid);

%- Look for Mex files
[pathstr,name] = fileparts(mfile);
samename = dir(fullfile(pathstr,[name	'.*']));
samename = {samename.name};
ext = {};
for i=1:length(samename)
	[dummy, dummy, ext{i}] = fileparts(samename{i});
end
s.ismex = ismember(mexexts,ext);
```
