## Supplemental File 1 for "A biophysical model of striatal microcircuits suggests delta/theta-rhythmically interleaved gamma and beta oscillations mediate periodicity in motor control": openfile.m

```
function fid = openfile(filename,permission)
%OPENFILE Open a file in read/write mode, catching errors
%  FID = OPENFILE(FILENAME,PERMISSION) opens file FILENAME
%  in PERMISSION mode ('r' or 'w') and return a file identifier FID.
%  File is opened in text mode: no effect on Unix but useful on Windows.

%  Copyright (C) 2004 Guillaume Flandin <>
%  $Revision: 1.1 $Date: 2004/05/05 17:14:09 $

[fid, errmsg] = fopen(filename,[permission 't']);
if ~isempty(errmsg)
	switch permission
		case 'r'
			error(sprintf('Cannot open %s in read mode.',filename));
		case 'w'
			error(sprintf('Cannot open %s in write mode.',filename));
		otherwise
			error(errmsg);
	end
end
```
