## Supplemental File 1 for "A biophysical model of striatal microcircuits suggests delta/theta-rhythmically interleaved gamma and beta oscillations mediate periodicity in motor control": searchindex.m

```
function [s, freq] = searchindex(mfile, szmin)
%SEARCHINDEX Compute keywords statistics of an M-file
%  S = SEARCHINDEX(MFILE) returns a cell array of char S containing
%  all the keywords (variables, function names, words in comments or
%  char arrays) found in M-file MFILE, of more than 2 characters.
%  S = SEARCHINDEX(MFILE, SZMIN) allows to specify the minimum size
%  SZMIN of the keywords.
%  [S, FREQ] = SEARCHINDEX(...) also returns the occurency frequence
%  of each keyword in the M-file.
%
%  See also M2HTML

%  Copyright (C) 2003 Guillaume Flandin <>
%  $Revision: 1.0 $Date: 2003/04/10 18:32:48 $

narginchk(1,2);
if nargin == 1, szmin = 2; end

%- Delimiters used in strtok
strtok_delim = sprintf(' \t\n\r(){}[]<>+-*^$~#!|\\@&/.,:;="''%%');

%- Open for reading the M-file
fid = openfile(mfile,'r');

%- Initialize keywords list
s = {};

%- Loop over lines
while 1
	tline = fgetl(fid);
	if ~ischar(tline), break, end
	
	%- Extract keywords in each line
	while 1
		[w, tline] = strtok(tline,strtok_delim);
		if isempty(w), break, end;
		%- Check the length of the keyword
		if length(w) > szmin
			s{end+1} = w;
		end
	end
end

%- Close the M-file
fclose(fid);

%- Remove repeted keywords
[s, i, j] = unique(s);

%- Compute occurency frenquency if required
if nargout == 2,
	if ~isempty(s)
		freq = histc(j,1:length(i));
	else
		freq = [];
	end
end
```
