## Supplemental File 1 for "A biophysical model of striatal microcircuits suggests delta/theta-rhythmically interleaved gamma and beta oscillations mediate periodicity in motor control": splitcode.m

```
function splitc = splitcode(code)
%SPLITCODE Split a line of Matlab code in string, comment and other
%  SPLITC = SPLITCODE(CODE) splits line of Matlab code CODE into a cell
%  array SPLITC where each element is either a character array ('...'),
%  a comment (%...), a continuation (...) or something else.
%  Note that CODE = [SPLITC{:}]
%
%  See also M2HTML, HIGHLIGHT

%  Copyright (C) 2003 Guillaume Flandin <>
%  $Revision: 1.0 $Date: 2003/29/04 17:33:43 $

%- Label quotes in {'transpose', 'beginstring', 'midstring', 'endstring'}
iquote = findstr(code,'''');
quotetransp = [double('_''.)}]') ...
			   double('A'):double('Z') ...
			   double('0'):double('9') ...
			   double('a'):double('z')];
flagstring = 0;
flagdoublequote = 0;
jquote = [];
for i=1:length(iquote)
	if ~flagstring
		if iquote(i) > 1 & any(quotetransp == double(code(iquote(i)-1)))
			% => 'transpose';
		else
			% => 'beginstring';
			jquote(size(jquote,1)+1,:) = [iquote(i) length(code)];
			flagstring = 1;
		end
	else % if flagstring
		if flagdoublequote | ...
		   (iquote(i) < length(code) & strcmp(code(iquote(i)+1),''''))
			% => 'midstring';
			flagdoublequote = ~flagdoublequote;
		else
			% => 'endstring';
			jquote(size(jquote,1),2) = iquote(i);
			flagstring = 0;
		end
	end
end

%- Find if a portion of code is a comment
ipercent = findstr(code,'%');
jpercent = [];
for i=1:length(ipercent)
	if isempty(jquote) | ...
	   ~any((ipercent(i) > jquote(:,1)) & (ipercent(i) < jquote(:,2)))
		jpercent = [ipercent(i) length(code)];
		break;
	end
end

%- Find continuation punctuation '...'
icont = findstr(code,'...');
for i=1:length(icont)
	if (isempty(jquote) | ...
		~any((icont(i) > jquote(:,1)) & (icont(i) < jquote(:,2)))) & ...
		(isempty(jpercent) | ...
		icont(i) < jpercent(1))
		jpercent = [icont(i) length(code)];
		break;
	end
end

%- Remove strings inside comments
if ~isempty(jpercent) & ~isempty(jquote)
	jquote(find(jquote(:,1) > jpercent(1)),:) = [];
end

%- Split code in a cell array of strings
icode = [jquote ; jpercent];
splitc = {};
if isempty(icode)
	splitc{1} = code;
elseif icode(1,1) > 1
	splitc{1} = code(1:icode(1,1)-1);
end
for i=1:size(icode,1)
	splitc{end+1} = code(icode(i,1):icode(i,2));
	if i < size(icode,1) & icode(i+1,1) > icode(i,2) + 1
		splitc{end+1} = code((icode(i,2)+1):(icode(i+1,1)-1));
	elseif i == size(icode,1) & icode(i,2) < length(code)
		splitc{end+1} = code(icode(i,2)+1:end);
	end
end
```
