## Supplemental File 1 for "A biophysical model of striatal microcircuits suggests delta/theta-rhythmically interleaved gamma and beta oscillations mediate periodicity in motor control": strtok.m

```
function [token, remainder, quotient] = strtok(string, delimiters)
%Modified version of STRTOK to also return the quotient
%  string = [quotient token remainder]
%STRTOK Find token in string.
%   STRTOK(S) returns the first token in the string S delimited
%   by "white space".   Any leading white space characters are ignored.
%
%   STRTOK(S,D) returns the first token delimited by one of the 
%   characters in D.  Any leading delimiter characters are ignored.
%
%   [T,R] = STRTOK(...) also returns the remainder of the original
%   string.
%   If the token is not found in S then R is an empty string and T
%   is same as S. 
%
%   Copyright 1984-2002 The MathWorks, Inc. 
%   $Revision: 5.14 $  $Date: 2002/04/09 00:33:38 $

token = []; remainder = []; quotient = string;

len = length(string);
if len == 0
    return
end

if (nargin == 1)
    delimiters = [9:13 32]; % White space characters
end

i = 1;
while (any(string(i) == delimiters))
    i = i + 1;
    if (i > len), return, end
end
start = i;
while (~any(string(i) == delimiters))
    i = i + 1;
    if (i > len), break, end
end
sfinish = i - 1;

token = string(start:sfinish);

if (nargout >= 2)
    remainder = string(sfinish + 1:length(string));
end

if (nargout == 3 & start > 1)
	quotient = string(1:start-1);
else
	quotient = [];
end
```
