## Supplemental File 1 for "A biophysical model of striatal microcircuits suggests delta/theta-rhythmically interleaved gamma and beta oscillations mediate periodicity in motor control": README

 M2HTML - Documentation System for Matlab .m files in HTML
 =========================================================

 Copyright (C) 2003-2005 Guillaume Flandin <>
 http://www.artefact.tk/software/matlab/m2html/

 This toolbox is intended to provide automatic generation of M-files 
 documentation in HTML. It reads each M-file in a set of directories
 (eventually recursively) to produce a corresponding HTML file containing
 synopsis, H1 line, help, function calls and called functions with 
 hypertext links, syntax highlighted source code with hypertext, ...
 
 Here is a summary of the features of the toolbox:
  o extraction of H1 lines and help of each function
  o hypertext documentation with functions calls and called functions
  o extraction of subfunctions (hypertext links)
  o ability to work recursively over the subdirectories of a file tree
  o ability to choose whether the source code must be included or not
  o syntax highlighting of the source code (as in the Matlab Editor)
  o ability to choose HTML index file name and extension
  o automatic creation of a TODO list (using % TODO % syntax)
  o "skins": fully customizable output thanks to HTML templates (see below)

 M2HTML may be particularly useful if you want to study code written by
 someone else (a downloaded toolbox, ...) because you will obtain an
 hypertext documentation in which you can easily walk through, thanks
 to your web browser.

 INSTALLATION
 ============
 
 Please read the INSTALL file for installation instructions.
 
 LICENSE
 =======
 
 Please read the LICENSE file for license details.
 
 TUTORIAL
 ========
 
 Note that a tutorial is available online at:
 	<http://www.artefact.tk/software/matlab/m2html/tutorial.php>
 as well as a Frequently Asked Questions repository:
    <http://www.artefact.tk/software/matlab/m2html/faq.php>
 
 An *important* thing to take care is the Matlab current directory: m2html 
 must be launched  one directory above the directory your wanting to generate 
 documentation for.
 For example, imagine your Matlab code is in the directory /home/foo/matlab/
 (or C:\foo\matlab\ on Windows), then you need to go in the foo directory:
 
   >> cd /home/foo  % (or cd C:\foo on Windows)
 Then you can launch m2html with the command:
 
   >> m2html('mfiles','matlab', 'htmldir','doc');
   
 It will populate all the m-files just within the 'matlab' directory, will parse
 them and then will write in the newly created 'doc' directory (/home/foo/doc/,
 resp., C:\foot\doc\) an HTML file for each M-file.
 
 You can also specify several subdirectories using a cell array of directories:

   >> m2html('mfiles',{'matlab/signal' 'matlab/image'}, 'htmldir','doc');
 
 If you want m2html to walk recursively within the 'matlab' directory then you 
 need to set up the recursive option:
 
   >> m2html('mfiles','matlab', 'htmldir','doc', 'recursive','on');
   
 You can also specify whether you want the source code to be displayed in the 
 HTML files (do you want the source code to be readable from everybody ?):
 
   >> m2html('mfiles','matlab', 'htmldir','doc', 'source','off');
 
 You can also specify whether you want global hypertext links (links among 
 separate Matlab directories). By default, hypertext links are only among 
 functions in the same directory (be aware that setting this option may 
 significantly slow down the process).
 
   >> m2html('mfiles','matlab', 'htmldir','doc', 'global','on');
 
 Other parameters can be tuned for your documentation, see the M2HTML help:
 
   >> help m2html
 
 CUSTOMIZATION
 =============
 
 This toolbox uses the HTML Template class so that you can fully customize the
 output. You can modify .tpl files in templates/blue/ or create new templates 
 in a new directory (templates/othertpl)
 You can then use the newly created template in specifying it:
 
   >> m2html( ... , 'template','othertpl');
 
 M2HTML will use your .tpl files (master, mdir, mfile, graph, search and 
 todo.tpl) and will copy all the other files (CSS, images, ....) in the root
 directory of the HTML documentation.
 
 See the template class documentation for more details.
 <http://www.artefact.tk/software/matlab/template/>

 
 Using templates, you can obtain a frame version of the documentation.
 (I don't like frames but it can be useful for documentation purpose with one
 frame with functions list and another with selected function description)
 To do so, use M2HTML like this (*delete* a previous documentation):
 
   >> m2html( ... , 'template','frame', 'index','menu');
 
 You need to specify a new HTML index basename because index.html is used
 by the template system as the frame index. So if your HTML extension is not
 '.html' you need to rename the file template/frame/index.html to your needs.
 Furthermore, 'menu' then becomes a M2HTML keyword and a function cannot have
 this name (if it is your case, use another keyword in the M2HTML syntax and
 modify the content of the file template/frame/index.html accordingly).
 
 -------------------------------------------------------------------------------
 Matlab is a Registered Trademark of The Mathworks, Inc.
 
 Copyright (C) 2003-2005 Guillaume Flandin <>
