## Supplemental File 1 for "A biophysical model of striatal microcircuits suggests delta/theta-rhythmically interleaved gamma and beta oscillations mediate periodicity in motor control": INSTALL

 =============================================================================
       3FRAMES - Template files for use with M2HTML documentation system
 =============================================================================

 0/ Requirements:
    - Matlab 5.3 or above
	  - Operating system: any
    - the source for M2HTML as provided by 
      <http://www.artefact.tk/software/matlab/m2html/>
      See the package's INSTALL and README files for more information
      Let's assume you installed the package to /home/foo/matlab/m2html
      (and lets call that directory M2HTML_PATH from now on

 2/ Extract the files for 3FRAMES into the template directory
       cd M2HTML_PATH/templates
       unzip 3frames.zip

 3/ That's it....
   
 A documentation is available in the README file.
   
===============================================================================
===============================================================================
 Matlab is a Registered Trademark of The Mathworks, Inc.
 M2HTML: Copyright (C) 2003 Guillaume Flandin <>
 3FRAMES: Copyright (C) 2003-2005 
          Guillaume Flandin <>
          Lorenz Gerstmayr <>
