## Supplemental File 1 for "A biophysical model of striatal microcircuits suggests delta/theta-rhythmically interleaved gamma and beta oscillations mediate periodicity in motor control": mdir.tpl

Index for Directory {MDIR}


### Index for directory "{MDIR}"

#### Matlab files in this directory:

- {NAME}     -  {H1LINE}


#### Other Matlab-specific files in this directory:

- {OTHERFILE}


#### Top directories:

- Toplevel directory
- Main directory

#### Subsequent directories:

- {SUBDIRECTORY}


#### Dependency Graph

- View the Graph.


#### TODO List

- View the TODO list.


---

Generated on {DATE} by **m2html** using template **3frames** © Guillaume Flandin, Lorenz Gerstmayr, 2003-2005
