## Supplemental File 1 for "A biophysical model of striatal microcircuits suggests delta/theta-rhythmically interleaved gamma and beta oscillations mediate periodicity in motor control": mfile.tpl

Description of {NAME}


### Documentation for "{NAME}"

#### PURPOSE

**{H1LINE}**

#### SYNOPSIS

**{SYNOPSIS}  This is a script file.**

#### DESCRIPTION

```
{DESCRIPTION}
```

#### CROSS-REFERENCE INFORMATION

This function calls:


- {NAME\_CALL} {H1LINE\_CALL}

This function is called by:

- {NAME\_CALLED} {H1LINE\_CALLED}

#### SUBFUNCTIONS

- {SUB}


#### SOURCE CODE

```
{SOURCECODE}
```


---

Generated on {DATE} by **m2html** using template **3frames** © Guillaume Flandin, Lorenz Gerstmayr, 2003-2005
