## Supplemental File 1 for "A biophysical model of striatal microcircuits suggests delta/theta-rhythmically interleaved gamma and beta oscillations mediate periodicity in motor control": README

 =============================================================================
       3FRAMES - Template files for use with M2HTML documentation system
 =============================================================================

 M2HTML:
=========
 The M2HTML toolbox is intended to provide automatic generation of M-files 
 documentation in HTML. It reads each M-file in a set of directories
 (eventually recursively) to produce a corresponding HTML file containing
 synopsis, H1 line, help, function calls and called functions with 
 hypertext links, syntax highlighted source code with hypertext, ...

 For more information see the documentation provided with the M2HTML package.
 It should be available at http://www.artefact.tk/software/matlab/m2html/

 3FRAMES:
==========
  3FRAMES is a set of templates for use with the M2HTML package to produce 
  documentation for MATLAB code. It is based on the FRAME template provided by 
  M2HTML. 3FRAMES divides the browser window into 3 frames. The left frame 
  always shows a list of MATLAB-code directories. The directories are linked to
  a detailed function list (much like a CONTENTS.M file) of the corresponding 
  directory shown in the upper right frame. The function list also contains a
  short description (H1LINE) for each function as well as links to subsequent
  and toplevel directories. The detailed documentation for each function is 
  shown in the lower right frame.
 
 INSTALLATION:
===============
 
 Please read the INSTALL file provided with 3FRAMES for installation 
 instructions.
 
 TUTORIAL:
===========
 
 Using 3FRAMES is fairly easy once you installed it. Just tell M2HTML to use
 the 3FRAMES templates by setting its template option to '3frames'.

 Here is an example call to M2HTML using 3FRAMES:

 -----------------------------------------------------------------------------
 htmldir = ['/home/' getenv('USER') '/develop/matlab/doc'];
 mdir = 'matlab/';

 m2html('mfiles', mdir,...
        'htmldir', htmldir,...
        'recursive', 'on',...
        'template', '3frames',...
        'index', 'menu',...
        'global', 'on');
 -----------------------------------------------------------------------------

 3FRAMES is supposed to work with all other options provided by M2HTML. See 
 the README file delivered with M2HTML for a comprehensive list of options or
 type HELP M2HTML.

 KNOWN ISSUES:
===============
 None (so far...)

 TODO:
=======
 Have fun using the package...
 
===============================================================================
===============================================================================
 Matlab is a Registered Trademark of The Mathworks, Inc.
 M2HTML: Copyright (C) 2003 Guillaume Flandin <>
 3FRAMES: Copyright (C) 2003-2005 
          Guillaume Flandin <>
          Lorenz Gerstmayr <>
