## Supplemental File 1 for "A biophysical model of striatal microcircuits suggests delta/theta-rhythmically interleaved gamma and beta oscillations mediate periodicity in motor control": todo.tpl

To Do List for {MDIR}


 Master index 

### TODO list for {MDIR}

#### {MFILE}:

- line {NBLINE}:  {COMMENT}


---

Generated on {DATE} by **m2html** using template **3frames** © Guillaume Flandin, Lorenz Gerstmayr, 2003-2005
