## Supplemental File 1 for "A biophysical model of striatal microcircuits suggests delta/theta-rhythmically interleaved gamma and beta oscillations mediate periodicity in motor control": doxysearch.php

```
<?php
/******************************************************************************
 *
 * $Id:$
 *
 * Copyright (C) 1997-2003 by Dimitri van Heesch.
 *
 * Permission to use, copy, modify, and distribute this software and its
 * documentation under the terms of the GNU General Public License is hereby 
 * granted. No representations are made about the suitability of this software 
 * for any purpose. It is provided "as is" without express or implied warranty.
 * See the GNU General Public License for more details.
 *
 */

function readInt($file)
{
  $b1 = ord(fgetc($file)); $b2 = ord(fgetc($file));
  $b3 = ord(fgetc($file)); $b4 = ord(fgetc($file));
  return ($b1<<24)|($b2<<16)|($b3<<8)|$b4;
}

function readString($file)
{
  $result="";
  while (ord($c=fgetc($file))) $result.=$c;
  return $result;
}

function readHeader($file)
{
    $header =fgetc($file); $header.=fgetc($file);
    $header.=fgetc($file); $header.=fgetc($file);
    return $header;
}

function computeIndex($word)
{
  if (strlen($word)<2) return -1;
  // high char of the index
  $hi = ord($word{0});
  if ($hi==0) return -1;
  // low char of the index
  $lo = ord($word{1});
  if ($lo==0) return -1;
  // return index
  return $hi*256+$lo;
}

function search($file,$word,&$statsList)
{
  $index = computeIndex($word);
  if ($index!=-1) // found a valid index
  {
    fseek($file,$index*4+4); // 4 bytes per entry, skip header
    $index = readInt($file);
    if ($index) // found words matching first two characters
    {
      $start=sizeof($statsList);
      $count=$start;
      fseek($file,$index);
      $w = readString($file);
      while ($w)
      {
        $statIdx = readInt($file);
        if ($word==substr($w,0,strlen($word)))
        { // found word that matches (as substring)
          $statsList[$count++]=array(
              "word"=>$word,
              "match"=>$w,
              "index"=>$statIdx,
              "full"=>strlen($w)==strlen($word),
              "docs"=>array()
              );
        }
        $w = readString($file);
      }
      $totalFreq=0;
      for ($count=$start;$count<sizeof($statsList);$count++)
      {
        $statInfo = &$statsList[$count];
        fseek($file,$statInfo["index"]); 
        $numDocs = readInt($file);
        $docInfo = array();
        // read docs info + occurrence frequency of the word
        for ($i=0;$i<$numDocs;$i++)
        {
          $idx=readInt($file); 
          $freq=readInt($file); 
          $docInfo[$i]=array("idx"=>$idx,"freq"=>$freq,"rank"=>0.0);
          $totalFreq+=$freq;
          if ($statInfo["full"]) $totalfreq+=$freq;
        }
        // read name an url info for the doc
        for ($i=0;$i<$numDocs;$i++)
        {
          fseek($file,$docInfo[$i]["idx"]);
          $docInfo[$i]["name"]=readString($file);
          $docInfo[$i]["url"]=readString($file);
        }
        $statInfo["docs"]=$docInfo;
      }
      for ($count=$start;$count<sizeof($statsList);$count++)
      {
        $statInfo = &$statsList[$count];
        for ($i=0;$i<sizeof($statInfo["docs"]);$i++)
        {
          $docInfo = &$statInfo["docs"];
          // compute frequency rank of the word in each doc
          $statInfo["docs"][$i]["rank"]=
            (float)$docInfo[$i]["freq"]/$totalFreq;
        }
      }
    }
  }
  return $statsList;
}

function combine_results($results,&$docs)
{
  foreach ($results as $wordInfo)
  {
    $docsList = &$wordInfo["docs"];
    foreach ($docsList as $di)
    {
      $key=$di["url"];
      $rank=$di["rank"];
      if (in_array($key, array_keys($docs)))
      {
        $docs[$key]["rank"]+=$rank;
        $docs[$key]["rank"]*=2; // multiple matches increases rank 
      }
      else
      {
        $docs[$key] = array("url"=>$key,
            "name"=>$di["name"],
            "rank"=>$rank
            );
      }
      $docs[$key]["words"][] = array(
               "word"=>$wordInfo["word"],
               "match"=>$wordInfo["match"],
               "freq"=>$di["freq"]
               );
    }
  }
  return $docs;
}

function normalize_ranking(&$docs)
{
  $maxRank = 0.0000001;
  // compute maximal rank
  foreach ($docs as $doc) 
  {
    if ($doc["rank"]>$maxRank)
    {
      $maxRank=$doc["rank"];
    }
  }
  reset($docs);
  // normalize rankings
  while (list ($key, $val) = each ($docs)) 
  {
    $docs[$key]["rank"]*=100/$maxRank;
  }
}

function filter_results($docs,&$requiredWords,&$forbiddenWords)
{
  $filteredDocs=array();
  while (list ($key, $val) = each ($docs)) 
  {
    $words = &$docs[$key]["words"];
    $copy=1; // copy entry by default
    if (sizeof($requiredWords)>0)
    {
      foreach ($requiredWords as $reqWord)
      {
        $found=0;
        foreach ($words as $wordInfo)
        { 
          $found = $wordInfo["word"]==$reqWord;
          if ($found) break;
        }
        if (!$found) 
        {
          $copy=0; // document contains none of the required words
          break;
        }
      }
    }
    if (sizeof($forbiddenWords)>0)
    {
      foreach ($words as $wordInfo)
      {
        if (in_array($wordInfo["word"],$forbiddenWords))
        {
          $copy=0; // document contains a forbidden word
          break;
        }
      }
    }
    if ($copy) $filteredDocs[$key]=$docs[$key];
  }
  return $filteredDocs;
}

function compare_rank($a,$b)
{
  return ($a["rank"]>$b["rank"]) ? -1 : 1; 
}

function sort_results($docs,&$sorted)
{
  $sorted = $docs;
  usort($sorted,"compare_rank");
  return $sorted;
}

function report_results(&$docs)
{
  echo "<table cellspacing=\"2\">\n";
  echo "  <tr>\n";
  echo "    <td colspan=\"2\"><h2>Search Results</h2></td>\n";
  echo "  </tr>\n";
  $numDocs = sizeof($docs);
  if ($numDocs==0)
  {
    echo "  <tr>\n";
    echo "    <td colspan=\"2\">".matches_text(0)."</td>\n";
    echo "  </tr>\n";
  }
  else
  {
    echo "  <tr>\n";
    echo "    <td colspan=\"2\">".matches_text($numDocs);
    echo "\n";
    echo "    </td>\n";
    echo "  </tr>\n";
    $num=1;
    foreach ($docs as $doc)
    {
      echo "  <tr>\n";
      echo "    <td align=\"right\">$num.</td>";
      echo     "<td><a class=\"el\" href=\"".$doc["url"]."\">".$doc["name"]."</a></td>\n";
      echo "  <tr>\n";
      echo "    <td></td><td class=\"tiny\">Matches: ";
      foreach ($doc["words"] as $wordInfo)
      {
        $word = $wordInfo["word"];
        $matchRight = substr($wordInfo["match"],strlen($word));
        echo "<b>$word</b>$matchRight(".$wordInfo["freq"].") ";
      }
      echo "    </td>\n";
      echo "  </tr>\n";
      $num++;
    }
  }
  echo "</table>\n";
}

function matches_text($num)
{
  if ($num==0)
  {
    return 'Sorry, no documents matching your query.';
  }
  else if ($num==1)
  {
    return 'Found 1 document matching your query.';
  }
  else // $num>1
  {
    return 'Found '.$num.' documents matching your query. Showing best matches first.';
  }
}

function main($idxfile)
{
  if(strcmp('4.1.0', phpversion()) > 0) 
  {
    die("Error: PHP version 4.1.0 or above required!");
  }
  if (!($file=fopen($idxfile,"rb"))) 
  {
    die("Error: Search index file could NOT be opened!");
  }
  if (readHeader($file)!="DOXS")
  {
    die("Error: Header of index file is invalid!");
  }
  $query="";
  if (array_key_exists("query", $_GET))
  {
    $query=$_GET["query"];
  }
  $results = array();
  $requiredWords = array();
  $forbiddenWords = array();
  $foundWords = array();
  $word=strtolower(strtok($query," "));
  while ($word) // for each word in the search query
  {
    if (($word{0}=='+')) { $word=substr($word,1); $requiredWords[]=$word; }
    if (($word{0}=='-')) { $word=substr($word,1); $forbiddenWords[]=$word; }
    if (!in_array($word,$foundWords))
    {
      $foundWords[]=$word;
      search($file,$word,$results);
    }
    $word=strtolower(strtok(" "));
  }
  $docs = array();
  combine_results($results,$docs);
  // filter out documents with forbidden word or that do not contain
  // required words
  $filteredDocs = filter_results($docs,$requiredWords,$forbiddenWords);
  // normalize rankings so they are in the range [0-100]
  normalize_ranking($filteredDocs);
  // sort the results based on rank
  $sorted = array();
  sort_results($filteredDocs,$sorted);
  // report results to the user
  report_results($sorted);
  fclose($file);
}

?>
```
