## Supplemental File 1 for "A biophysical model of striatal microcircuits suggests delta/theta-rhythmically interleaved gamma and beta oscillations mediate periodicity in motor control": master.tpl

Matlab Index


### Matlab Index

#### Matlab Directories

- {DIR}

#### Matlab Files found in these Directories

{IDNAME}

#### Search Engine

Search for 


#### Dependency Graph

- View the Graph.


---

Generated on {DATE} by **m2html** © 2005
