## Supplemental File 1 for "A biophysical model of striatal microcircuits suggests delta/theta-rhythmically interleaved gamma and beta oscillations mediate periodicity in motor control": mdir.tpl

Index for Directory {MDIR}


|  |  |
| --- | --- |
| Master index | Index for {MDIR} |

### Index for {MDIR}

#### Matlab files in this directory:

|  |  |
| --- | --- |
| {NAME} | {H1LINE} |

#### Other Matlab-specific files in this directory:

- {OTHERFILE}


#### Subsequent directories:

- {SUBDIRECTORY}


#### Dependency Graph

- View the Graph.


#### TODO List

- View the TODO list.


---

Generated on {DATE} by **m2html** © 2005
