## Supplemental File 1 for "A biophysical model of striatal microcircuits suggests delta/theta-rhythmically interleaved gamma and beta oscillations mediate periodicity in motor control": mdir.tpl

Index for Directory {MDIR}


 Master index 

### Index for {MDIR}

#### Matlab files in this directory:

- {NAME}

#### Other Matlab-specific files in this directory:

- {OTHERFILE}


#### Subsequent directories:

- {SUBDIRECTORY}


#### Dependency Graph

- View the Graph.


#### TODO List

- View the TODO list.


---

Generated by **m2html** © 2005
