## Supplemental File 1 for "A biophysical model of striatal microcircuits suggests delta/theta-rhythmically interleaved gamma and beta oscillations mediate periodicity in motor control": TODO

 M2HTML - Documentation System for Matlab .m files in HTML
 =========================================================
 
 Wish list:
 ==========
   o special handling of Matlab classes and private directories
   o special handling of Java code
   o special handling of MEX files (C/C++ or Fortran) and P-files
   o special handling of Simulink (*.mdl) files (see load_system,open_system,get/set_param)
   o PDF output (LaTeX), Rich Text Format (RTF) output ?
   o search engine (in PHP, following Doxygen)
   o Javascript menu for lists of files (http://www.treemenu.com/ or http://www.treeview.net/)
   o detect urls (http, ftp) and mailto in source code and create links
   o 'previous function' and 'next function' links in templates
   o special template compatible with Matlab helpwin
   o create another template example looking like Chora (http://cvs.php.net/)
   o do not display any list in cross-references if empty
   o set H1 to the first *not empty* line in help
   o hierarchical graph of directories ?
   o list of used keywords using <fparser.m> ?
   o use <depfun> in <private/mfileparse> to distinguish functions or variables 
     with same names and other subtilities
   o 'end' is sometimes badly identified (end,:) or (2:end,1)
   o detect 'Usage' or 'Syntax' in help
   o if 'download' on and 'recursive' on, if m2html launched twice, if htmldir
     in mfiles, then it will also document m-files to be downloaded...
   o copyfile may wait for a <return> if the file still exists (on Linux)
   + optional link to original M-file (implemented with copies of M-files, maybe
     a link to their original location would be enough (but not 'portable').
   o creation of the output directories may be buggy because of <exist> which
     also check the existence in the Matlab path (never observed bug at that time)
   o add a 'rootdir' option like in <zip> to specify the output dir (pwd).
   + if 'global' and 'graph' are 'on', then there should be a link in master
     directory file for a full dependency graph with hypertext links
   o build a graphical interface for the fill in of the arguments.
   o toolbbar icons for the GUI should be saved in m2htmltoolbarimages.mat
     in a struct array with fields as help, new, open, save, wheel, ...
	 (as in camtoolbarimages.mat)
   o lines should have an anchor <a name="nbligne"></a> so that ToDO links go 
     directly to the requested line
   o H1Lines in A HREF title are printed without using <entity> to escape reserved
     characters (this bug may appear everywhere text is sent to output without calling
	 <entity> on it.
   o would be great if using written functions, it would be possible to write a function
     that would return a .tex file containing documentation of a single file, or maybe
	 there should be a M2LaTeX that would do the same as m2html
   o add a page with source code statistics : nb folders, classes, files, functions, lines
   o allow to add options when calling <dot> (resize image output, etc)
   o support different languages (english, french, german, ...)
   o remove <private/doxyread.m> : only used for debug purpose, replaced by <private/doxysearch.m>
   o explain in the online tutorial / FAQ how does the PHP search engine work
   o Bugs in the search index with upper/lower case ?
   o explain in the online tutorial the several ways to give parameters : struct, key/value
     and the possibility of using short keys as in get/set
   + .map file created with the full dependency graph does not handle directory changes
   + prevent or display a warning when an html matlab file overwrite the 'index' file
   o explain in FAQ what to do when a warning index file overwritten appears...
   o creating a ToDo list including all the directories rather than only one by directory
   o See Also functions are with hypertext only if in the same directory : should be hypertexted
     when 'global' is on
   o Add an option to automatically generate a Contents.m file in M-files directory
   o ability to parse template headers as proposed by Denis Gilbert ?
   o See if <setdiff.m> or <setxor.m> or <unique.m> or <intersect.m> might be useful for 'search'
   o Modify <copyfile.m> call to overcome the Matlab bug:
     http://www.mathworks.com/support/solutions/data/1-1B5JY.html
   o See what to do with new R14 functions : <publish.m>, ...
   o should I move <mdot.m> in the private folder ?
   o extract LateX string in code and display them as images using <teximage.m> from MatlabCentral
     (or use MathML)
   o see http://www.mathworks.com/support/solutions/data/1-18U6Q.html?solution=1-18U6Q
     for adding third-party documentation to the MATLAB Help System and generate index.db
   + check that the command line invoking dot works with directories with spaces => added ""
   + Matlab 7 has an old dot version (1.8) which does not handle png : add in FAQ what to do...
   o there's a bug remaining with space handling in directories name
     on Unix, one can use '\' to escape them. See what to do on Windows.
   o get dependencies from callbacks in GUI ('Callback', 'ClickedCallback', 'CreateFcn', 'DeleteFcn', 
     'OffCallback', 'OnCallback')...
   o replace 'exist(c) == 2' by 'exist(c,'file')' and 'exist(d,'dir')'
     to work with files and directories which have same name as built-in
	 (thanks to Alois Schloegl). Need to do it everywhere <exist> is used ?
	 Reverted for the moment because it then didn't work properly for unknown reasons.
   o propose to split long lines with '...' (see <breaklines.m> from Michael Wild)
   o have a look to 'SubFuns' in MatlabCentral, and hyperlinks in the command window (matlab7)
   o handle preprocessor statements used by the Matlab Compiler (from Robert van Kuijk)
     %#function checkconn checkinput checknargin
   o remove the last <what> call (to find other Matlab-specific files)
   o <doc> use 'matlab:' as a protocol to launch Matlab code from a browser
   o create a <helptoc.xml> to integrate in the Matlab help system (request from Stefan Mangard)
   o label object directories '@something' a class and inside functions methods.
   o group functions using structure provided in <Contents.m> (request from Stefan Mangard)
   o bug in handling directories having spaces in them (on windows)?
   o having list of all files with the frame version? (request from Brian Wandell)
   o in the frame version, can only go to the top or down but not one level ahead
   + specify a list of directory names one can exclude from the file generation (request from Brian Wandell)
   o hyperlinks should be present for the whole 'Contents.m' file instead of after 
     'see also' only (request of Piotr Dollar)
   o changing two pages in a frame at the same time with http://goldray.com/frames/chng2-nest.htm
   o if only a file change (or detected automatically) then don't reparse everything.
     like the make command or using md5 hash (request from Hitashyam Maka)
   o exclude a directory when generating cross-referenced html pages of m-files (request from Marcus Wilms)
   o the field purpose might display again the name of the function (from h1line) (request from M. Buehren)
     Sol: replace tpl = set(tpl,'var','H1LINE', entity(h1line{j})); by
     >> h1linetmp = entity(h1line{j});
     >> if ~isempty(strmatch(lower(names{j}), lower(h1linetmp)))
     >>    h1linetmp = h1linetmp(length(names{j})+1:end);
     >> end
     >> tpl = set(tpl,'var','H1LINE', h1linetmp);
   o have an "ignore" feature to exclude certain directories or files from the documentation. (request from M. Buehren)
   o full dependency graph (with global option) may display several identical arrows between two functions if
     two directories have files with same name (typical for classes) (detected by Tomasz Reczek)
   o function call graph for which files/functions share the same globals (request from Bj�rn Gustavsson)
   o It would be good if the menu.html /index.html could use that file as a guide-line to sort the links.
     Now one has to do that resorting by hand after running m2html. (request from Bj�rn Gustavsson)
   
 -------------------------------------------------------------------------------
 Matlab is a Registered Trademark of The Mathworks, Inc.

 Copyright (C) 2003-2005 Guillaume Flandin <>
