## Supplemental File 1 for "A biophysical model of striatal microcircuits suggests delta/theta-rhythmically interleaved gamma and beta oscillations mediate periodicity in motor control": MapN.m

```
classdef MapN < handle
    % A multidimensional map container
    %
    % A MapN object contains key lists and values. A value is some unit of
    % data that you want stored in the MapN object, and a key list is a
    % unique reference to that data.
    %
    % MapN is modelled on containers.Map, but whereas a containers.Map
    % object is indexed by a single scalar or string key, a MapN object is
    % indexed by an ordered set of such keys, expressed as a
    % comma-separated list of arguments.
    %
    % MapN is a handle class.
    %
    % Description - basic outline
    % ---------------------------
    %
    % A MapN object is constructed like this:
    %
    %   M = MapN();
    %
    % Values are stored using M(key1, key2, ...) = value, for example:
    %
    %   M(1, 'a')     = 'a string value';
    %   M(1, 'b')     = 287.2;
    %   M(2)          = [1 2 3; 4 5 6]; 
    %   M(2, 'x', pi) = {'a' 'cell' 'array'};
    %
    % and values are retrieved using M(key1, key2, ...), for example
    %
    %   v = M(1, 'b');
    %   u = M(2);
    %
    % Key lists and key types
    % -----------------------
    %
    % At least one key must be given in a key list. (The empty key list
    % could logically be allowed, were it not that expressions of the form
    % M() = v generate syntax errors.)
    %
    % Apart from this, a key list may be of any length. Key lists of
    % different lengths are independent, so M('a') and M('a', 'b') may both
    % be given values, and will not interact.
    %
    % Each key must be of a type supported by containers.Map. This includes
    % numerical scalars and strings. A given key list may contain a mixture
    % of different key types; however, type consistency between different
    % key lists is required.
    %
    % A simple rule is that there will be no error if the same key type is
    % used consistently at each position in the key list. In the example
    % above, position 1 uses a double, position 2 uses a string, and
    % position 3 uses a double. An attempt to use a string as a key in
    % position 1 is inconsistent and throws an exception:
    %
    %   M('a', 'c') = 'v';       % error!
    %
    % The type of the key at a given position is established by the first
    % use of a key at that position. (There is no equivalent to the
    % 'KeyType' parameter of containers.Map.)
    %
    % In fact, the restriction on key types is more liberal than the simple
    % rule implies. An exact statement is that if two key lists have the
    % same length and are identical in the first N positions, then the two
    % keys at position N+1 must have the same type.
    %
    % Value types
    % -----------
    %
    % By default, values may be of any type and value types may be mixed
    % freely in a MapN object.
    %
    % There may be an efficiency advantage in restricting the value types
    % if a large number of strings, or scalar values of the same type, are
    % to be stored. The restriction is put in place in the constructor
    % thus:
    %
    %   Muni = MapN('uniformvalues', true);
    %
    % If this is done, stored values must be scalars or strings, and they
    % cannot be arbitrarily mixed. The rule is that if two key lists are
    % identical apart from the final key in each, then the two
    % corresponding values must such that they can both be assigned to a
    % containers.Map object which has had 'uniformvalues' set to true. The
    % type is determined by the first value stored. (There is no equivalent
    % to the 'ValueType' parameter of containers.Map.) Numerical values may
    % be automatically converted to the type of previously stored values.
    %
    % The method uniformValues returns the setting of the 'uniformvalues'
    % parameter.
    %
    % Invalid key lists and default values
    % ------------------------------------
    %
    % It is possible to test whether a key list has been stored using
    % isKey. For example
    %
    %   b = isKey(M, 2, 'x');
    %
    % assigns true to b if M(2, 'x') holds a value, and false otherwise.
    % isKey returns false if a key has an invalid type (as for
    % containers.Map).
    %
    % By default, an exception will be thrown if an attempt is made to
    % access a MapN object using a key list which has not been stored. For
    % example:
    %
    %   M = MapN();
    %   M(2, 'a') = 123;
    %   v = M(2, 'a');     % OK
    %   u = M(2, 'b');     % error!
    %
    % This may be changed by using the 'default' parameter in the
    % constructor:
    %
    %   Mdef = MapN('default', 'zzz');
    %
    % If a default is set, then the default value, which may have any type,
    % is returned for any key list for which no value has been stored:
    %
    %   u = Mdef(2, 'b');    % u gets value 'zzz', no error
    %
    % The types of keys in the key list must still be consistent with the
    % types of keys in the key lists that have been stored.
    %
    % The default value may be set or changed for an existing MapN object
    % using
    %
    %   setDefault(M, 'defaultvalue');
    %
    % and the default may be removed (so access with invalid keys will
    % cause errors) using:
    %
    %   noDefault(M);
    %
    % The method hasDefault(M) tells whether M has a default, and if it
    % has, getDefault(M) returns the current default value.
    %
    % Updating and removing entries
    % -----------------------------
    %
    % The value for a given key list is updated using the usual assignment;
    % the previous value is overwritten.
    %
    %   M(pi, 'x') = 1;     % 1 is current value
    %   M(pi, 'x') = 2;     % 2 replaces 1 as the value for this key list
    %
    % A key list and its corresponding value may be deleted using the
    % remove method, for example:
    %
    %   remove(M, pi, 'x');
    %   u = M(pi, 'x');     % u gets default value, or there's an error
    %
    % If an attempt is made to remove a key which is not stored, an
    % exception is thrown. (This is different to containers.Map, which
    % issues a warning. The advantage of an exception is that it can be
    % caught.)
    %
    % Multiple key lists and values
    % -----------------------------
    %
    % For the methods in this section, a key list is represented as a cell
    % array of keys. A set of key lists is a cell array of cell arrays.
    %
    % A set of key lists and a set of values can be supplied to the
    % constructor to set up some initial associations. For example
    %
    %   M = MapN({{1 'a'} {1 'b'} {2} {2 'x' pi}}, ...
    %    {'a string' 287.2 [1 2 3; 4 5 6] {'a' 'cell' 'array'}});
    %
    % is equivalent to
    %
    %   M = MapN();
    %   M(1, 'a')     = 'a string';
    %   M(1, 'b')     = 287.2;
    %   M(2)          = [1 2 3; 4 5 6];
    %   M(2, 'x', pi) = {'a' 'cell' 'array'};
    %
    % Multiple values may be retrieved with the values method. For example:
    %
    %   vals = values(M, {{2}, {1, 'a'}});
    %
    % assigns a cell array to vals such that vals{1} is [1 2 3; 4 5 6] and
    % vals{2} is 'a string'.
    %
    % If the second argument is omitted, values returns all the values:
    %
    %   vals = values(M);   % vals is a 1x4 cell array
    % 
    % Multiple values can be stored with storeValues; for example, the
    % first two assignments above are equivalent to:
    %
    %   storeValues(M, {{1 'a'} {1 'b'}}, {'a string' 287.2});
    %
    % All the key lists stored in an object may be got with:
    %
    %   klists = keys(M);
    %
    % The order of the result is consistent with the order of values, so
    % values(M) is the same as values(M, keys(M)).
    %
    % The isKey function can be used with a set of key lists, and returns a
    % logical array with true in the positions of the valid key lists:
    %
    %   b = isKey(M, {{1 'a'} {'spqr' 17 pi} {2}});
    %
    % sets b to logical([1 0 1]).
    %
    % Other methods
    % -------------
    %
    % The number of key lists stored may be obtained with the length
    % method. (The Count property available in containers.Map is not
    % supported; use length(M) instead.)
    %
    % The size method returns the number of key lists stored as the first
    % dimension, and 1 for all other dimensions.
    %
    % Vertical concatenation is supported (as for containers.Map). If M1,
    % M2 and M3 are MapN objects, and we execute, for example
    %
    %   Mcomb = [M1; M2; M3];
    %
    % then Mcomb is a single MapN object which stores all the key lists
    % from M1, M2 and M3 and their associated values. If a key list is
    % duplicated, then the last value associated with it by M1, M2 or M3,
    % in that order, is stored in Mcomb. If any of M1, M2 or M3 has a
    % default value, then Mcomb has a default whose value is the last one
    % encountered in M1, M2 or M3.
    %
    % Mcomb is constructed with an 'uniformvalues' parameter equal to that
    % of M1. If M1 has 'uniformvalues' set to true, and any of the input
    % maps contain values that have inconsistent types, an error will
    % result.
    %
    % Horizontal concatenation is not supported.
    %
    % Copying a MapN object
    % ---------------------
    %
    % Because this is a handle class, Mnew = M does not make a true copy,
    % so updates to M will affect Mnew, and vice versa. To make an
    % independent copy of M, you can use:
    %
    %   Mnew = copy(M);
    %
    % Method call syntax
    % ------------------
    %
    % Methods of MapN must be called using the syntax func(MapNobj, ...),
    % not MapNobj.func(...).
    %
    % Methods and properties
    % ----------------------
    %
    % MapN methods:
    %   MapN        - constructor for MapN objects
    %   subsref     - implements value = Mobj(keylist)
    %   subsasgn    - implements M(keylist) = value
    %   isKey       - determine whether given key lists are stored
    %   keys        - get all the stored keys
    %   values      - get stored values
    %   storeValues - store multiple keys and values
    %   remove      - remove stored values
    %   hasDefault  - determine whether a MapN has a default value
    %   getDefault  - get the default value
    %   noDefault   - unset any default value
    %   setDefault  - set the default value
    %   uniformValues - get the value of the 'uniformvalues' property
    %   length      - get the number of key lists stored
    %   size        - get the number of key lists stored
    %   isempty     - determine whether any key lists are stored
    %   vertcat     - implement [Mobj1; Mobj2]
    %   horzcat     - error on [Mobj1, Mobj2]
    %   copy        - copy a MapN object
    %
    % This class has no publicly accessible properties.
    %
    % See also: containers.Map
    
    % Copyright David Young 2011
    
    
    properties (Access = private)
        % Root map implemented as a cell array rather than a Map
        % as indexed by no. of arguments, and m() = v is forbidden by
        % syntax restriction
        rootMap = {};
        
        % This is a private property accessed via a method, rather than a
        % publicly accessible property because of the complexity of
        % providing correct obj.name indexing when subsref/subsasgn are
        % overriden.
        Ct = 0;
        
        doDefault;  % boolean, whether to supply a default value
        default;    % value of default
        
        univals;    % 'uniformvalues' parameter for new Maps 
    end
    
    methods
        
        function M = MapN(varargin)
            % constructs MapN objects
            %
            % M = MapN() constructs an empty MapN object.
            %
            % M = MapN(KEYLISTS, VALUES) constructs a MapN object which
            % contains the specified key lists and values. KEYLISTS is a
            % cell array; each element of this is a cell array containing
            % the keys making up a key list. VALUES is a cell array
            % containing the values. KEYLISTS and VALUES must have the same
            % number of elements and VALUES{k} is associated with
            % KEYLISTS{k} in M.
            %
            % M = MapN(..., 'default', d) sets the default value returned
            % by M to d.
            %
            % M = MapN(..., 'uniformvalues', tf), where tf is a logical
            % scalar, sets the 'uniformvalues' properties of the underlying
            % containers.Map objects. This restricts variation of the types
            % of the values stored in M.
            %
            % Additional values are added to M using
            %
            %   M(key1, key2, ...) = val;
            %
            % and values are retrieved from M using
            %
            %   val = M(key1, key2, ...);
            %
            % See the class information for details and examples.
            %
            % See also: MapN
            
            ip = inputParser;
            ip.addOptional('keylist', {}, @iscell);
            ip.addOptional('vallist', {}, @iscell);
            ip.addParamValue('default', []);
            ip.addParamValue('uniformvalues', false, ...
                @(x) islogical(x) && isscalar(x));
            ip.parse(varargin{:});
            
            M.doDefault = ~ismember('default', ip.UsingDefaults);
            M.default = ip.Results.default;
            
            M.univals = ip.Results.uniformvalues;
            
            if ~isempty(ip.Results.keylist)
                storeValues(M, ip.Results.keylist, ip.Results.vallist);
            end
        end
                
        
        function v = subsref(M, S)
            % returns value associated with key list
            %
            % Implements the syntax
            %
            %   value = MapNobj(key1, key2, ...)
            %
            % See also: MapN, MapN/subsasgn, MapN/values
            
            if ~isscalar(S) || ~strcmp(S.type, '()')
                error('MapN:Subsref:LimitedIndexing', ...
                    'Only ''()'' indexing is supported by a MapN');
            end
            
            try
                v = subsrefAction(M, S.subs);
            catch me
                % default is handled in subsrefError for efficiency
                v = subsrefError(M, me, S.subs);
            end
        end
        
        
        function M = subsasgn(M, S, v)
            % sets value associated with key list
            %
            % Implements the syntax
            %
            %   MapNobj(key, key2, ...) = value
            %
            % See also: MapN, MapN/subsref, MapN/storeValues
            
            if ~isscalar(S) || ~strcmp(S.type, '()')
                error('MapN:Subsasgn:LimitedIndexing', ...
                    'Only ''()'' indexing is supported by a MapN');
            end
            
            try
                M = subsasgnAction(M, S.subs, v);
            catch me
                subsasgnError(M, me, S.subs);
            end            
        end
        
        
        function remove(M, varargin)
            % deletes keylist/value pairs
            %
            % remove(MAPNOBJ, KEY1, KEY2, ...) removes the key list and its
            % associated value from MAPNOBJ.
            %
            % remove(MAPNOBJ, KEYLISTS) removes multiple key lists
            % and values. KEYLISTS is a cell array of cell arrays, each of
            % which contains a list of keys.
            %
            % See also: MapN
            
            ss = varargin;
            nargs = length(ss);
            
            if nargs == 1 && iscell(ss{1})
                
                c = ss{1};
                for ii = 1:numel(c)
                    remove(M, c{ii}{:});
                end
                
            else
                
                try
                    m = M.rootMap{nargs};
                    sx(m, 1);
                    if isempty(m)
                        M.rootMap{nargs} = [];
                    end
                    
                    M.Ct = M.Ct - 1;
                catch me
                    removeError(M, me, ss);
                end
                
            end
            
            function sx(m, ind)
                if ind == nargs
                    m(ss{ind});  % prompt error if not a valid key
                    remove(m, ss{ind});
                else
                    mm = m(ss{ind});
                    sx(mm, ind+1);
                    if isempty(mm)
                        remove(m, ss{ind});
                    end
                end
            end
            
        end
        
        
        function b = isKey(M, varargin)
            % tests whether keys are stored
            %
            % B = isKey(MAPNOBJ, KEY1, KEY2, ...) returns a logical scalar
            % value indicating whether the keylist is stored in the MapN
            % object.
            %
            % B = (MAPNOBJ, KEYLISTS) tests multiple key lists. KEYLISTS is
            % a cell array of cell arrays, each of which contains a list of
            % keys. B is a logical array such that B(k) is true if
            % KEYLISTS{k} is stored in MAPNOBJ.
            %
            % See also: MapN
            
            nargs = length(varargin);
            ss = varargin;
            
            if nargs == 0
                error('MapN:IsKey:emptyKeys', ...
                    'At least one key must be supplied.');
            end
            
            if nargs == 1 && iscell(ss{1})
            
                c = ss{1};
                b = false(size(c));
                for ii = 1:numel(c)
                    b(ii) = isKey(M, c{ii}{:});
                end
                
            else
                
                if nargs > length(M.rootMap) || isempty(M.rootMap{nargs})
                    b = false;
                else
                    b = si(M.rootMap{nargs}, 1);
                end
                
            end
            
            function b = si(m, ind)
                if ind == nargs
                    b = isKey(m,ss{ind});
                elseif isKey(m, ss{ind})
                    b = si(m(ss{ind}), ind+1);
                else
                    b = false;
                end
            end
        end
        
        
        function c = keys(M)
            % returns all the stored key lists
            %
            % KEYLISTS = keys(MAPNOBJ) returns a cell array containing all
            % the key lists stored in MAPNOBJ. KEYLISTS is a cell array of
            % cell arrays, each of which contains a list of keys.
            %
            % See also: MapN, MapN/values
            
            c = {};
            for nargs = 1:length(M.rootMap)
                if ~isempty(M.rootMap{nargs})
                    c = [c sk(M.rootMap{nargs}, nargs-1)]; %#ok<AGROW>
                end
            end
            
            function c = sk(m, argsleft)
                ks = keys(m);
                if argsleft
                    c = {};
                    for ii = 1:length(ks)
                        newks = sk(m(ks{ii}), argsleft-1);
                        newks = cellfun(@(newk) [ks(ii) newk], newks, ...
                            'UniformOutput', false);
                        c = [c newks]; %#ok<AGROW>
                    end
                else
                    c = cellfun(@(k) {k}, ks, 'UniformOutput', false);
                end
                
            end
            
        end
        
        
        function c = values(M, keysin)
            % returns stored values
            %
            % VALS = values(MAPNOBJ) returns all the stored values. VALS is
            % a cell array of values, stored in the same order as the
            % corresponding key lists in the result of keys(MAPNOBJ).
            %
            % VALS = values(MAPNOBJ, KEYLISTS) returns the values
            % corresponding to the given key lists. KEYLISTS is a cell
            % array of cell arrays, each of which contains a list of keys.
            %
            % See also: MapN, MapN/keys, MapN/subsref
            
            if nargin == 2
                
                c = cell(size(keysin));
                for kk = 1:numel(keysin)
                    try
                        c{kk} = subsrefAction(M, keysin{kk});
                    catch me
                        c{kk} = subsrefError(M, me, keysin{kk});
                    end
                end
                
            else
                
                c = {};
                for nargs = 1:length(M.rootMap)
                    if ~isempty(M.rootMap{nargs})
                        c = [c sv(M.rootMap{nargs}, nargs-1)]; %#ok<AGROW>
                    end
                end
                
            end
            
            function c = sv(m, argsleft)
                if argsleft
                    c = {};
                    ks = keys(m);
                    for ii = 1:length(ks)
                        c = [c sv(m(ks{ii}), argsleft-1)]; %#ok<AGROW>
                    end
                else
                    c = values(m);
                end
                
            end
            
        end
        
        
        function storeValues(M, keysin, valsin)
            % stores multiple values
            %
            % storeValues(MAPNOBJ, KEYLISTS, VALUES) stores the given key
            % lists and their corresponding values in MAPNOBJ. KEYLISTS is
            % a cell array of cell arrays, each of which contains a list of
            % keys. VALUES is a cell array of the corresponding values.
            %
            % See also: MapN, MapN/subsasgn
            
            if ~isequal(numel(keysin), numel(valsin))
                error('MapN:StoreValues:argLength', ...
                    'Keys and Values arguments must be same length');
            end
            
            for ii = 1:numel(keysin)
                try
                    M = subsasgnAction(M, keysin{ii}, valsin{ii});
                catch me
                    subsasgnError(M, me, keysin{ii});
                end
            end
        end
        
        
        function b = hasDefault(M)
            % tests for a default value
            %
            % B = hasDefault(MAPNOBJ) returns true if MAPNOBJ(BADKEYLIST)
            % returns a default value when BADKEYLIST is not stored, and
            % false if MAPNOBJ(BADKEYLIST) throws an exception.
            %
            % See also: MapN, MapN/getDefault, MapN/setDefault,
            % MapN/noDefault
            
            b = M.doDefault;
        end
   
        
        function d = getDefault(M)
            % returns the default value
            %
            % V = getDefault(MAPNOBJ) returns the default value, provided
            % that hasDefault(MAPNOBJ) returns true.
            %
            % See also: MapN, MapN/hasDefault, MapN/setDefault,
            % MapN/noDefault
            
            d = M.default;
        end
        
        
        function setDefault(M, d)
            % sets the default value
            %
            % setDefault(MAPNOBJ, V) sets the default value to D. This also
            % sets the value returned by hasDefault(MAPNOBJ) to true.
            %
            % See also: MapN, MapN/getDefault, MapN/hasDefault,
            % MapN/noDefault

            M.doDefault = true;
            M.default = d;
        end
        
        
        function noDefault(M)
            % unsets the default value
            %
            % noDefault(MAPNOBJ) removes any default value from MAPNOBJ,
            % causing it to throw an exception if an attempt is made to
            % index it with a key list that has not been stored. This also
            % sets the value returned by hasDefault(MAPNOBJ) to false.
            %
            % See also: MapN, MapN/getDefault, MapN/hasDefault,
            % MapN/setDefault

            M.doDefault = false;
        end
        
        
        function b = uniformValues(M)
            % returns the value of the 'uniformvalues' parameter
            %
            % B = uniformValues(MAPNOBJ) returns true only if
            % 'uniformvalues' was set to true in the constructor when
            % MAPNOBJ was created.
            %
            % See also: MapN
            
            b = M.univals;
        end
           
        
        function l = length(M)
            % returns the number of stored keylist/value pairs
            %
            % L = length(MAPNOBJ) returns the number of keylist/value pairs
            % stored in MAPNOBJ.
            
            l = M.Ct;
        end
        
        
        function b = isempty(M)
            % tests if any data is stored
            %
            % B = isempty(M) returns true only if no keylist/value pairs
            % are currently stored in MAPNOBJ.
            %
            % See also: MapN
            
            b = M.Ct == 0;
        end
        
        
        function varargout = size(M, dim)
            % returns a version of the size
            %
            % S = size(MAPNOBJ) returns the number of keylist/value pairs
            % stored in MAPNOBJ. Other syntaxes, such as [S1, S2] =
            % size(MAPNOBJ) and S = size(MAPNOBJ, DIM) return the number of
            % pairs for the first dimension and 1 for the second and
            % subsequent dimensions.
            %
            % Although MapN objects are multidimensional, it is not
            % possible to compute a meaningful size for each dimension
            % because the keys are not restricted to a small set of
            % integers. The size method therefore behaves as if MapN
            % objects were 1-dimensional, with a length equal to the number
            % of stored associations.
            %
            % See also: MapN, MapN/length

            if nargin == 2
                if isequal(dim, 1)
                    varargout{1} = M.Ct;
                else    % containers.Map doesn't check dim is valid either!
                    varargout{1} = 1;
                end
            else
                if nargout < 2
                    varargout{1} = [M.Ct 1];
                else
                    varargout{1} = M.Ct;
                    varargout(2:nargout) = num2cell(ones(1, nargout-1));
                end
            end
        end
        
        
        function Mout = horzcat(varargin) %#ok<STOUT>
            % throws an exception
            %
            % Horizontal concatenation is not supported.
            %
            % See also: MapN
            
            error('MapN:HorzcatNotSupported', ...
                'Horizontal concatenation is not supported by a MapN');
        end
                
        
        function Mout = vertcat(M, varargin)
            % combines MapN objects into a single object
            %
            % MAPNOBJ_OUT = [MAPNOBJ1; MAPNOBJ2; ...] combines MAPNOBJ1
            % etc. into a single object which contains all the keys and a
            % value for each key.
            %
            % See also: MapN, MapN/copy
            
            Mout = copy(M);
            
            for ii = 1:length(varargin)
                M = varargin{ii};
                storeValues(Mout, keys(M), values(M));
                if M.doDefault
                    Mout.doDefault = true;
                    Mout.default = M.default;
                end
            end
        end
        
        
        function Mout = copy(M)
            % makes a copy of a MapN object
            %
            % MAPNOBJ_OUT = copy(MAPNOBJ_IN) makes a copy of the input
            % object, such that updates to MAPNOBJ_OUT do not affect
            % MAPNOBJ_IN, and vice versa.
            %
            % Because MapN is a handle class, writing MAPNOBJ_OUT =
            % MAPNOBJ_IN does not make an independent object, only a new
            % handle to it, and updates to one affect the other.
            %
            % See also: MapN
            
            Mout = MapN(keys(M), values(M), 'uniformvalues', M.univals);
            Mout.doDefault = M.doDefault;
            Mout.default = M.default;
        end            
            
    end
    
    
    methods (Access = private)
        
        function v = subsrefAction(M, ss)
            % Implements indexed access by traversing a tree of Maps. Each
            % key in a key list is used as an index into a Map whose values
            % are the Maps for the next position in the list, except for
            % the last Map whose values are to be returned. At the root of
            % the tree is a cell array indexed by the number of keys in the
            % key list.
            %
            % The default is handled at a higher level by catching an
            % exception. This is inelegant, but is forced on us because
            % containers.Map does not provide a default mechanism at all,
            % and we would need to call containers.Map.isKey before every
            % access to avoid exceptions. Our strategy prioritises
            % efficient access for valid key lists.
            
            nargs = length(ss);
            v = sr(M.rootMap{nargs}, 1);
            
            function v = sr(m, ind)
                if ind == nargs
                    v = m(ss{ind});
                else
                    v = sr(m(ss{ind}), ind+1);
                end
            end
            
        end

        
        function M = subsasgnAction(M, ss, v)
            % Implements indexed update by building a tree of Maps. New
            % Maps are constructed and added to the tree as necessary. See
            % subsrefAction.
            
            nargs = length(ss);
            rmap = M.rootMap;
            
            if length(rmap) >= nargs && ~isempty(rmap{nargs})
                sa(rmap{nargs}, 1);
            else
                M.rootMap{nargs} = sa([], 1);
            end
                       
            function m = sa(m, ind)
                import containers.Map
                
                key = ss{ind};
                
                if isempty(m) % need to make a new Map
                    
                    % Need to check that an array has not been passed, in order
                    % to pass the key to the Map constructor
                    if ~(isscalar(key) || ischar(key))
                        error('MapN:Subsasgn:arrayKey', ...
                            'Key must be a scalar or a string');
                    end
                    
                    if ind == nargs
                        m = Map(key, v, 'uniformvalues', M.univals);
                        M.Ct = M.Ct + 1;
                    else
                        m = Map(key, sa([], ind+1));
                    end
                    
                else  % update existing Map
                    
                    if ind == nargs
                        hadKey = isKey(m, key);
                        m(key) = v;
                        if ~hadKey
                            M.Ct = M.Ct + 1;
                        end
                    else
                        if isKey(m, key)
                            sa(m(key), ind+1);
                        else
                            m(key) = sa([], ind+1);
                        end
                    end
                    
                end
            end
                        
        end

    end
    
    methods (Access = private)
        
        function v = subsrefError(M, me, ss)
            % handles exceptions on access. Includes default mechanism -
            % see comments below and in subsrefAction.
            
            if isempty(ss) % must come first, as identifier same as next case
                throwAsCaller(MException('MapN:Subsref:emptyKeys', ...
                    'At least one key must be supplied.'));
                
            elseif ismember(me.identifier, ...
                    {'MATLAB:badsubscript', 'MATLAB:Containers:Map:NoKey'})
                % Return default if one is set. Unpleasant to do it via
                % exception handling, but since containers.Map does not
                % allow a default, the alternative is to call isKey before
                % every Map access, costing time for normal access.
                if M.doDefault
                    v = M.default;
                else
                    throwAsCaller(MException('MapN:Subsref:NoKey', ...
                        'The specified key list is not present in this container.'));
                end
                
            elseif strcmp(me.identifier, 'MATLAB:Containers:TypeMismatch')
                throwAsCaller(MException('MapN:Subsref:KeyTypeMismatch', ...
                    'A key has a different type to that expected'));
                
            else
                rethrow(me);
            end
        end
        
        
        function subsasgnError(~, me, ss)
            % handles exceptions on updates
            
            if isempty(ss)   % should be trapped by syntax checking
                throwAsCaller(MException('MapN:Subsasgn:emptyKeys', ...
                    'At least one key must be supplied.'));
                
            elseif strcmp(me.identifier, 'MATLAB:Containers:TypeMismatch')
                % Map has same id but different messages for key and val
                % mismatch! To distinguish need to look at the message.
                if isempty(strfind(me.message, 'key'))
                    throwAsCaller(MException('MapN:Subsasgn:ValueTypeMismatch', ...
                        'A value has a different type to that expected'));
                else
                    throwAsCaller(MException('MapN:Subsasgn:KeyTypeMismatch', ...
                        'A key has a different type to that expected'));
                end
              
            elseif ~isempty(strfind(me.identifier, 'MapN:Subsasgn')) || ...
                    strcmp(me.identifier, 'MATLAB:Containers:Map:UniformNonScalar')
                throwAsCaller(me);
                
            else
                rethrow(me);
            end
        end
        
        
        function removeError(~, me, ss)
            % handles exceptions on key removals
            
            if  isempty(ss)  % must come first
                throwAsCaller(MException('MapN:Remove:emptyKeys', ...
                    'At least one key must be supplied.'));
                
            elseif ismember(me.identifier, ...
                    {'MATLAB:badsubscript', 'MATLAB:Containers:Map:NoKey'})
                throwAsCaller(MException('MapN:Remove:NoKey', ...
                    'The specified key list is not present in this container.'));
                
            elseif strcmp(me.identifier, 'MATLAB:Containers:TypeMismatch')
                throwAsCaller(MException('MapN:Remove:TypeMismatch', ...
                    'A key has a different type to that expected at its position'));
                
            else
                rethrow(me);
            end
        end
        
    end
    
end
```
