## Supplemental File 1 for "A biophysical model of striatal microcircuits suggests delta/theta-rhythmically interleaved gamma and beta oscillations mediate periodicity in motor control": memoize.m

```
function fmem = memoize(f)
%MEMOIZE stores function results to avoid recalculation
%   FMEM = MEMOIZE(F) accepts a function handle, F, and returns a function
%   handle to a memoized version of the same function. The following
%   restrictions apply to the input function:
%
%   - each argument to F must be a scalar numerical value or a string
%   - F must be called with at least one argument
%   - each argument should have a consistent type (see MapN for the
%       detailed restrictions on types in the argument list)
%   - nargout(F) must be positive, and when called F must actually return
%       this number of results
%   - F should not have side-effects (that matter)
%
% F may have a variable number of arguments.
%
% The first time FMEM is called with a given argument list, F is called to
% compute the results, which are returned and also stored. If FMEM is
% called again with the same argument list, the stored results are
% returned.
%
% Example
%
%   existM = memoize(@exist);
%
%   existM(matlabroot)
%   existM(matlabroot, 'dir')   % existM is used just like exist
%
% Calling existM may be faster than calling exist, especially for finding
% out out about disk files, but the results from existM will be out of date
% if there are changes between calls with the same arguments.
%
% See also: MapN

% Copyright David Young 2011

store = MapN;
nout = nargout(f);

if nout == 1
    fmem = @memo1;
elseif nout > 1
    fmem = @memoN;
else
    error('Memoize:variableNargout', ...
        'Function must have definite number of outputs');
end

    function v = memo1(varargin)
        % One result returned, so can be stored as is
        if isKey(store, varargin{:})
            v = store(varargin{:});
        else
            v = f(varargin{:});
            store(varargin{:}) = v;
        end
    end

    function varargout = memoN(varargin)
        % Wraps multiple results up into a cell array for storage
        if isKey(store, varargin{:})
            result = store(varargin{:});
        else
            [result{1:nout}] = f(varargin{:});
            store(varargin{:}) = result;
        end
        varargout = result(1:nargout);
    end

end
```
