## Supplemental File 1 for "A biophysical model of striatal microcircuits suggests delta/theta-rhythmically interleaved gamma and beta oscillations mediate periodicity in motor control": calcClasses.m

```
function [out, outsimple] = calcClasses(obj,input,field_type)
    % For internal use in importing data in to nDDict.

    switch field_type
        case 'data'
            
            % Returns class type of entries in field.
            if isnumeric(input)
                out = 'numeric';
            elseif iscell(input)
                if iscellnum(input)
                    out = 'cellnum';
                elseif all(cellfun(@(s) isa(s,'nDDict'),input(:))) || all(cellfun(@(s) isa(s,'xPlt'),input(:)))
                    out = 'cellnDDict';
                else
                    out = 'cell';
                end
            else
                out = 'unknown';
            end


        case 'axis_values'
            % Validate input
            if ~iscell(input); error('Input must be cell array'); end
            
            % Create dummy axis handle to get access to its functions
            nda = nDDictAxis;
            
            Na = length(input);
            out = cell(1,Na);
            
            % Returns class type of entries in obj.axis.values
            for i = 1:Na
                out{i} = nda.calcClasses(input{i},'values');
            end
            
        case 'axis_name'
            % Validate input
            if ~iscell(input); error('Input must be cell array'); end
            
            % Create dummy axis handle to get access to its functions
            nda = nDDictAxis;
            
            Na = length(input);
            out = cell(1,Na);
            
            % Returns class type of entries in obj.axis.values
            for i = 1:Na
                out{i} = nda.calcClasses(input{i},'name');
            end
            
        otherwise
            error('Unrecognized input foramt');

    end
    
    % If out is a cell, just call it a cell and dont give advanced details
    outsimple = out;
    outsimple = strrep(outsimple,'cellnum','cell');
    outsimple = strrep(outsimple,'cellnDDict','cell');


end


function OUT = iscellnum(IN)
% ISCELLNUM(S) returns 1 if IN is a cell array of numerics and 0
%   otherwise.

    OUT = all(cellfun(@isnumeric,IN(:)));

end
```
