## Supplemental File 1 for "A biophysical model of striatal microcircuits suggests delta/theta-rhythmically interleaved gamma and beta oscillations mediate periodicity in motor control": importLinearData.m

```
function obj = importLinearData(obj,X,varargin)
%% obj = importLinearData(obj,X,varargin)
%     Purpose:
%     Imports a linear array of data, converts it into a matrix
%     based on the supplied axislabels, and stores it in xp.data_pr.
%     Also populates the xp.axis_pr.values appropriately.
% 
%     Forms:
%     xp = importLinearData(X,axislabels1,...,axislabelsN)
% 
%     Inputs:
%     X - vector containing input data. Can be numeric or cell array.
%     axislabels1 - vector containing data labels for dimension1 in X
%     ...
%     axislabelsN - vector containing containing data labels for dimensionN in X
%     NOTE: axislabels1...N must be either numeric or cell arrays of character vectors

    % Initialize
    axeslinear = varargin;


% %     Don't need this option for now.
%     % Check if final argument in varargin is a name/value pair
%     if ischar(varargin{end-1})
%         if strcmp(varargin{end-1},'format')
%             outputformat = varargin{end};
%             axeslinear = axeslinear(1:end-2);       % Remove the name-value pair from axeslinear
%         end
%     end

    % Error checking - X must be linear
    if ~isvector(X); error('X must be linear'); end

    % Error checking - X must be cell or numeric
    obj2 = obj.reset;
    [temp1,Xformat] = obj2.calcClasses(X,'data');
    if strcmp(Xformat,'unknown'); error('X must be numeric or cell array'); end

    % Error checking - each entry in axislinear must be either numeric or
    % cell. If it's a cell, all entries must char.
    temp = obj2.calcClasses(axeslinear, 'axis_values');
    if any(strcmp(temp,'unknown')); error('Axislabels must be numeric or cell array of all chars'); end

    N = length(X);
    Ndims = length(axeslinear);

    % Set up xp.axis_pr
    for j = 1:Ndims
        obj.axis_pr(j).values = unique(axeslinear{j});
        sz(j) = length(obj.axis_pr(j).values);

        if isnumeric(axeslinear{j}(1))
            if any(isnan(axeslinear{j})) || any(isinf(axeslinear{j}))
                error('Axis cannot contain NaNs or Infs');
            end
        end
    end

    % Set up target matrix
    switch Xformat
        case 'cell'
            obj.data_pr=cell(sz);
%         case 'string'
%             xp.data_pr = repmat(string(''),sz);
        case 'numeric'
            obj.data_pr = zeros(sz);
        otherwise
            error('Case not implemented');
    end

    % Set up xp.data_pr -> Convert linear data into a multi dimensional matrix
    for i = 1:N
        % Get subscripts
        subs = cell(1,Ndims);
        for j = 1:Ndims
            if iscellstr(axeslinear{j})
                subs{j} = find(strcmp(axeslinear{j}{i},obj.axis_pr(j).values));
            else
                subs{j} = find(axeslinear{j}(i) == obj.axis_pr(j).values);
            end
        end

        % Add data to sparse cell array or matrix based on subscripts
            % Note: Need to find a good way for dealing with duplicate
            % rows. Right now, default behavior is to overwrite
        obj.data_pr(subs{:}) = X(i);

    end

end
```
