## Supplemental File 1 for "A biophysical model of striatal microcircuits suggests delta/theta-rhythmically interleaved gamma and beta oscillations mediate periodicity in motor control": nDDict.m

```
%% % % % % % % % % % % % % % % % % % % % % % % % % % % % % % % 
% % % % % % % % % % % MAIN CLASS DEF % % % % % % % % % % %
% % % % % % % % % % % % % % % % % % % % % % % % % % % % % % % %

classdef nDDict

    properties
        meta = struct;     % Metadata about stuff that's stored in data
    end
    
    properties (Access = private)
        data_pr               % Storing the actual data (multi-dimensional matrix or cell array)
        axis_pr = nDDictAxis  % 1xNdims - array of nDDictAxis classes for each axis. Ndims = ndims(data)
    end
    
    properties (Dependent)
        data
        axis
    end
    
    
    methods
        %% % % % % % % % % % % % % % % % % % % % % % % % % % % % % % % 
        % % % % % % % % % % % Getter and Setters % % % % % % % % % % % % %
        % % % % % % % % % % % % % % % % % % % % % % % % % % % % % % % %        
        function obj = set.data(obj,value)
            obj.data_pr = value;
            obj.checkDims;
        end
        function value = get.data(obj)
            value = obj.data_pr;
        end
        
        function obj = set.axis(obj,value)
            obj.axis_pr = value;
            obj.checkDims;
        end
        
        function varargout = get.axis(obj)
            [varargout{1:nargout}] = obj.axis_pr;
        end
        
        %% % % % % % % % % % % % % % % % % % % % % % % % % % % % % % % 
        % % % % % % % % % % % CLASS SETUP % % % % % % % % % % % % % % %
        % % % % % % % % % % % % % % % % % % % % % % % % % % % % % % % %
        function obj = nDDict
            
            obj.axis_pr = repmat(nDDictAxis,1,ndims(obj.data_pr));     % For a 2D matrix
            obj = obj.fixAxes;
            
        end
        
        function [obj] = reset(obj)
            obj.data_pr = [];
            obj.axis_pr = nDDictAxis;
            obj.meta = struct;
        end
        
        % TO DO: Fix this.
        function obj_xp = xPlt(obj)
            % Converter nDDict -> xPlt.
            obj_xp = xPlt;
            obj_xp = importData(obj_xp, obj.data);
            obj_xp.axis = obj.axis;
            obj_xp.meta = obj.meta;
            
        end
        
        %% % % % % % % % % % % % % % % % % % % % % % % % % % % % % % % 
        % % % % % % % % % % % INDEXING/SEARCHING DATA % % % % % % % % % % %
        % % % % % % % % % % % % % % % % % % % % % % % % % % % % % % % %
        
        function [selection_out, startIndex] = findaxis(obj,str)
            % Returns the index of the axis with name matching str
            allnames = {obj.axis_pr.name};
            [selection_out, startIndex] = regex_lookup(allnames, str);
        end

        function [obj2, ro] = subset(obj,varargin)
            % Define variables and check that all dimensions are consistent
            % ro - if regular expressions are used, returns the index
            % values discovered by the regular expression.
            
            % Verify that size of obj is correct
            checkDims(obj);
            
            % Validate that "selection" is the right size
            selection = varargin(:);
            Na = length(obj.axis_pr);
            Nd = ndims(obj.data_pr);
            
            % Fill out selection with empties if too short. For example, say
            % size(obj.data_pr) is MxNx1x1. The user might enter a selection with
            % length(selection) = 2, such as selection = {[A],[B]}. In this
            % case, we convert it from to selection = {[A],[B],[],[]}.
            Ns = length(selection);
            if Ns < Na && Ns >= Nd
                selection2 = repmat({[]},1,Na);
                selection2(1:Ns) = selection;
                selection = selection2;
                clear selection2
            end
            
            % If Ns is still wrong dimensions, return error
            Ns = length(selection);
            if Ns ~= Na
                error('Number of inputs must match dimensionality of nDDict.data_pr');
            end
            
            % Convert selection to index if using regular expressions
            ro = {};
            for i = 1:Ns
                if ischar(selection{i})
                    ro{i}(1,:) = obj.axis_pr(i).values;
                    [selection{i} ro{i}(2,:)] = regex_lookup(obj.axis_pr(i).values, selection{i});
                    
                end
            end
            
            % Make sure that size of selection doesnt exceed size of data
            sz = size(obj);
            for i = 1:Ns
                if selection{i} > sz(i); error('Selection index exceeds dimensions'); end
            end
            
            % Initialize
            
            obj2 = obj;             % Create new class of same type as original
            obj2 = obj2.reset;
            obj2.meta = obj.meta;
            
            % First update each axis with the corresponding selection and
            % convert empty cells to code for full range.
            for i = 1:Ns

                if isempty(selection{i})
                    selection{i} = 1:sz(i);
                end
                
                obj2.axis_pr(i) = obj.axis_pr(i);       % Import axis information
                obj2.axis_pr(i).values = obj.axis_pr(i).values(selection{i});   % Overwrite values field; leave everything else the same.
            end
            
            % Update the data
            obj2.data_pr = obj.data_pr(selection{:});
            
            % Corrects number of axes. The above code automatically
            % converts obj.data_pr from MxNx1x1 to MxN, whereas axis will stay
            % as MxNx1x1 (e.g. length of 4). Thus, fixAxes corrects this.
            % This should no longer be necessary!!!
            % obj2 = fixAxes(obj2);
            
        end
        
        %% % % % % % % % % % % % % % % % % % % % % % % % % % % % % % % 
        % % % % % % % % % % % % IMPORT DATA  % % % % % % % % % % % % % %
        % % % % % % % % % % % % % % % % % % % % % % % % % % % % % % % %
        
        function obj = importAxisNames(obj,ax_names)
            Nd = ndims(obj.data_pr);
            Na = length(obj.axis_pr);
            
            if nargin < 2
                ax_names = cellfun(@num2str,num2cell(1:Nd),'UniformOutput',0);
                ax_names = cellfun(@(s) ['Dim ' s],ax_names,'UniformOutput',0);
            end
            
            if ~iscellstr(ax_names); error('ax_names must be a cell array of chars'); end
            
            if length(ax_names) > Na
                error('Mismatch between number of axis names supplied and number of axes in object.'); end

            for i = 1:length(ax_names)
                obj.axis_pr(i).name = ax_names{i};
            end
        end
        
        function obj = importMeta(obj,meta_struct)
            obj.meta = meta_struct;
        end
        
        obj = importLinearData(obj,X,varargin)            % Function for importing data in a linear format
        
        function obj = importData(obj,data,axis_vals,axis_names)
            
            obj.data_pr = data;
            obj = obj.fixAxes;
            
            if nargin > 2
                if ~isempty(axis_vals)
                    if ~iscell(axis_vals); error('axis_vals must be a cell array.'); end
                    for i = 1:length(axis_vals);
                        obj.axis_pr(i).values = axis_vals{i};
                    end
                    obj.checkDims;
                end
            end
            
            if nargin > 3
                if ~isempty(axis_names)
                    obj = obj.importAxisNames(axis_names);
                end
            end
            
        end
        
        function out = exportAxisVals(obj)
            Na = length(obj.axis);
            out = cell(1,Na);
            for i = 1:Na
                out{i} = obj.axis(i).values;
            end
        end
        function out = exportAxisNames(obj)
            Na = length(obj.axis);
            out = cell(1,Na);
            for i = 1:Na
                out{i} = obj.axis(i).name;
            end
        end
        
        
        %% % % % % % % % % % % % % % % % % % % % % % % % % % % % % % % 
        % % % % % % % % % % % REARRANGING DATA % % % % % % % % % % %
        % % % % % % % % % % % % % % % % % % % % % % % % % % % % % % % %
        
        function obj = mergeDims(obj,dims2pack)
            
            obj.checkDims;
            Nd2p = length(dims2pack);
            %sz = size(obj.data_pr);
            sz = size(obj);
            N = ndims(obj);
            Nmerged = prod(sz(dims2pack));       % Final number entries in the merged dimension
            
            % % First, do axis names
            % Get cell array of all linearized axis values.
            inds = 1:Nmerged;
            [subs{1:Nd2p}] = ind2sub(sz(dims2pack),inds);
            temp = cell(1,Nd2p);
            for i = 1:Nd2p
                for j = 1:Nmerged
                    ax = obj.axis_pr(dims2pack(i));
                    currval = ax.values(subs{i}(j));
                    temp{i}(j) = currval;
                end
            end
            
            % Compress these into a single string to be used as new value name
            tempstr = {};
            for i = 1:Nd2p
                for j = 1:Nmerged
                    if iscellstr(temp{i}); currval = temp{i}{j};
                    else
                        currval = num2str(temp{i}(j));  % If it's not a string, convert it to one.
                    end
                    if i == 1; tempstr{j} = currval;
                    else tempstr{j} = [tempstr{j} '_' currval];
                    end
                end
            end
            
            % Finally drop this into the values entry for the new "merged"
            % axis
            obj.axis_pr(dims2pack(1)).values = tempstr;
            obj.axis_pr(dims2pack(1)).astruct.premerged_values = temp;
            obj.axis_pr(dims2pack(1)).astruct.premerged_names = {obj.axis_pr(dims2pack).name};
            
            % Give it a new axis name, reflecting the merger of all the
            % others
            allnames = {obj.axis_pr(dims2pack).name};
            allnames = cat(1,allnames(:)',repmat({'_'},1,length(allnames)));
            obj.axis_pr(dims2pack(1)).name = strcat(allnames{1:end-1});
            
            % Clear the remaining axes names
            for i = 2:Nd2p
                obj.axis_pr(dims2pack(i)).name = ['Dim ' num2str(dims2pack(i))];
                obj.axis_pr(dims2pack(i)).values = 1;
            end

            % % Now, work on obj.data_pr
            dat = obj.data_pr;
            obj.data_pr = [];       % Clear obj.data_pr for now, to save memory.
            
            % Figure out which dimensions were *NOT* Targeted for the merge
            Nd = ndims(dat);
            alldims = 1:Nd;
            ind_chosen = false(size(alldims));
            for i = 1:length(dims2pack)
                ind_chosen = ind_chosen | alldims == dims2pack(i);
            end
            ind_unchosen = ~ind_chosen;
            dims_remaining = find(ind_unchosen);
            
            % Bring the dims to be merged to the front
            dat = permute(dat,[dims2pack,dims_remaining]);
            
            % REshape these into a single dim
            %sz = size(dat);
            sz = arrayfun(@(x) size(dat,x), 1:N);       % Need to use this extended size command to get extra railing 1's for certain use cases.
            dat = reshape(dat,[ prod(sz(1:Nd2p)), ones(1,Nd2p-1), sz(Nd2p+1:end) ]);
            
            % Undo the earlier permute, and put back into obj.data_pr
            dat = ipermute(dat,[dims2pack,dims_remaining]);
            obj.data_pr = dat;
        end
        
        function obj = packDim(obj,dim_src,dim_target)
            
            if isstr(dim_src)
                dim_src_string = dim_src;
                dim_src = obj.axis.findAxes(dim_src_string);
                if ~isscalar(dim_src) || isempty(dim_src)
                    error('Multiple or zero dimensions matching %s.', dim_src_string)
                end
            end
            
            if ~isscalar(dim_src) || isempty(dim_src) || dim_src == 0
                error('Dimension to pack must be a nonempty, nonzero scalar.')
            end
            
            if nargin < 3
                % Should pack dimension as last dimension of obj.data.
                sizes = cellfun(@(x) length(size(x)), obj.data);
                dim_target = max(sizes(:)) + 1; 
            end
            
            checkDims(obj);
            
            % Make sure that obj.data_pr is a cell array
            if ~iscell(obj.data_pr); error('nDDict.data_pr must be a cell array.'); end
            
            % Make sure that obj.data_pr is a numeric
            temp = cellfun(@isnumeric,obj.data_pr);
            if any(temp(:) ~= 1); error('nDDict.data_pr must contain only numerics'); end      % Can redo this in the future to work with nDDicts containing matrices
            % % To do: implement this so it works with cell arrays and nDDict
            % classes in the future too
            
            % Make sure target dimension in nDDict.data_pr is a singleton
            temp = cellfun(@(x) size(x,dim_target),obj.data_pr);
            if any(temp(:) ~= 1); error('Target dimension in nDDict.data_pr needs to be size 1. Try reshaping contents of nDDict.data_pr or choosing a different target dimension.'); end
            clear sz_targets
            
            % Bring chosen dimension to the front. Thus, we will be
            % merging along rows.
            Nd = ndims(obj.data_pr);
            obj.data_pr = permute(obj.data_pr,[dim_src, 1:dim_src-1, dim_src+1:Nd]);
            
            % Temporarily linearize all other dimensions.
            sz0 = size(obj.data_pr);
            obj.data_pr = reshape(obj.data_pr,sz0(1),prod(sz0(2:end)));
            
            % Add NaNs where needed
                % Note: to understand what this is doing, it really helps
                % to draw a diagram!
            sz = size(obj.data_pr);
            empties = cellfun(@isempty,obj.data_pr);    % 2D matrix with 1's marking empty cells
            se = sum(empties,1);                    % Number of empties per column in this matrix
            bad_inds = se ~= 0 & se ~= sz(1);     % Good columns must be either all empty or all non-empty
            
            if any(bad_inds)
                fprintf('Note: Empty entries found along collapsing dim. Using NaNs as placeholders to fill out the matrix. \n');
                bi = find(bad_inds);
                for j = 1:length(bi)                    % Sweep through bad columns
                    curr_bad = find(empties(:,j));      % Empties along this particular column
                    curr_good = find(~empties(:,j));    % Non-empties along this particular column.
                    for i = 1:length(curr_bad)
                        % Populate the empty cells with matrices of NaNs
                        % that are the same dimensionality as the first
                        % good entry.
                        obj.data_pr{curr_bad(i),bi(j)} = NaN*ones(size(obj.data_pr{curr_good(1),bi(j)}));
                    end
                end
            end
            
            % Check that sizes and dimensionalities are compatible
            data_ndims = cellfun(@ndims,obj.data_pr,'UniformOutput',true);
            if any(any(data_ndims ~= repmat(data_ndims(1,:),sz(1),1),1),2)
                error('Dimensions of nDDict.data_pr not uniform along packing dimensions.');
            end
           
            data_sz = cellfun(@size,obj.data_pr,'UniformOutput',false);
            data_sz_firsts = repmat(data_sz(1,:),sz(1),1);
            myfunc = @(x,y) any(x(:) ~= y(:));
            bool_size_mismatch = cellfun(myfunc,data_sz,data_sz_firsts);
            if any(bool_size_mismatch(:))
                error('Sizes of nDDict.data_pr are not uniform along packing dimension. (This usually results form trying to combine populations with different numbers of cells.');
            end
        
            for j = 1:sz(2)
                obj.data_pr{1,j} = cat(dim_target,obj.data_pr{:,j});
                obj.data_pr(2:end,j) = cell(sz(1)-1,1);     % Set remainder to empty
            end
            
            obj.data_pr = obj.data_pr(1,:);         % Keep only 1st dimension;
            sz0(1) = 1;

            % Lastly, restore original dimensions
            % of nDDict.data_pr
            obj.data_pr = reshape(obj.data_pr,sz0);
            obj.data_pr = permute(obj.data_pr,[2:dim_src, 1, dim_src+1:Nd]);
            
            % Also, clear obj.axis_pr
            ax_src = obj.axis_pr(dim_src);
            obj = setAxisDefaults(obj,dim_src);
            obj.axis_pr(dim_src).name = ['Dim ' num2str(dim_src)];
            
            % Store obj.axis_pr(dim_src) as meta data.
            obj.meta.(['matrix_dim_', num2str(dim_target)]) = ax_src;
            
            % If obj.data_pr is a nDDict object itself, update axes labels
            % appropriately
            for i = 1:numel(obj.data_pr)
                if isa(obj.data_pr{i},'nDDict')
                    warning('This mode doesnt work yet');
                    obj.data_pr{i}.axis_pr(dim_target) = ax_src;
                end
            end
            
        end
        
        function obj_out = merge(obj1,obj2)
            % This might be slow when working with huge matrices. Perhaps do
            % alternate approach for them.
            names = {obj1.axis_pr.name};
            
            % Merge two objects together
            Nd1 = ndims(obj1);
            obj1 = obj1.mergeDims(1:Nd1);
            X1 = obj1.data_pr;
            axislabels1 = obj1.axis_pr(1).astruct.premerged_values;
            
            
            Nd2 = ndims(obj2);
            obj2 = obj2.mergeDims(1:Nd2);
            X2 = obj2.data_pr;
            axislabels2 = obj2.axis_pr(1).astruct.premerged_values;
            
            X = vertcat(X1(:),X2(:));
            for i = 1:length(axislabels1)
                axl{i} = vertcat(axislabels1{i}(:),axislabels2{i}(:));
            end
            
            obj_out = obj1.reset;
            obj_out = importLinearData(obj_out,X,axl{:});
            
            obj_out = obj_out.importAxisNames(names);
        end
        
        function obj_new = unpackDim(obj, dim_src, dim_target, dim_name, dim_values)
            
            % Temporarily linearize obj.data_pr.
            sz0 = size(obj);
            dim0 = length(sz0);
            obj.data_pr = reshape(obj.data_pr,prod(sz0),1);
            
            % Get sizes of dimension dim_src for matrices inside obj.data_pr.
            sizes = cellfun(@(x) size(x, dim_src), obj.data_pr);
            max_size = max(sizes);
            
            % Calculate size of new nDDict object with dim_src unpacked.
            % The unpacked dimension will be the new first dimension.
            sz_new = [max_size, sz0];
            
            % Initialize new nDDict object which will have dim_src
            % unpacked.
            obj_new = obj; % eval(['obj_new = ', class(obj), ';'])
            obj_new = obj_new.reset;
            
            % % Creating obj_new.data_pr. % % % % % % % % % % % % % % % % % % 
            % Loop over linearized indices of old obj.data_pr cell array.
            for data_index = 1:size(obj.data_pr, 1)
                
                % Retrieve matrix from obj.data_pr at data_index.
                temp_matrix = obj.data_pr{data_index};
                [temp_size, slice_size] = deal(size(temp_matrix));
                
                % Create indices for an arbitrary slice from dimension
                % dim_src, padding out with ':' if temp_matrix has fewer
                % dimensions than dim_src.
                temp_effective_dimensions = max(length(temp_size), dim_src);
                slice_indices = cell(1, temp_effective_dimensions);
                slice_indices(:) = {':'};
                
                % Loop over slices of dim_src.
                for slice_index = 1:max_size
                    
                    % Find linearized index in obj_new that this slice will inhabit.
                    new_index = (data_index - 1)*max_size + slice_index;
                    
                    if slice_index <= sizes(data_index)
                        slice_indices{dim_src} = slice_index; % Get correct slice indices.
                        slice = temp_matrix(slice_indices{:});
                    else
                        slice = [];
                    end
                    
                    obj_new.data_pr{new_index} = slice;
                    
                end
                
            end
            
            % Reshape obj_new to be multidimensional, with the unpacked
            % dimension as dimension 1.
            obj_new.data_pr = reshape(obj_new.data_pr, sz_new);
            
            %  % Creating obj_new.axis_pr and obj_new.meta. % % % % % % % % % 
            % Setting default axis name and values.
            % If none are given, let names and values for the dimension be empty.
            if nargin == 4
                dim_values = [];
            elseif nargin < 4
                dim_name = []; dim_values = [];
            end
            
            % Save obj.meta to pass to obj_new.
            meta = obj.meta;
            
            % If names and/or values are empty, search for them in
            % meta; if they are found, remove them from meta.
            dim_src_name = ['matrix_dim_' num2str(dim_src)];
            if isfield(meta, dim_src_name)
                if isempty(dim_name)
                    dim_name = meta.(dim_src_name).name;
                end
                if isempty(dim_values)
                    dim_values = meta.(dim_src_name).values;
                end
                meta = rmfield(meta, dim_src_name);
            end
            
            % If names and/or values are still empty, replace them with
            % defaults.
            if isempty(dim_name), dim_name = dim_src_name; end
            if isempty(dim_values), dim_values = (1:max_size)'; end
            
            % Pass meta to obj_new.
            obj_new = importMeta(obj_new, meta);
            
            % Checking axis dimensions against data dimensions.
            if length(dim_values) < max_size
                warning('dimension %d is longer for some cells than dim_values.\n', dim_src)
                % dim_values((end + 1):max_size) = (length(dim_values) +
                % 1):max_size; % No longer necessary since we're running
                % fixAxes on obj_new.
            elseif length(dim_values) > max_size
                warning('dimension %d is shorter for all cells than dim_values.\n', dim_src)
            end
            
            % Creating unpacked axis.
            unpacked_axis = nDDictAxis;
            unpacked_axis.name = dim_name;
            unpacked_axis.values = dim_values;
            
            % Make new axis last one, then move to front.
            axis_new = obj.axis_pr;
            axis_new(end + 1) = unpacked_axis;
            axis_new = axis_new([dim0 + 1, 1:dim0]);
            obj_new.axis_pr = axis_new;
            
            % Putting unpacked dimension in dim_target, if given.
            if nargin < 3, dim_target = []; end
            if isempty(dim_target), dim_target = 1; end
            if dim_target ~= 1
                new_dim_order = [2:dim_target 1 (dim_target + 1):(dim0 + 1)];
                obj_new = permute(obj_new, new_dim_order);
            end
            
            % Fix axes.
            obj_new = fixAxes(obj_new);
                
        end
        
        %% % % % % % % % % % % % % % % % % % % % % % % % % % % % % % % 
        % % % % % % % % % % % HOUSEKEEPING FUNCTIONS % % % % % % % % % % %
        % % % % % % % % % % % % % % % % % % % % % % % % % % % % % % % %
        
        function out = getaxisinfo(obj,showclass)
            % If no output arguments, prints axis info to the screen. If
            % output arguments are supplied, returns this information as a
            % string
            
            if nargin < 2
                showclass = 1;
            end
            
            if nargout > 0
                out = '';
            end

            for i = 1:length(obj.axis_pr)
                
                out1 = obj.axis_pr(i).getaxisinfo(showclass);
                spacer = '';
                
                if nargout > 0
                    out = [out, spacer, out1, '; ' ];
                else
                    spacer = ['Axis ', num2str(i), ': '];
                    fprintf([spacer, out1, '\n']);
                end
            end
            
            if isempty(obj.data_pr)
                if nargout > 0; out = 'obj.data_pr is empty';
                else fprintf('obj.data_pr is empty \n');
                end
                return;
            end
            
            % Lastly output a summary of dimensionality comparing nDDict.axis_pr
            % and nDDict.data_pr. These should match up.
            if nargout == 0
                fprintf(['nDDict.axis_pr dimensionality ' num2str(cellfun(@length,{obj.axis_pr.values})) '\n']);
                fprintf(['nDDict.data_pr dimensionality ' num2str(size(obj.data_pr)) '\n']);
            end
        end
        
        function obj = fixAxes(obj)
            % This function forces the nDDict axis data to be updated to
            % match the dimensions of the data structure.
            % The convention of nDDict is to follow MATLAB
            % conventions for dimensionality. Thus, the size(obj.data_pr)
            % command is used to determine the correct number of axes, and
            % axis is adjusted to match, adding or removing dimensions as
            % needed. If you are getting errors when running checkDims, you
            % should run this command.
            %
            % The one exception to MATLAB conventions is that there are
            % allowed to be more axes than there are dimensions in obj.data_pr
            % as long as the number of entries in each of these axes is 1.
            % This allows you to store axis labels and names for trailing
            % singleton dimensions (e.g. dimensions of greater number than
            % ndims(obj.data_pr) would return).

            Nd = ndims(obj.data_pr);
            Na = length(obj.axis_pr);
            
            % Make sure obj.data_pr, obj.axis_pr.name, and obj.axis_pr.values have the right data types
            if strcmp(getclass_obj_data(obj),'unknown'); error('Obj.data_pr must be either numeric or cell array'); end
            if any(strcmp(getclass_obj_axis_values(obj),'unknown')); error('Obj.axis_pr.values must be of type numeric or cell array of character vectors.'); end
            if any(strcmp(getclass_obj_axis_name(obj),'unknown')); error('Obj.axis_pr.name must be of type char.'); end

            % Sweep through all axes and make sure dimensions are correct.
            % Add new axes if needed, up to Nd.
            for i = 1:Nd
                obj = setAxisDefaults(obj,i);  % Sets axis #i to the default name
            end

            % Trim away excess values in axes
            if Na > Nd
                for i = Nd+1:Na
                    if length(obj.axis_pr(i).values) > 1
                        obj.axis_pr(i).values = obj.axis_pr(i).values(1);
                        fprintf(['Extra values found in axis #' num2str(i) ' ' obj.axis_pr(i).name '. Trimming \n']);
                    end
                end
                
            end
            
        end
        
        function checkDims(obj)
            % We enforce that size(obj.data_pr) must always match up to 
            % length(obj.axis_pr(i).values) for all i. We allow there to be
            % more axis than ndims(obj.data_pr), but only if these axes have
            % lengths of 1.
            
            % Note, fixAxes fixes everything automatically.
            % Only call checkDims if you want to
            % be alerted to mismatches, but not to correct them. Use fixAxes to
            % automatically correct everything.
            
            %if isempty(obj); error('Object is empty. Input some data first!'); return; end
            
            % Make sure obj.data_pr, obj.axis_pr.name, and obj.axis_pr.values have the right data types
            if strcmp(getclass_obj_data(obj),'unknown'); error('Obj.data_pr must be either numeric or cell array'); end
            if any(strcmp(getclass_obj_axis_values(obj),'unknown')); error('Obj.axis_pr.values must be of type numeric or cell array of character vectors.'); end
            if any(strcmp(getclass_obj_axis_name(obj),'unknown')); error('Obj.axis_pr.name must be of type char.'); end
            
            sza = arrayfun(@(x) length(x.values),obj.axis_pr);
            szd = size(obj.data_pr);
            
            Nd = ndims(obj.data_pr);
            Na = length(obj.axis_pr);

            if Nd > Na
                error('checkDims: Error found! Number of dimensions in nDDict.data_pr does not equal number of axes. Try using method importData or importLinearData if you want to alter objects dimensions.');
            end

            % For all dimensions in obj.data_pr
            for i = 1:Nd
                if sza(i) ~= szd(i)
                    obj.getaxisinfo
                    error('Mismatch between obj.data and obj.axis dimensionality. Use importData to make modifications like this.');
                end
            end
            
            % For additional axes beyond ndims(obj.data_pr)
            ind = sza > 1;
            if any(ind(Nd+1:Na))
                ind2 = find(ind);
                ind2 = ind2(ind2 > Nd);
                fprintf(['checkDims: Error found! ndims(obj.data_pr)=' num2str(Nd) ' but axis obj.axis_pr(' num2str(ind2) ').values has ' num2str(sza(ind2)) ' entries. Try using method importData or importLinearData if you want to alter objects dimensions.\n']);
                error(' ');
            end
            
            % All good if make it to here
        end

        %% % % % % % % % % % % % % % % % % % % % % % % % % % % % % % % 
        % % % % % % % % % % % OVERLOADED FUNCTIONS % % % % % % % % % % %
        % % % % % % % % % % % % % % % % % % % % % % % % % % % % % % % %
        function A = isempty(obj)
            A = isempty(obj.data_pr);
        end
        
        function varargout = size(obj,varargin)
            % Returns size of obj. This function is basically the same as
            % running size(obj.data_pr) except we base it off of the dimensions
            % of obj.axis_pr rather than obj.data_pr. This has the effect of
            % returning 1's if length(obj.axis_pr) > ndims(obj.data_pr)
            
            checkDims(obj);
            
            [varargout{1:nargout}] = size(obj.data_pr,varargin{:});
            
            % If function is called in the form sz = size(obj) OR size(obj),
            % return the length of each axis.
            if nargout <= 1 && nargin == 1
                Na = length(obj.axis_pr);
                sz = zeros(1,Na);
                for i = 1:Na
                    sz(i) = length(obj.axis_pr(i).values);
                end
                if nargout == 1; varargout{1} = sz; end
            end
        end
        
        function Nd = ndims(obj)
            checkDims(obj);
            Nd = length(obj.axis_pr);
        end
        
        function obj = permute(obj,order)
            checkDims(obj);
            obj.data_pr = permute(obj.data_pr,order);
            obj.axis_pr = obj.axis_pr(order);
        end
        
        function obj = ipermute(obj,order)
            checkDims(obj);
            inverseorder(order) = 1:numel(order);
            obj.data_pr = permute(obj.data_pr,inverseorder);
            obj.axis_pr = obj.axis_pr(inverseorder);
        end
        
        function obj = transpose(obj)
            checkDims(obj);
            Nd = ndims(obj.data_pr);
            
            if Nd > 2; error('Can only transpose data with at most 2 dimensions');
            end
            
            obj.data_pr = obj.data_pr';
            obj.axis_pr([1,2]) = obj.axis_pr([2,1]);        % Axis should always be at least length=2.
        end
        
        function obj = squeeze(obj)
            % This is just like MATLAB's normal squeeze command. However,
            % there is one key difference:
            % Normally, if squeeze operates on a 1xN matrix, it will leave
            % it as 1xN. This function forces it to always return as Nx1
            
            checkDims(obj);
            
            % If data is bigger than a matrix, squeeze out dimensions that
            % are of size 1.
            sz = size(obj.data_pr);
            if length(sz) > 2
                ind = sz~=1;
                obj.axis_pr = obj.axis_pr(ind);

                % Now squeeze obj.data_pr
                obj.data_pr = squeeze(obj.data_pr);         % Normal squeeze command

%                 % Lastly, if the result is a row vector, force it to be a
%                 % column vector
%                 if isvector(obj.data_pr) && ~iscolumn(obj.data_pr)
%                     obj.data_pr = obj.data_pr';
%                 end
            else
                % Otherwise, if data is a matrix, remove all axis beyond
                % the first two. These should only be size 1 (e.g. "name"
                % axes anyways)
%                 szA = cellfun(@length,{obj.axis_pr.values});
%                 ind = szA~=1;
%                 ind(1:2) = true;
                obj.axis_pr = obj.axis_pr(1:2);
            end
            
            % Make sure everything is good before returning.
            obj = obj.fixAxes;
            checkDims(obj);
        end
        
        function obj = abs(obj)
            checkDims(obj);
            obj.data = cellfun(@(x) abs(x), obj.data_pr, 'UniformOutput', 0);
        end
        
        %% % % % % % % % % % % % % % % % % % % % % % % % % % % % % % % 
        % % % % % % % % % % % OVERLOADED OPERATORS % % % % % % % % % % %
        % % % % % % % % % % % % % % % % % % % % % % % % % % % % % % % %
        function varargout = subsref(varargin)
            
%             % Default settings for everything
%             [varargout{1:nargout}] = builtin('subsref',varargin{:});
            
            obj = varargin{1};
            S = varargin{2};
            
            if length(S) == 1               % This discounts cases like obj.subset(1,2,3,4)
                switch S.type
                    case '()'
                        %[varargout{1:nargout}] = builtin('subsref',varargin{:});
                        %varargout{1} = builtin('subsref',obj.data_pr,S);
                        
                        % Convert colon operators to empties, which subset
                        % uses to denote "take everything"
                        for i = 1:length(S.subs)
                            if strcmp(S.subs{i},':')
                                S.subs{i} = [];
                            end
                        end
                        
                        varargout{1} = obj.subset(S.subs{:});
                    case '{}'
                        %[varargout{1:nargout}] = builtin('subsref',varargin{:});
                        S2 = S;
                        S2.type = '()';
                        [varargout{1:nargout}] = builtin('subsref',obj.data_pr,S2,varargin{3:end});
                    case '.'
                        [varargout{1:nargout}] = builtin('subsref',varargin{:});
                    otherwise
                        error('Unknown indexing method. Should never reach this.');
                end
            else
                [varargout{1:nargout}] = builtin('subsref',varargin{:});
            end
             
        end
    end
    
    %% % % % % % % % % % % % % % % % % % % % % % % % % % % % % % % 
    % % % % % % % % % % % PRIVATE FUNCTIONS % % % % % % % % % % %
    % % % % % % % % % % % % % % % % % % % % % % % % % % % % % % % %
    methods (Access = private)
        [out, outsimple] = calcClasses(xp,input,field_type)     % Used by importLinearData and other importData functions
    end
    
    %% % % % % % % % % % % % % % % % % % % % % % % % % % % % % % % 
    % % % % % % % % % % % HELPER FUNCTIONS % % % % % % % % % % %
    % % % % % % % % % % % % % % % % % % % % % % % % % % % % % % % %
    methods
        % These functions should only be called by other nDDict methods.
        % Make these public for now for testing, but set to private
        % eventually.
        function [out, outsimple] = getclass_obj_data(obj)
            [out, outsimple] = obj.calcClasses(obj.data_pr,'data');
        end
        
        function out = getclass_obj_axis_values(obj)
            % Returns class type of entries in obj.axis_pr.values
            Na = length(obj.axis_pr);
            out = cell(1,Na);
            for i = 1:Na
                out{i} = obj.axis_pr(i).getclass_values;
            end
        end
        
        function out = getclass_obj_axis_name(obj)
            % Returns class type of entries in obj.axis_pr.values            
            Na = length(obj.axis_pr);
            out = cell(1,Na);
            for i = 1:Na
                out{i} = obj.axis_pr(i).getclass_name;
            end
        end

    end
end

%% % % % % % % % % % % % % % % % % % % % % % % % % % % % % % % 
% % % % % % % % % % % LOCAL FUNCTIONS % % % % % % % % % % %
% % % % % % % % % % % % % % % % % % % % % % % % % % % % % % % %

function output = inheritObj(output,input)
    % Merges contents of input into output.
    C = metaclass(input);
    P = C.Properties;
    for k = 1:length(P)
        if ~P{k}.Dependent
            output.(P{k}.Name) = input.(P{k}.Name);
        end
    end
end

function obj = setAxisDefaults(obj,dim)
    % Sets obj.axis_pr(i) to default values
    
    % Get desired size of dataset
    sz_dim = size(obj.data_pr,dim);
    
    % If axis doesn't already exist, create it. Otherwise, copy existing.
    if length(obj.axis_pr) < dim
        ax_curr = nDDictAxis;
    else
        ax_curr = obj.axis_pr(dim);
    end
    
    % Name it if necessary
    if isempty(ax_curr.name)
        ax_curr.name = ['Dim ' num2str(dim)];
    end
    
    % If values is empty, add default values.
    if isempty(ax_curr.values)
        %ax_curr.values = cellfun(@num2str,num2cell(1:sz(i)),'UniformOutput',0);     % Populate with strings
        ax_curr.values = 1:sz_dim;                                                   % Populate with numerics
    else
        % Otherwise, make sure dimensionality is correct. If not, update it
        % missing entries with default names.
        N = length(ax_curr.values);
        
        % If too short
        if N < sz_dim
            if isnumeric(ax_curr.values)
                for j = (N + 1):sz_dim; ax_curr.values(j) = j; end
            elseif iscellstr(ax_curr.values)
                for j = (N + 1):sz_dim; ax_curr.values{j} = num2str(j); end
            else
                error('axis.values must be either type numeric or cell array of strings');
            end
        end
        
        % If too long
        if N > sz_dim
            %ax_curr.values = ax_curr.values(1:sz(dim));
            ax_curr.values = 1:sz_dim;                                                   % Populate with genetic numerics
        end
    end
    
    % Assign our new axis to the current dimension
    obj.axis_pr(dim) = ax_curr;
end

function [selection_out, startIndex] = regex_lookup(vals, selection)
    if ~iscellstr(vals); error('nDDict.axis_pr.values must be strings when using regular expressions');
    end
    if ~ischar(selection); error('Selection must be string when using regexp');
    end
    
    startIndex = regexp(vals,selection);
    selection_out = logical(~cellfun(@isempty,startIndex));
    selection_out = find(selection_out);
    
end

% function varargout = size2(varargin)
%     [varargout{1:nargout}] = size(varargin{:});
%     if nargout == 1
%         sz = varargout{1};
%         if length(sz) == 2 && sz(2) == 1
%             sz = sz(1);
%         end
%         varargout{1} = sz;
%     end
% end
```
