## Supplemental File 1 for "A biophysical model of striatal microcircuits suggests delta/theta-rhythmically interleaved gamma and beta oscillations mediate periodicity in motor control": recursivePlot.m

```
function varargout = recursivePlot(xp,function_handles,dimensions,function_arguments)
    % function_arguments - cell array of argument cell arrays to pass to
    % function_handles. Must have one cell array for each function_handle
    % passed. Use empty cell arrays for no arguments. E.g.
    % function_arguments = { {}, {}, {1} } for nothing, nothing, and
    % 1 as arguments.
    
    if nargin < 4
        function_arguments = cell(size(function_handles));
        for i = 1:length(function_arguments)
            function_arguments{i} = {};
        end
    end
    
    % Validate inputs
    Nfh = length(function_handles);
    Nd = length(dimensions);
    Nfha = length(function_arguments);
    
    if Nfh ~= Nd; error('Number of cells in dimensions must equal the number of function handles supplied'); end
    if Nfh ~= Nfha; error('Number of cells in function_arguments must equal number of function handles supplied'); end
    
    
    % Convert any regular expressions in dimensions into their
    % corresponding indices if necessary
    dimensions = dimensions_regex_2_index(xp,dimensions);
    
    
%     Na = ndims(xp);
%     N_dims_supplied = sum(cellfun(@length,dimensions));
%     if Na ~= N_dims_supplied
%         error('Total number of entries supplied in dimensions must equal ndims(xp)');
%     end
    

    sz = size(xp);
    if length(function_handles) > 1
        
        xp2 = xPlt;
        selection_curr = cell(1,length(sz));
        % Just cardcode in the various cases for dimensionality. It's
        % unlikely they will ever want to go above showing 4D in a single
        % plot.
        switch length(dimensions{1})        
            case 1                          % 1D
                dim1 = dimensions{1}(1);
                for i = 1:sz(dim1)
                    selection_curr{dim1} = i;
                        % Note: need to make it a row vector!
                    mydata{i,1} = @() recursivePlot(xp.subset(selection_curr{:}),function_handles(2:end),dimensions(2:end),function_arguments(2:end));
                end
                
            case 2                          % 2D
                dim1 = dimensions{1}(1);
                dim2 = dimensions{1}(2);
                for i = 1:sz(dim1)
                    for j = 1:sz(dim2)
                        selection_curr{dim1} = i;
                        selection_curr{dim2} = j;
                        mydata{i,j} = @() recursivePlot(xp.subset(selection_curr{:}),function_handles(2:end),dimensions(2:end),function_arguments(2:end));
                    end
                end
                
            case 3                          % 3D
                dim1 = dimensions{1}(1);
                dim2 = dimensions{1}(2);
                dim3 = dimensions{1}(3);
                for i = 1:sz(dim1)
                    for j = 1:sz(dim2)
                        for k = 1:sz(dim3)
                            selection_curr{dim1} = i;
                            selection_curr{dim2} = j;
                            selection_curr{dim3} = k;
                            mydata{i,j,k} = @() recursivePlot(xp.subset(selection_curr{:}),function_handles(2:end),dimensions(2:end),function_arguments(2:end));
                        end
                    end
                end
                
            case 4                          % 4D
                dim1 = dimensions{1}(1);
                dim2 = dimensions{1}(2);
                dim3 = dimensions{1}(3);
                dim4 = dimensions{1}(4);
                for i = 1:sz(dim1)
                    for j = 1:sz(dim2)
                        for k = 1:sz(dim3)
                            for l = 1:sz(dim4)
                                selection_curr{dim1} = i;
                                selection_curr{dim2} = j;
                                selection_curr{dim3} = k;
                                selection_curr{dim4} = l;
                                mydata{i,j,k,l} = @() recursivePlot(xp.subset(selection_curr{:}),function_handles(2:end),dimensions(2:end),function_arguments(2:end));
                            end
                        end
                    end
                end
                
                
        end
        
        % Update axes
        for i = 1:length(dimensions{1})
            dim_curr = dimensions{1}(i);
            myvalues{i} = xp.axis(dim_curr).values;
            ax_names{i} = xp.axis(dim_curr).name;
        end
        
        xp2 = xp2.importData(mydata,myvalues,ax_names);
        xp2 = xp2.fixAxes;
        
        [varargout{1:nargout}] = function_handles{1}(xp2,function_arguments{1}{:});
        
    else
        [varargout{1:nargout}] = function_handles{1}(xp,function_arguments{1}{:});
    end
end
          

function dimensions = dimensions_regex_2_index(xp,dimensions)
    for i = 1:length(dimensions)
        dim_curr = dimensions{i};
        if iscell(dim_curr)
            
            for j = find(cellfun(@ischar,dim_curr))
                
                dim_curr{j} = findaxis_mod(xp,dim_curr{j});
            end
            
            if any(cellfun(@length,dim_curr) > 1); error('Ambiguous dimension supplied'); end
            
            dim_curr = cell2mat(dim_curr);
        elseif ischar(dim_curr)
            dim_curr = findaxis_mod(xp,dim_curr);
        end
        dimensions{i} = dim_curr;
    end
end

function out = findaxis_mod(xp,str)

    if strcmp(str,'data')
        out = 0;
    else
        out = xp.findaxis(str);
    end

end
```
