## Supplemental File 1 for "A biophysical model of striatal microcircuits suggests delta/theta-rhythmically interleaved gamma and beta oscillations mediate periodicity in motor control": recursivePlot_2.m

```
function varargout = recursivePlot_2(xp,function_handles,dimensions,function_arguments)
%% varargout = recursivePlot(xp,function_handles,dimensions,function_arguments)
%     Purpose: Takes in multidimensional data and a set of function handles for
%     plotting. Function handles operate on low-dimensional subspaces. 
%     recursivePlot assigns the high dimensional data in xp to the plotting
%     functions until all dimensions have been used up.
% 
%     Forms:
%         varargout = recursivePlot_2(xp,function_handles,dimensions)
%         varargout = recursivePlot_2(xp,function_handles,dimensions,function_arguments)
% 
%     Inputs:
%       xp                          : xPlt structure with Nd dimensions
% 
%       function_handles            : 1xNd+1 cell array of function handles. See below
%                                     for function handle specifications.
% 
%       dimensions                  : 1xNd cell array of vectors. Each vector corresponds
%                                     to a function handle. Specifies which dimensions
%                                     in xp to assign to each function handle.
% 
%       function_arguments   : (optional) 1xNd cell array of cell arrays, each containing
%                                     additional arguments to pass to each function
%                                     handle. There should be one cell array for
%                                     each function handle.
% 
%     Details:
%       The function_handles cell array should point to functions of the form:
%             varargout = function func (xp,varargin)
%       where xp is an xPlt object. Each function handle
% warning('finish this');
% 
%     Algorithm:
%     recursivePlot plots the data in a recursive manner. It does the following
%         1. Creates a new xPlt structure called xp2, which is a low-dimensional
%            version of xp. The dimensions of xp2 are given by dimensions{1}.
% 
%         2. recusrivePlot then passes xp2 to the first function handle,
%            function_handles{1}.
%            
%            Each entry in xp2.data contains a function handle that recursively
%            calls back recursivePlot. This serves as simply an instruction
%            to handle plotting the remainder of the dimensions in xp (e.g.
%            dimensions(2:end)).
%            
%            function_handles{1} loops through xp2.data and executes these
%            function handles, thus producing plots of xp's remaining dimensions.
%            
%         3. The stopping conditions for the recursion are reached when xp runs
%            out of dimensions. In this case, the final function handle receives
%            a xp object with size(xp) = 1. Thus, the final function handle should
%            actually plot the data.
%            
%     Example
%         See demos_xPlt.m
%            
%            for plotting the 
%                function_handles{1} cycles through all the entries in xp2
%            The dimensions of xp2 are sz(dimensions{1}), where
%            sz = size(xp). xp2.data contains a function handle. This function
%            handle is a recursive call to 
%         2. Calls function_handles{1} and passes it xp2.
%         3. 
%         
%         Takes dimensions{1} from xp and creates a new 
% 
%         Examples:
%             See demos file.


%     function_arguments - cell array of argument cell arrays to pass to
%     function_handles. Must have one cell array for each function_handle
%     passed. Use empty cell arrays for no arguments. E.g.
%     function_arguments = { {}, {}, {1} } for nothing, nothing, and
%     1 as arguments.
    
    if nargin < 4
        function_arguments = cell(size(function_handles));
        for i = 1:length(function_arguments)
            function_arguments{i} = {};
        end
    end
    
    % Validate inputs
    Nfh = length(function_handles);
    Nd = length(dimensions);
    Nfha = length(function_arguments);
    
    if Nfh ~= Nd; error('Number of cells in dimensions must equal the number of function handles supplied'); end
    if Nfh ~= Nfha; error('Number of cells in function_arguments must equal number of function handles supplied'); end
    
    
    % Convert any regular expressions in dimensions into their
    % corresponding indices if necessary
    dimensions = dimensions_regex_2_index(xp,dimensions);
    
    
%     Na = ndims(xp);
%     N_dims_supplied = sum(cellfun(@length,dimensions));
%     if Na ~= N_dims_supplied
%         error('Total number of entries supplied in dimensions must equal ndims(xp)');
%     end

    sz = size(xp);
    if length(function_handles) > 1
        
        % xp_temp = permute(xp, [dimensions{1}, cell2mat(dimensions{2:end})]);
        
        % Set up a new xPlt object with dimensions matching current
        % function handle
        xp2 = xp.reset;                         % Create a new xPlt object        
        sz2 = sz(dimensions{1});
        ndims2 = length(dimensions{1});
        total_calls = prod(sz2);
        
        selection_curr = cell(1,length(sz));
        
        clear output_indices
        [xp2_indices{1:ndims2}] = ind2sub(sz2, 1:total_calls);      % xp2 will loop through chosen dimensions
        
        xp2_indices = num2cell(cat(1, xp2_indices{:})');
        
        xp_indices = cell(1, length(xp.axis));                      % Initialize xp_indices
        
        for call = 1:total_calls
            
            xp_indices(dimensions{1}) = xp2_indices(call, :);
            
            mydata{xp2_indices{call, :}} = @() recursivePlot_2(xp.subset(xp_indices{:}),...
                function_handles(2:end),dimensions(2:end),function_arguments(2:end));
            
        end
        
        % Update axes
        for i = 1:length(dimensions{1})
            dim_curr = dimensions{1}(i);
            myvalues{i} = xp.axis(dim_curr).values;
            mynames{i} = xp.axis(dim_curr).name;
        end
        
        xp2 = xp2.importData(mydata,myvalues,mynames);
        
        [varargout{1:nargout}] = function_handles{1}(xp2,function_arguments{1}{:});
        
    else
        [varargout{1:nargout}] = function_handles{1}(xp,function_arguments{1}{:});
    end
end
    


function dimensions = dimensions_regex_2_index(xp,dimensions)
    for i = 1:length(dimensions)
        dim_curr = dimensions{i};
        if iscell(dim_curr)
            
            for j = find(cellfun(@ischar,dim_curr))
                
                dim_curr{j} = findaxis_mod(xp,dim_curr{j});
            end
            
            if any(cellfun(@length,dim_curr) > 1); error('Ambiguous dimension supplied'); end
            
            dim_curr = cell2mat(dim_curr);
        elseif ischar(dim_curr)
            dim_curr = findaxis_mod(xp,dim_curr);
        end
        dimensions{i} = dim_curr;
    end
end

function out = findaxis_mod(xp,str)

    if strcmp(str,'data')
        out = 0;
    else
        out = xp.findaxis(str);
    end

end
```
