## Supplemental File 1 for "A biophysical model of striatal microcircuits suggests delta/theta-rhythmically interleaved gamma and beta oscillations mediate periodicity in motor control": xPlt.m

```
classdef xPlt < nDDict
    
    % xPlt class inherets from the nDDict class (AKA multidimensional
    % dictionaries). xPlt adds some plotting functionality (see
    % xPlt.RecursivePlot)
    
%     methods
%     
%         function obj = nDDict(obj)
%             % Converter xPlt/nDDict
%             obj = obj;
%         end
%             
%     end
    
end
```
