## Supplemental File 1 for "A biophysical model of striatal microcircuits suggests delta/theta-rhythmically interleaved gamma and beta oscillations mediate periodicity in motor control": nDDictAxis.m

```
classdef nDDictAxis
    % Helper class for handling axis in nDDict class

    properties
        name = ''         % (optional) 1x1 - string naming the axis in question (e.g. time, voltage, cell number, etc)
        values = []       % (optional) 1xN - can be numerical matrix or cell array of strings describing axis in question
        astruct = struct   % (optional) 1xN structure array of additional information
    end
    
    methods
        function out = getaxisinfo(obj,show_class)
            if nargin < 2
                show_class = 1;
            end
            
            max_values_to_display = 10;
            
            if isempty(obj.values); out = 'Empty axis'; return; end
            
            % Add type
            values_class = obj.getclass_values;
            
            if show_class
                temp = [obj.name, ' (' values_class ')  -> '];
            else
                temp = [obj.name, ' -> '];
            end
            
            Nvals = length(obj.values);
            Nvals = min(Nvals,max_values_to_display);          % Limit to displaying 10 values
            
            for i = 1:Nvals-1
                temp = [temp,obj.getvaluestring(i),', '];
            end
            
            if length(obj.values) > max_values_to_display
                temp = [temp,obj.getvaluestring(Nvals),', ...'];
            else
                temp = [temp,obj.getvaluestring(Nvals)];
            end
            
            if nargout > 0
                out = temp;
            else
                fprintf([temp, '\n'])
            end
        end
        
        function out = findAxes(obj, axis_string_ref)
            % Finds numbers of axis matching a given regular expression.
            allnames = {obj.name};
            out = regex_lookup(allnames, axis_string_ref);
        end
        
        function out = getvaluenoncell(obj,i)
            % Looks at entry obj.value(i) and returns its output as a
            % numeric, regardless of whether it is actually cell array.
            if length(i) > 1; error('i must be singleton'); end
            if iscell(obj.values)
                out = obj.values{i};
            else
                out = obj.values(i);
            end
        end
        
        function out = getvaluestring(obj,i)
            % Looks at entry obj.value(i) and returns its output as a
            % string, regardless of what data type it actually is (string,
            % cell, numeric, etc).
            if length(i) > 1; error('i must be singleton'); end
            out = obj.getvaluenoncell(i);
            if isnumeric(out)
                out = num2str(out);
            end
        end
        
        function out = getvaluescellstring(obj)
            % Looks at entry obj.value(i) and returns its output as a
            % cell array of strings
            out = cell(1,length(obj.values));
            for i = 1:length(obj.values)
                out{i} = num2str(obj.getvaluenoncell(i));
            end
        end
        
        function out = getclass_values(obj)
            out = obj.calcClasses(obj.values,'values');
        end
        
        function out = getclass_name(obj)
            out = obj.calcClasses(obj.name,'name');
        end
        
        function out = calcClasses(obj,input,field)
            switch field
                case 'values'
                    % Returns class type of obj.values.
                    if isnumeric(input)
                        out = 'numeric';
                    elseif iscellstr(input)
                        out = 'cellstr';
                    else
                        out = 'unknown';
                    end
                case 'name'
                    % Returns class type of obj.name.
                    if ischar(input)
                        out = 'char';
                    else
                        out = 'unknown';
                    end
                otherwise
                    error('Unrecognized input foramt');
            end
        end
        
        %% % % % % % % % % % % % % % % % % % % % % % % % % % % % % % % 
        % % % % % % % % % % % OVERLOADED OPERATORS % % % % % % % % % % %
        % % % % % % % % % % % % % % % % % % % % % % % % % % % % % % % %
        function varargout = subsref(varargin)
            
%             % Default settings for everything
%             [varargout{1:nargout}] = builtin('subsref',varargin{:});
            
            obj = varargin{1};
            S = varargin{2};
            
            switch S(1).type
                
                case '()'
%                     % Default
%                     [varargout{1:nargout}] = builtin('subsref',varargin{:});                        
                    
                    allnames = {obj.name};
                    if iscellstr(S(1).subs)
                        [selection_out, startIndex] = regex_lookup(allnames, S(1).subs{1});
                        S(1).subs{1} = selection_out;
                        [varargout{1:nargout}] = builtin('subsref',obj,S,varargin{3:end});
                    else
                        % Default
                        [varargout{1:nargout}] = builtin('subsref',varargin{:});                        
                    end
                    
                case '{}'
                    % Default
                    [varargout{1:nargout}] = builtin('subsref',varargin{:});
                case '.'
                    % Default
                    [varargout{1:nargout}] = builtin('subsref',varargin{:});
                otherwise
                    error('Unknown indexing method. Should never reach this');
            end
             
        end

    end
end


function [selection_out, startIndex] = regex_lookup(vals, selection)
    if ~iscellstr(vals); error('nDDictAxis.values must be strings when using regular expressions');
    end
    if ~ischar(selection); error('Selection must be string when using regexp');
    end
    
    startIndex = regexp(vals,selection);
    selection_out = logical(~cellfun(@isempty,startIndex));
    selection_out = find(selection_out);
    
end
```
