## Supplemental File 1 for "A biophysical model of striatal microcircuits suggests delta/theta-rhythmically interleaved gamma and beta oscillations mediate periodicity in motor control": debug_testing_script.m

```
% This is a script that I run through every time I want to test the code.
% Assumes I've just run one of the scripts in demos.m, namely the section
% titled "%% Save data from a set of simulations"
% Then, I load demo_sPING_3:
% data=ImportData('demo_sPING_3');
%% Run simulation - Sparse Pyramidal-Interneuron-Network-Gamma (sPING)
% Save both figures and data.

dynasim_path='~/src/DynaSim';                    
% add DynaSim toolbox to Matlab path
addpath(genpath(dynasim_path)); % comment this out if already in path
%%
cd outputs

% define equations of cell model (same for E and I populations)
eqns={ 
  'dv/dt=Iapp+@current+noise*randn(1,N_pop)';
  'monitor iGABAa.functions, iAMPA.functions'
};
% Tip: monitor all functions of a mechanism using: monitor MECHANISM.functions

% create DynaSim specification structure
s=[];
s.populations(1).name='E';
s.populations(1).size=80;
s.populations(1).equations=eqns;
s.populations(1).mechanism_list={'iNa','iK'};
s.populations(1).parameters={'Iapp',5,'gNa',120,'gK',36,'noise',40};
s.populations(2).name='I';
s.populations(2).size=20;
s.populations(2).equations=eqns;
s.populations(2).mechanism_list={'iNa','iK'};
s.populations(2).parameters={'Iapp',0,'gNa',120,'gK',36,'noise',40};
s.connections(1).direction='I->E';
s.connections(1).mechanism_list={'iGABAa'};
s.connections(1).parameters={'tauD',10,'gSYN',.1,'netcon','ones(N_pre,N_post)'};
s.connections(2).direction='E->I';
s.connections(2).mechanism_list={'iAMPA'};
s.connections(2).parameters={'tauD',2,'gSYN',.1,'netcon',ones(80,20)};

% Vary two parameters (run a simulation for all combinations of values)
vary={
  'E'   ,'Iapp',[0 10 20];      % amplitude of tonic input to E-cells
  'I->E','tauD',[5 10 15]       % inhibition decay time constant from I to E
  };
SimulateModel(s,'save_data_flag',1,'study_dir','demo_sPING_3b',...
                'vary',vary,'verbose_flag',1, ...
                'save_results_flag',1,'plot_functions',@PlotData,'plot_options',{'format','png'} );

cd ..

%% Load the data and import into xPlt class
% ...Assumes we have some DynaSim data already loaded...
cd outputs
data=ImportData('demo_sPING_3b');
cd ..

% Load the data linearly
[data_linear,ax,ax_names,time] = DynaSimExtract (data);


% Import into an xPlt class
xp = xPlt;
xp = xp.importLinearData(data_linear,ax{:});
xp = xp.importAxisNames(ax_names);
meta = struct;
meta.datainfo(1:2) = xPltAxis;      % Use xPltAxis here, because why not?
meta.datainfo(1).name = 'time(ms)';
meta.datainfo(1).values = time;
meta.datainfo(2).name = 'cells';
meta.datainfo(2).values = [];
xp = xp.importMeta(meta);

%% Try selecting a bunch of different subsets, squeezing, and testing for errors
% This should catch most bugs that crop up.
xp2 = xp.subset(2,2,[],7:8);    % 1x1x2x2
    xp2.checkDims;
    xp2.getaxisinfo
    xp2 = xp2.squeeze;
    xp2.getaxisinfo
    xp2.checkDims
    %%
clc
xp2 = xp.subset(2,2,[],8);      % 1x1x2x1
    xp2.checkDims;
    xp2.getaxisinfo
    xp2 = xp2.squeeze;
    xp2.getaxisinfo
    xp2.checkDims
    %%
clc
xp2 = xp.subset(2,2,2,[]);      % 1x1x1x8
    xp2.checkDims;
    xp2.getaxisinfo
    xp2 = xp2.squeeze;
    xp2.getaxisinfo
    xp2.checkDims
    %%
clc
xp2 = xp.subset(1,[],2,1);      % 1x2x1x1
    xp2.checkDims;
    xp2.getaxisinfo
    xp2 = xp2.squeeze;
    xp2.getaxisinfo
    xp2.checkDims
    %%
clc
xp2 = xp.subset([],1,2,1);      % 2x1x1x1
    xp2.checkDims;
    xp2.getaxisinfo
    xp2 = xp2.squeeze;
    xp2.getaxisinfo
    xp2.checkDims
 
    %%
clc
xp3 = xp;
xp3.data = xp.data(:,:,:,5:8);      % Brute force selection of the xp.data.    
                                    % Should produce an error when run
                                    % xp3.squeeze since axes dimensions
                                    % mismatch.
xp3.getaxisinfo;
% xp3.checkDims;      % Should produce an error
xp3 = xp3.fixAxes;
xp3.checkDims;
xp3.getaxisinfo;
                                    
xp6 = xp.subset(1,[],2,1);


%% Run a recursive plot

clear xp2 xp3
xp4 = (xp.subset([],[],[],8));
xp4.getaxisinfo

% recursivePlot(xp4,{@xp_subplot,@xp_subplot,@xp_matrix_basicplot},{1:2,3},{{[],1},{1,1},{}});
% recursivePlot(xp4,{@xp_subplot_grid3D,@xp_subplot,@xp_matrix_basicplot},{1:2,3},{{},{0,1},{}});
% recursivePlot(xp4,{@xp_subplot_grid3D,@xp_matrix_basicplot},{[3,1,2]},{{},{}});
recursivePlot(xp4,{@xp_subplot_grid3D,@xp_subplot_grid3D,@xp_matrix_basicplot},{[1,2,4],3},{{},{0,1},{}});


%% Run another recursive plot
clear xp2 xp3
xp4 = (xp.subset([],[],[],8));
xp4.getaxisinfo

% recursivePlot(xp4,{@xp_subplot,@xp_subplot,@xp_matrix_basicplot},{1:2,3},{{[],1},{1,1},{}});
% recursivePlot(xp4,{@xp_subplot_grid3D,@xp_subplot,@xp_matrix_basicplot},{1:2,3},{{},{0,1},{}});
% recursivePlot(xp4,{@xp_subplot_grid3D,@xp_matrix_basicplot},{[3,1,2]},{{},{}});
recursivePlot(xp4,{@xp_subplot_grid3D,@xp_matrix_basicplot},{[3,1,2]},{{},{}});


%% Run another recursive plot
clear xp2 xp3
xp4 = (xp.subset([],[],1,8));
xp4.getaxisinfo

% recursivePlot(xp4,{@xp_subplot,@xp_subplot,@xp_matrix_basicplot},{1:2,3},{{[],1},{1,1},{}});
% recursivePlot(xp4,{@xp_subplot_grid3D,@xp_subplot,@xp_matrix_basicplot},{1:2,3},{{},{0,1},{}});
% recursivePlot(xp4,{@xp_subplot_grid3D,@xp_matrix_basicplot},{[3,1,2]},{{},{}});
recursivePlot(xp4,{@xp_subplot,@xp_matrix_basicplot},{[1,2]},{{0,0},{}});


%% Plot for 3D data (larger one)

xp4 = xp(1:2,1:2,1,:);
xp4 = xp4.squeeze;
xp4.getaxisinfo

% recursivePlot(xp4,{@xp_subplot,@xp_subplot,@xp_matrix_basicplot},{1:2,3},{{[],1},{1,1},{}});
% recursivePlot(xp4,{@xp_subplot_grid3D,@xp_subplot,@xp_matrix_basicplot},{1:2,3},{{},{0,1},{}});
% recursivePlot(xp4,{@xp_subplot_grid3D,@xp_matrix_basicplot},{[3,1,2]},{{},{}});
recursivePlot(xp4,{@xp_subplot_grid3D,@xp_matrix_basicplot},{[3,1,2],0},{{},{}});


%% Test subset selection using regular expressions
xp5 = xp.subset([],[],[1],'iNa*');
xp5.getaxisinfo

    %%
    xp5 = xp.subset([],[],[1],'_s');
    xp5.getaxisinfo

%% Test packDims
clear xp2 xp3 xp4 xp5
% xp2 = xp.subset(2,2,[],[1,3,5:8]);      % Selection based on index locations
xp2 = xp.subset(2,2,[],'(v|^i||ISYN$)');  % Same thing as above using regular expression. Selects everything except the _s terms. "^" - beginning with; "$" - ending with
xp2 = xp2.squeeze;
xp2.getaxisinfo;
xp2 = xp2.packDim(2,3);
xp2.getaxisinfo;

%% Test mergeDims
xp2 = xp.mergeDims([3,4]);


%% Load xPlt structure of images

cd outputs
file = 'demo_sPING_3b';
data = ImportPlots(file);
cd ..

[data_linear,ax,ax_names] = DynaSimPlotExtract (data);

xp = xPlt;
xp = xp.importLinearData(data_linear,ax{:});
xp = xp.importAxisNames(ax_names);


% to do: rename DynaSimExtrat to DynaSimDataExtract

%% Run recursive plot of images
recursivePlot(xp,{@xp_subplot_grid3D,@xp_plotimage},{[1,2]},{{},{.25}});


%% Run PlotData2
% ...Assumes we have some DynaSim data already loaded...
cd outputs
data=ImportData('demo_sPING_3b');
cd ..


PlotData2(data);
PlotData2(data,'population','E','variable','iNa','varied',{'E_Iapp',1:2});
PlotData2(data,'population','E','variable','iNa','varied',{{'E_Iapp',1:2},{'I_E_tauD',3}});


%% To implement
% 
% 
% Implement the following:
% 1. Example of averaging across cells
% 2. Example of averaging synaptic currents (e.g. LFP estimate)
% 3. Plotting - plots with embedded images
% 4. Plotting - load images directly from DynaSim 
% 5. Starting work on PlotData2 - any new requests?
```
