## Supplemental File 1 for "A biophysical model of striatal microcircuits suggests delta/theta-rhythmically interleaved gamma and beta oscillations mediate periodicity in motor control": demos_xPlt.m

```
%% % % % % % % % % % % % % % % xPlt DEMO - Requires DynaSim % % % % % % % %
% % % % % % % % % % % % % % % % % % % % % % % % % % % % % % % % % % % % % 

%% Set up paths 
% Get ready...

% Format
format compact
restoredefaultpath

% Check if in right folder
[parentfolder,currfolder] = fileparts(pwd);
if ~strcmp(currfolder,'MDD'); error('Should be in MDD folder to run this code.'); end

% Set path to your copy of the DynaSim toolbox
dynasim_path = fullfile(parentfolder,'..');

% add DynaSim toolbox to Matlab path
addpath(genpath(dynasim_path)); % comment this out if already in path

% Set where to save outputs
output_directory = fullfile(parentfolder,'..','outputs');
study_dir = fullfile(output_directory,'demo_sPING_3b');

% move to root directory where outputs will be saved
mkdir_silent(output_directory);
% cd(fullfile(dynasim_path, output_directory));

%% Run simulation - Sparse Pyramidal-Interneuron-Network-Gamma (sPING)

% define equations of cell model (same for E and I populations)
eqns={ 
  'dv/dt=Iapp+@current+noise*randn(1,N_pop)';
  'monitor iGABAa.functions, iAMPA.functions'
};
% Tip: monitor all functions of a mechanism using: monitor MECHANISM.functions

% create DynaSim specification structure
s=[];
s.populations(1).name='E';
s.populations(1).size=80;
s.populations(1).equations=eqns;
s.populations(1).mechanism_list={'iNa','iK'};
s.populations(1).parameters={'Iapp',5,'gNa',120,'gK',36,'noise',40};
s.populations(2).name='I';
s.populations(2).size=20;
s.populations(2).equations=eqns;
s.populations(2).mechanism_list={'iNa','iK'};
s.populations(2).parameters={'Iapp',0,'gNa',120,'gK',36,'noise',40};
s.connections(1).direction='I->E';
s.connections(1).mechanism_list={'iGABAa'};
s.connections(1).parameters={'tauD',10,'gSYN',.1,'netcon','ones(N_pre,N_post)'};
s.connections(2).direction='E->I';
s.connections(2).mechanism_list={'iAMPA'};
s.connections(2).parameters={'tauD',2,'gSYN',.1,'netcon',ones(80,20)};

% Vary two parameters (run a simulation for all combinations of values)
vary={
  'E'   ,'Iapp',[0 10 20];      % amplitude of tonic input to E-cells
  'I->E','tauD',[5 10 15]       % inhibition decay time constant from I to E
  };
SimulateModel(s,'save_data_flag',1,'study_dir',study_dir,...
                'vary',vary,'verbose_flag',1, ...
                'save_results_flag',1,'plot_functions',@PlotData,'plot_options',{'format','png'} );


%% Load the data and import it into xPlt class

% Load data in traditional DynaSim format
data=ImportData(study_dir);

% Extract the data in a linear table format
[data_table,column_titles,time] = Data2Table (data);

% Preview the contents of this table
%     Note: We cannot make this one big cell array since we want to allow
%     axis labels to be either strings or numerics.
previewTable(data_table,column_titles);

% Import the linear data into an xPlt object
xp = xPlt;
X = data_table{1};                          % X holds the data that will populate the multidimensional array. Must be numeric or cell array.
axislabels = data_table(2:end);             % Each entry in X has an associated set of axis labels, which will define its location in multidimensional space. **Must be numeric or cell array of chars only**
xp = xp.importLinearData(X,axislabels{:});
xp = xp.importAxisNames(column_titles(2:end));  % There should be 1 axis name for every axis, of type char.


% xPlt objects are essentially cell arrays (or matricies), but with the
% option to index using strings instead of just integers. 
% Thus, they are analogous to dictionaries in Python.
% (This core functionality is implemented by the multidimensional
% dictionaries (nDDict), which xPlt inherits, and to which xPlt adds
% plotting functionality.)
disp(xp);


% At its core, xPlt has 3 fields. xp.data stores the actual data (either a 
% matrix or a cell array). 
disp(xp.data);
size(xp.data)

% Next, xp.axis stores axis labels associated with each dimension in
% xp.data.
disp(xp.axis(1));

% Axis.values stores the actual axis labels. These can be numeric...
disp(xp.axis(1).values);

% ...or string type. As we shall see below, these axis labels can be
% referenced via index or regular expression.
disp(xp.axis(4).values);

% Axis.name field stores the name of the dimension. In this case, it is
% named after the parameter in the model that was varied.
disp(xp.axis(1).name)

% Axis.astruct is for internal use and is currently empty.
xp.axis(1).astruct

% All of this information can be obtained in summary form by running
% getaxisinfo
xp.getaxisinfo

% xp.meta stores meta data for use by the user as they see fit.
% Here we will add some custom info to xp.metadata. This can be whatever
% you want. Here, I will use this to provide information about what is
% stored in each of the matrices in the xp.data cell array. (Alternatively,
% we could also make each of these matrices an xPlt object!)
meta = struct;
meta.datainfo(1:2) = nDDictAxis;
meta.datainfo(1).name = 'time(ms)';
meta.datainfo(1).values = time;
meta.datainfo(2).name = 'cells';
meta.datainfo(2).values = [];
xp.meta = meta;
clear meta


%% % % % % % % % % % % % % % % xPlt BASICS % % % % % % % % % % % % 
% % % % % % % % % % % % % % % % % % % % % % % % % % % % % % % % % % 

%% Importing data

% xPlt objects are just matrices or cell arrays with extra functionality to
% keep track of what each dimension is storing. 

% Above, we imported data linearized data into an xPlt object. But
% it is also possible to import a high dimensional matrix directly.

% For example let's say we have a random cell array.
data = xp.data;

% We can import the data as follows.
xp2 = xPlt;
xp2 = xp2.importData(data);

% We didn't supply any axis names/values, so default values were assgined
xp2.getaxisinfo;

% We can instead import axis values along with the data
ax_vals = xp.exportAxisVals;
clear xp2
xp2 = xPlt;
xp2 = xp2.importData(data,ax_vals);
xp2.getaxisinfo

% Axis names can be assigned in this way as well
ax_names = xp.exportAxisNames;
xp2 = xp2.importData(data,ax_vals,ax_names);
xp2.getaxisinfo


%% xPlt Indexing

% Indexing works just like with normal matrices and cell arrays and axis
% labels are updated appropriately.
clc
xp4 = xp(:,:,1,8);                  % ## Update - This does the same thing as xp.subset([],[],1,8), which was the old way
                                    % of pulling data subsets. Note that [] and : are equivalent.
xp4.getaxisinfo

% Similarly, can index axis values using regular expressions
% Pull out sodium mechs only
xp5 = xp(:,:,1,'iNa*');
xp5.getaxisinfo

% Pull out synaptic state variables
xp5 = xp(:,:,1,'_s');
xp5.getaxisinfo

% Can also reference a given axis based on its index number or based on its
% name
disp(xp.axis(4))
disp(xp.axis('populations'))

% Lastly, you can reference xp.data with the following shorthand
% (This is the same as xp.data(:,:,1,8). Regular expressions dont work in this mode)
mydata = xp{:,:,1,8};              
mydata2 = xp.data(:,:,1,8);
disp(isequal(mydata,mydata2));

clear mydata mydata2 xp4 xp5


%% % % % % % % % % % % % % % % PLOTTING EXAMPLES % % % % % % % % % % % % 
% % % % % % % % % % % % % % % % % % % % % % % % % % % % % % % % % % % % % 

%% Plot 2D data
% Tip: don't try to understand what recursivePlot is doing - instead, try
% putting break points in the various function handles to see how this
% command works.

% Pull out a 2D subset of the data
clc
xp4 = xp(:,:,'E','v');
xp4.getaxisinfo

% Set up plotting arguments
function_handles = {@xp_subplot_grid,@xp_matrix_basicplot};   % Specifies the handles of the plotting functions
dimensions = {[1,2],0};                                       % Specifies which dimensions of xp each function handle
                                                                % will operate on. Note that dimension "0" refers to the 
                                                                % the contents of each element in xp.data (e.g. the matrix of
                                                                % time series data). If specified, it must come last.
function_arguments = {{},{}};	% This allows you to supply input arguments to each of the 
                                % functions in function handles. For
                                % now we'll leave this empty.
                                                                
% Run the plot. Note the "+" icons next to each plot allow zooming. 
figl; recursivePlot(xp4,function_handles,dimensions,function_arguments);

% Alternatively, dimensions can be specified as axis names instead of
% indices. The last entry, data, refers to the contents of xp.data (e.g.
% dimension 0 above).
dimensions = {{'E_Iapp','I_E_tauD'},{'data'}}; 
figl; recursivePlot(xp4,function_handles,dimensions,function_arguments);

%% Plot 3D data 

% Pull out a 3D subset of data (parameter sweeps and the 2 cell
% types)
clc
xp4 = xp(:,1:2,:,'v');
xp4.getaxisinfo

% This will plot E cells and I cells (axis 3) each in separate figures and
% the parameter sweeps (axes 1 and 2) as subplots.
dimensions = {{'populations'},{'I_E_tauD','E_Iapp'},{'data'}};
recursivePlot(xp4,{@xp_handles_newfig,@xp_subplot_grid,@xp_matrix_imagesc},dimensions);

% Note that here we produced rastergrams instead of time series by
% submitting a different function to operate on dimension zero.

%% Plot 3D data re-ordered

% Alternatively, we can put E and I cells in the same figure. This
% essentially swaps the population and tauD axes.
dimensions = {{'I_E_tauD'},{'populations','E_Iapp'},'data'};
recursivePlot(xp4,{@xp_handles_newfig,@xp_subplot_grid,@xp_matrix_imagesc},dimensions);

%% Plot 4D data

% Pull out sodium channel state variables for E and I cells.
clc
xp4 = xp(1:2,1:2,:,6:7);
xp4.getaxisinfo

dimensions = {'populations',{'E_Iapp','I_E_tauD'},'variables',0};       % Note - we can also use a mixture of strings and index locations to specify dimensions

% Note that here we will supply a function argument. This tells the second
% subplot command to write its output to the axis as an RGB image, rather than
% as subplots. This "hack" enables nested subplots.
function_arguments = {{},{},{1},{}};

if verLessThan('matlab','8.4'); error('This will not work on earlier versions of MATLAB'); end
recursivePlot(xp4,{@xp_handles_newfig,@xp_subplot_grid,@xp_subplot_grid,@xp_matrix_basicplot},dimensions,function_arguments);

%% Plot multiple dimensions adaptively.

% Another option is to use @xp_subplot_grid_adaptive, which will plot the data using axes in
% descending order of the size of the axis values, and plot remaining
% combinations of axis values across figures.

recursivePlot(xp4,{@xp_subplot_grid_adaptive,@xp_matrix_basicplot},{1:4,0});

%% Plot two xPlt objects combined
clc
xp3 = xp(2,:,'E','v');
xp3.getaxisinfo

xp4 = xp(:,3,'E','v');
xp4.getaxisinfo

xp5 = merge(xp3,xp4);

dimensions = {[1,2],0};
figl; recursivePlot(xp5,{@xp_subplot_grid,@xp_matrix_imagesc},dimensions);

%% Plot saved figures rather than raw data

% Import plot files
data_img = ImportPlots(study_dir);

% Load into DynaSim structure
[data_table,column_titles] = DataField2Table (data_img,'plot_files');

% Preview the contents of this table
previewTable(data_table,column_titles);

% The entries in the first column contain the paths to the figure files.
% There can be multiple figures associated with each simulation, which is
% why these are cell arrays of strings.
disp(data_table{1}{1})
disp(data_table{1}{2})

% Import the linear data into an xPlt object
xp_img = xPlt;
X = data_table{1}; axislabels = data_table(2:end);
xp_img = xp_img.importLinearData(X, axislabels{:});
xp_img = xp_img.importAxisNames(column_titles(2:end));

dimensions = {[1,2],[0]};
func_arguments = {{},{.5}};         % The 0.5 argument tells xp_plotimage to
                                    % scale down the resolution of its
                                    % plots by 0.5. This increases speed.

figl; recursivePlot(xp_img,{@xp_subplot_grid,@xp_plotimage},dimensions,func_arguments);


%% % % % % % % % % % % % % % % ADVANCED xPlt / nDDict USAGE % % % % % % % 
% % % % % % % % % % % % % % % % % % % % % % % % % % % % % % % % % % % % % 

%% Modifying xPlt.data directly

% While it is encouraged to use importData, xPlt.data can also be written
% to directly. For example, we can take the average across all cells by
% doing the following.
xp2 = xp;
for i = 1:numel(xp2.data)
    xp2.data{i} = mean(xp2.data{i},2);
end
disp(xp2.data);         % (Note the size is 10001x1 instead of by 10001x20 or 10001x80)

% However, you cannot make modifications that would destroy the 1:1 matchup
% between data and axis (commenting out error producing commands below).
xp2 = xp;
xp2.data{5} = randn(100);       % Ok
% xp2.axis = xp2.axis(1:2);       % Not ok
% mydata = reshape(xp2.data,[3,3,16]);
% xp2.data = mydata;              % Not ok.

%% Method packDim
% Analogous to cell2mat.
clear xp2 xp3 xp4 xp5

% Start by taking a smaller subset of the original xp object.
% xp2 = xp.subset(2,2,[],[1,3,5:8]);      % Selection based on index locations
xp2 = xp.subset(2,2,:,'(v|^i||ISYN$)');  % Same thing as above using regular expression. Selects everything except the _s terms. "^" - beginning with; "$" - ending with
xp2 = xp2.squeeze;
xp2.getaxisinfo;

% Note that xp2 is sparse (there are some empty cells)
disp(xp2.data);         % (E cells don't receive AMPA synapses, and I cells don't receive GABAA synapses)

% Now pack dimension two (columns) of xp2 into xp2.data.
src = 2;                    % Take 2nd dimension in xp2
dest = 3;                   % Pack into 3rd dimension in xp2.data matrix
xp3 = xp2.packDim(src,dest);

% Check dimensionality of xp3.data
disp(xp3.data)             % The dimension "variables", which was dimension 2
                           % in xp2, is now dimension 3 in xp3.data.
                           % Now xp3.data is time x cells x variables

% View axis of xp3
xp3.getaxisinfo;            % The dimension "variables" is now missing

% Alternatively, you can use a regular expression to select the dimension
% you want to pack; if the destination dimension is left empty, the first
% dimension which is not occupied in any of the matrix entries of xp.data
% will be used.
xp3 = xp2.packDim('var');
xp3.getaxisinfo;

% Note some of this data is sparse! We can see this sparseness by plotting
% as follows (note the NaNs)
temp1 = squeeze(xp3.data{1}(100,:,:));  % Pick out a random time point
temp2 = squeeze(xp3.data{2}(100,:,:));  % Pick out a random time point
figure; 
subplot(211); imagesc(temp1);
ylabel('Cells');
xlabel(xp2.axis(2).name); 
set(gca,'XTick',1:length(xp2.axis(2).values)); set(gca,'XTickLabel',strrep(xp2.axis(2).values,'_',' '));

subplot(212); imagesc(temp2);
ylabel('Cells');
xlabel(xp2.axis(2).name); 
set(gca,'XTick',1:length(xp2.axis(2).values)); set(gca,'XTickLabel',strrep(xp2.axis(2).values,'_',' '));

%% Method unpackDim (undoing packDims)
% When packDim is applied to an xPlt object, say to pack dimension 3, the
% information from the packed axis is stored in the nDDictAxis
% matrix_dim_3, a field of xp3.meta.
xp3.meta.matrix_dim_3.getaxisinfo

% If dimension 3 of each cell in xp3.data is unpacked using unpackDim,
% xp3.meta.matrix_dim_3 will be used to provide axis info for the new
% xPlt object.
xp4 = xp3.unpackDim(dest, src);
xp4.getaxisinfo;

% Unless new axis info is provided, that is.
xp4 = xp3.unpackDim(dest, src, 'New_Axis_Names'); % The values can also be left empty, as in xp4 = xp3.unpackDim(dest, src, 'New_Axis_Names', []);
xp4.getaxisinfo;

xp4 = xp3.unpackDim(dest, src, 'New_Axis_Names', {'One','Two','Three','Four','Five','Six'});
xp4.getaxisinfo;

%% Use packDim to average across cells

xp2 = xp;
xp2 = xp(:,:,:,'v');  % Same thing as above using regular expression. Selects everything except the _s terms. "^" - beginning with; "$" - ending with
xp2 = xp2.squeeze;
%
% Average across all cells
xp2.data = cellfun(@(x) mean(x,2), xp2.data,'UniformOutput',0);

% % Convert xp2.data from a matrix into an xPlt object as well. This is
% % useful for keeping track of axis names. 
% mat_ax_names = {'Time','Cell Number'};
% mat_ax_values = {1:10001, []};
% 
% % xp2.data = Cell_2_nDDict(xp2.data,mat_ax_names,mat_ax_values);

% Pack E and I cells together
src=3;
dest=2;
xp3 = xp2.packDim(src,dest);


% Plot 
figl; recursivePlot(xp3,{@xp_subplot_grid,@xp_matrix_basicplot},{[1,2],[]},{{},{}});

%% Use packDim to average over synaptic currents
% Analogous to cell2mat
% See also plotting material by Hadley Wickham

% First, pull out synaptic current variables
xp2 = xp(:,:,:,'(ISYN$)');  % Same thing as above using regular expression. Selects everything except the _s terms. "^" - beginning with; "$" - ending with
xp2.getaxisinfo;

% Second, put this into matrix form, so we can average over them
xp3 = xp2.packDim(4,3);
disp(xp3.data)              % xp3.data is now 3D, with the 3rd dim denoting synaptic current
xp3 = xp3.squeeze;
xp3.getaxisinfo;

% Average across membrane currents
xp3.data = cellfun(@(x) nanmean(x,3), xp3.data,'UniformOutput',0);

% Plot 
recursivePlot(xp3,{@xp_handles_newfig,@xp_subplot_grid,@xp_matrix_basicplot},{[3],[1,2],[0]});

%% Test mergeDims
% Analogous to Reshape.

% This command combines two (or more) dimensions into a single dimension.
xp2 = xp.mergeDims([3,4]);
xp2.getaxisinfo;

%% Use mergeDims to convert to Jason's DynaSim format
% Analogous to Reshape

% This combines the 2D "vary" sweep into a single dimension. It also
% combines all populations and variables into a single 1D list. Thus, Axis
% 1 is equivalent to Jason's structure array - data(1:9). Axis 4 is
% equivalent to the structure fields in Jason's DynaSim structure.
xp2 = xp.mergeDims([3,4]);
xp2 = xp2.mergeDims([1,2]);
xp3 = squeeze(xp2); % Squeeze out the empty dimensions.
xp3.getaxisinfo;


% Note that the variable names are not sorted the same was as in Jason's
% DynaSim structure, in that they alternate E and I cells
disp(xp3.axis(2).values);
% Can easily sort them as follows.
[~,I] = sort(xp3.axis(2).values);
xp4 = xp3(:,I);
disp(xp4.axis(2).values);

%% To implement
% 
% 
% Implement the following:
% + Make DynaSimPlotExtract more general
% + Starting work on PlotData2 - any new requests?
```
