## Supplementary figures and images for "A biophysical model of striatal microcircuits suggests delta/theta-rhythmically interleaved gamma and beta oscillations mediate periodicity in motor control"

### demoicon.gif

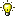

### matlabicon.gif

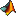

### simulinkicon.gif

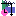
